# *Daphnia magna* modifies its gene expression extensively in response to caloric restriction revealing a novel effect on haemoglobin isoform preference

**DOI:** 10.1101/2020.05.24.113381

**Authors:** Jack Hearn, Jessica Clark, Philip J. Wilson, Tom J. Little

## Abstract

Caloric restriction (CR) produces clear phenotypic effects within and between generations of the model crustacean *Daphnia magna*. We have previously established that micro RNAs and cytosine methylation change in response to CR in this organism, and we demonstrate here that CR has a dramatic effect on gene expression. Over 6000 genes were differentially expressed between CR and well-fed *D. magna*, with a bias towards up-regulation of genes under caloric restriction. We identified a highly expressed haemoglobin gene that responds to CR by changing isoform proportions. Specifically, a transcript containing three erythrocruorin domains was strongly down-regulated under CR in favour of transcripts containing fewer or no such domains. This change in the haemoglobin mix is similar to the response to hypoxia in *Daphnia,* which is mediated through the transcription factor hypoxia-inducible factor 1, and ultimately the mTOR signalling pathway. This is the first report of a role for haemoglobin in the response to CR. We also observed high absolute expression of super-oxide dismutase (SOD) in normally-fed individuals, which contrasts with observations of high SOD levels under in CR in other taxa. However, key differentially expressed genes, like SOD, were not targeted by differentially expressed micro-RNAs. Whether the link between Haemoglobin and CR is the case in other organisms, or is related to the aquatic lifestyle, remains to be tested. It suggests that one response to CR may be to simply transport less oxygen and lower respiration.

## Introduction

Caloric restriction (CR) is the reduction in dietary intake of calories without undernutrition (Koubova & Guarente, 2003). CR induces marked phenotypic changes in many organisms. Most notably an increase in longevity has been observed in various arthropods, rodents, yeast, and possibly also humans (Heilbronn & Ravussin, 2003; Kapahi, Kaeberlein, & Hansen, 2017; Lakowski & Hekimi, 1998; Redman & Ravussin, 2011; Sohal & Weindruch, 1996; Walford, Harris, & Weindruch, 1987). This occurs through CR-mediated delays in the onset of processes and diseases associated with ageing (Koubova & Guarente, 2003; Most, Tosti, Redman, & Fontana, 2017). As a result, moderate CR could well be beneficial to human health, although this remains to be confirmed by sufficiently long-running clinical trials (Most et al., 2017).

The transcriptomic response to CR has been investigated in a variety of model organisms spanning mammals, invertebrates and yeasts (Choi et al., 2018; Ding et al., 2014; Dobson et al., 2018; Heintz et al., 2017; Kapahi et al., 2017; Kim et al., 2016; Matthews et al., 2017; Regan et al., 2016; Wood et al., 2015). Neuroprotective qualities of CR have been attributed to an altered gene expression profile of CR rats due to an altered response to oxidative stress and histone deacetlyase activity, in addition to changes to insulin-signalling pathway and longevity-associated gene levels (Wood et al., 2015). In *Drosophila* CR responses have been characterised at the whole-organism and tissue-level (Ding et al., 2014; Dobson et al., 2018; Kapahi et al., 2017; Regan et al., 2016). There is an overall down-regulation of genes under CR (Ding et al., 2014), which is a hallmark of CR (Russell & Kahn, 2007). However, despite the popularity of *Daphnia magna* for studying life-history traits and the effects of environmental stressors (Boersma, Spaak, & De Meester, 1998; Garbutt & Little, 2014, 2017; Lampert, 1987; Latta IV, Frederick, & Pfrender, 2011; Mitchell & Lampert, 2000; Orsini et al., 2016), the gene-level response to CR has not previously been studied in this species.

### Environmental stressors and caloric restriction in *Daphnia magna*

There is a wealth of existing phenotypic data in *Daphnia*, including demonstrations of CR-mediated lifespan increase (Latta IV et al., 2011), which we confirm and expand upon here. There are also clear maternal effects due to CR where *D. magna* mothers exposed to CR produce offspring that are (1) larger at birth, (2) feed at a slower rate than offspring of abundantly fed mothers, and are (3) more resistant to parasitism (Garbutt & Little, 2014, 2017). Furthermore, *Daphnia* are aquatic invertebrates, and it is possible that they respond to CR in a manner that is partially distinct from terrestrial invertebrates. We explore this novel perspective in the context of haemoglobins here.

### *Daphnia* Haemoglobins and the link to CR

Haemoglobin (henceforward “Hb”) production was observed to increase in response to food levels in *D. magna* (Fox, Gilchrist, & Phear, 1951; Zeis, 2020). However, no further studies considered the relationship between nutrition and Hb in *Daphnia*. By contrast, it is well established that the response to hypoxia and hyper-thermal stress in *D. magna* and *D. pulex* causes changes in Hb concentration and subunit components to occur (Cuenca Cambronero, Zeis, & Orsini, 2018; Gerke, Börding, Zeis, & Paul, 2011; Lai et al., 2016; Lyu et al., 2015; Zeis et al., 2003). This suggests that there is phenotypic plasticity in the *Daphnia* Hb response to stress. Interestingly, acclimation to hypoxic conditions in *D. magna* results in smaller adults without impacting on clutch size during the first five broods (Seidl, Paul, & Pirow, 2005). A phenotype which is similar to that for the offspring of CR individuals (Garbutt & Little, 2014). Links between nutrition and hypoxia have not been explored widely, but in *Drosophila* a specifically low-protein CR regime partially reverses the reduction in lifespan caused by hypoxia (Vigne & Frelin, 2006).

*Daphnia magna* encodes 12 di-domain Hb genes [as annotated in the gene-set: (Orsini et al., 2016)] that result from a complex series of duplications (Colbourne et al., 2011; Zeis, 2020). Sixteen Hb di-domains are aggregated to form the free-circulating *D. magna* Hb molecule (Zeis, 2020). Structural differences in Hb molecules comprised of the products of different Hb isoforms is correlated to changes in oxygen-binding characteristics (Zeis, 2020). The induction of Hb genes in *Daphnia* is mediated by the transcription factor hypoxia-inducible factor (HIF) (Gorr, Cahn, Yamagata, & Bunn, 2004; Zeis, 2020), a heterodimer formed from HIF-1α and HIF-1β proteins. The HIF-1 complex is an important factor in extending longevity of *Caenorhabditis elegans* under CR (Di Chen & Kapahi, 2009; Lee, Hwang, & Kenyon, 2010; Y. Zhang, Shao, Zhai, Shen, & Powell-Coffman, 2009). Increased expression of HIF-1 alone was responsible for this and was triggered by elevated concentrations of mitochondrial reactive oxygen species (Lee et al., 2010). HIF-1 itself is regulated by the mTOR pathway (Land & Tee, 2007) and in *C. elegans* is dependent on the inositol-requiring protein-1 (IRE-1) of the unfolded protein stress response in the endoplasmic reticulum (Di Chen & Kapahi, 2009; Kapahi et al., 2017).

### Master control mechanisms underpinning CR responses

The free radical theory of ageing has been linked to CR (Gladyshev, 2014; Liochev, 2013; Viña, Borras, Abdelaziz, Garcia-Valles, & Gomez-Cabrera, 2013). The theory hypothesises that ageing results from accumulated oxidative damage due to reactive oxygen species. Super-oxide dismutase (SOD), a key component of the theory (Gladyshev, 2014), defends against reactive oxygen species (ROS) by converting abundant superoxide into hydrogen peroxide and oxygen. Evidence that lacking SOD genes results in reduced lifespans across taxa supports this hypothesis (Muid, Karakaya, & Koc, 2014; Muller et al., 2006; Oka, Hirai, Yasukawa, Nakahara, & Inoue, 2015).

Four nutrient-sensing, signalling pathways are most often implicated in lifespan extension as a result of CR (Kenyon, 2010). They are 1) the insulin/insulin-like growth factor (IGF-1), 2) the mechanistic target of rapamycin (mTOR) signalling pathway, 3) 5' adenosine monophosphate-activated protein kinase (AMP Kinase), and 4) sirtuin signalling protein-modulated pathways. These pathways are complex and interlinking, with mTOR signalling potentially acting as the ultimate modulator of the other three (Johnson, Rabinovitch, & Kaeberlein, 2013).

**The present study adds expression data to data on *D. magna* miRNA and DNA methylation responses to CR**

To characterise phenotypic and genotypic ageing in a single organism, we performed a whole organism longevity experiment on a set of eight *D. magna* genotypes each subjected to a calorically restricted and normal diet. To contrast gene expression between food levels, we selected one genotype that demonstrated increased longevity here. We have previously established that life history traits (Garbutt & Little, 2014, 2017), DNA methylation (Hearn, Pearson, Blaxter, Wilson, & Little, 2019), and miRNA expression in the selected strain (Hearn et al., 2018) respond to CR; indeed the miRNA was sequenced from the same set of RNA extractions. Using gene and transcript level expression change of this CR-responsive genotype of *D. magna* we now identify known and novel gene families and metabolic pathways implicated in CR.

## Materials and Methods

### Longevity experiment

This experiment used eight genotypes from geographically dispersed populations. These were obtained from the *D. magna* diversity panel held in Basel, Switzerland (http://evolution.unibas.ch/ebert/research/referencepanel/). They are (clone ID, Country of Origin): BEK22 (Belgium), Clone 32 (UK), FIFAV1 (Finland), GBEL75 (UK), GG8 (Germany), ILPS1 (Italy), MNDM1 (Mongolia), RUYAK1-6 (Russia). Prior to the experiments, replicates of all genotypes were put through three generations of acclimation to harmonise environmental effects arising from variation in stock conditions. During this period, each individual was maintained in a 60ml glass jar filled with artificial pond medium (Kluttgen et al., 1994). This was changed twice weekly and when offspring were produced. Each individual was fed ×6.25 × 10^6^ *Chlorella vulgaris* cells daily and was maintained on a 12:12 L:D cycle at 20°C. We estimate cell numbers by measuring the daily optical absorbance of 650nm white light by the *Chlorella* culture, with 1.0 absorbance being equivalent to approximately 5 × 10^6^ algal cells. Offspring from the second clutch initiated each generation, including the experimental generation.

From the acclimated females of all eight genotypes, two offspring from clutch two were taken. One was assigned to normal food (NF: ×6.25 × 10^6^ cells as per acclimation) and one was assigned to a caloric restriction (CR: ×1.4 × 10^6^ cells) treatment, which is approximately 20% of the amount of food available to NF replicates. Each food treatment and genotype combination were replicated 24 times. Date of birth and date of death were recorded. They were otherwise maintained identically to the acclimation period.

### Survival Analysis

A Cox’s proportional hazards model was used to test for differences in longevity between the two food treatments for all eight genotypes. This was done using the survival package in R (code used and model outputs: Supplementary File S1, clone longevity input data: Supplementary File S2). The response variable was days alive, with genotype, food treatment and their interaction (days alive = clone + food + clone*food), as fixed effects. There was no censoring as all individuals were followed from their day of birth to the day of death. We present the results of the analysis of deviance (ANOVA) for the Cox’s model which performs *χ*^2^-tests of likelihood ratios of each model factor sequentially, which in this case was food followed by clone followed by the interaction between them.

### Material for RNA harvesting

Clone 32 (UK) (Auld, Hall, Housley Ochs, Sebastian, & Duffy, 2014) was selected for RNA sequencing because it showed a longevity response in the above experiment, and it shows clear maternal effects under variation in maternal food - larger offspring are produced under CR (Garbutt & Little, 2014, 2017). This clone was also the focus of more detailed analysis of food and longevity (Clark, Wilson, McNally and Little, *In revision*) where it again showed lifespan extension in response to food restriction. To generate RNA, maternal lines of C32 were first acclimatized for three generations in artificial pond medium at 20°C and on a 12h:12h light:dark cycle and fed 2.5 × 10^6^ cells of the single-celled green algae *Chlorella vulgaris* daily. The treatment generation [G_0_ in (Hearn et al., 2018)] was then split into two groups of eight replicates paired by mother and fed either a normal diet of 5 × 10^6^ algal cells/day or a caloric restricted diet of 1 × 10^6^ algal cells/day per individual. Each replicate was formed of five *D. magna* reared in the same jar from birth.

After the birth of the first clutch of offspring in a replicate jar, the jar was treated for microbial contamination with tetracycline and ampicillin (as described in Hearn et al., 2018) over 24 hours, then *D. magna* individuals were homogenised in 700ul Qiazol Qiagen reagent ID: 79306), and stored at −70°C until further processed. RNA was extracted using miRNeasy mini kits (Qiagen Cat No./ID: 74106) and integrity and quality checked by Qubit (Thermo Fisher) fluorometer, nanodrop (Thermo Fisher) and Bioanalyzer (Agilent). No degradation was observed on Bioanalyzer total RNA traces. Extractions were subsequently halved and one half was used to create small RNA libraries for miRNA expression analysed in (Hearn et al., 2018). TruSeq stranded mRNA-seq (Illumina, San Diego, USA) libraries were prepared by Edinburgh Genomics from the remaining RNA for eight normal food and caloric-restricted replicates as for the previous study. All libraries were multiplexed and sequenced on one lane of HiSeq 4000 to at a targeted depth of 15 million read-pairs per sample yielding a total of at least 290 million read-pairs. Raw sequencing data generated by this project was deposited in the European Nucleotide Archive under Bioproject PRJEB25137.

### Differential gene expression and transcript usage analysis

Reads were adapter and quality trimmed with Cutadapt (version 1.16, options: −q 15––trim-n −m 36) (Martin, 2011) and then Trimmommatic (version 0.36, default options) (Bolger, Lohse, & Usadel, 2014) to remove remaining residual adapter sequence. Fastq files were inspected using FastQC (Andrews, 2010) before and after quality filtering to confirm the removal of adapter-derived and low-quality sequences and reports combined using multiQC (versions 1.8). We used a transcript-driven approach to quantify gene expression as (1) an excellent gene-set is available for *D. magna* (Orsini et al., 2016, 2018), (2) ‘alignment-free’ mapping approaches to transcripts are at least the equal of genome-based alignments for RNASeq (C. Zhang, Zhang, Lin, & Zhao, 2017), and (3) the *D. magn*a genome is still in draft form with many genes overlapping and/or spread across multiple scaffolds. Gene expression per replicate was quantified using Salmon v0.13.1 (Patro, Duggal, Love, Irizarry, & Kingsford, 2017) with parameters “salmon quant −-dumpEq −-validateMappings −-rangeFactorizationBins 4 -l A --seqBias -gcBias” against the *D. magna* reference transcriptome (Orsini et al., 2016, 2018). The reference transcriptome was created by combining principal and alternative transcripts (downloaded from http://arthropods.eugenes.org/EvidentialGene/daphnia/daphnia_magna/Genes/earlyaccess) and indexed with a k-mer size of 25.

The salmon results were converted into a gene-by-replicate expression matrix for input to DESeq2 (Love, Huber, & Anders, 2014) with the R package tximport (Soneson, Love, & Robinson, 2015). Read mapping rates per replicate were taken from the resulting Salmon log files (Supplementary Table S1). Differential gene expression between caloric restriction and normal food replicates was tested using DESeq2, with ‘mother’ fit as a blocking factor (× mother + condition). We incorporated log_2_-fold changes into our significance test and report genes significant at log_2_-fold change thresholds of 1, and 2 as this is more robust than post-hoc filtering of genes by log_2_-fold change alone. This method results in s-values for the non-zero log_2_-fold change analyses (i.e log_2_-fold change 1 and 2) which are analogous to q-values (Stephens, 2016; Zhu, Ibrahim, & Love, 2019). S-values are a measure of the chance that the sign (+ or −) of the log_2_-fold change for the gene is question is incorrect. We applied a significance threshold of 0.005 to the s-values as recommended by DESeq2 and apeglm (Zhu et al., 2019) package authors, as s-values are less conservative than q-values. We also performed significance testing without imposing a log_2_-fold change threshold (referred to as 0 log_2_-fold change) in DESeq2, with a q-value threshold of 0.05. We advise caution on the use of log_2_-fold change thresholds, however. Although they helped us to identify a general trend in the data, the method penalised high expression genes. This is because the variance in gene-expression increases with the level of gene expression, meaning that high-expression genes have wider confidence intervals on their predicted log_2_-fold changes. Therefore, we also considered genes differentially expressed at log_2_-fold change 0 with overall expression of greater than 10,000 length-scaled transcript per million (TPM) when interpreting the results. Differential transcript usage (DTU) was tested in DRIMSeq (Nowicka & Robinson, 2016) and DEXSeq (Anders, Reyes, & Huber, 2012) and the overall false discovery rate (OFDR) for a two-step procedure (here referring to gene- and transcript-level) was calculated in stageR following (Love, Soneson, & Patro, 2018), with ‘mother’ fit as a blocking factor as for DESeq2. Only those genes that exhibited significant DTU in both DRIMSeq and DEXSeq after OFDR were considered further. Differential gene and transcript usage scripts are given in Supplementary File S1.

### TopGO and gene set enrichment analysis

Gene ontology term enrichment was tested in topGO (Alexa & Rahnenfuhrer, 2016) for each category of significant gene from gene- and transcript-level analyses using *D. magna* GO term annotations following (Hearn et al., 2018; Hearn, Pearson, et al., 2019). A conservative p-value threshold of 0.01 was applied with no multiple testing correction in line with topGO author recommendation (Section 6.2. The adjustment of p-values, topGO manual: https://bioconductor.org/packages/release/bioc/html/topGO.html). We considered enriched GO terms of the biological process (BP) and molecular function (MF) sub-categories for discussion. We clustered the significantly enriched GO terms in two-dimensional semantic space and treemaps of higher-order processes using REVIGO (Supek, Bošnjak, Škunca, & Šmuc, 2011) following (Hearn, Blaxter, et al., 2019) to identify patterns in enrichment and reduce redundancy in GO terms. We adapted the REVIGO webserver-produced R-script for each category of enriched GO terms to combine groups of terms by colour according to the REVIGO treemap categories (R script, Supplementary File S1).

Gene set enrichment analysis (GSEA) of *D. magna* KEGG (Kyoto Encyclopedia of Genes and Genomes) orthologs (Kanehisa & Goto, 2000; Kanehisa, Sato, Kawashima, Furumichi, & Tanabe, 2015) was applied to identify KEGG gene pathways up- or down-regulated under CR. GSEA was run for all genes in the expression experiment using their log_2_-fold changes from the log_2_-fold change zero differential gene expression experiment. To obtain KEGG annotations for each gene the *D. magna* gene-set was annotated with trinotate (Haas et al., 2013) and gene-to-KEGG annotations were input to the universal enrichment protocol of clusterProfiler (Yu, Wang, Han, & He, 2012), a q-value cut-off of 0.05 was used to determine significance (R scripts: Supplementary File S1). We ran 1000 iterations of GSEA as the results were variable between runs and selected only those KEGG terms found significant in >95% of runs.

### Interaction between expression level and differential methylation

Average TPM gene expressions calculated in tximport and normalised in DESeq for CR and NF were correlated with the proportion of methylation within genes for the corresponding treatment using data taken from Hearn, Pearson, et al., (2019). Proportion methylated was defined as the count of methylated reads over the total reads at CpG sites within a genic region (exons + introns) aligned against from the *D. magna* genome assembly (version 2.4) using Bismark (Krueger & Andrews, 2011), see Hearn, Pearson, et al., (2019) for further detail. Genic regions were defined from the *D. magna* reference annotation (downloaded from http://arthropods.eugenes.org/EvidentialGene/daphnia/daphnia_magna/Genes/earlyaccess/dmagset7finloc9c.puban.gff.gz). CpG counts for CR or NF were extracted by combining CpG counts of the six replicates per treatment from Hearn, Pearson, et al., (2019) from Bismark bam files. Average methylation rates within genic regions per treatment were calculated using bedtools (Quinlan & Hall, 2010). We then combined CpG averages with corresponding average TPM expression for each gene, calculated from the normalised Salmon count matrix. Genes that had expression greater than 10 TPM, at least 5% CpG methylation, and did not overlap another gene in *D. magna* annotation were retained for the correlation analysis. We applied Spearman’s rho and Kendall’s tau calculated in base R to the results, because of the lack of normality in the proportions of methylation and mean expression data. Furthermore, we intersected the list of differentially expressed genes at each log_2_-fold threshold and differential transcript usage with the differentially-methylated regions identified in response to CR from Hearn, Pearson, et al., (2019).

### miRNA target prediction and mRNA-miRNA expression correlation

MiRNA targets of differentially expressed miRNAs identified in Hearn et al., (2018) were predicted in the 3’ untranslated regions (3’ UTRs, following Graham & Barreto, 2019) of differentially expressed genes using PITA (Kertesz, Iovino, Unnerstall, Gaul, & Segal, 2007), RNAhybrid (Krüger & Rehmsmeier, 2006), miRanda (Enright et al., 2003), MicroTar (Thadani & Tammi, 2006) and rna22 (Miranda et al., 2006) implemented on the tools4miRs platform (Lukasik, Wójcikowski, & Zielenkiewicz, 2016). Only those 3’ UTRs predicted as targets by at least four programs were considered further. Pearson’s correlation performed in miRLAB (Le, Zhang, Liu, Liu, & Li, 2015). The DESeq2 normalised count matrices for differentially expressed miRNAs from (Hearn et al., 2018) and mRNAs at log_2_-fold change 0 were supplied as input to miRLAB, and the results were intersected with the list of predicted miRNA to mRNA targets. MiRNA-mRNA pairs with greater than 0.5 or less than −0.5 correlation in expression level were considered further.

## Results

### Effect of CR on longevity

Cox’s proportional hazards ANOVA (see Supplementary File S1 for R code and model output) of the survivorship data (Supplementary File S2) showed that CR *D. magna* lived longer on average than NF *D. magna* (*χ*^2^ = 13.26, p = 0.0003; Figure 1, part A). Clone 32 was one of the clones that demonstrated increased longevity under CR (Figure 1, Part B). Genotypes also differed in their lifespans (*χ*^2^ = 155.4, p < 0.0001), and there was a significant interaction between genotype and food level for differences in longevity (Figure 1; *χ*^2^ = 29.4, p = 0.0001).

**Figure 1.**
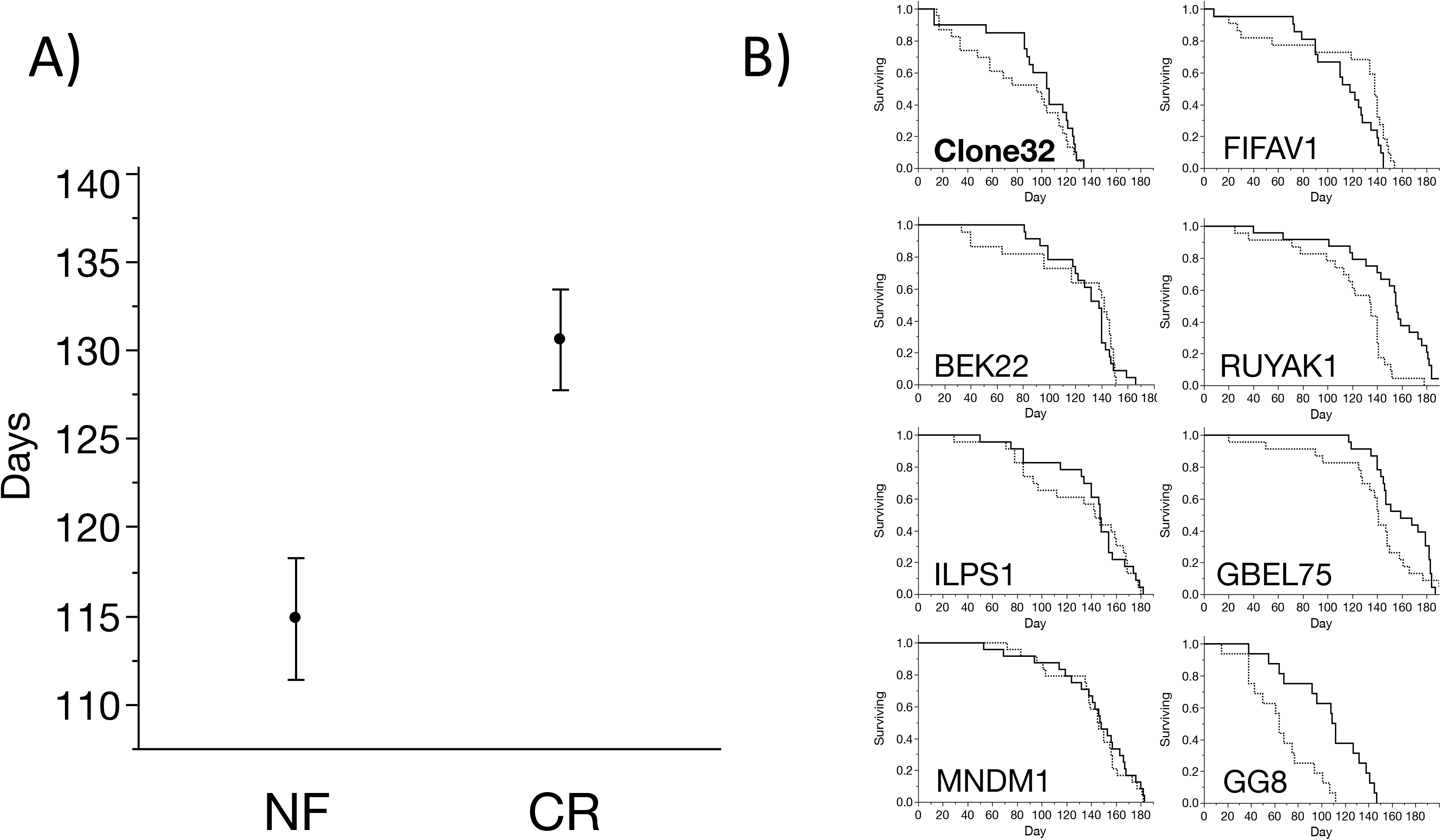
CR *Daphnia magna* lived approximately 15 days longer than NF *D. magna* on average. A) Point estimates are mean longevity across clones for each treatment and whiskers are standard errors. B) Survivorship between treatments varied among the eight different *D. magna* genotypes (dashed line = NF, solid line = CR); Clone 32 is labelled in bold in the upper-left plot.

### Expression experiment

A median of 23,365,302 read-pairs were sequenced per sample with a range of 21,452,900-26,752,873, and FastQC revealed no issues with read quality after trimming (Supplementary Table S1: read filtering and alignment rates, Supplementary File S3: multiQC report). Read mapping rates of greater than 80% were recorded for all replicates, except one (3H2, normal food) that had a mapping rate of 74% (Supplementary Table S1). Similar differences in gene expression occur at the 0 log_2_-fold change analysis at 3345 up-regulated and 3064 down-regulated under CR (Table 1, Supplementary Tables S2-S4). This global response was reflected in principal components analysis of replicates (Figure 2), in which the first component explains 49% of the variance and separates all sample pairs by treatment. This was also true for miRNAs derived from the same RNA extractions (Hearn et al., 2018). However, with increasing log_2_-fold change thresholds there was a bias towards genes being up-regulated in CR (Table 1).

**Table 1.**
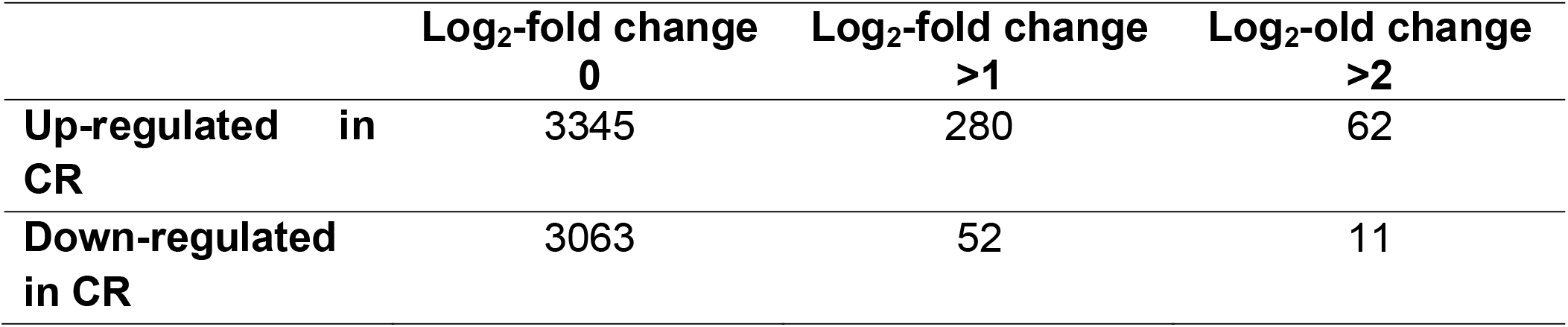
Significantly differentially expressed genes at each log_2_-fold change threshold. More genes were up-regulated in CR replicates.

**Figure 2.**
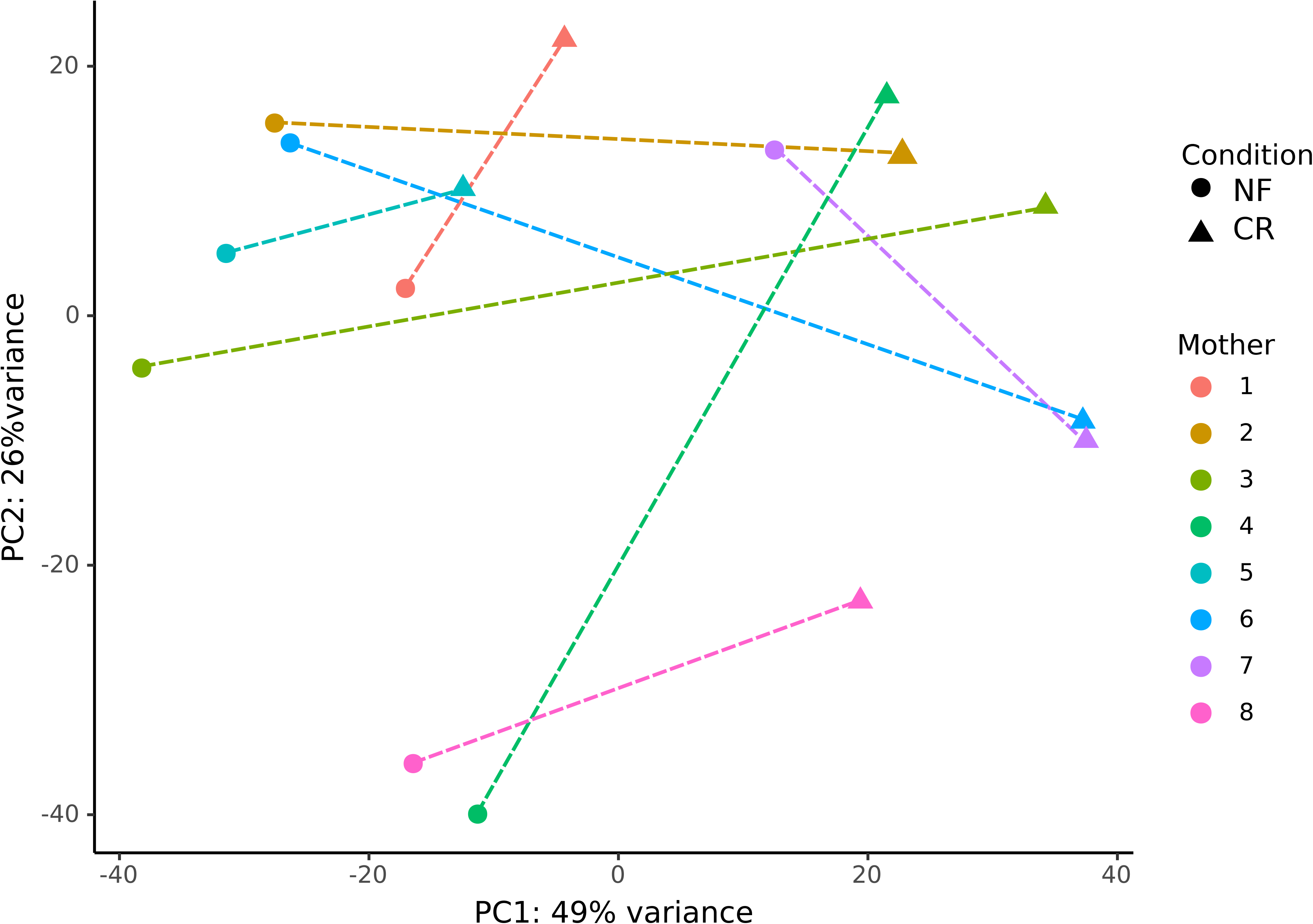
DESeq2 generated PCA plots of regularized logarithm transformation of data of replicates in the nutrition experiment. Shapes define treatment, and colour defines mother. Figure legend: NF = normal food, CR = caloric restriction. Dashed lines indicate relationship between replicates paired by mother; these are always divided along the ×-axis in the same direction.

There were 75 genes with a mean expression greater than 10,000 TPM that were significant at the log_2_-fold change 0 threshold up-regulated in CR, versus 15 that were down-regulated. Four of the 15 genes down-regulated under CR were super-oxide dismutases, including the most expressed gene (Dapma7bEVm009708) at a mean expression of 847468.627 TPM. This was over 6-fold greater than the next gene, a Vitellogenin-1 precursor (Dapma7bEVm014991) (Supplementary Table S5: gene descriptions and expression levels). Two further Vitellogenin-1 precursor genes were in the top five most expressed genes (Dapma7bEVm018415 and Dapma7bEVm029595), as was a di-domain hemoglobin precursor (Dapma7bEVm029622). By contrast, the gene lists for the log_2_-fold changes 1 and 2 were composed of genes at relatively modest expression levels and contained a high proportion of uncharacterised genes (Supplementary Table S5).

### Differential Transcript Usage and Hb related genes

For differential transcript usage, 498 transcripts corresponding to 294 genes were significant in the DEXSeq analysis and 327 transcripts from 187 genes for the DRIMSeq analysis. There was an overlap of 181 transcripts and 112 genes between the two methods (Supplementary File S4: DRIMSeq transcript expression levels for DTU significant genes). Of these, 32 genes have a mean gene expression normalised count level greater than 1000 TPM and five greater than 10000 TPM. The most highly-expressed DTU exhibiting gene was a di-domain haemoglobin precursor (Figure 3, Dapma7bEVm014981) with a TPM of 65,675. The isoform most abundant in CR replicates (“transcript 21”, Figure 3) does not contain an erythrocruorin domain, whereas an isoform containing three such domains (”transcript 12”, Figure 3), is down-regulated in CR. A further di-domain haemoglobin was also in this high expression group (Dapma7bEVm015367, TPM 13,532). Other notable examples of DTU included a Neurocalcin in which transcript 2 is heavily up-regulated in CR (Supplementary File S4: Dapma7bEVm018721), a Carboxypeptidase A4 (Dapma7bEVm015011), a Lactosylceramide (Dapma7bEVm007442), and an Ezrin-moesin-radixin (Dapma7bEVm009772).

**Figure 3.**
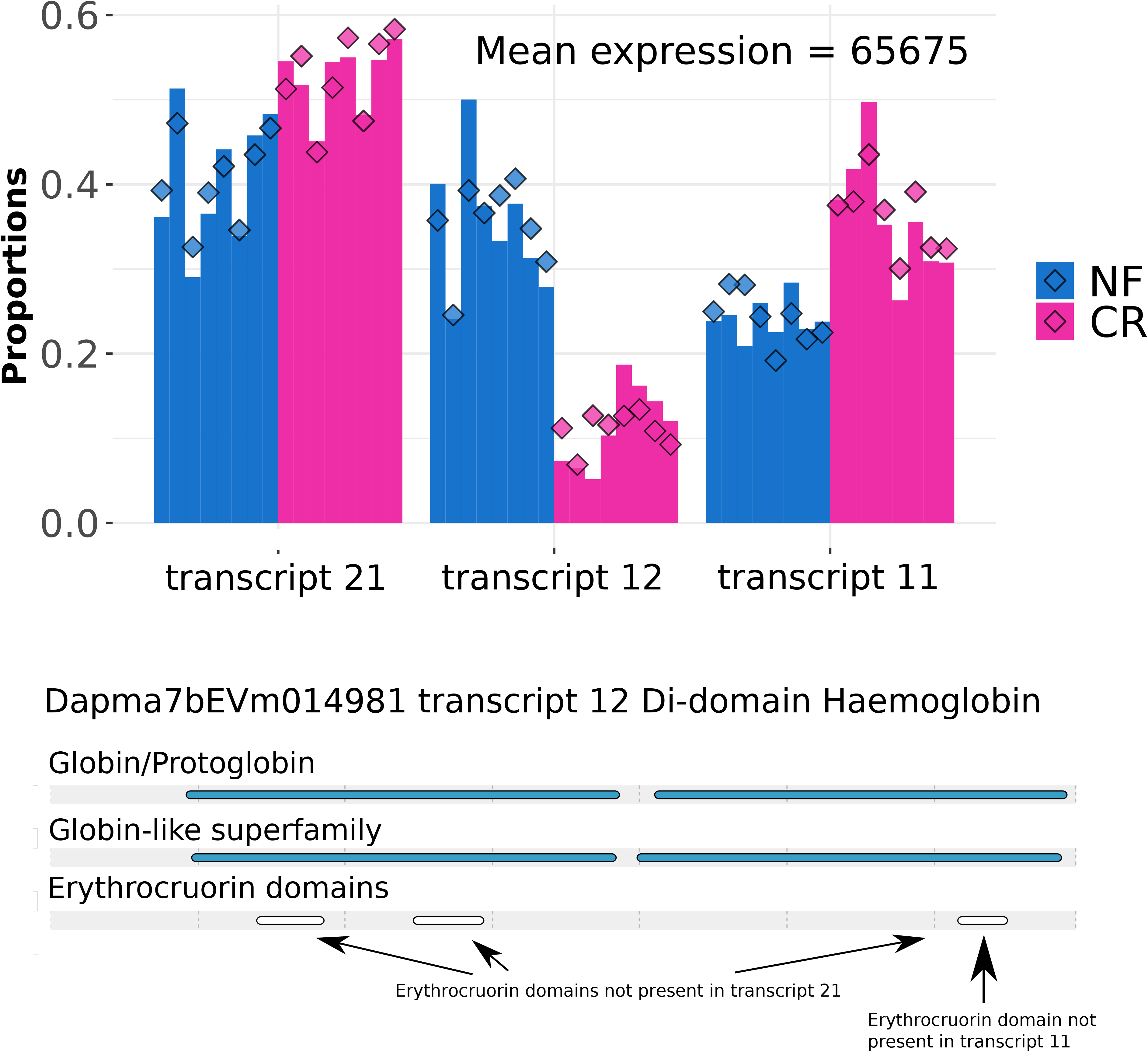
Transcript proportion plots and protein domains for differentially used transcripts of an up-regulated di-domain haemoglobin. Transcripts 11, 12, and 21 respectively of gene Dapma7bEVm014981 show DTU across conditions in DEXSeq and transcript 12 in DRIMSeq adjusted results. Bars represent proportions of total gene expression for that transcript per replicate and diamonds DRIMSeq fitted values: NF = normal food and CR = caloric restriction. Di-domain haemoglobin and erythrocruorin domains for transcript 12 are shown; arrows below indicate one erythrocruorin domain is missing in transcript 11 and none occur in transcript 21, although both still encode a di-domain haemoglobin like transcript 12.

Following from this result we found that one Hb trans-inducer HIF-1α (Dapma7bEVm009543) was up-regulated significantly in CR (log_2_-fold change 0) at a mean expression of 6903 TPM versus 4558 in NF. Three of the HIF-1 co-dimer HIF-1β genes were DE at this threshold, the highest expressed gene was down-regulated in CR at a mean 2316 TPM versus 2667 in NF. The two other HIF-1β genes were up-regulated in CR, but had lower in average overall expression at 464 and 391 TPM respectively. An mTOR protein kinase (Dapma7bEVm000341) was also up-regulated under CR at log_2_-fold change 0 albeit with a modest log_2_-fold change overall (0.14) and mean expression of 2434 TPM.

### Gene ontology and gene set enrichment analyses

For molecular function GO term enrichment approximately equal numbers of terms at 20 and 21 significant terms up- and down-regulated at log_2_-fold change 0 respectively (Supplementary Table S6). Biological process GO terms differ in that 25 terms are enriched for genes down-regulated in CR at log_2_-fold change 0, versus 8 that were up-regulated (REVIGO clustering of GO terms, Figure 4 and Supplementary Table S7).

**Figure 4.**
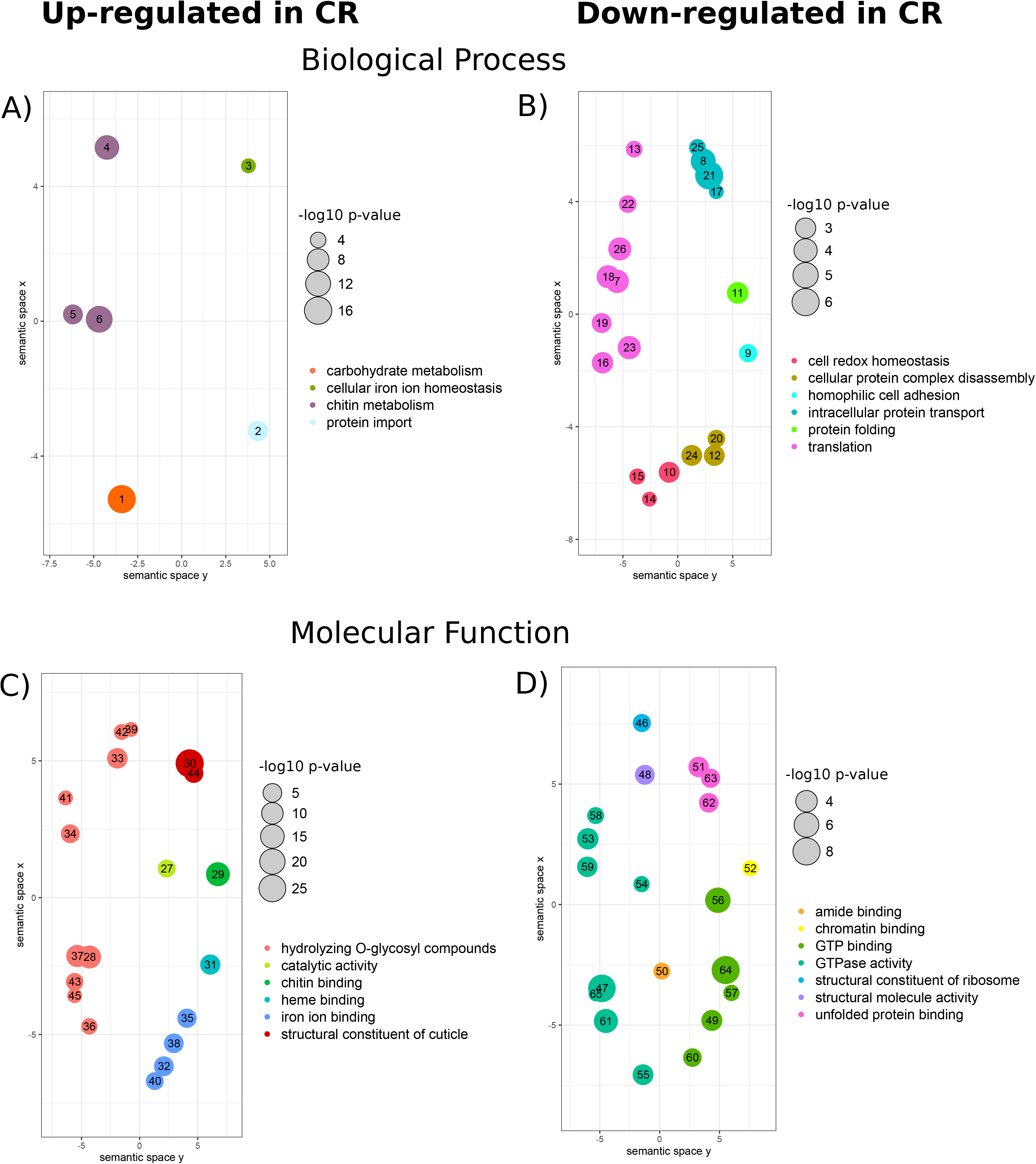
REVIGO scatter plots of enriched biological process GO terms for Caloric Restriction *versus* Normal Food. GO terms are grouped by REVIGO into broader categories indicated by colour and legend, Circle size is −log_10_ of the topGO enrichment p-value with scales inset next to each plot. A) Higher expression in CR Biological Process GO terms, B) Higher expression in NF Biological Process GO terms C) Higher expression in CR molecular function GO terms C) Higher expression in CR molecular function GO terms, D) Higher expression in NF molecular function GO terms. GO terms associated with numbers can be found in Supplementray Table S10.

For the GSEA analysis of KEGG terms, five of the six enriched terms that occur in at least 95% of 1000 iterations were groups of genes up-regulated under CR (representative GSEA result, Figure 5). The five KEGG terms significantly enriched in CR were for serine/threonine-protein kinase/endoribonuclease IRE1 (K08852), preprotein translocase subunit SecA (K03070), serine/threonine-protein kinase/endoribonuclease IRE2 (K11715), chitinase (K01183) and cytochrome P450 family 4 (K15001). The single term enriched under normal food was for apolipoprotein D and lipocalin family protein (K03098).

**Figure 5.**
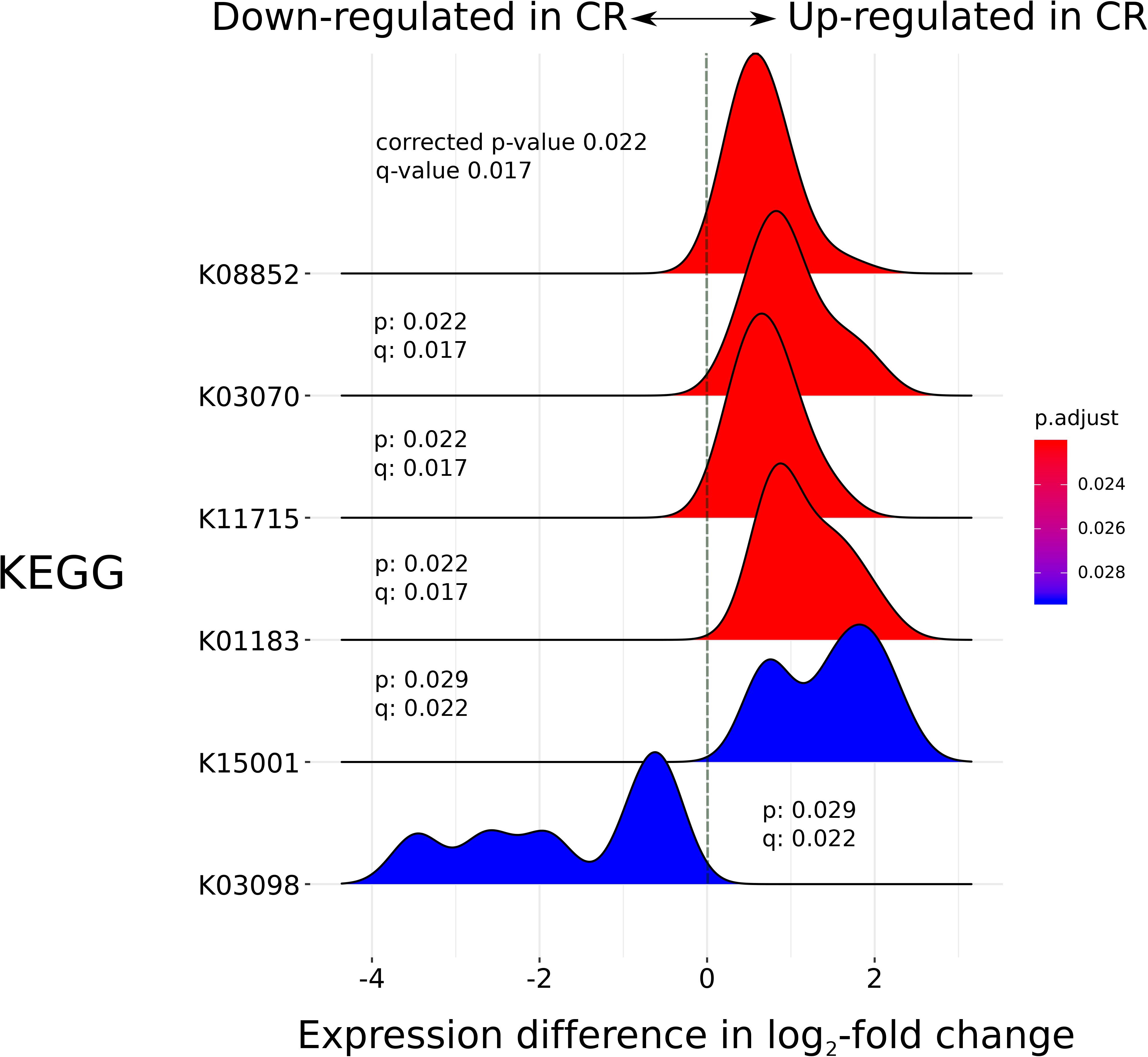
Gene Set Enrichment Analysis ridge plot for significant KEGG terms. Ridge plots are density plots of the frequency of log_2_ fold-change values per gene within each enriched KEGG group, which helps to interpret the up- or down-regulation of that KEGG category. The plot was created in clusterProfiler using KEGG orthologue annotations and log_2_-fold changes per gene calculated by DESeq2 during differential expression analysis. ×-axis is log_2_-fold change in expression for genes present in each KEGG category plotted, with positive values indicating increased expression in CR replicates and negative values in NF replicates. Peaks are coloured by corrected p-value as shown by the legend and corrected p-value and q-values are shown per KEGG category. KEGG term definitions: K08852=serine/threonine-protein kinase/endoribonuclease IRE1; K03070=preprotein translocase subunit secA; K11715=serine/threonine-protein kinase/endoribonuclease IRE2; K01183=chitinase; K15001=cytochrome P450 family 4; K03098=apolipoprotein D and lipocalin family protein.

### Overlap with differentially methylated regions

Over 2000 genes that met our criteria (i.e., expression greater than 10 and at least 5% methylation) were included in the correlations with CpG methylation rate for each of the NF and CR comparisons. Weak, significant and highly-concordant negative correlations were found for both comparisons for Kendall’s tau and Spearman’s rho (Supplementary File S5). Previously we identified 115 and 192 genes containing regions of hypo- or hypermethylated at CpG sites under CR respectively (Hearn, Pearson, et al., 2019). For both categories of methylation there was little signal of co-regulation with gene-expression. The majority of overlap occurs at the log_2_-fold 0 level for which thousands of genes were significantly up- and down-regulated under CR implying a high-degree of overlap by chance (Table 2). Only two genes up-regulated in CR with a log_2_-fold change of 1 or greater intersected with differentially methylated regions. One was an uncharacterised gene (Dapma7bEVm028334) which was identified as hypo-methylated under CR in Hearn, Pearson, et al., (2019), and the other an integral membrane protein (Dapma7bEVm027395) which was hyper-methylated under CR.

**Table 2.** Overlap between significantly expressed genes at log_2_-fold change 0 and hypo- or hyper-methylated genes. Percentages were calculated from total hypo- and hyper-methylated genes.

|  | Up-regulated in CR | Down-regulated in CR |
| --- | --- | --- |
| Hypo-methylated | 20 (10%) | 48 (25%) |
| Hyper-methylated | 16 (14%) | 28 (24%) |

### miRNA interaction

In total 240 miRNA-mRNA pairs were predicted by the combined target-site method, which was composed of 117 genes and 34 miRNAs (miRNA-mRNA pairs and miRbase homologs: Supplementary Table S8). Many miRNAs were predicted by our combined approach to target multiple genes, hence the discrepancy between number of genes and miRNAS. As for the expression analyses these miRNA-mRNA pairs showed a bias toward genes expressed more highly under CR (Table 3). The strongest negative correlation for genes up-regulated in CR was −0.71 and the strongest positive correlation was 0.87. There was little overlap between these genes and other genes of interest, but a Histone-lysine N-methyltransferase Suv4-20 (Dapma7bEVm018601, mean expression = 10277 TPM) up-regulated under CR had a negative correlation with nine miRNAs (Supplementary Table S8: gene description for gene numbers included in Table 3). Four of the genes exhibiting DTU were predicted to have miRNA targets, of these only a Para-nitrobenzyl esterase (Dapma7bEVm000405), was annotated.

**Table 3.** Overlap between significantly expressed genes at log_2_-fold change 0 (and log_2_-fold change 2) and predicted correlation in expression between miRNA-mRNA pairs. A positive correlation means miRNA-mRNA pairs were both highly or lowly expressed, and a negative correlation that they exhibit reciprocal expression. Annotations for the genes in each category are given in Supplementary Table S9.

|  | Up-regulated in CR | Down-regulated in CR |
| --- | --- | --- |
| **Pearson's r > 0.5** | 33 (6) | 5 (1) |
| **Pearson's r < -0.5** | 33 (7) | 4 (0) |

## Discussion

Firstly, We established that CR has an effect on average lifespan across eight different genotypes of *D. magna*, in line with previous results (Garbutt & Little, 2014, 2017; Latta IV et al., 2011). Focussing on Clone 32 we explored the molecular basis of CR through difference in gene expression with normal food levels. This difference was global, as over 6000 genes respond to treatment (Table 1. Log-fold change 0). Differential methylation identified previously in response to CR in Clone 32 (Hearn, Pearson, et al., 2019) did not impact upon gene expression. The abundance of several miRNAs that respond to CR in Clone 32 (Hearn et al., 2018) correlated with gene expression, and these correlations were biased towards higher expression in the CR treatment. Generally, we showed that more genes are significantly up-regulated under CR (Table 1) and for the remainder of the discussion we focus on the responses of particular genes and pathways to CR.

### A haemoglobin response to CR in *D. magna*

Fewer genes had differential transcript usage detected than differential gene expression. This may reflect biological reality as the eight replicates per condition give reasonable power in DTU analyses (by comparison to simulated data in Love et al., 2018). Alternatively, this may be because we were conservative in our approach to filtering DTUs by only taking the union of DEXSeq and DRIMSeq. Gene isoforms may also be under-annotated, however the gene-set used as a reference results from a comprehensive bioinformatic construction after exposure of *D. magna* to twelve environmental stressors (Orsini et al., 2016).

DTU analysis identified a highly-expressed Hb gene that responded to CR by changing isoform proportions (Figure 3). We believe it is the first time haemoglobin genes have been implicated in CR in *D. magna*. This is concordant with the observed changing Hb protein levels in response to food (Fox et al., 1951). The gene involved, Dapma7bEVm014981, has many different isoforms with the three most highly expressed in this experiment exhibiting varied proportions of expression between CR and NF. Most strikingly, transcript 12, which encodes three erythrocruorin (IPR002336) domains, occurs at much higher expression in NF than CR (Figure 3). These erythrocruorin domain provides the scaffold for a Haem-binding pocket (http://www.ebi.ac.uk/interpro/entry/InterPro/IPR002336/) and hence oxygen affinity of the haemoglobin gene-product. Based on these results we hypothesise that *D. magna* moderates its haemoglobin mix in response to CR by reducing respiration leading to a lower proportion of haemoglobin containing erythrocruorin domains under CR. This is evidenced by a change in the proportions of expressed isoforms of gene Dapma7bEVm014981 under CR in favour of isoforms without (transcript 21) an erythrocruorin domain or two domains only (transcript 11).

Thus variation in erythrocruorin domains could underly the observed correlation between oxygen-binding capacity and structurally distinct Hb-gene isoforms (Zeis, 2020). Future studies of Hb response to environmental stressors in *Daphnia* should consider differential isoform as well as Hb gene-copy usage following our insights here. The Hb mix of *D. magna* is known to change in response to hypoxia due to the action of the dimeric hypoxia-inducible factor 1 (Cuenca Cambronero et al., 2018; Zeis, 2020). We identified that the HIF-1α component of the Hb gene transcription factor HIF-1 is up-regulated in CR, as are two HIF-1β genes. However, the most highly-expressed HIF-1β is down-regulated in CR. The discordant patterns of HIF-1 component expression suggests HIF-1 regulation is not straightforward. Indeed, in *C. elegans* HIF-1 can promote or limit longevity (Leiser & Kaeberlein, 2010). The involvement of HIF-1 suggests a previously unidentified overlap in hypoxia and CR response in *Daphnia*. This may explain the similar phenotypic effects on body size in offspring under hypoxia and CR, with little associated impact on reproduction of CR and hypoxia (Garbutt & Little, 2017; Hearn et al., 2018; Seidl et al., 2005). It also indicates that the mTOR pathway is active in the *D. magna* response to CR (Land & Tee, 2007), which was supported here by up-regulation under CR of an mTOR protein kinase (Dapma7bEVm000341). An alternative to the HIF-1 induction pathway exists and it is triggered by juvenile hormone (Rider, Gorr, Olmstead, Wasilak, & LeBlanc, 2005). We did not see evidence for juvenile hormone involvement in the gene expression results, although this would require hormone assays to discard rigorously.

### The endoplasmic reticulum stress response under CR

The serine/threonine-protein kinase/endoribonuclease inositol-requiring enzyme 1 KEGG category (IRE1, K08852) was up-regulated under CR in clusterProfiler GSEA analysis (Figure 5). IRE1 is a sensor protein in the unfolded protein response (UPR) that lowers stress in the endoplasmic reticulum. When activated it initiates a transcription factor (X-box binding protein 1) that up-regulates endoplasmic reticulum associated degradation genes (ERAD) (Calfon et al., 2002). HIF-1 mediated CR lifespan increases in *C. elegans* depends on IRE1 (Di Chen & Kapahi, 2009). Transient CR-derived stress in *C. elegans* larvae causes a robust IRE1-dependent UPR to be maintained into adulthood, which is an example of hormesis (Matai et al., 2019). The up-regulation of IRE1 reported here links the CR response in *D. magna* to protein homeostasis in the ER, the dysregulation of which is strongly linked to ageing in general (Brown et al., 2014; Chadwick & Lajoie, 2019; Cohen, Bieschke, Perciavalle, Kelly, & Dillin, 2006; Steinkraus et al., 2008). This gene has further roles in CR in other organisms. It regulates the increased usage of intestinal triacyglycerol in *Drosophila* which mediates the metabolic adaptation of midgut epithelium to CR (Luis et al., 2016). While in mice the upregulation of IRE1 in response to a reduced protein diet protects against cancer (Rubio-Patiño et al., 2018).

### Superoxide dismutase was down-regulated under CR

Several of the well-known candidate response genes to CR were significantly up-regulated under CR, including sirtuins, IGF, and the mTOR protein kinase. None of these genes showed large differences in their mean expression and log_2_-fold changes were modest.

Copper-zinc super-oxide dismutase (SOD) was down-regulated under CR in *D. magna,* it also showed varying responses to CR (or links to longevity) in other systems such as yeast (Mesquita et al., 2010). Over-expression of copper-zinc and manganese SOD in *Drosophila* did not increase lifespan (Orr, Mockett, Benes, & Sohal, 2003), and in termites increased longevity of queens was associated with enzyme activity and not expression level (Tasaki, Kobayashi, Matsuura, & Iuchi, 2018). The effect of SOD disruption on lifespan varies by experimental context in *Drosophila* (Wang, Branicky, Noë, & Hekimi, 2018). In *D. magna* copper-zinc SOD is known to increase in expression in response to copper, ammonia, and hypoxia levels as they are considered to be general stress-response factors (Lyu, Zhu, Wang, Chen, & Yang, 2013). We hypothesise that up-regulated SOD production under NF was due to greater ROS production from higher-respiration levels than CR which was compensated for by greater SOD expression. An alternative explanation is that dissolved oxygen content was lowered in NF rearing jars by increased respiration-levels versus CR jars leading to a hypoxia-induced stress-response known to occur in *D. magna* and *D. pulex* in low oxygen conditions (Klumpen et al., 2017; Lyu et al., 2013). This could also explain the Dapma7bEVm014981 Hb gene expressing differential transcripts in response to hypoxia through oxygen-depletion in NF versus CR.

### Several processes were repressed under CR

Guanosine triphosphate (GTP) related molecular function GO terms dominated enrichment in genes down-regulated under CR. The down-regulation of a GTPase has a key role in life-extension due to CR in *C. elegans* (Hansen et al., 2008). This is hypothesized to stimulate recycling organelles and cytoplasmic proteins (autophagy), which promotes increased lifespans through down-stream mechanisms (Hansen et al., 2008).

Apolipoprotein D and other lipocalin genes (K03098) were down-regulated under CR. In *Drosophila* overexpression of Apolipoprotein D increased lifespan by 18% and tolerance to starvation, and could act as a scavenger of toxic products in defence against oxidative stress (Sanchez et al., 2006; Walker, Muffat, Rundel, & Benzer, 2006). Why this class of genes would be reduced in expression under CR in *D. magna* rather than up-regulated is unclear. Perhaps NF individuals are more developmentally advanced due to their rapid growth than CR individuals and are combating the effects of accumulated oxidative stress earlier in life.

### A weak correlation between gene methylation and expression

We observed a negative correlation between genes with a methylated CpG portion greater than 5% and gene expression. The highly concordant results for CR and NF comparisons indicate there is no effect of CR on this relationship. In *D. magna,* links between methylation and alternative splicing have been observed previously (Asselman et al., 2017; Kvist et al., 2018) and DNA methylation is enriched in gene-bodies. We saw no overlap between the DTU analysis and previously identified genes overlapping differentially methylated regions (DMRs), which is perhaps not surprising given the expression-methylation correlations between CR and NF expression and CpG methylation are essentially the same. These results contrast with (Kvist et al., 2018) in which methylation at exons two to four exhibits significant positive correlations with gene expression in *D. magna*. Our focus on the response to a treatment differs form Kvist et al., (2018) which defined the evolutionary conservation of methylation across gene bodies in *D. magna*.

### miRNA correlations are biased towards CR

A robust miRNA response was observed to CR in Hearn et al., (2018) originating from the same total RNA samples included in this study. Here we observed greater miRNA targeting of genes up-regulated in CR at log_2_-fold change 0 for both positive and negative correlations (33 in each case, Table 3). This is in keeping with the general bias towards up-regulation in CR across this experiment; we do not see overlap in the miRNA correlated gene-lists with Hb-related genes or SOD discussed above. Of note, there is a negative correlation between nine miRNAs and Histone-lysine N-methyltransferase Suv4-20 (Dapma7bEVm018601). This gene is significantly up-regulated at log_2_-fold change 1 under CR (average TPM 15063 versus 5278 in NF). Suv4-20 trimethylates histone H4 lysine 20 and has an important role in DNA repair and genomic stability (Jørgensen, Schotta, & Sørensen, 2013). However, we must interpret the results with caution as computational miRNA-mRNA target inference is prone to false-positives (Fridrich, Hazan, & Moran, 2019; Pinzón et al., 2017). This is because animal miRNA seed binding is ‘wobbly’ and does not require perfect complementarity between miRNA and mRNA. Because of this, Fridrich et al., (2019) recommend biological interpretation only when further experimental support is available. This included when taking the overlap of multiple prediction programs as we have done.

Demonstrating that an mRNA is regulated by specific miRNAs will require the development of a crosslinking, ligation and sequencing of hybrids (CLASH) protocol for the *D. magna* Argonaute (AGO) proteins (Helwak, Kudla, Dudnakova, & Tollervey, 2013). The CLASH method isolates the AGO-miRNA-mRNA complexes that form during mRNA silencing by miRNA. Sequencing of the interacting RNA in the AGO protein can then be used to identify which miRNAs are bound to what mRNAs under the experimental condition surveyed. Even if the link between miRNA targeting and Suv4-20 is not borne out in future such experiments, the differential expression result indicates a potential link between CR, the histone epigenome, and DNA repair.

## Conclusions

We first showed that caloric restriction increases the lifespan of *D. magna* across multiple genotypes. We then chose a CR-responsive genotype to survey the transcriptome of young-adults after their first clutch and detected a number of canonical stress- and CR responses. By contrasting differential transcript usage between CR and NF we also showed that the haemoglobin isoform mix of a highly-expressed *D. magna* Hb gene is reduced for isoforms containing erythrocruorin domains. We speculate that this may reduce the overall respiration-levels of CR individuals and partially explain the observed increased lifespan under CR. Components of the transcription factor that controls Hb-gene transcription, HIF-1, are also differentially expressed linking the response to hypoxia with that for CR. An mTOR protein kinase is also differentially expressed and is known to modulate HIF-1. The mTOR pathway is implicated in CR responses of diverse organisms, which is also the case in *D. magn*a. Future work should test if respiration is depressed under CR through changes to the haemoglobin mix in *D. magna*, and if this response is conserved in non-aquatic organisms.

## Acknowledgements

This research was funded by the Wellcome Trust Institutional Strategic Support Fund (Round 2, University of Edinburgh). The funder was not involved in study design, and collection, analysis, or interpretation of data and in writing the manuscript.

## Data Accessibility

Raw read data generated for this study has been deposited in the European Nucleotide Archive, Bioproject PRJEB25137.

## Author Contributions

JH and TJL designed this research and wrote the paper. JH, TJL, PJW and JC performed research. PJW and JC maintained *D. magna* genotypes used in this study.

## Supplementary Tables

**Supplementary Table S1.** Raw and filtered reads per replicate included in this experiment.

**Supplementary Table S2.** DESeq2 output for significant genes at log_2_-fold change

Column 1 = gene name, column 2 = mean expression level of gene, column 3 = log in base two fold change, column 4 = log_2_-fold change standard error, column 5 = p-value, column 6 = false discovery rate adjusted p-value.

**Supplementary Table S3.** DESeq2 output for significant genes at log_2_-fold change

Column 1 = gene name, column 2 = mean expression level of gene, column 3 = log in base two fold change, column 4 = log_2_-fold change standard error, column 5 = s-value.

**Supplementary Table S4.** DESeq2 output for significant genes at log_2_-fold change

Column 1 = gene name, column 2 = mean expression level of gene, column 3 = log in base two fold change, column 4 = log_2_-fold change standard error, column 5 = s-value.

**Supplementary Table S5.** Gene annotations and expression levels in TPM for significant genes with higher expression under caloric restriction and higher under normal food respectively: 1 and 2 = genes expressed at greater than 10000 mean for log_2_-fold change 0; 3 and 4 = all expression levels for log_2_-fold change 1; 5 and 6 = all expression levels for log_2_-fold change 2.

**Supplementary Table S6.** Molecular function GO terms enriched in differentially expressed genes at each log_2_-fold change level after enrichment testing in topGO at alpha = 0.05. GO.ID = GO term identifier, Term = description of GO term, Annotated

= number of genes annotated with this term, Significant = number of significant genes for the specified contrast annotated with this term, Expected = expected number of significant genes for the specified contrast annotated with this term, weight01 = p-value for GO term.

**Supplementary Table S7.** Biological Process GO terms enriched in differentially expressed genes at each log_2_-fold change level after enrichment testing in topGO at alpha = 0.05. GO.ID = GO term identifier, Term = description of GO term, Annotated

= number of genes annotated with this term, Significant = number of significant genes for the specified contrast annotated with this term, Expected = expected number of significant genes for the specified contrast annotated with this term, weight01 = p-value for GO term.

**Supplementary Table S8.** mRNA-miRNA pairs predicted by at least four of five tools. mrna = *Daphnia* gene, mirna = *Daphnia* miRNA, microtar/miranda/pita/rna22/rnahybrid = target prediction tool, binding_sites = number of binding sites in 3' UTRs of mRNA, no_tools = number of tools that predicted a binding site, closest miRbase homolog = as identified in supplementary table S4 of Hearn et al (2018).

**Supplementary Table S9.** Gene descriptions for genes included in Table 3, annotations for genes in each of the four possible combinations are given. CR = caloric restriction.

**Supplementary Table S10.** GO terms for each numbered circle in Figure 4.

**Supplementary Files**

**Supplementary File S1**. R code used to analyse clone survival at each food level and model outputs in part 1, R code to identify differential gene expression using DESeq2 in part 2, and differential transcript usage using DEXSeq, DRIMSeq and stageR in part 3, topGo functional enrichment in part 4, and clusterProfiler gene set enrichment analysis in part 5, R code for creating REVIGO-derived scatter plots of enriched GO terms in part 6.

**Supplementary File S2.** Raw clone longevity data for all eight clones input into R survival analysis; R code given in Supplementary File S1, part 1.

**Supplementary File S3.** MultiQC results for FastQC read data-quality assessment, this file is in html format.

**Supplementary File S4**. DRIMSeq proportion plots for the 112 genes with significant differential transcript usage in common between DESeq and DRIMSeq. There are two plots per gene, one a barplot and the other a line plot.

**Supplementary File S5. Correlation of methylation proportion and mean expression for caloric restriction and normal food datasets.** Methylation data adapted from (Hearn, Pearson, et al., 2019). For CR and NF 2029 and 2033 genes respectively with a methylation rate greater than 5%, a TPM expression mean over 10 and non-overlapping genome annotation were included, and the Y-axis was limited to a TPM expression of 10000. Kendall’s tau and Spearman’s rho values are inset, Bonferroni-Holm corrected p-values for all correlations were highly significant at p < 1 × 10^−6^. The regression line in blue with grey confidence intervals is for Kendall’s tau correlations.

## Notes

### Competing Interest Statement

The authors have declared no competing interest.

