## Supplementary File 4 for "*Daphnia magna* modifies its gene expression extensively in response to caloric restriction revealing a novel effect on haemoglobin isoform preference"

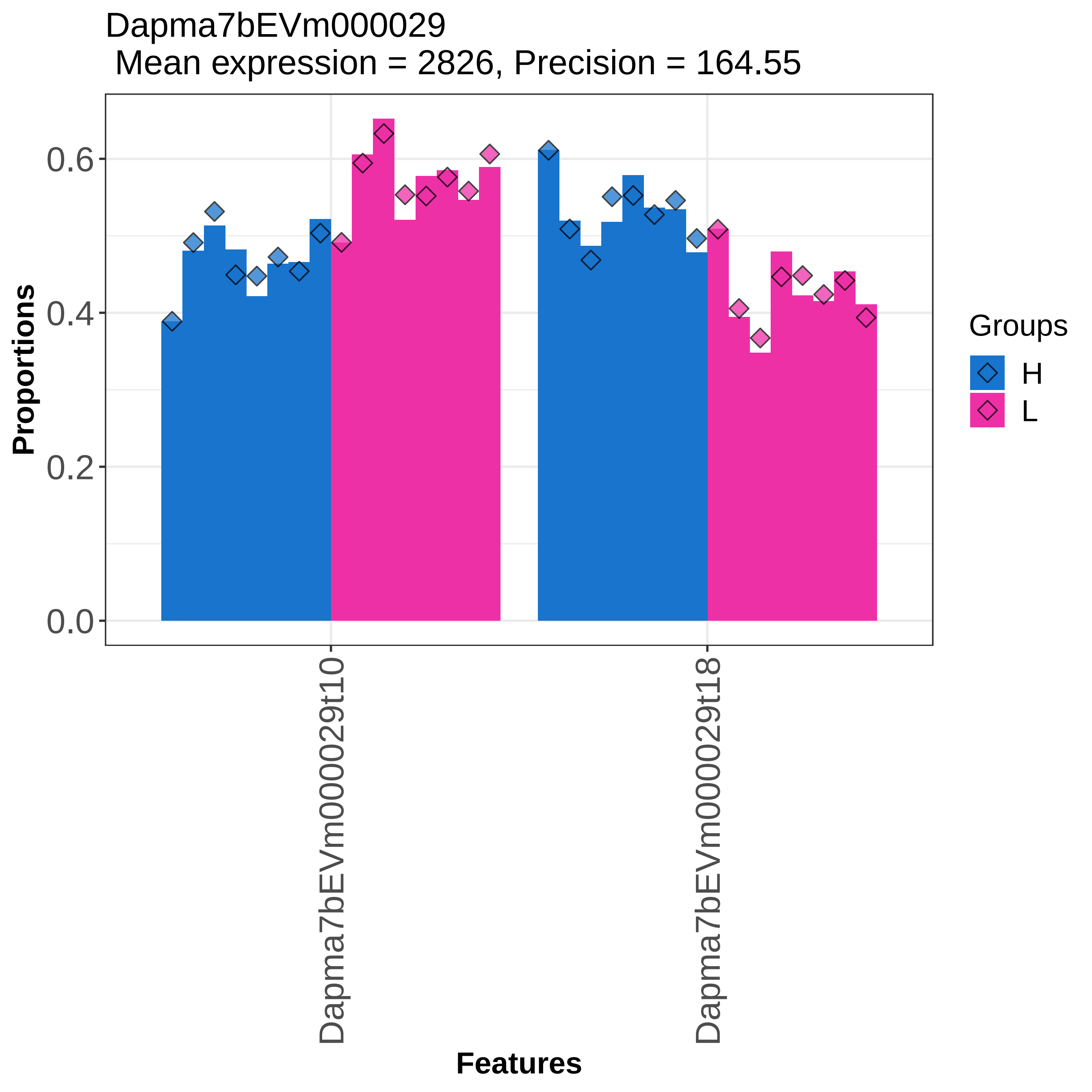
Dapma7bEVm000029
Mean expression = 2826, Precision = 164.55
Proportions
0.0
0.2
0.4
0.6
Dapma7bEVm000029t10
Dapma7bEVm000029t18
Features
Groups
H
L

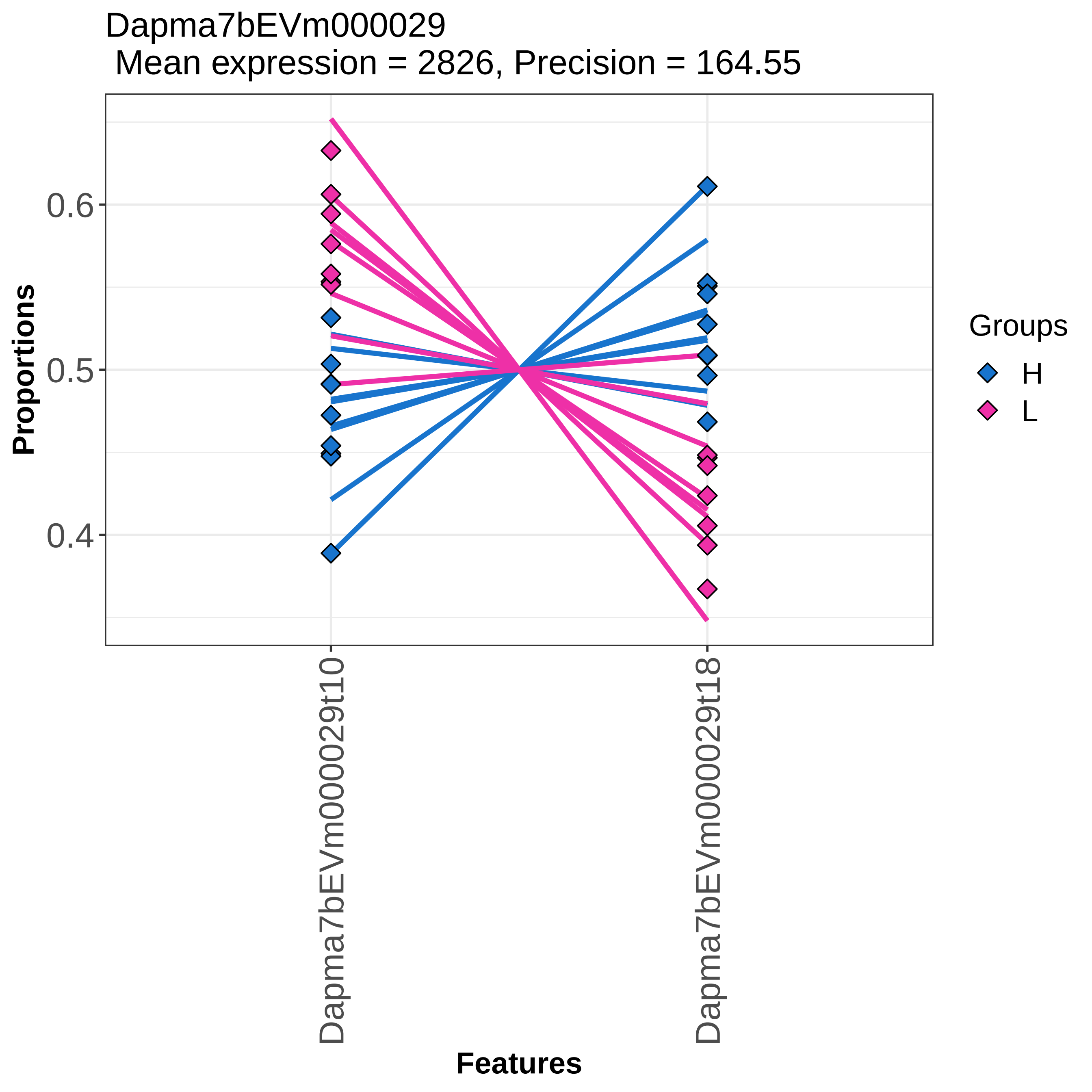

Dapma7bEVm000029
Mean expression = 2826, Precision = 164.55
Proportions
0.6
0.5
0.4
Dapma7bEVm000029t10
Dapma7bEVm000029t18
Features
Groups
H
L

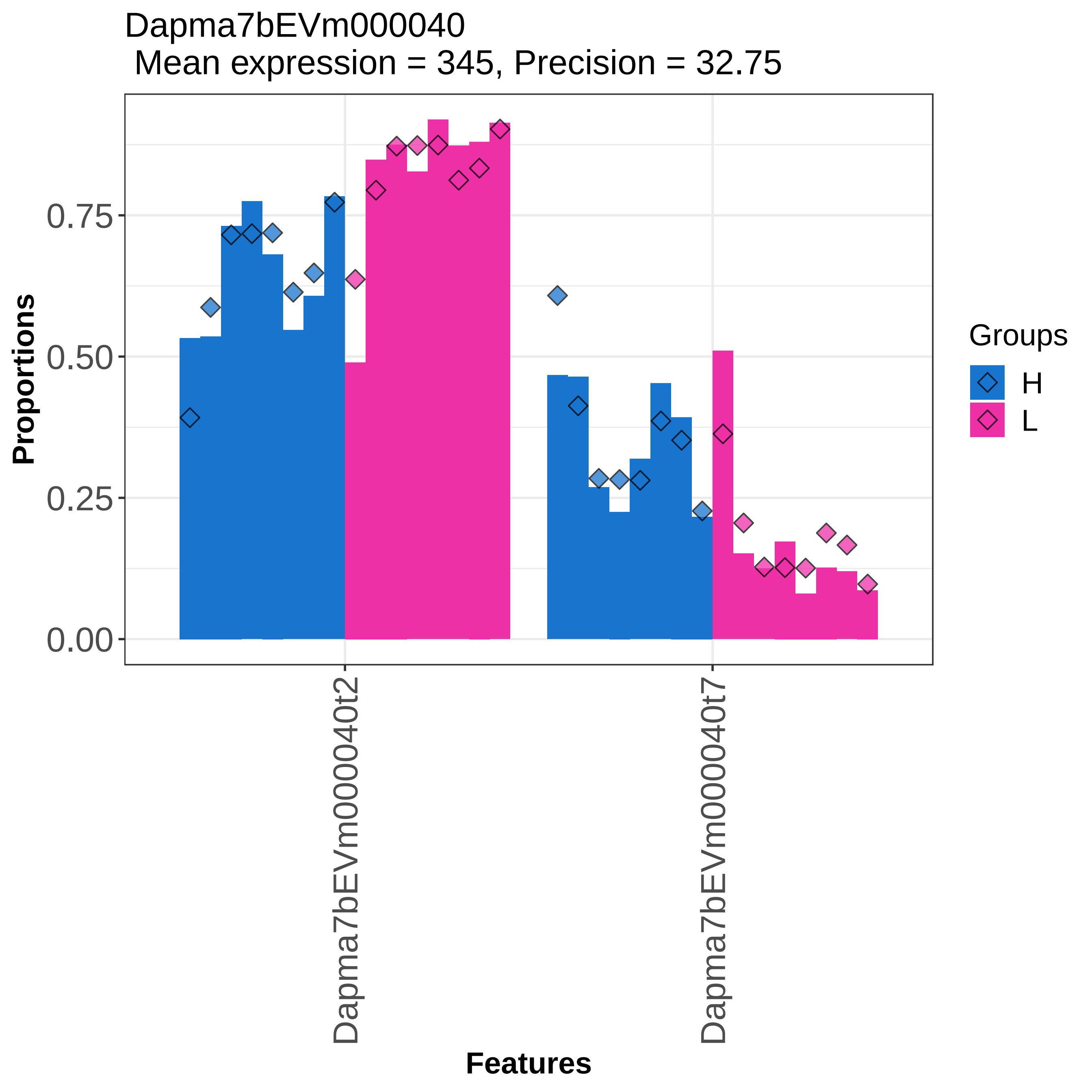

Dapma7bEVm000040
Mean expression = 345, Precision = 32.75
Proportions
0.00
0.25
0.50
0.75
Dapma7bEVm000040t2
Dapma7bEVm000040t7
Features
Groups
H
L

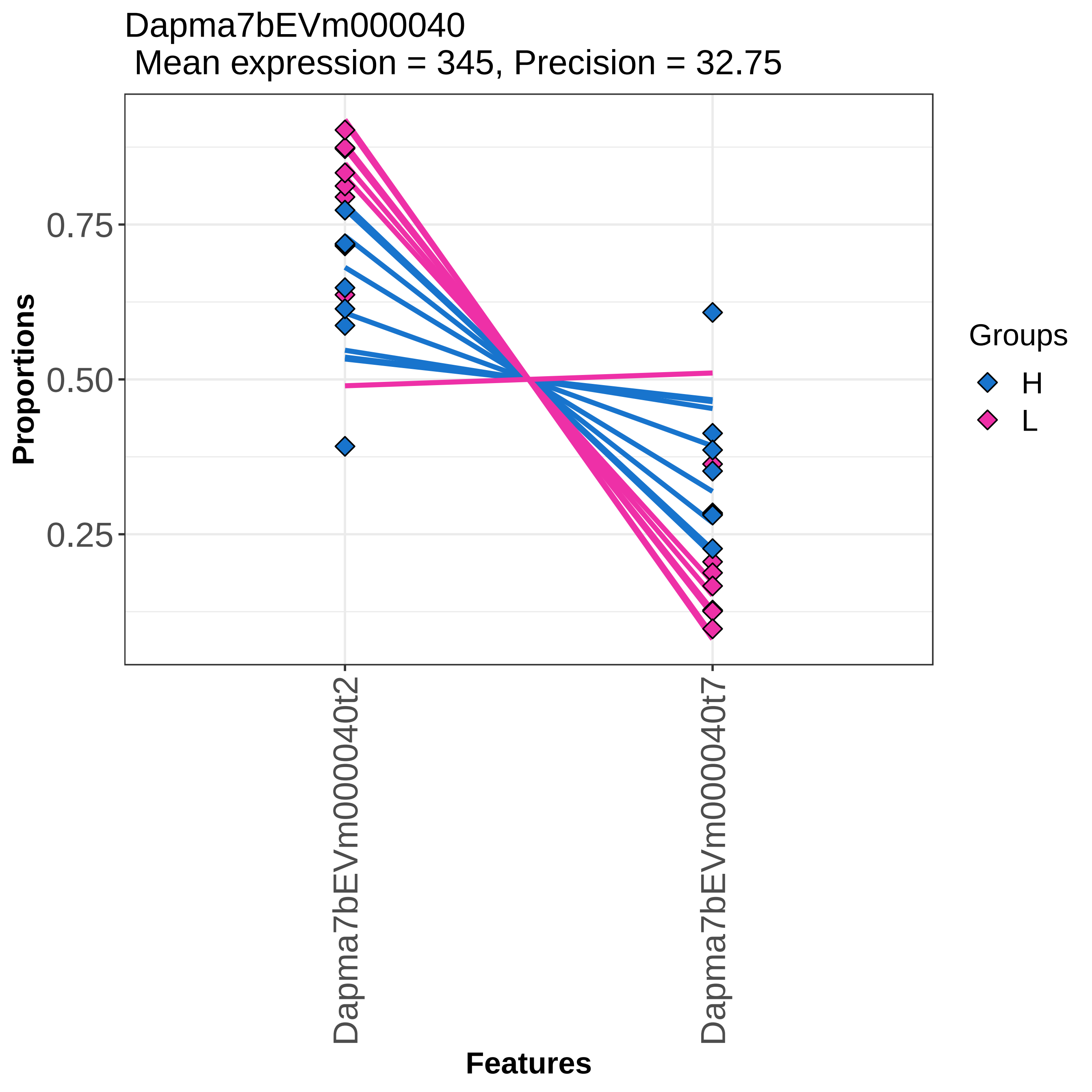
Dapma7bEVm000040
Mean expression = 345, Precision = 32.75
Proportions
0.75
0.50
0.25
Dapma7bEVm000040t2
Dapma7bEVm000040t7
Features
Groups
H
L

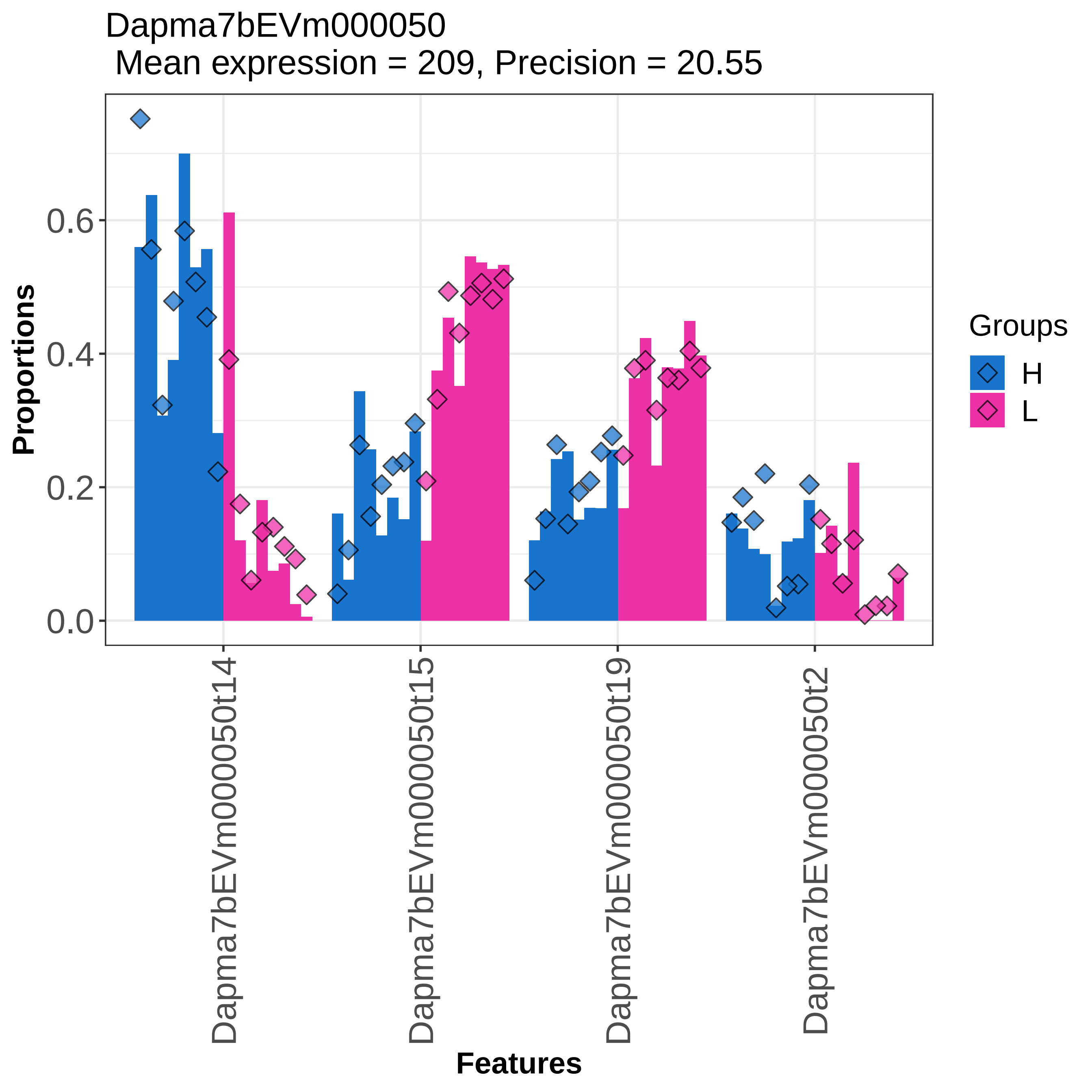

Dapma7bEVm000050
Mean expression = 209, Precision = 20.55
Proportions
0.0
0.2
0.4
0.6
Dapma7bEVm000050t14
Dapma7bEVm000050t15
Dapma7bEVm000050t19
Dapma7bEVm000050t2
Features
Groups
H
L

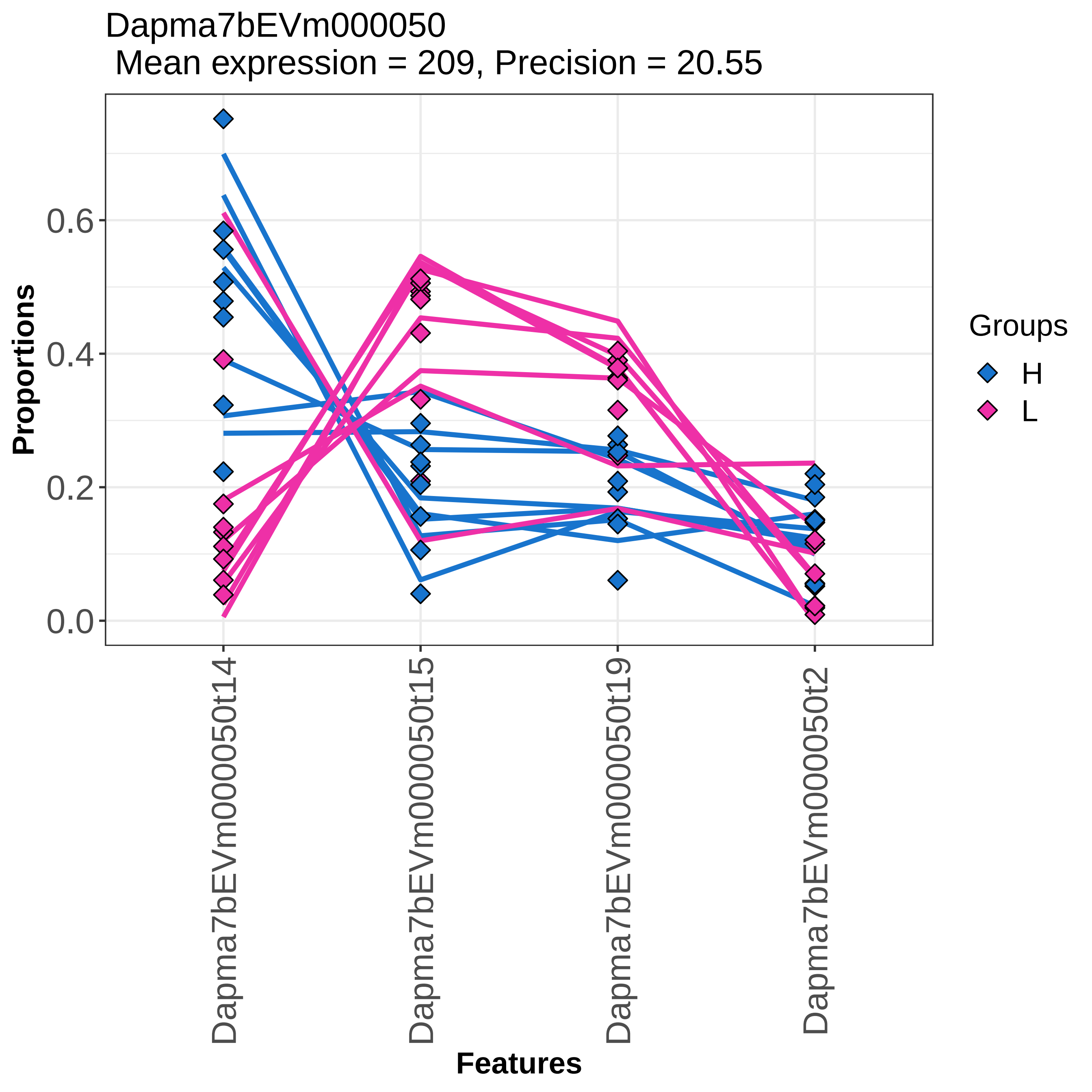
Dapma7bEVm000050
Mean expression = 209, Precision = 20.55
Proportions
0.0
0.2
0.4
0.6
Dapma7bEVm000050t14
Dapma7bEVm000050t15
Dapma7bEVm000050t19
Dapma7bEVm000050t2
Features
Groups
H
L

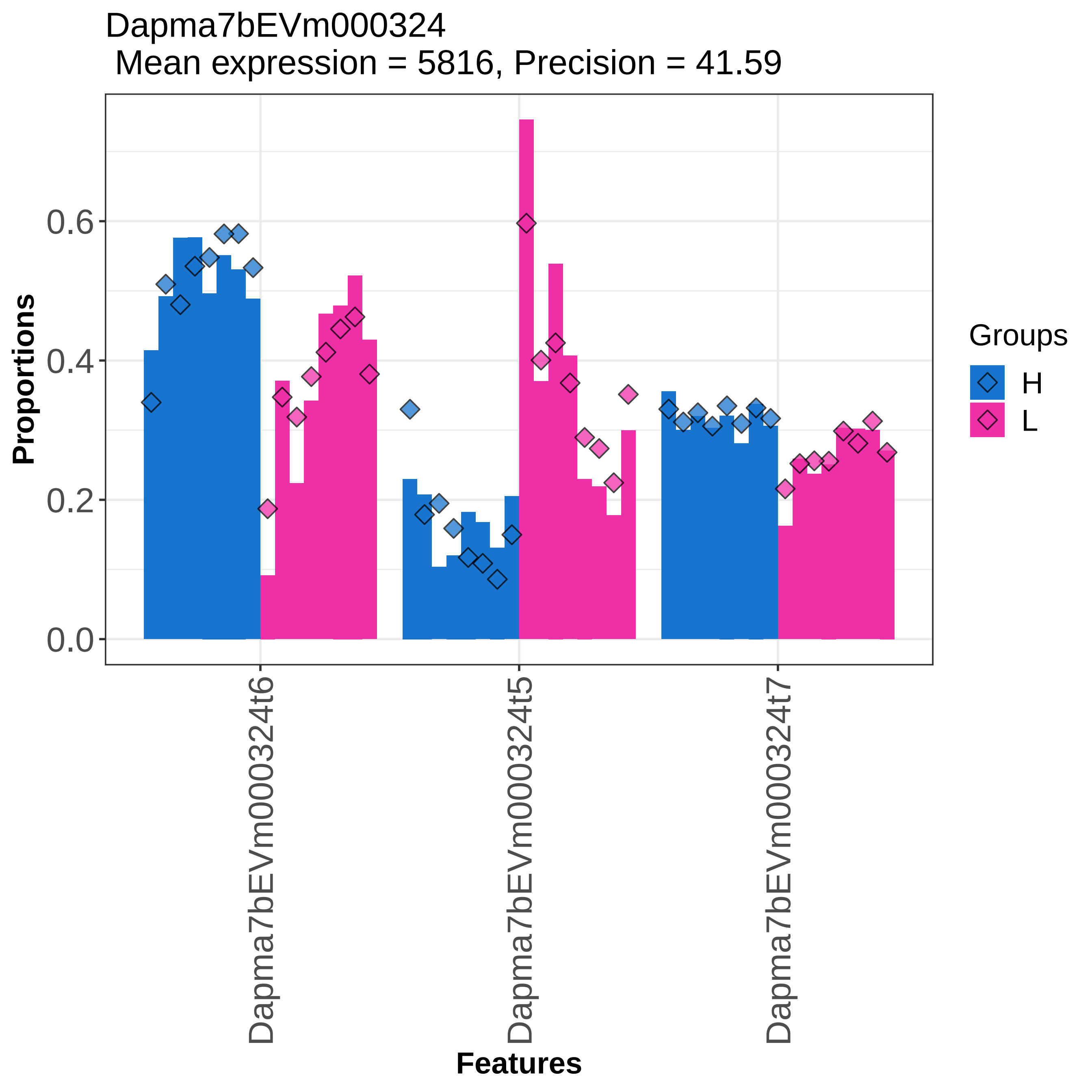

Dapma7bEVm000324
Mean expression = 5816, Precision = 41.59
Proportions
0.0
0.2
0.4
0.6
Dapma7bEVm000324t6
Dapma7bEVm000324t5
Dapma7bEVm000324t7
Features
Groups
H
L

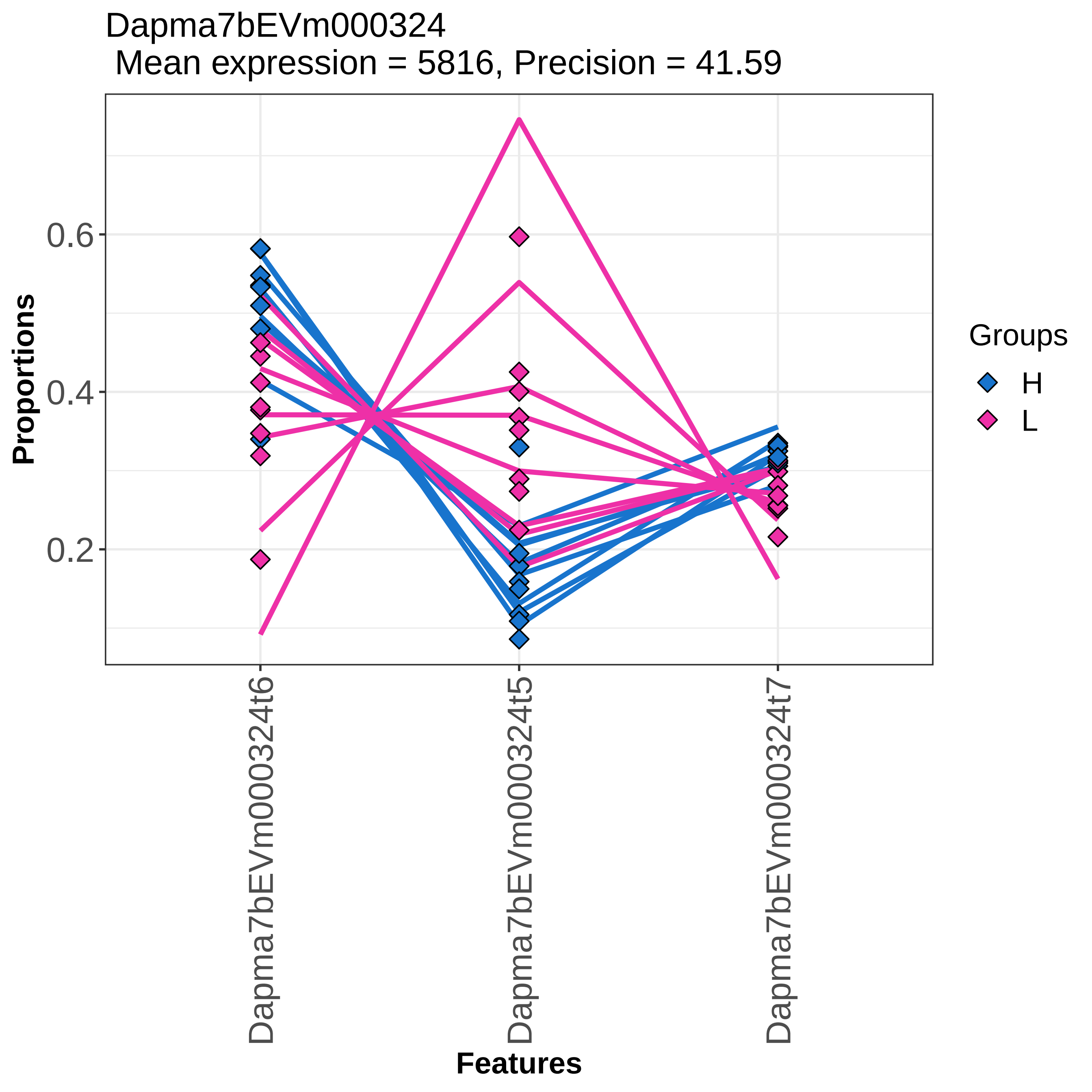

Dapma7bEVm000324
Mean expression = 5816, Precision = 41.59
Proportions
0.6
0.4
0.2
Dapma7bEVm000324t6
Dapma7bEVm000324t5
Dapma7bEVm000324t7
Features
Groups
H
L

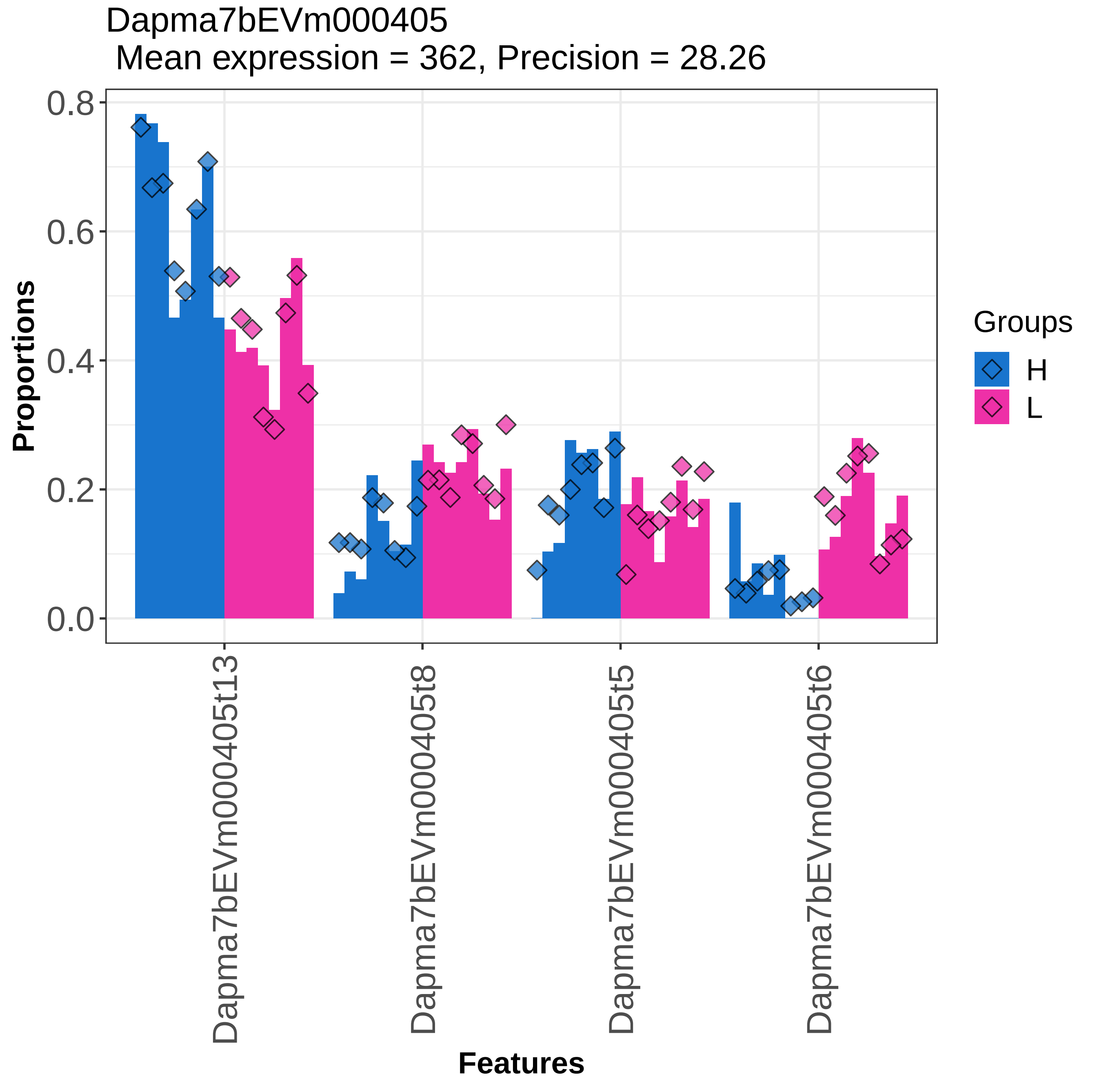

Dapma7bEVm000405
Mean expression = 362, Precision = 28.26
Proportions
0.0
0.2
0.4
0.6
0.8
Dapma7bEVm000405t13
Dapma7bEVm000405t8
Dapma7bEVm000405t5
Dapma7bEVm000405t6
Features
Groups
H
L

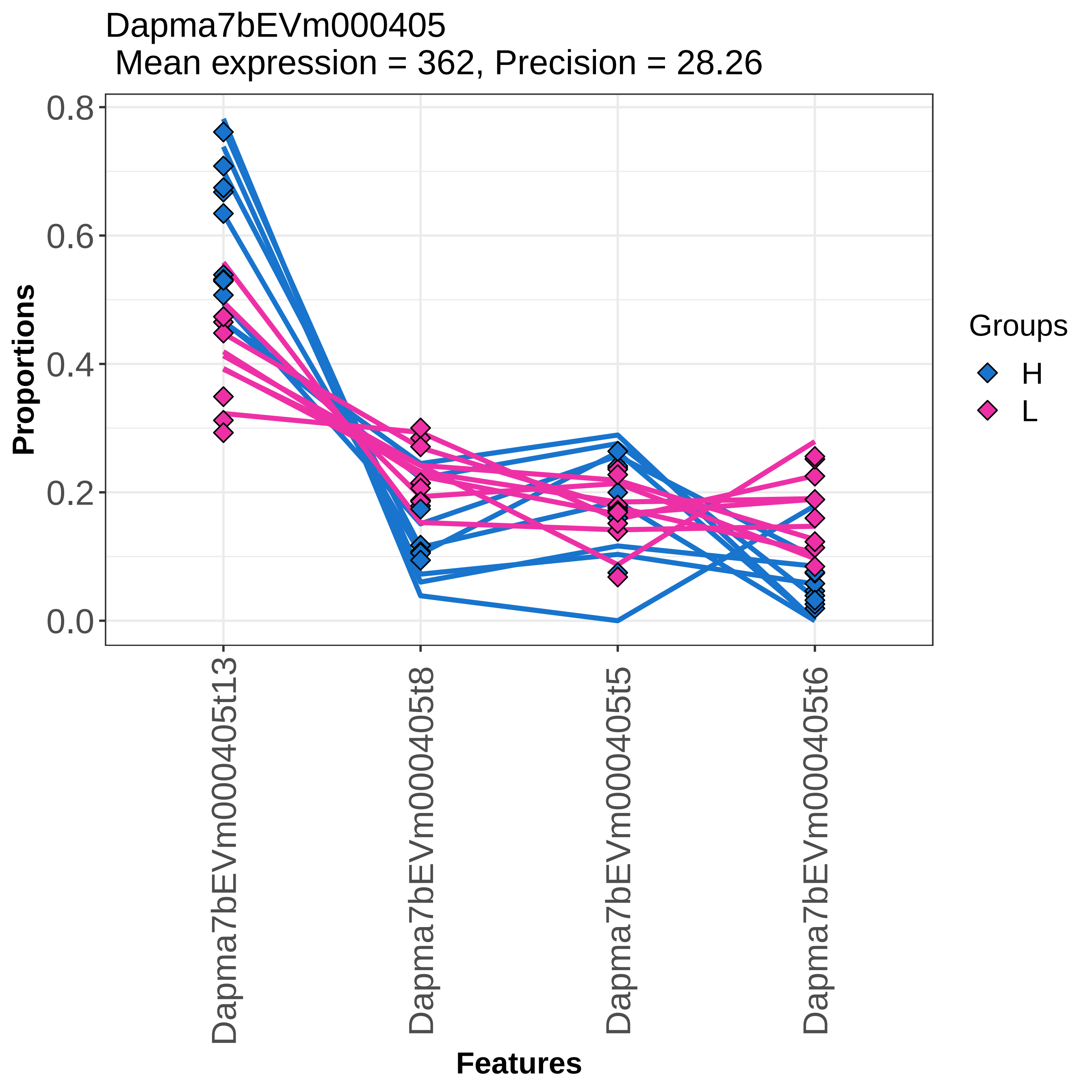

Dapma7bEVm000405
Mean expression = 362, Precision = 28.26
Proportions
0.8
0.6
0.4
0.2
0.0
Dapma7bEVm000405t13
Dapma7bEVm000405t8
Dapma7bEVm000405t5
Dapma7bEVm000405t6
Features
Groups
H
L

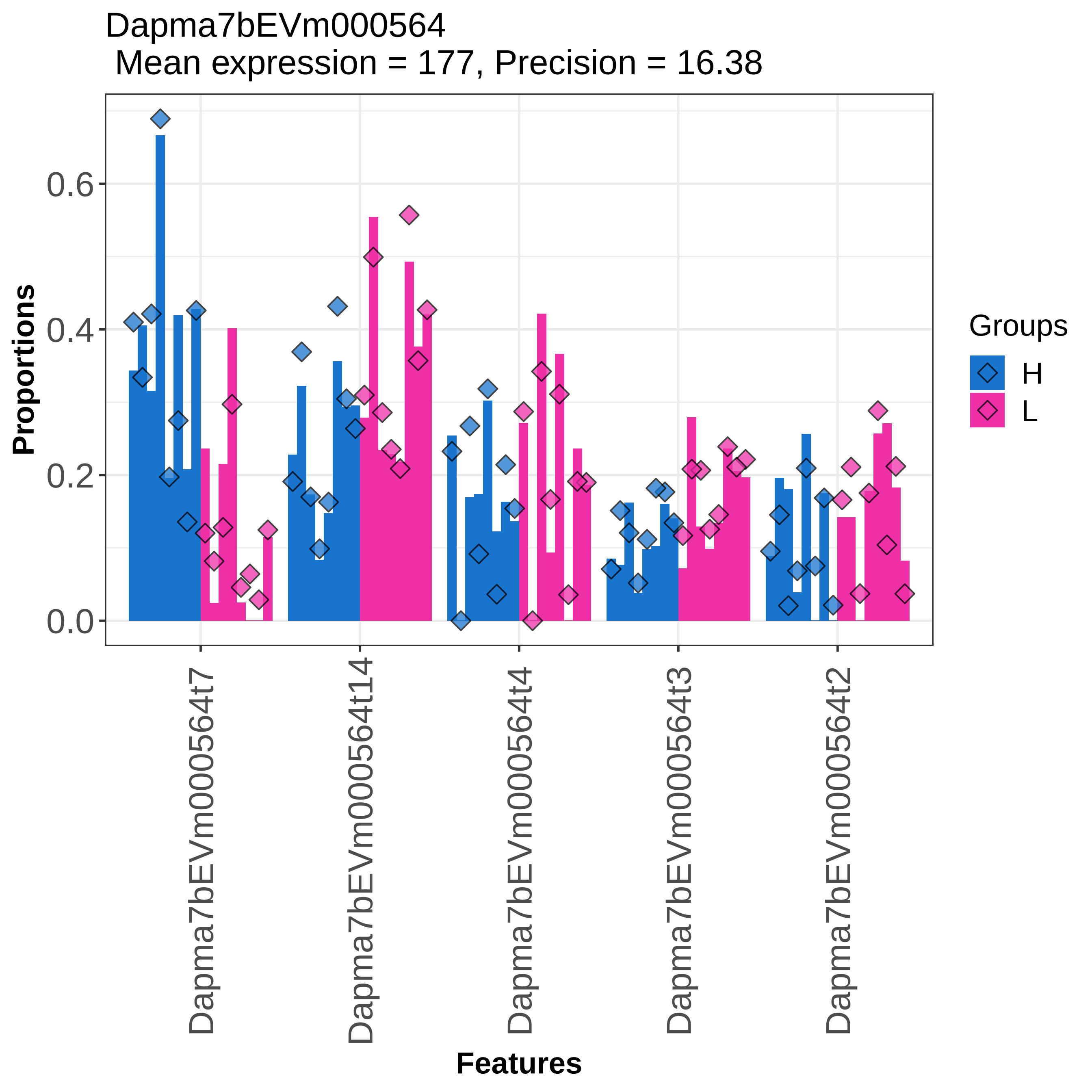

Dapma7bEVm000564
Mean expression = 177, Precision = 16.38
Proportions
0.0
0.2
0.4
0.6
Dapma7bEVm000564t7
Dapma7bEVm000564t14
Dapma7bEVm000564t4
Dapma7bEVm000564t3
Dapma7bEVm000564t2
Features
Groups
H
L

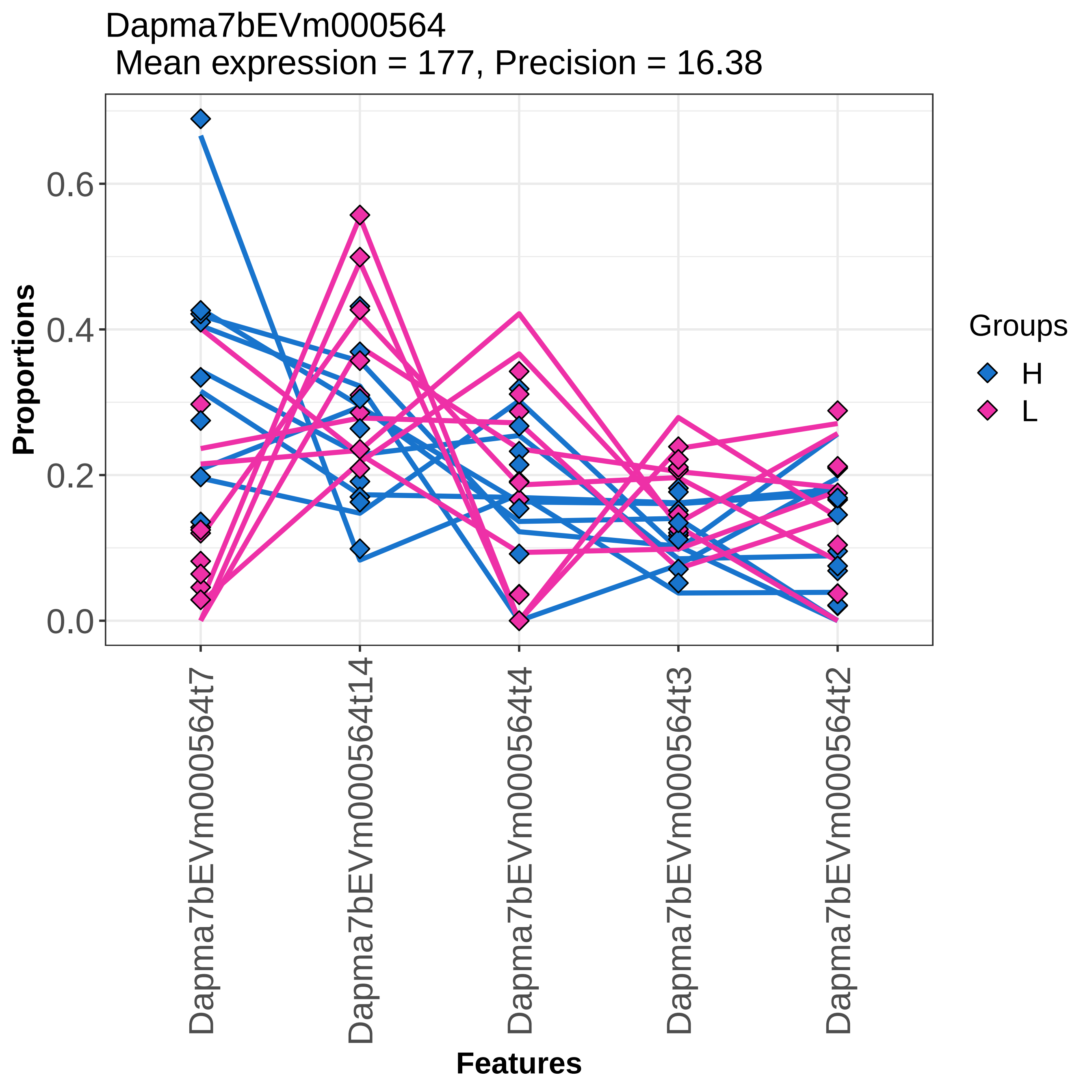
Dapma7bEVm000564
Mean expression = 177, Precision = 16.38
Proportions
0.0
0.2
0.4
0.6
Features
Dapma7bEVm000564t7
Dapma7bEVm000564t14
Dapma7bEVm000564t4
Dapma7bEVm000564t3
Dapma7bEVm000564t2
Groups
H
L

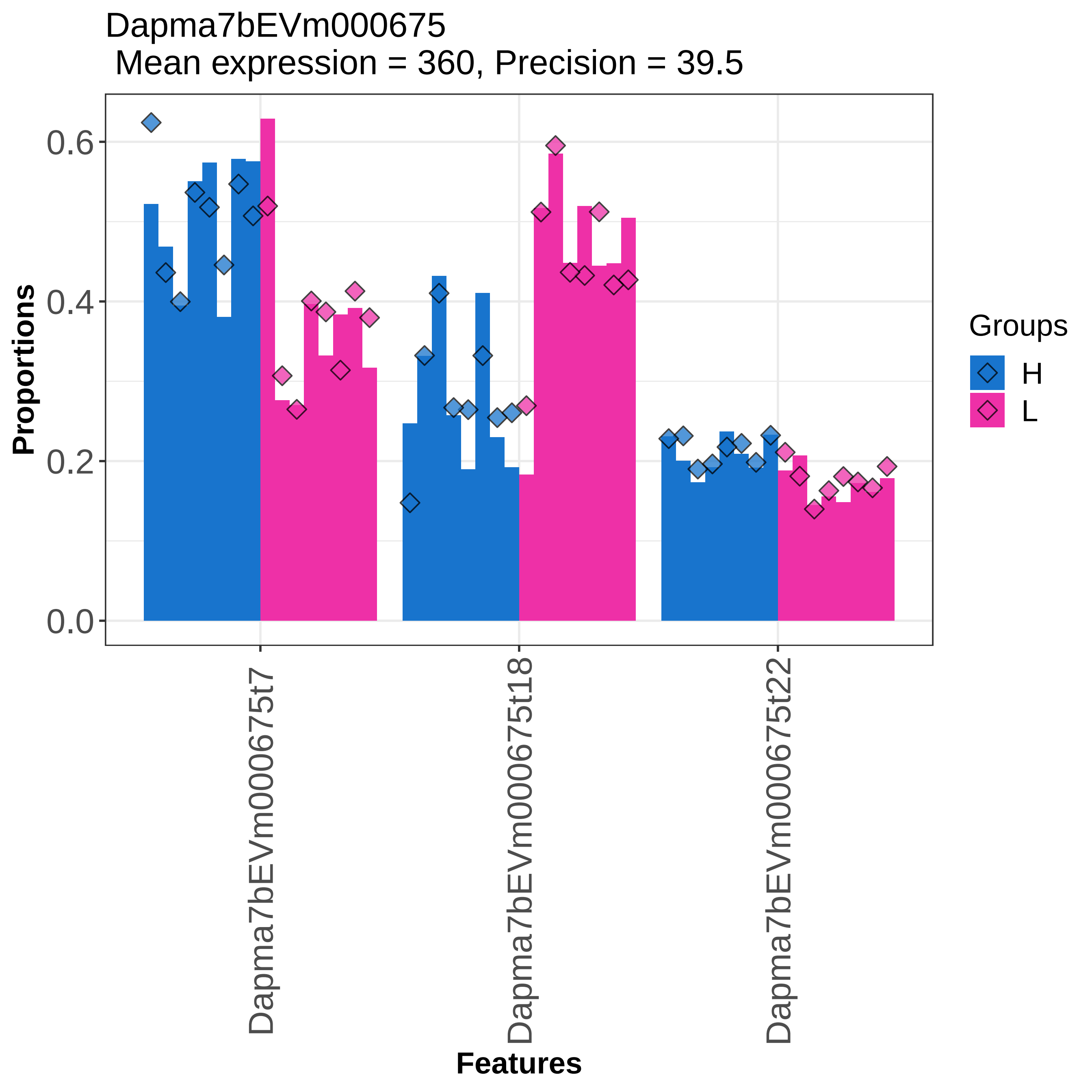
Dapma7bEVm000675
Mean expression = 360, Precision = 39.5
Proportions
0.0
0.2
0.4
0.6
Dapma7bEVm000675t7
Dapma7bEVm000675t18
Dapma7bEVm000675t22
Features
Groups
H
L

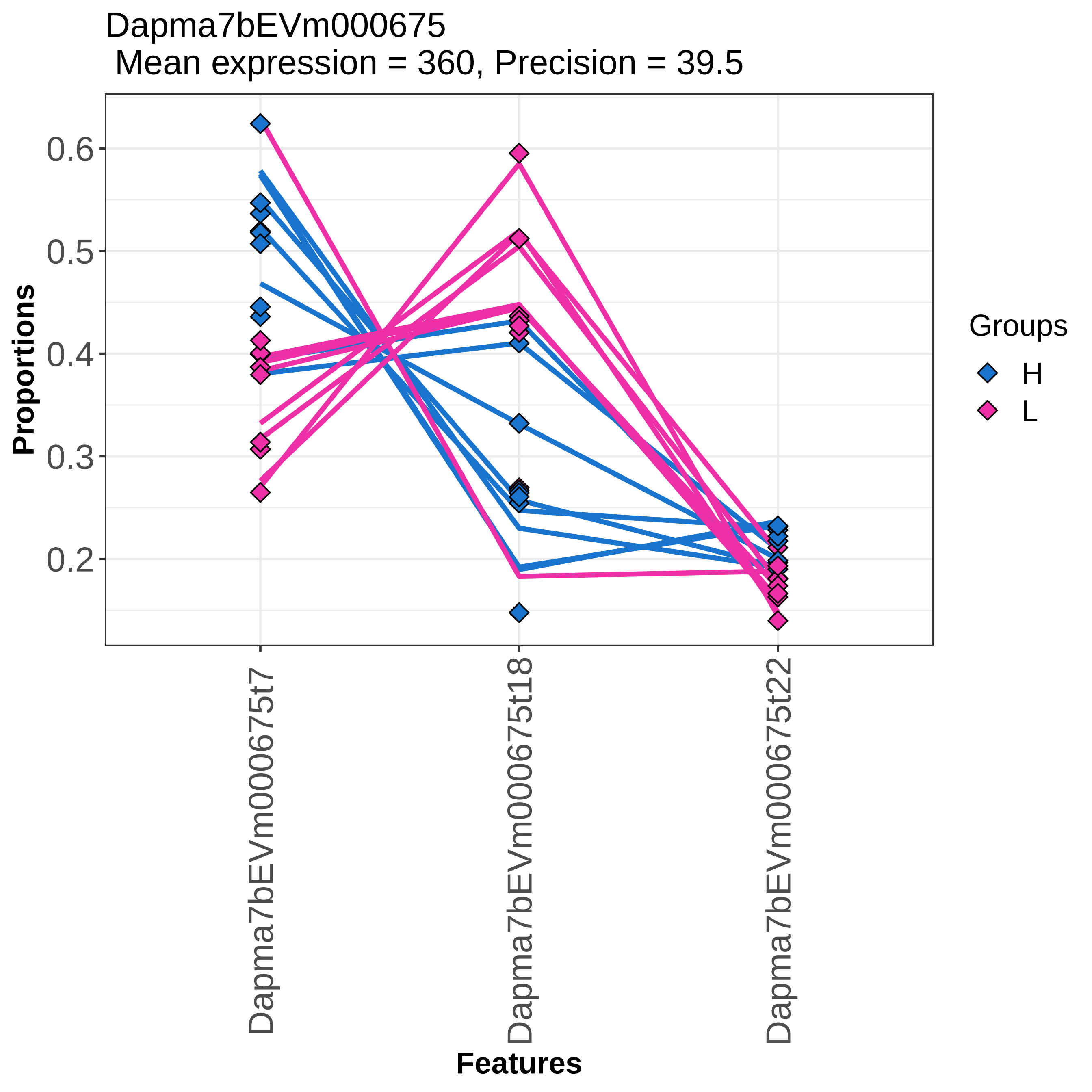

Dapma7bEVm000675
Mean expression = 360, Precision = 39.5
Proportions
0.6
0.5
0.4
0.3
0.2
Dapma7bEVm000675t7
Dapma7bEVm000675t18
Dapma7bEVm000675t22
Features
Groups
H
L

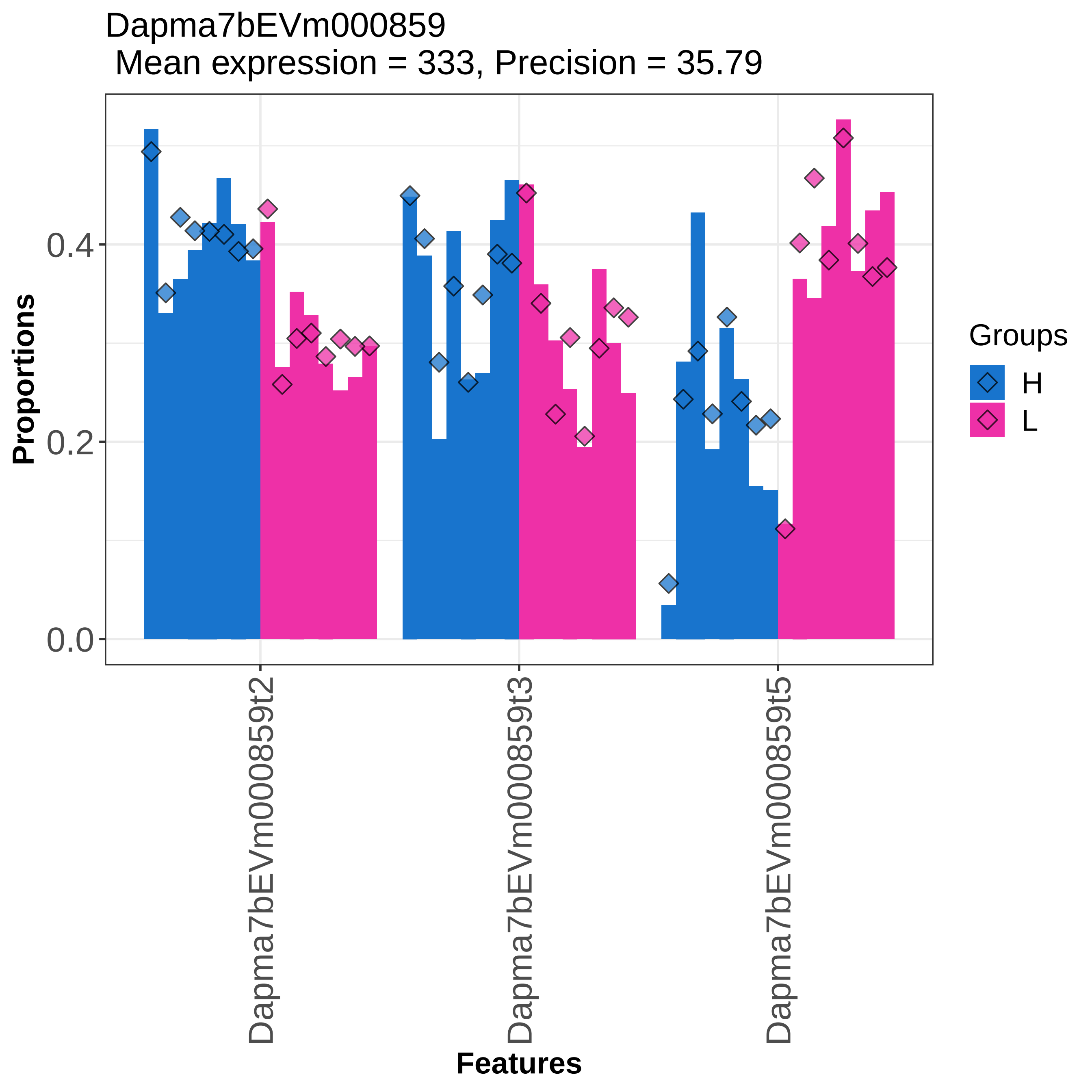
Dapma7bEVm000859
Mean expression = 333, Precision = 35.79
Proportions
0.0
0.2
0.4
Dapma7bEVm000859t2
Dapma7bEVm000859t3
Dapma7bEVm000859t5
Features
Groups
H
L

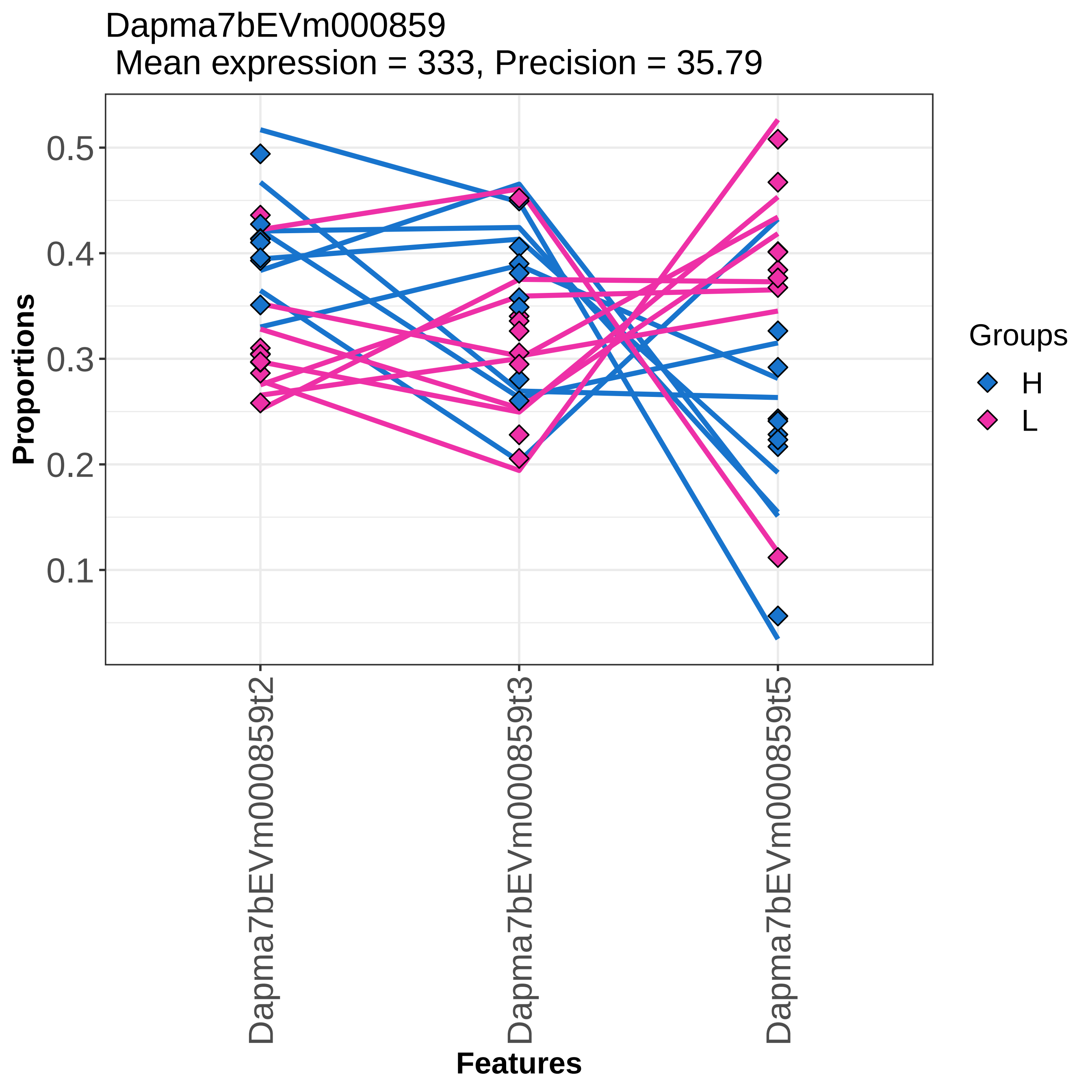

Dapma7bEVm000859
Mean expression = 333, Precision = 35.79
Proportions
0.5
0.4
0.3
0.2
0.1
Dapma7bEVm000859t2
Dapma7bEVm000859t3
Dapma7bEVm000859t5
Features
Groups
H
L

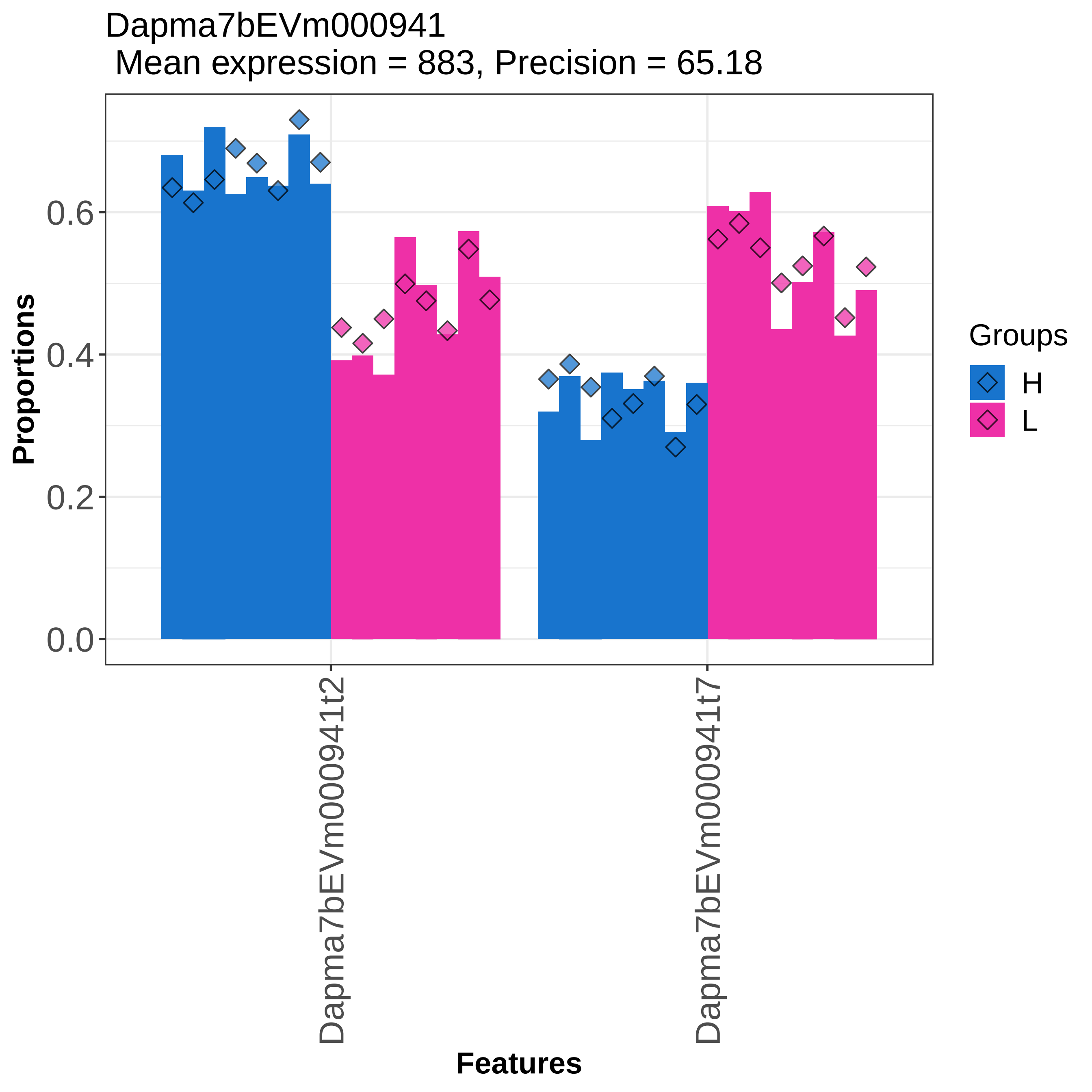

Dapma7bEVm000941
Mean expression = 883, Precision = 65.18
Proportions
0.0
0.2
0.4
0.6
Dapma7bEVm000941t2
Dapma7bEVm000941t7
Features
Groups
H
L

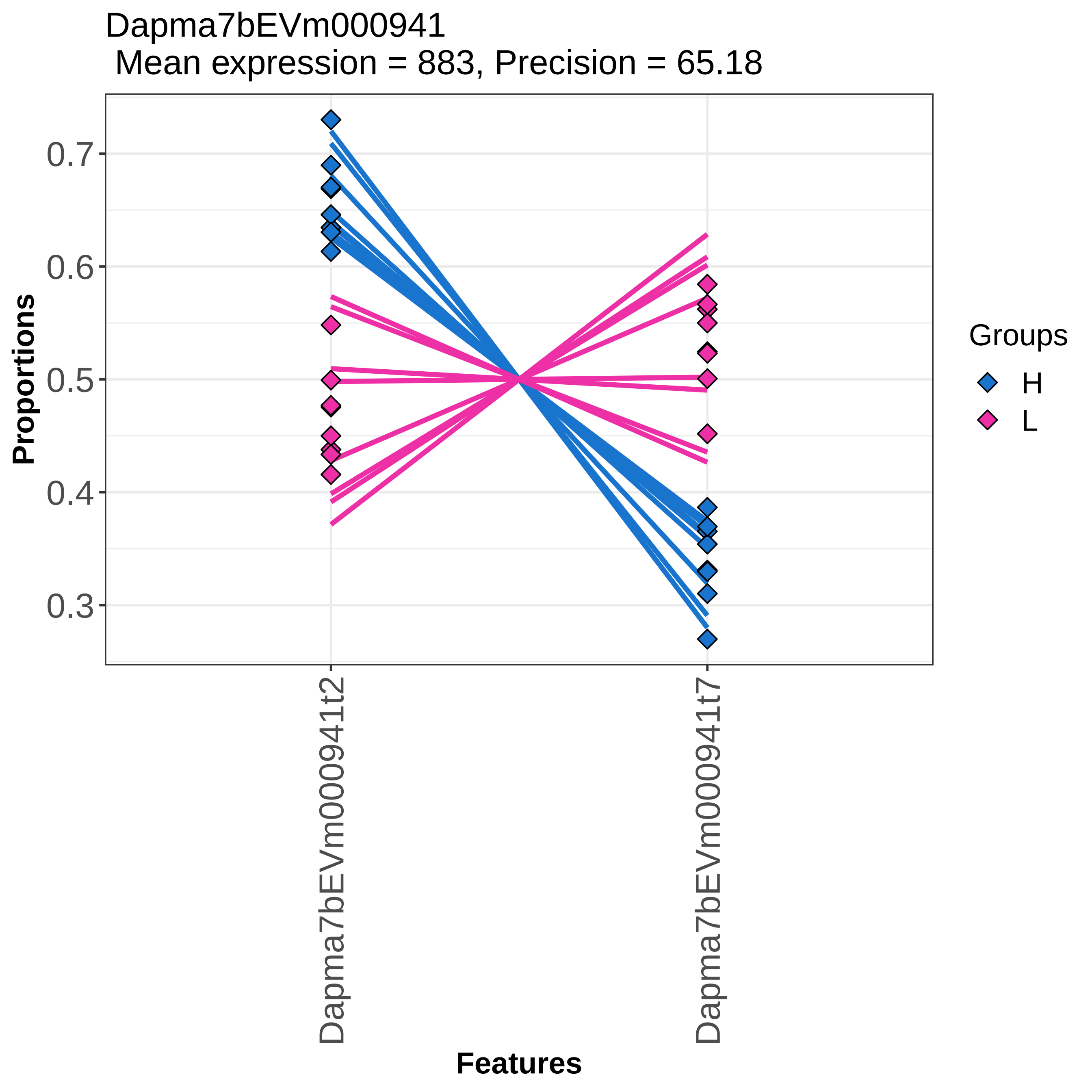

Dapma7bEVm000941
Mean expression = 883, Precision = 65.18
Proportions
0.7
0.6
0.5
0.4
0.3
Dapma7bEVm000941t2
Dapma7bEVm000941t7
Features
Groups
H
L

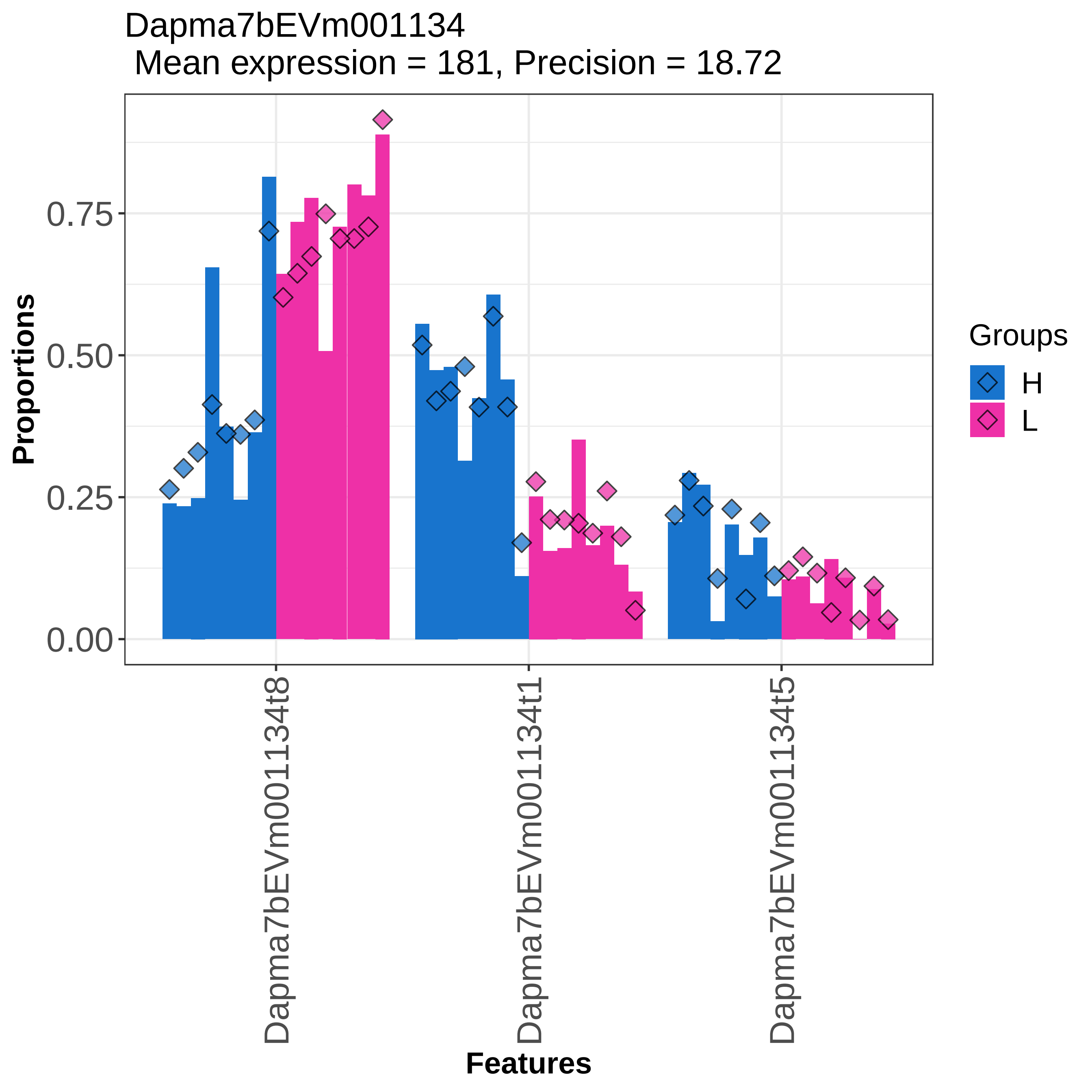

Dapma7bEVm001134
Mean expression = 181, Precision = 18.72
Proportions
0.00
0.25
0.50
0.75
Dapma7bEVm001134t8
Dapma7bEVm001134t1
Dapma7bEVm001134t5
Features
Groups
H
L

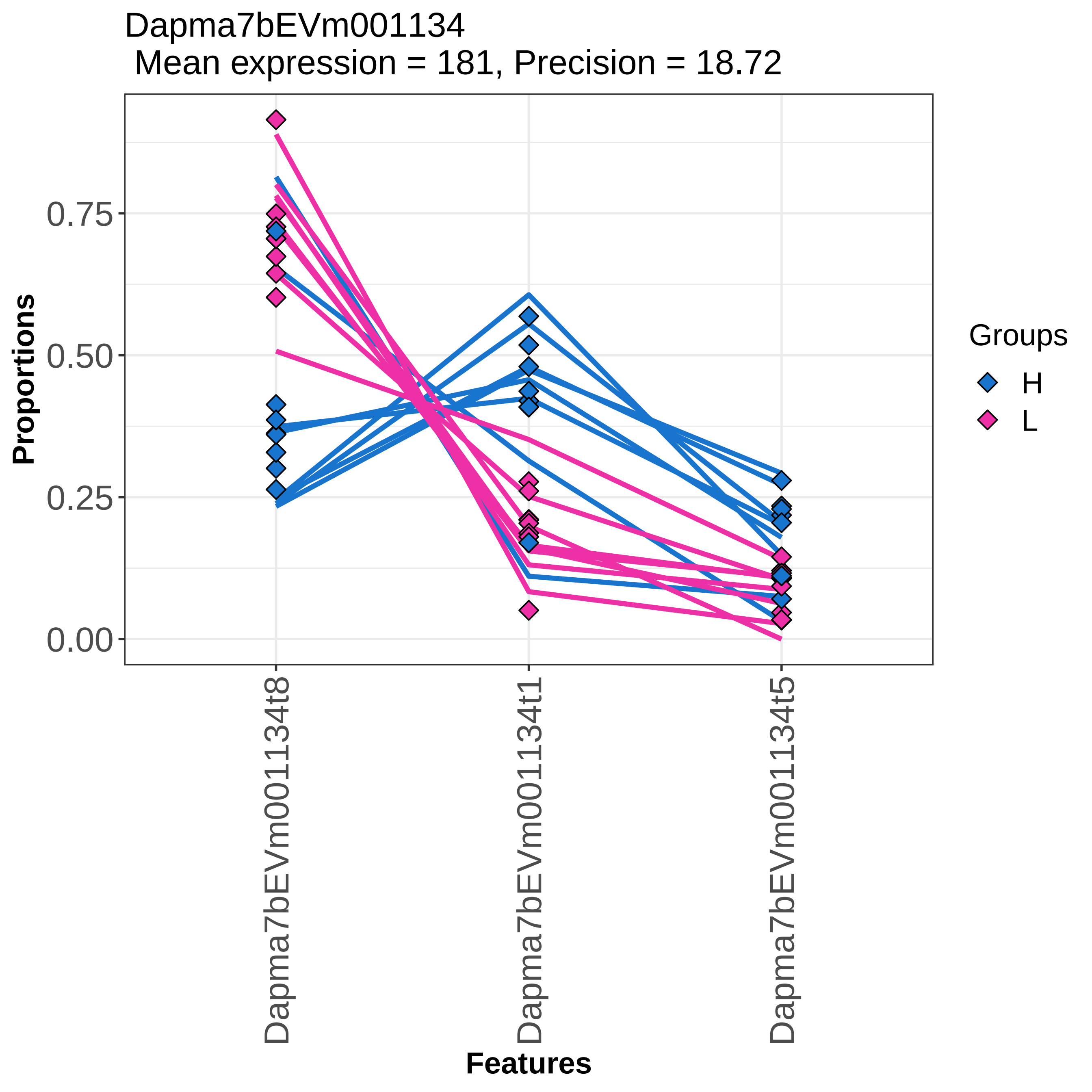

Dapma7bEVm001134
Mean expression = 181, Precision = 18.72
Proportions
0.75
0.50
0.25
0.00
Dapma7bEVm001134t8
Dapma7bEVm001134t1
Dapma7bEVm001134t5
Features
Groups
H
L

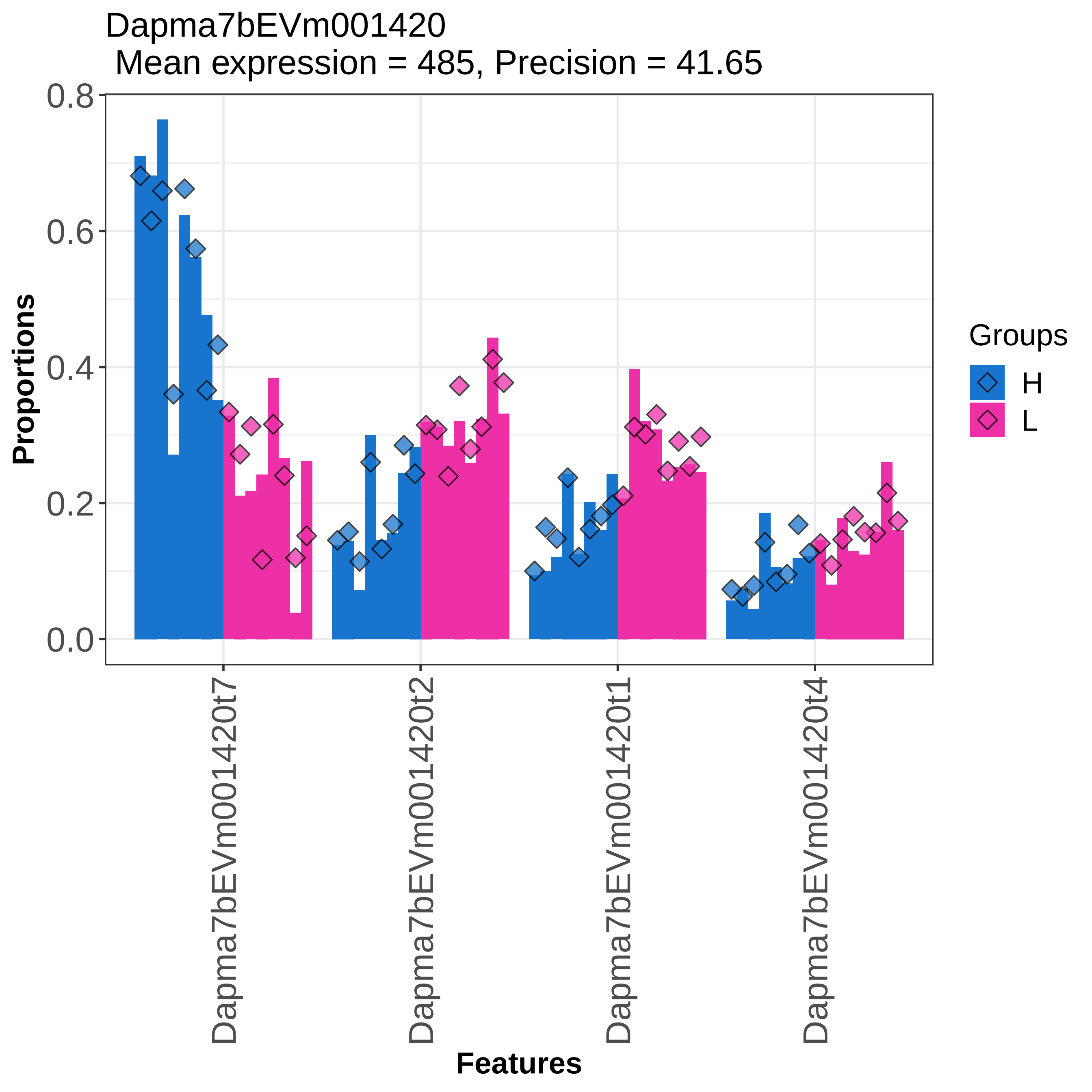
Dapma7bEVm001420
Mean expression = 485, Precision = 41.65
Proportions
0.8
0.6
0.4
0.2
0.0
Dapma7bEVm001420t7
Dapma7bEVm001420t2
Dapma7bEVm001420t1
Dapma7bEVm001420t4
Features
Groups
H
L

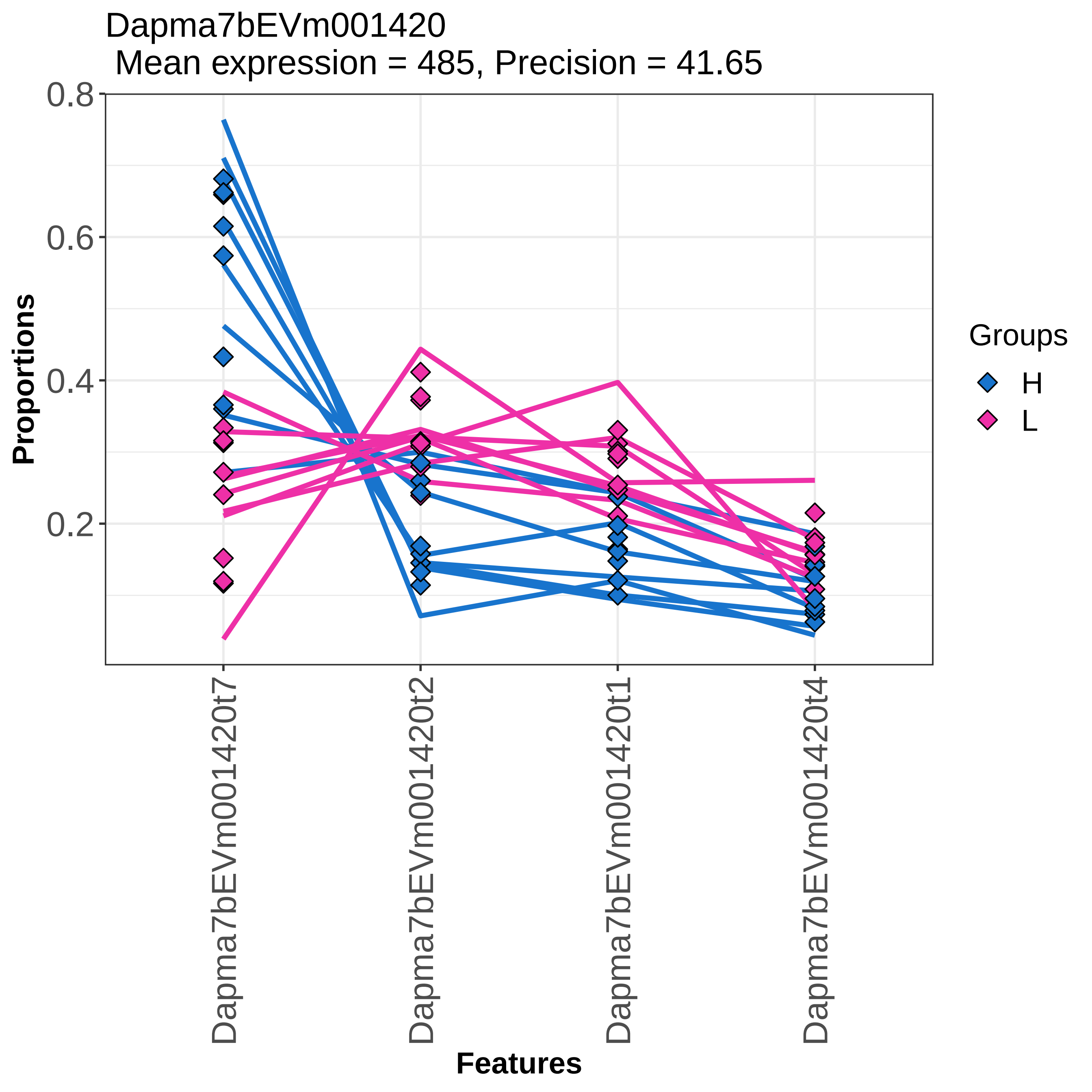

Dapma7bEVm001420
Mean expression = 485, Precision = 41.65
0.8
0.6
0.4
0.2
Proportions
Dapma7bEVm001420t7
Dapma7bEVm001420t2
Dapma7bEVm001420t1
Dapma7bEVm001420t4
Features
Groups
H
L

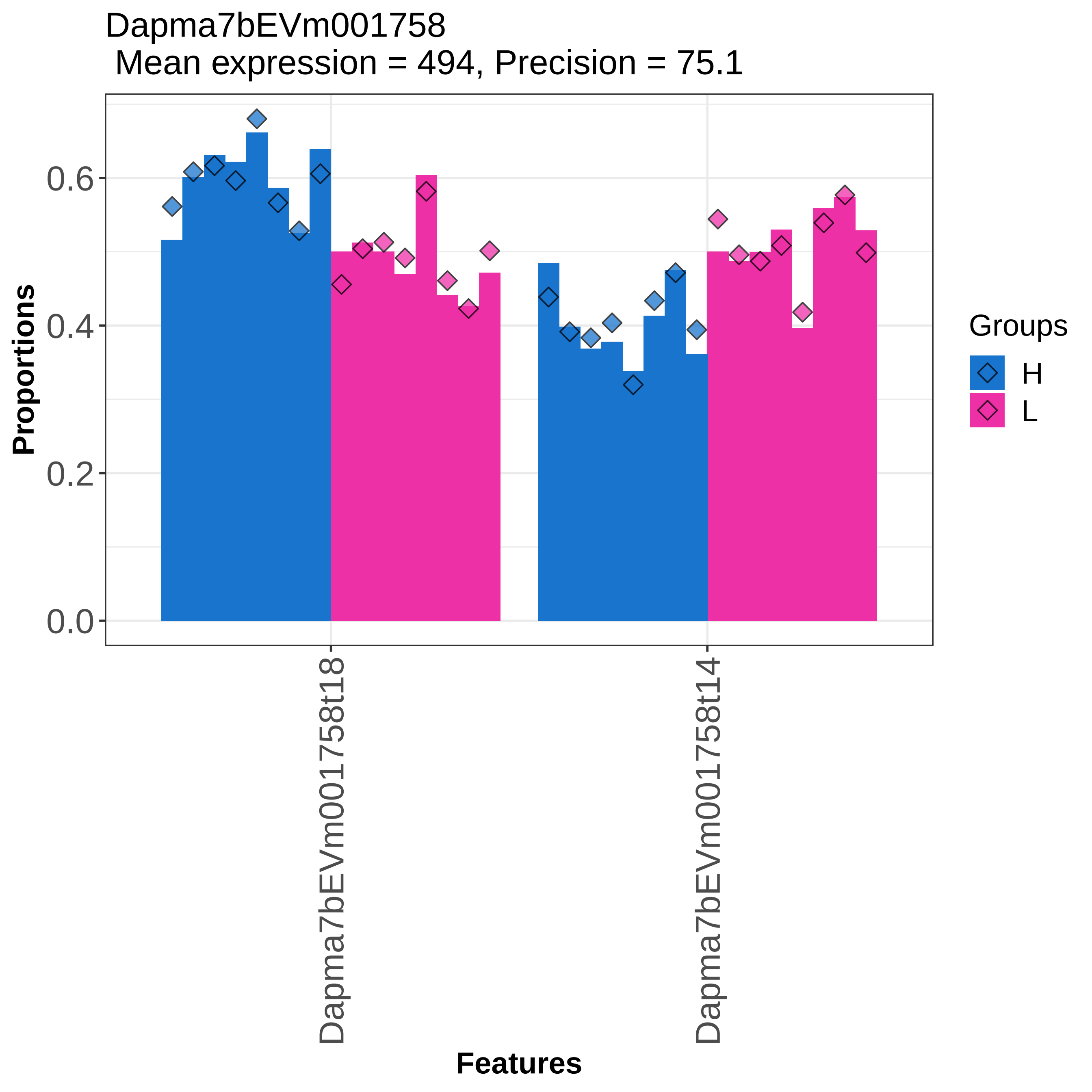

Dapma7bEVm001758
Mean expression = 494, Precision = 75.1
Proportions
0.0
0.2
0.4
0.6
Dapma7bEVm001758t18
Dapma7bEVm001758t14
Features
Groups
H
L

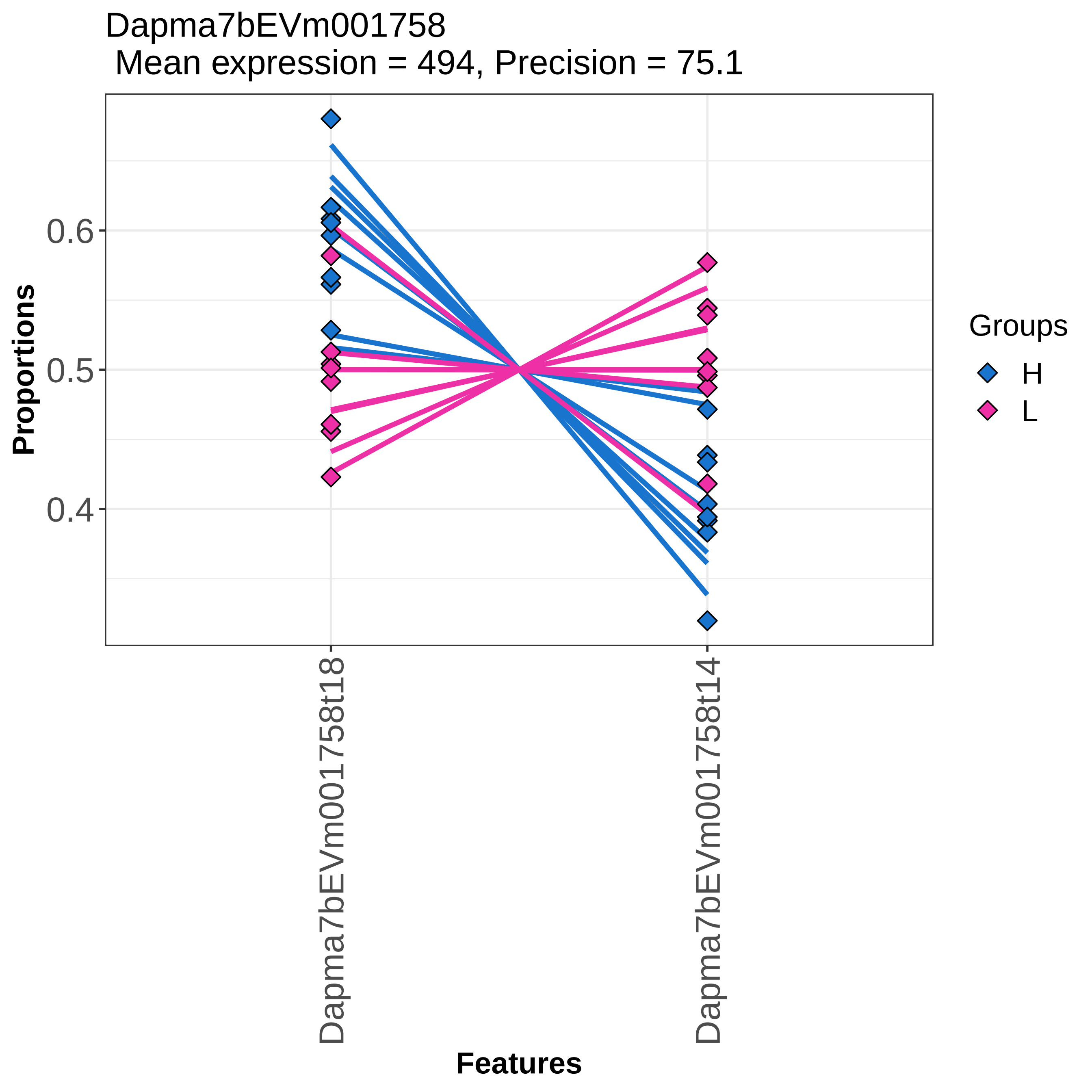

Dapma7bEVm001758
Mean expression = 494, Precision = 75.1
Proportions
0.6
0.5
0.4
Features
Dapma7bEVm001758t18
Dapma7bEVm001758t14
Groups
H
L

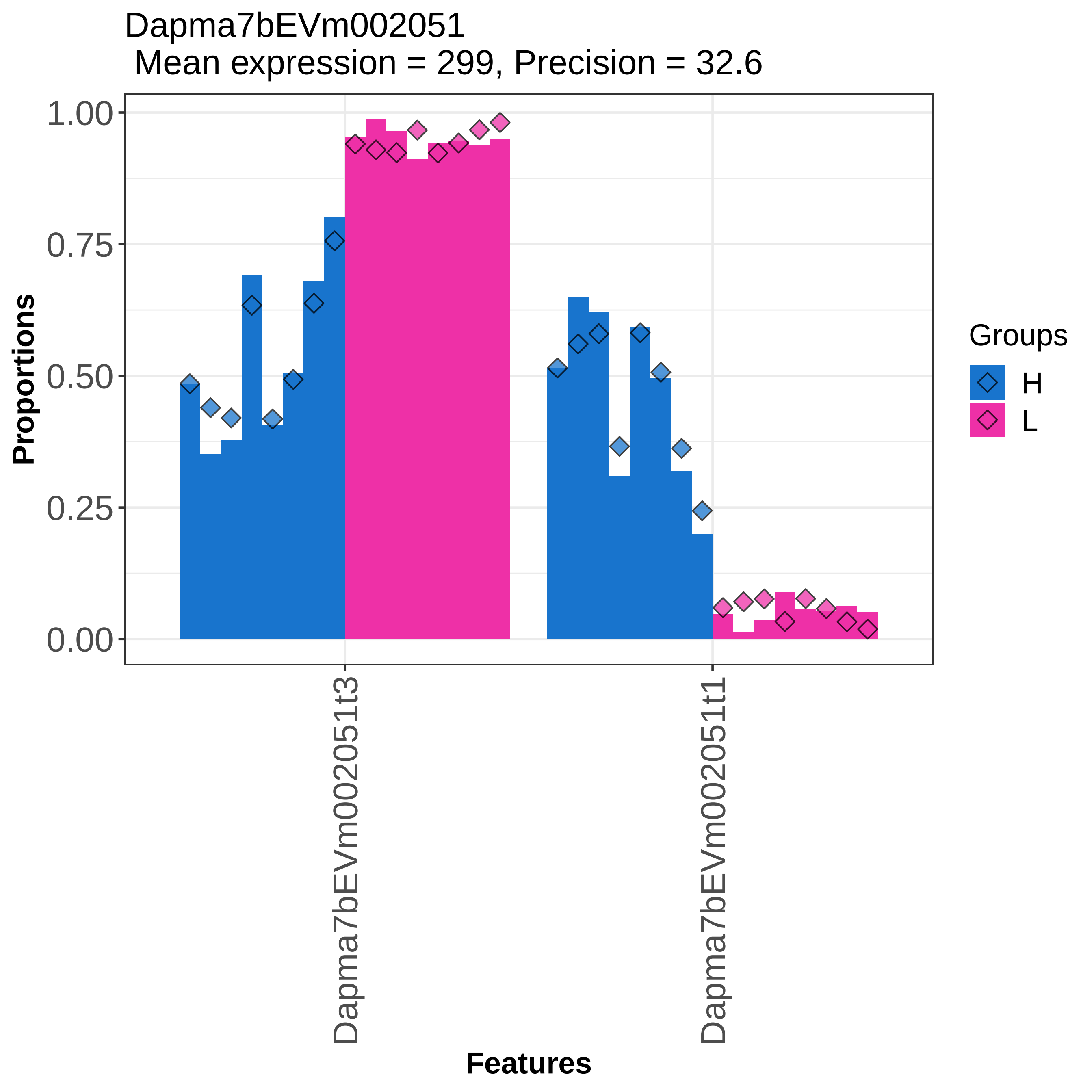

Dapma7bEVm002051
Mean expression = 299, Precision = 32.6
Proportions
1.00
0.75
0.50
0.25
0.00
Dapma7bEVm002051t3
Dapma7bEVm002051t1
Features
Groups
H
L

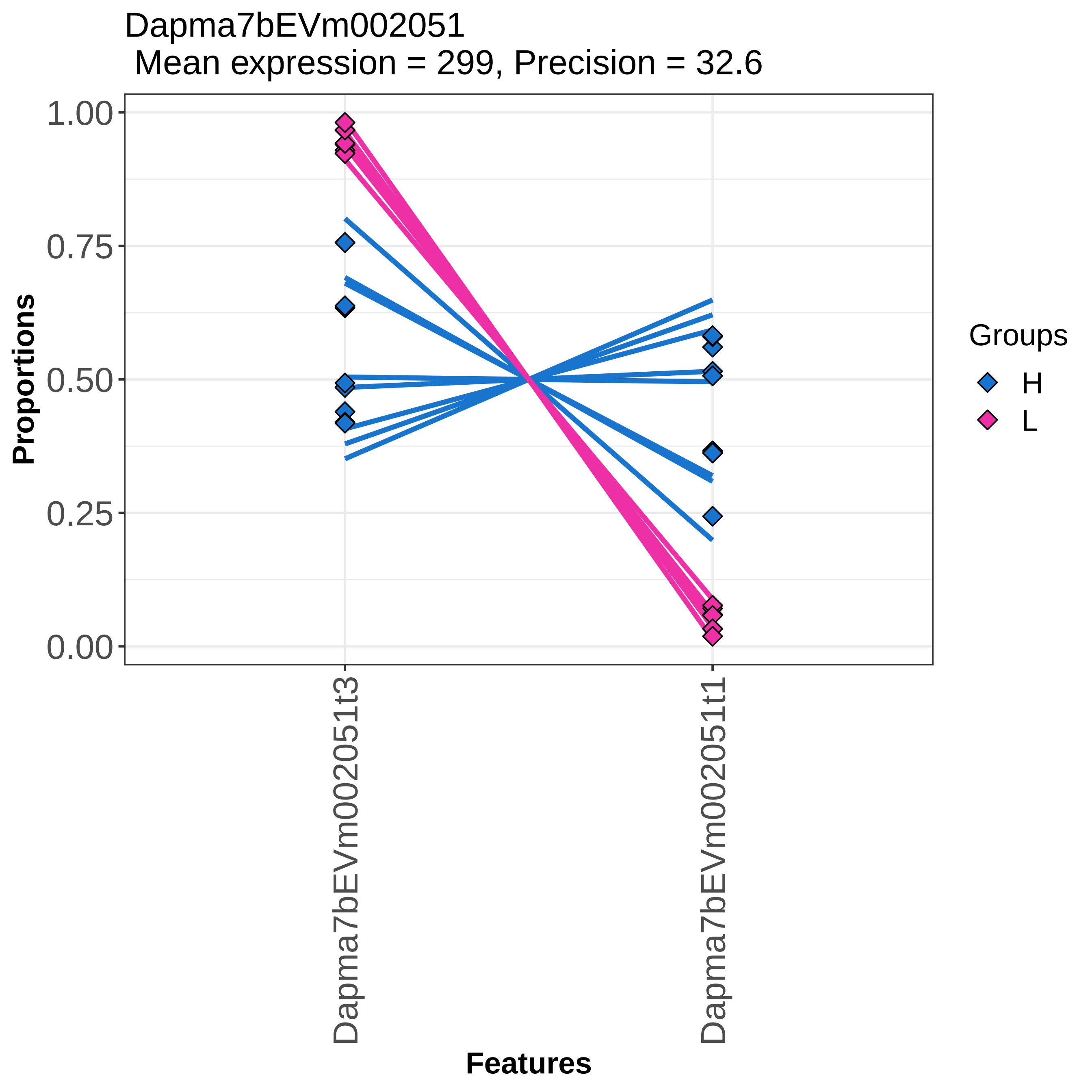

Dapma7bEVm002051
Mean expression = 299, Precision = 32.6
Proportions
1.00
0.75
0.50
0.25
0.00
Dapma7bEVm002051t3
Dapma7bEVm002051t1
Features
Groups
H
L

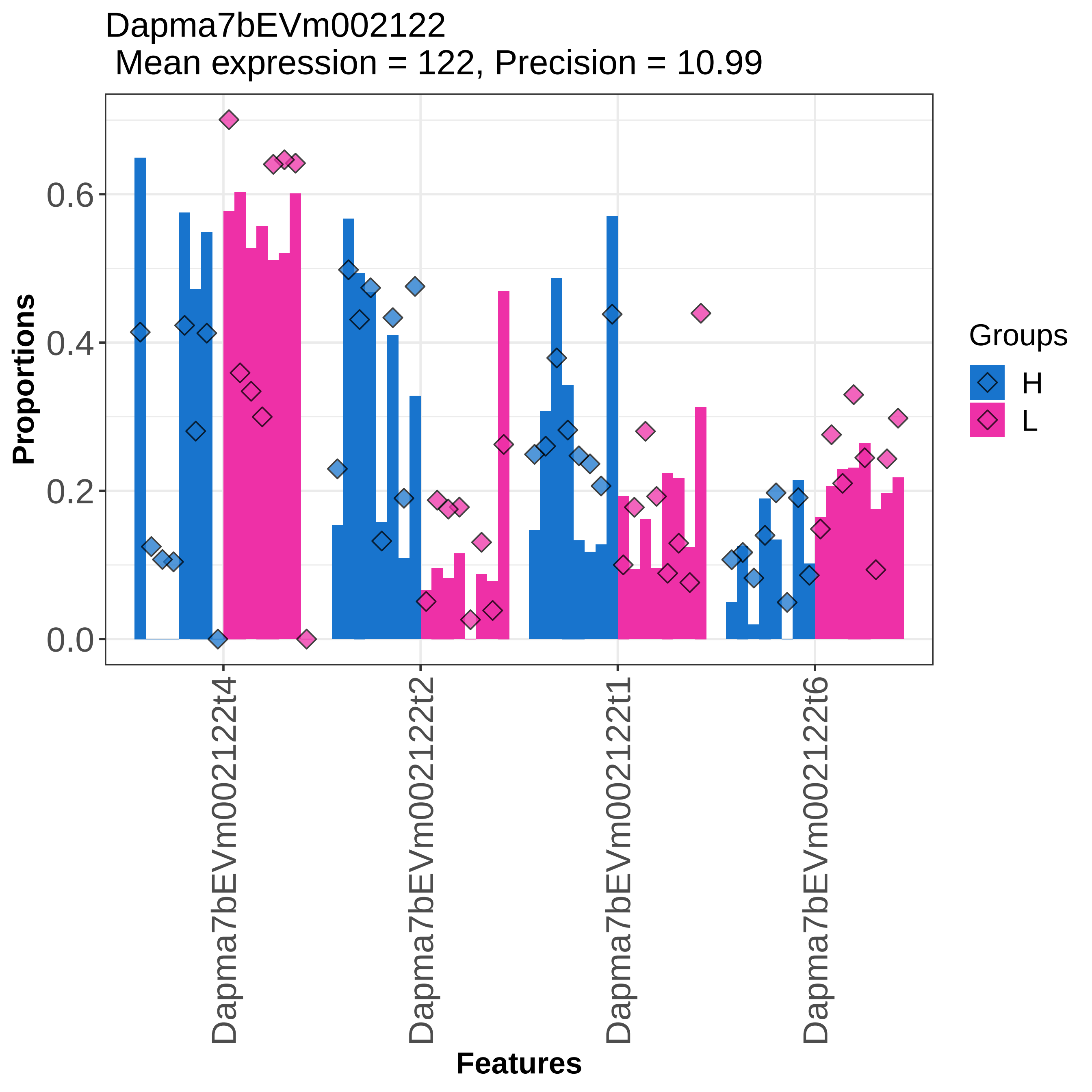
Dapma7bEVm002122
Mean expression = 122, Precision = 10.99
Proportions
0.0
0.2
0.4
0.6
Dapma7bEVm002122t4
Dapma7bEVm002122t2
Dapma7bEVm002122t1
Dapma7bEVm002122t6
Features
Groups
H
L

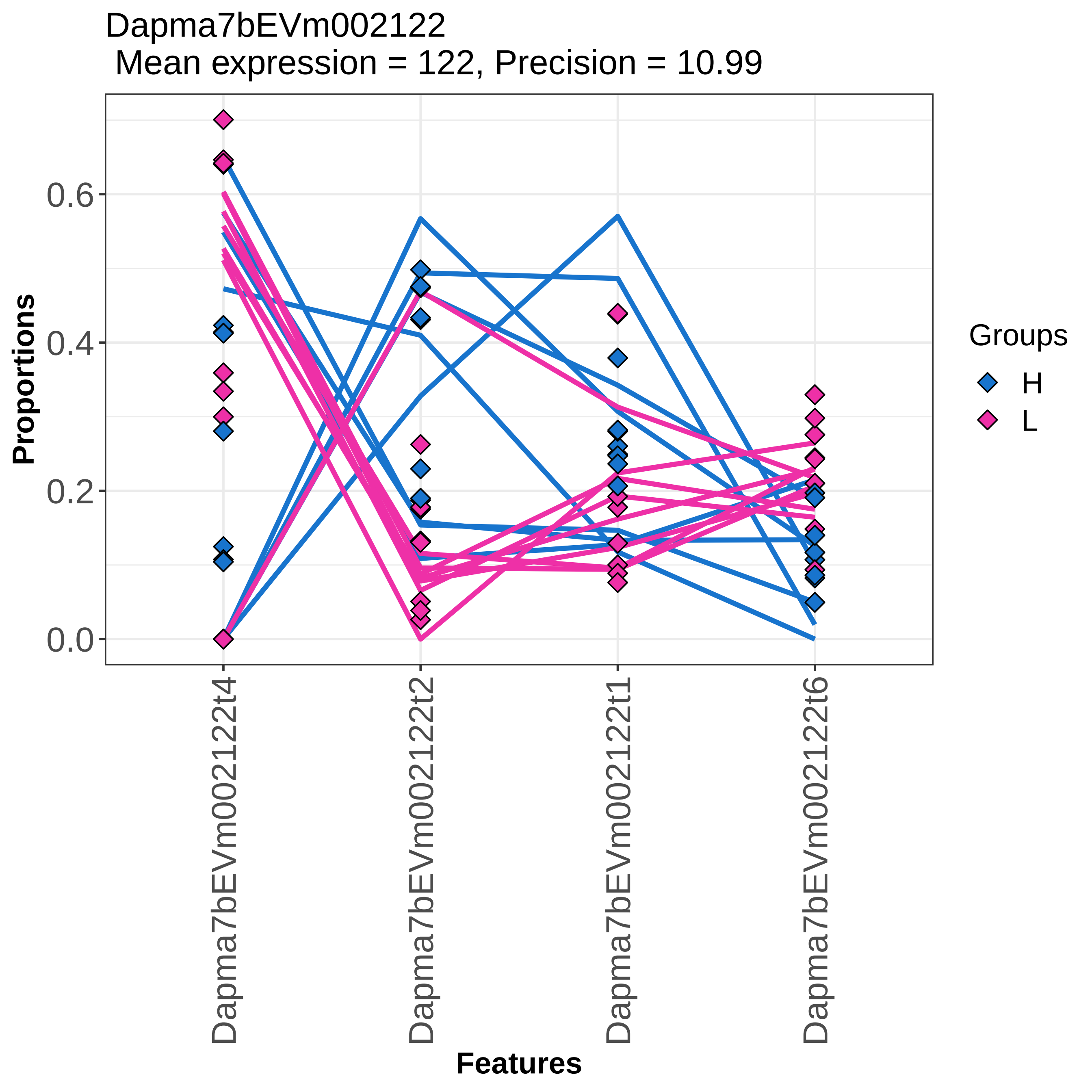
Dapma7bEVm002122
Mean expression = 122, Precision = 10.99
Proportions
0.0
0.2
0.4
0.6
Dapma7bEVm002122t4
Dapma7bEVm002122t2
Dapma7bEVm002122t1
Dapma7bEVm002122t6
Features
Groups
H
L

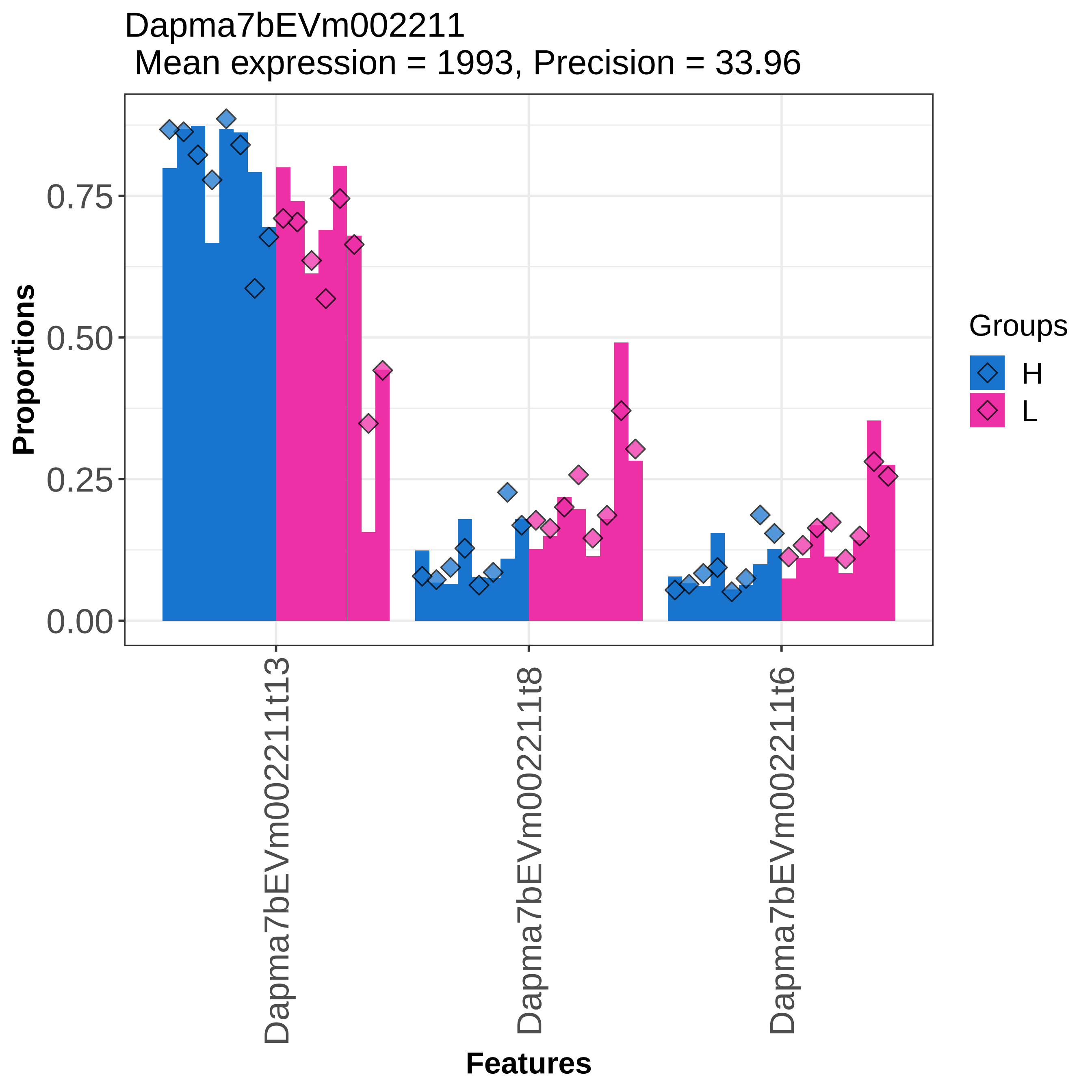

Dapma7bEVm002211
Mean expression = 1993, Precision = 33.96
Proportions
0.75
0.50
0.25
0.00
Dapma7bEVm002211t13
Dapma7bEVm002211t8
Dapma7bEVm002211t6
Features
Groups
H
L

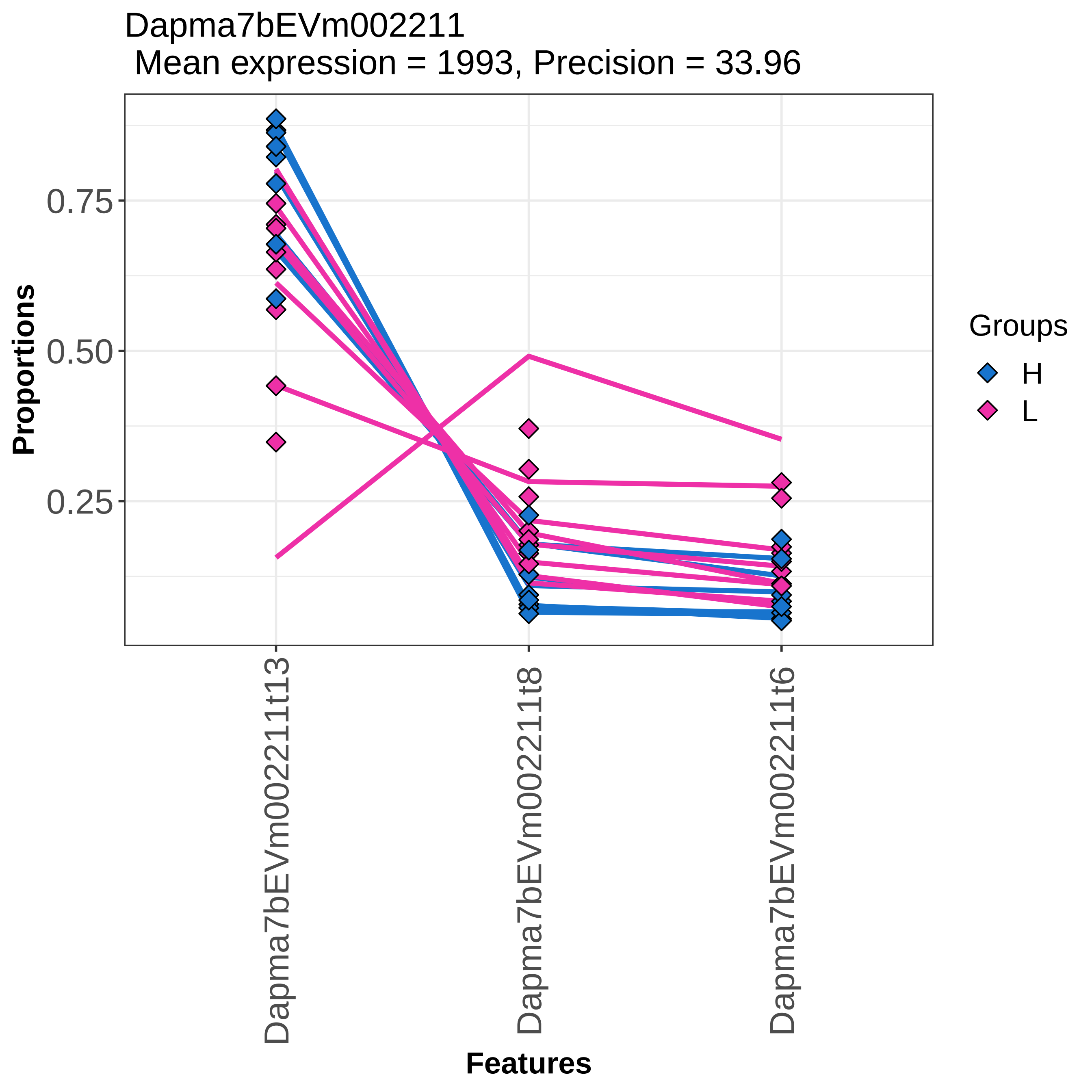
Dapma7bEVm002211
Mean expression = 1993, Precision = 33.96
Proportions
0.75
0.50
0.25
Dapma7bEVm002211t13
Dapma7bEVm002211t8
Dapma7bEVm002211t6
Features
Groups
H
L

Dapma7bEVm002306
Mean expression = 284, Precision = 7.56

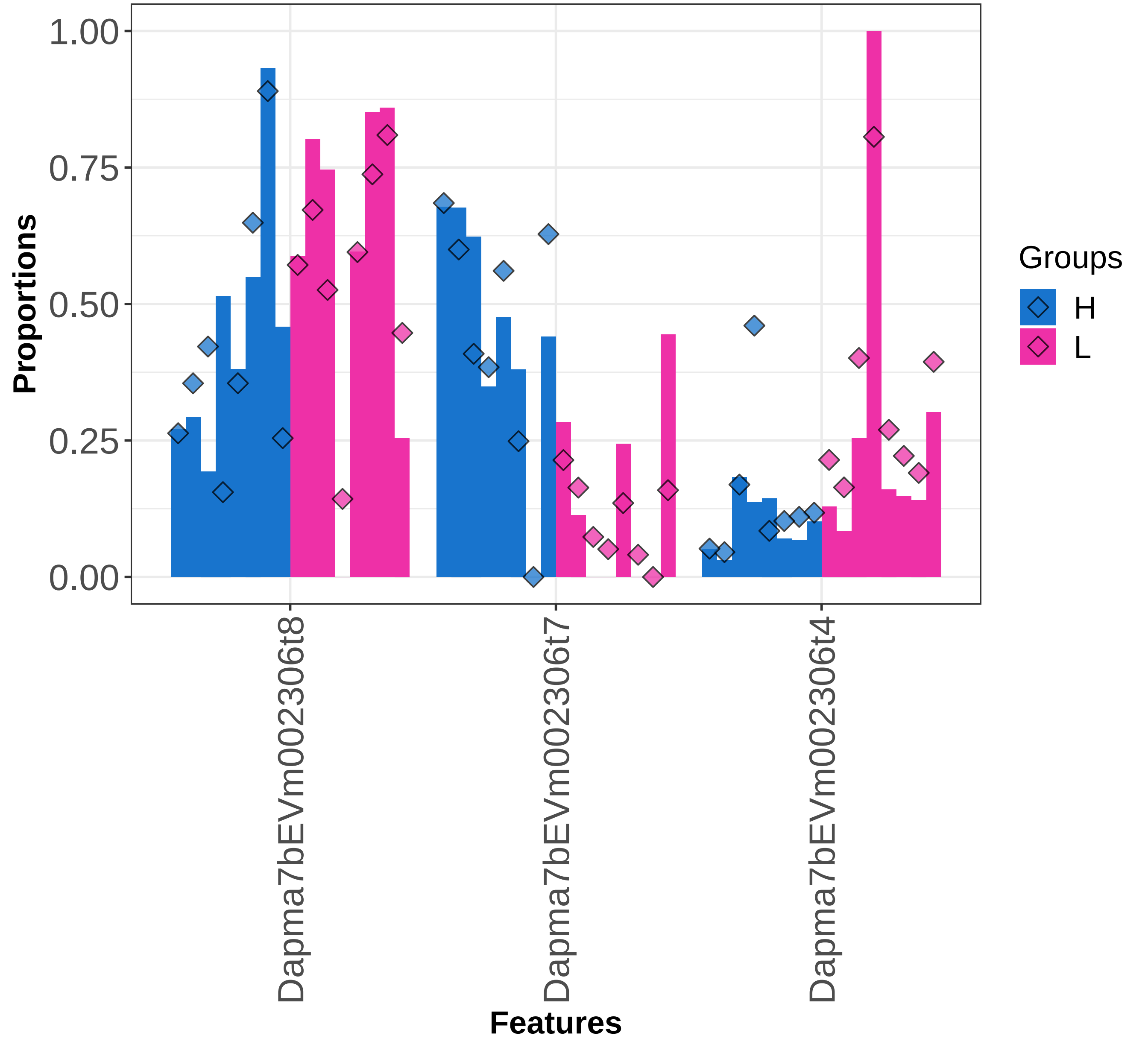

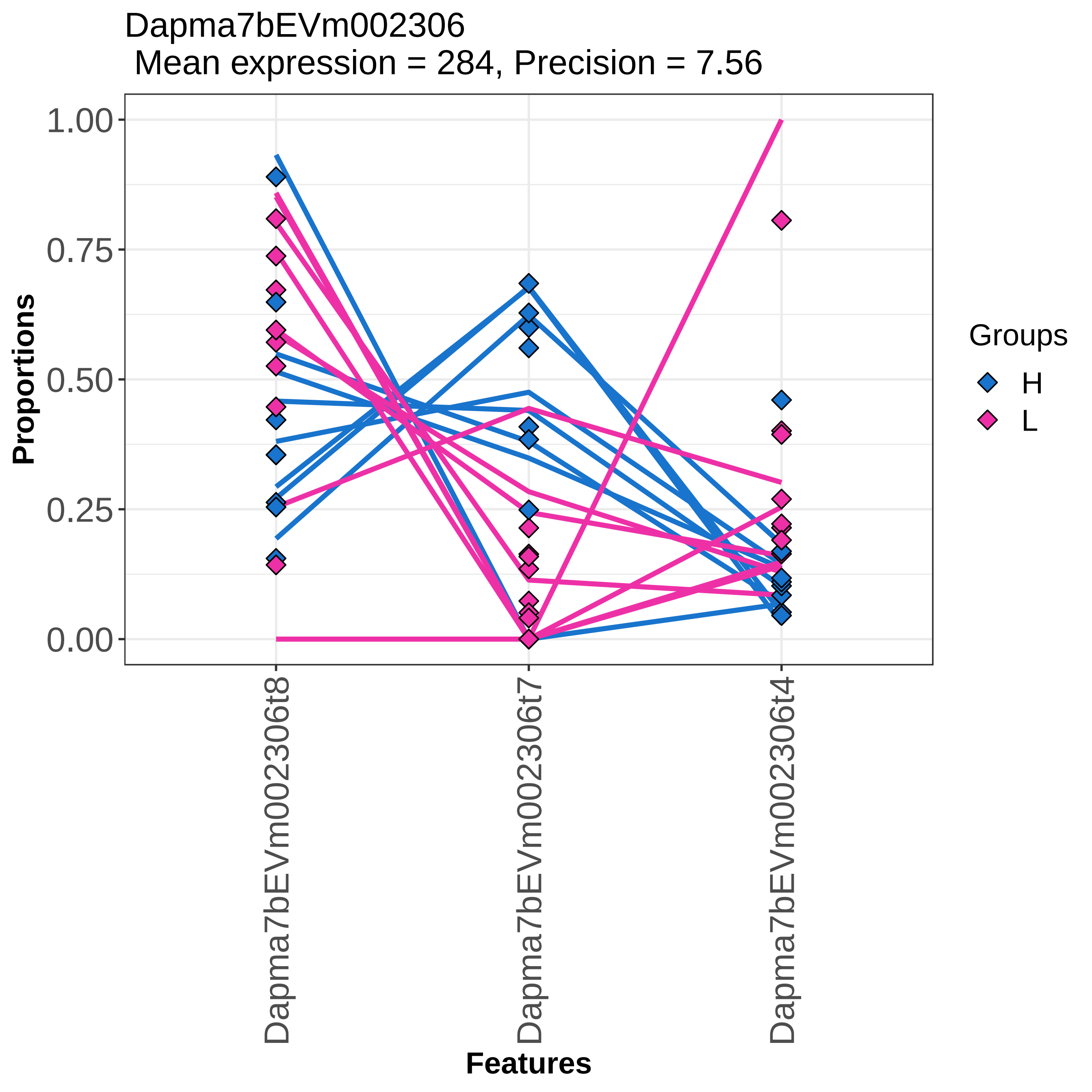

Dapma7bEVm002306
Mean expression = 284, Precision = 7.56
Proportions
1.00
0.75
0.50
0.25
0.00
Dapma7bEVm002306t8
Dapma7bEVm002306t7
Dapma7bEVm002306t4
Features
Groups
H
L

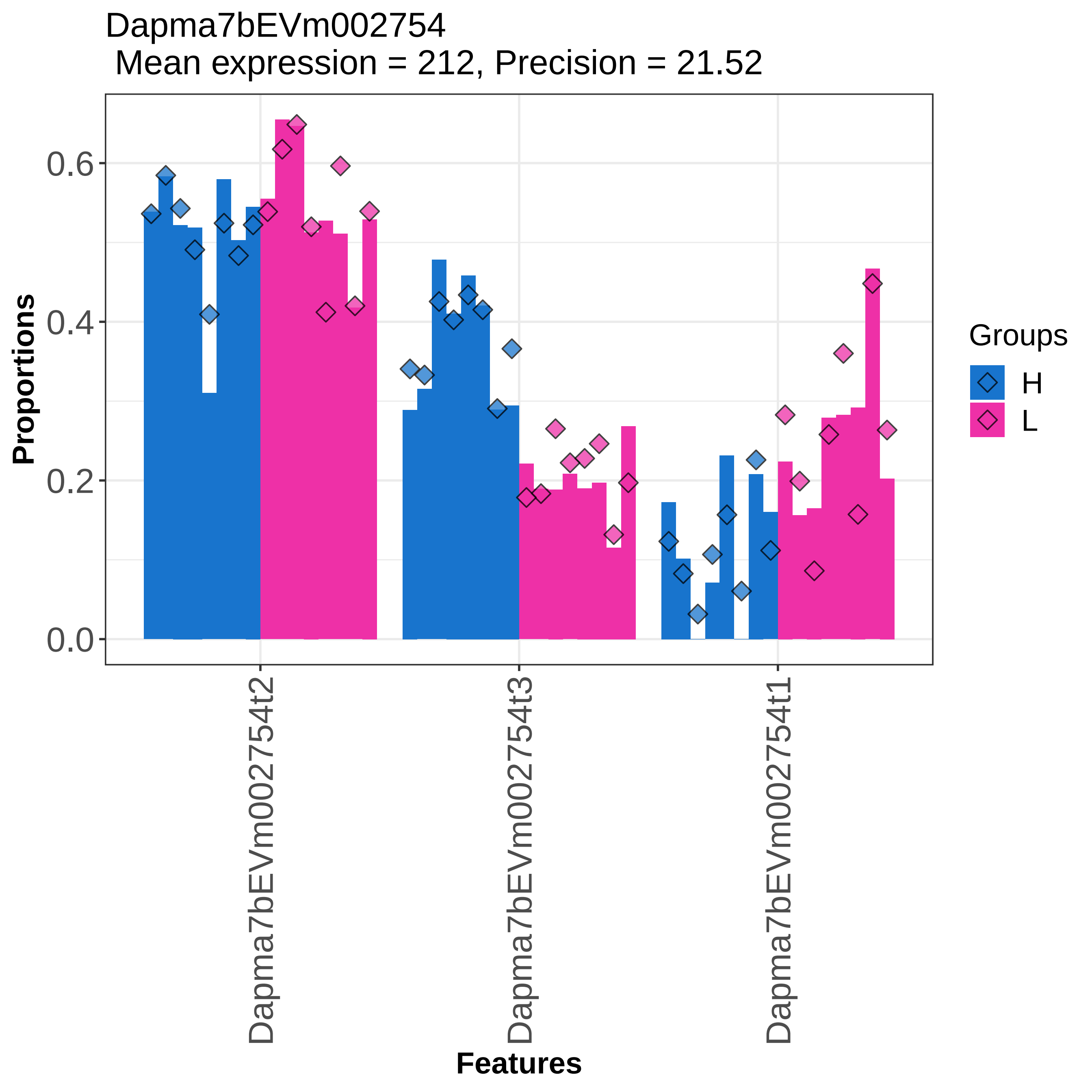

Dapma7bEVm002754
Mean expression = 212, Precision = 21.52
Proportions
0.0
0.2
0.4
0.6
Features
Dapma7bEVm002754t2
Dapma7bEVm002754t3
Dapma7bEVm002754t1
Groups
H
L

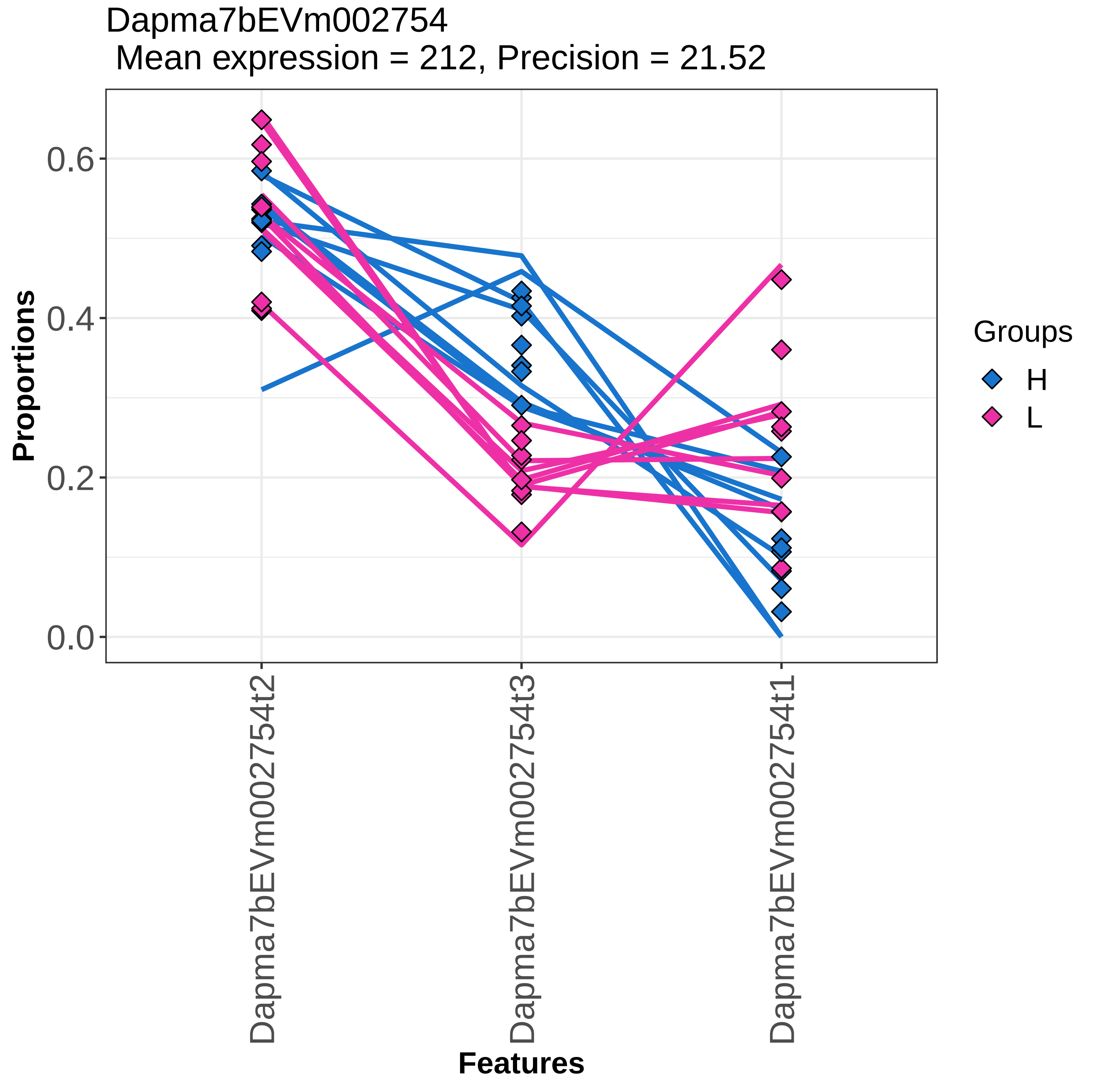
Dapma7bEVm002754
Mean expression = 212, Precision = 21.52
Proportions
0.0
0.2
0.4
0.6
Dapma7bEVm002754t2
Dapma7bEVm002754t3
Dapma7bEVm002754t1
Features
Groups
H
L

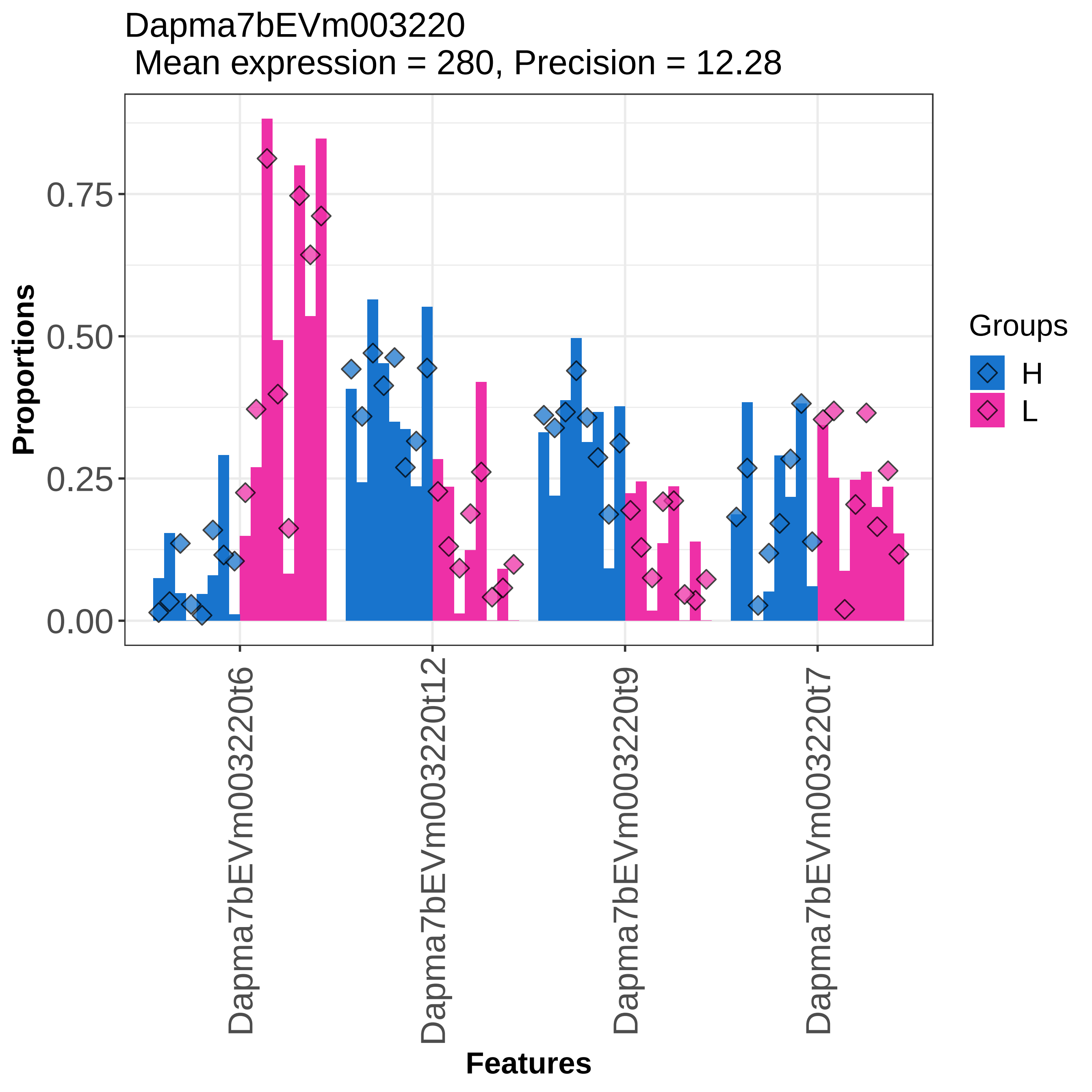

Dapma7bEVm003220
Mean expression = 280, Precision = 12.28
Proportions
0.75
0.50
0.25
0.00
Dapma7bEVm003220t6
Dapma7bEVm003220t12
Dapma7bEVm003220t9
Dapma7bEVm003220t7
Features
Groups
H
L

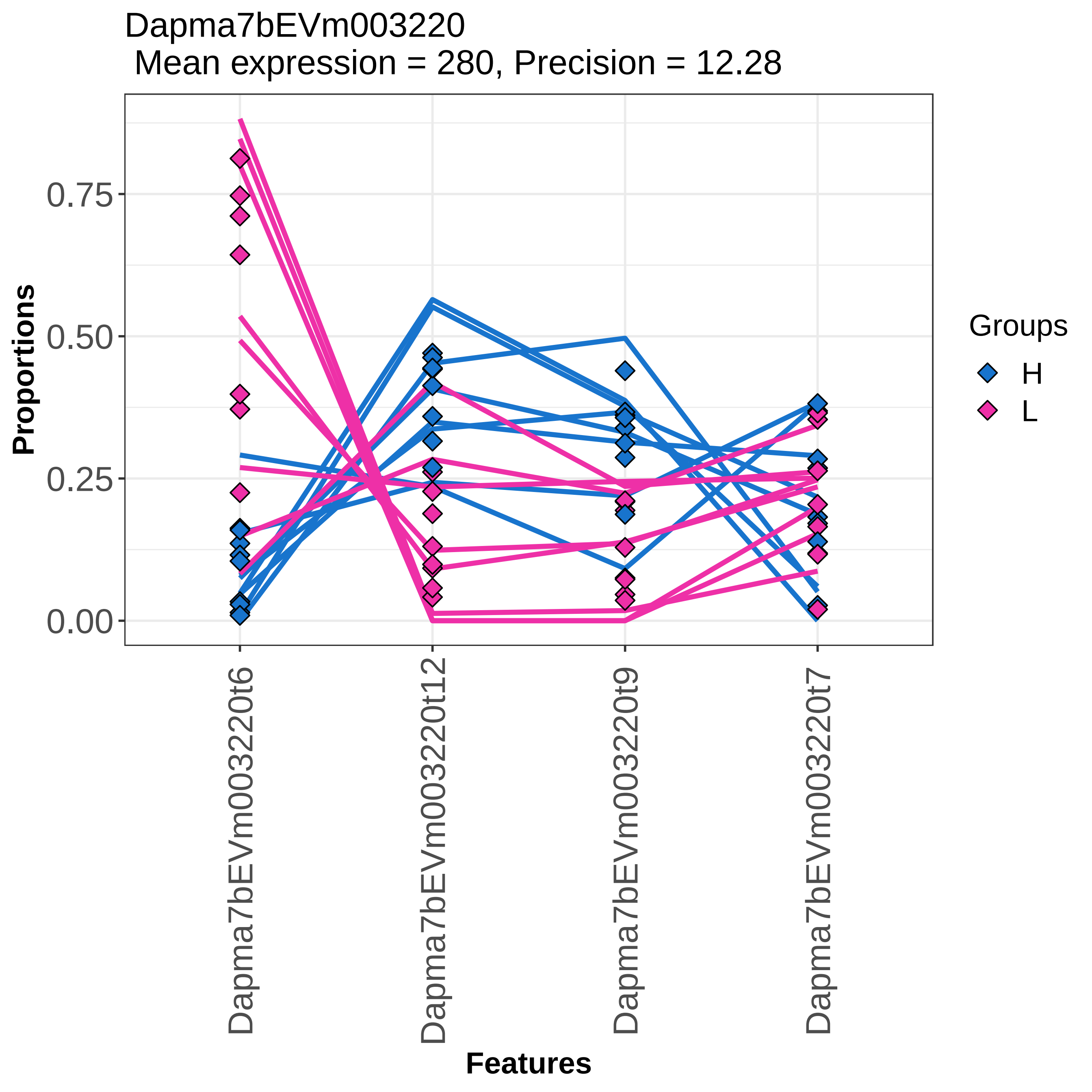

Dapma7bEVm003220
Mean expression = 280, Precision = 12.28
Proportions
0.75
0.50
0.25
0.00
Dapma7bEVm003220t6
Dapma7bEVm003220t12
Dapma7bEVm003220t9
Dapma7bEVm003220t7
Features
Groups
H
L

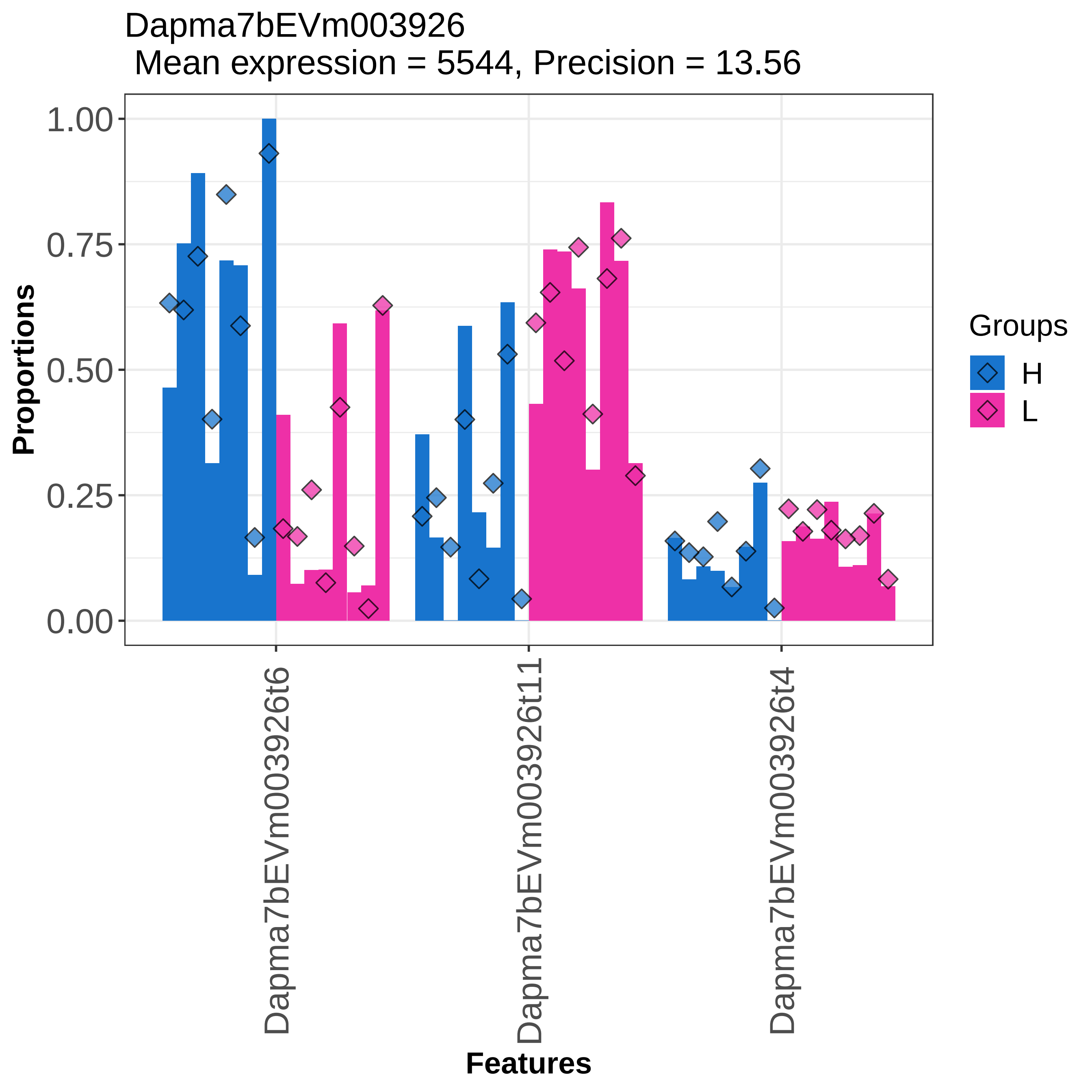
Dapma7bEVm003926
Mean expression = 5544, Precision = 13.56
Proportions
1.00
0.75
0.50
0.25
0.00
Dapma7bEVm003926t6
Dapma7bEVm003926t11
Dapma7bEVm003926t4
Features
Groups
H
L

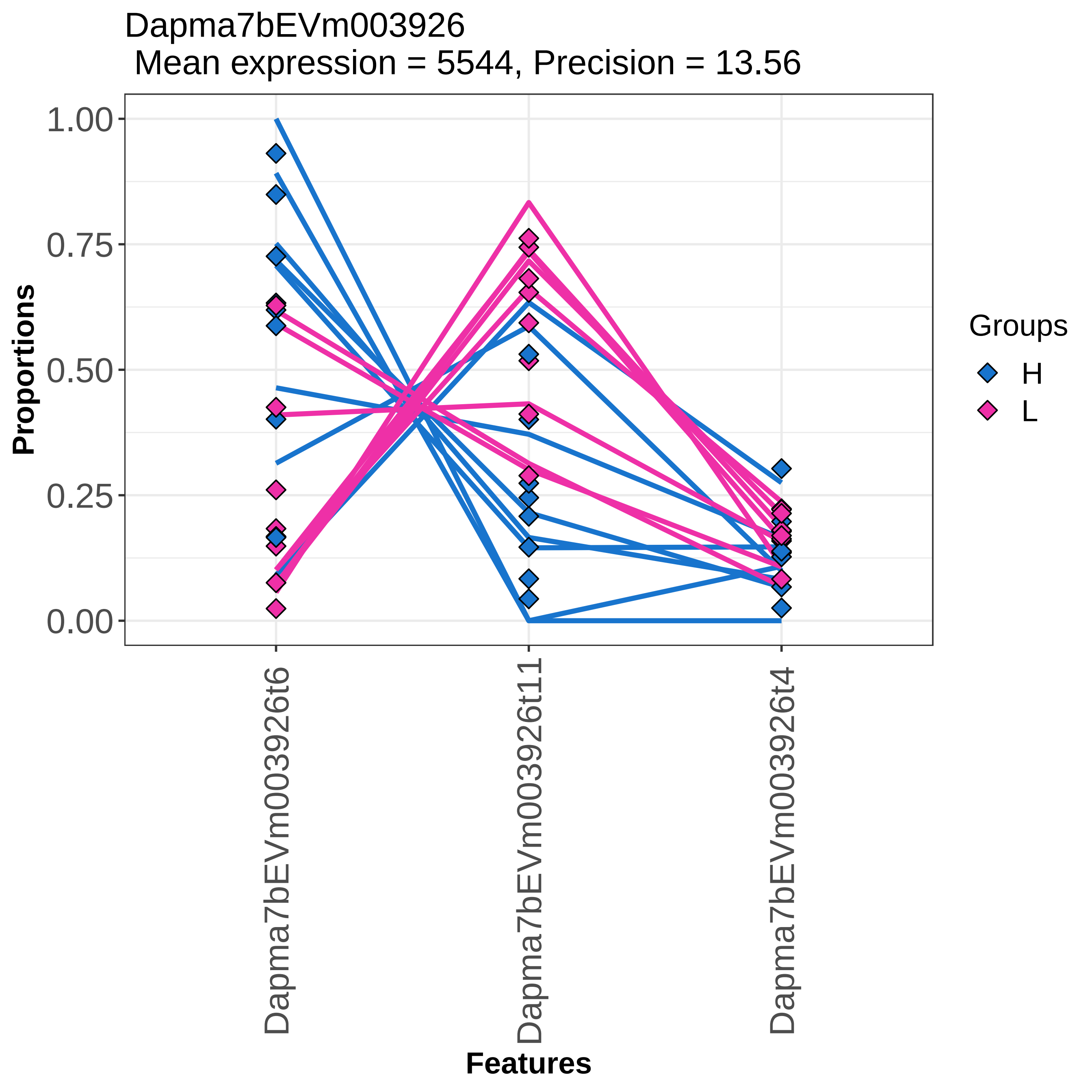

Dapma7bEVm003926
Mean expression = 5544, Precision = 13.56
Proportions
1.00
0.75
0.50
0.25
0.00
Dapma7bEVm003926t6
Dapma7bEVm003926t11
Dapma7bEVm003926t4
Features
Groups
H
L

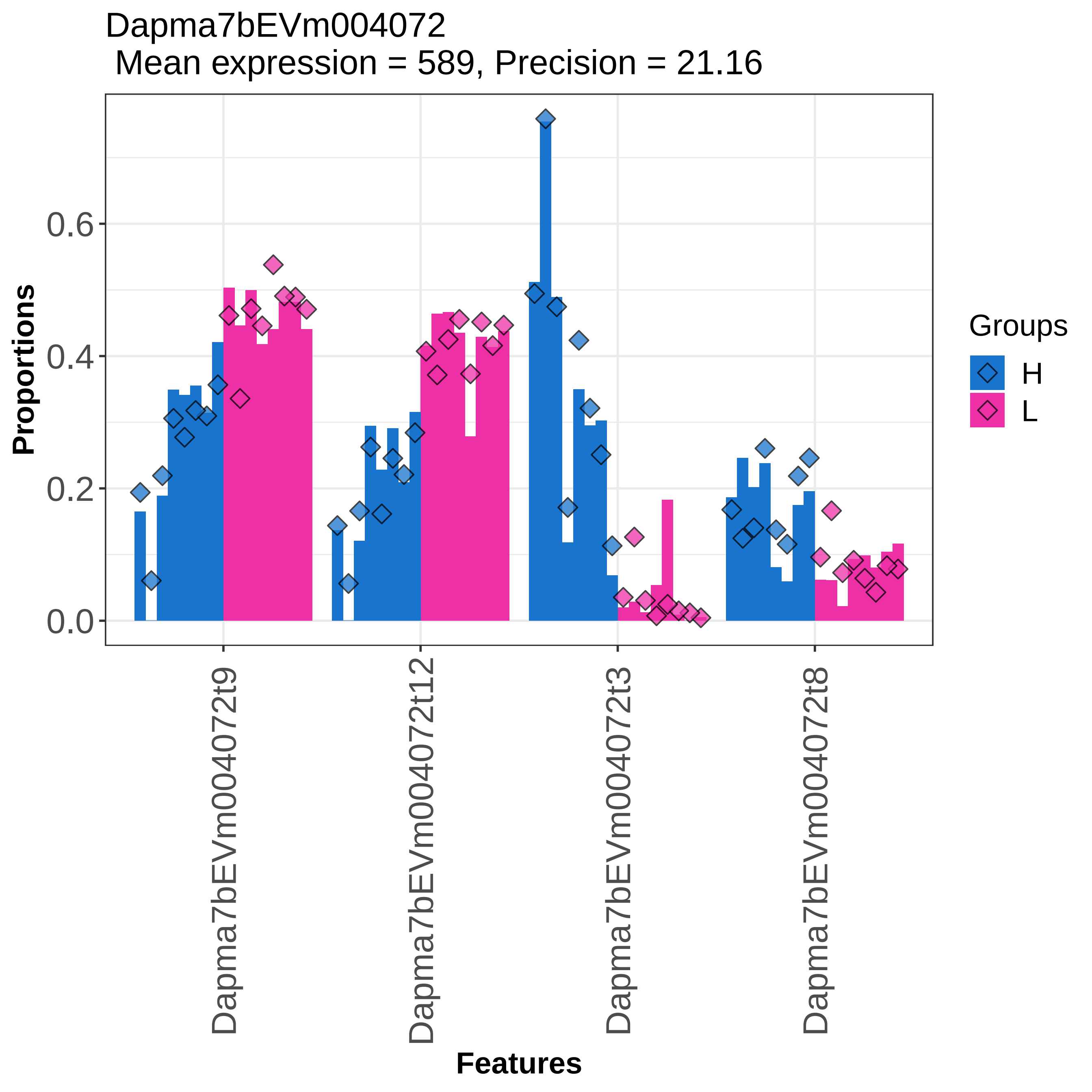
Dapma7bEVm004072
Mean expression = 589, Precision = 21.16
Proportions
0.0
0.2
0.4
0.6
Dapma7bEVm004072t9
Dapma7bEVm004072t12
Dapma7bEVm004072t3
Dapma7bEVm004072t8
Features
Groups
H
L

Dapma7bEVm004072
Mean expression = 589, Precision = 21.16

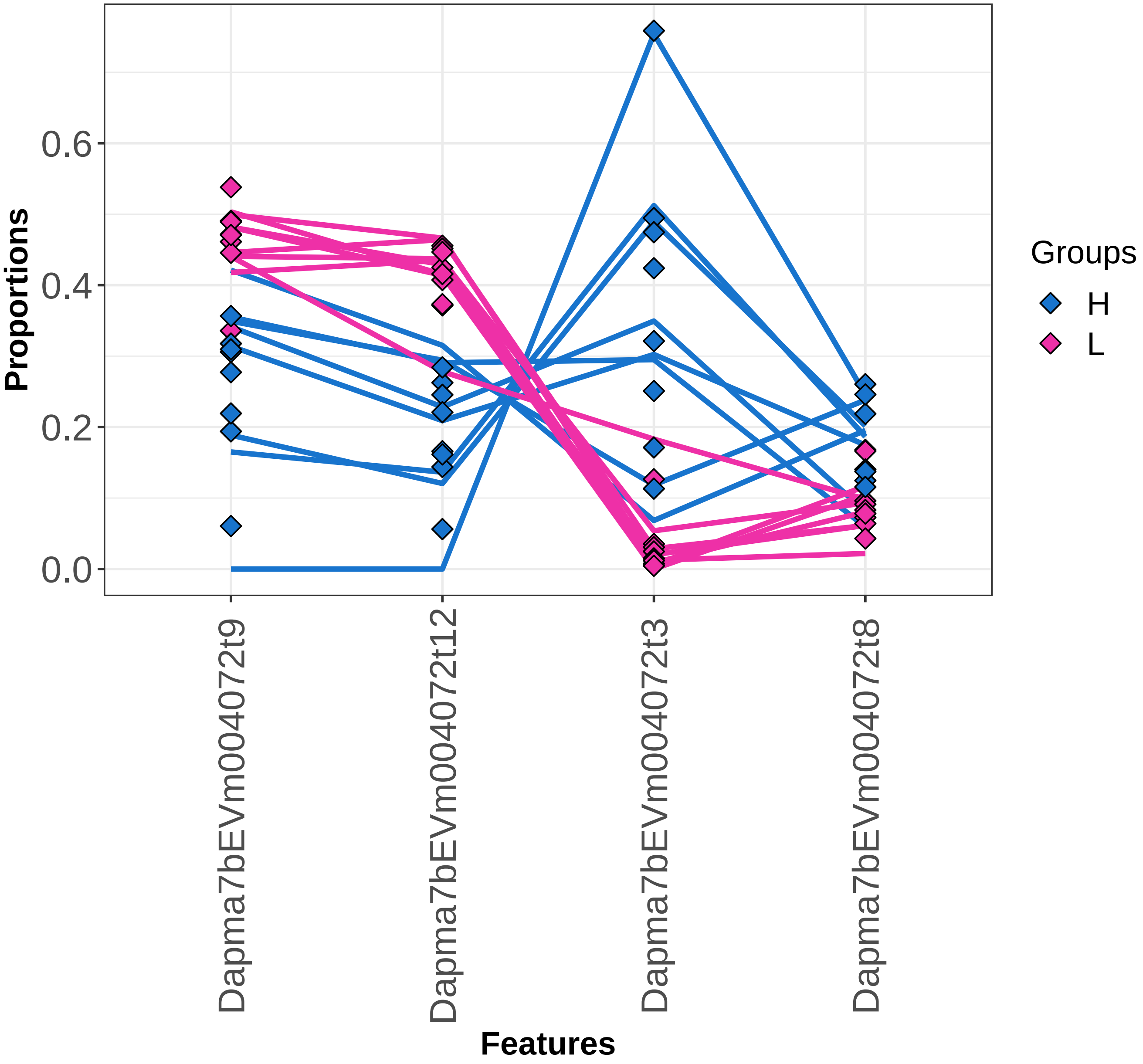

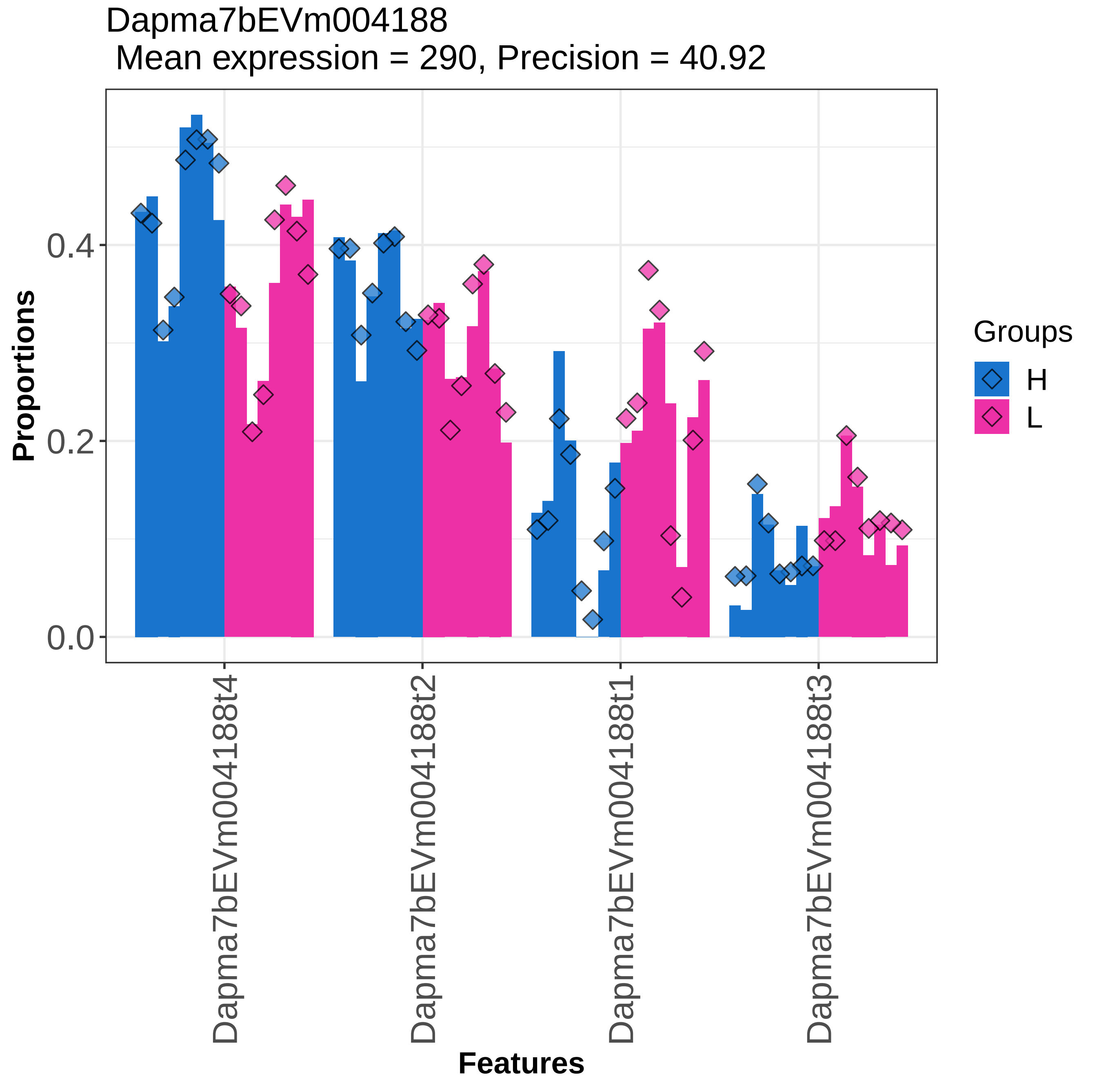
Dapma7bEVm004188
Mean expression = 290, Precision = 40.92
Proportions
0.0
0.2
0.4
Dapma7bEVm004188t4
Dapma7bEVm004188t2
Dapma7bEVm004188t1
Dapma7bEVm004188t3
Features
Groups
H
L

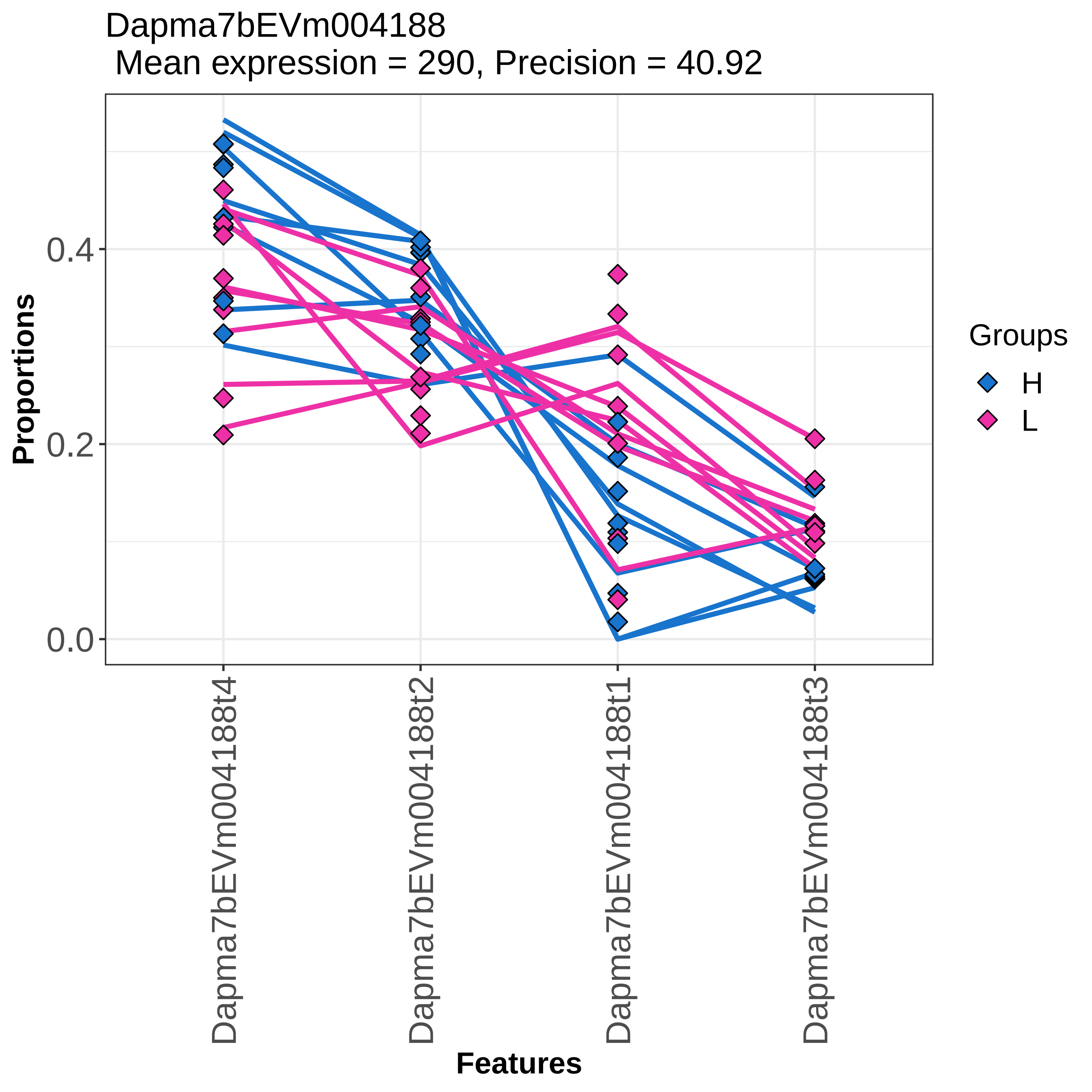

Dapma7bEVm004188
Mean expression = 290, Precision = 40.92
Proportions
0.4
0.2
0.0
Dapma7bEVm004188t4
Dapma7bEVm004188t2
Dapma7bEVm004188t1
Dapma7bEVm004188t3
Features
Groups
H
L

Dapma7bEVm004206
Mean expression = 340, Precision = 44.81

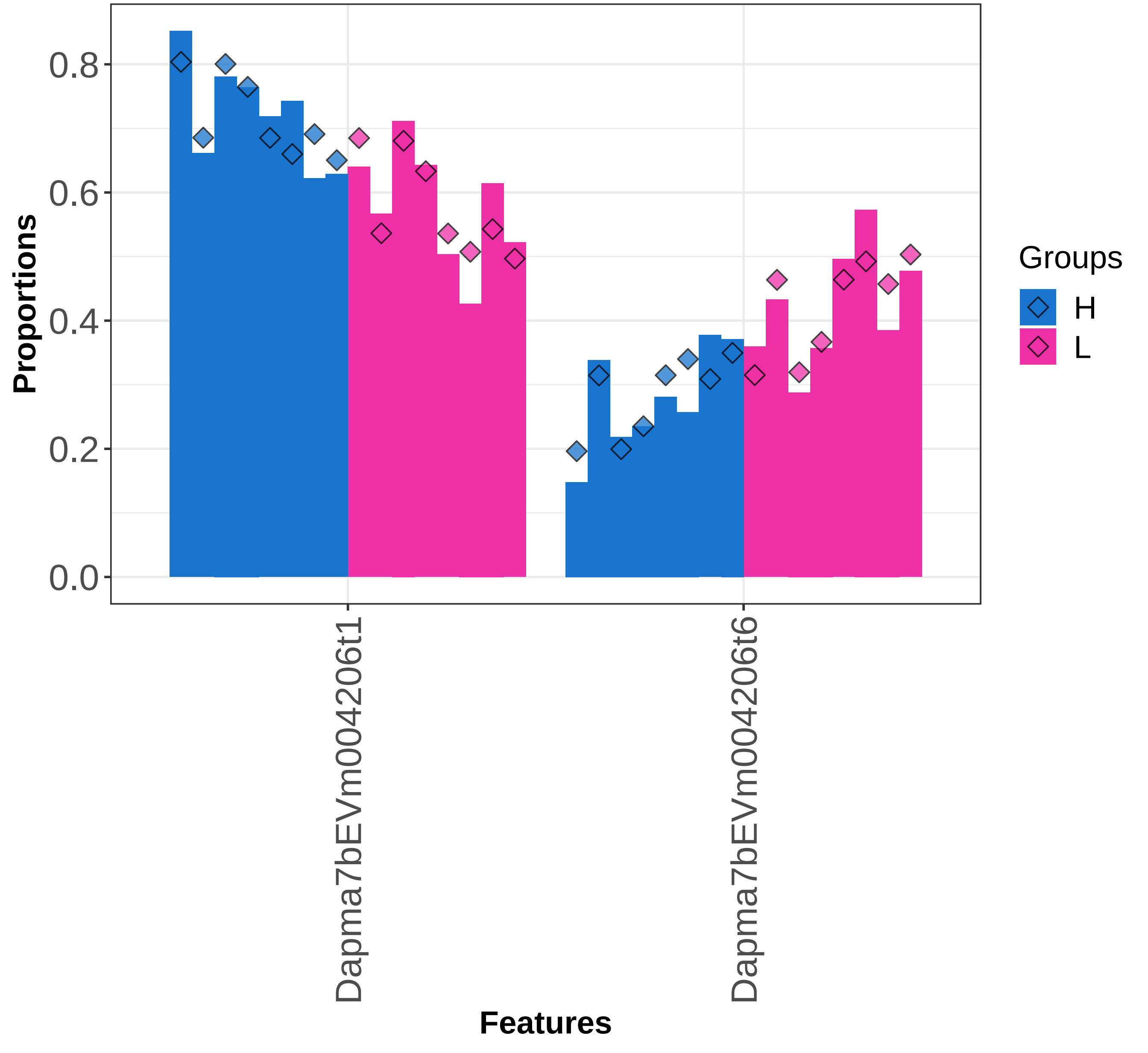

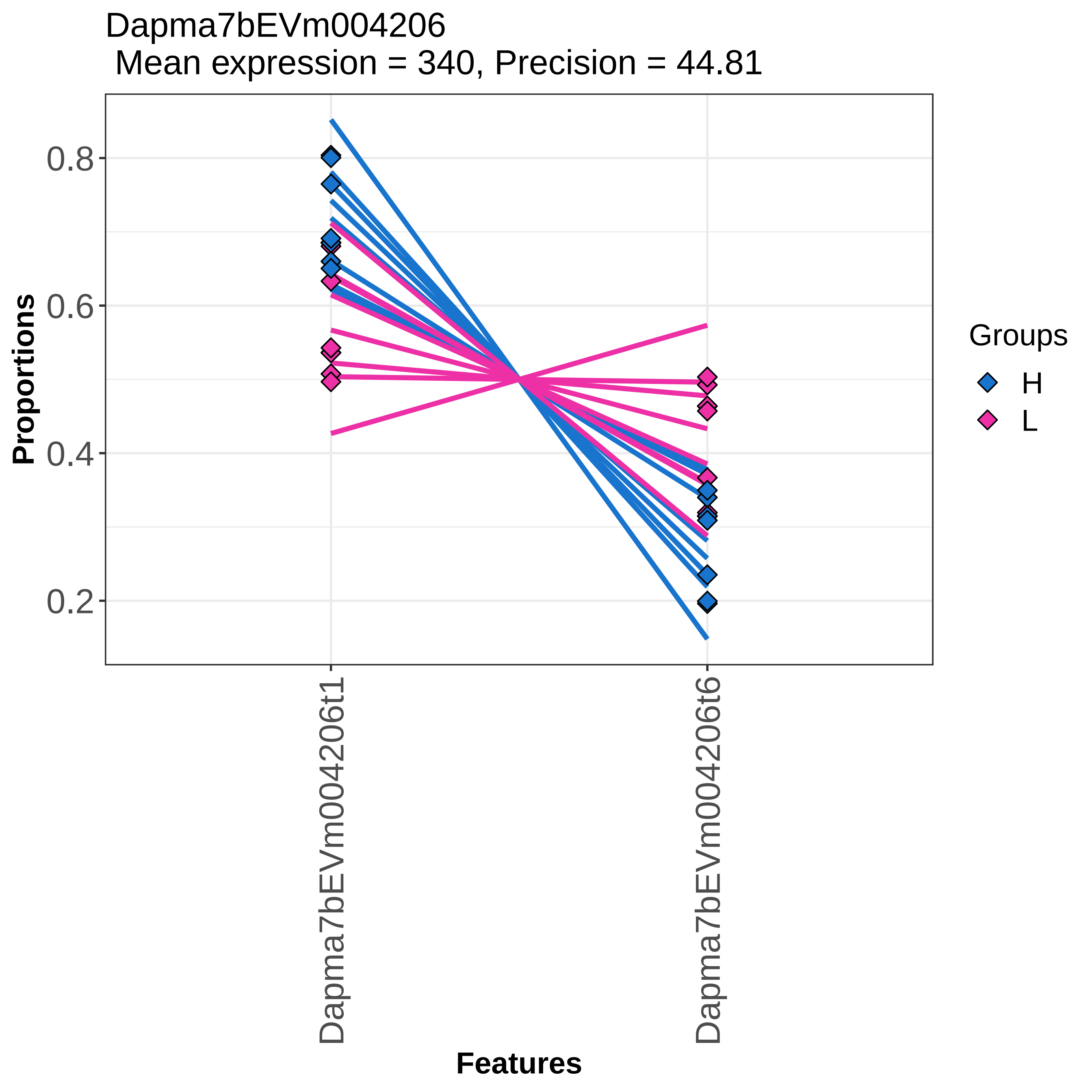

Dapma7bEVm004206
Mean expression = 340, Precision = 44.81
Proportions
0.8
0.6
0.4
0.2
Dapma7bEVm004206t1
Dapma7bEVm004206t6
Features
Groups
H
L

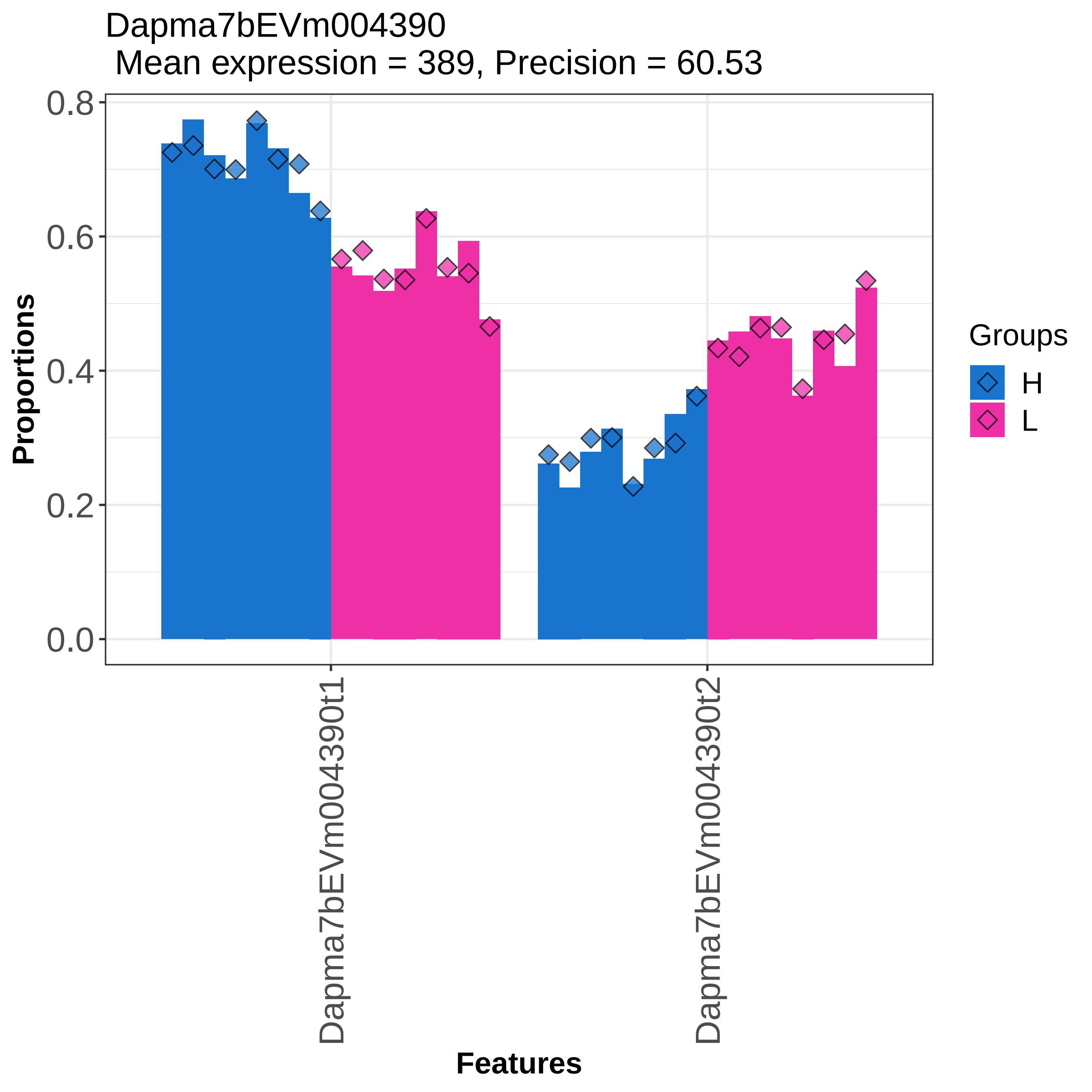
Dapma7bEVm004390
Mean expression = 389, Precision = 60.53
Proportions
0.8
0.6
0.4
0.2
0.0
Dapma7bEVm004390t1
Dapma7bEVm004390t2
Features
Groups
H
L

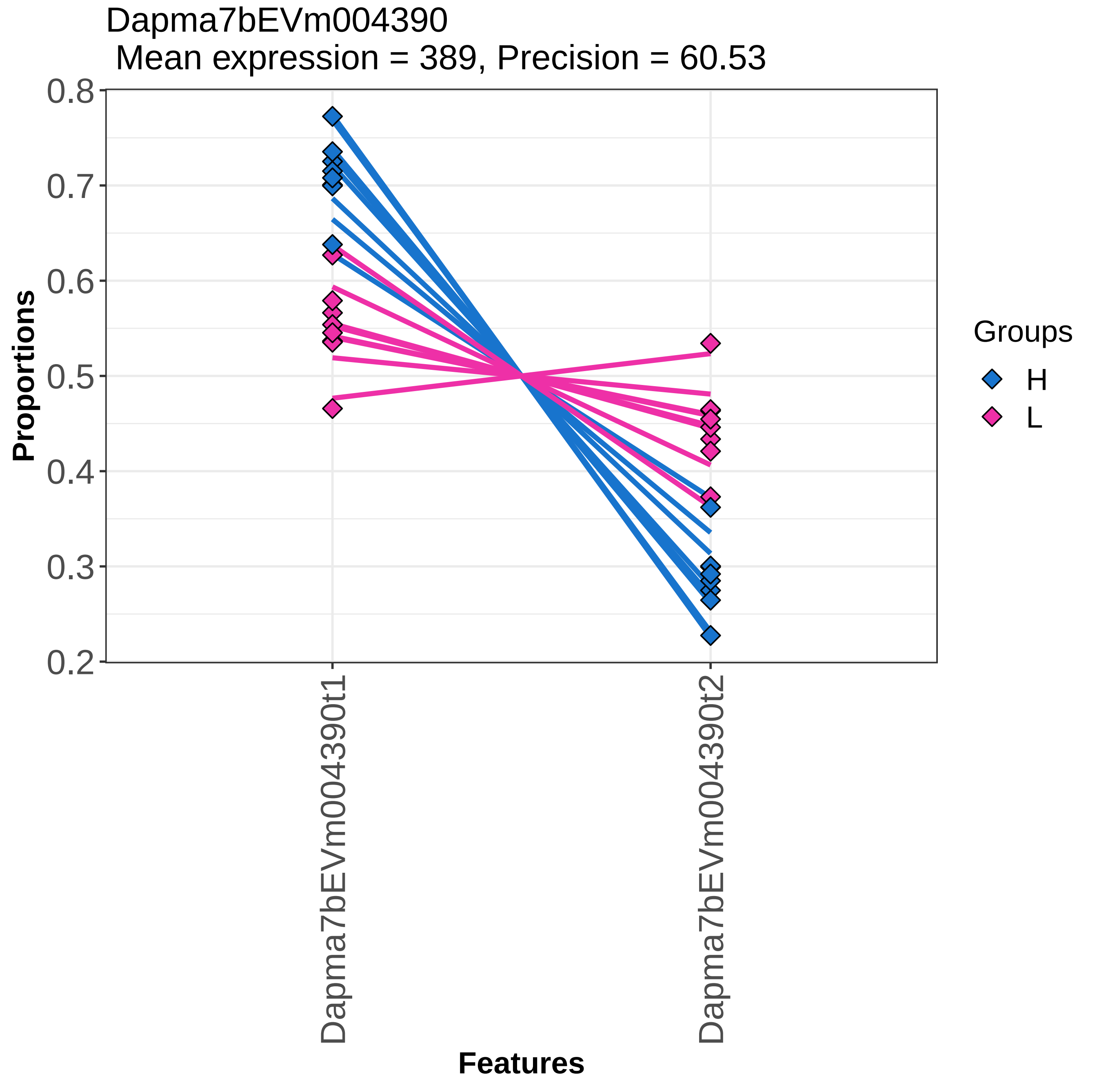

Dapma7bEVm004390
Mean expression = 389, Precision = 60.53
Proportions
0.8
0.7
0.6
0.5
0.4
0.3
0.2
Dapma7bEVm004390t1
Dapma7bEVm004390t2
Features
Groups
H
L

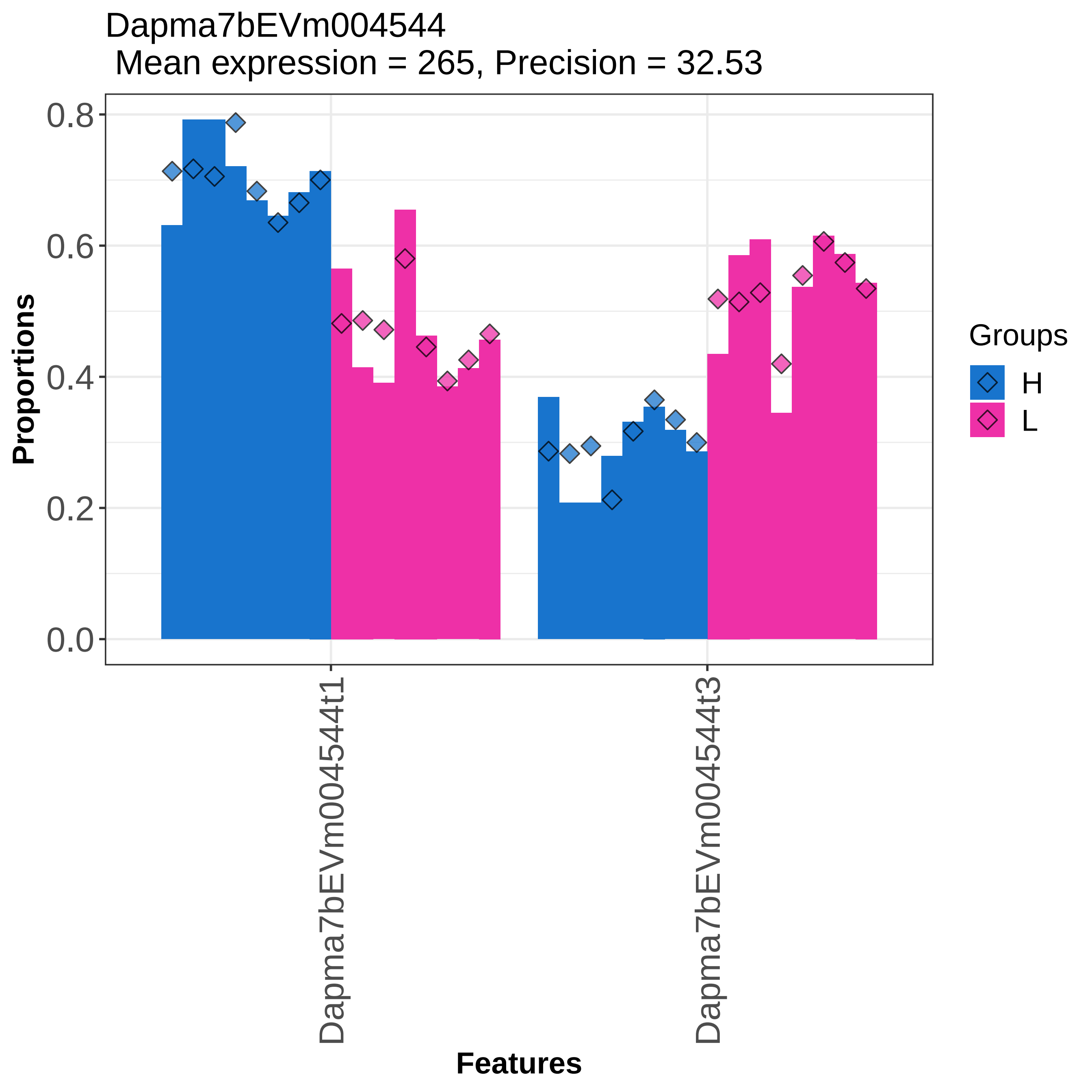

Dapma7bEVm004544
Mean expression = 265, Precision = 32.53
Proportions
0.0
0.2
0.4
0.6
0.8
Dapma7bEVm004544t1
Dapma7bEVm004544t3
Features
Groups
H
L

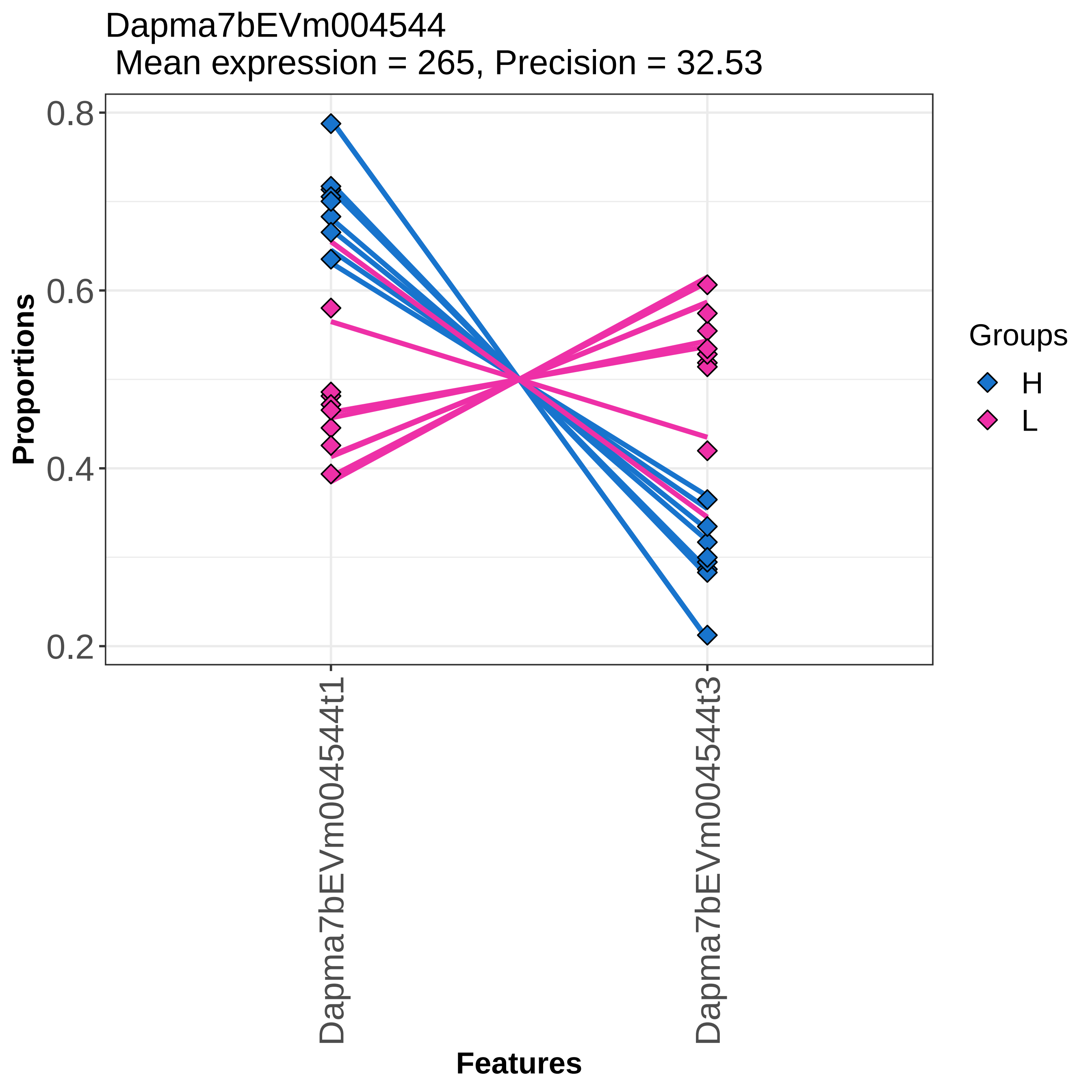
Dapma7bEVm004544
Mean expression = 265, Precision = 32.53
Proportions
0.8
0.6
0.4
0.2
Dapma7bEVm004544t1
Dapma7bEVm004544t3
Features
Groups
H
L

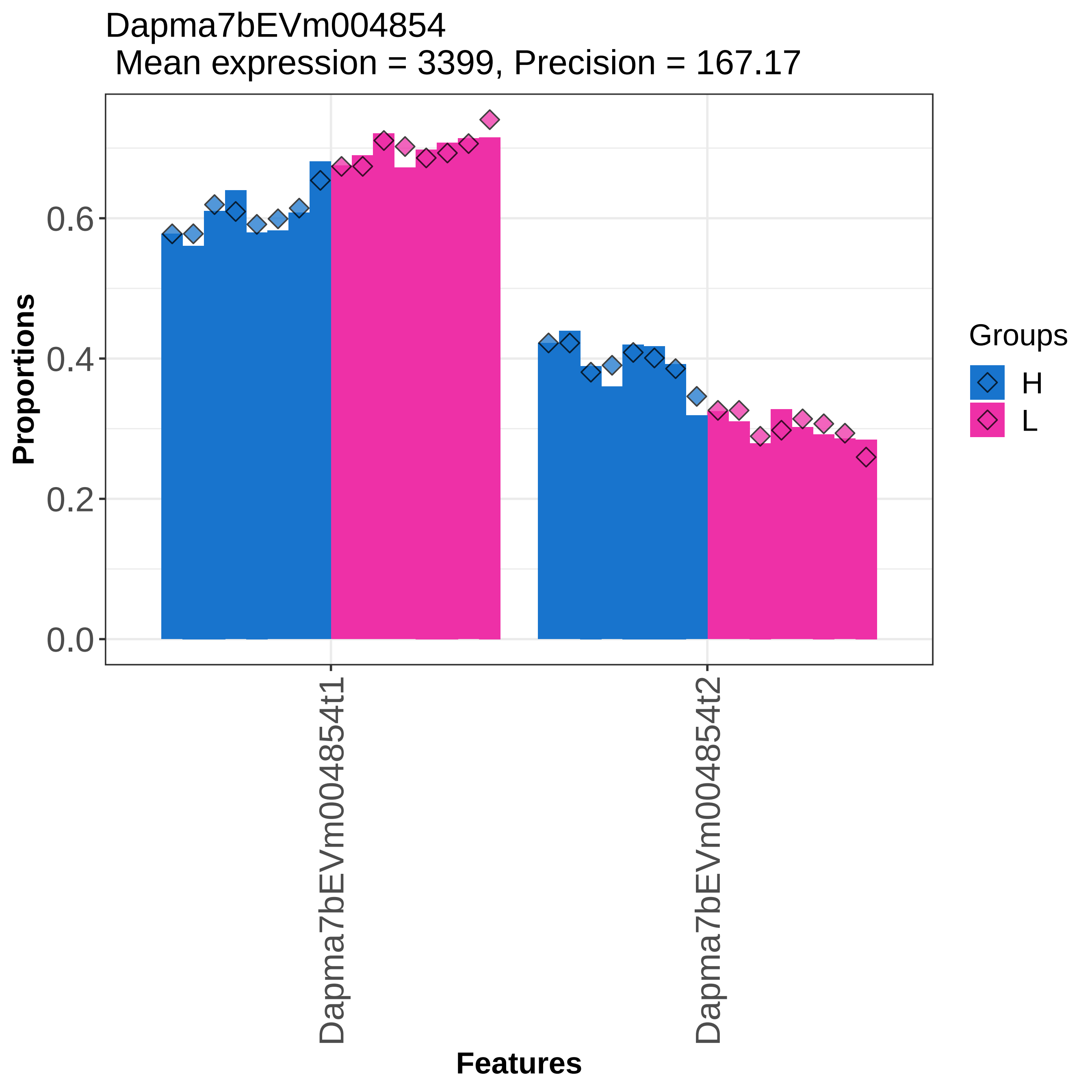

Dapma7bEVm004854
Mean expression = 3399, Precision = 167.17
Proportions
0.0
0.2
0.4
0.6
Dapma7bEVm004854t1
Dapma7bEVm004854t2
Features
Groups
H
L

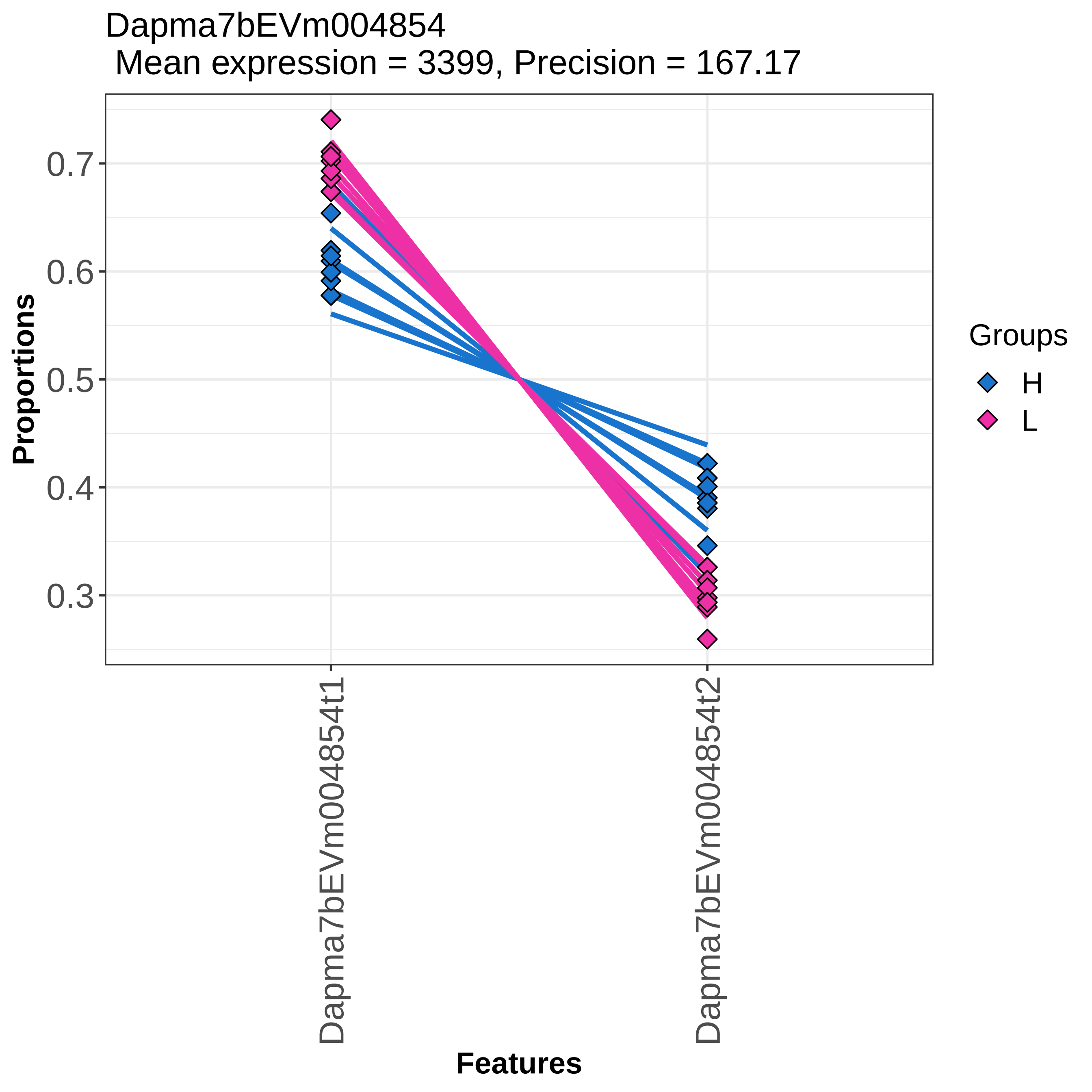

Dapma7bEVm004854
Mean expression = 3399, Precision = 167.17
Proportions
0.7
0.6
0.5
0.4
0.3
Dapma7bEVm004854t1
Dapma7bEVm004854t2
Features
Groups
H
L

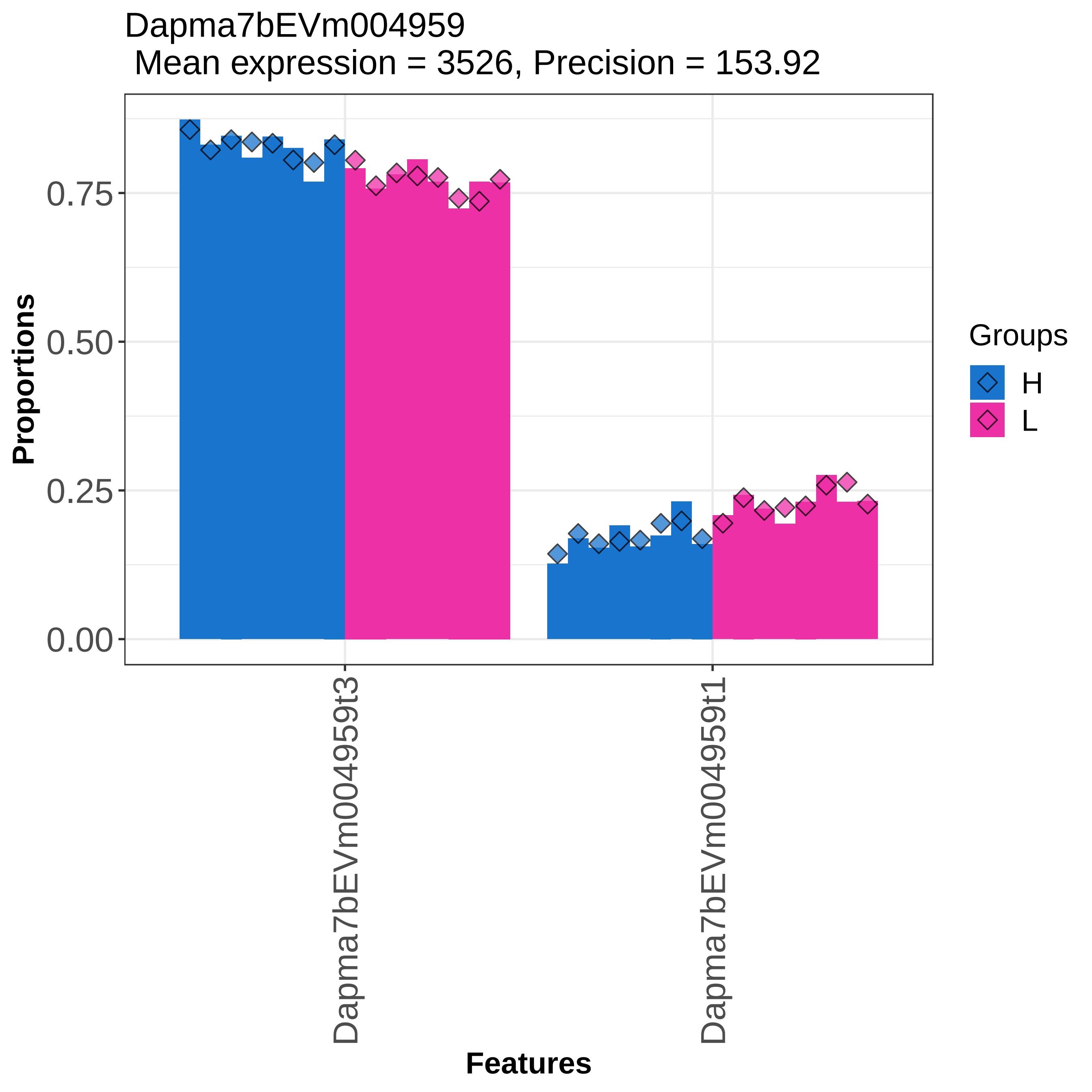
Dapma7bEVm004959
Mean expression = 3526, Precision = 153.92
Proportions
0.00
0.25
0.50
0.75
Dapma7bEVm004959t3
Dapma7bEVm004959t1
Features
Groups
H
L

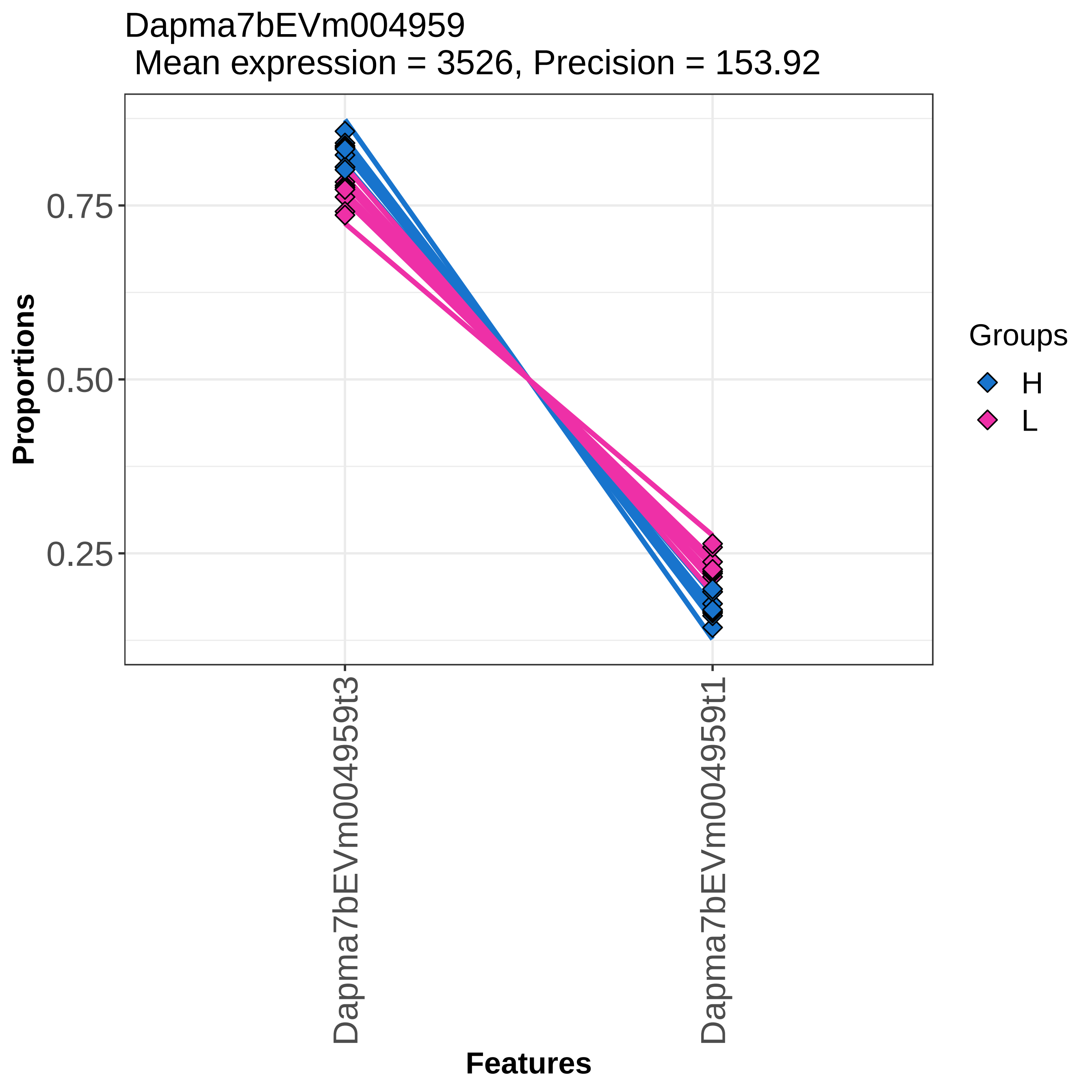

Dapma7bEVm004959
Mean expression = 3526, Precision = 153.92
Proportions
0.75
0.50
0.25
Dapma7bEVm004959t3
Dapma7bEVm004959t1
Features
Groups
H
L

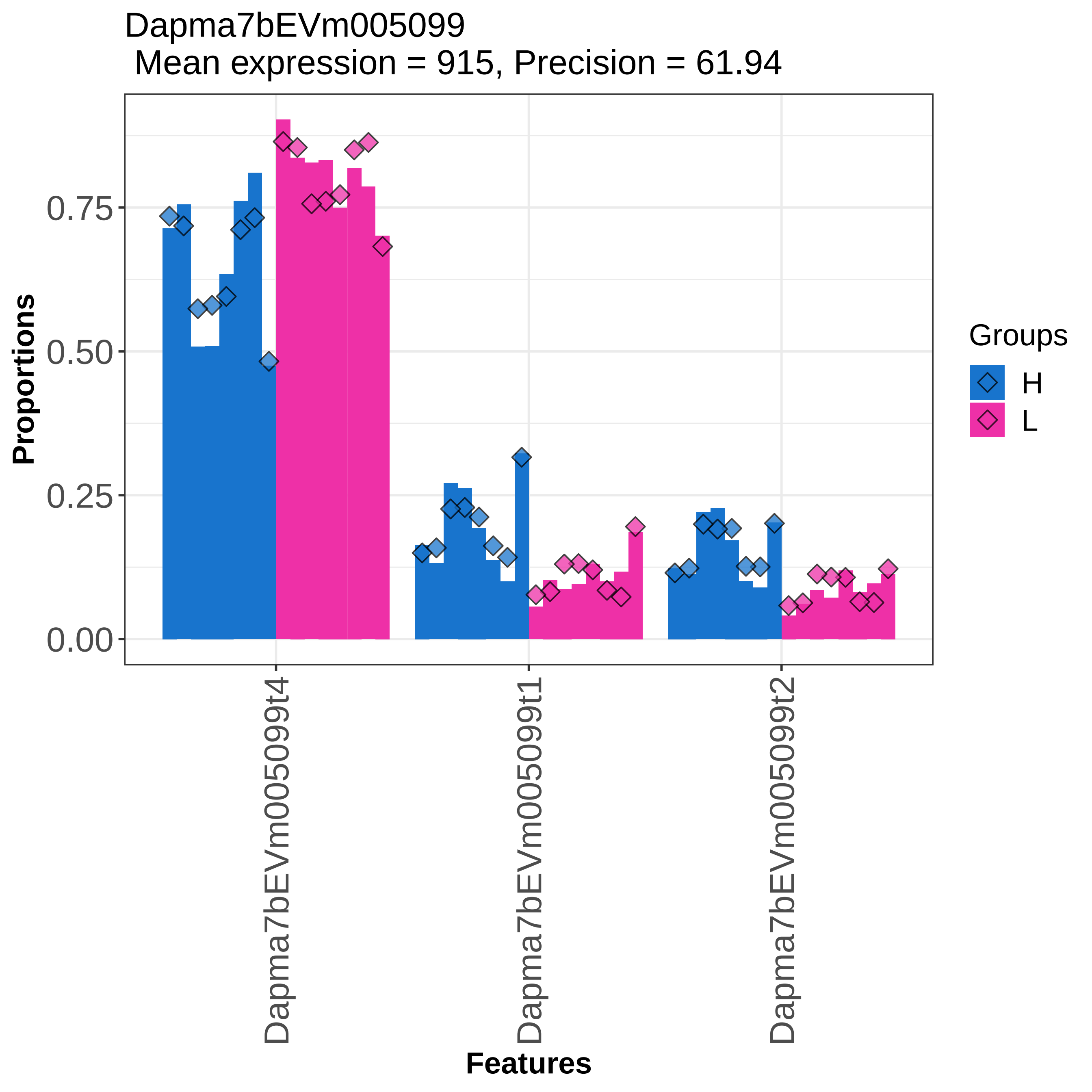
Dapma7bEVm005099
Mean expression = 915, Precision = 61.94
Proportions
0.75
0.50
0.25
0.00
Dapma7bEVm005099t4
Dapma7bEVm005099t1
Dapma7bEVm005099t2
Features
Groups
H
L

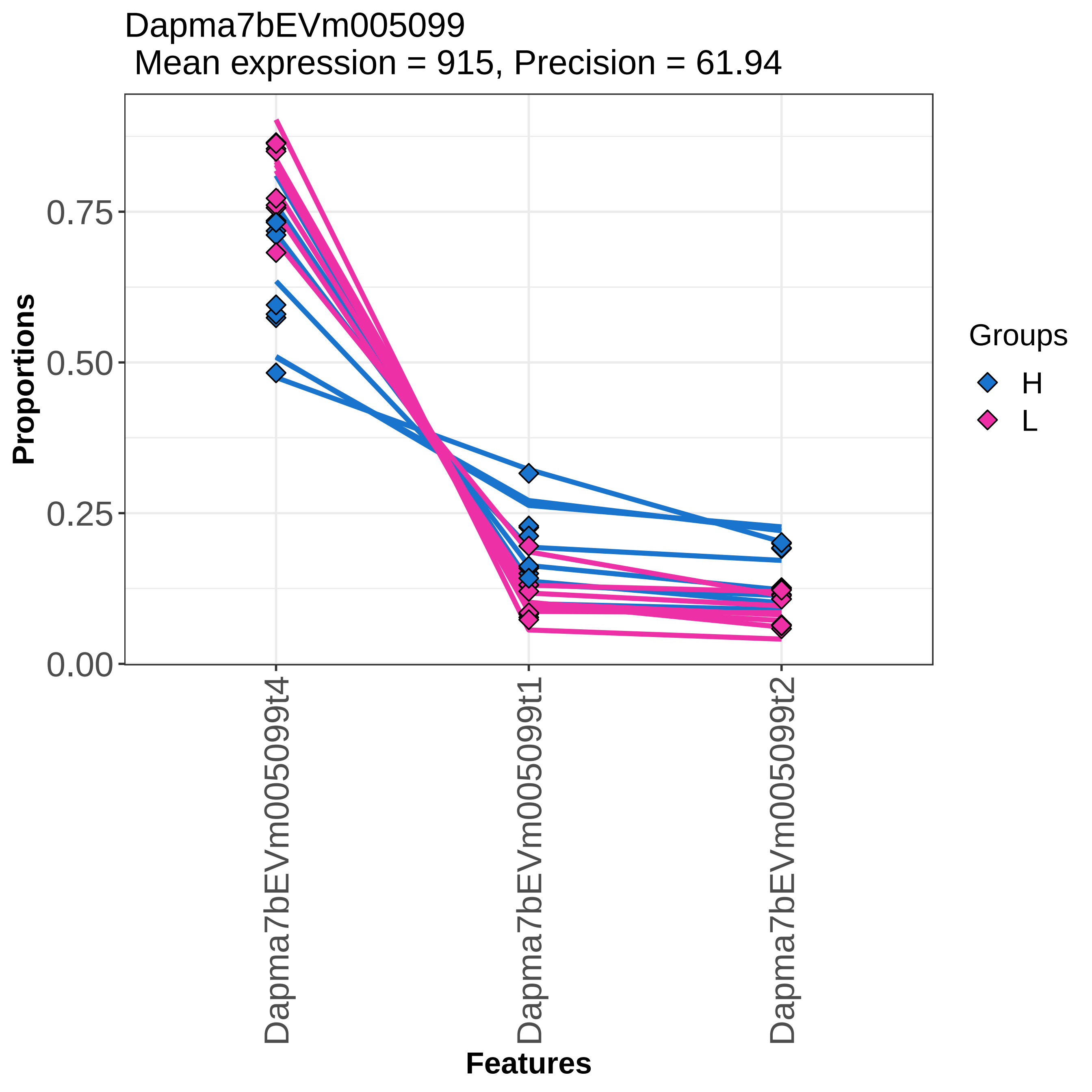
Dapma7bEVm005099
Mean expression = 915, Precision = 61.94
Proportions
0.00
0.25
0.50
0.75
Features
Dapma7bEVm005099t4
Dapma7bEVm005099t1
Dapma7bEVm005099t2
Groups
H
L

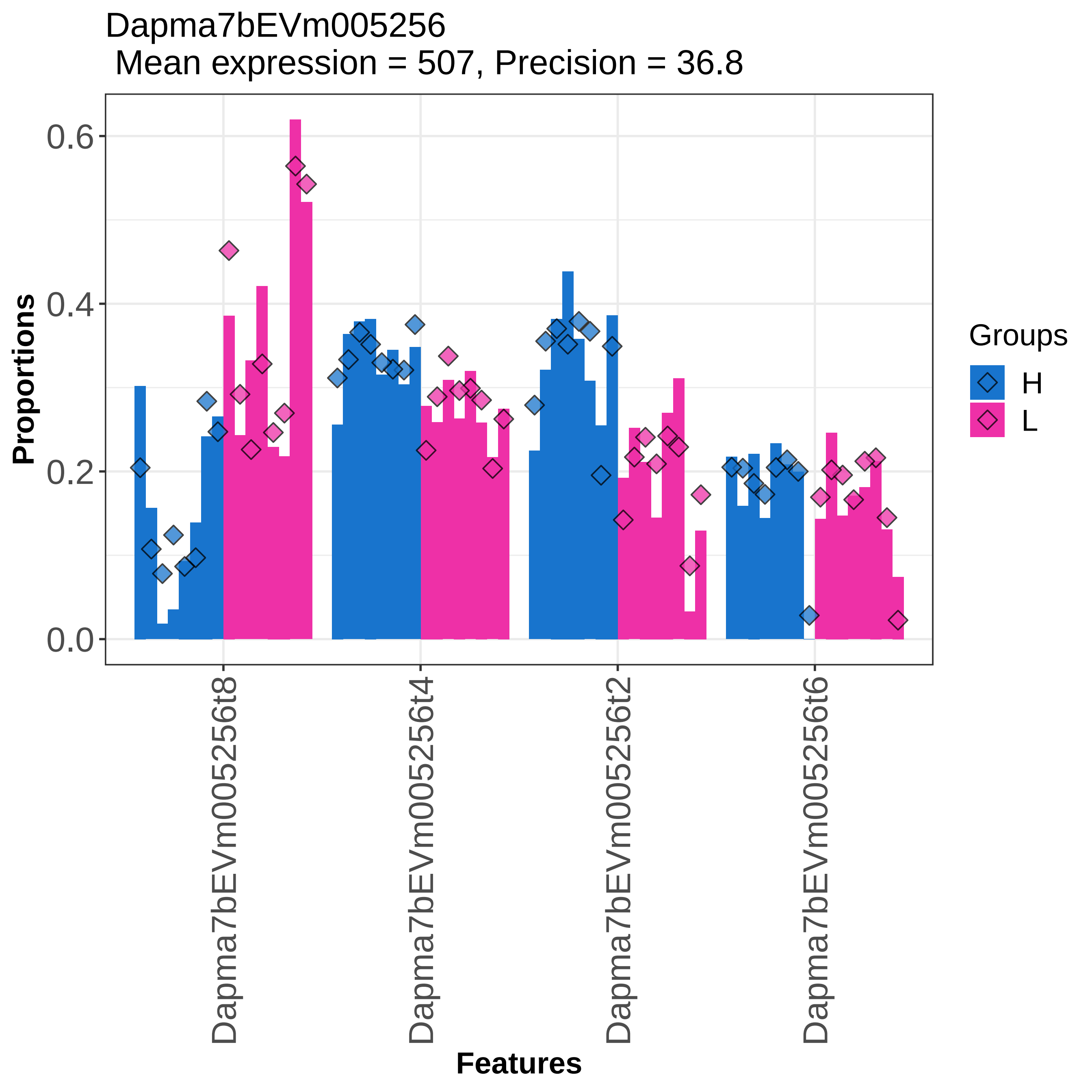
Dapma7bEVm005256
Mean expression = 507, Precision = 36.8
Proportions
0.0
0.2
0.4
0.6
Dapma7bEVm005256t8
Dapma7bEVm005256t4
Dapma7bEVm005256t2
Dapma7bEVm005256t6
Features
Groups
H
L

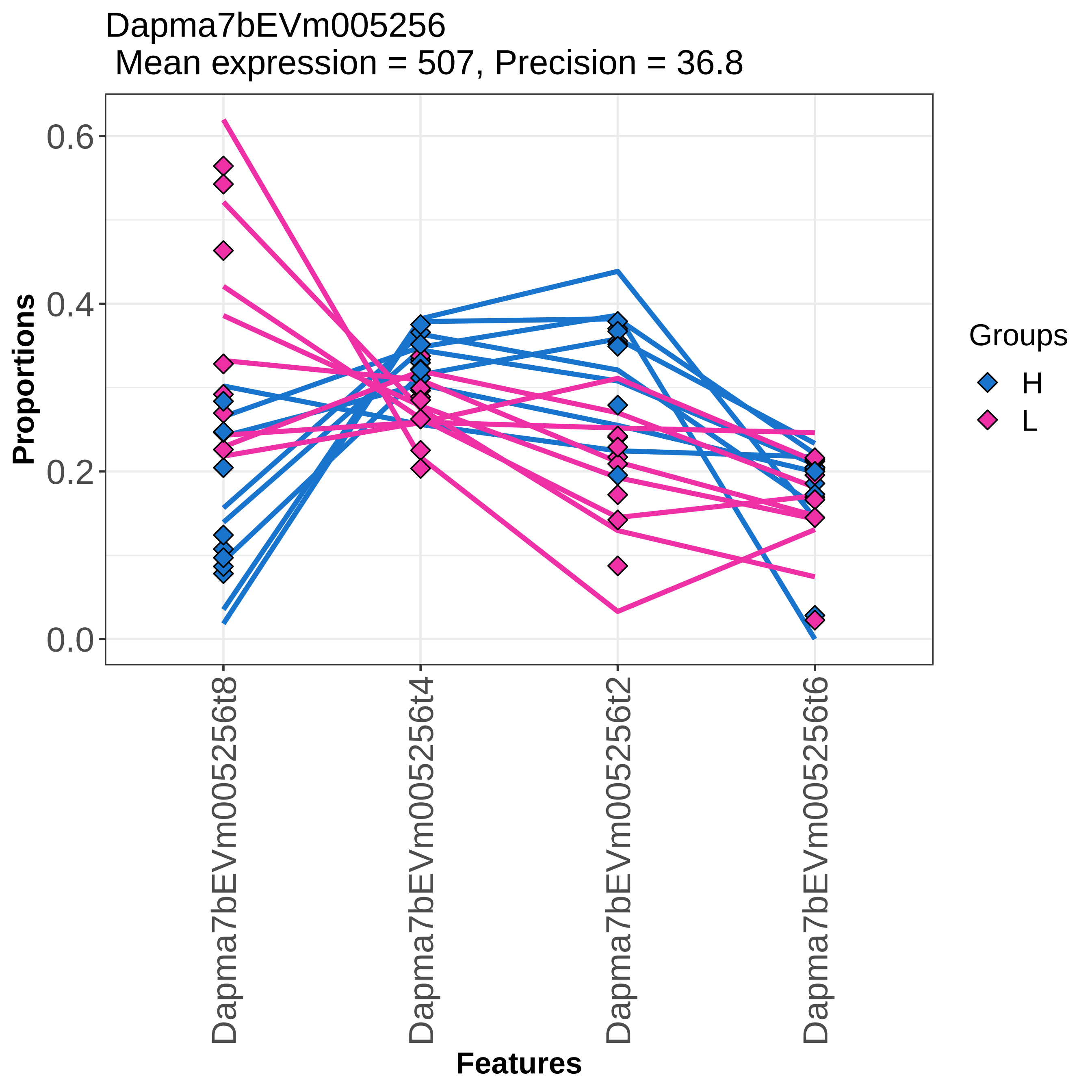
Dapma7bEVm005256
Mean expression = 507, Precision = 36.8
Proportions
0.6
0.4
0.2
0.0
Dapma7bEVm005256t8
Dapma7bEVm005256t4
Dapma7bEVm005256t2
Dapma7bEVm005256t6
Features
Groups
H
L

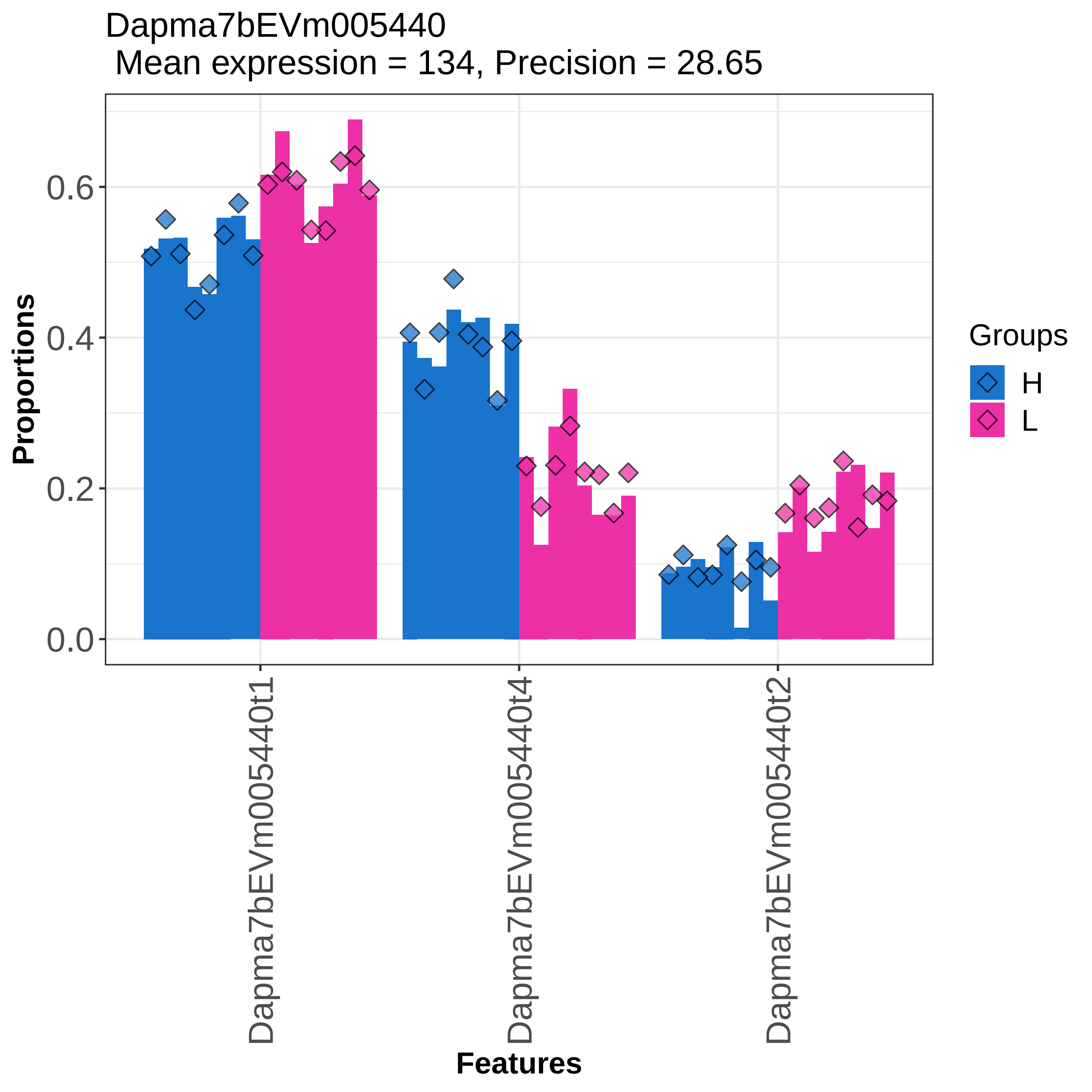

Dapma7bEVm005440
Mean expression = 134, Precision = 28.65
Proportions
0.0
0.2
0.4
0.6
Dapma7bEVm005440t1
Dapma7bEVm005440t4
Dapma7bEVm005440t2
Features
Groups
H
L

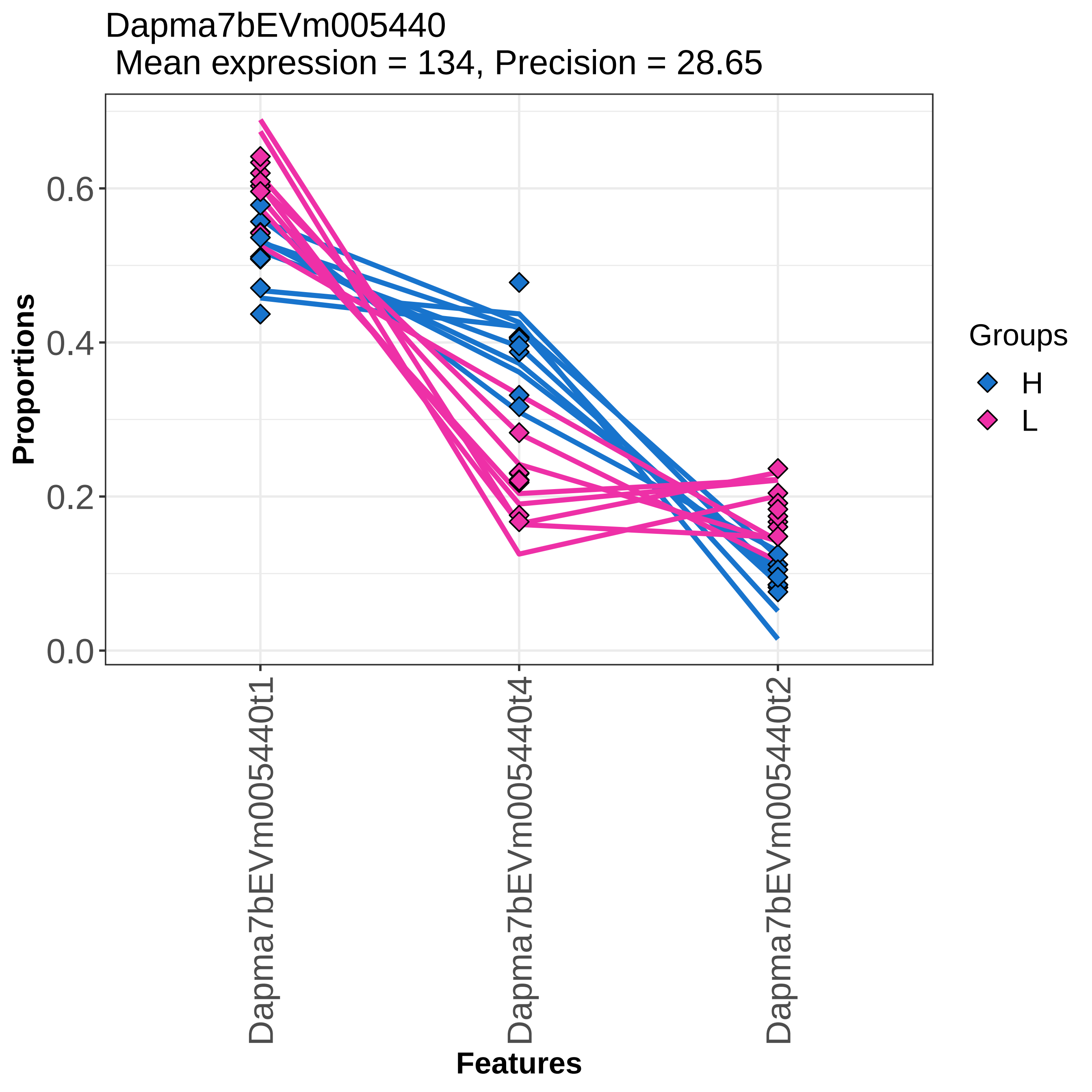

Dapma7bEVm005440
Mean expression = 134, Precision = 28.65
Proportions
0.0
0.2
0.4
0.6
Features
Dapma7bEVm005440t1
Dapma7bEVm005440t4
Dapma7bEVm005440t2
Groups
H
L

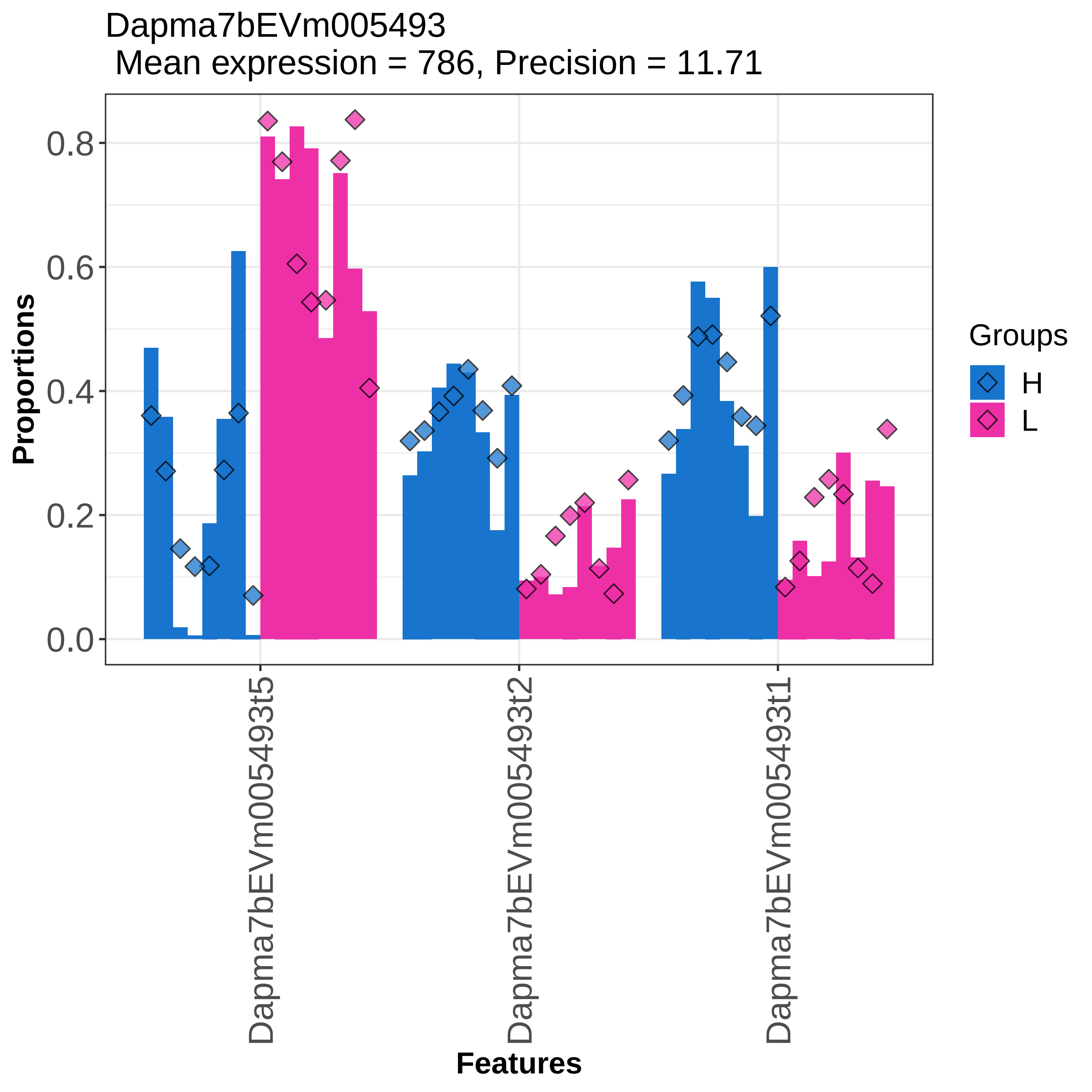

Dapma7bEVm005493
Mean expression = 786, Precision = 11.71
Proportions
0.0
0.2
0.4
0.6
0.8
Dapma7bEVm005493t5
Dapma7bEVm005493t2
Dapma7bEVm005493t1
Features
Groups
H
L

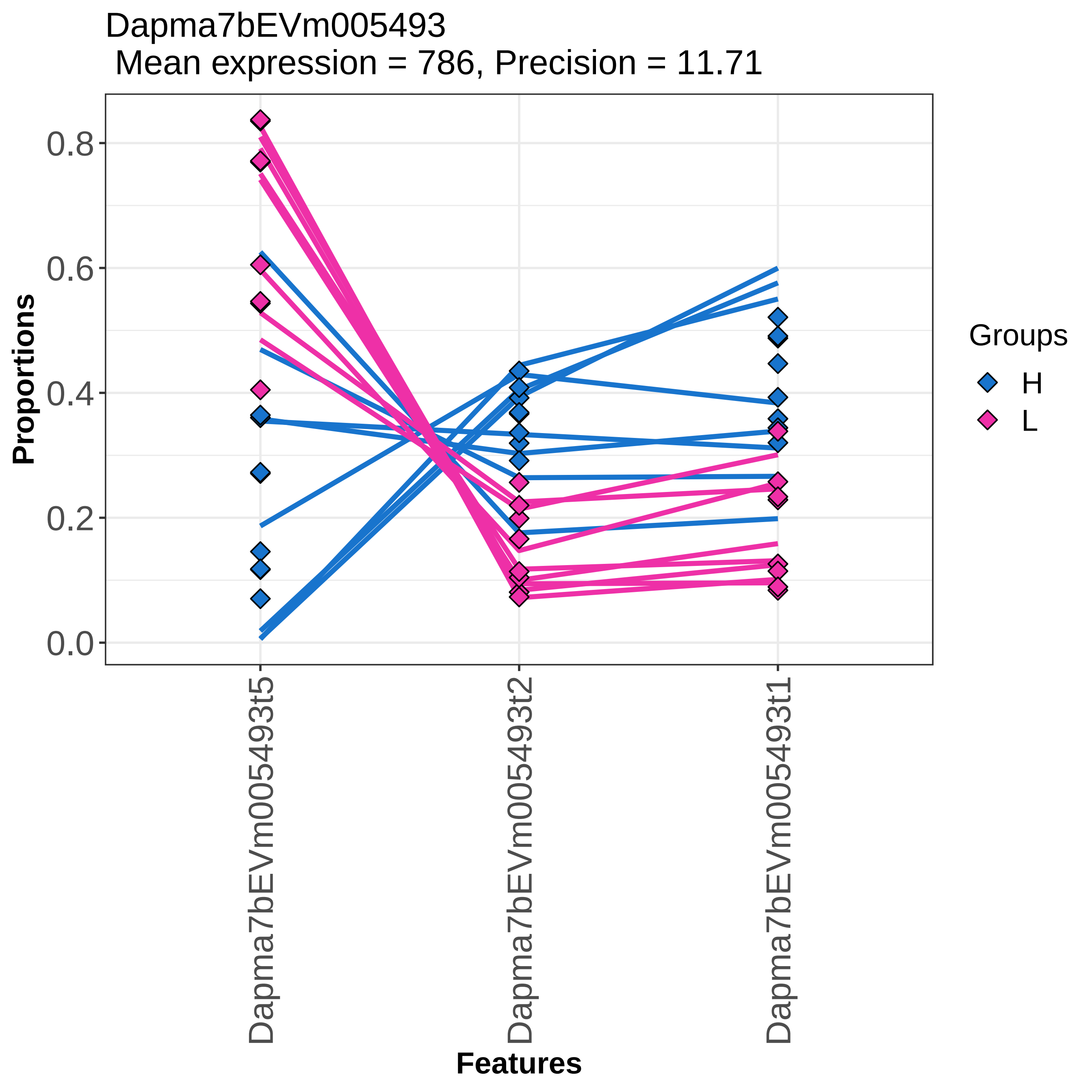

Dapma7bEVm005493
Mean expression = 786, Precision = 11.71
Proportions
0.0
0.2
0.4
0.6
0.8
Dapma7bEVm005493t5
Dapma7bEVm005493t2
Dapma7bEVm005493t1
Features
Groups
H
L

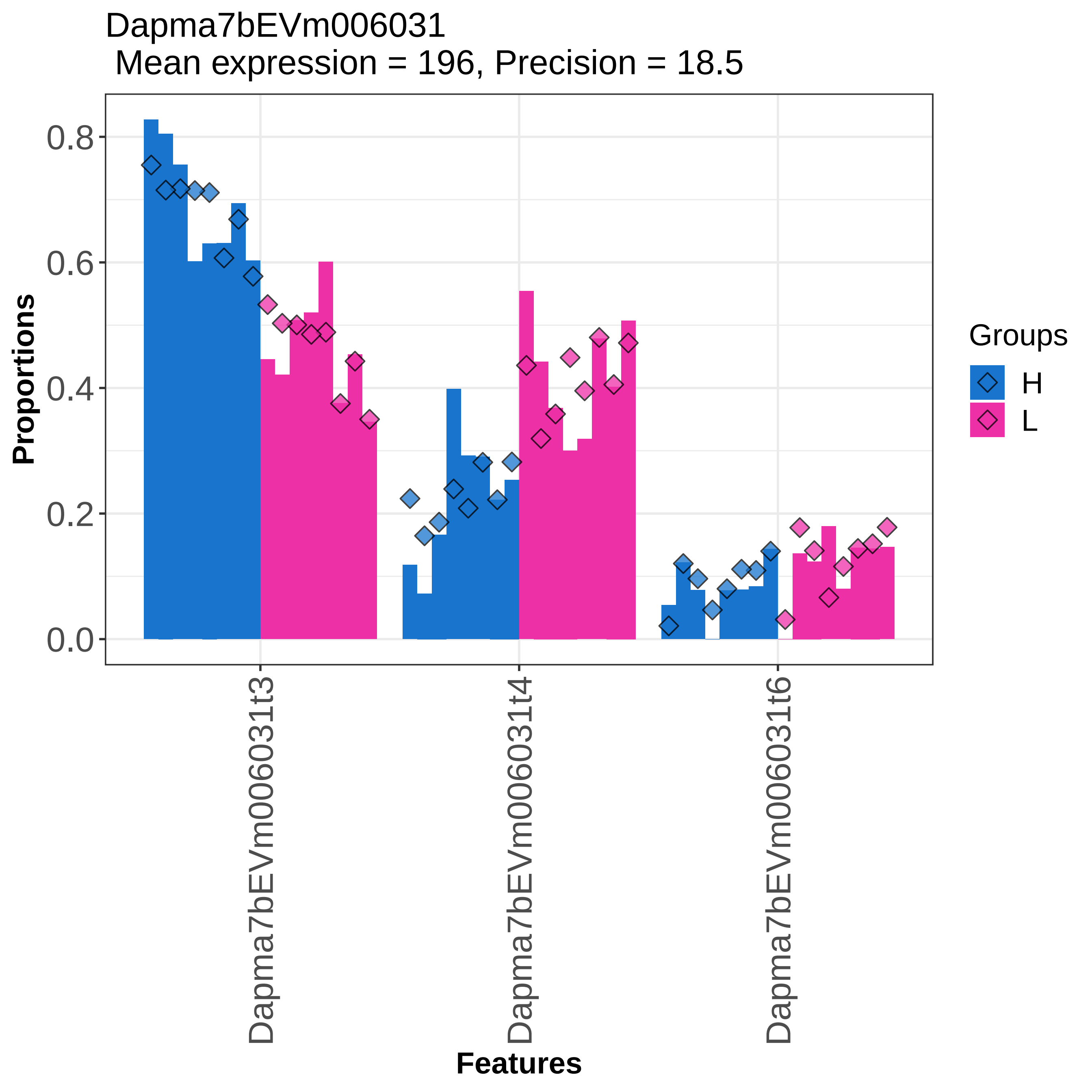
Dapma7bEVm006031
Mean expression = 196, Precision = 18.5
Proportions
0.0
0.2
0.4
0.6
0.8
Dapma7bEVm006031t3
Dapma7bEVm006031t4
Dapma7bEVm006031t6
Features
Groups
H
L

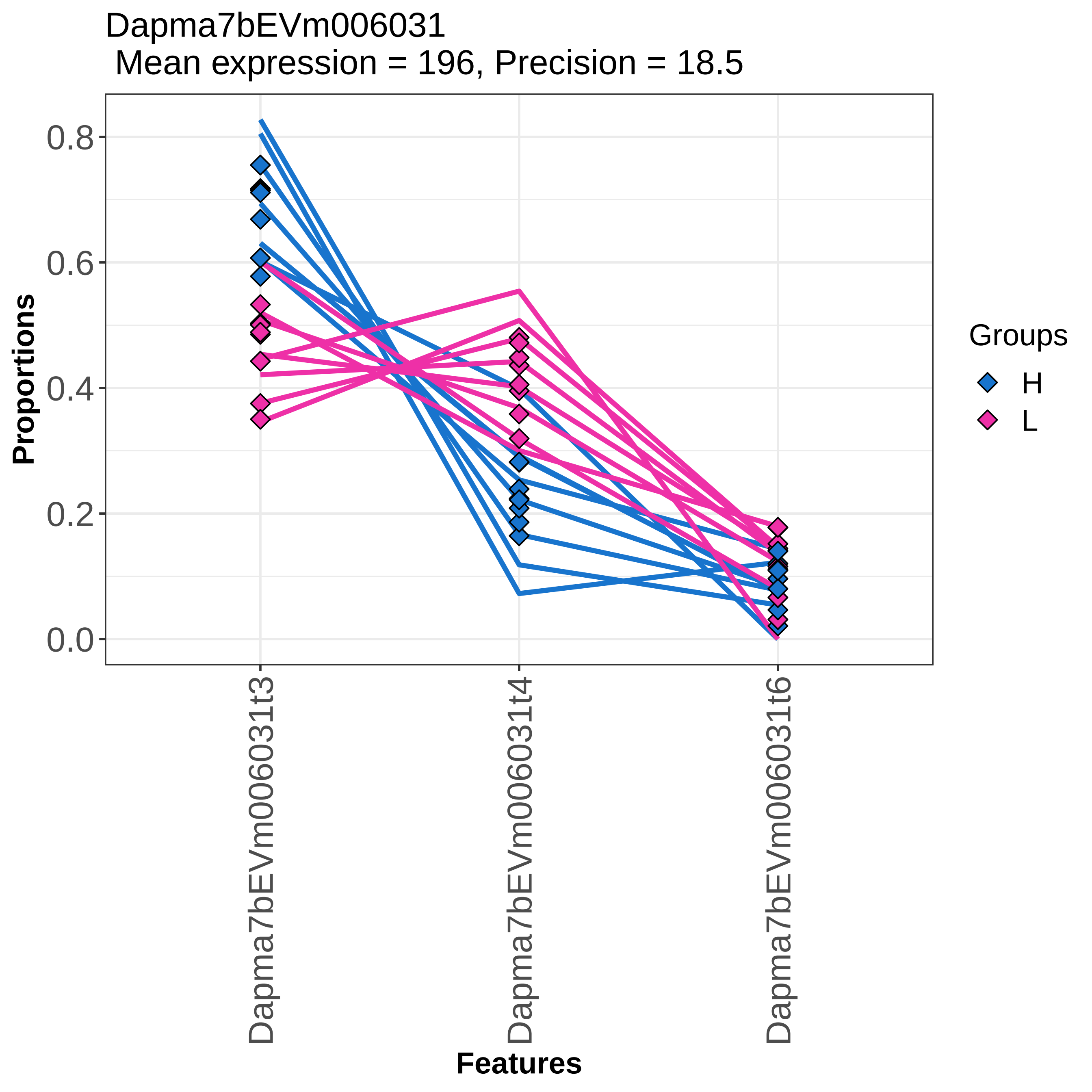
Dapma7bEVm006031
Mean expression = 196, Precision = 18.5
Proportions
0.8
0.6
0.4
0.2
0.0
Dapma7bEVm006031t3
Dapma7bEVm006031t4
Dapma7bEVm006031t6
Features
Groups
H
L

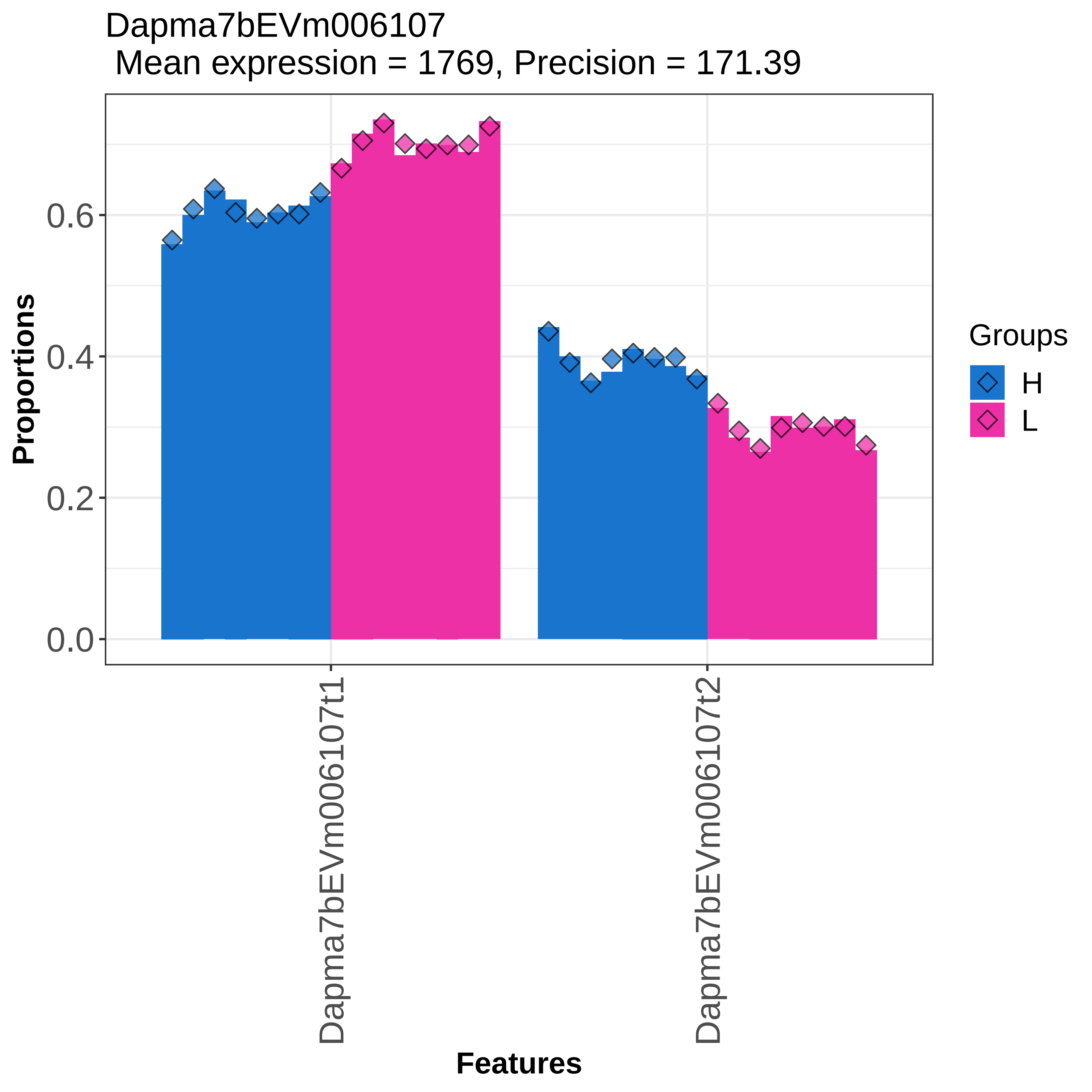
Dapma7bEVm006107
Mean expression = 1769, Precision = 171.39
Proportions
0.0
0.2
0.4
0.6
Features
Dapma7bEVm006107t1
Dapma7bEVm006107t2
Groups
H
L

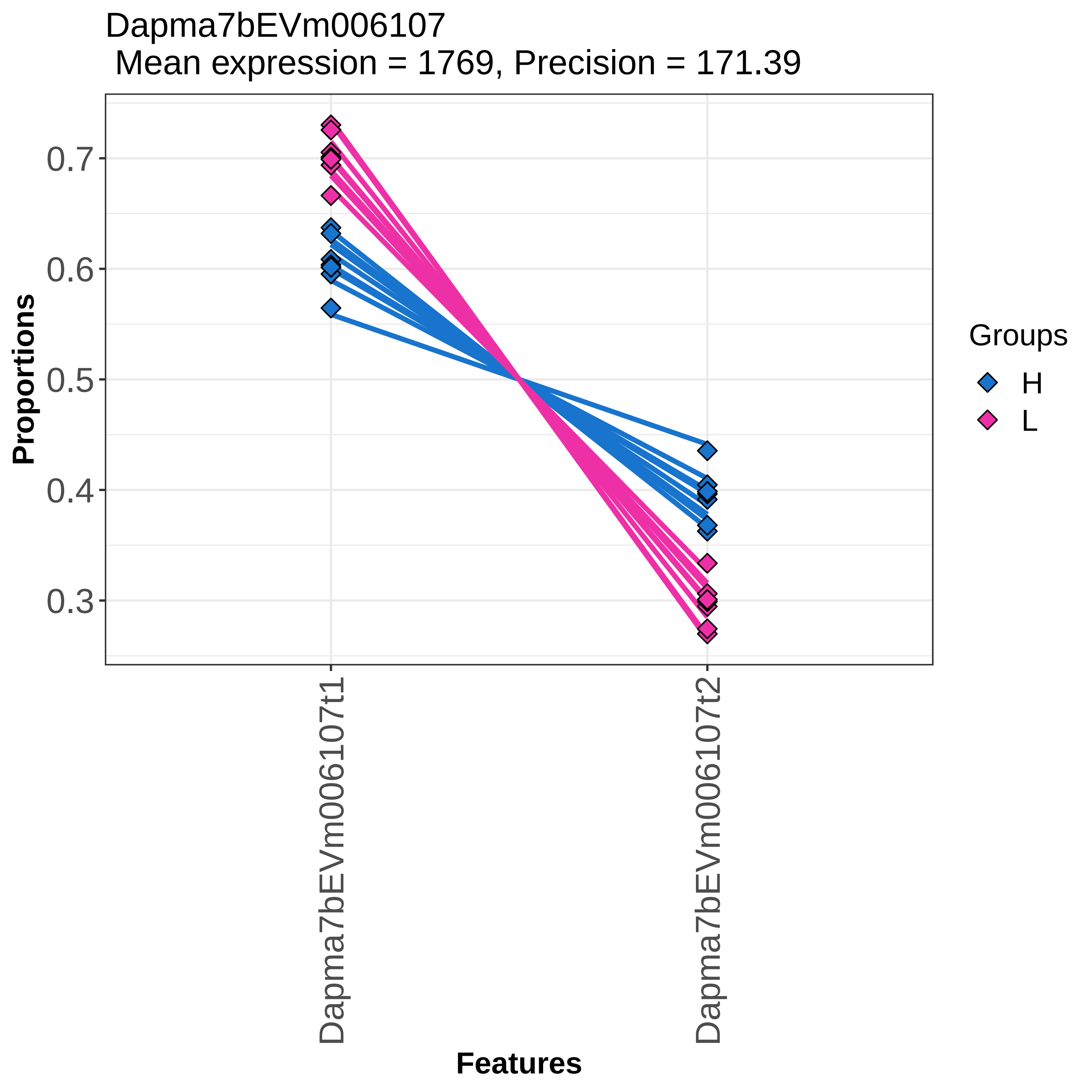

Dapma7bEVm006107
Mean expression = 1769, Precision = 171.39
Proportions
0.7
0.6
0.5
0.4
0.3
Features
Dapma7bEVm006107t1
Dapma7bEVm006107t2
Groups
H
L

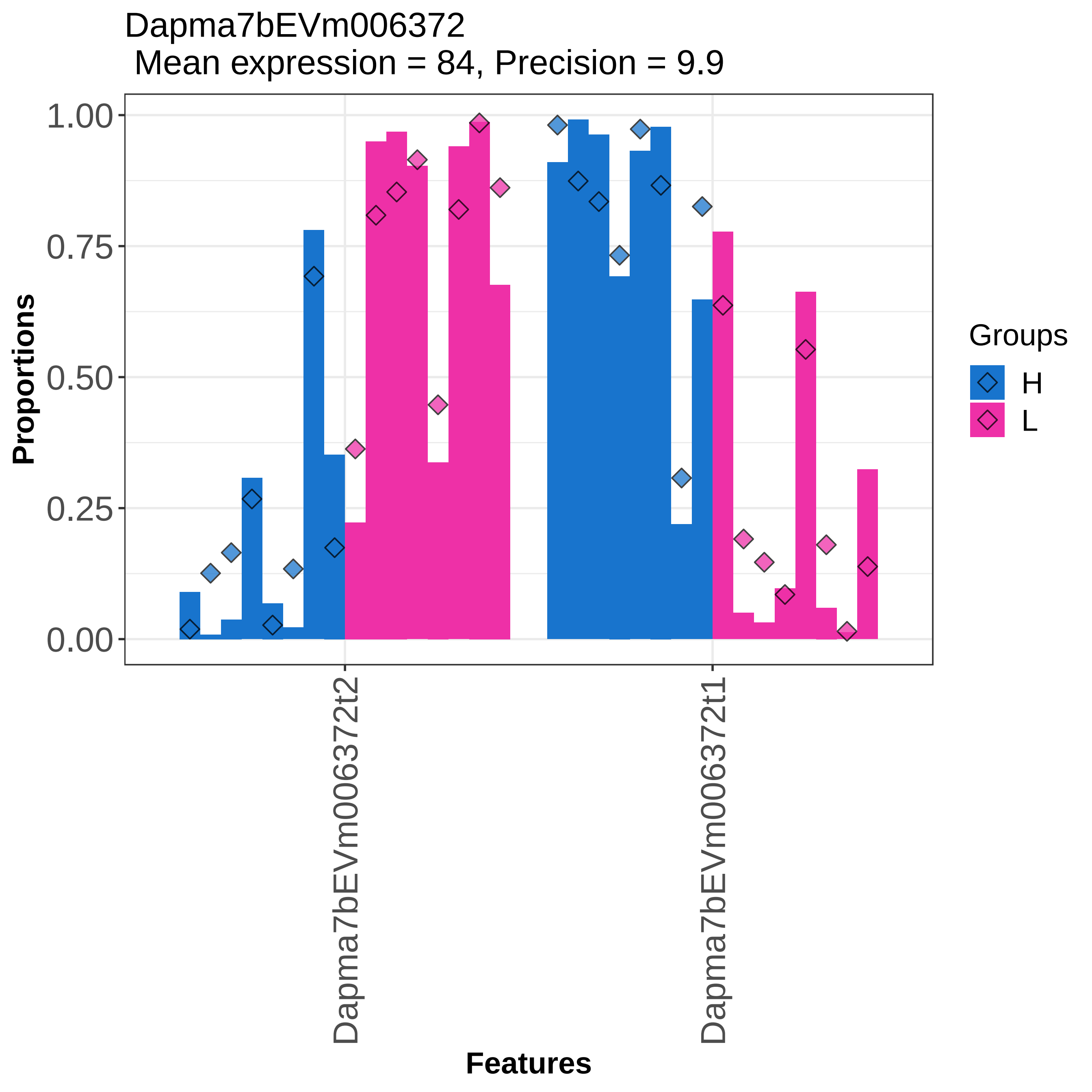
Dapma7bEVm006372
Mean expression = 84, Precision = 9.9
Proportions
1.00
0.75
0.50
0.25
0.00
Features
Dapma7bEVm006372t2
Dapma7bEVm006372t1
Groups
H
L

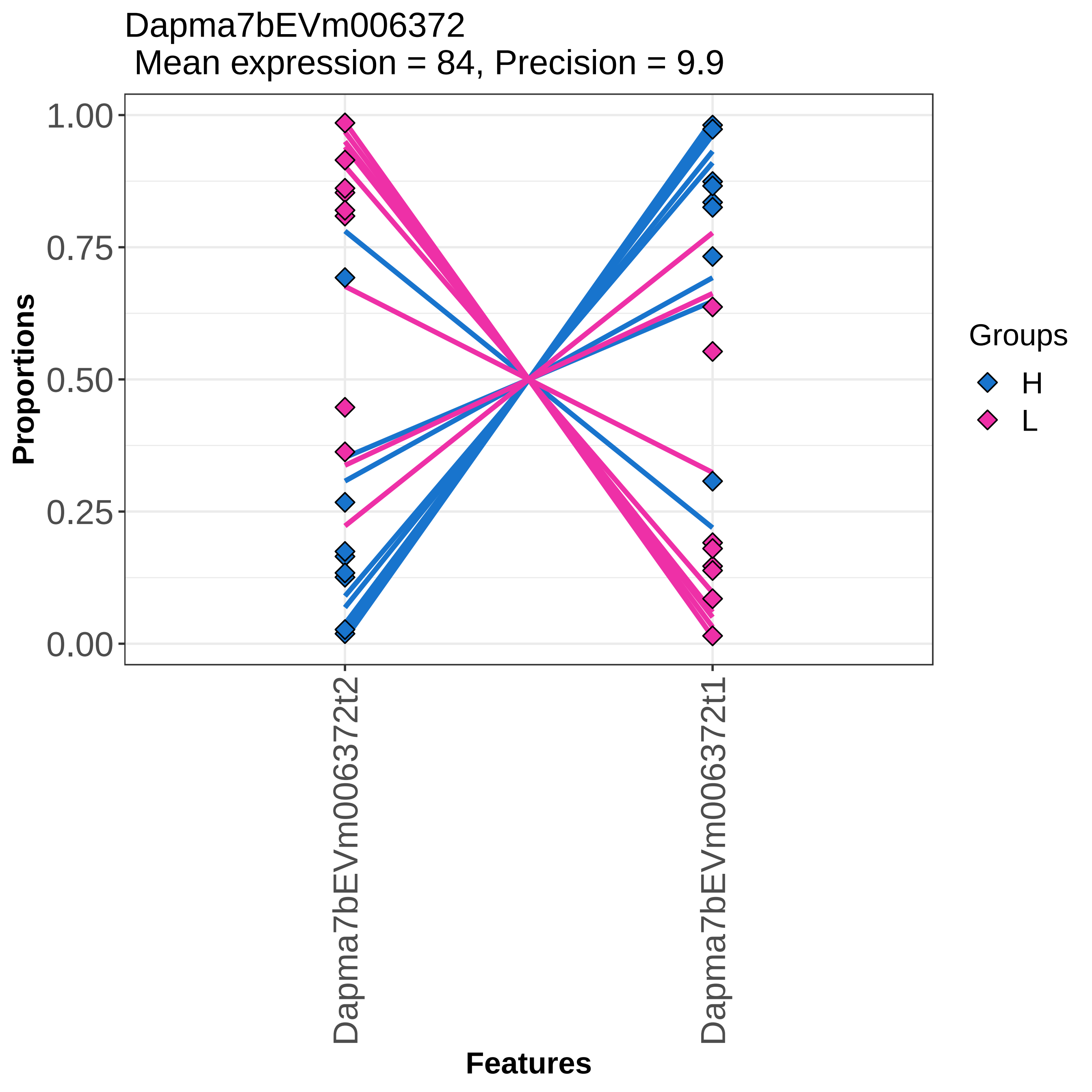
Dapma7bEVm006372
Mean expression = 84, Precision = 9.9
Proportions
1.00
0.75
0.50
0.25
0.00
Dapma7bEVm006372t2
Dapma7bEVm006372t1
Features
Groups
H
L

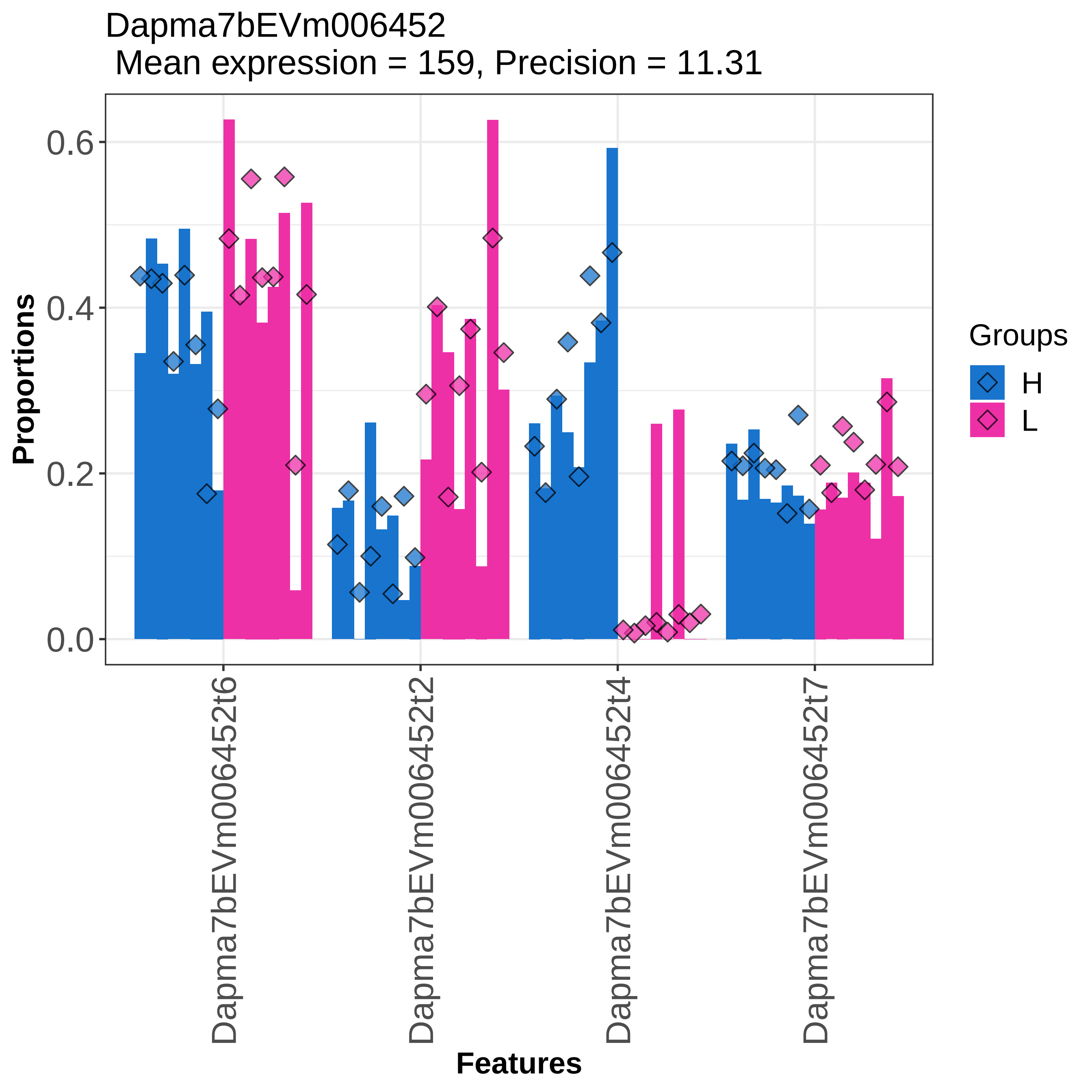

Dapma7bEVm006452
Mean expression = 159, Precision = 11.31
Proportions
0.0
0.2
0.4
0.6
Dapma7bEVm006452t6
Dapma7bEVm006452t2
Dapma7bEVm006452t4
Dapma7bEVm006452t7
Features
Groups
H
L

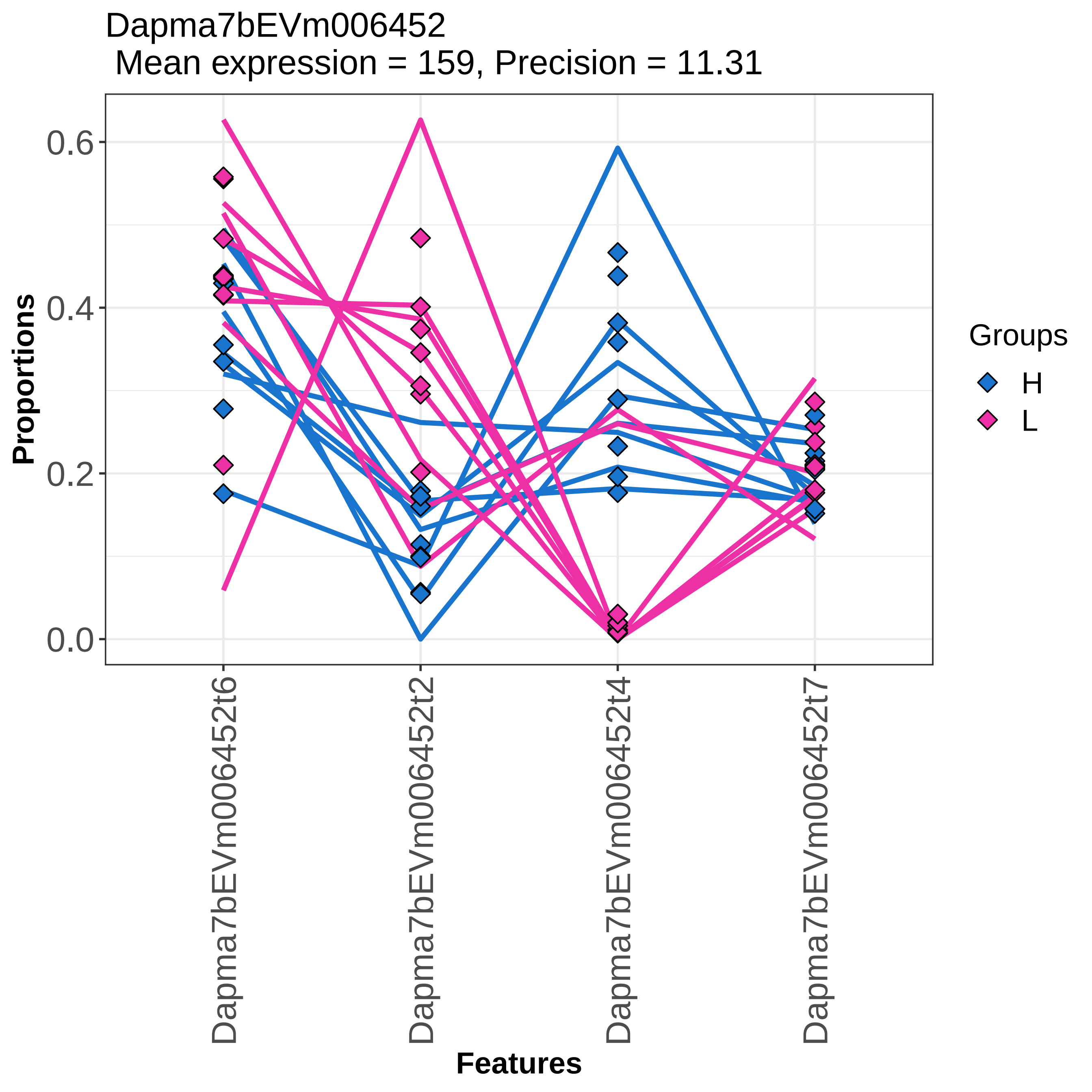
Dapma7bEVm006452
Mean expression = 159, Precision = 11.31
Proportions
0.6
0.4
0.2
0.0
Dapma7bEVm006452t6
Dapma7bEVm006452t2
Dapma7bEVm006452t4
Dapma7bEVm006452t7
Features
Groups
H
L

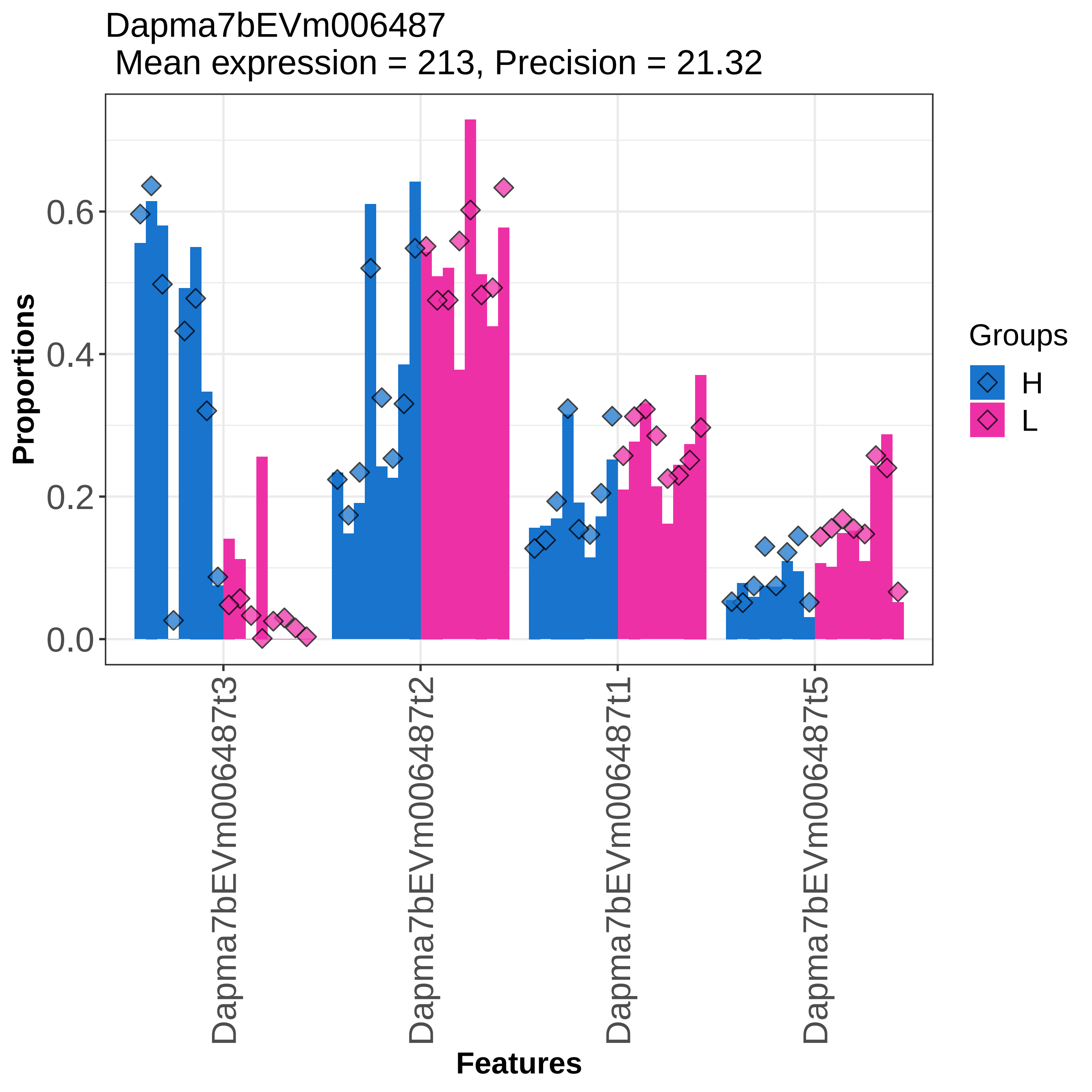

Dapma7bEVm006487
Mean expression = 213, Precision = 21.32
Proportions
0.0
0.2
0.4
0.6
Dapma7bEVm006487t3
Dapma7bEVm006487t2
Dapma7bEVm006487t1
Dapma7bEVm006487t5
Features
Groups
H
L

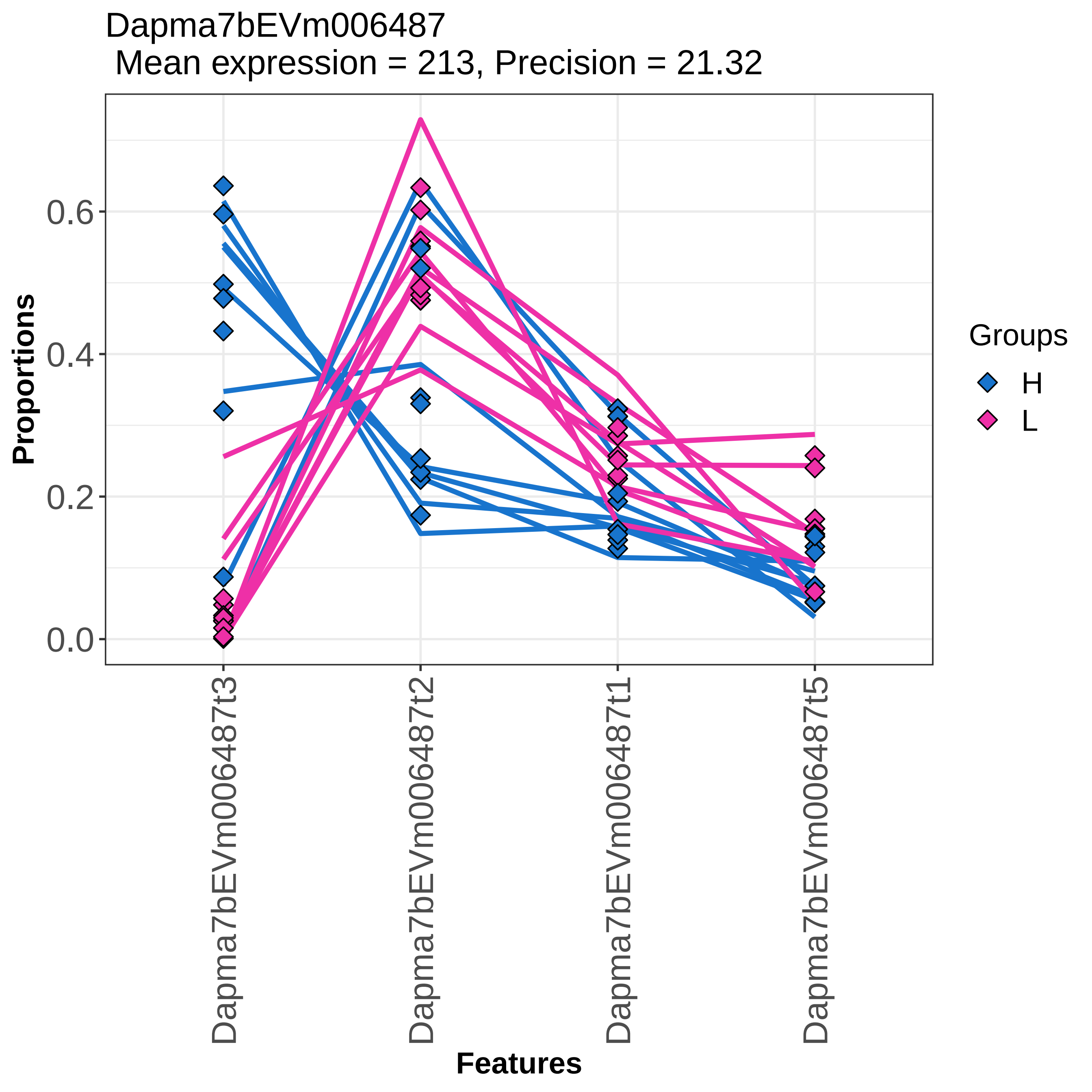

Dapma7bEVm006487
Mean expression = 213, Precision = 21.32
Proportions
0.0
0.2
0.4
0.6
Dapma7bEVm006487t3
Dapma7bEVm006487t2
Dapma7bEVm006487t1
Dapma7bEVm006487t5
Features
Groups
H
L

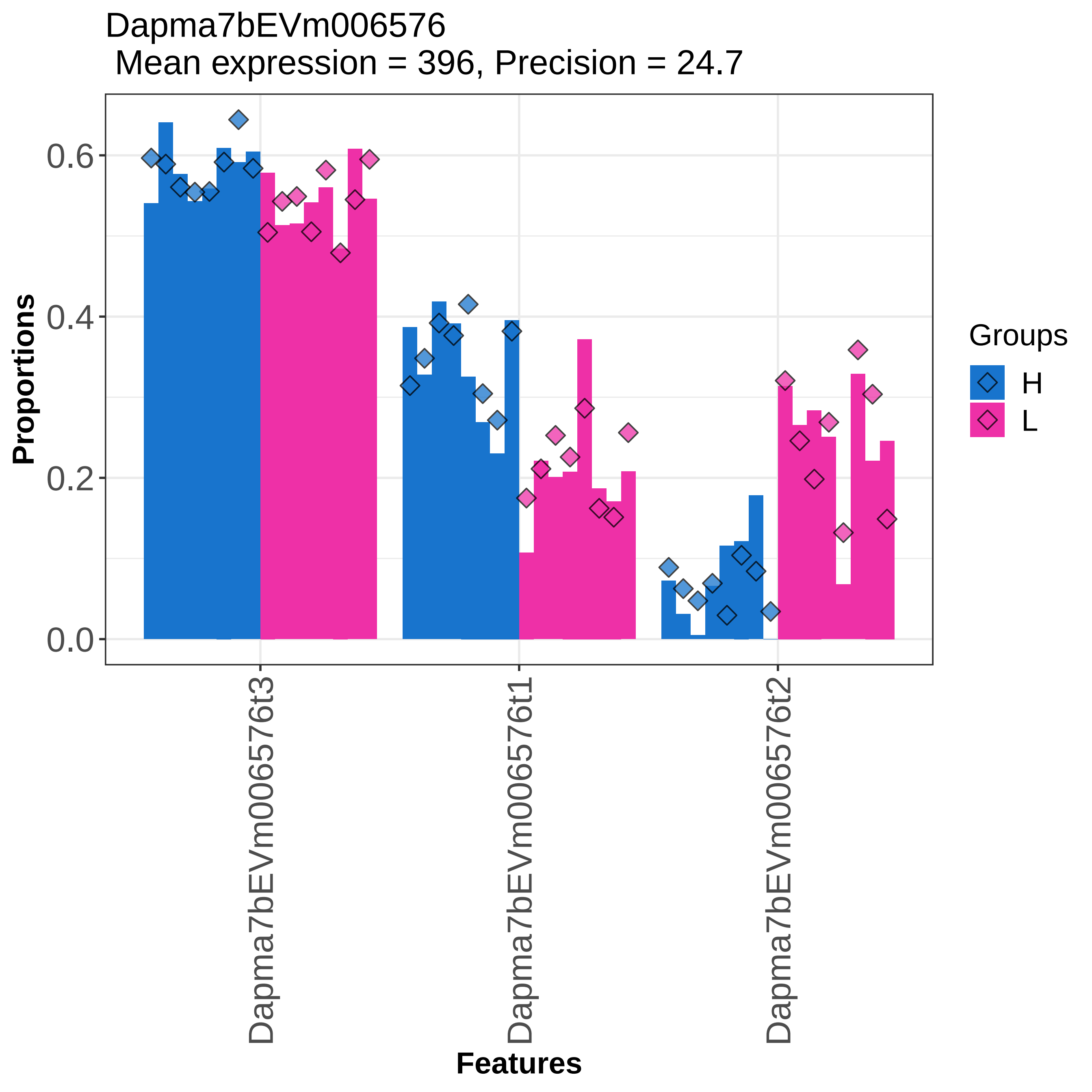

Dapma7bEVm006576
Mean expression = 396, Precision = 24.7
Proportions
0.6
0.4
0.2
0.0
Dapma7bEVm006576t3
Dapma7bEVm006576t1
Dapma7bEVm006576t2
Features
Groups
H
L

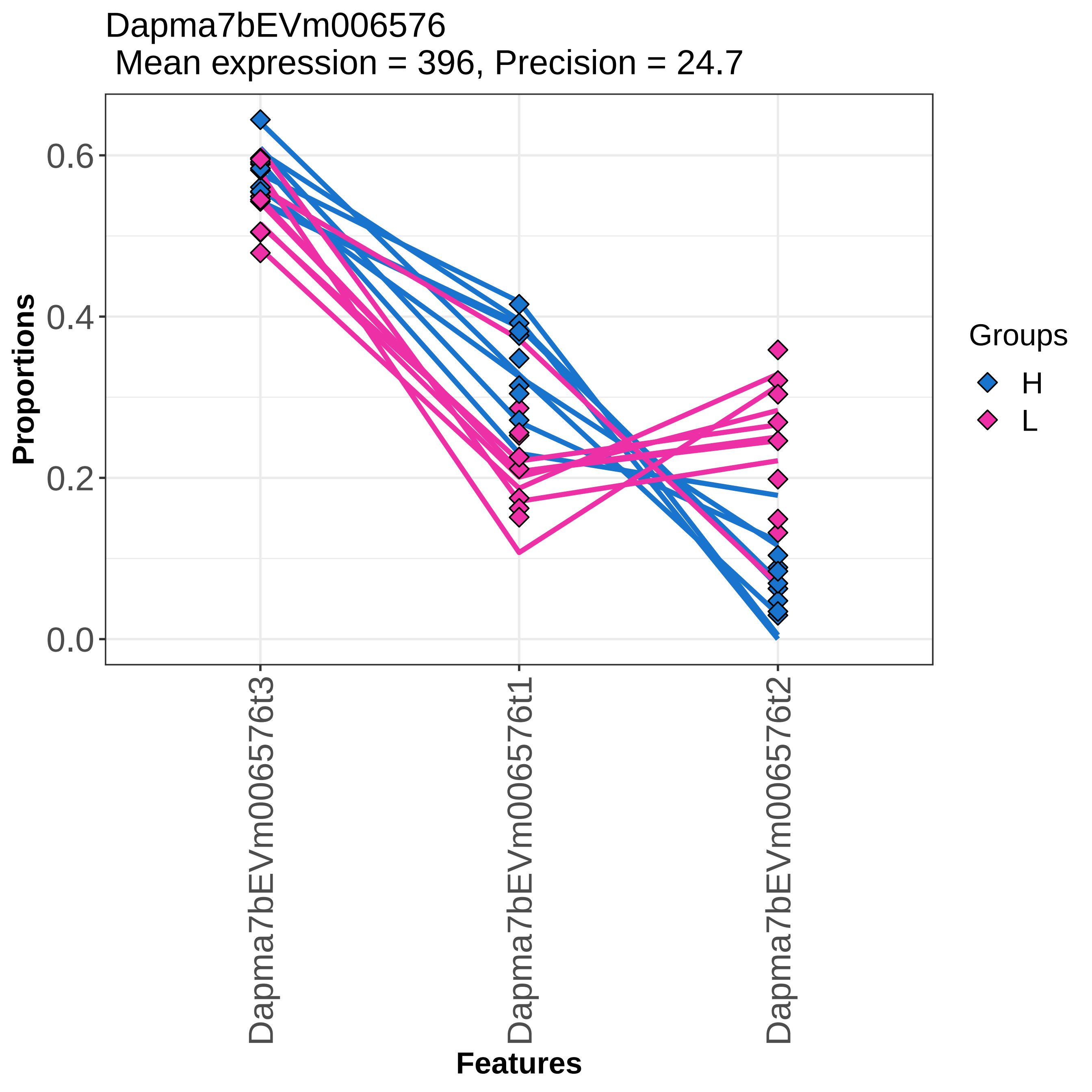
Dapma7bEVm006576
Mean expression = 396, Precision = 24.7
Proportions
0.0
0.2
0.4
0.6
Dapma7bEVm006576t3
Dapma7bEVm006576t1
Dapma7bEVm006576t2
Features
Groups
H
L

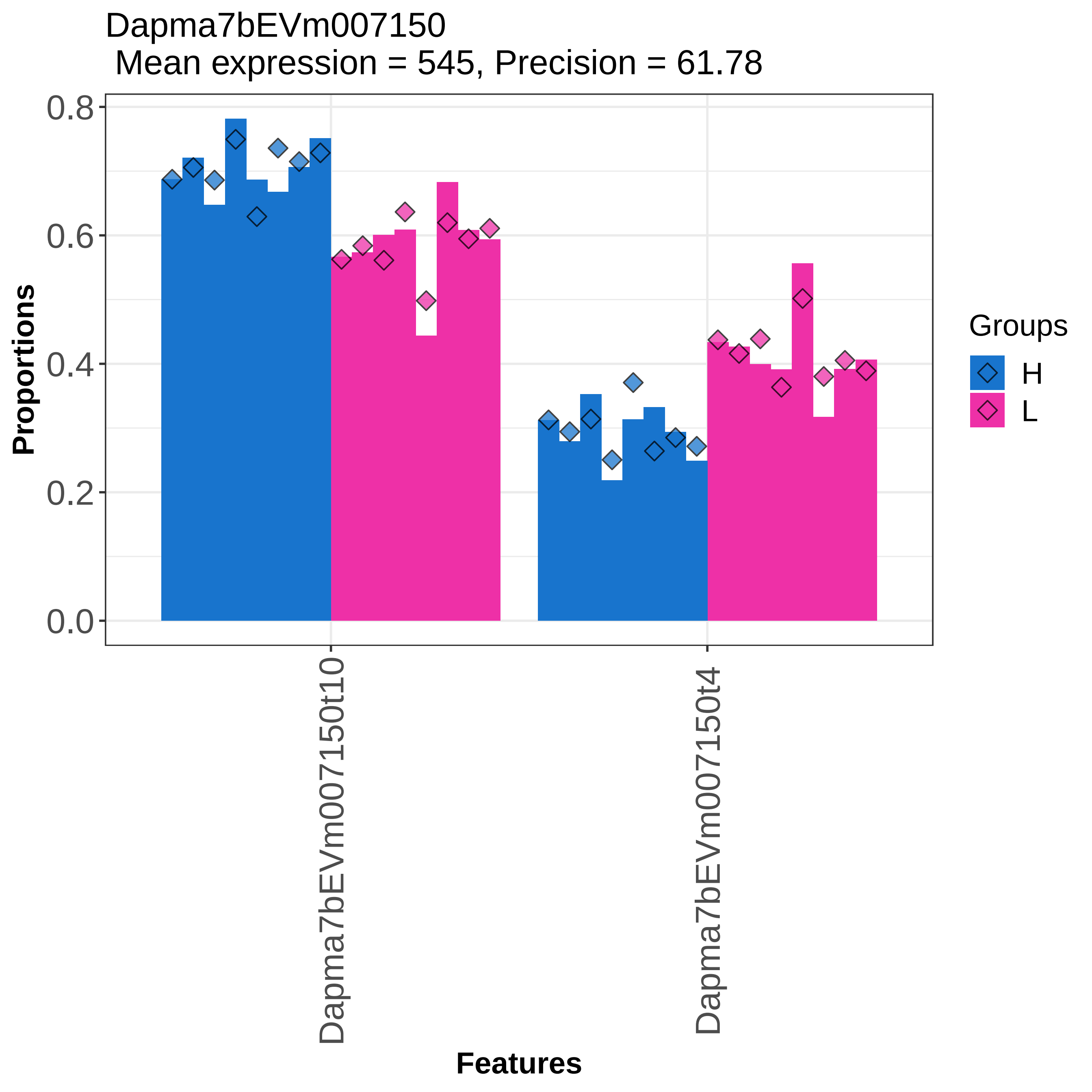

Dapma7bEVm007150
Mean expression = 545, Precision = 61.78
Proportions
0.0
0.2
0.4
0.6
0.8
Dapma7bEVm007150t10
Dapma7bEVm007150t4
Features
Groups
H
L

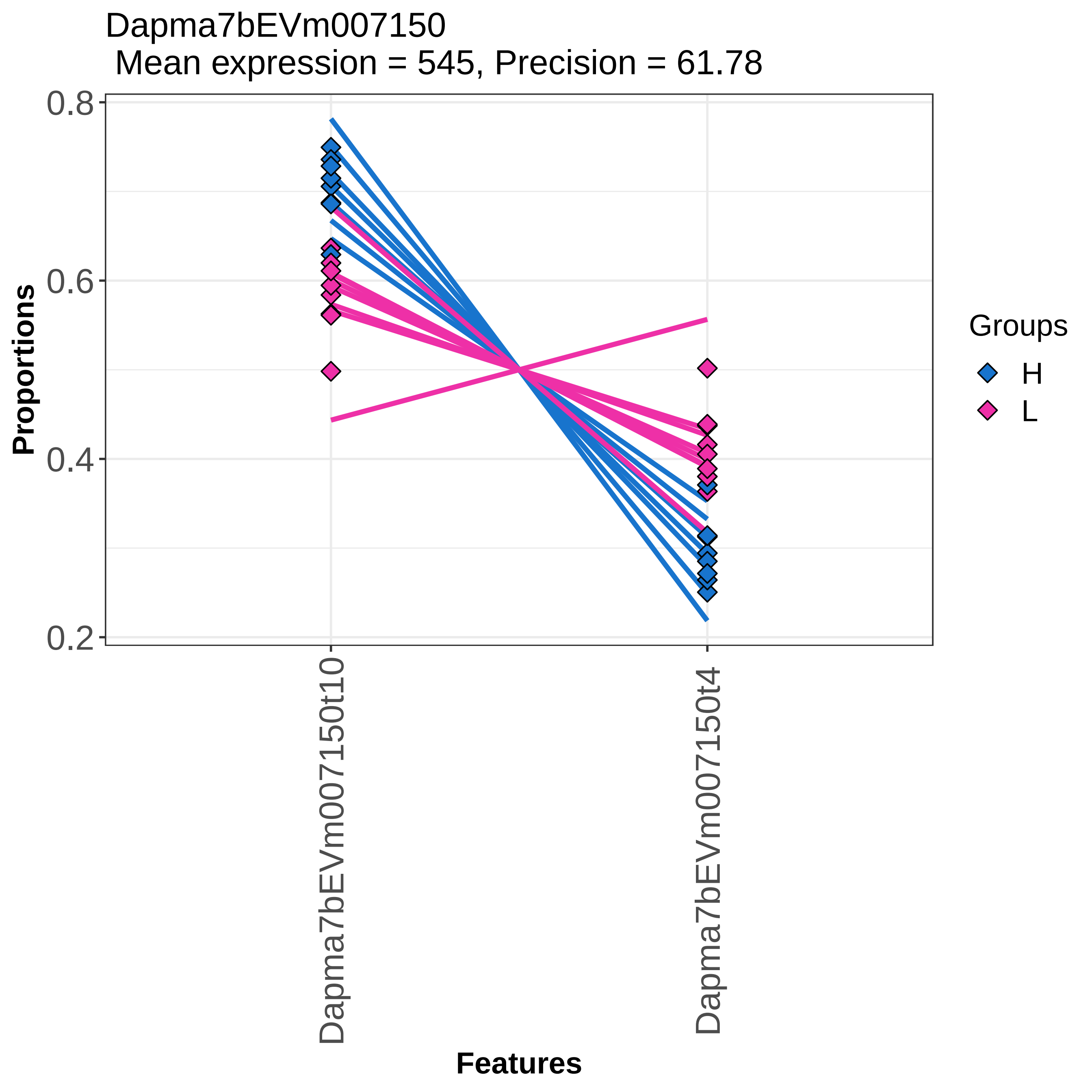

Dapma7bEVm007150
Mean expression = 545, Precision = 61.78
Proportions
0.8
0.6
0.4
0.2
Dapma7bEVm007150t10
Dapma7bEVm007150t4
Features
Groups
H
L

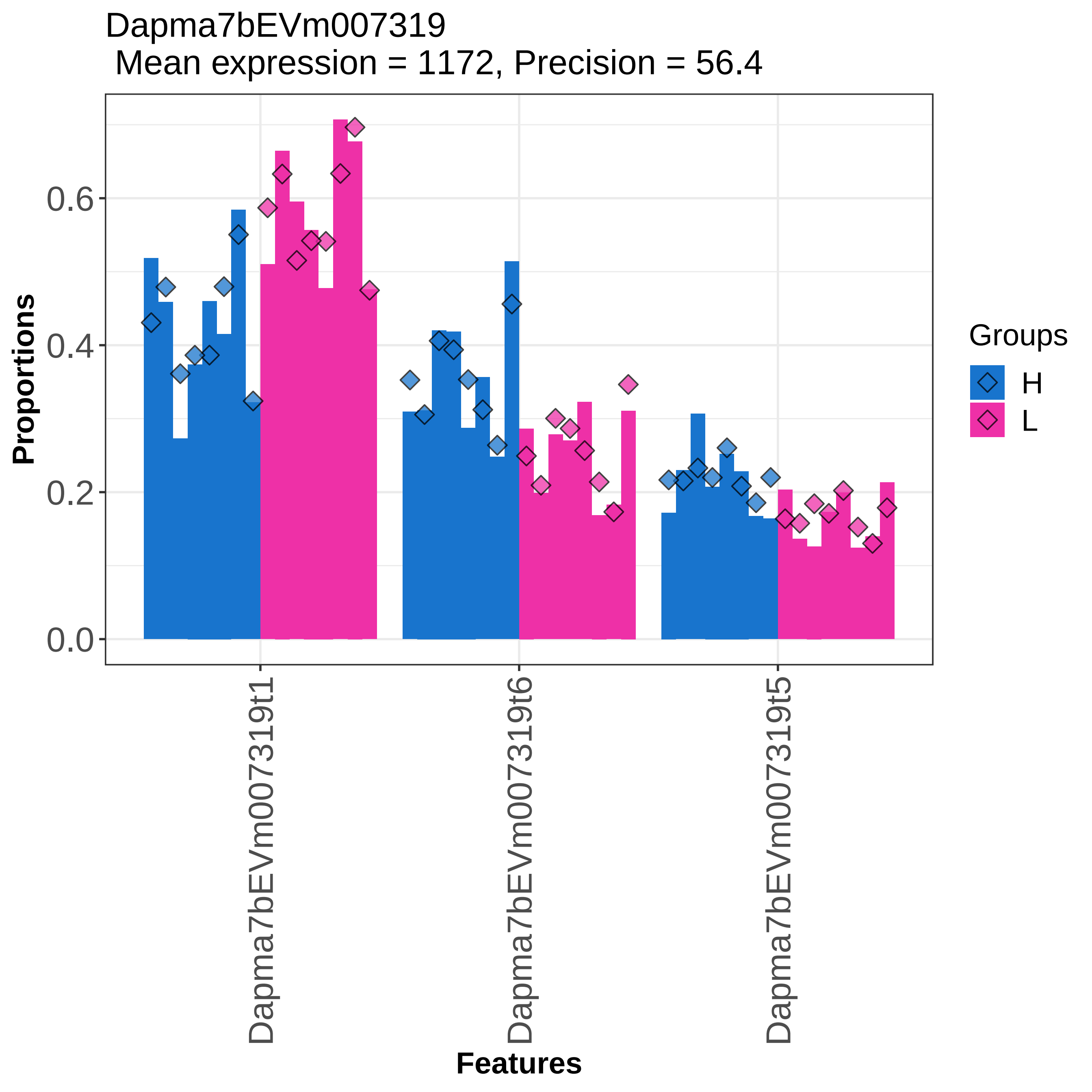

Dapma7bEVm007319
Mean expression = 1172, Precision = 56.4
Proportions
0.0
0.2
0.4
0.6
Dapma7bEVm007319t1
Dapma7bEVm007319t6
Dapma7bEVm007319t5
Features
Groups
H
L

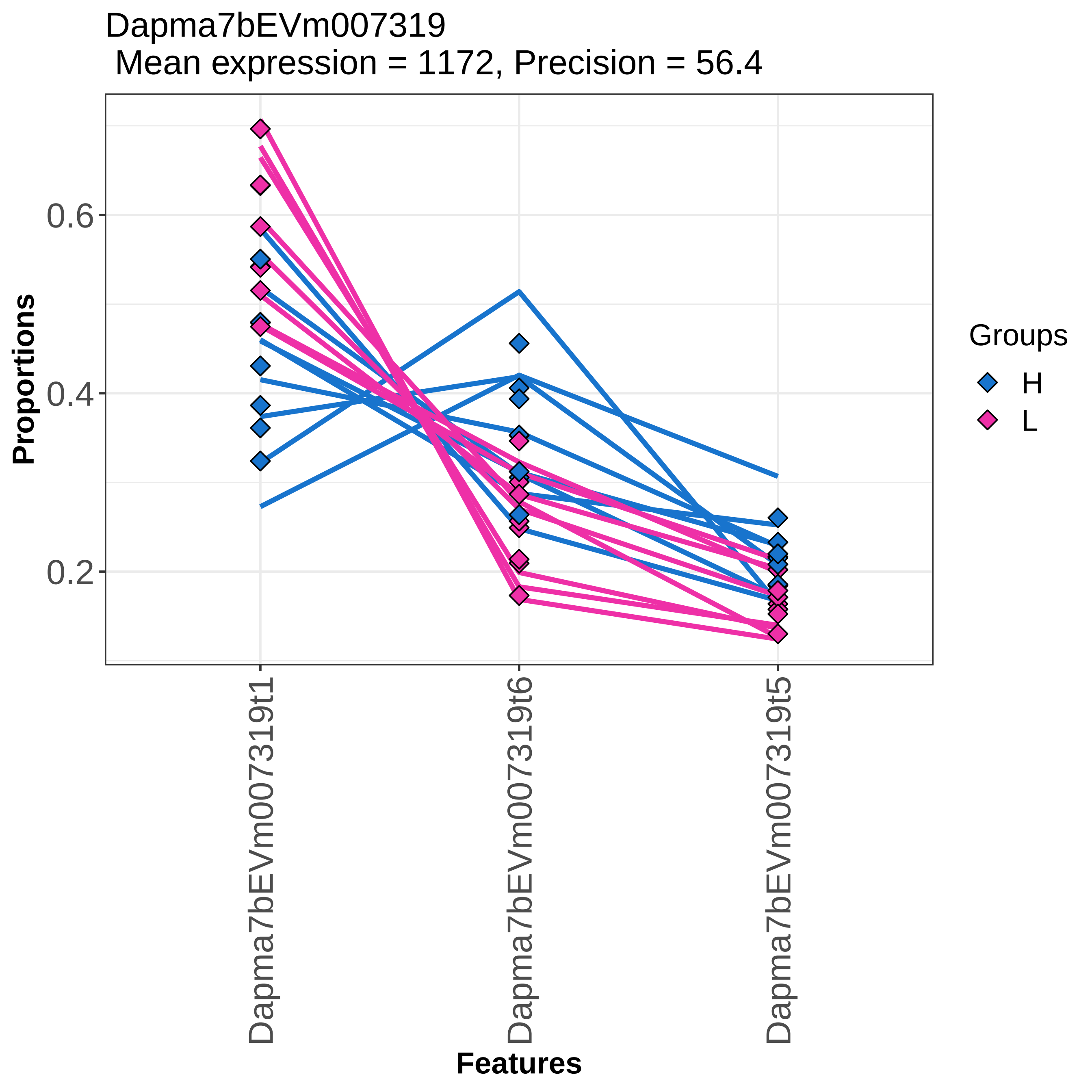

Dapma7bEVm007319
Mean expression = 1172, Precision = 56.4
Proportions
0.6
0.4
0.2
Dapma7bEVm007319t1
Dapma7bEVm007319t6
Dapma7bEVm007319t5
Features
Groups
H
L

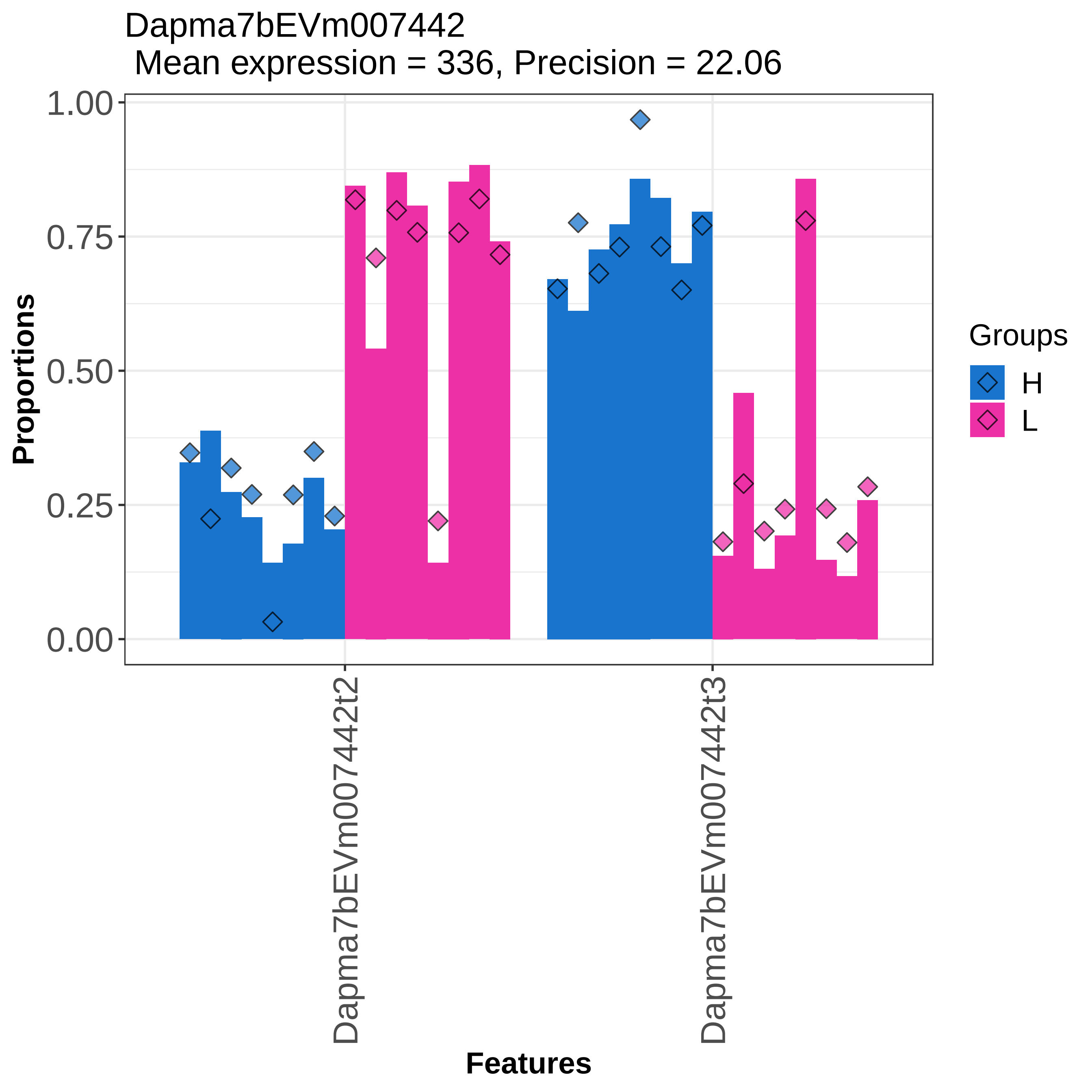

Dapma7bEVm007442
Mean expression = 336, Precision = 22.06
Proportions
1.00
0.75
0.50
0.25
0.00
Dapma7bEVm007442t2
Dapma7bEVm007442t3
Features
Groups
H
L

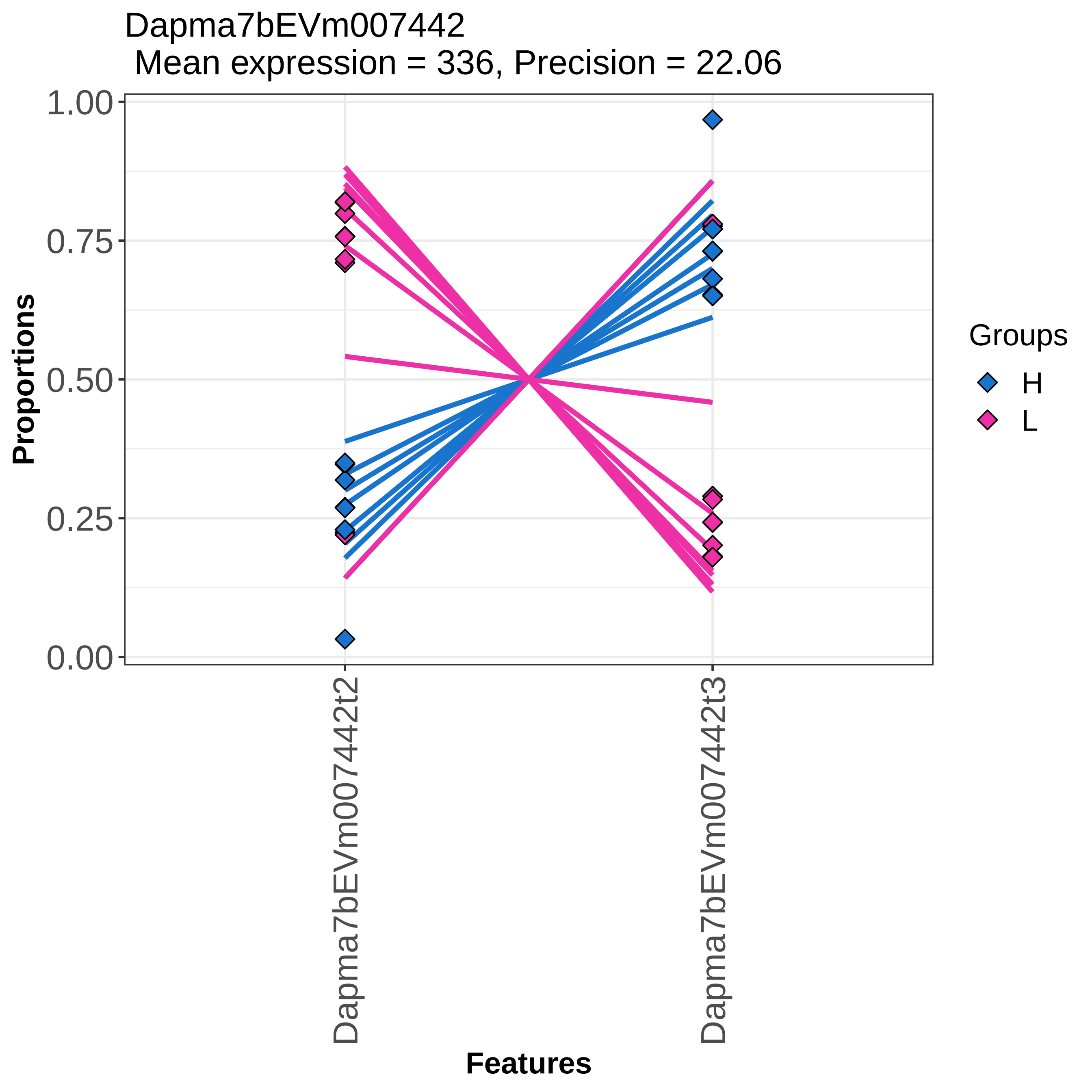

Dapma7bEVm007442
Mean expression = 336, Precision = 22.06
Proportions
1.00
0.75
0.50
0.25
0.00
Dapma7bEVm007442t2
Dapma7bEVm007442t3
Features
Groups
H
L

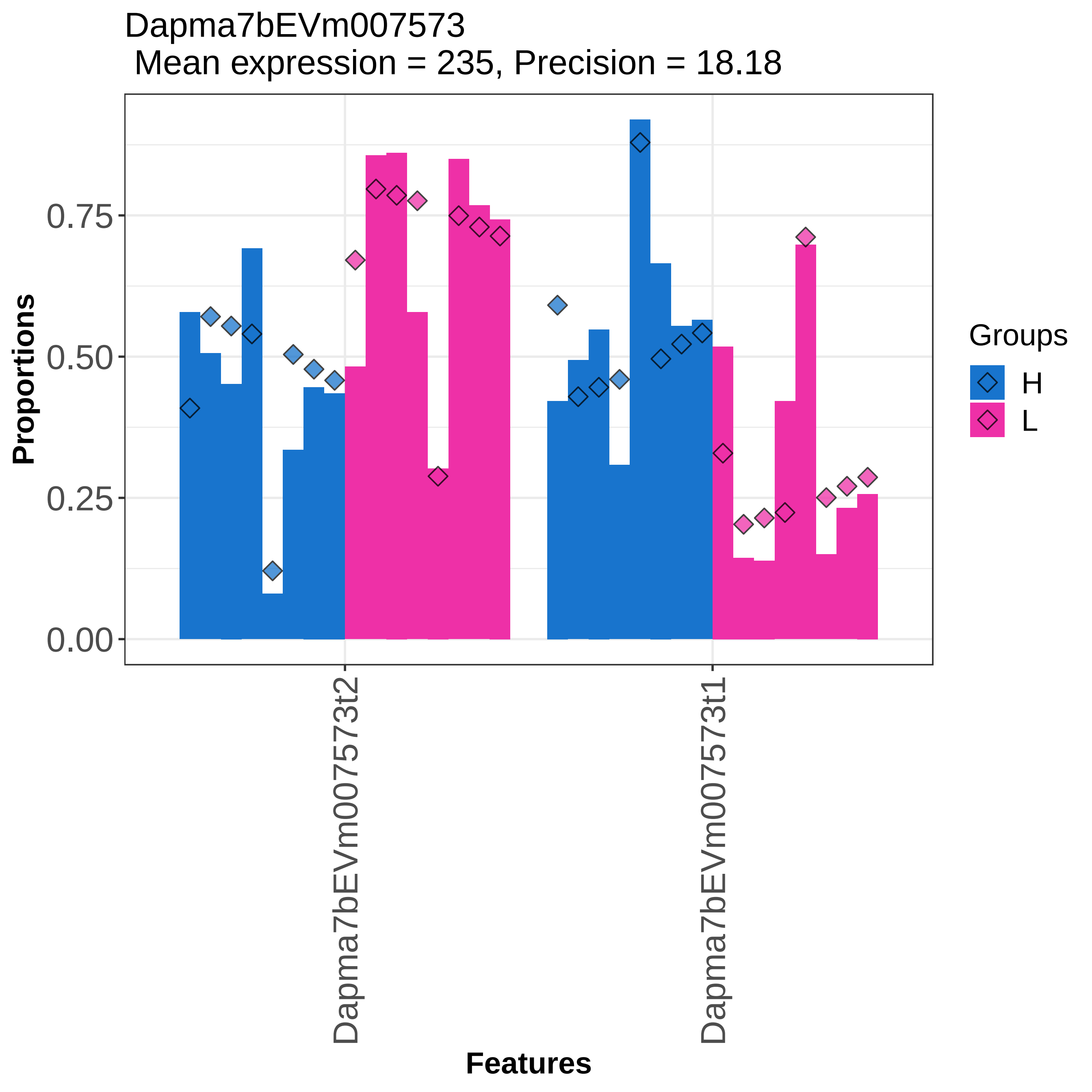

Dapma7bEVm007573
Mean expression = 235, Precision = 18.18
Proportions
0.75
0.50
0.25
0.00
Dapma7bEVm007573t2
Dapma7bEVm007573t1
Features
Groups
H
L

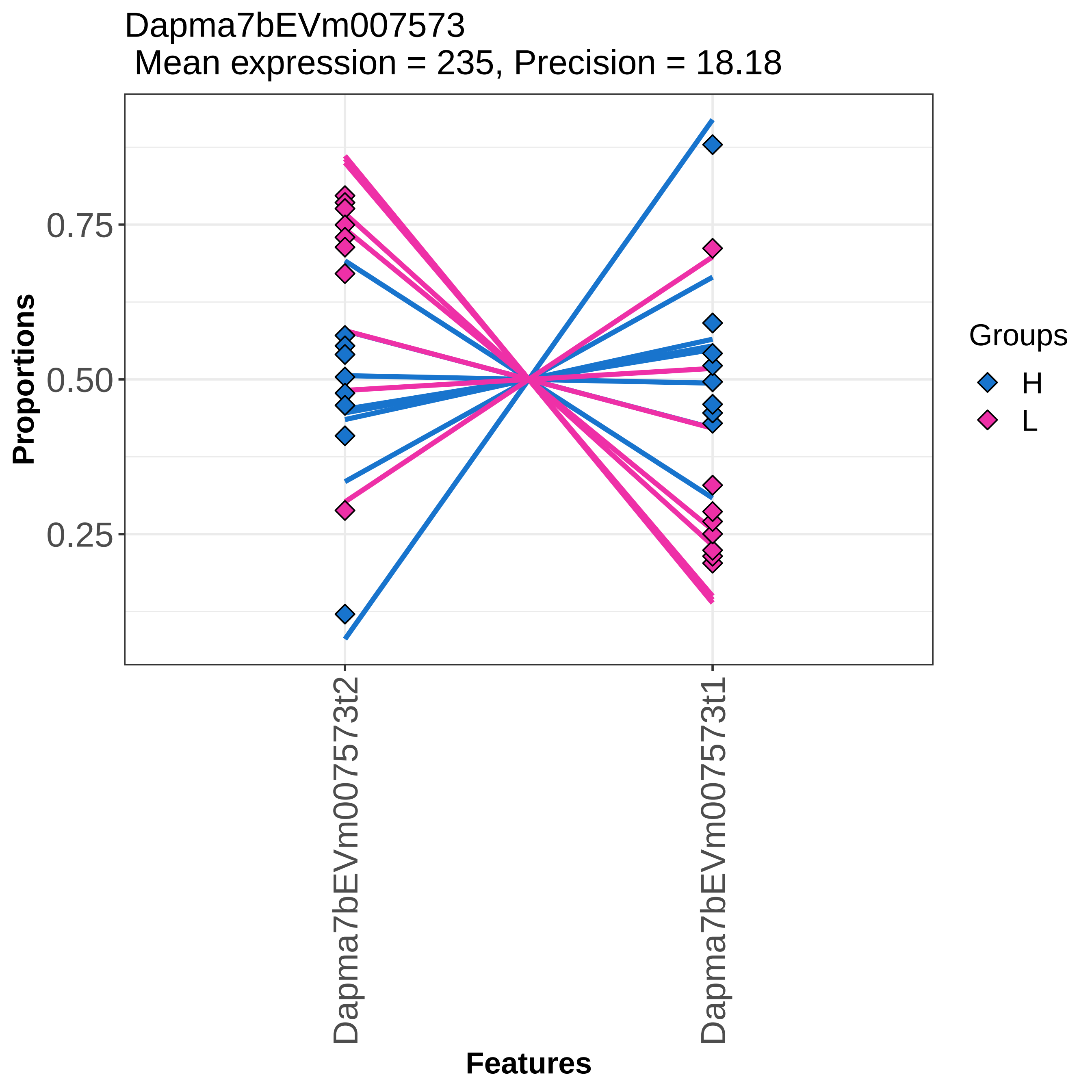

Dapma7bEVm007573
Mean expression = 235, Precision = 18.18
Proportions
0.75
0.50
0.25
Features
Dapma7bEVm007573t2
Dapma7bEVm007573t1
Groups
H
L

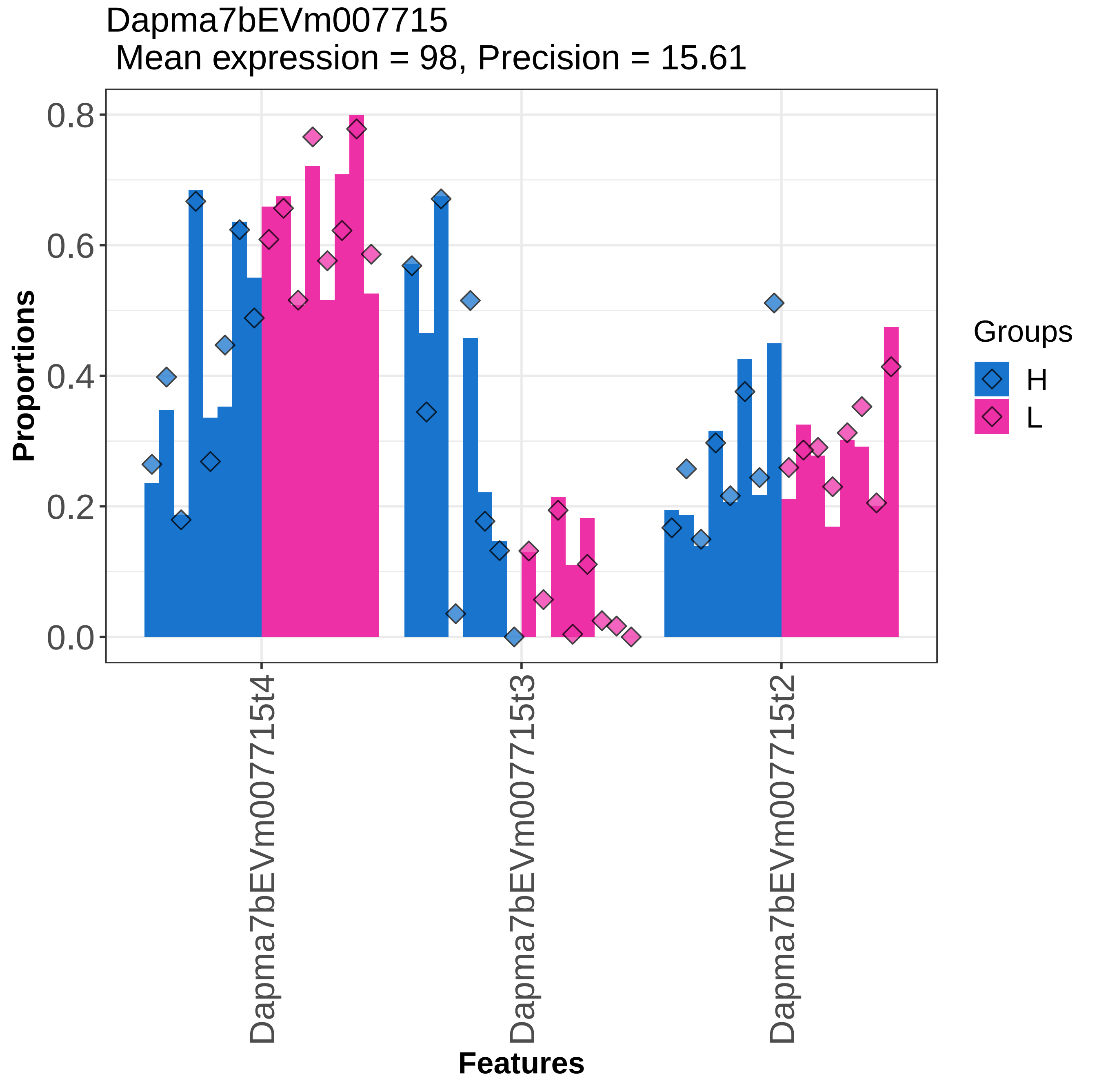

Dapma7bEVm007715
Mean expression = 98, Precision = 15.61
Proportions
0.8
0.6
0.4
0.2
0.0
Dapma7bEVm007715t4
Dapma7bEVm007715t3
Dapma7bEVm007715t2
Features
Groups
H
L

Dapma7bEVm007715
Mean expression = 98, Precision = 15.61

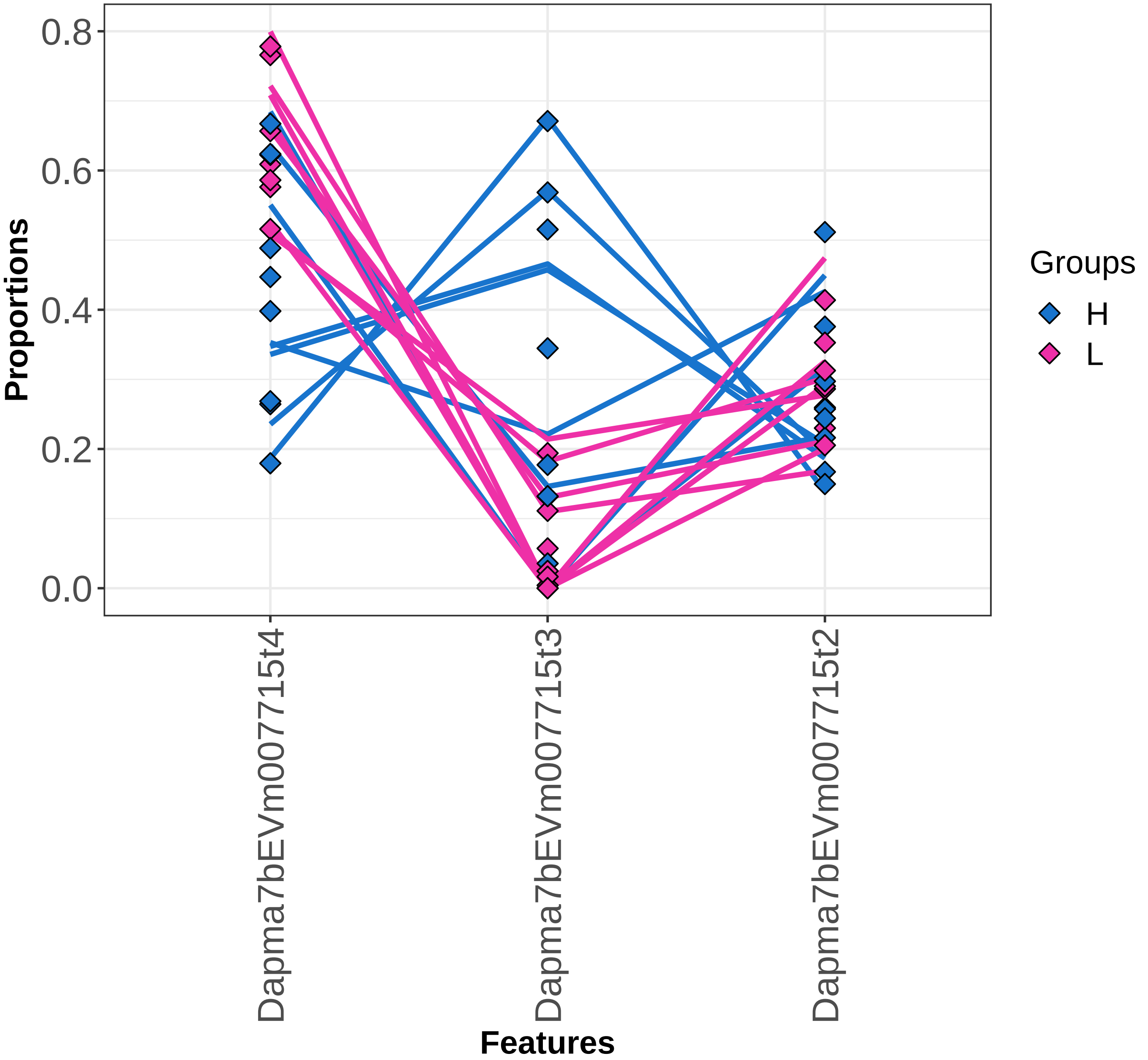

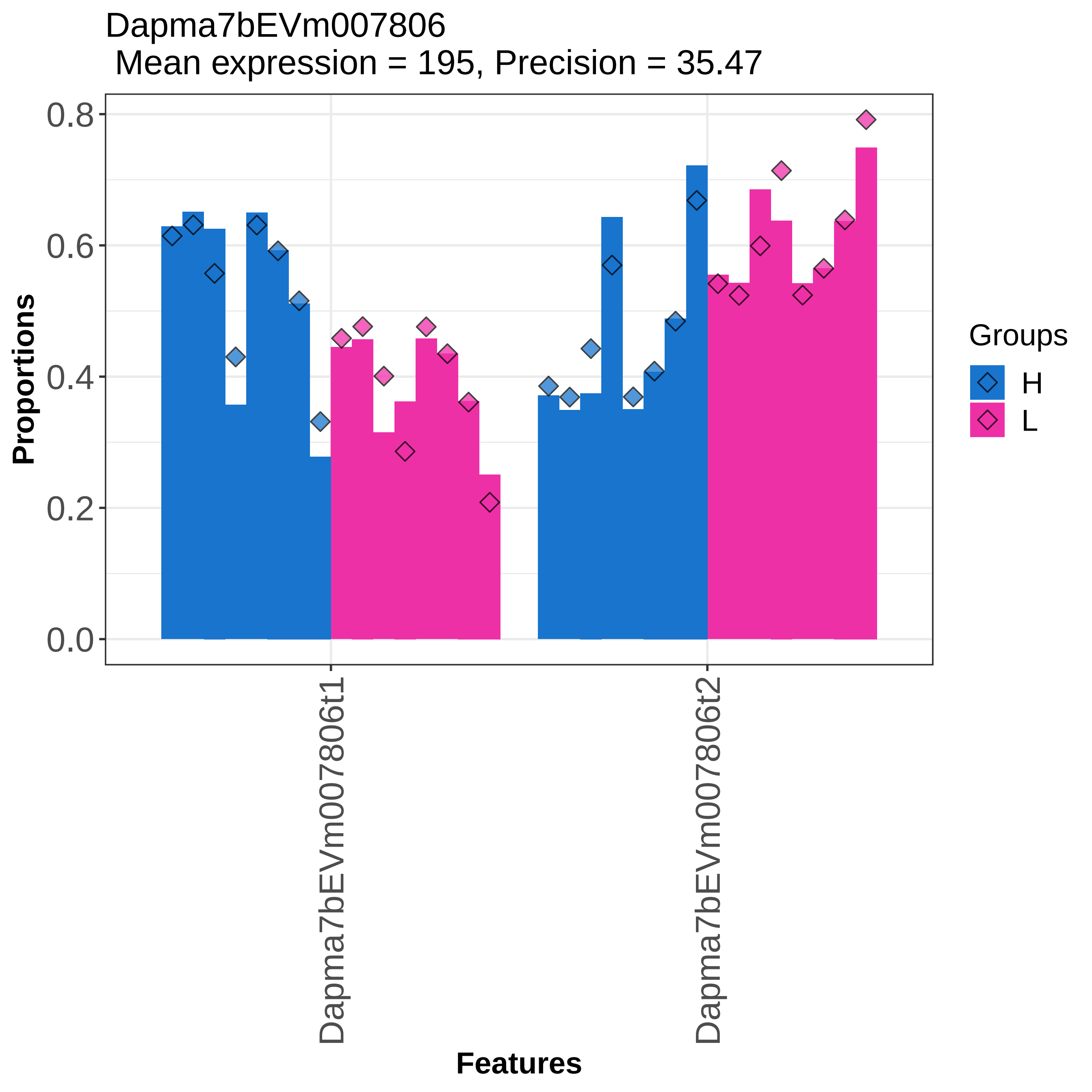
Dapma7bEVm007806
Mean expression = 195, Precision = 35.47
0.8
0.6
0.4
0.2
0.0
Proportions
Dapma7bEVm007806t1
Dapma7bEVm007806t2
Features
Groups
H
L

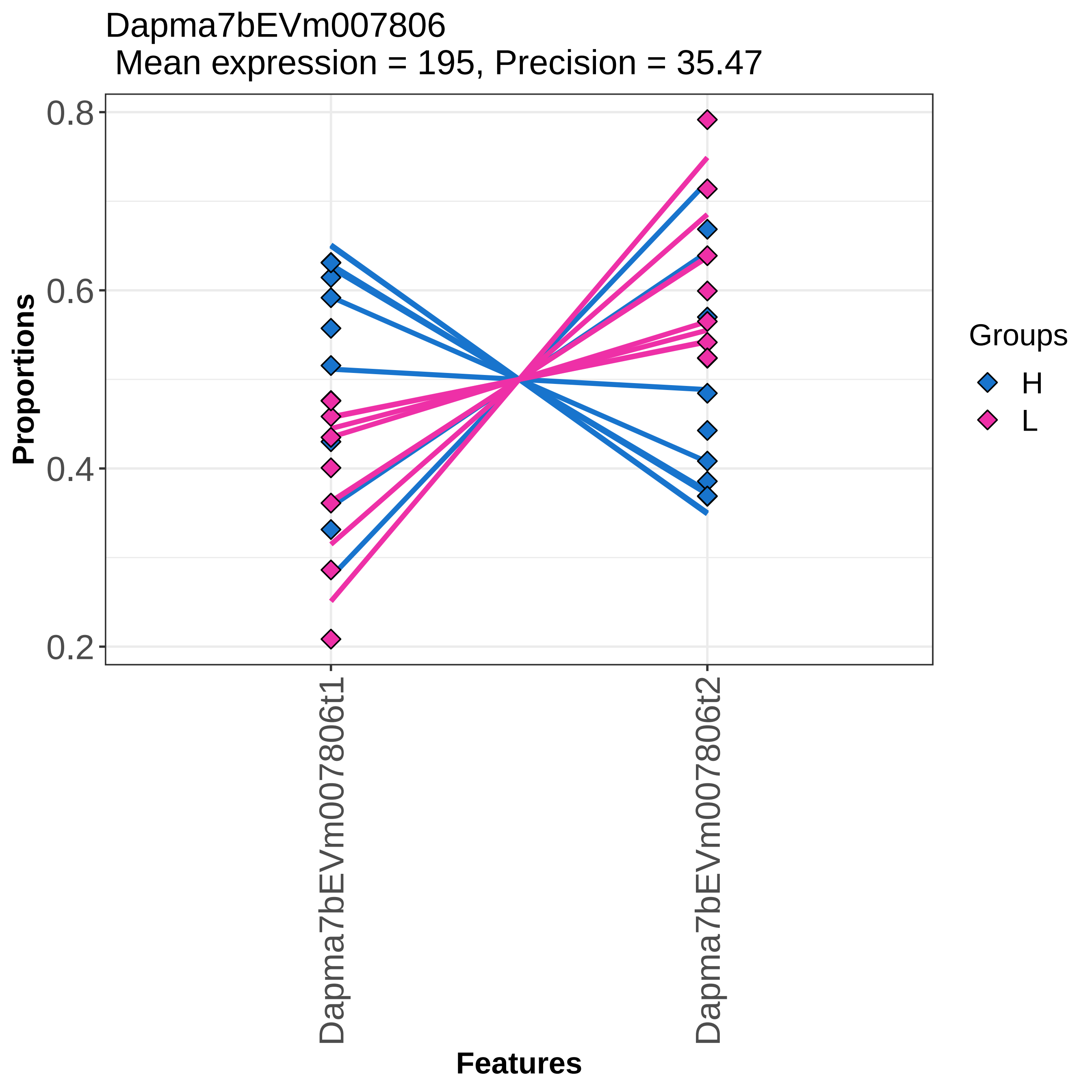
Dapma7bEVm007806
Mean expression = 195, Precision = 35.47
Proportions
0.8
0.6
0.4
0.2
Dapma7bEVm007806t1
Dapma7bEVm007806t2
Features
Groups
H
L

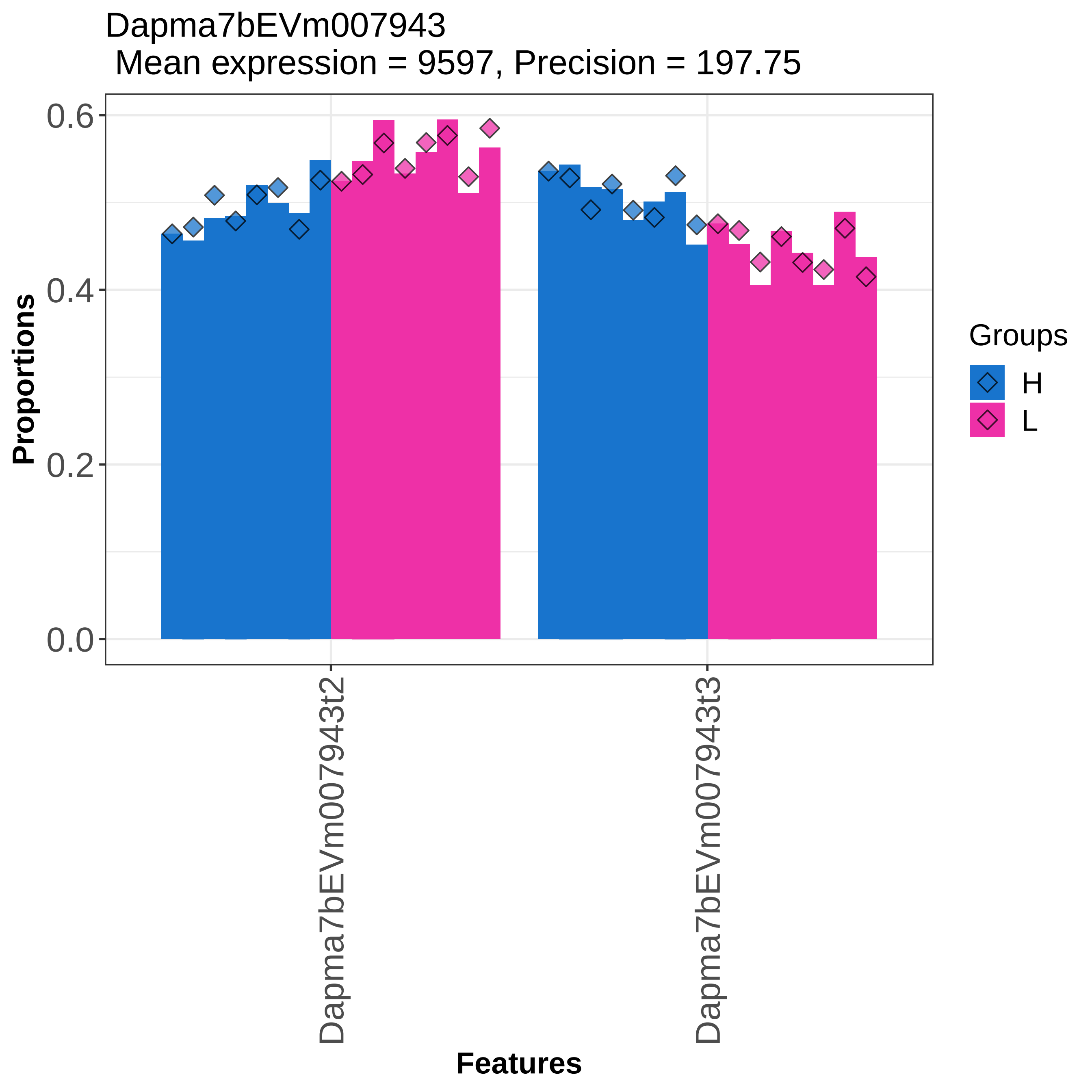
Dapma7bEVm007943
Mean expression = 9597, Precision = 197.75
0.6
0.4
0.2
0.0
Proportions
Dapma7bEVm007943t2
Dapma7bEVm007943t3
Features
Groups
H
L

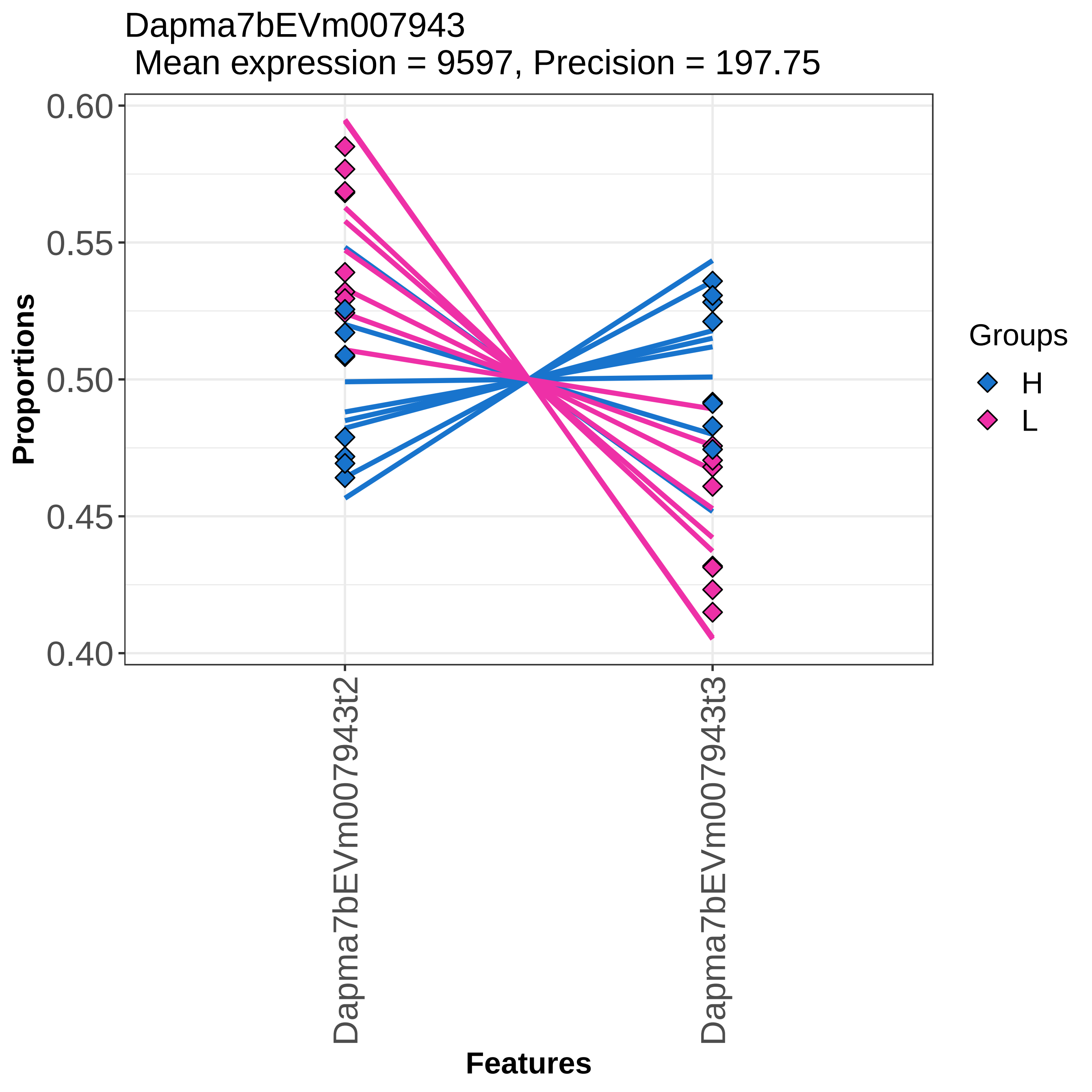

Dapma7bEVm007943
Mean expression = 9597, Precision = 197.75
0.60
0.55
0.50
0.45
0.40
Proportions
Dapma7bEVm007943t2
Dapma7bEVm007943t3
Features
Groups
H
L

Dapma7bEVm008122
Mean expression = 861, Precision = 34.28

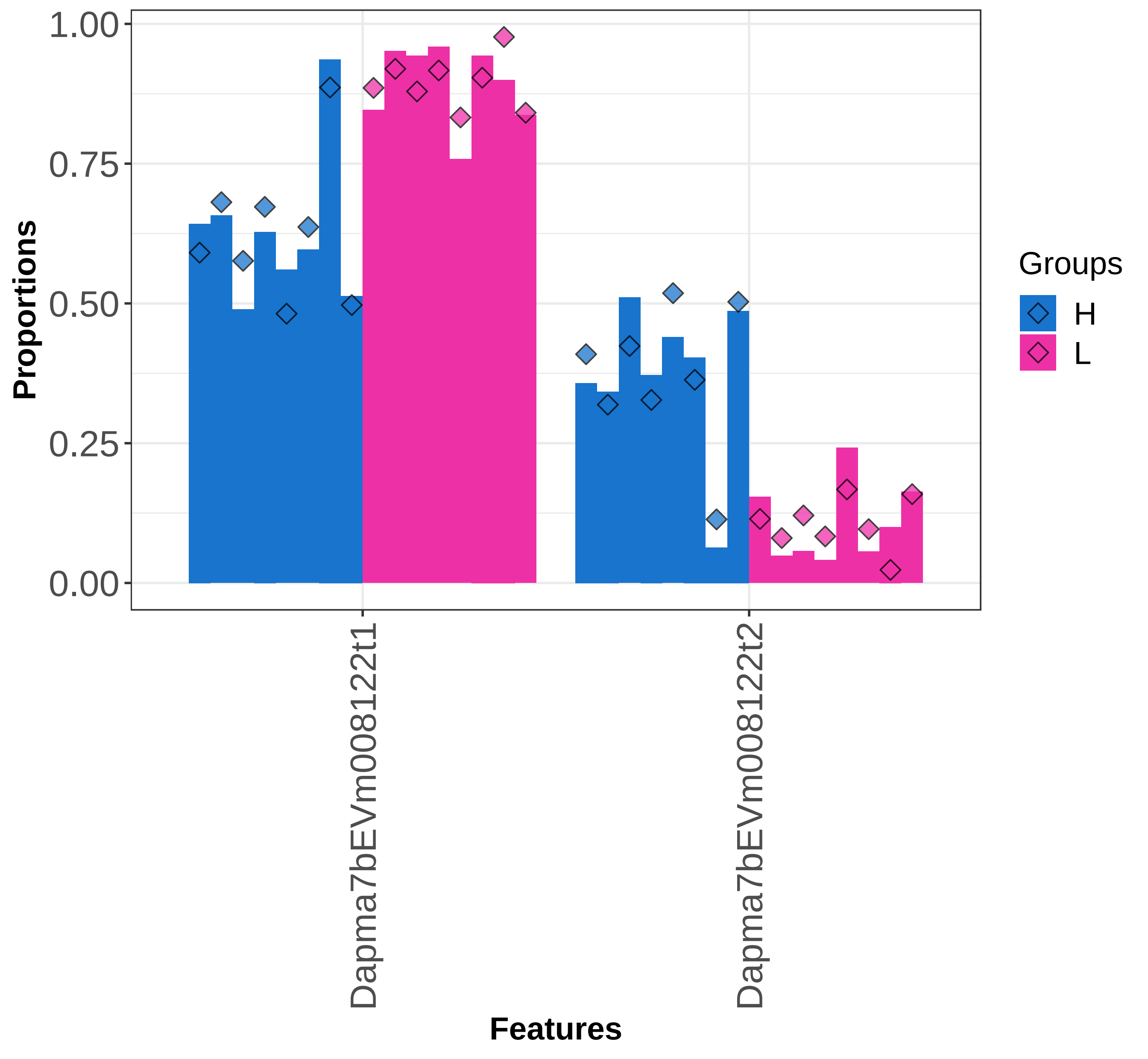

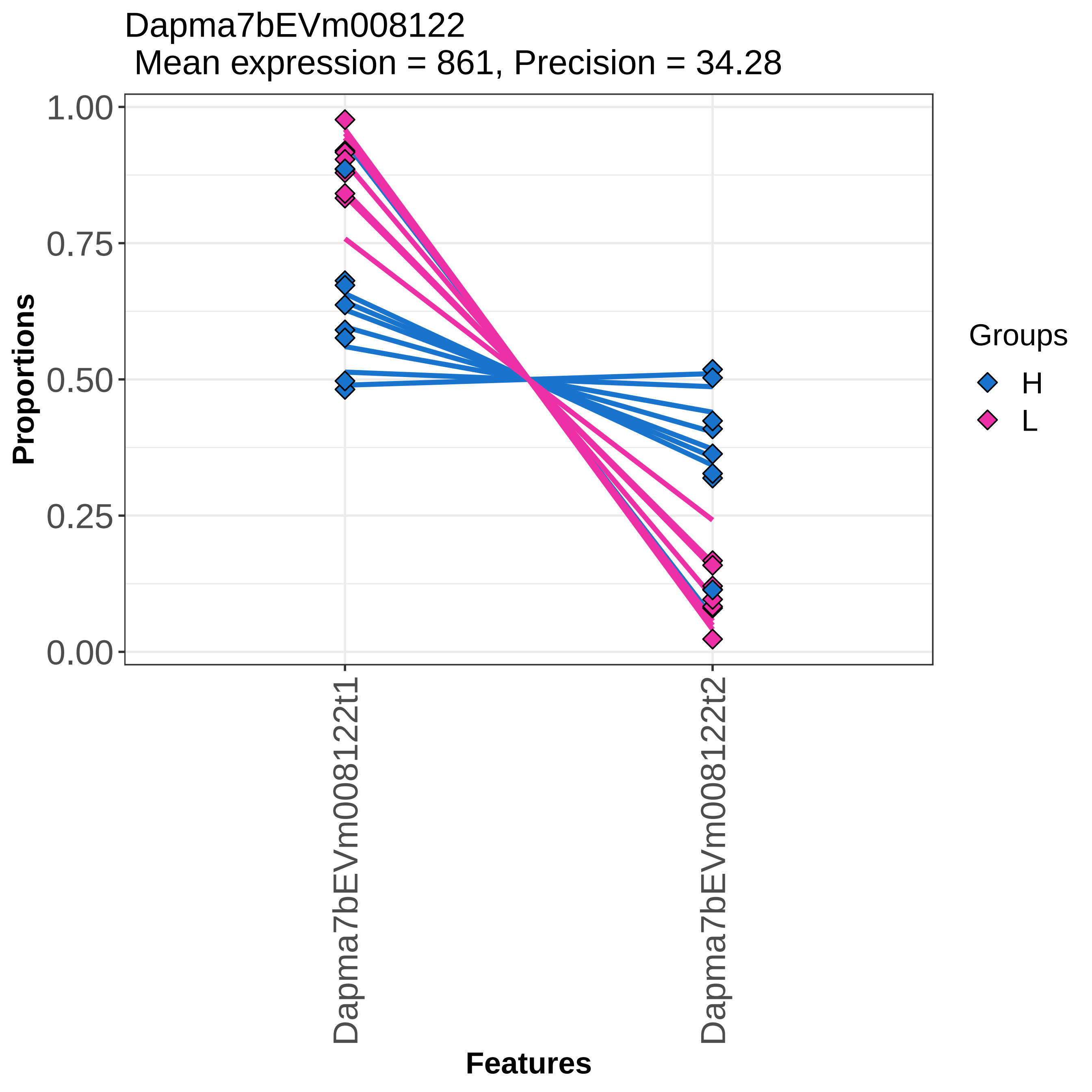
Dapma7bEVm008122
Mean expression = 861, Precision = 34.28
Proportions
1.00
0.75
0.50
0.25
0.00
Dapma7bEVm008122t1
Dapma7bEVm008122t2
Features
Groups
H
L

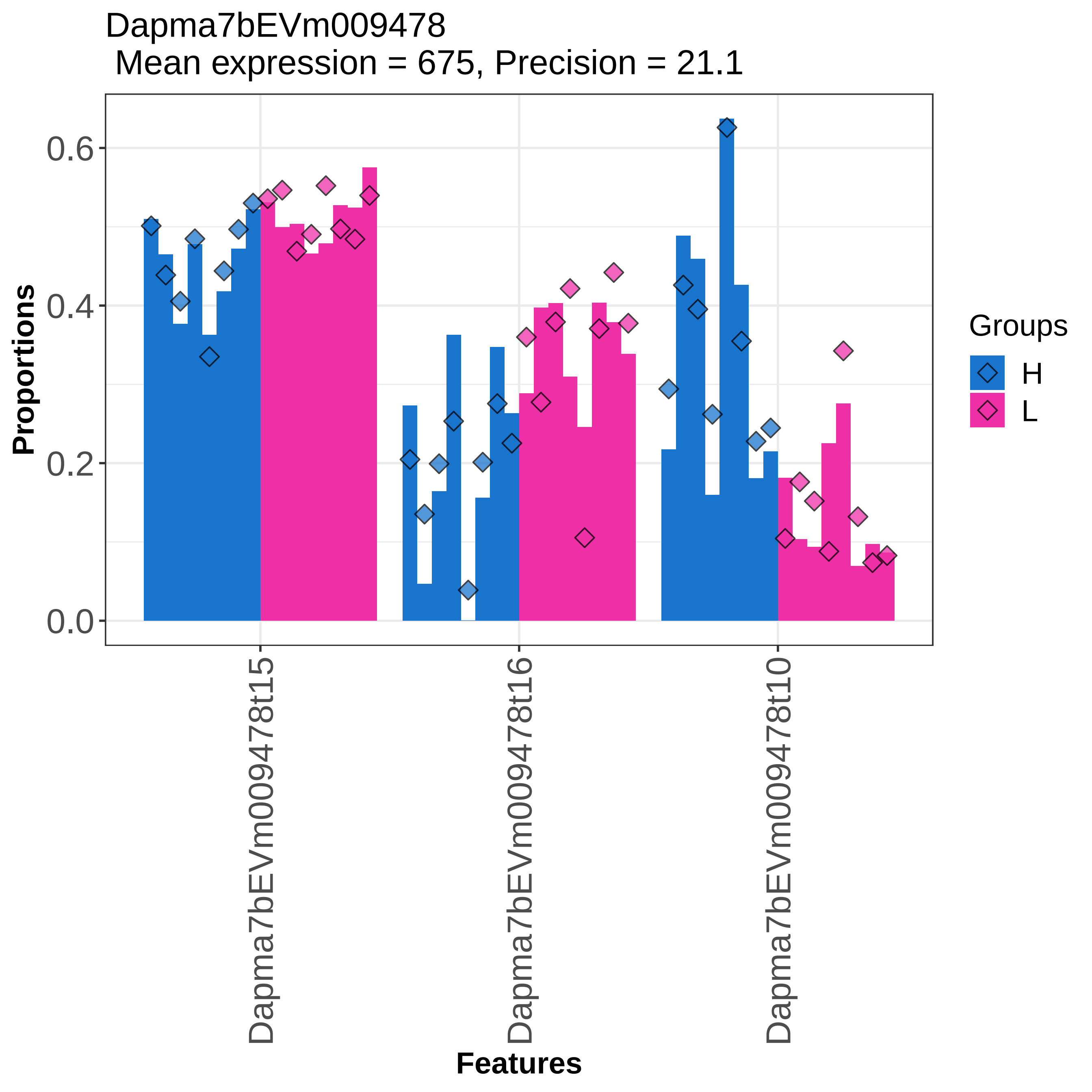
Dapma7bEVm009478
Mean expression = 675, Precision = 21.1
Proportions
0.0
0.2
0.4
0.6
Dapma7bEVm009478t15
Dapma7bEVm009478t16
Dapma7bEVm009478t10
Features
Groups
H
L

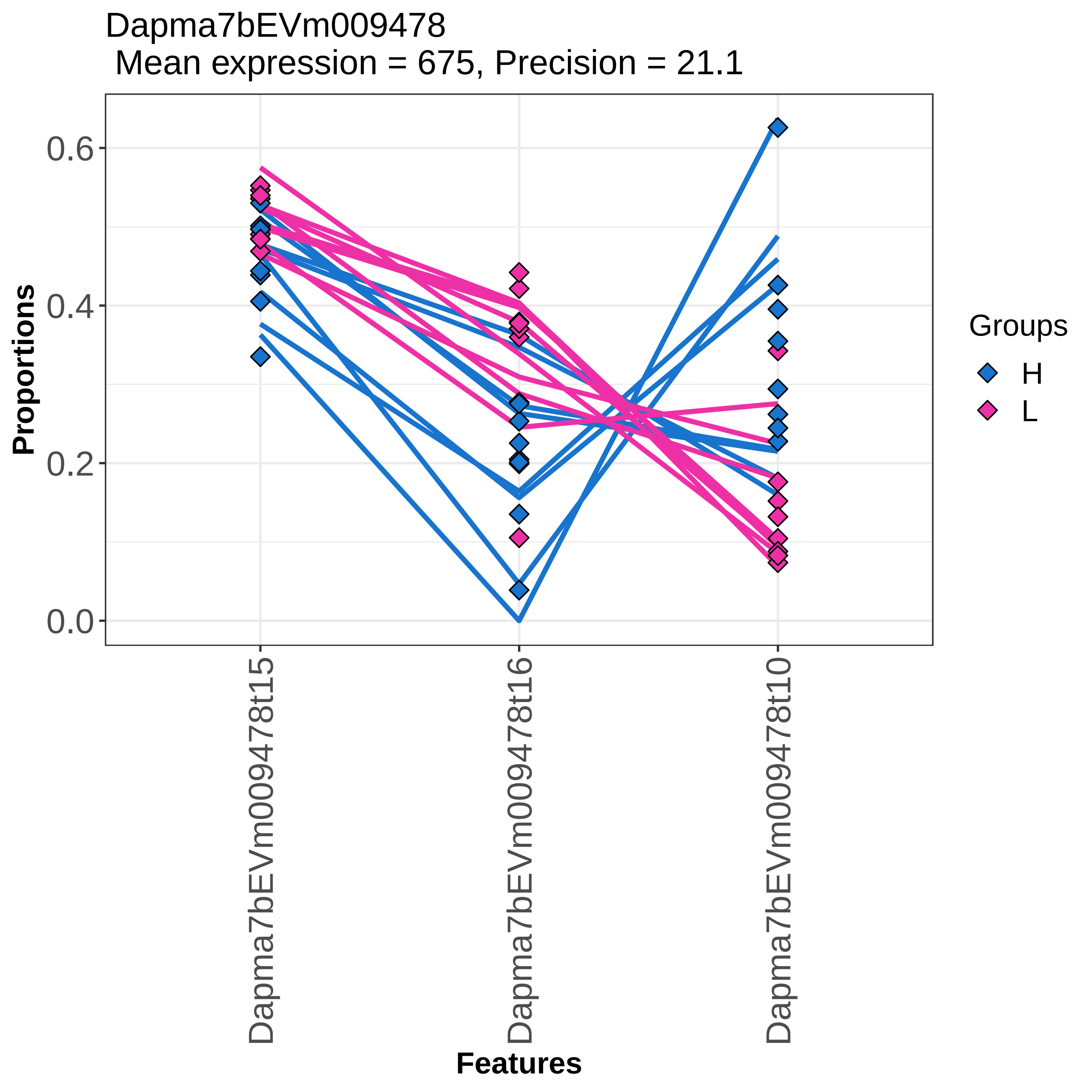

Dapma7bEVm009478
Mean expression = 675, Precision = 21.1
Proportions
0.0
0.2
0.4
0.6
Groups
H
L
Dapma7bEVm009478t15
Dapma7bEVm009478t16
Dapma7bEVm009478t10
Features

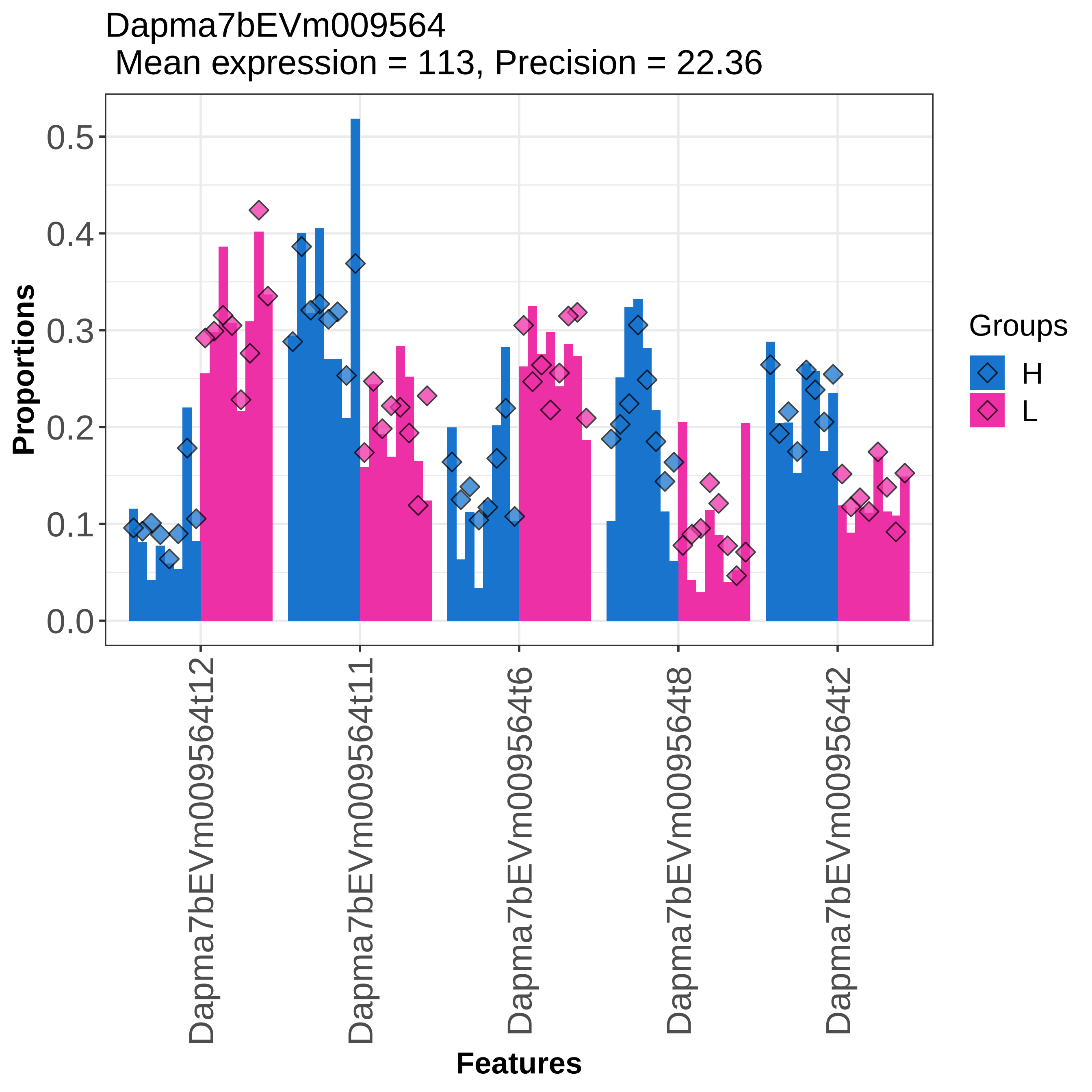
Dapma7bEVm009564
Mean expression = 113, Precision = 22.36
Proportions
0.0
0.1
0.2
0.3
0.4
0.5
Dapma7bEVm009564t12
Dapma7bEVm009564t11
Dapma7bEVm009564t6
Dapma7bEVm009564t8
Dapma7bEVm009564t2
Features
Groups
H
L

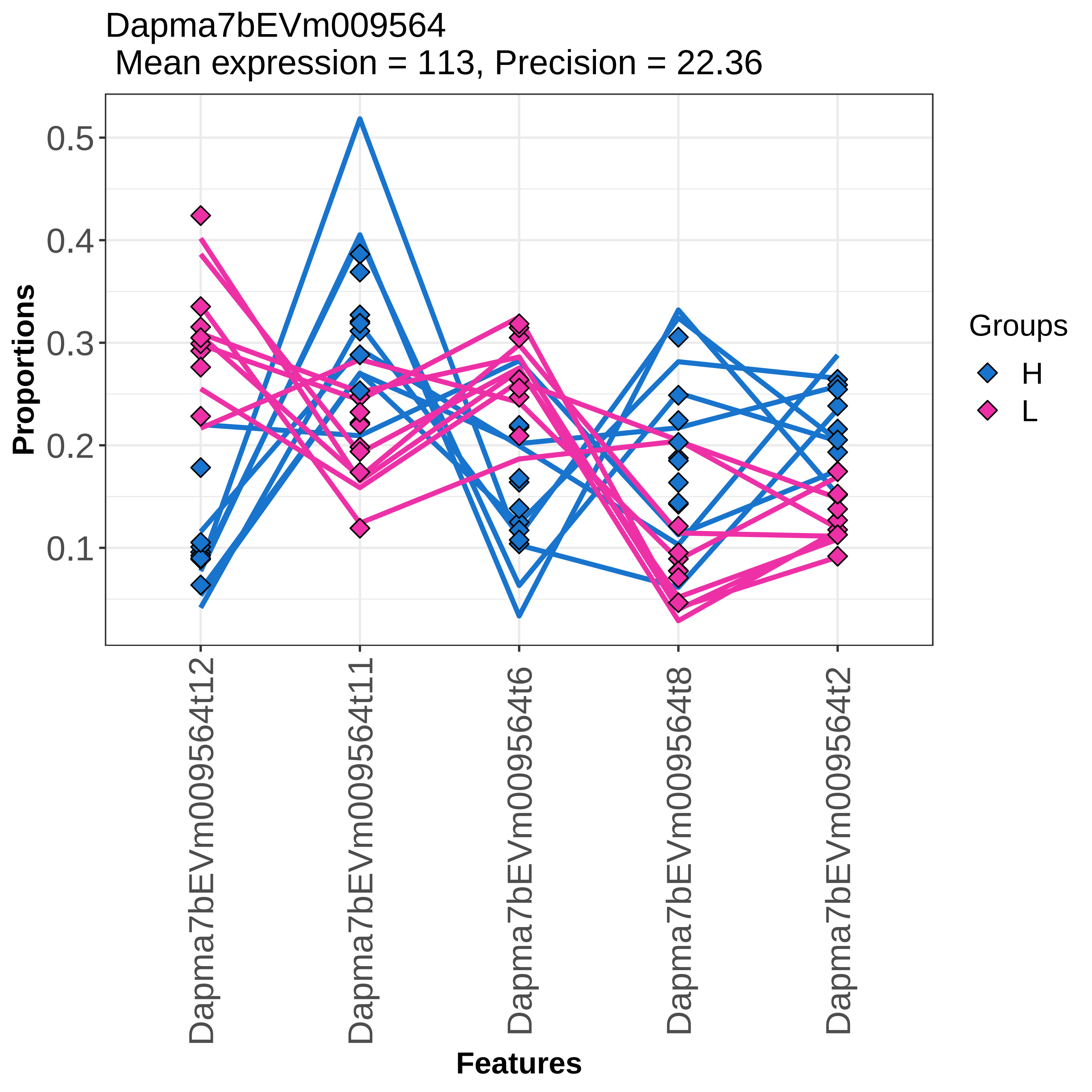

Dapma7bEVm009564
Mean expression = 113, Precision = 22.36
Proportions
0.5
0.4
0.3
0.2
0.1
Features
Dapma7bEVm009564t12
Dapma7bEVm009564t11
Dapma7bEVm009564t6
Dapma7bEVm009564t8
Dapma7bEVm009564t2
Groups
H
L

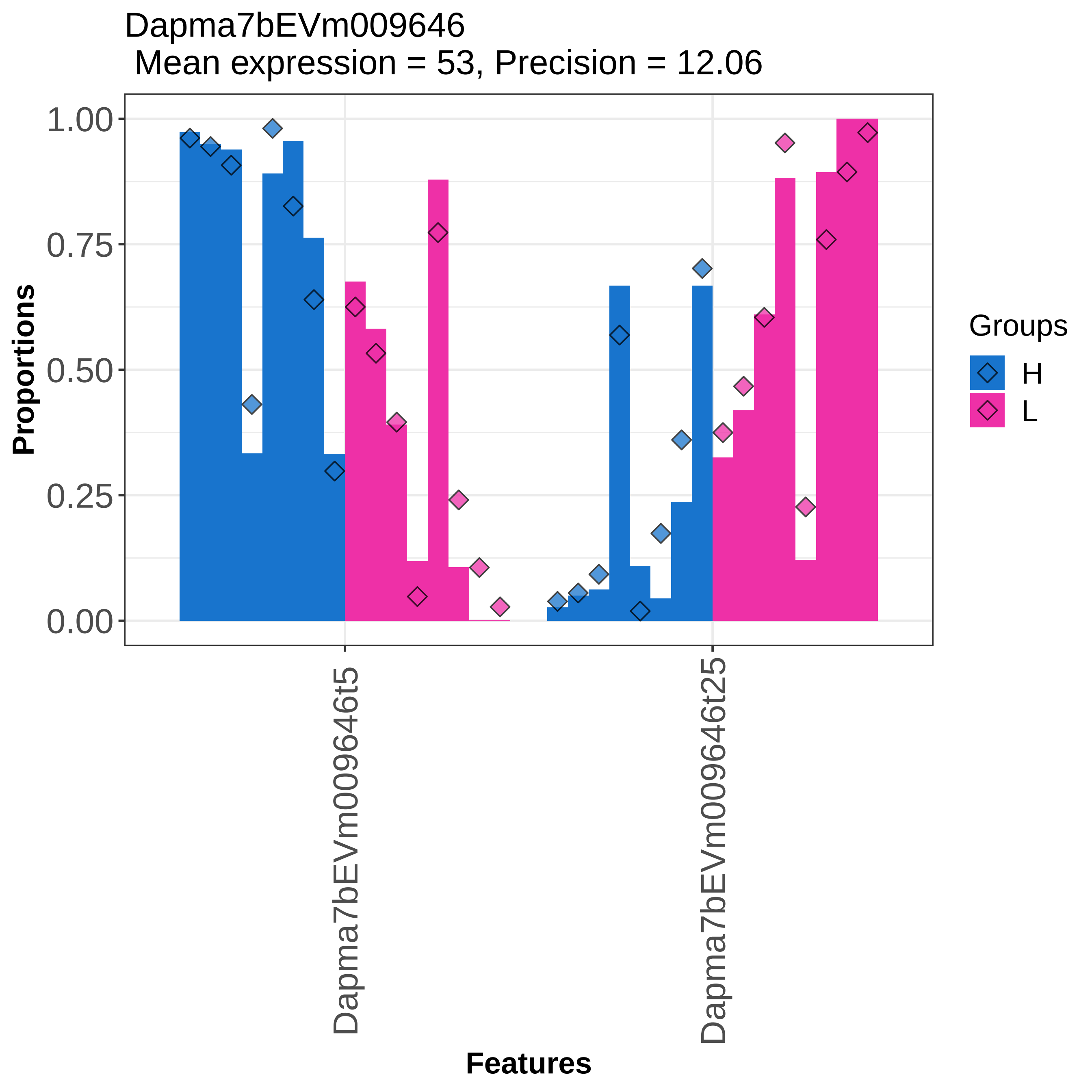

Dapma7bEVm009646
Mean expression = 53, Precision = 12.06
Proportions
1.00
0.75
0.50
0.25
0.00
Dapma7bEVm009646t5
Dapma7bEVm009646t25
Features
Groups
H
L

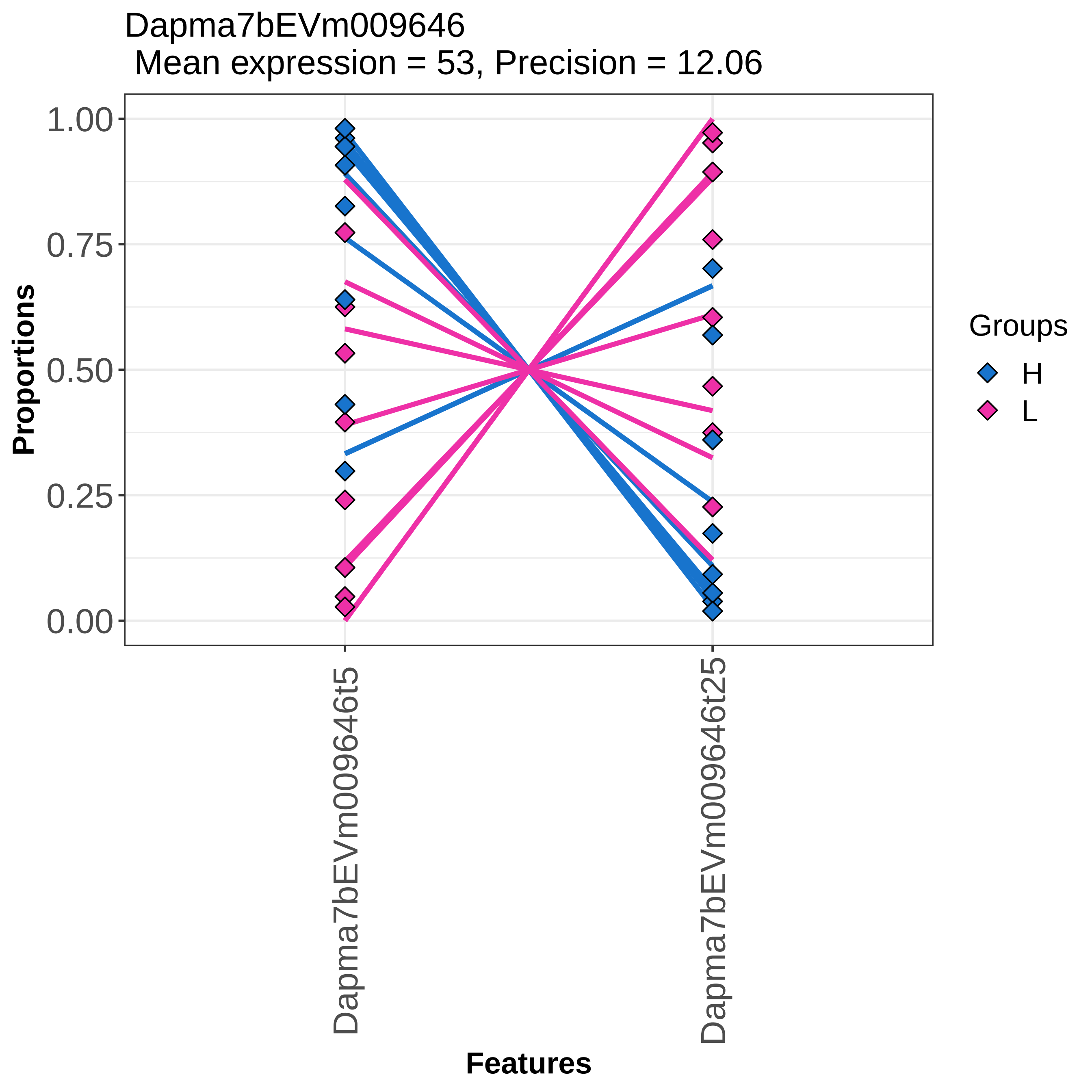

Dapma7bEVm009646
Mean expression = 53, Precision = 12.06
Proportions
1.00
0.75
0.50
0.25
0.00
Dapma7bEVm009646t5
Dapma7bEVm009646t25
Features
Groups
H
L

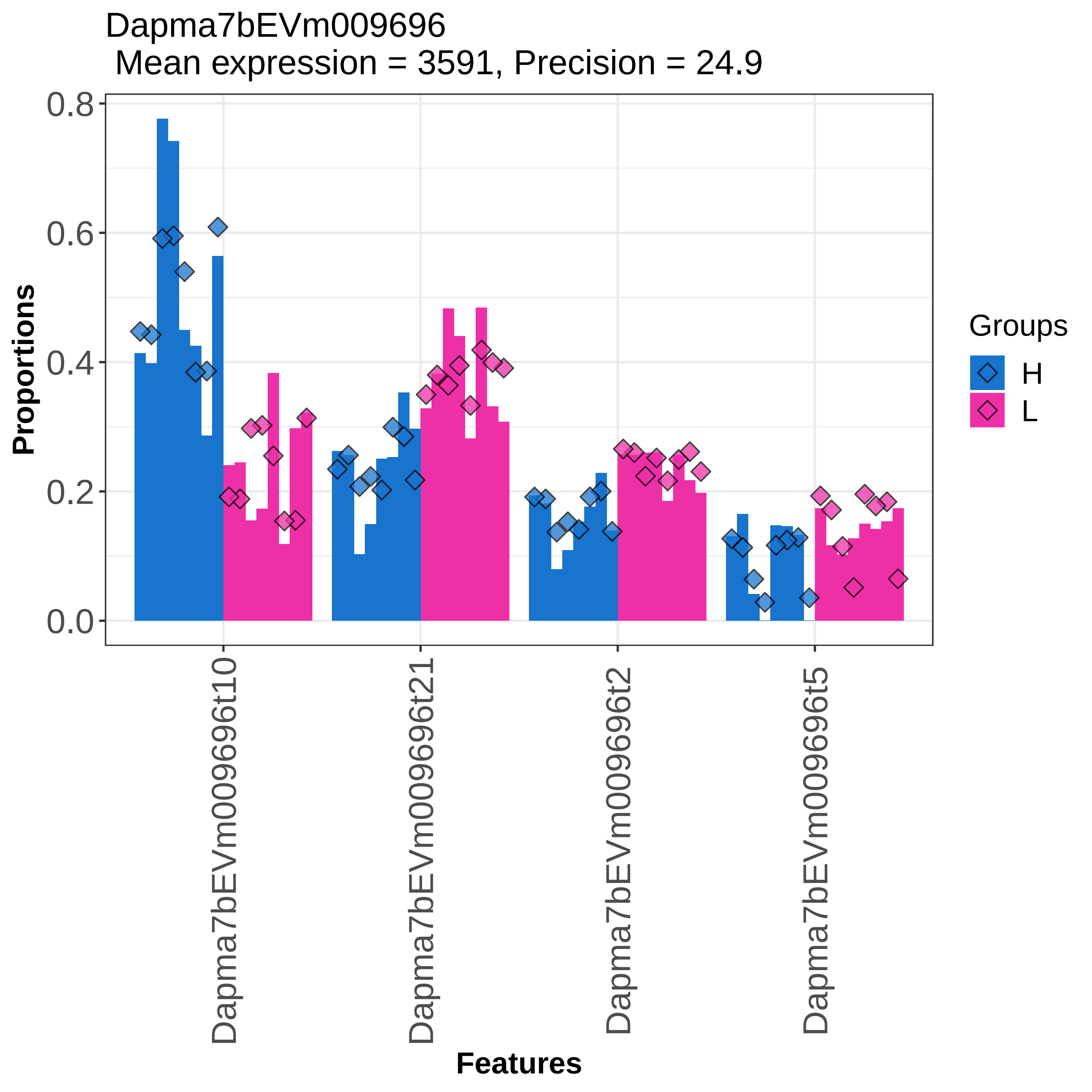

Dapma7bEVm009696
Mean expression = 3591, Precision = 24.9
Proportions
0.8
0.6
0.4
0.2
0.0
Features
Dapma7bEVm009696t10
Dapma7bEVm009696t21
Dapma7bEVm009696t2
Dapma7bEVm009696t5
Groups
H
L

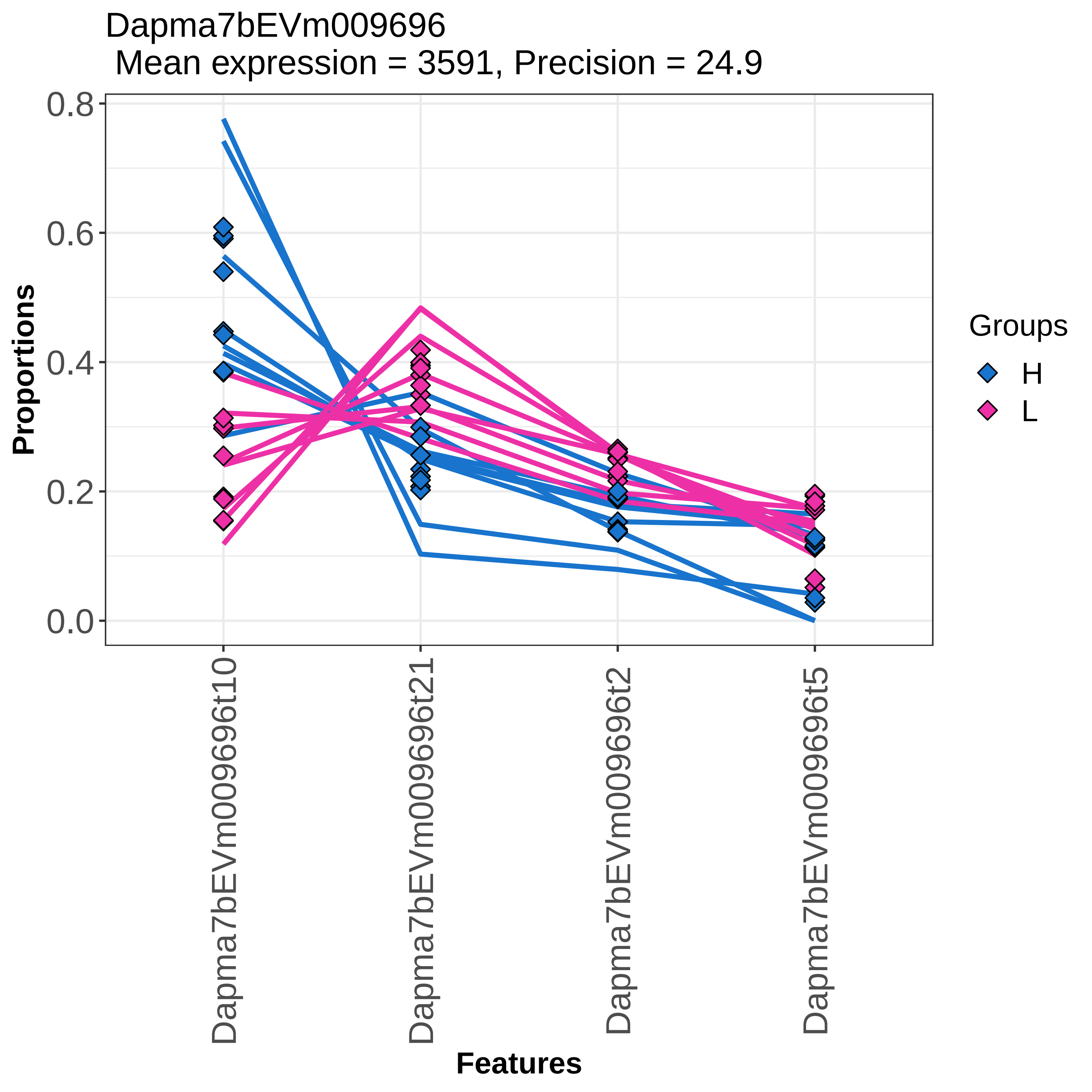
Dapma7bEVm009696
Mean expression = 3591, Precision = 24.9
Proportions
0.0
0.2
0.4
0.6
0.8
Dapma7bEVm009696t10
Dapma7bEVm009696t21
Dapma7bEVm009696t2
Dapma7bEVm009696t5
Features
Groups
H
L

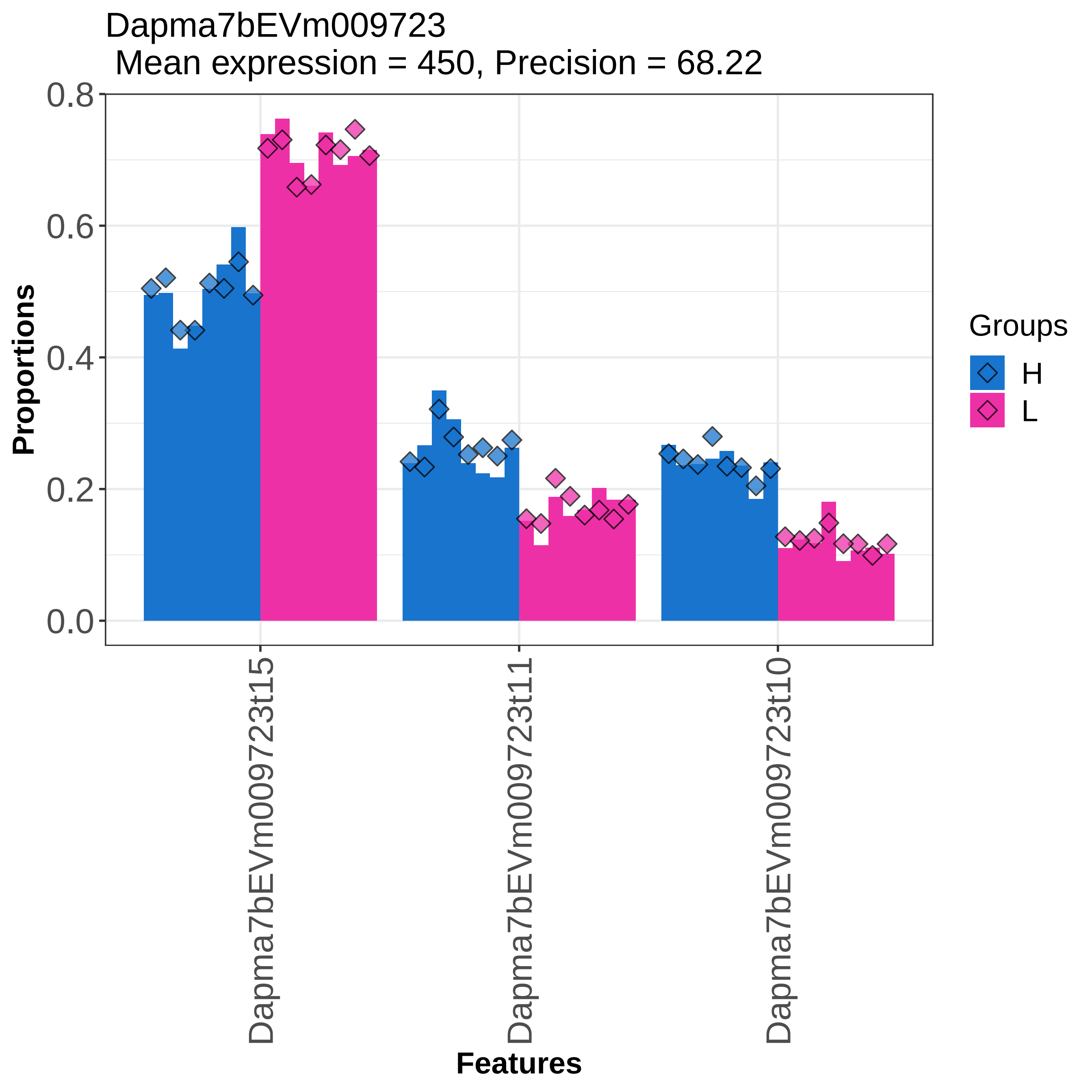
Dapma7bEVm009723
Mean expression = 450, Precision = 68.22
Proportions
0.0
0.2
0.4
0.6
0.8
Dapma7bEVm009723t15
Dapma7bEVm009723t11
Dapma7bEVm009723t10
Features
Groups
H
L

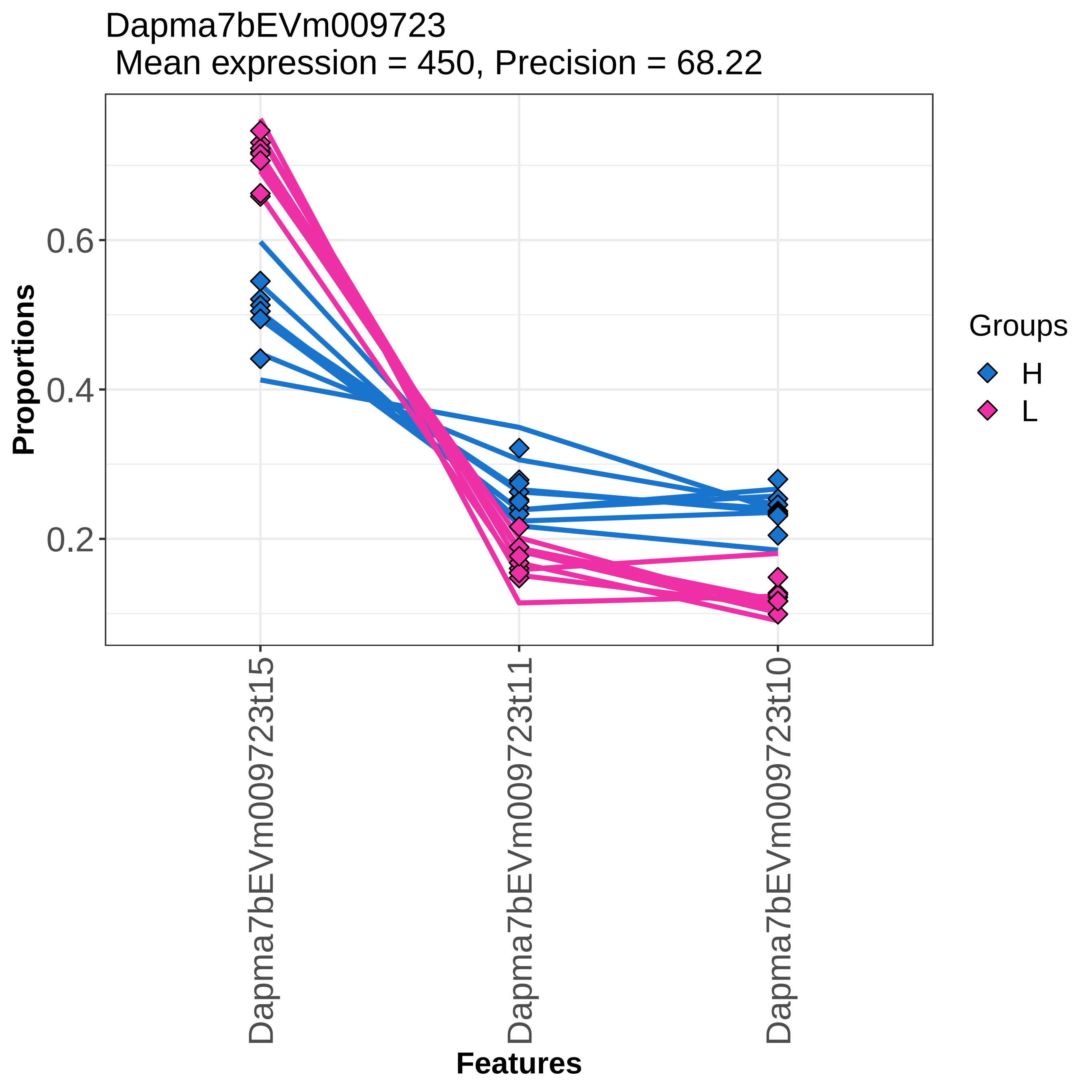

Dapma7bEVm009723
Mean expression = 450, Precision = 68.22
Proportions
0.6
0.4
0.2
Dapma7bEVm009723t15
Dapma7bEVm009723t11
Dapma7bEVm009723t10
Features
Groups
H
L

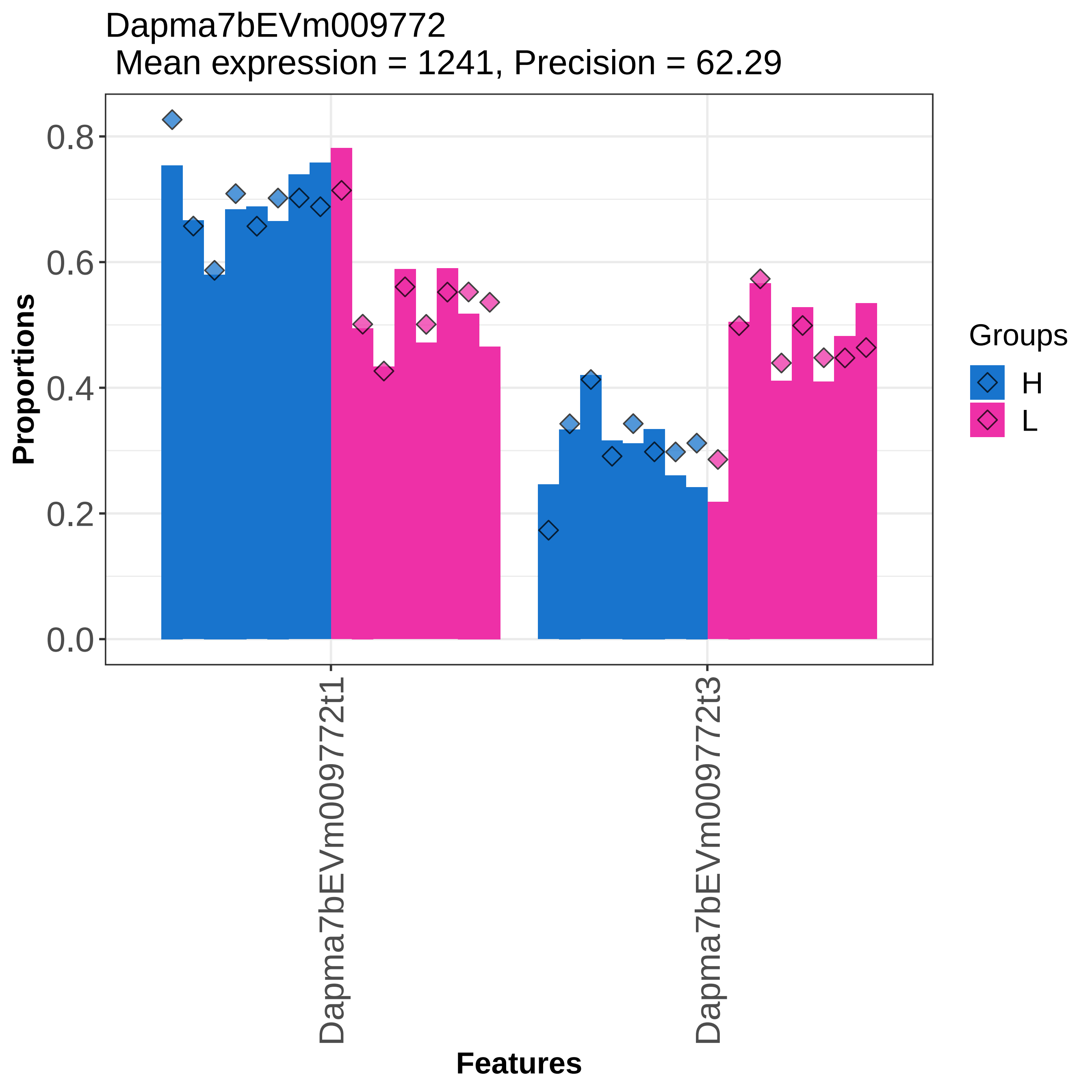

Dapma7bEVm009772
Mean expression = 1241, Precision = 62.29
Proportions
0.0
0.2
0.4
0.6
0.8
Dapma7bEVm009772t1
Dapma7bEVm009772t3
Features
Groups
H
L

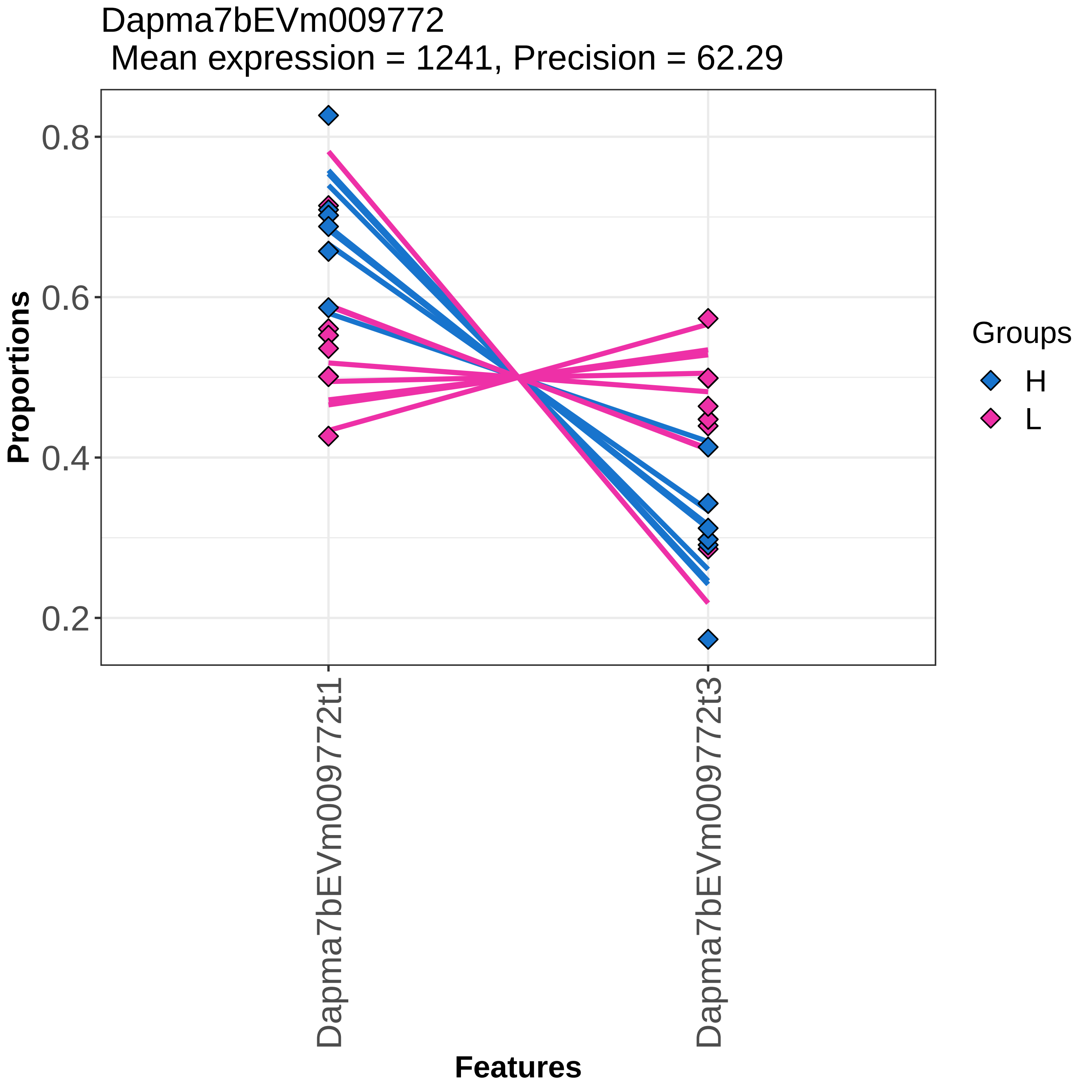
Dapma7bEVm009772
Mean expression = 1241, Precision = 62.29
Proportions
0.8
0.6
0.4
0.2
Dapma7bEVm009772t1
Dapma7bEVm009772t3
Features
Groups
H
L

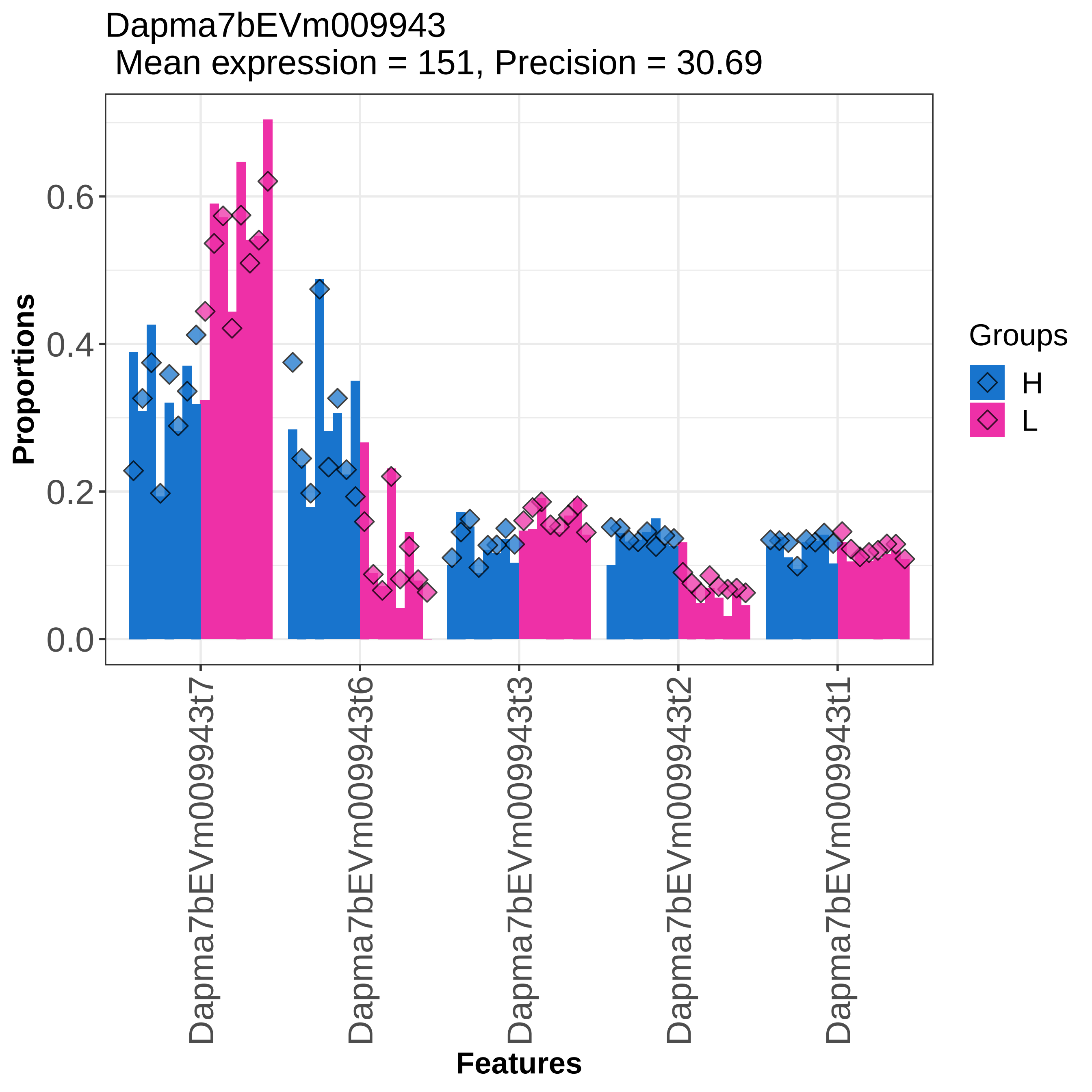
Dapma7bEVm009943
Mean expression = 151, Precision = 30.69
Proportions
0.0
0.2
0.4
0.6
Features
Dapma7bEVm009943t7
Dapma7bEVm009943t6
Dapma7bEVm009943t3
Dapma7bEVm009943t2
Dapma7bEVm009943t1
Groups
H
L

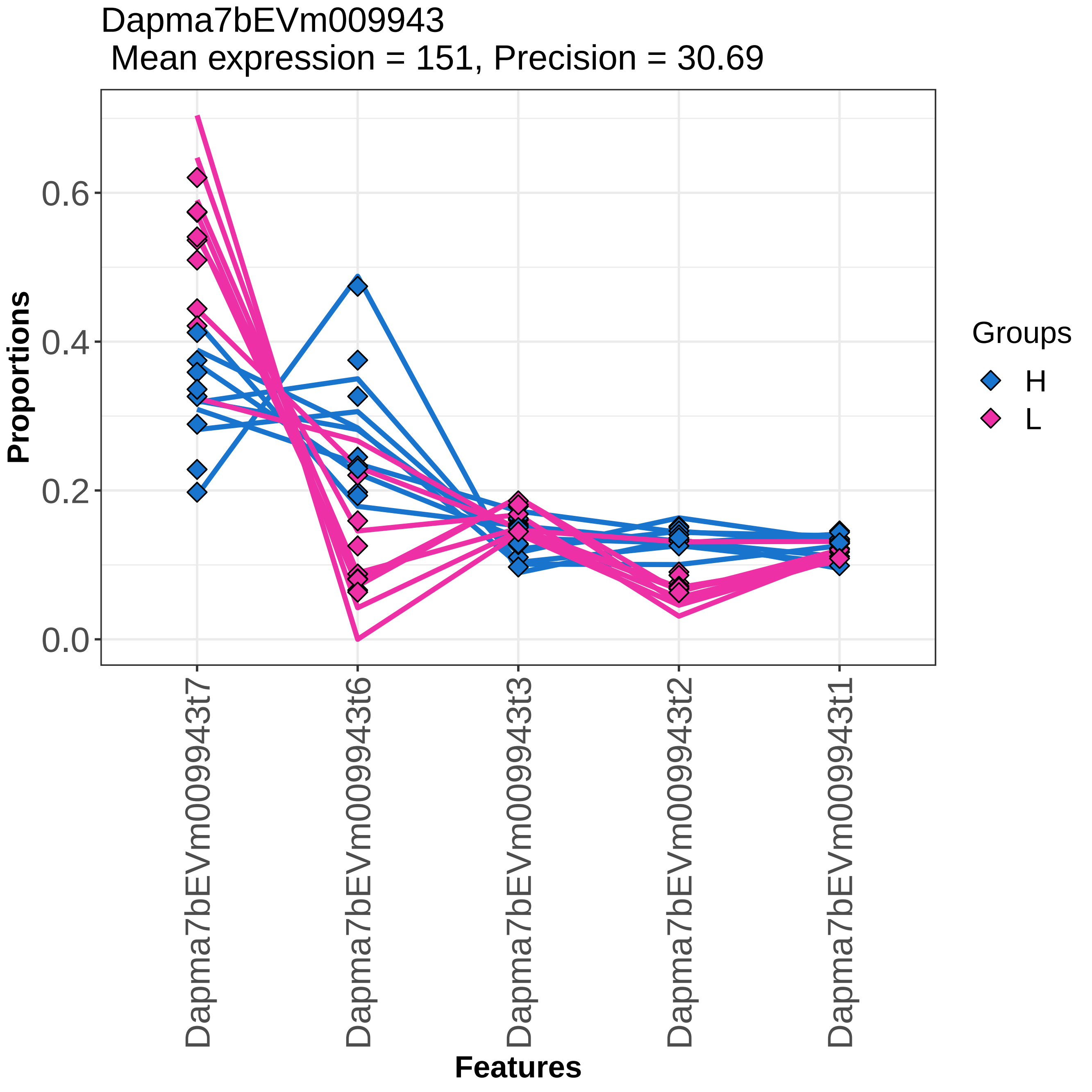
Dapma7bEVm009943
Mean expression = 151, Precision = 30.69
Proportions
0.0
0.2
0.4
0.6
Features
Dapma7bEVm009943t7
Dapma7bEVm009943t6
Dapma7bEVm009943t3
Dapma7bEVm009943t2
Dapma7bEVm009943t1
Groups
H
L

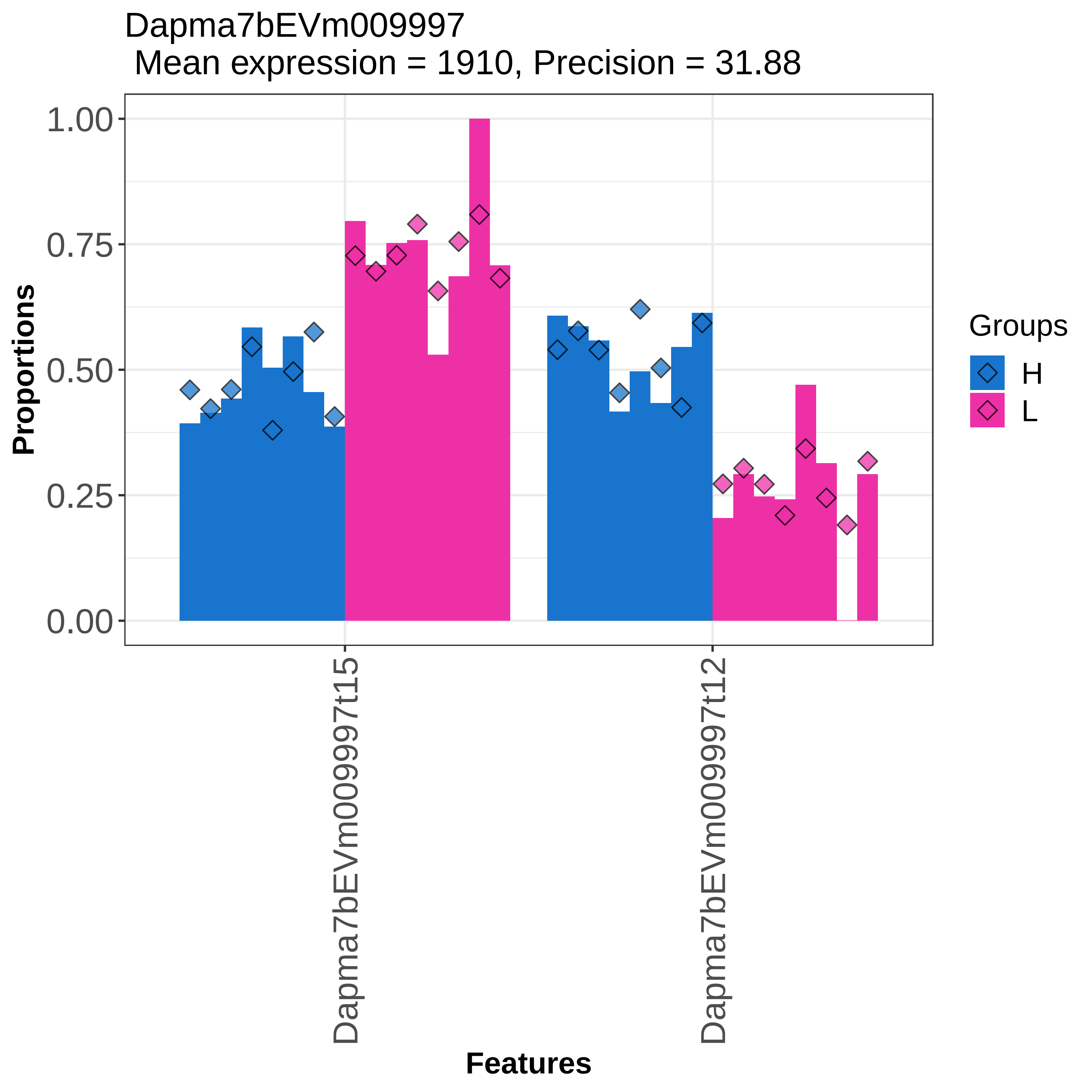

Dapma7bEVm009997
Mean expression = 1910, Precision = 31.88
Proportions
1.00
0.75
0.50
0.25
0.00
Dapma7bEVm009997t15
Dapma7bEVm009997t12
Features
Groups
H
L

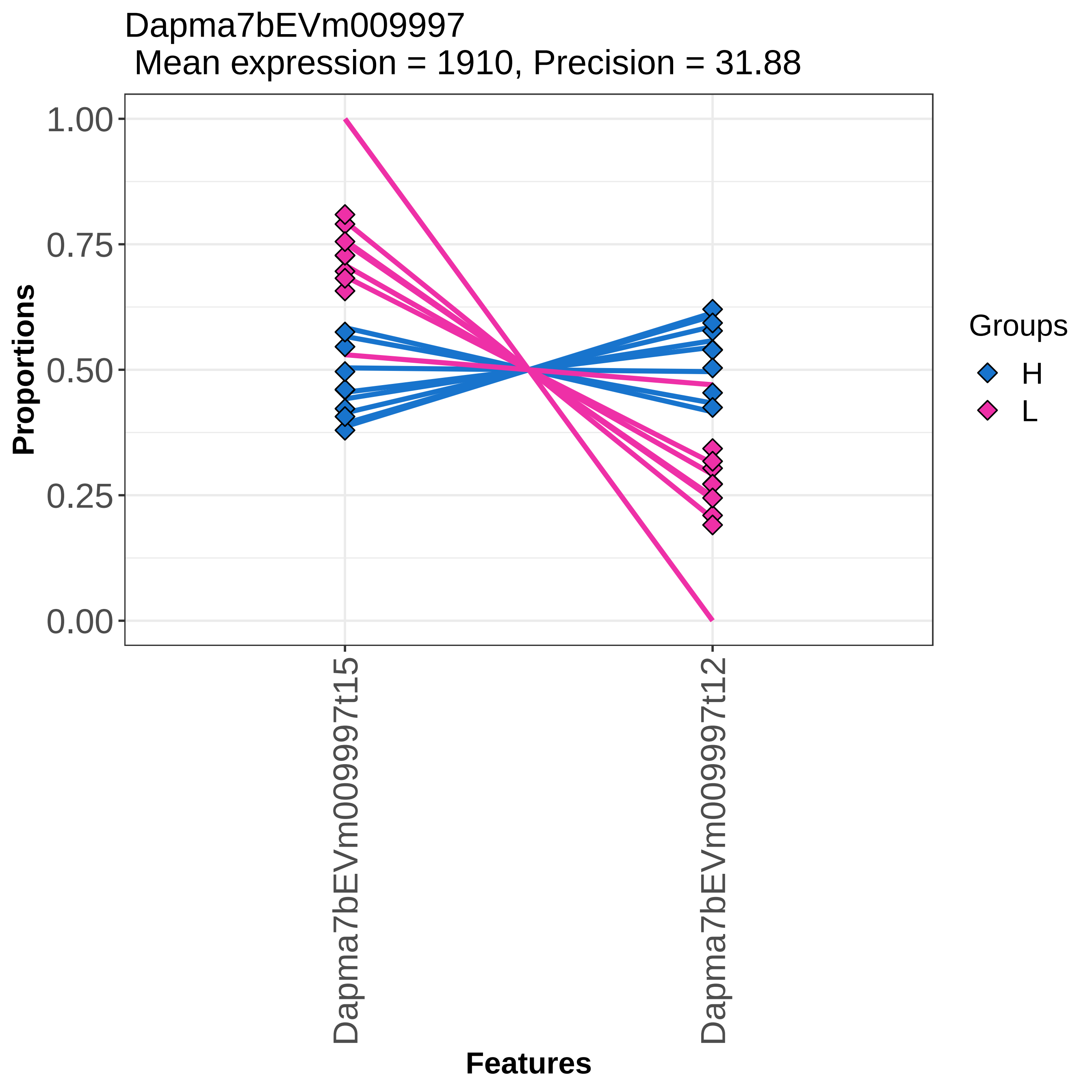

Dapma7bEVm009997
Mean expression = 1910, Precision = 31.88
Proportions
1.00
0.75
0.50
0.25
0.00
Dapma7bEVm009997t15
Dapma7bEVm009997t12
Features
Groups
H
L

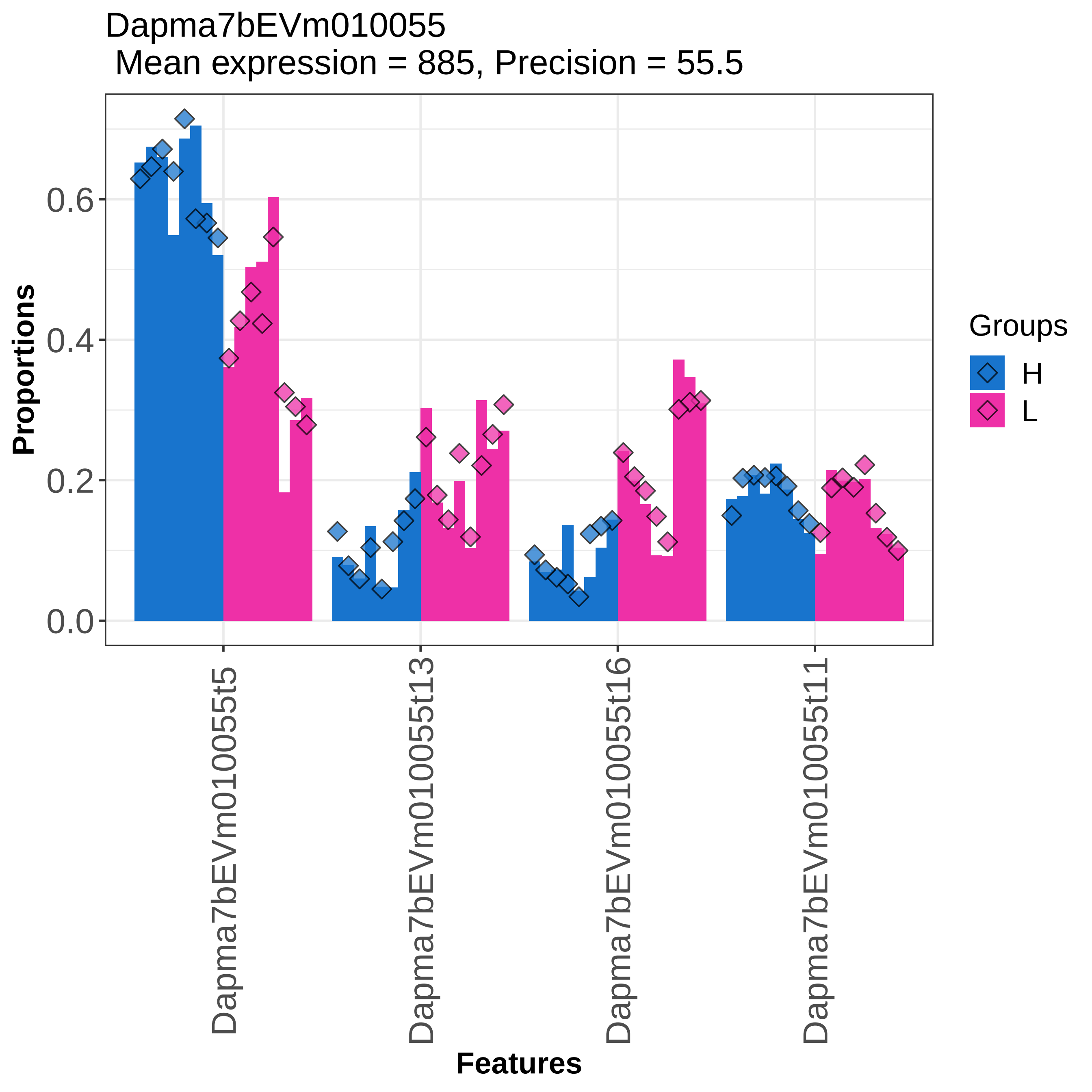

Dapma7bEVm010055
Mean expression = 885, Precision = 55.5
Proportions
0.0
0.2
0.4
0.6
Dapma7bEVm010055t5
Dapma7bEVm010055t13
Dapma7bEVm010055t16
Dapma7bEVm010055t11
Features
Groups
H
L

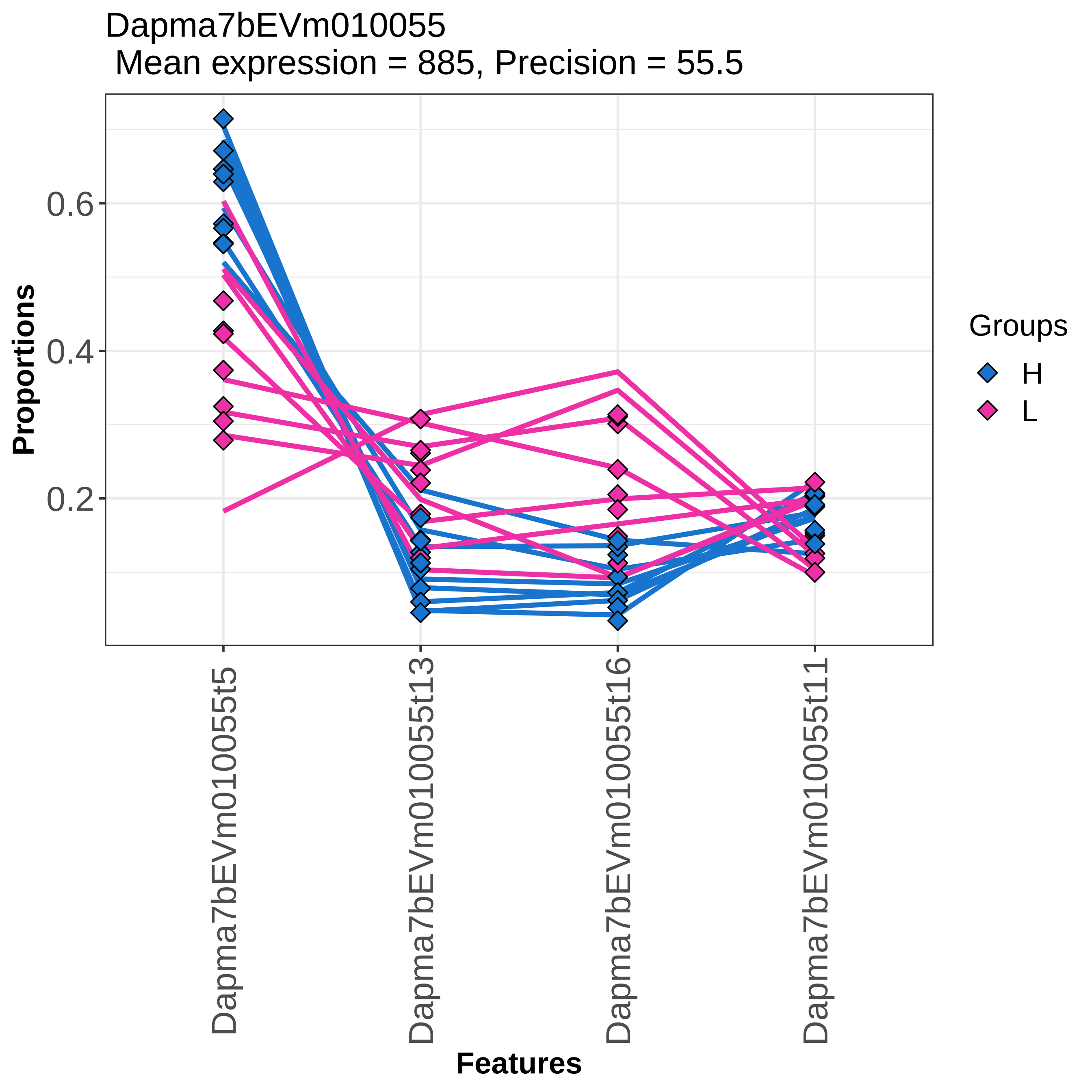

Dapma7bEVm010055
Mean expression = 885, Precision = 55.5
Proportions
0.6
0.4
0.2
Groups
H
L
Dapma7bEVm010055t5
Dapma7bEVm010055t13
Dapma7bEVm010055t16
Dapma7bEVm010055t11
Features

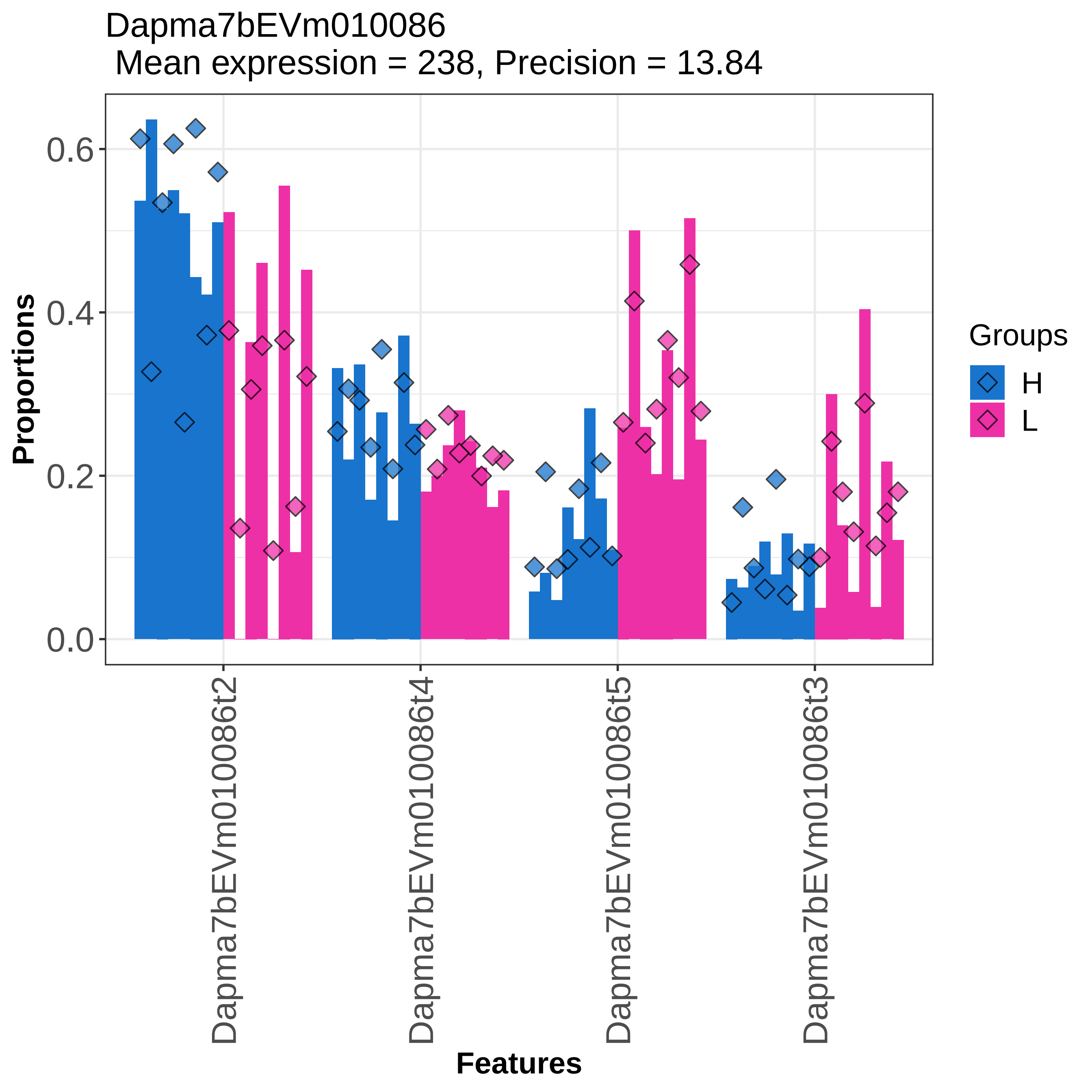

Dapma7bEVm010086
Mean expression = 238, Precision = 13.84
Proportions
0.0
0.2
0.4
0.6
Features
Dapma7bEVm010086t2
Dapma7bEVm010086t4
Dapma7bEVm010086t5
Dapma7bEVm010086t3
Groups
H
L

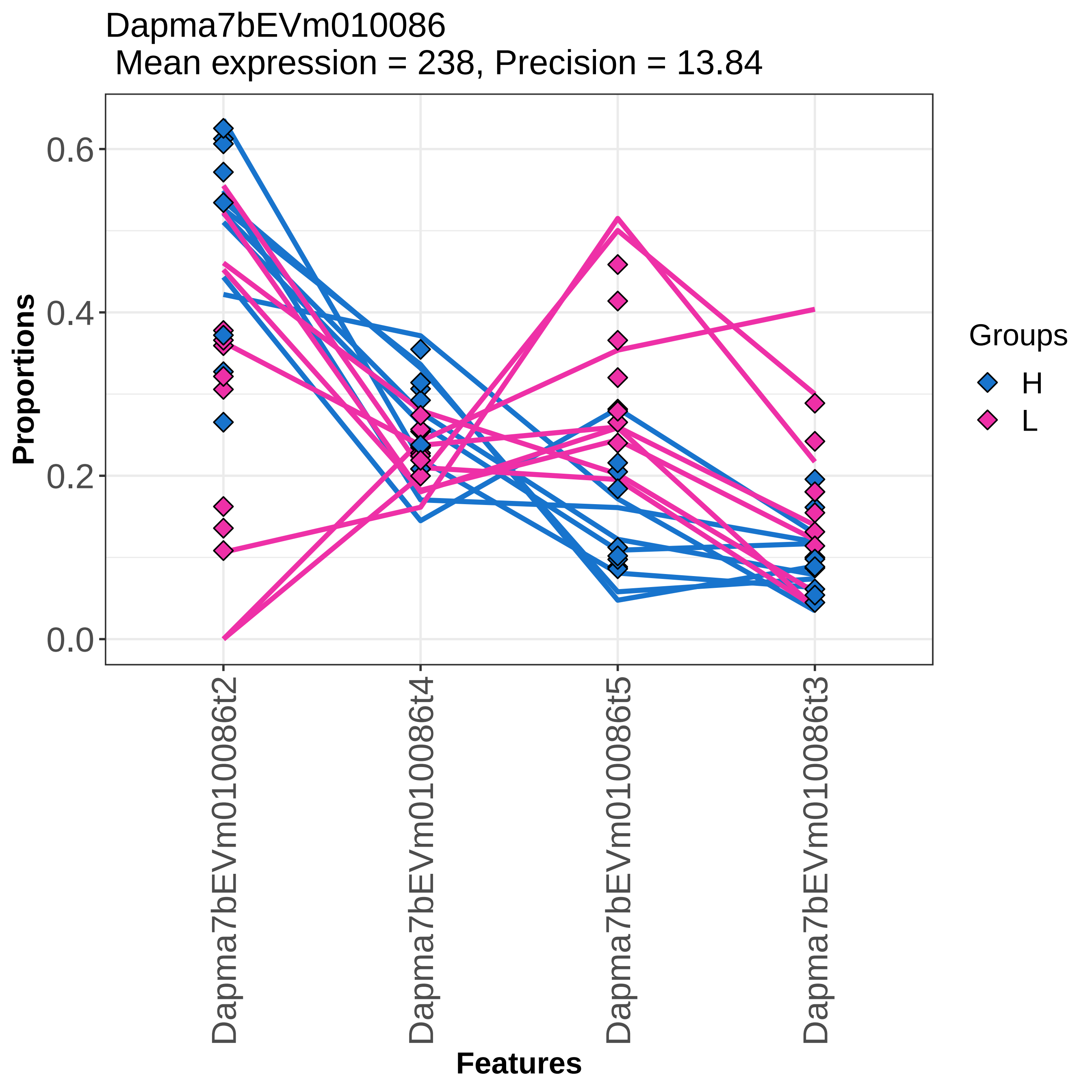
Dapma7bEVm010086
Mean expression = 238, Precision = 13.84
Proportions
0.0
0.2
0.4
0.6
Features
Dapma7bEVm010086t2
Dapma7bEVm010086t4
Dapma7bEVm010086t5
Dapma7bEVm010086t3
Groups
H
L

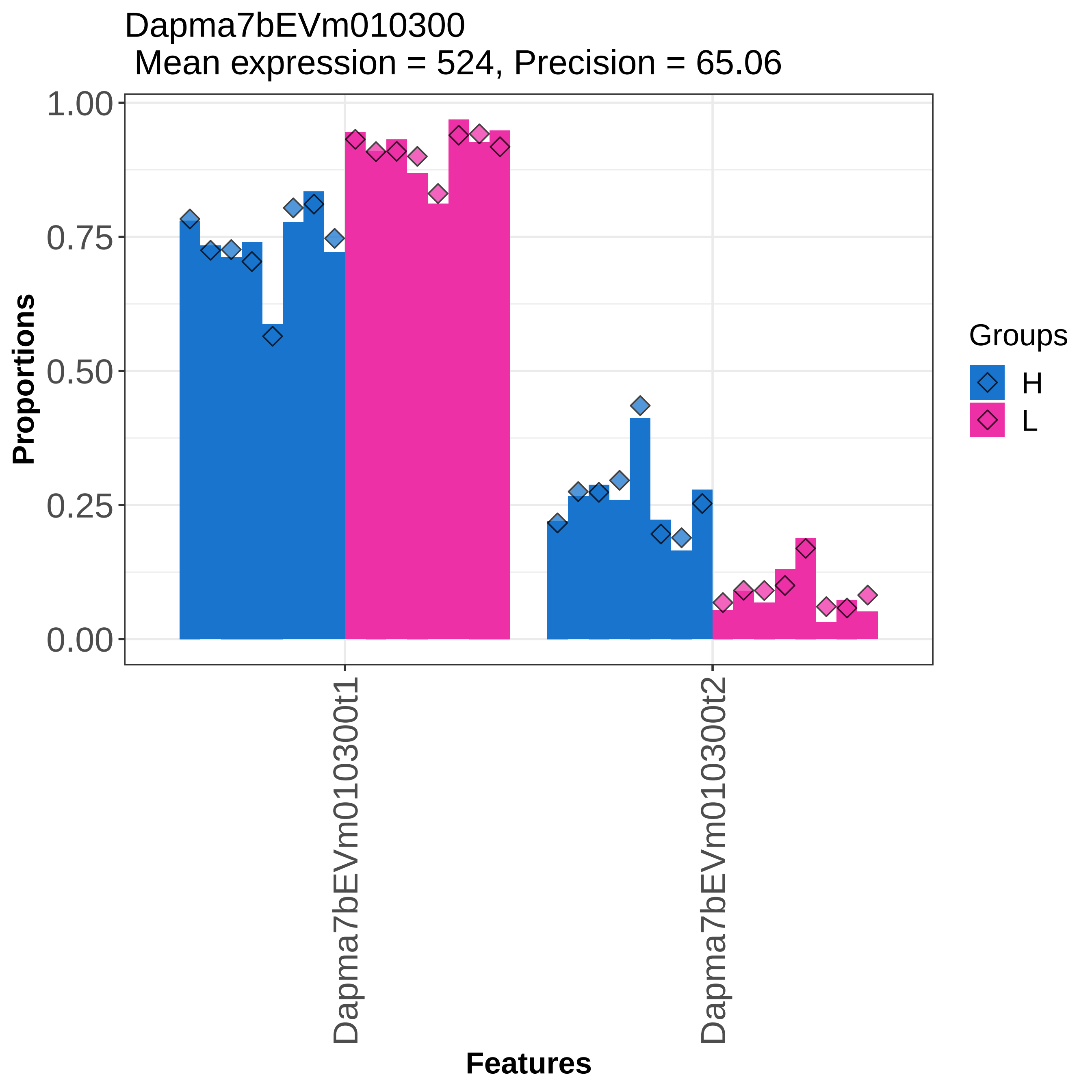

Dapma7bEVm010300
Mean expression = 524, Precision = 65.06
Proportions
1.00
0.75
0.50
0.25
0.00
Dapma7bEVm010300t1
Dapma7bEVm010300t2
Features
Groups
H
L

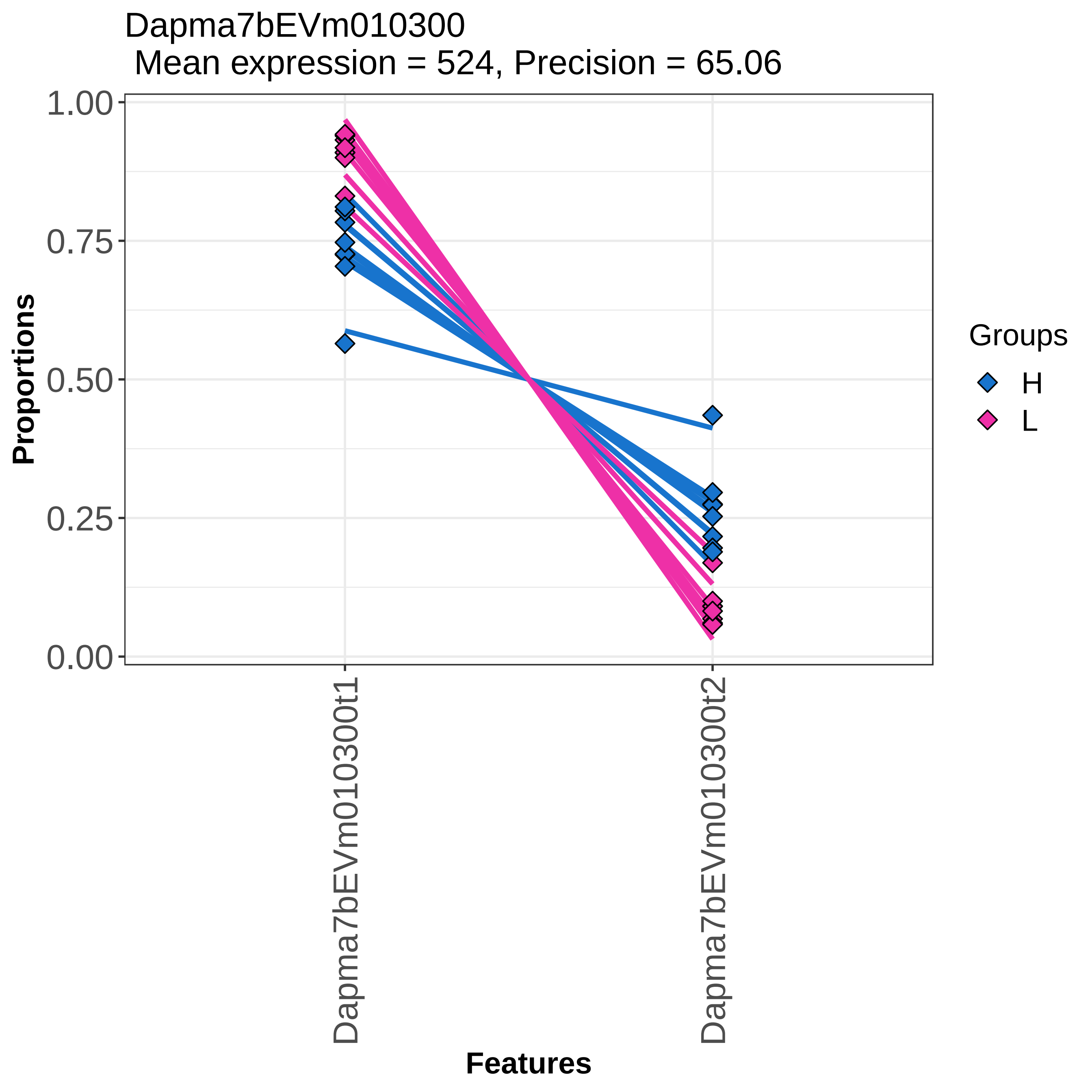

Dapma7bEVm010300
Mean expression = 524, Precision = 65.06
Proportions
1.00
0.75
0.50
0.25
0.00
Dapma7bEVm010300t1
Dapma7bEVm010300t2
Features
Groups
H
L

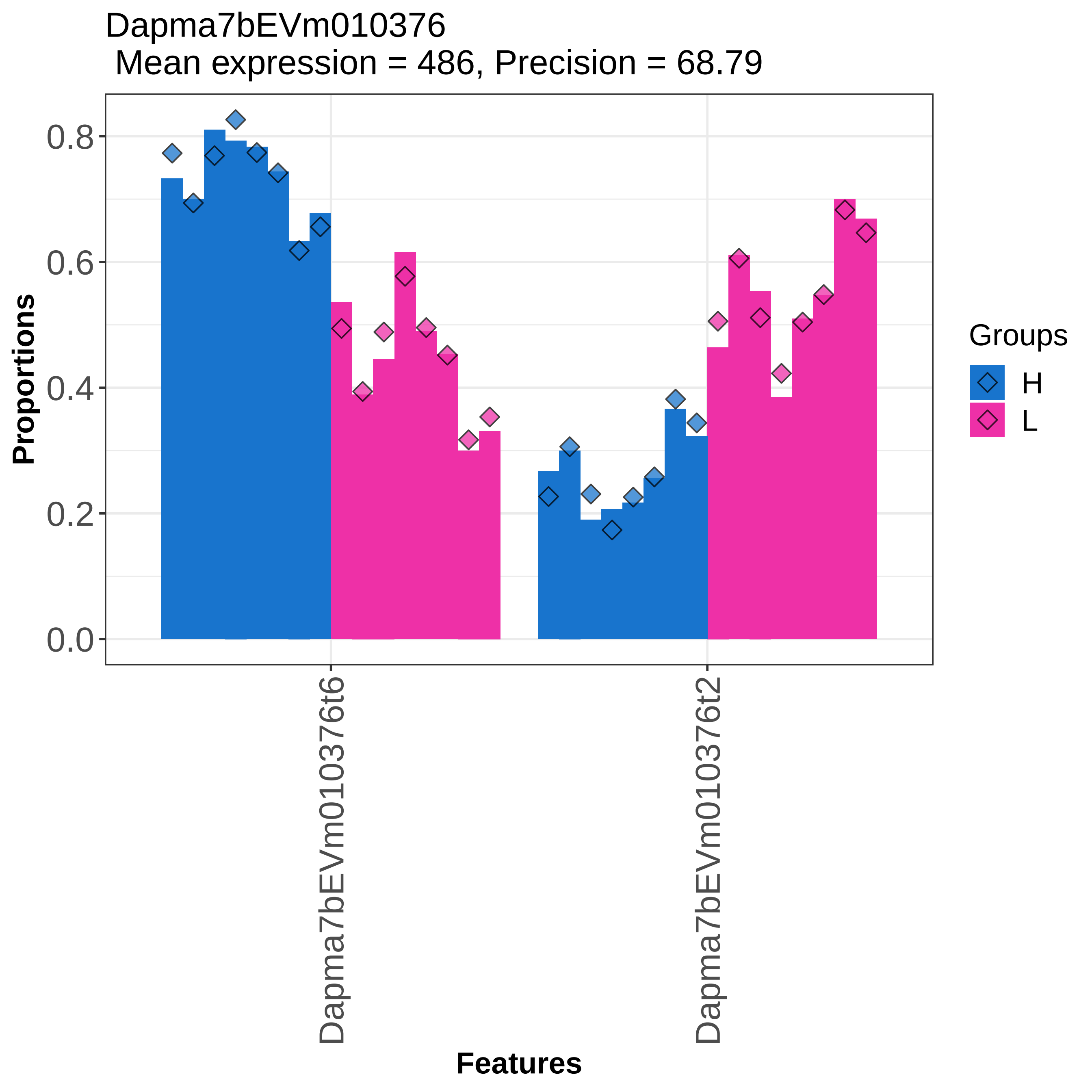
Dapma7bEVm010376
Mean expression = 486, Precision = 68.79
Proportions
0.0
0.2
0.4
0.6
0.8
Dapma7bEVm010376t6
Dapma7bEVm010376t2
Features
Groups
H
L

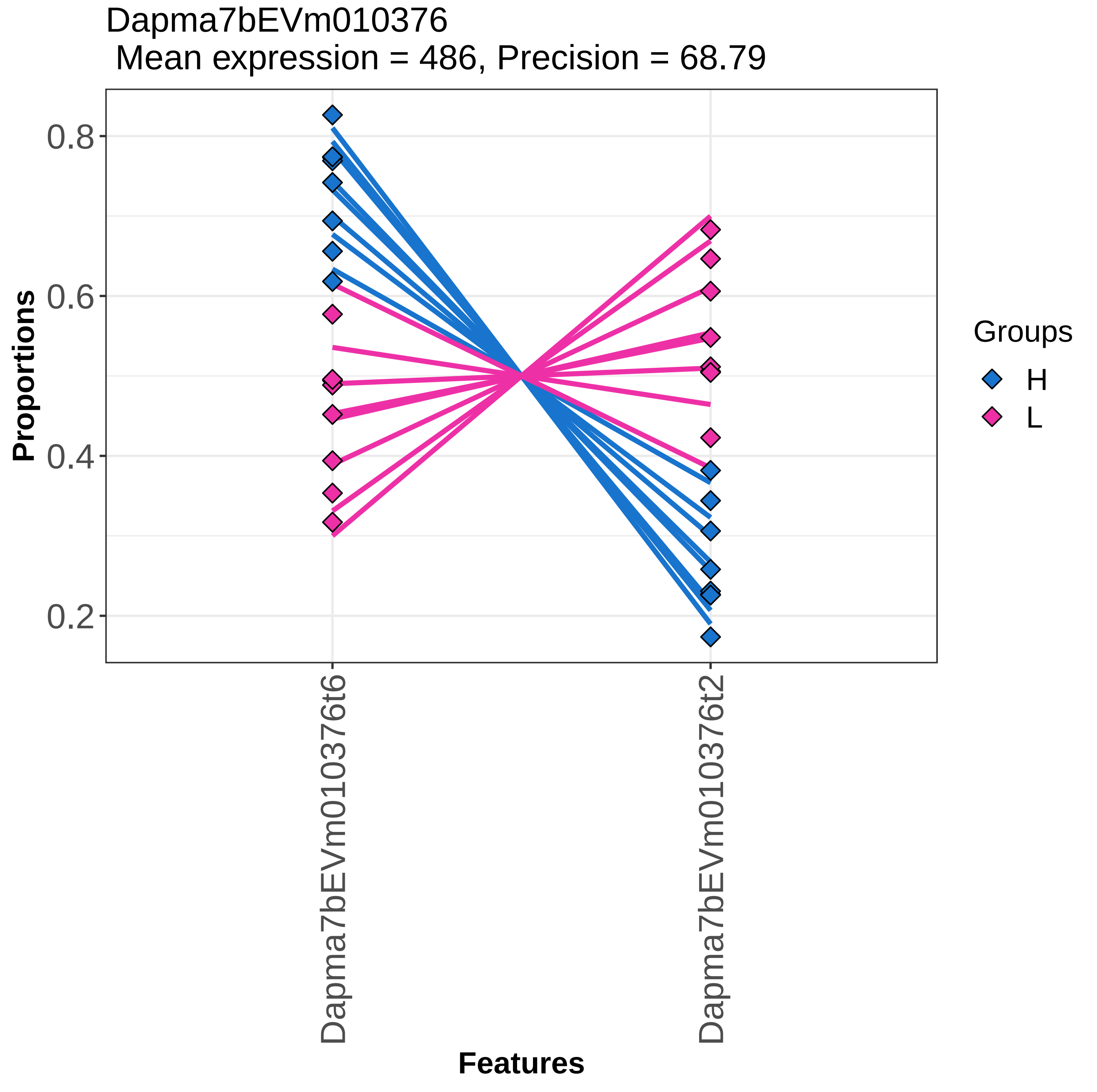
Dapma7bEVm010376
Mean expression = 486, Precision = 68.79
Proportions
0.8
0.6
0.4
0.2
Dapma7bEVm010376t6
Dapma7bEVm010376t2
Features
Groups
H
L

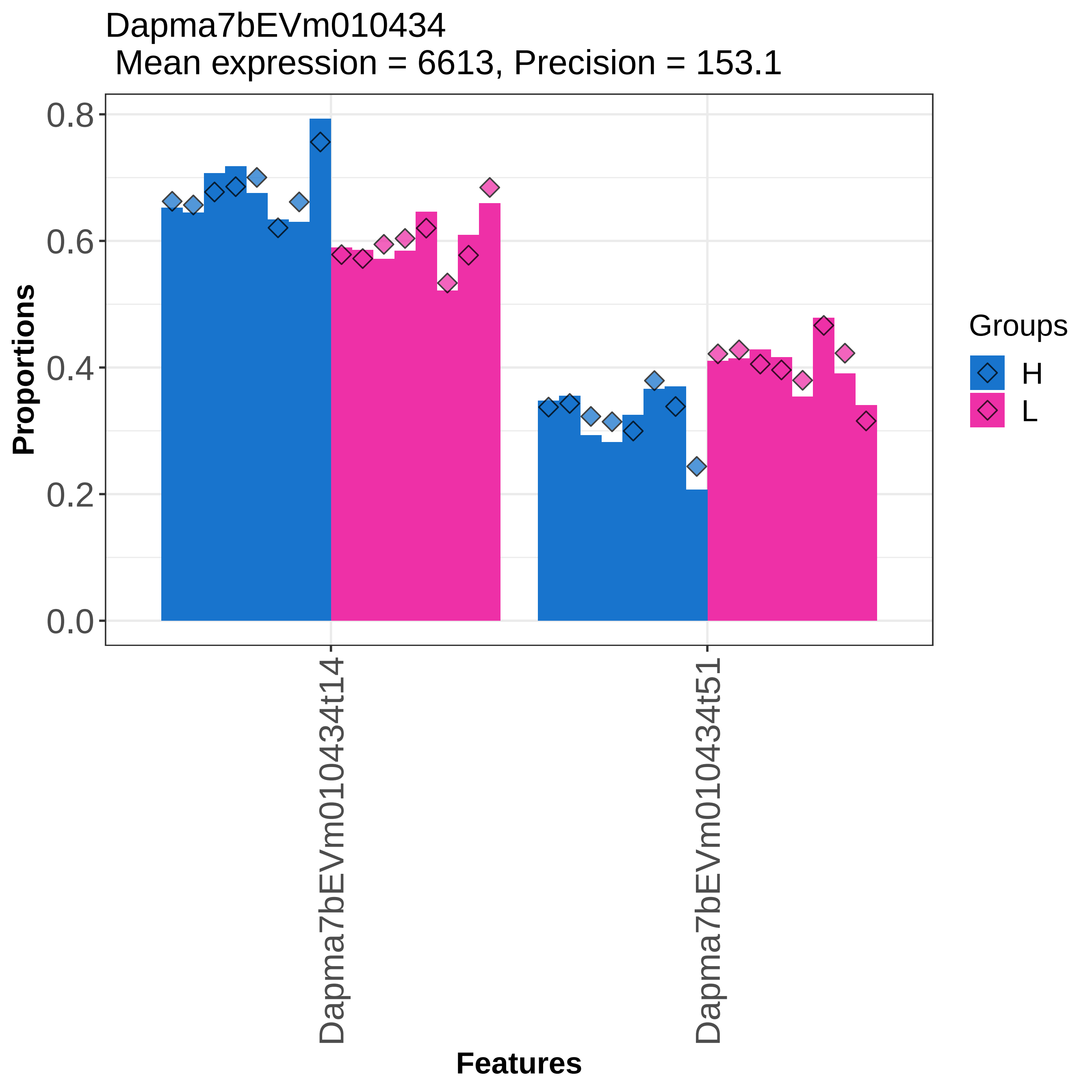

Dapma7bEVm010434
Mean expression = 6613, Precision = 153.1
Proportions
0.0
0.2
0.4
0.6
0.8
Dapma7bEVm010434t14
Dapma7bEVm010434t51
Features
Groups
H
L

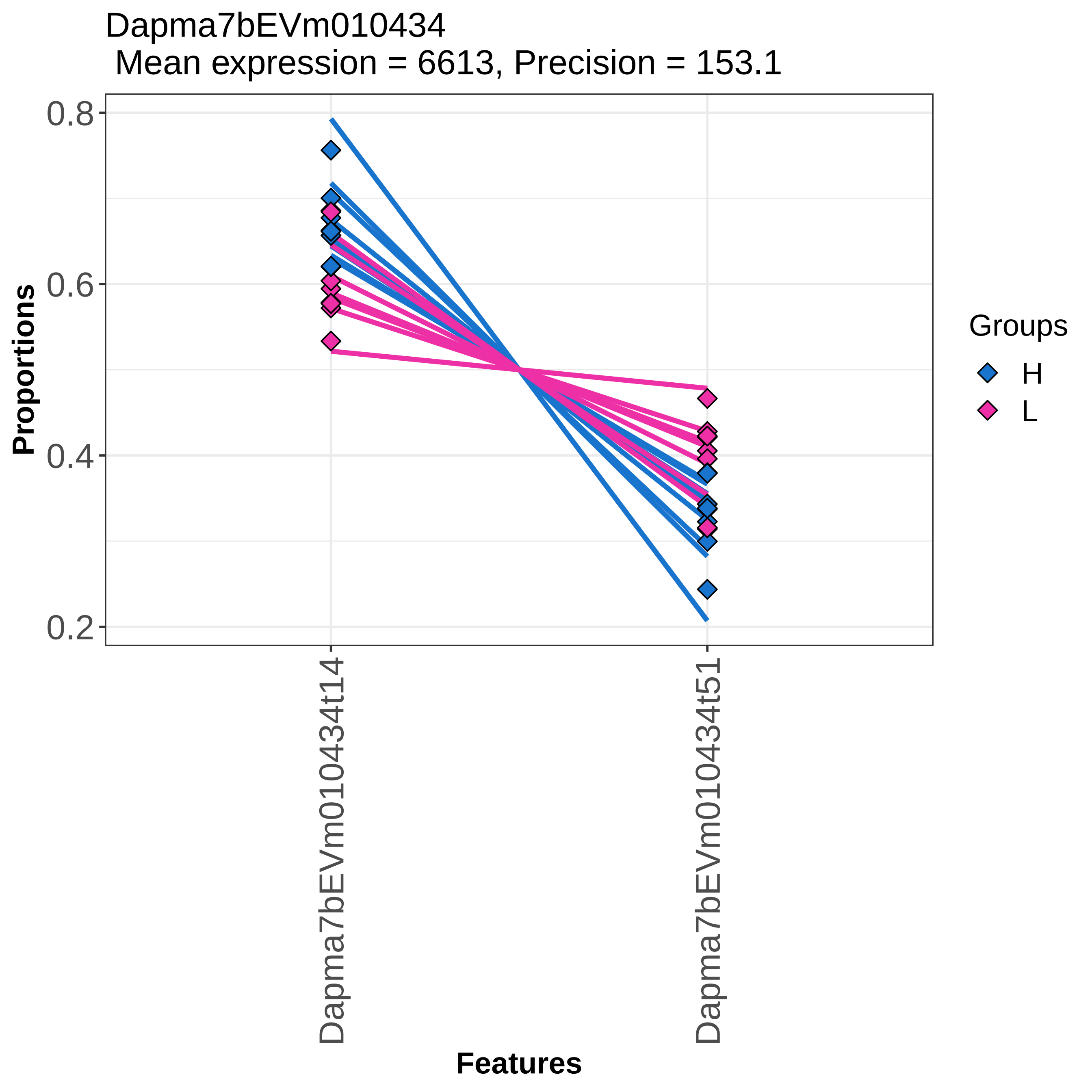
Dapma7bEVm010434
Mean expression = 6613, Precision = 153.1
0.8
0.6
0.4
0.2
Proportions
Dapma7bEVm010434t14
Dapma7bEVm010434t51
Features
Groups
H
L

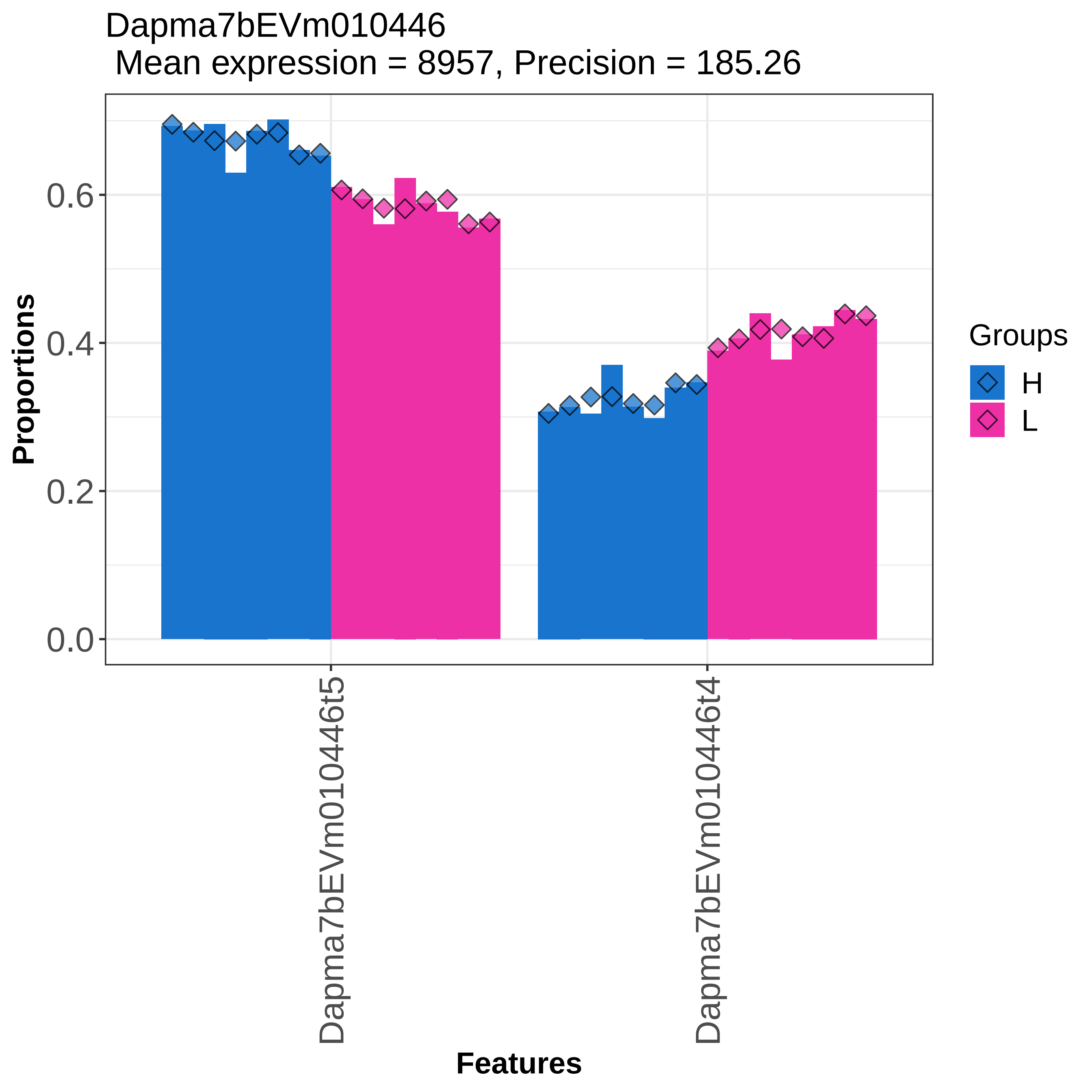

Dapma7bEVm010446
Mean expression = 8957, Precision = 185.26
Proportions
0.0
0.2
0.4
0.6
Dapma7bEVm010446t5
Dapma7bEVm010446t4
Features
Groups
H
L

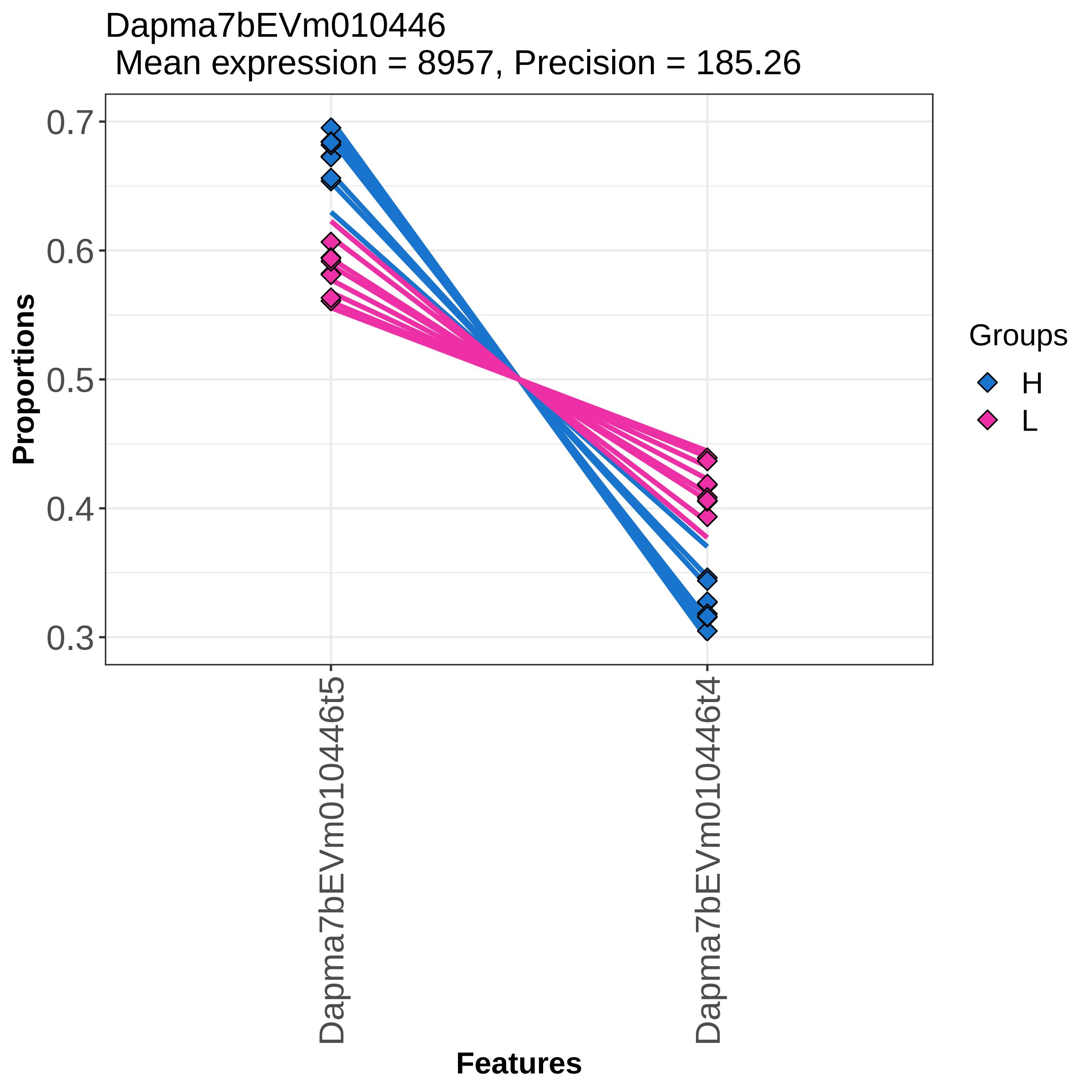

Dapma7bEVm010446
Mean expression = 8957, Precision = 185.26
Proportions
0.7
0.6
0.5
0.4
0.3
Dapma7bEVm010446t5
Dapma7bEVm010446t4
Features
Groups
H
L

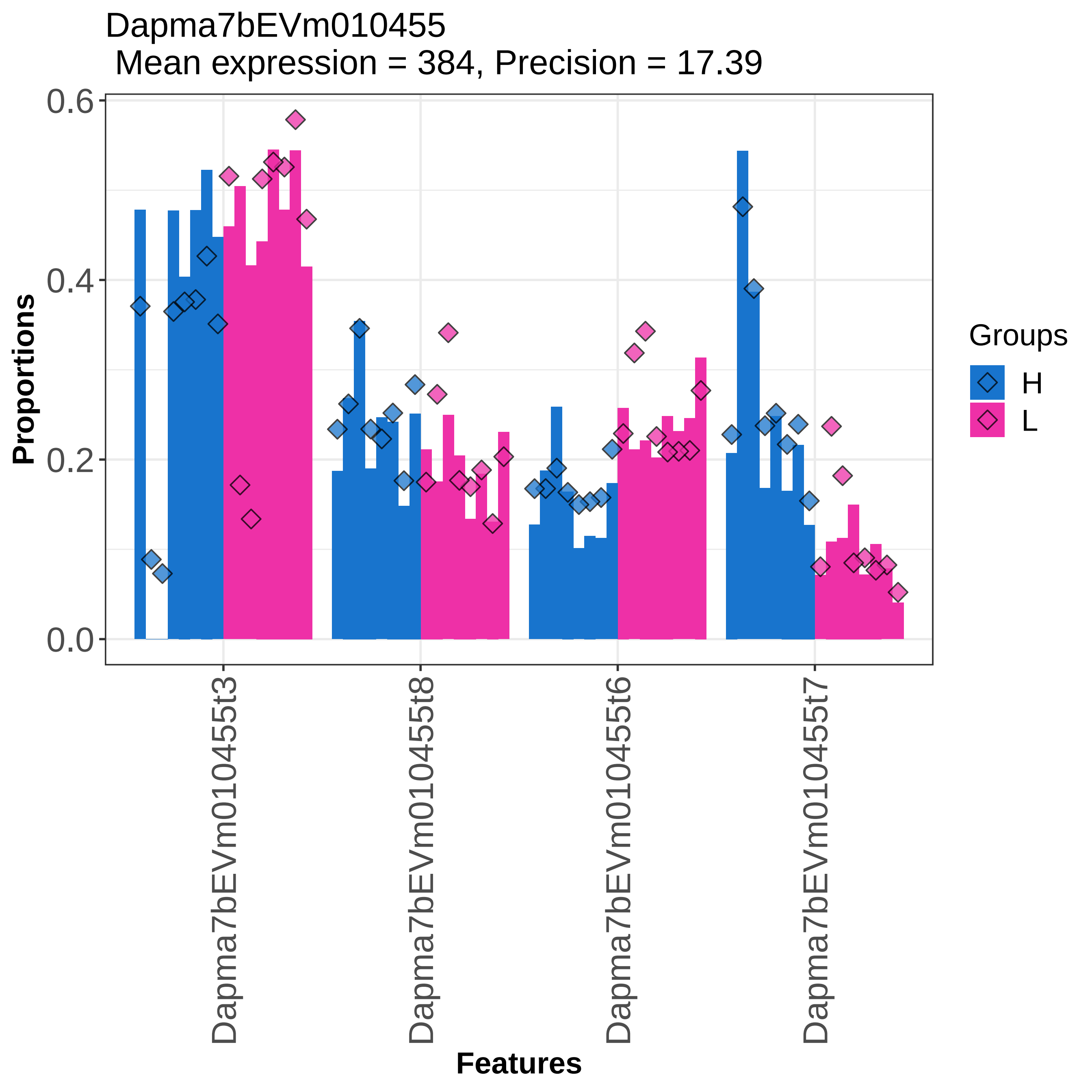
Dapma7bEVm010455
Mean expression = 384, Precision = 17.39
Proportions
0.6
0.4
0.2
0.0
Dapma7bEVm010455t3
Dapma7bEVm010455t8
Dapma7bEVm010455t6
Dapma7bEVm010455t7
Features
Groups
H
L

Dapma7bEVm010455
Mean expression = 384, Precision = 17.39

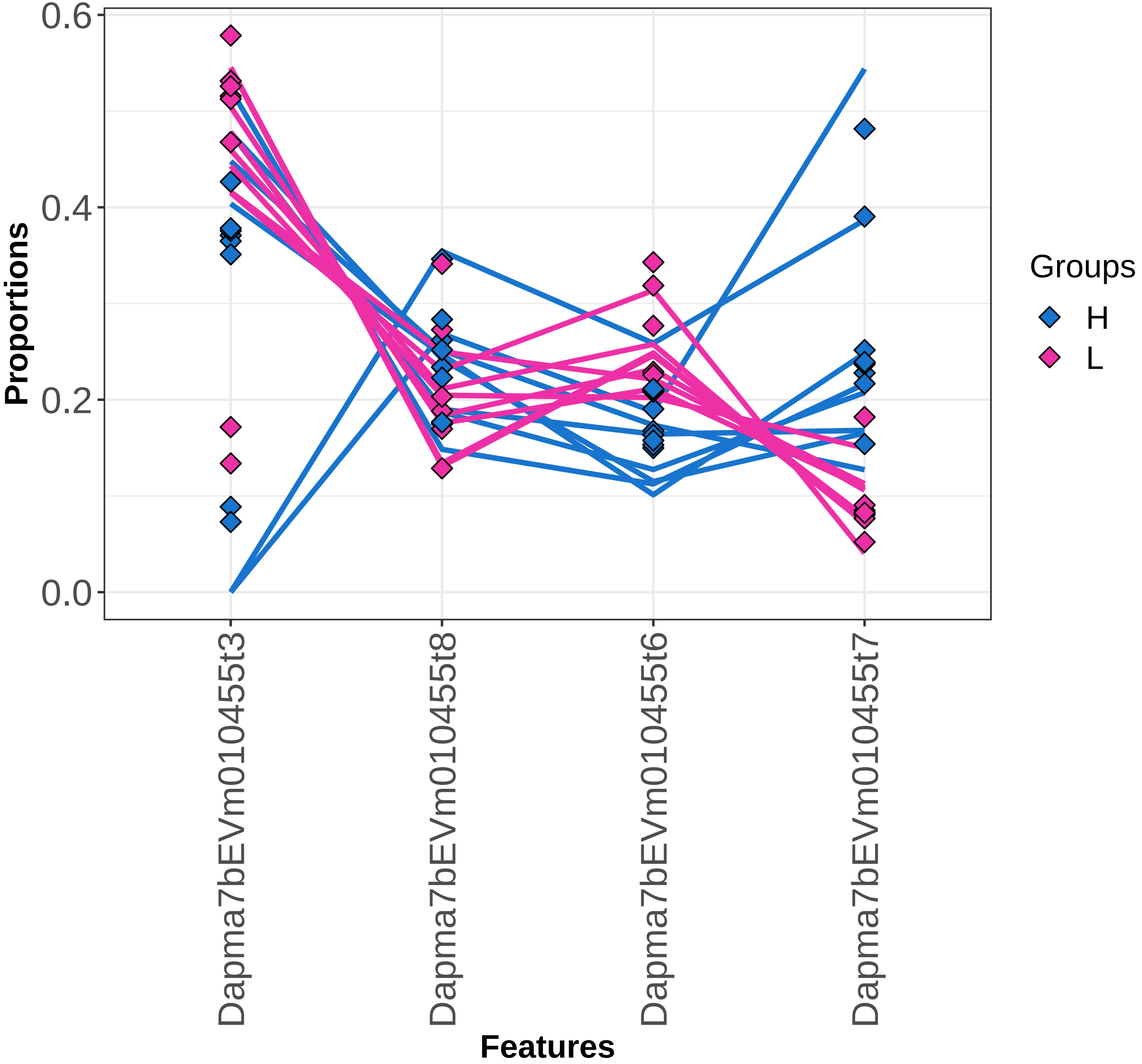

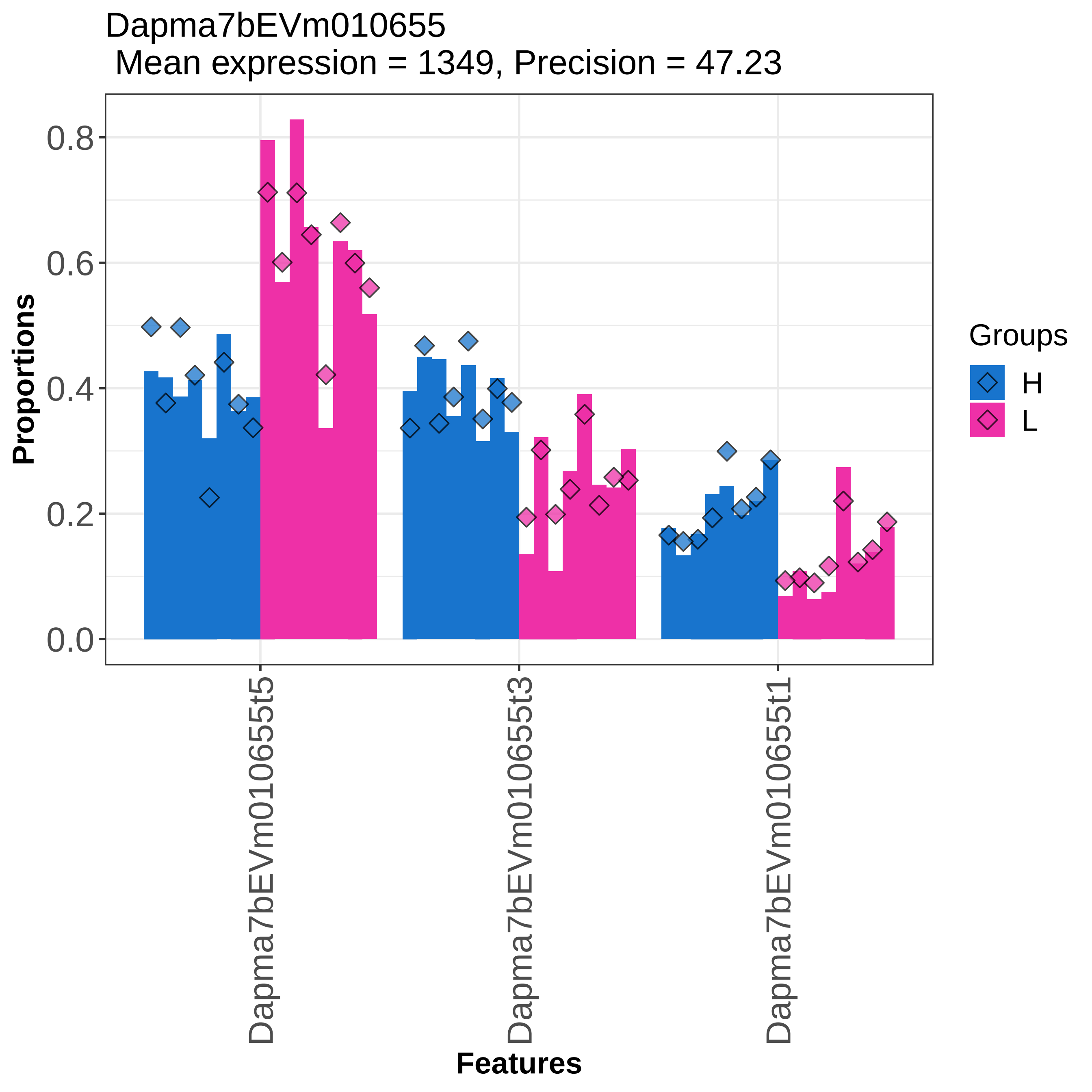
Dapma7bEVm010655
Mean expression = 1349, Precision = 47.23
Proportions
0.0
0.2
0.4
0.6
0.8
Dapma7bEVm010655t5
Dapma7bEVm010655t3
Dapma7bEVm010655t1
Features
Groups
H
L

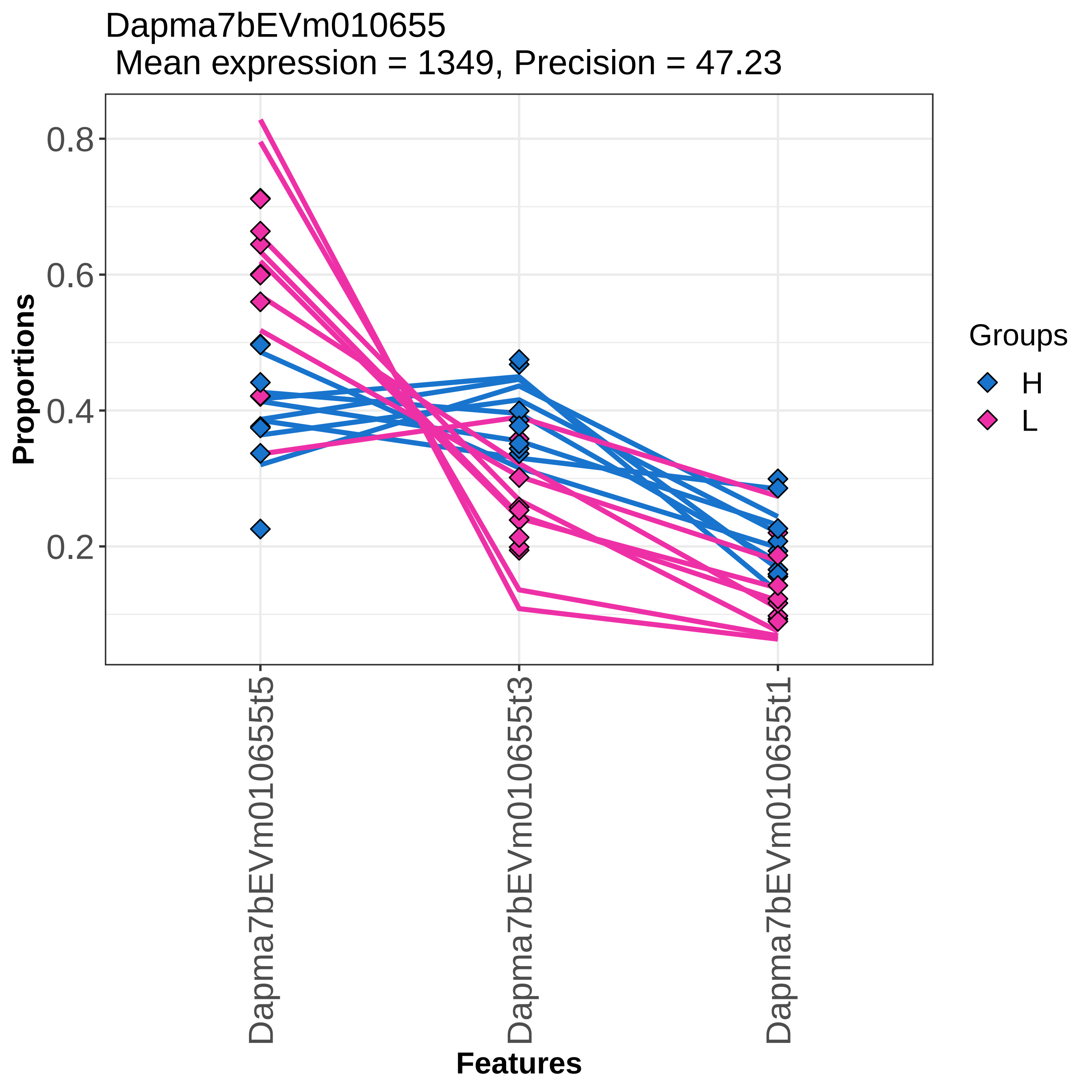
Dapma7bEVm010655
Mean expression = 1349, Precision = 47.23
Proportions
0.8
0.6
0.4
0.2
Features
Dapma7bEVm010655t5
Dapma7bEVm010655t3
Dapma7bEVm010655t1
Groups
H
L

Dapma7bEVm010714
Mean expression = 274, Precision = 17.85

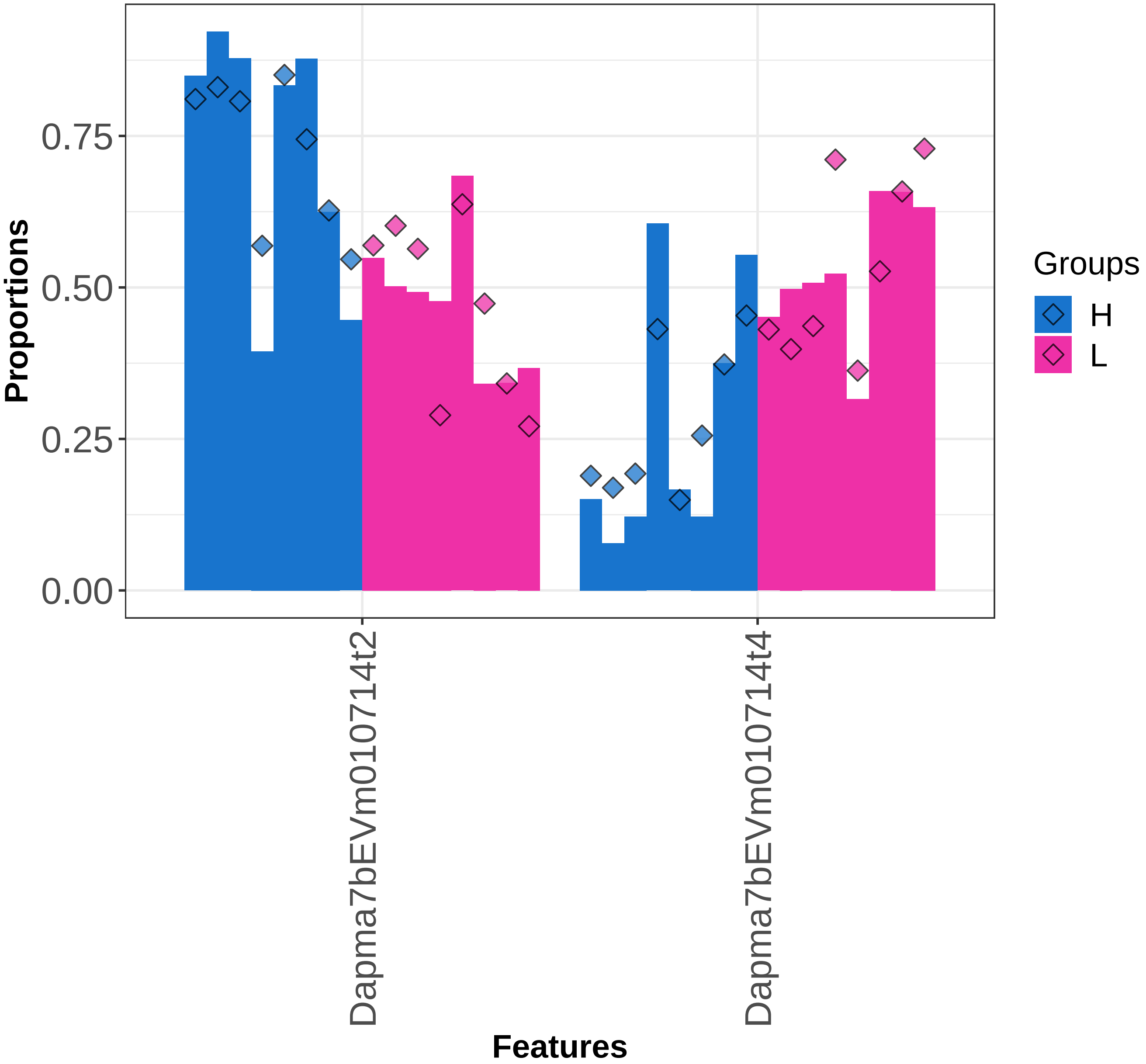

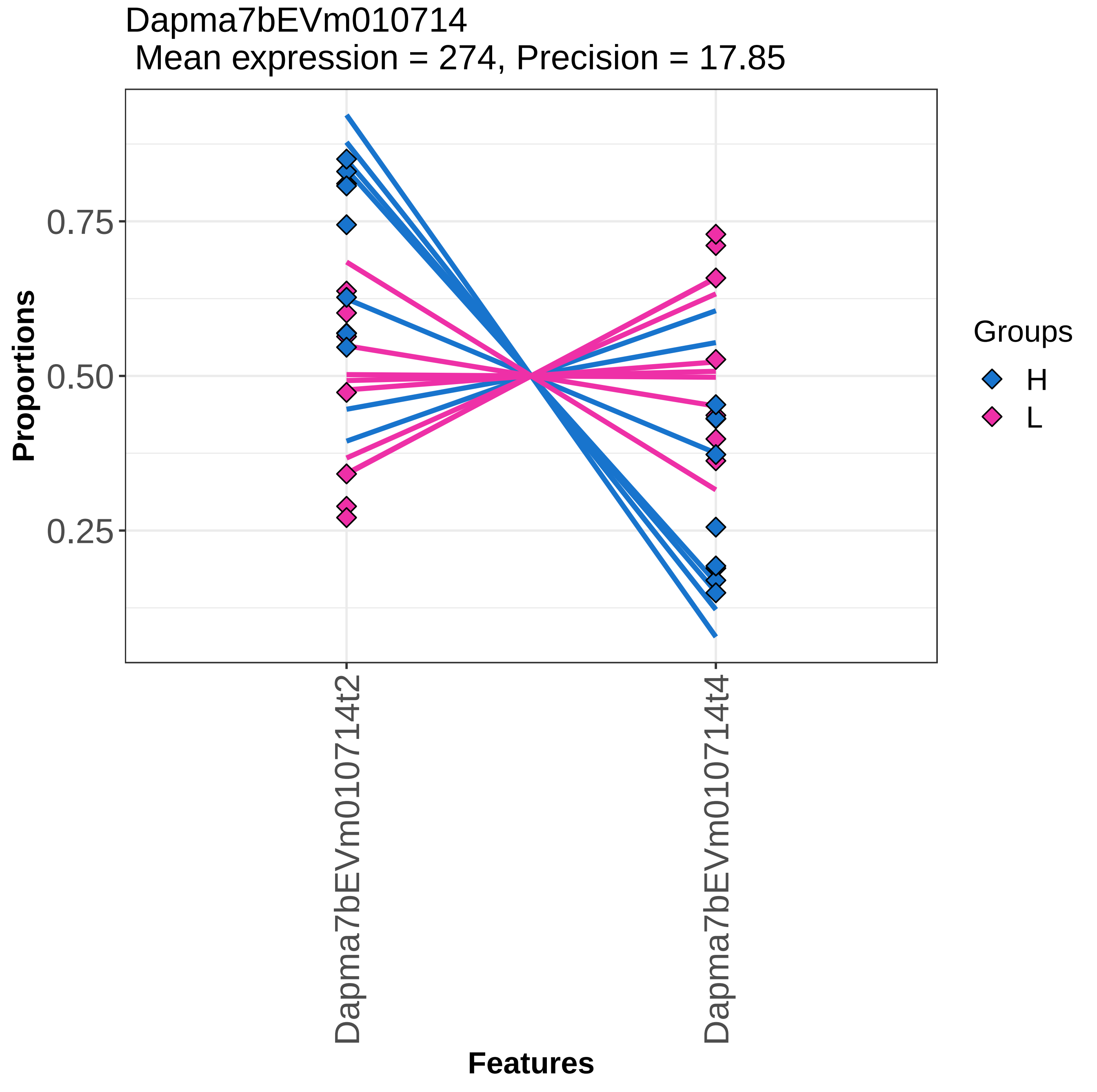

Dapma7bEVm010714
Mean expression = 274, Precision = 17.85
Proportions
0.75
0.50
0.25
Dapma7bEVm010714t2
Dapma7bEVm010714t4
Features
Groups
H
L

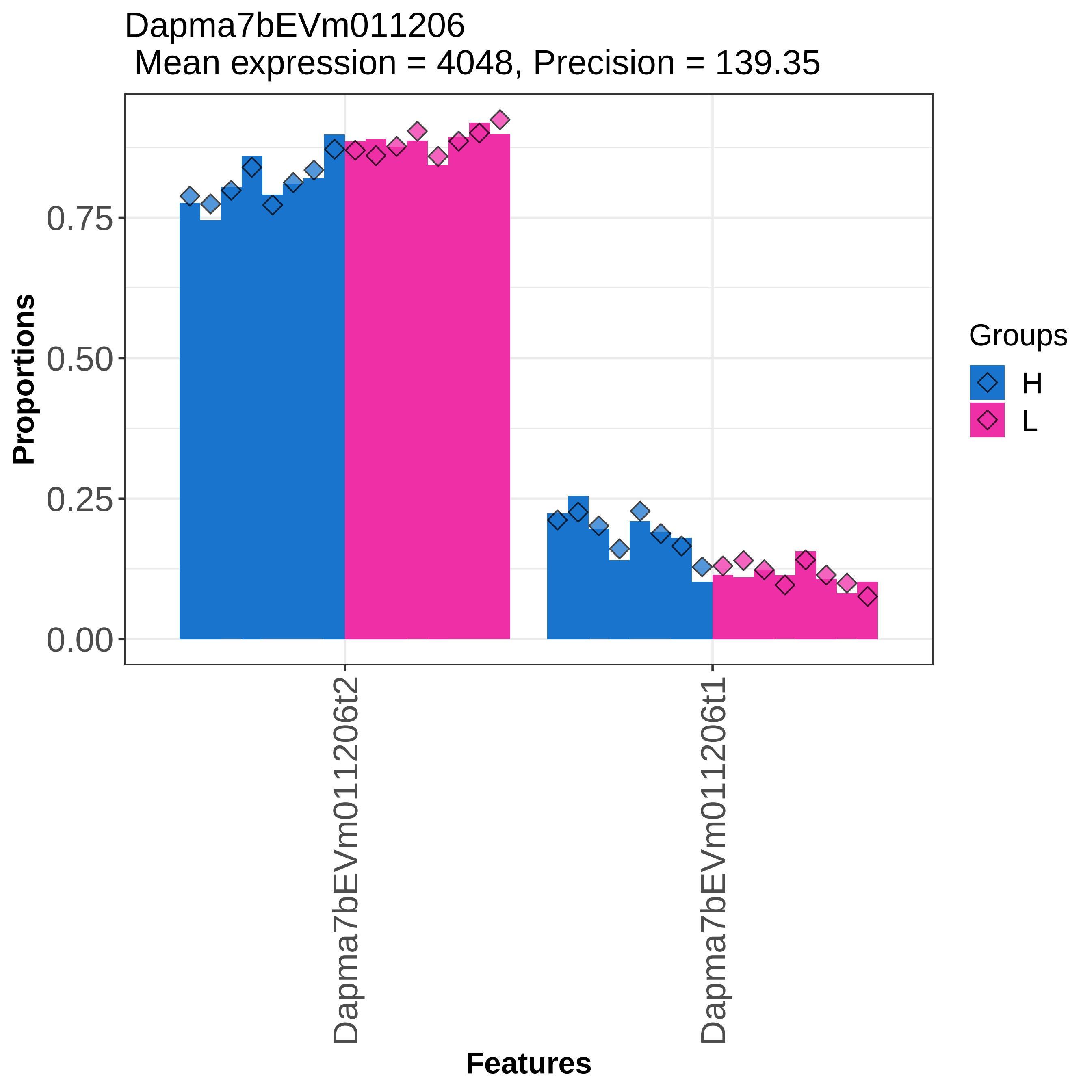
Dapma7bEVm011206
Mean expression = 4048, Precision = 139.35
Proportions
0.00
0.25
0.50
0.75
Dapma7bEVm011206t2
Dapma7bEVm011206t1
Features
Groups
H
L

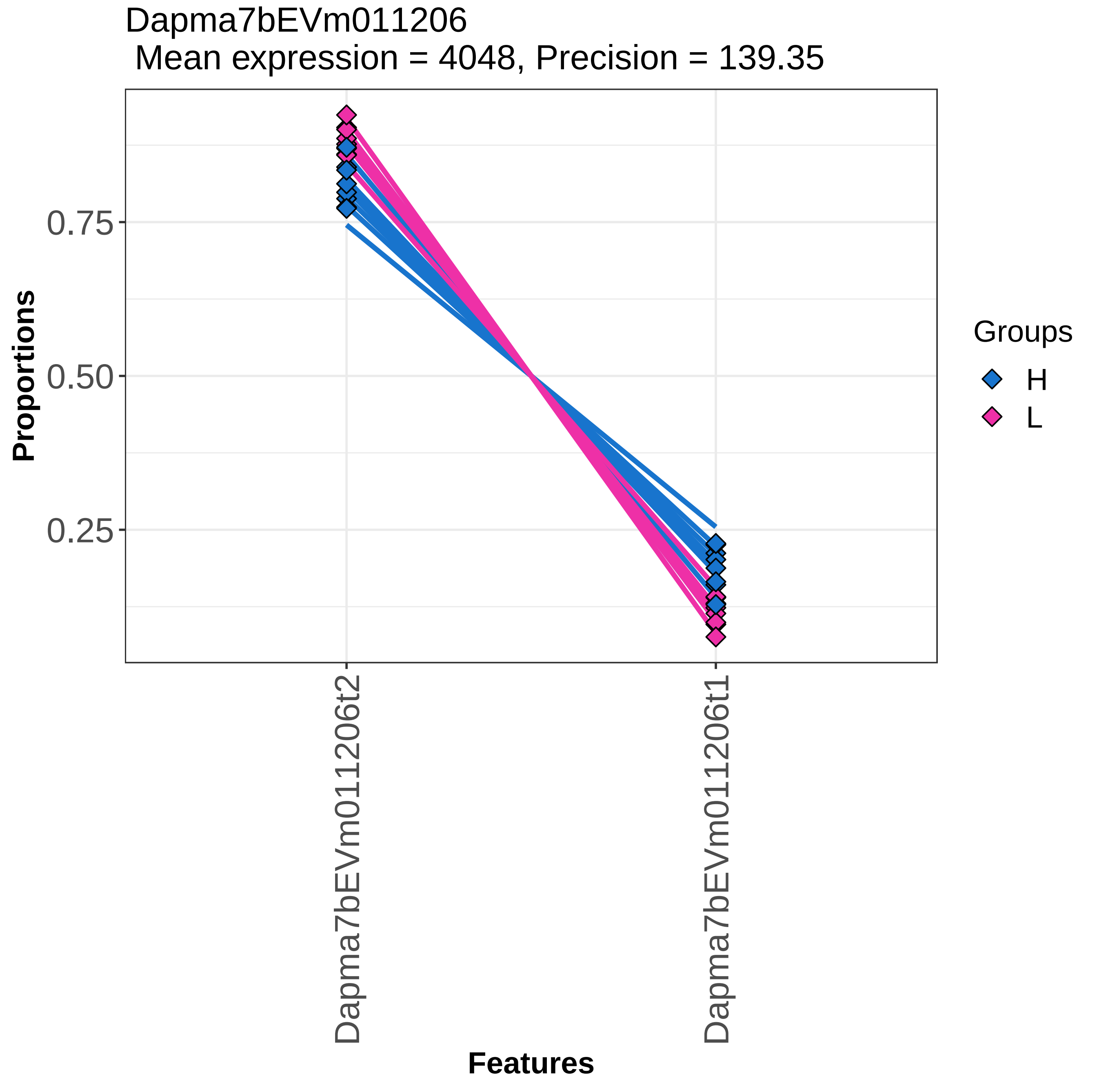
Dapma7bEVm011206
Mean expression = 4048, Precision = 139.35
Proportions
0.75
0.50
0.25
Dapma7bEVm011206t2
Dapma7bEVm011206t1
Features
Groups
H
L

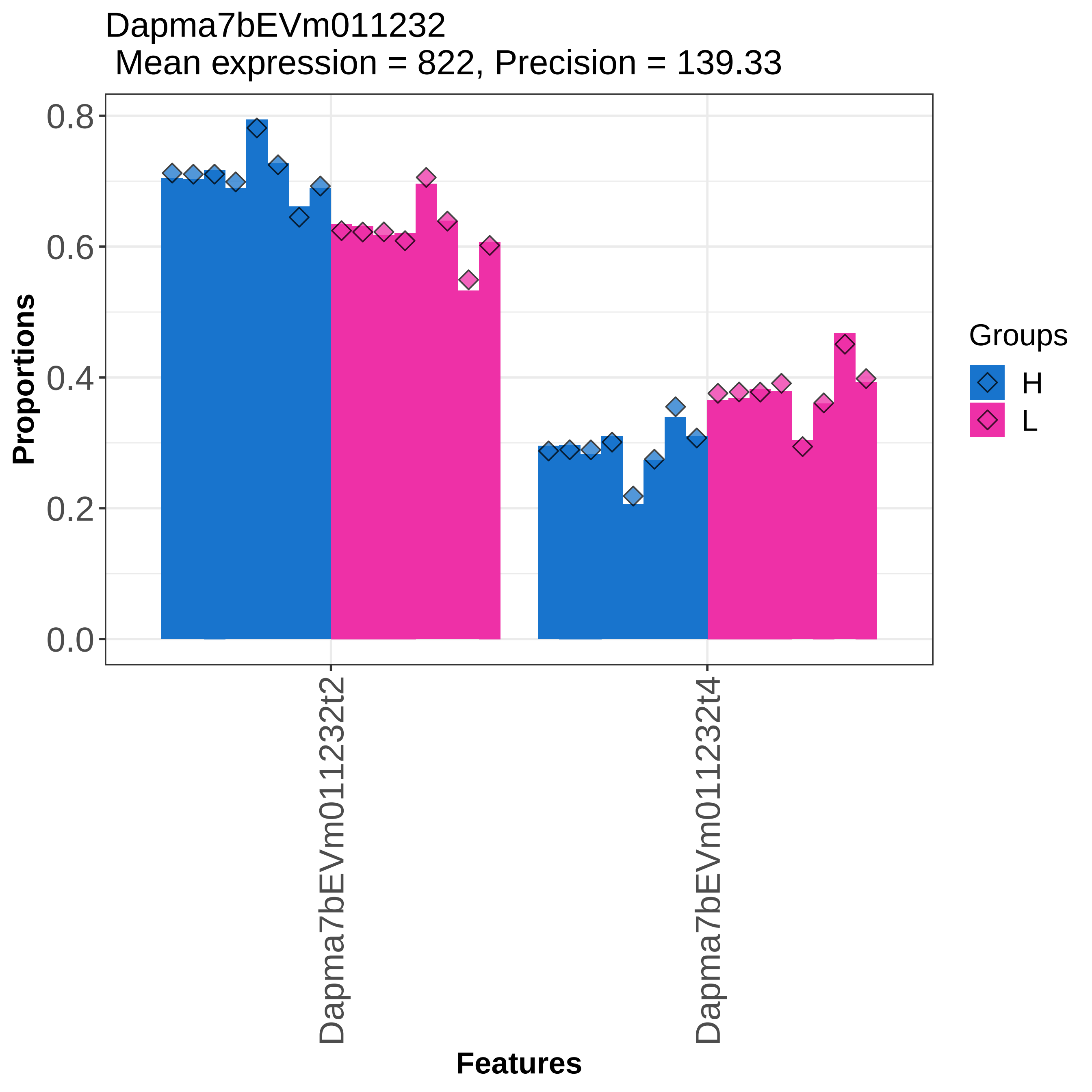
Dapma7bEVm011232
Mean expression = 822, Precision = 139.33
Proportions
0.0
0.2
0.4
0.6
0.8
Dapma7bEVm011232t2
Dapma7bEVm011232t4
Features
Groups
H
L

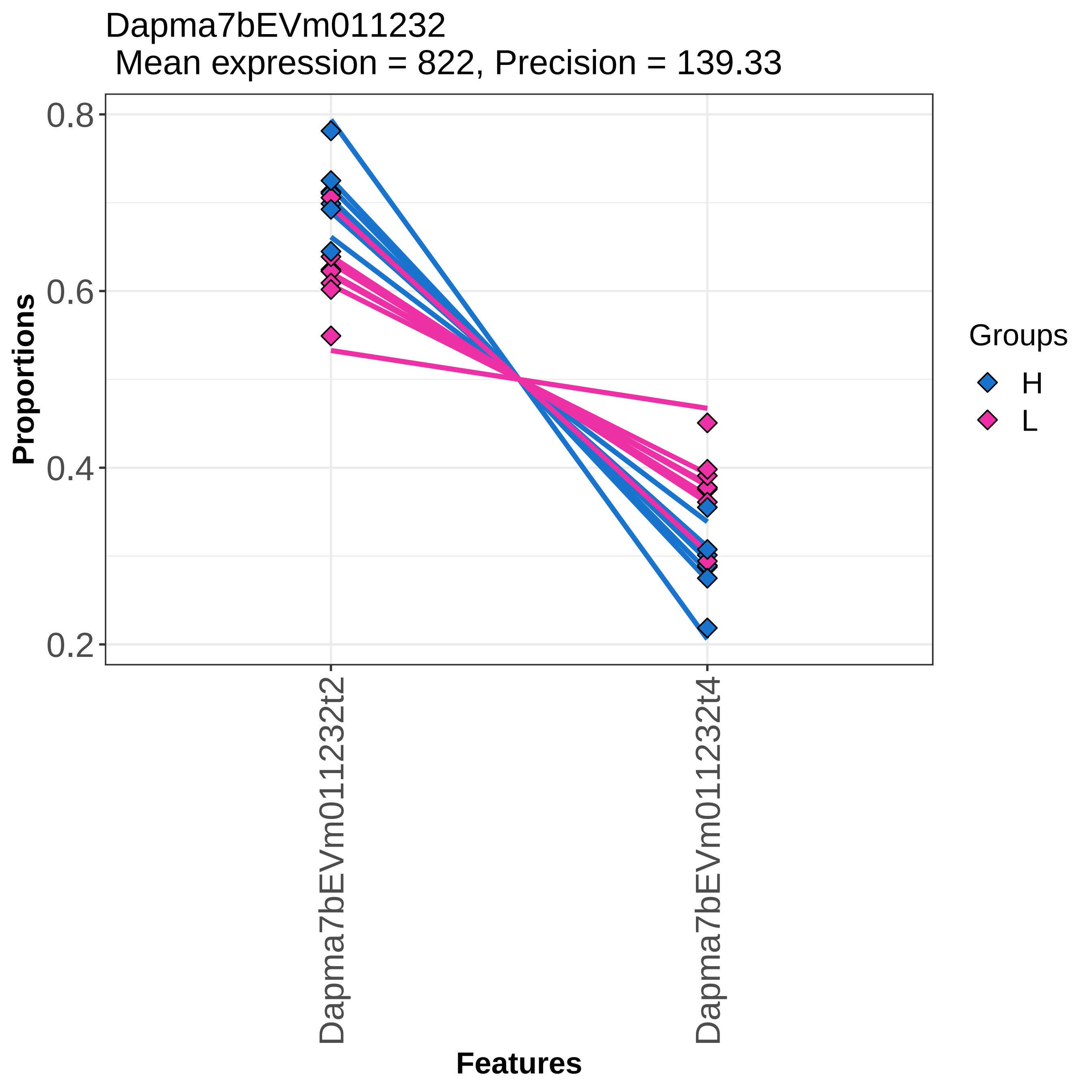
Dapma7bEVm011232
Mean expression = 822, Precision = 139.33
Proportions
0.8
0.6
0.4
0.2
Features
Dapma7bEVm011232t2
Dapma7bEVm011232t4
Groups
H
L

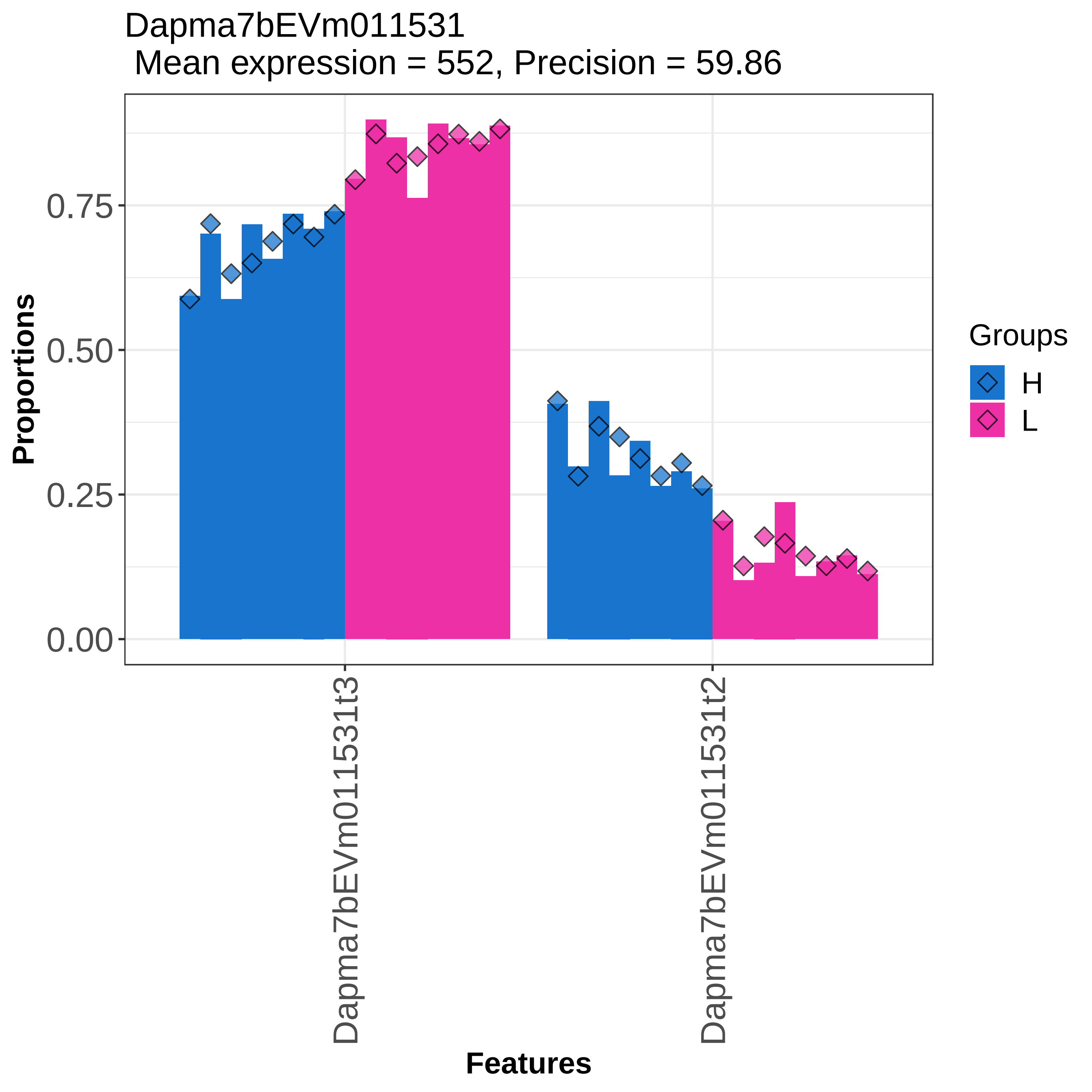

Dapma7bEVm011531
Mean expression = 552, Precision = 59.86
Proportions
0.00
0.25
0.50
0.75
Dapma7bEVm011531t3
Dapma7bEVm011531t2
Features
Groups
H
L

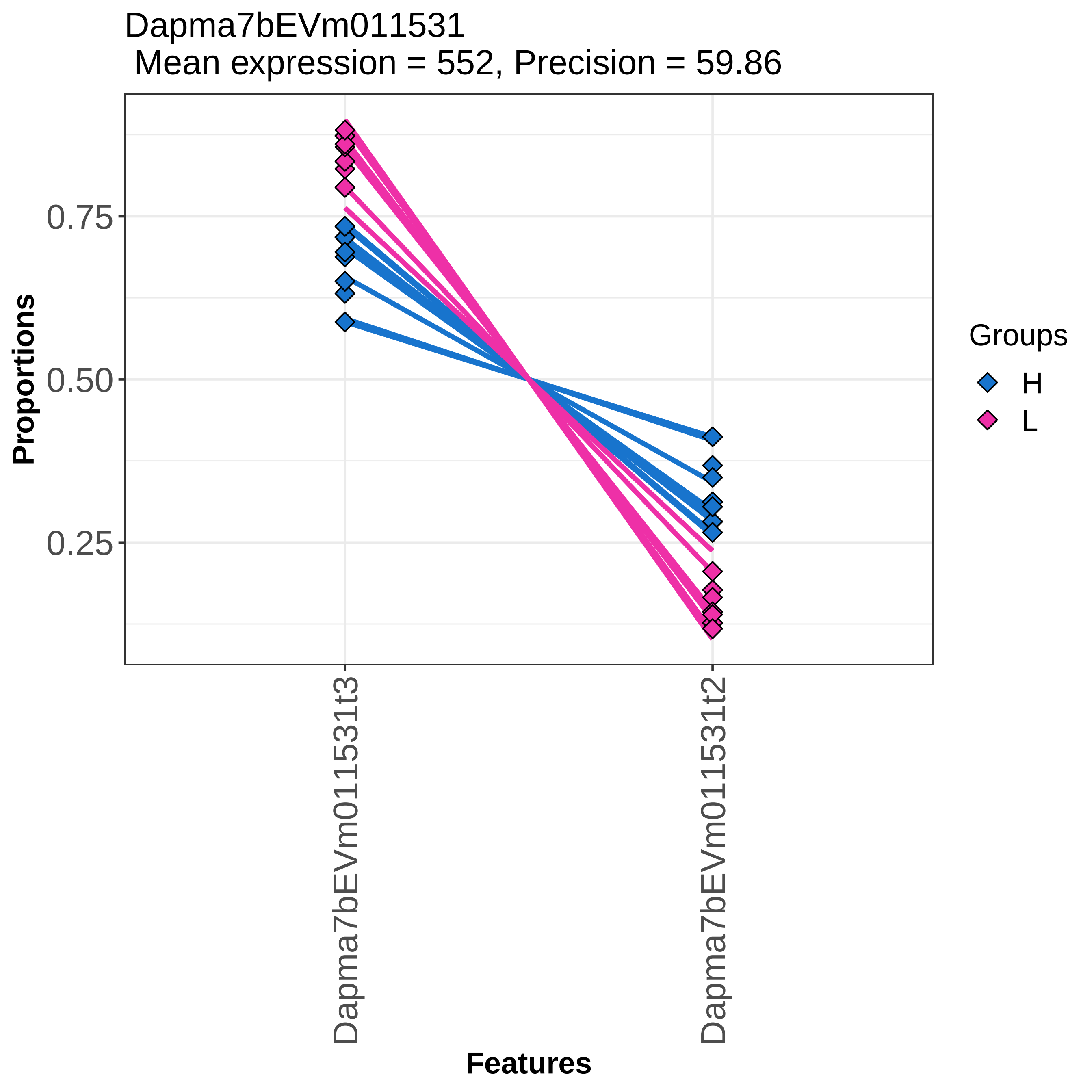
Dapma7bEVm011531
Mean expression = 552, Precision = 59.86
Proportions
0.75
0.50
0.25
Dapma7bEVm011531t3
Dapma7bEVm011531t2
Features
Groups
H
L

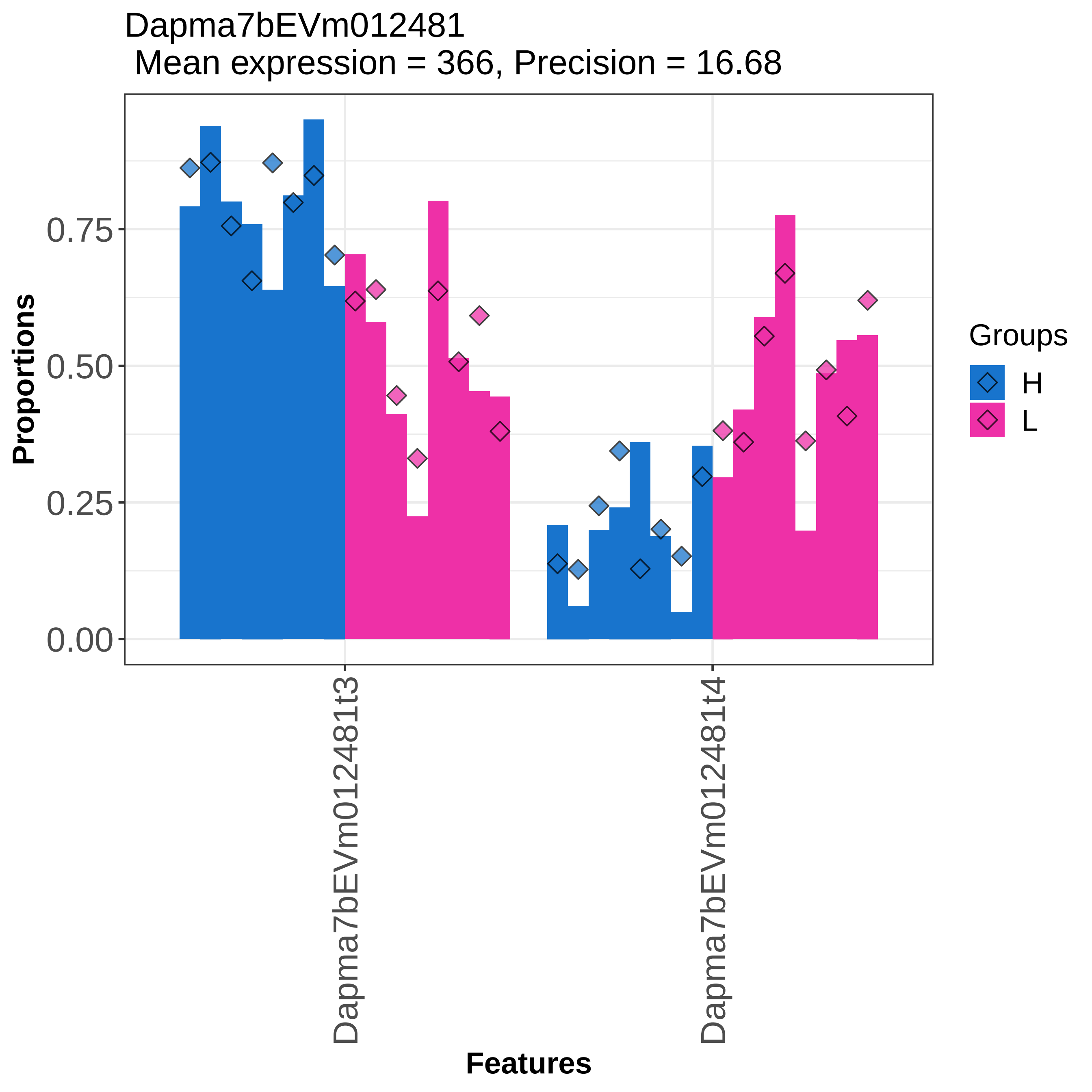
Dapma7bEVm012481
Mean expression = 366, Precision = 16.68
Proportions
0.00
0.25
0.50
0.75
Dapma7bEVm012481t3
Dapma7bEVm012481t4
Features
Groups
H
L

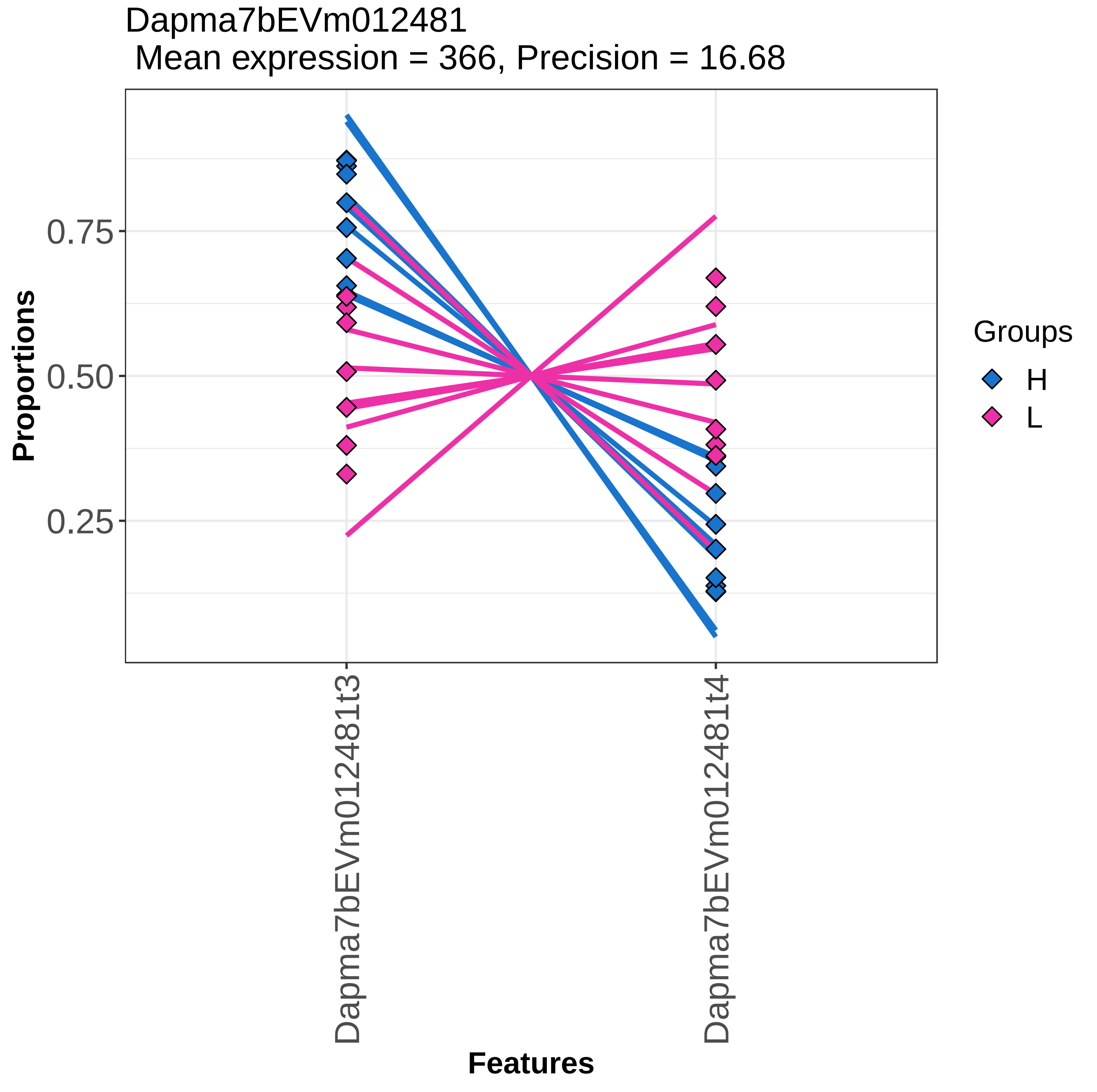
Dapma7bEVm012481
Mean expression = 366, Precision = 16.68
Proportions
0.75
0.50
0.25
Dapma7bEVm012481t3
Dapma7bEVm012481t4
Features
Groups
H
L

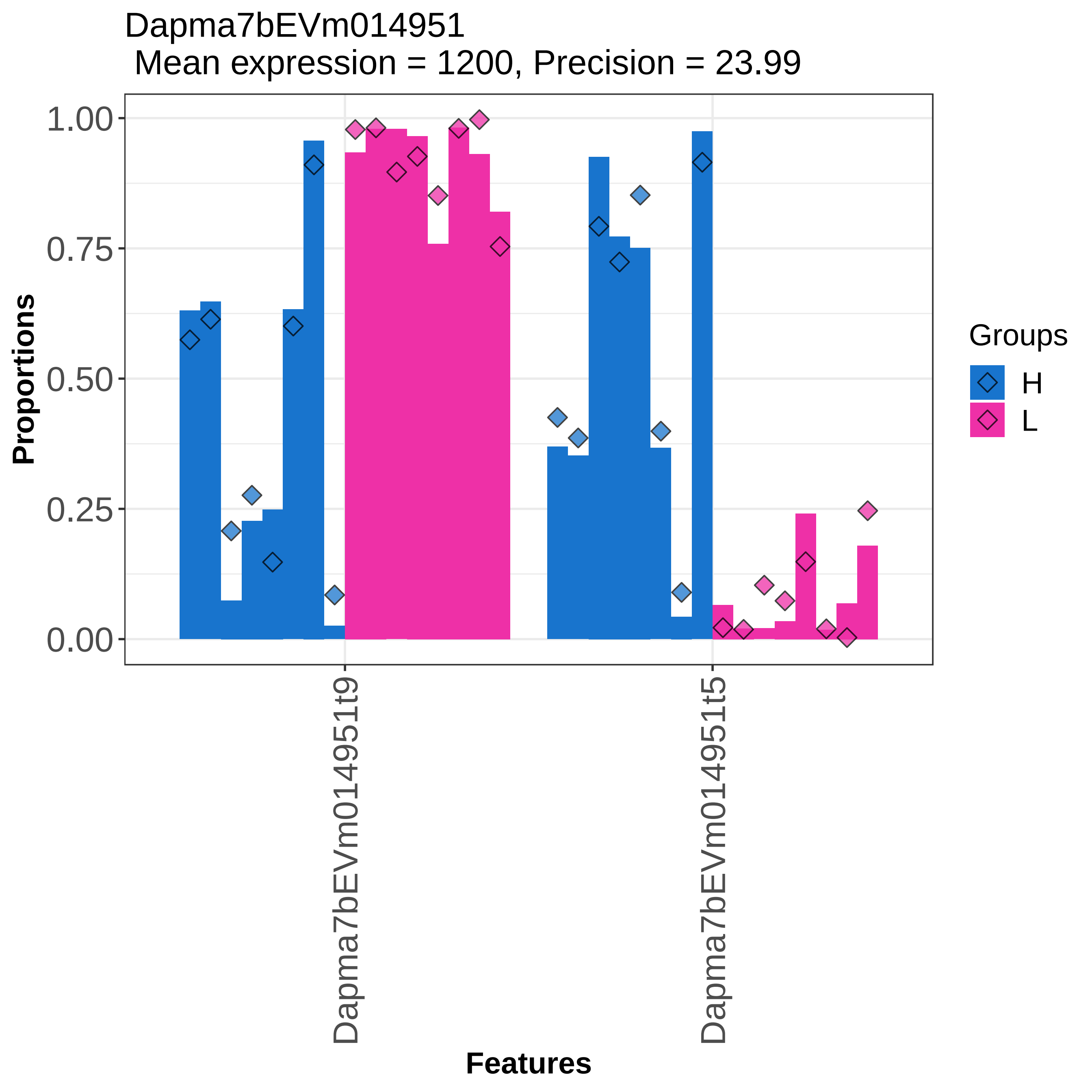
Dapma7bEVm014951
Mean expression = 1200, Precision = 23.99
Proportions
1.00
0.75
0.50
0.25
0.00
Dapma7bEVm014951t9
Dapma7bEVm014951t5
Features
Groups
H
L

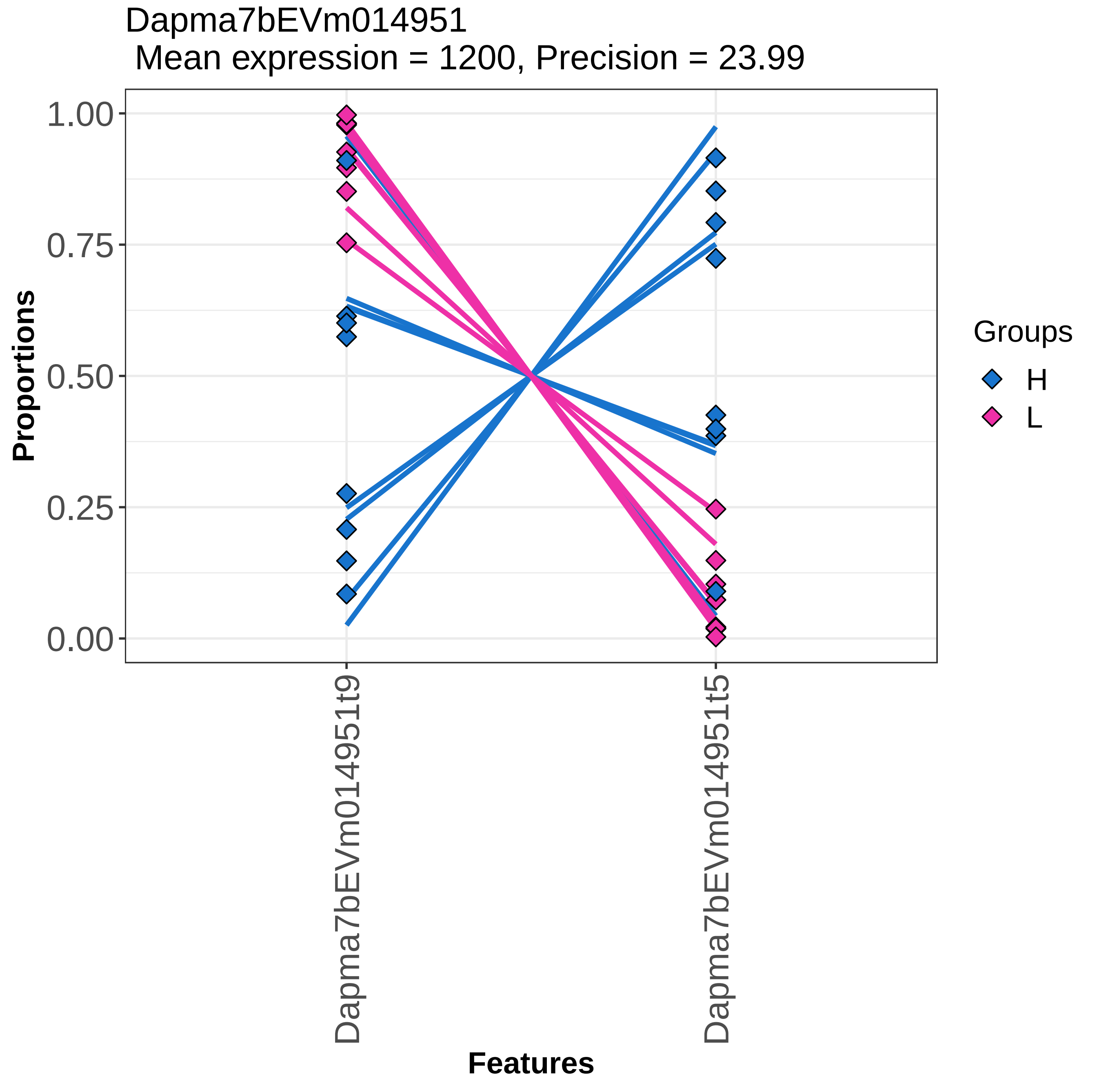
Dapma7bEVm014951
Mean expression = 1200, Precision = 23.99
Proportions
1.00
0.75
0.50
0.25
0.00
Dapma7bEVm014951t9
Dapma7bEVm014951t5
Features
Groups
H
L

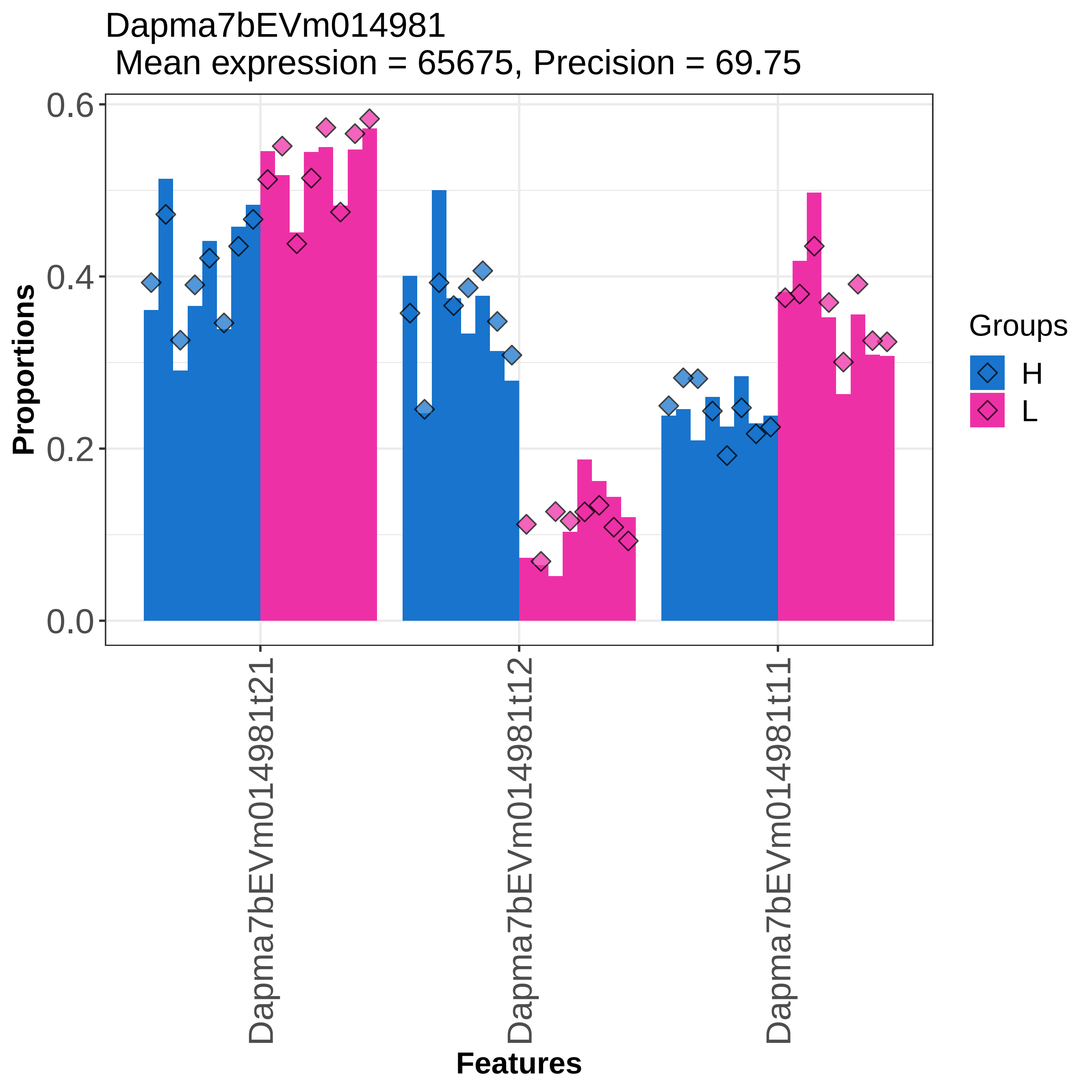
Dapma7bEVm014981
Mean expression = 65675, Precision = 69.75
Proportions
0.0
0.2
0.4
0.6
Dapma7bEVm014981t21
Dapma7bEVm014981t12
Dapma7bEVm014981t11
Features
Groups
H
L

Dapma7bEVm014981
Mean expression = 65675, Precision = 69.75

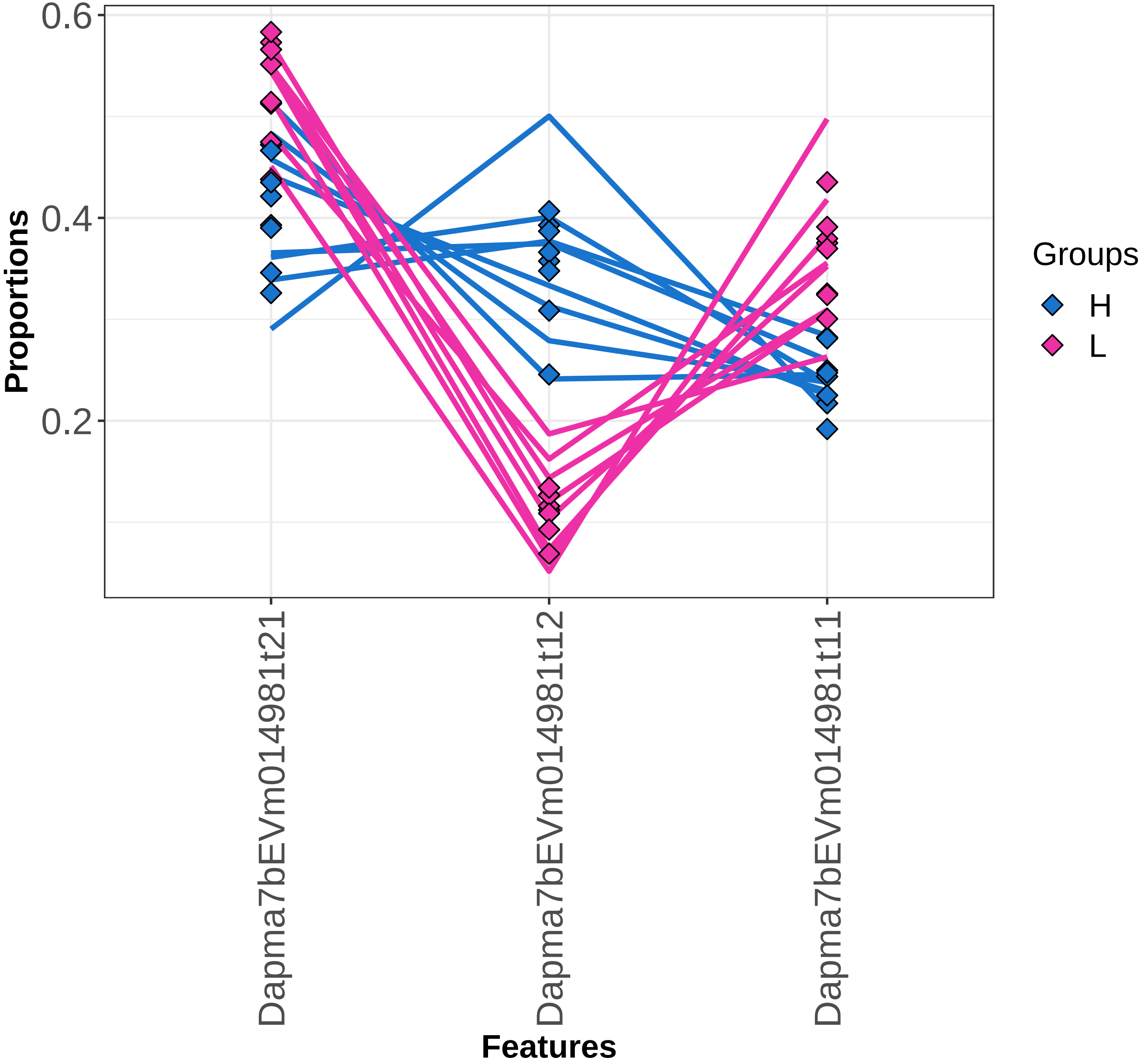

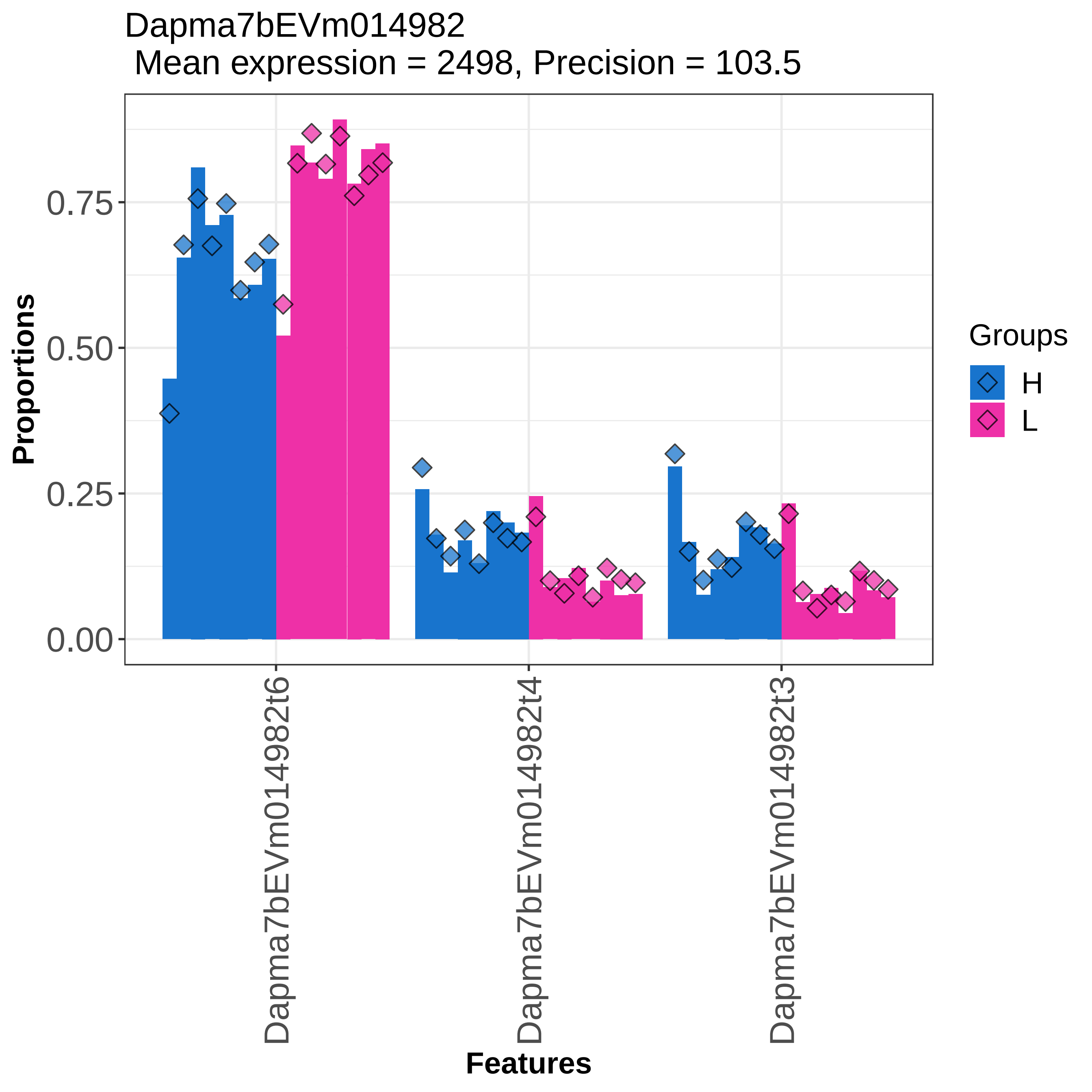
Dapma7bEVm014982
Mean expression = 2498, Precision = 103.5
Proportions
0.00
0.25
0.50
0.75
Dapma7bEVm014982t6
Dapma7bEVm014982t4
Dapma7bEVm014982t3
Features
Groups
H
L

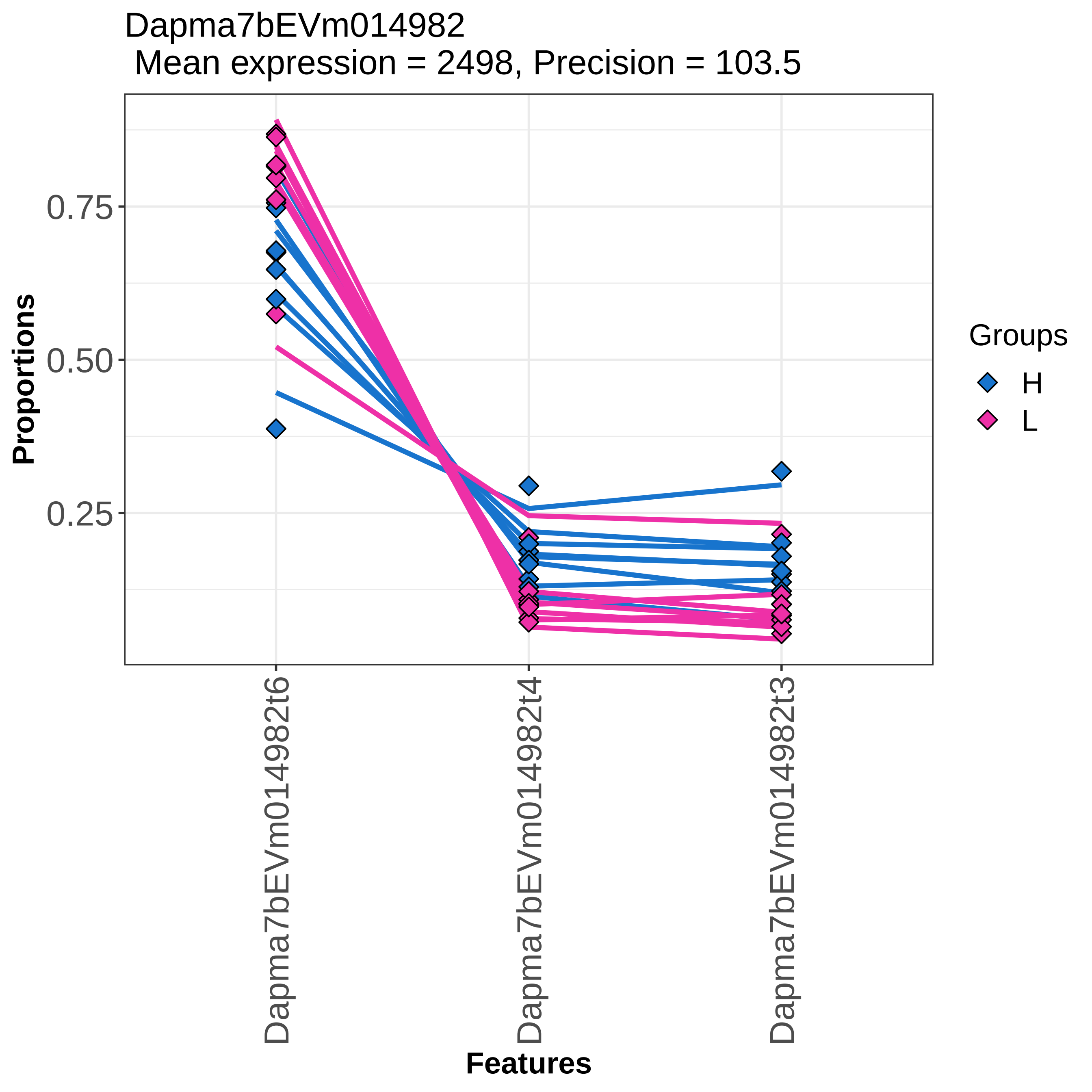

Dapma7bEVm014982
Mean expression = 2498, Precision = 103.5
Proportions
0.75
0.50
0.25
Features
Dapma7bEVm014982t6
Dapma7bEVm014982t4
Dapma7bEVm014982t3
Groups
H
L

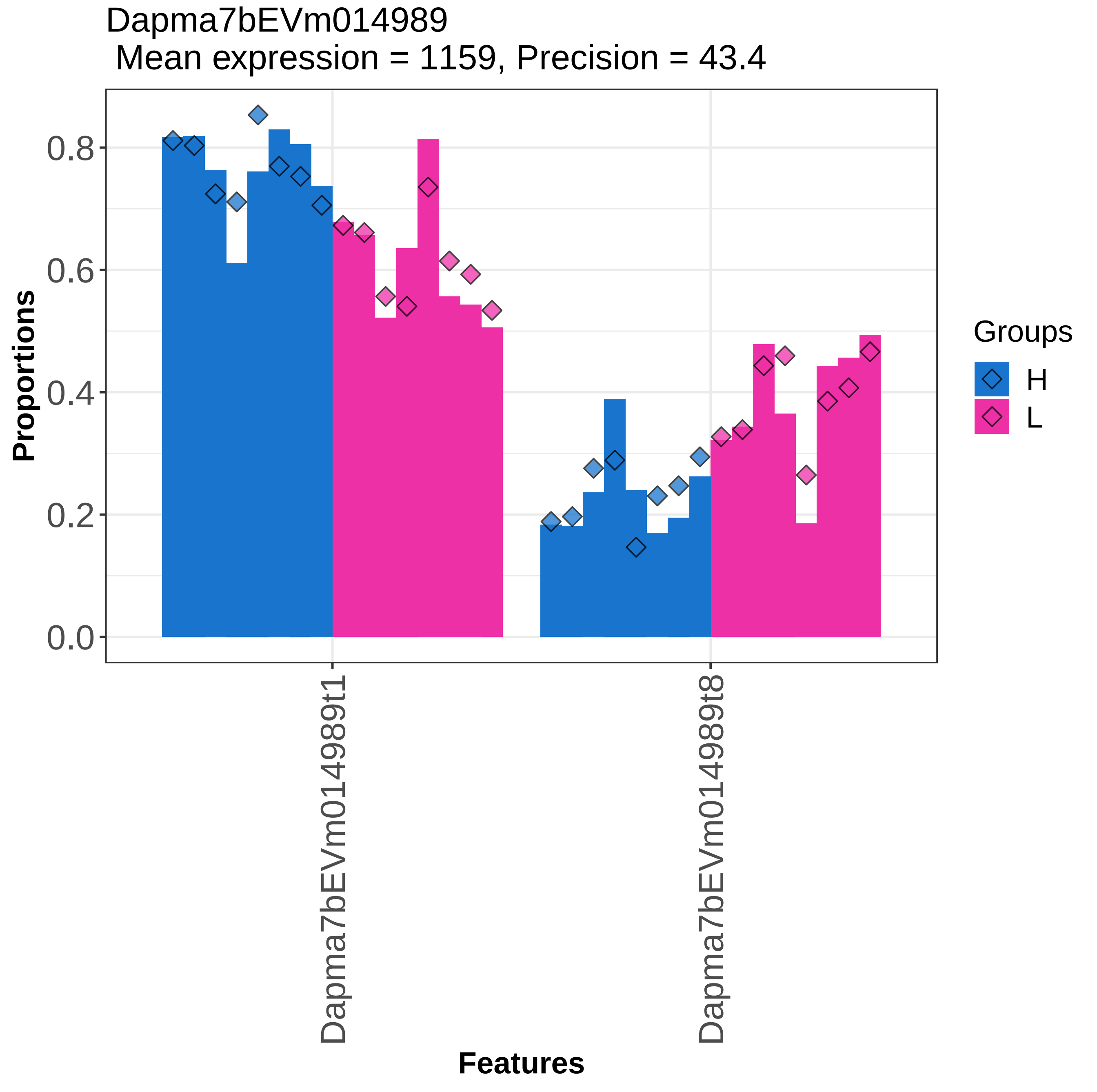

Dapma7bEVm014989
Mean expression = 1159, Precision = 43.4
Proportions
0.0
0.2
0.4
0.6
0.8
Features
Dapma7bEVm014989t1
Dapma7bEVm014989t8
Groups
H
L

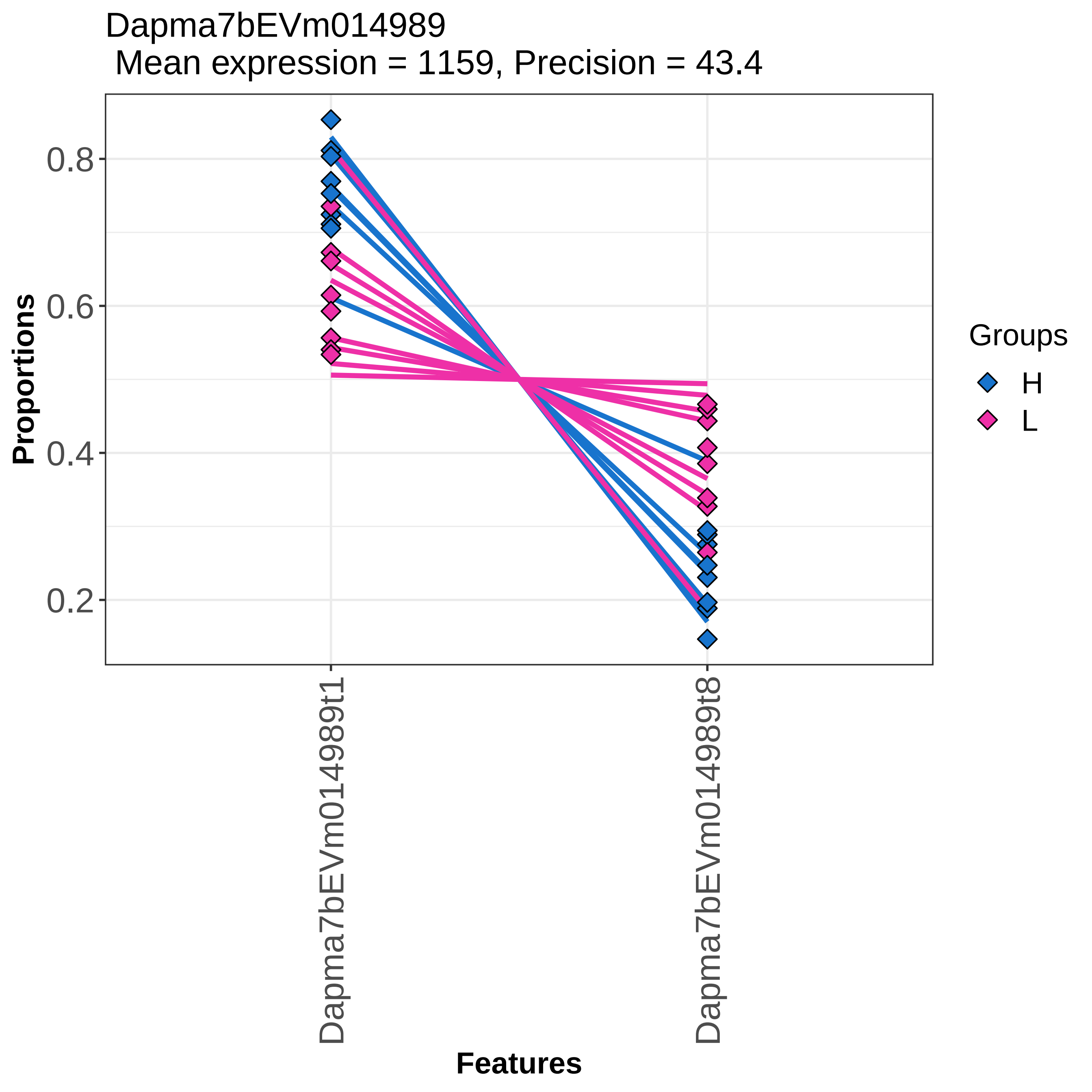

Dapma7bEVm014989
Mean expression = 1159, Precision = 43.4
Proportions
0.8
0.6
0.4
0.2
Dapma7bEVm014989t1
Dapma7bEVm014989t8
Features
Groups
H
L

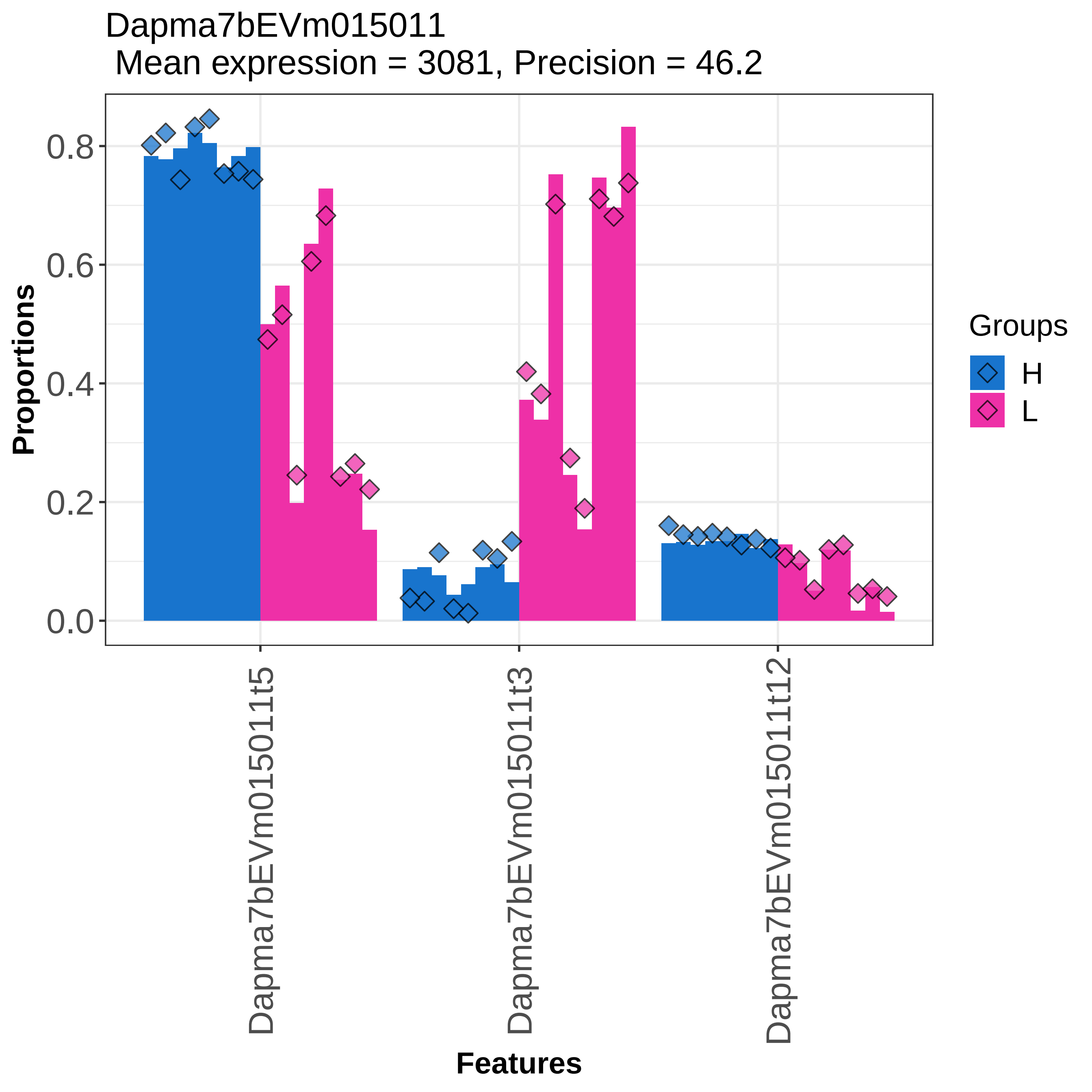
Dapma7bEVm015011
Mean expression = 3081, Precision = 46.2
Proportions
0.0
0.2
0.4
0.6
0.8
Dapma7bEVm015011t5
Dapma7bEVm015011t3
Dapma7bEVm015011t12
Features
Groups
H
L

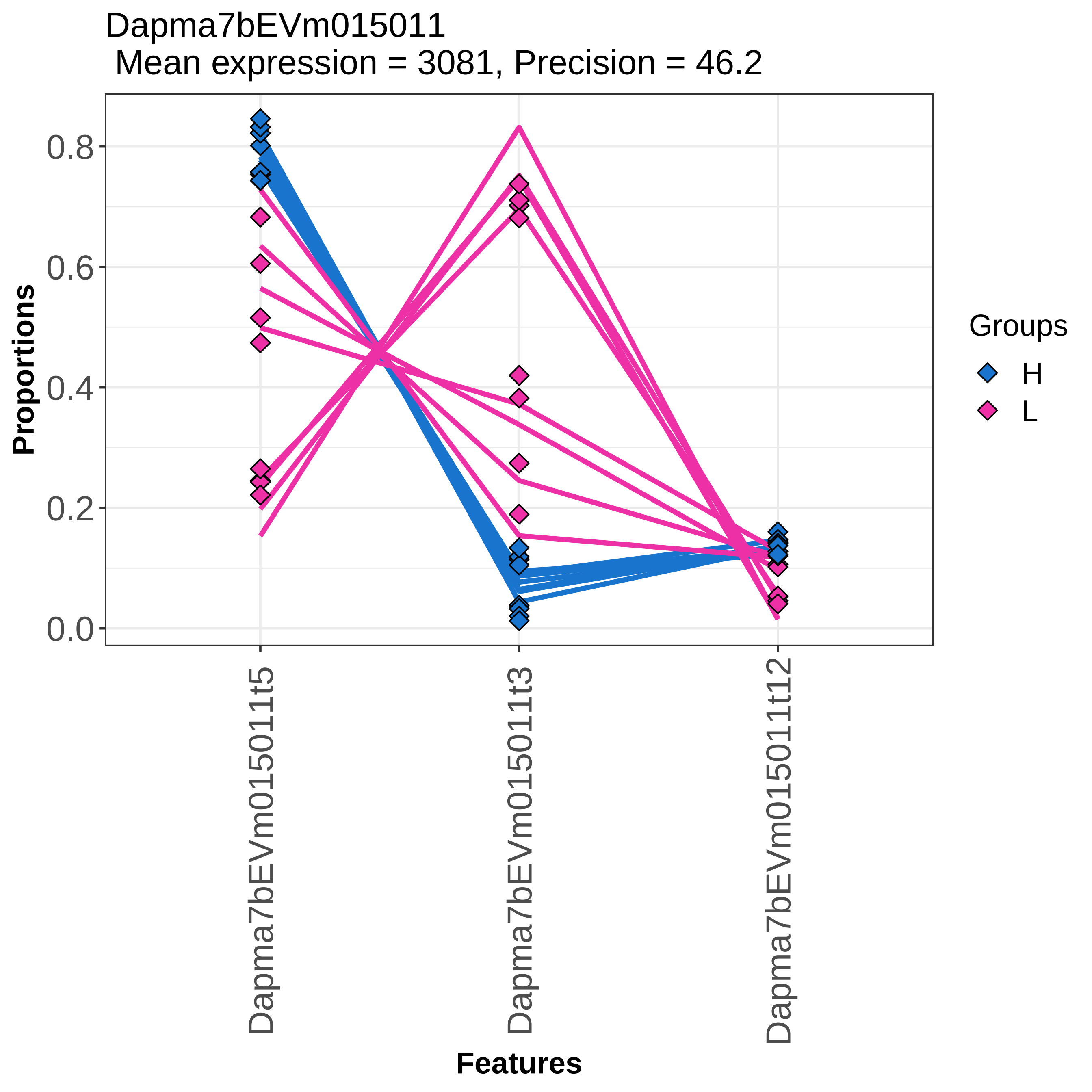

Dapma7bEVm015011
Mean expression = 3081, Precision = 46.2
Proportions
0.0
0.2
0.4
0.6
0.8
Features
Dapma7bEVm015011t5
Dapma7bEVm015011t3
Dapma7bEVm015011t12
Groups
H
L

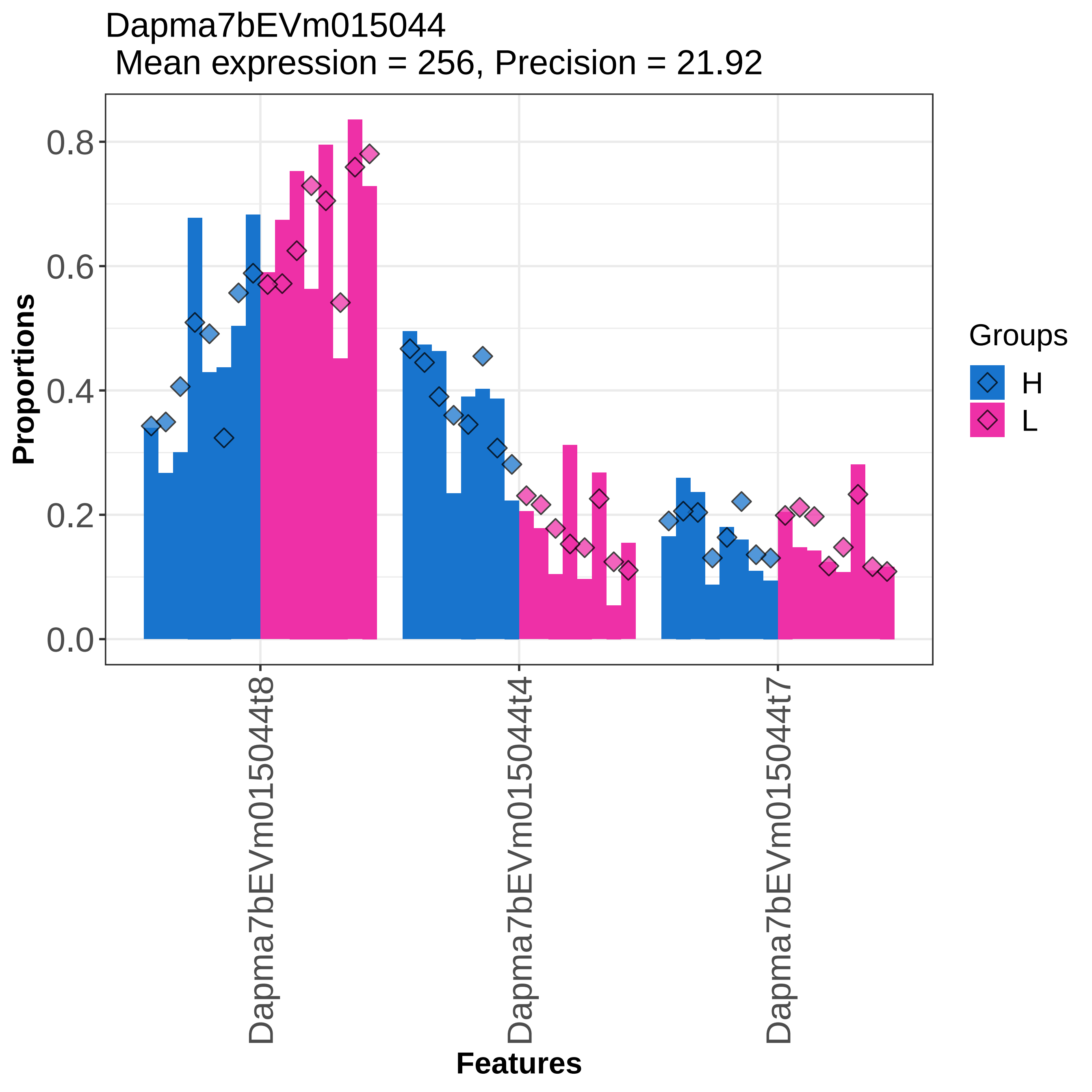
Dapma7bEVm015044
Mean expression = 256, Precision = 21.92
Proportions
0.0
0.2
0.4
0.6
0.8
Dapma7bEVm015044t8
Dapma7bEVm015044t4
Dapma7bEVm015044t7
Features
Groups
H
L

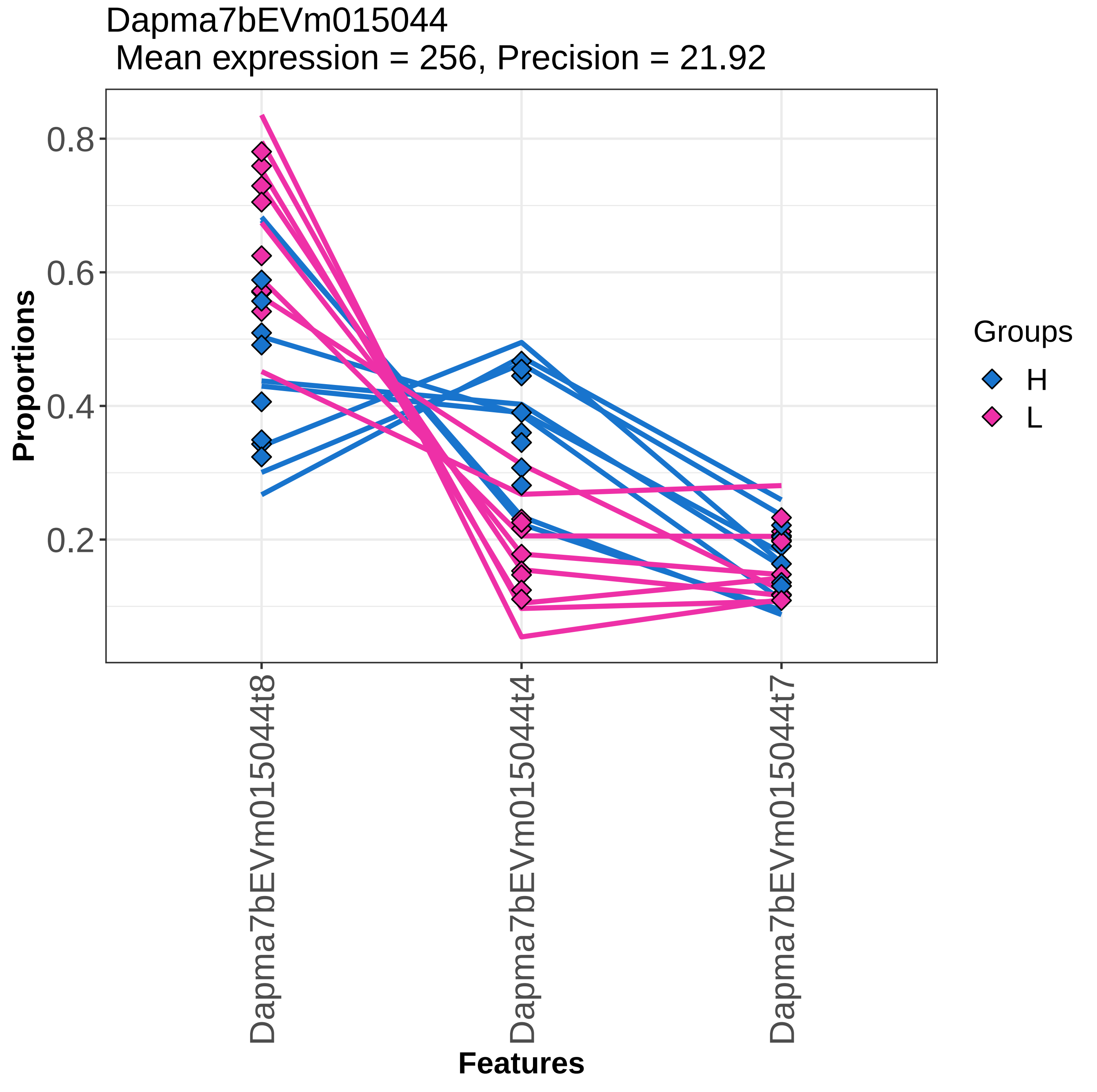

Dapma7bEVm015044
Mean expression = 256, Precision = 21.92
Proportions
0.8
0.6
0.4
0.2
Features
Dapma7bEVm015044t8
Dapma7bEVm015044t4
Dapma7bEVm015044t7
Groups
H
L

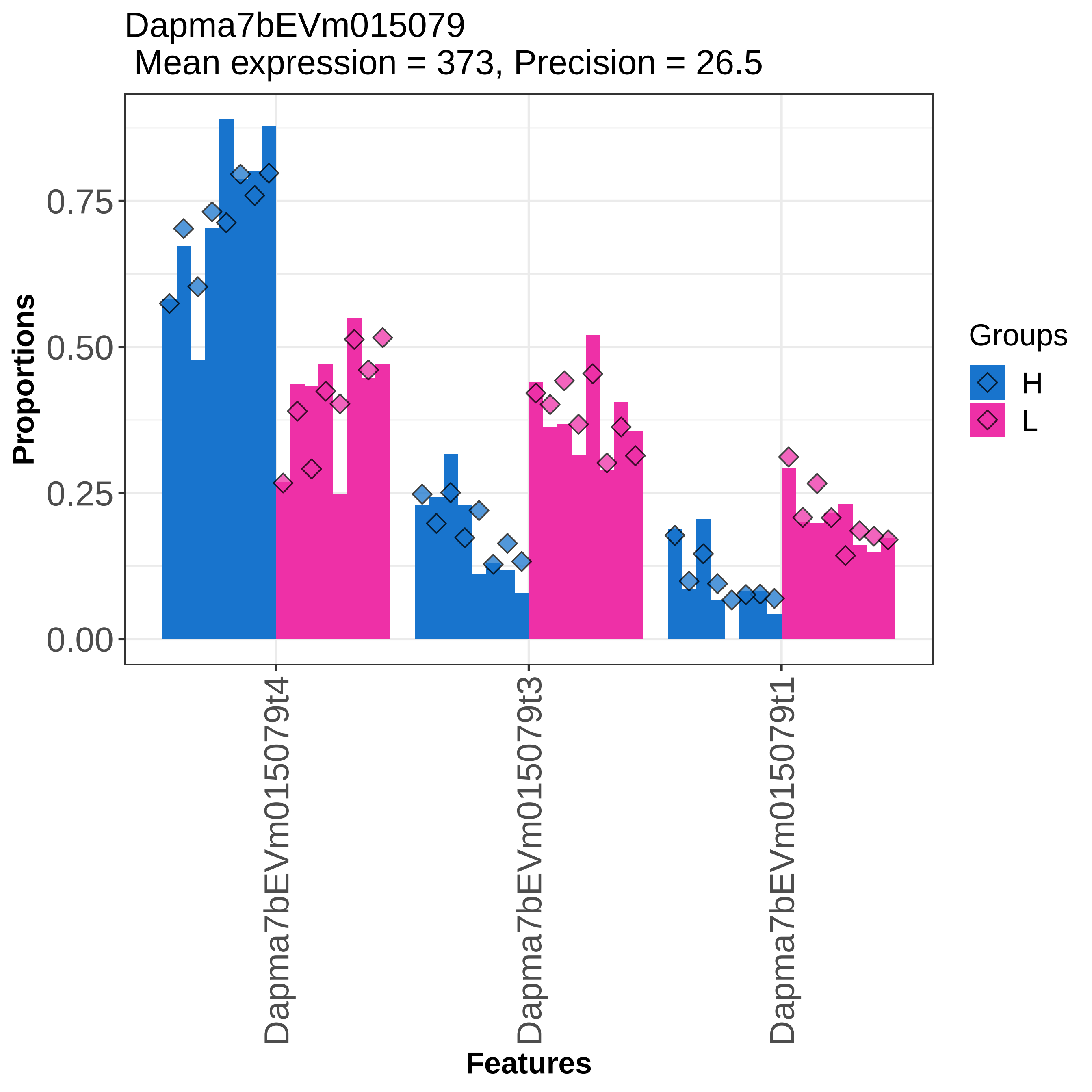
Dapma7bEVm015079
Mean expression = 373, Precision = 26.5
Proportions
0.00
0.25
0.50
0.75
Dapma7bEVm015079t4
Dapma7bEVm015079t3
Dapma7bEVm015079t1
Features
Groups
H
L

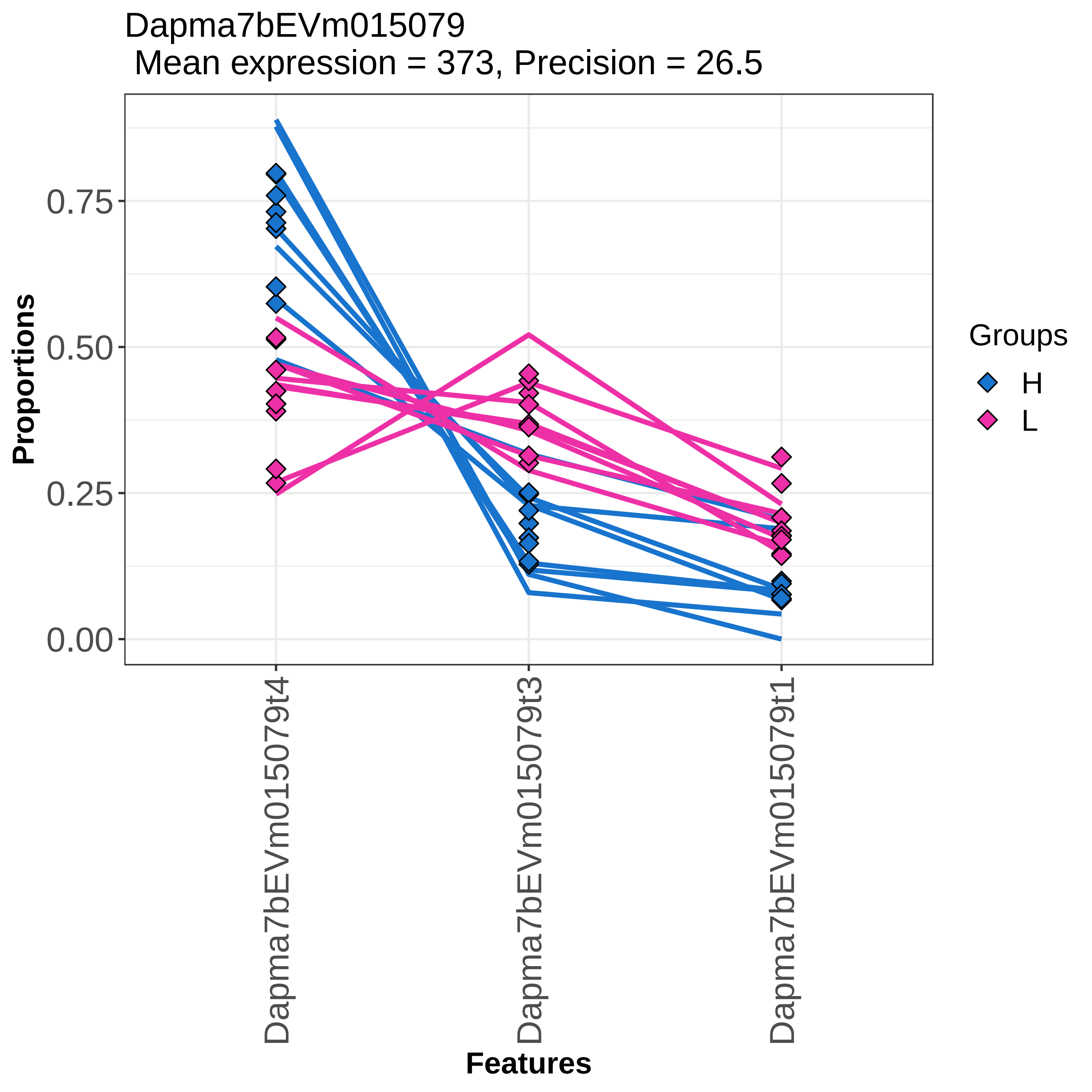
Dapma7bEVm015079
Mean expression = 373, Precision = 26.5
Proportions
0.75
0.50
0.25
0.00
Dapma7bEVm015079t4
Dapma7bEVm015079t3
Dapma7bEVm015079t1
Features
Groups
H
L

Dapma7bEVm015096
Mean expression = 75, Precision = 9.9

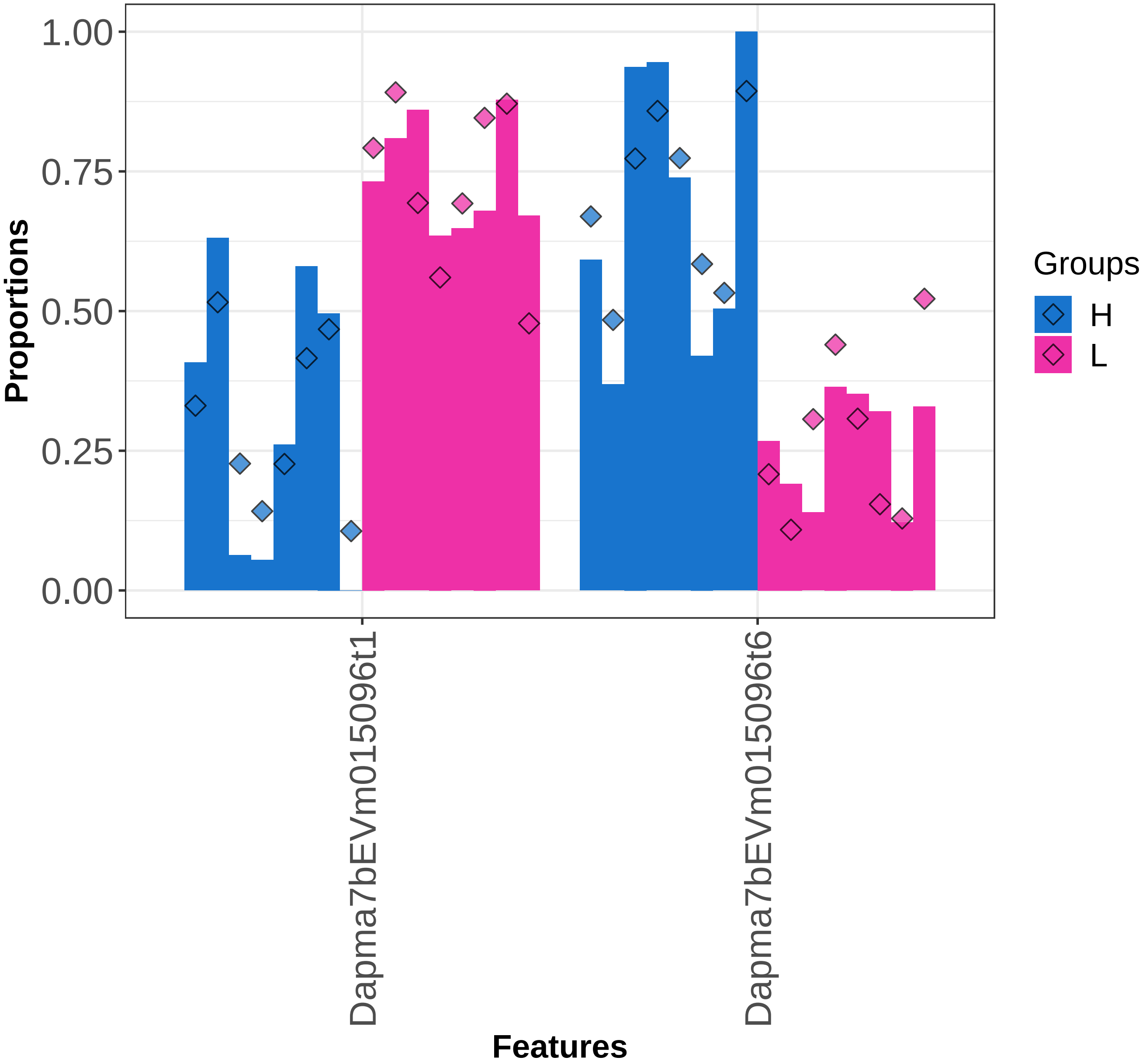

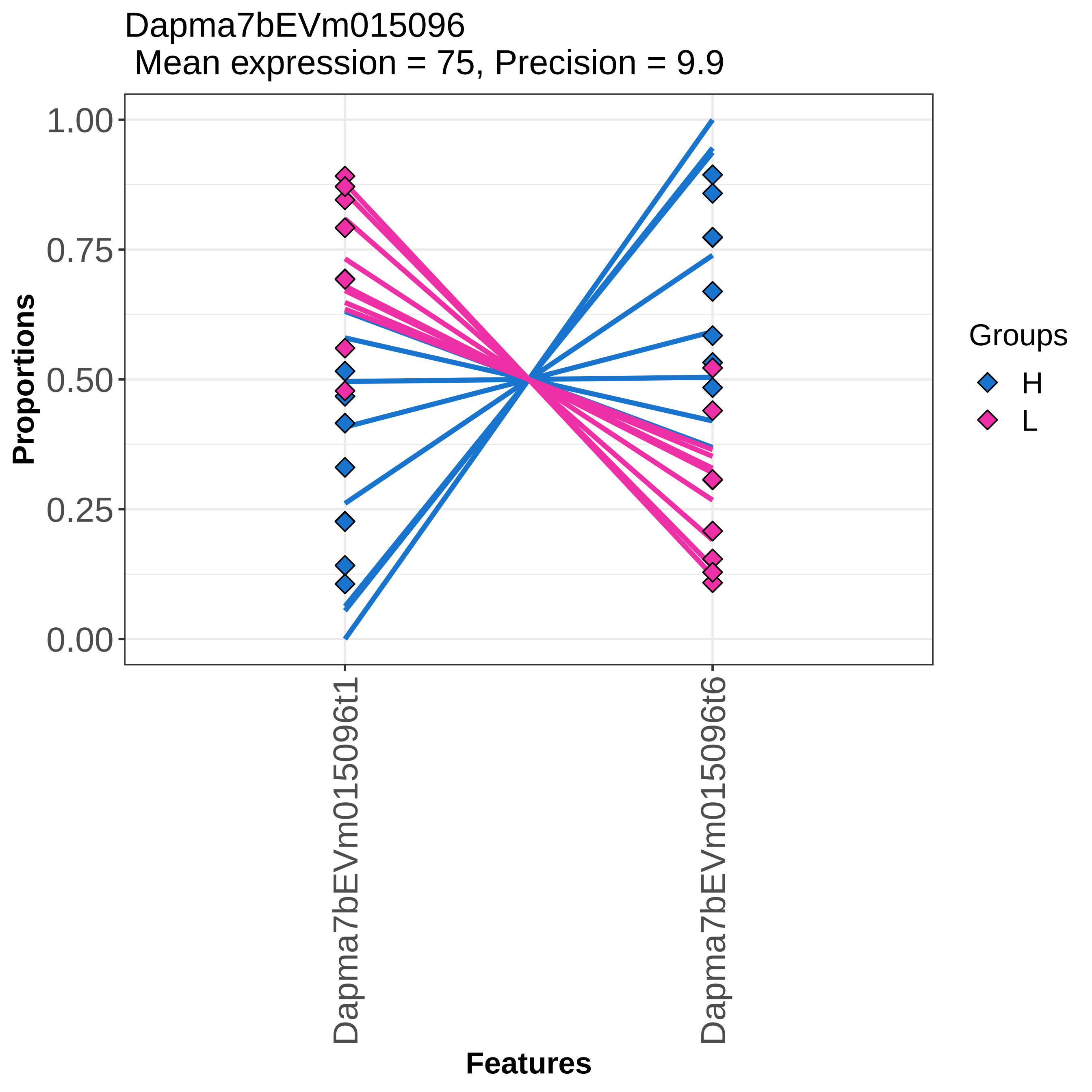

Dapma7bEVm015096
Mean expression = 75, Precision = 9.9
Proportions
1.00
0.75
0.50
0.25
0.00
Features
Dapma7bEVm015096t1
Dapma7bEVm015096t6
Groups
H
L

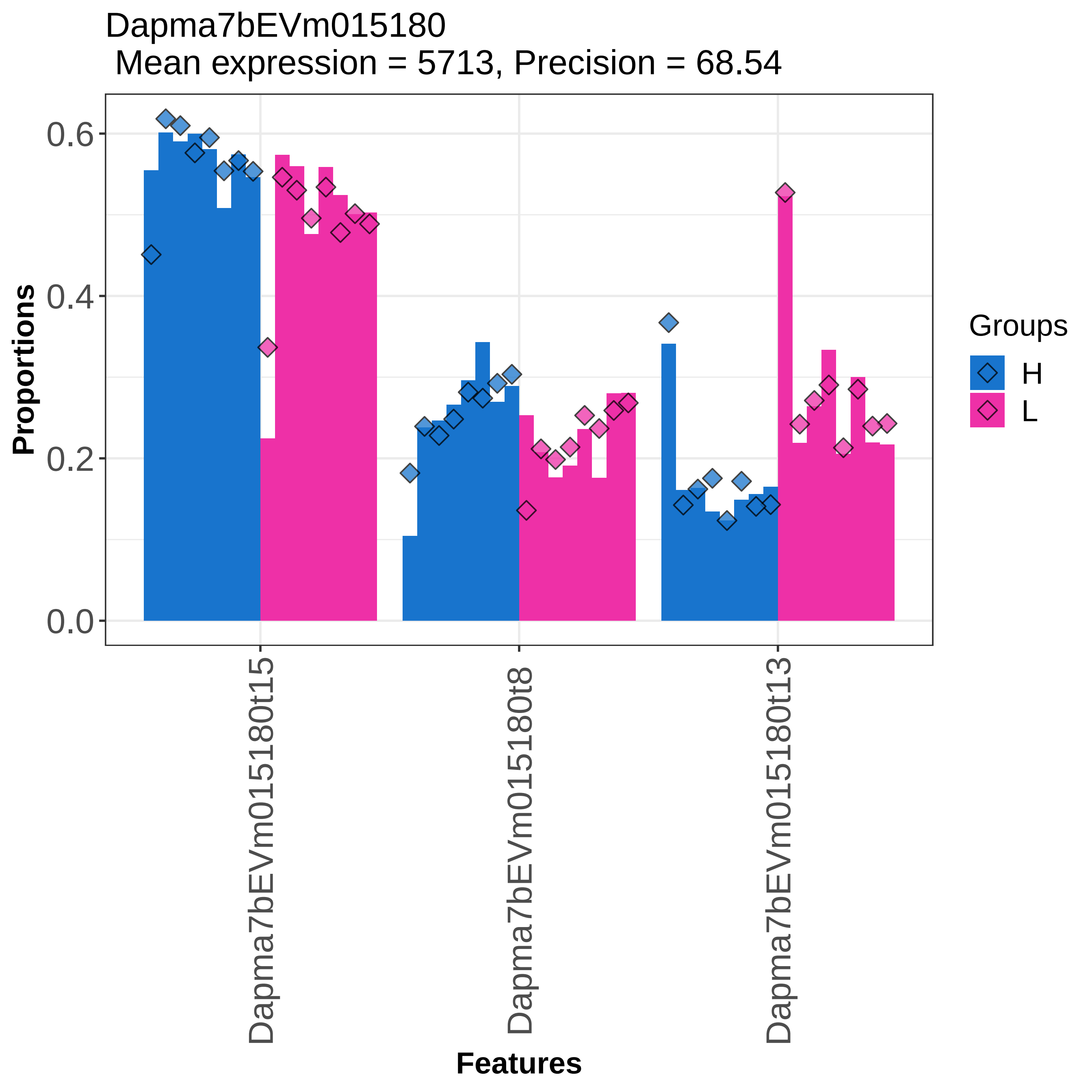

Dapma7bEVm015180
Mean expression = 5713, Precision = 68.54
Proportions
0.0
0.2
0.4
0.6
Dapma7bEVm015180t15
Dapma7bEVm015180t8
Dapma7bEVm015180t13
Features
Groups
H
L

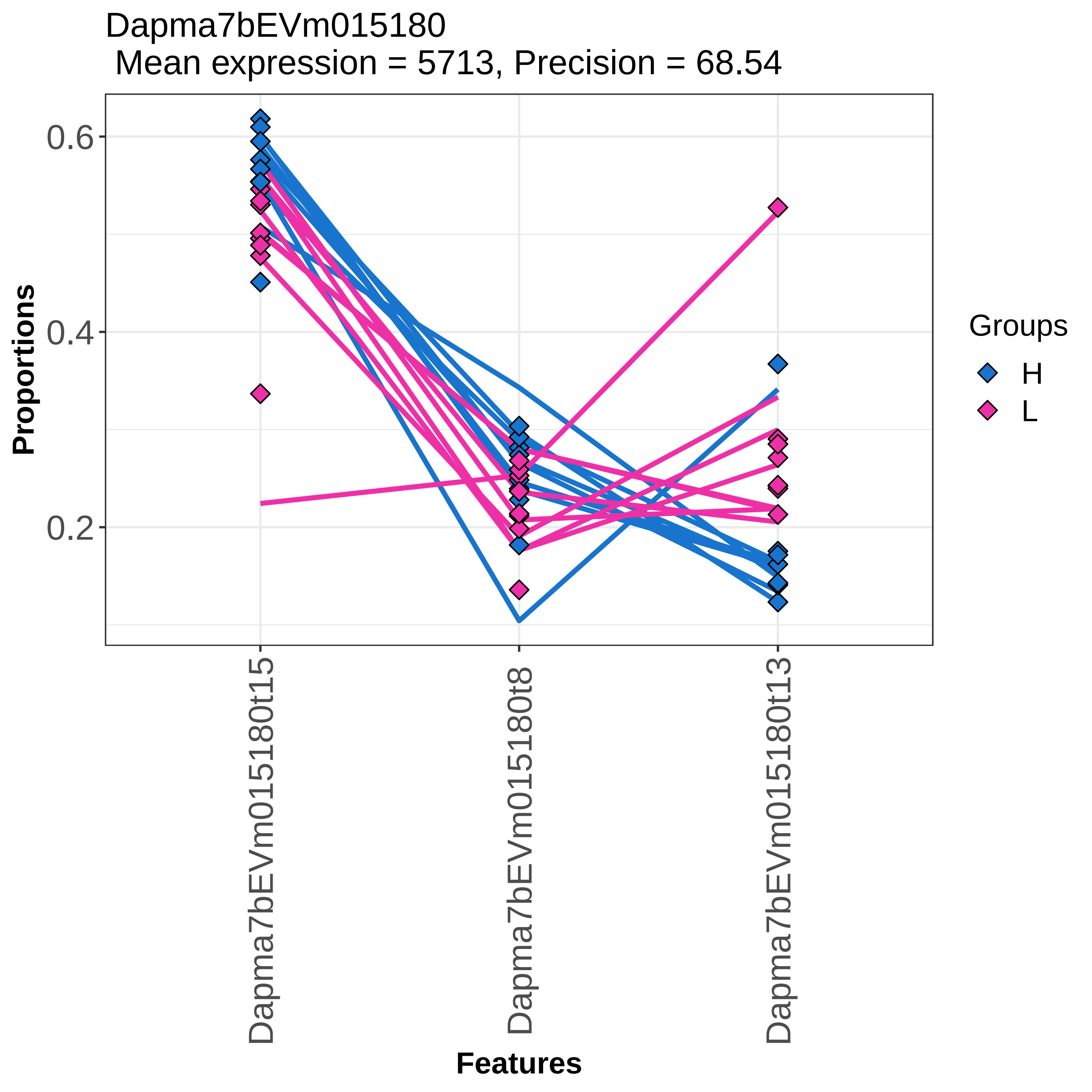
Dapma7bEVm015180
Mean expression = 5713, Precision = 68.54
Proportions
0.6
0.4
0.2
Dapma7bEVm015180t15
Dapma7bEVm015180t8
Dapma7bEVm015180t13
Features
Groups
H
L

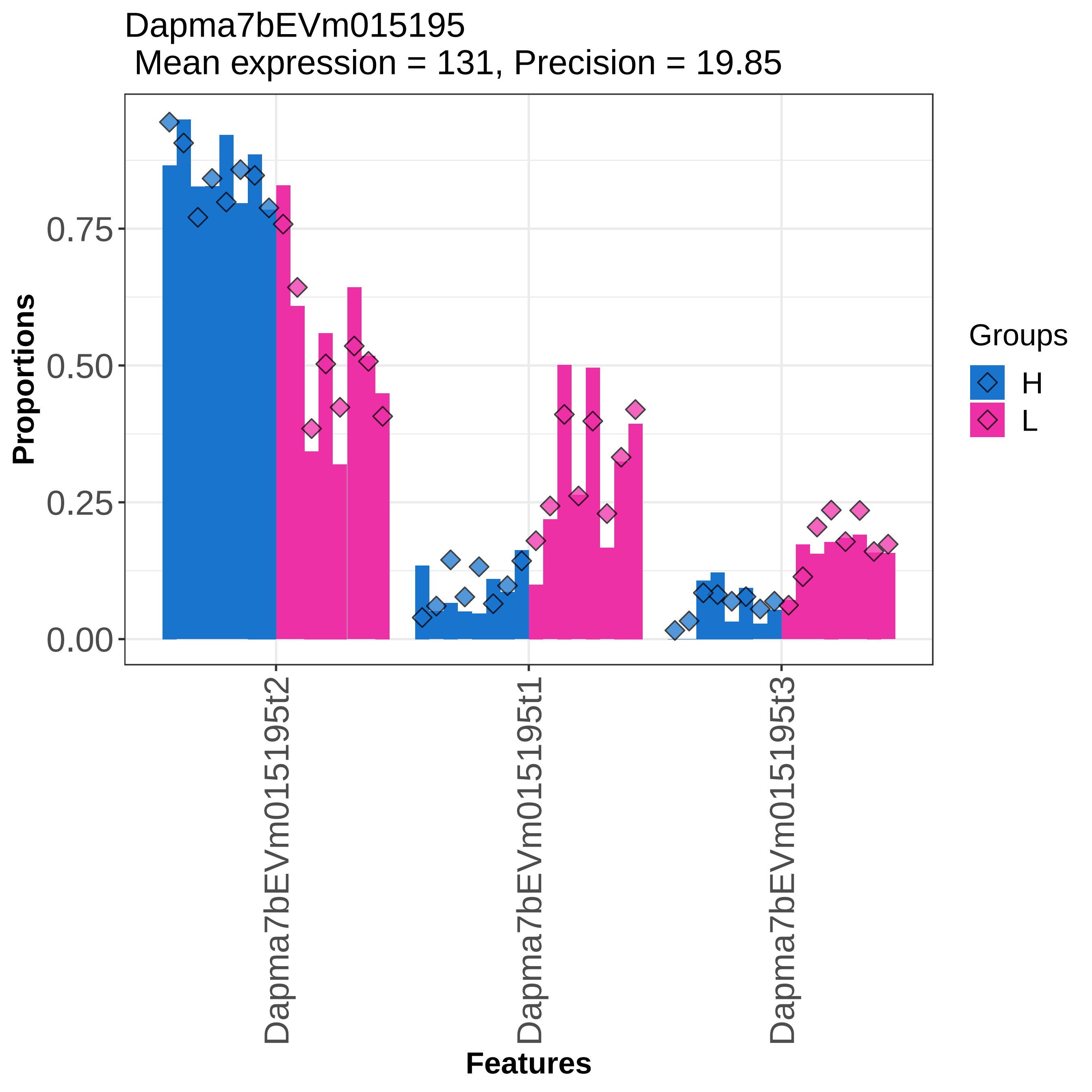

Dapma7bEVm015195
Mean expression = 131, Precision = 19.85
Proportions
0.00
0.25
0.50
0.75
Dapma7bEVm015195t2
Dapma7bEVm015195t1
Dapma7bEVm015195t3
Features
Groups
H
L

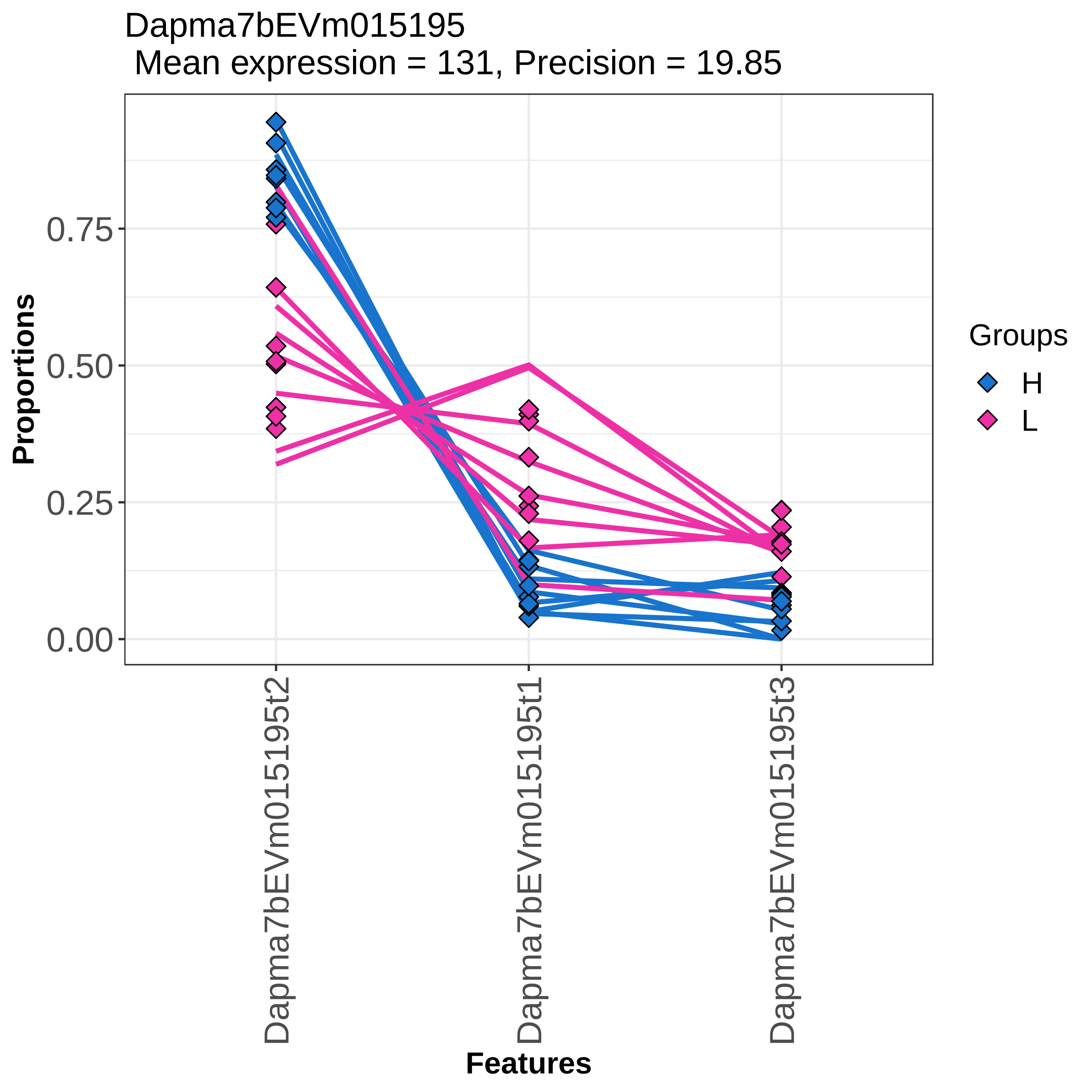

Dapma7bEVm015195
Mean expression = 131, Precision = 19.85
Proportions
0.00
0.25
0.50
0.75
Features
Dapma7bEVm015195t2
Dapma7bEVm015195t1
Dapma7bEVm015195t3
Groups
H
L

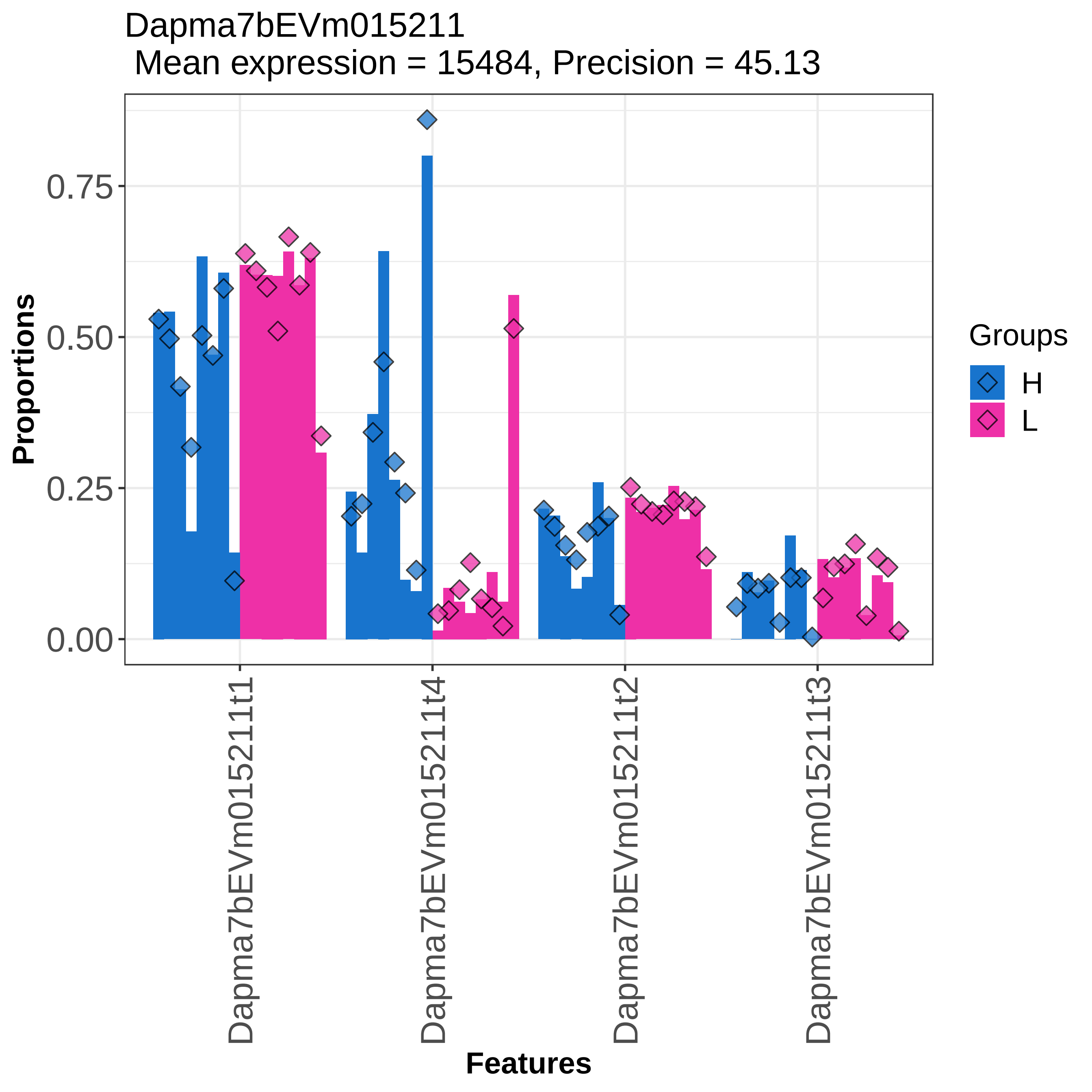

Dapma7bEVm015211
Mean expression = 15484, Precision = 45.13
Proportions
0.00
0.25
0.50
0.75
Features
Dapma7bEVm015211t1
Dapma7bEVm015211t4
Dapma7bEVm015211t2
Dapma7bEVm015211t3
Groups
H
L

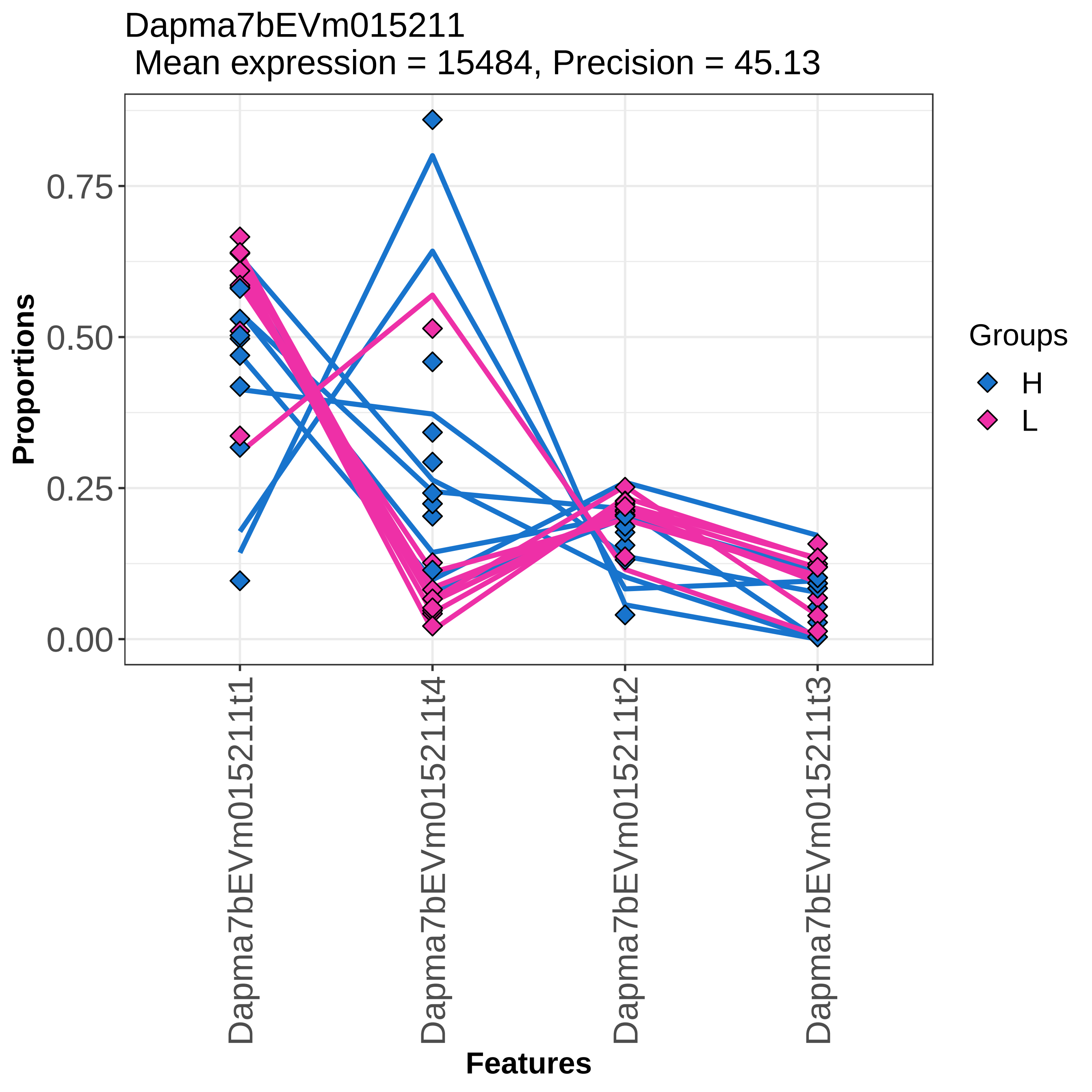

Dapma7bEVm015211
Mean expression = 15484, Precision = 45.13
Proportions
0.00
0.25
0.50
0.75
Features
Dapma7bEVm015211t1
Dapma7bEVm015211t4
Dapma7bEVm015211t2
Dapma7bEVm015211t3
Groups
H
L

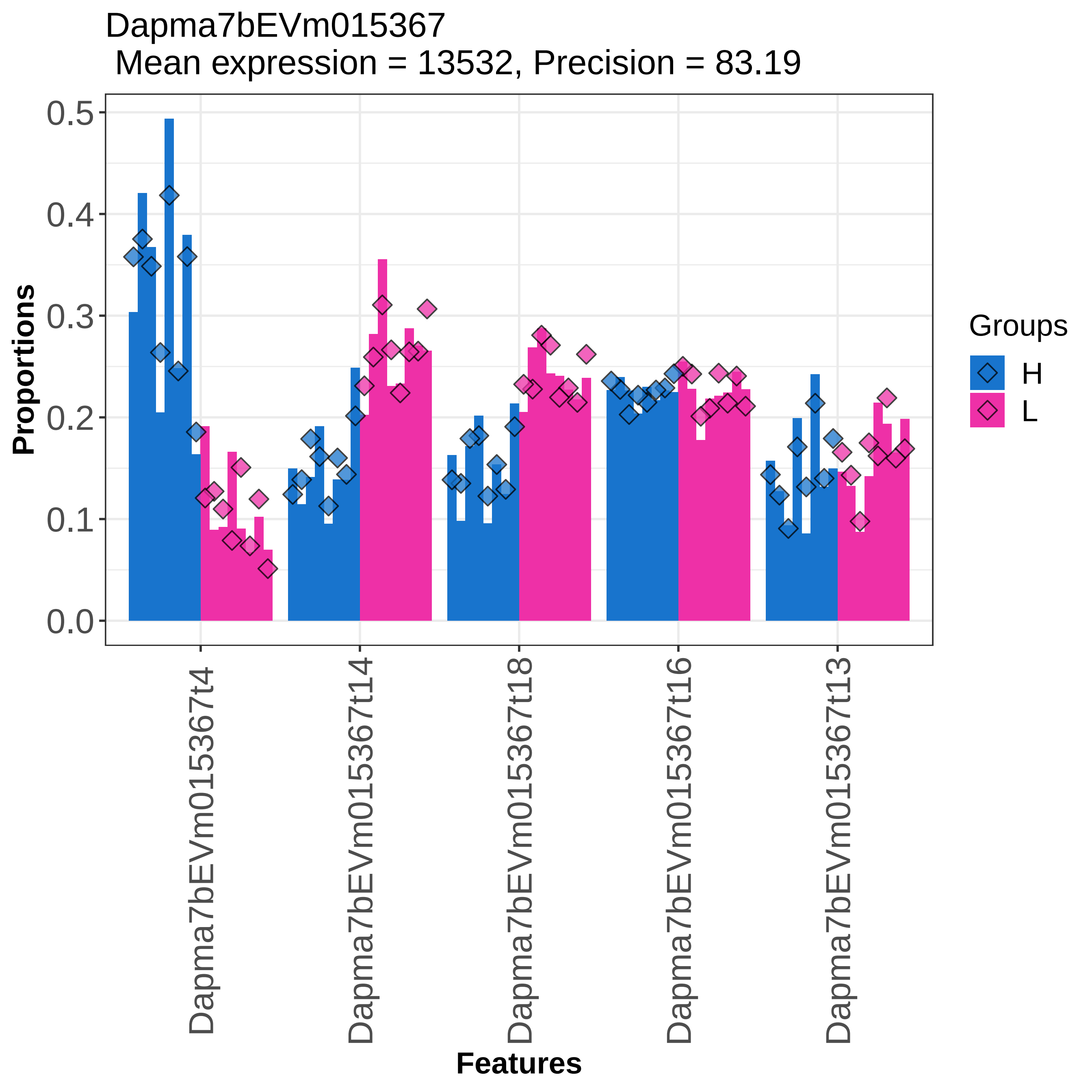

Dapma7bEVm015367
Mean expression = 13532, Precision = 83.19
Proportions
0.0
0.1
0.2
0.3
0.4
0.5
Dapma7bEVm015367t4
Dapma7bEVm015367t14
Dapma7bEVm015367t18
Dapma7bEVm015367t16
Dapma7bEVm015367t13
Features
Groups
H
L

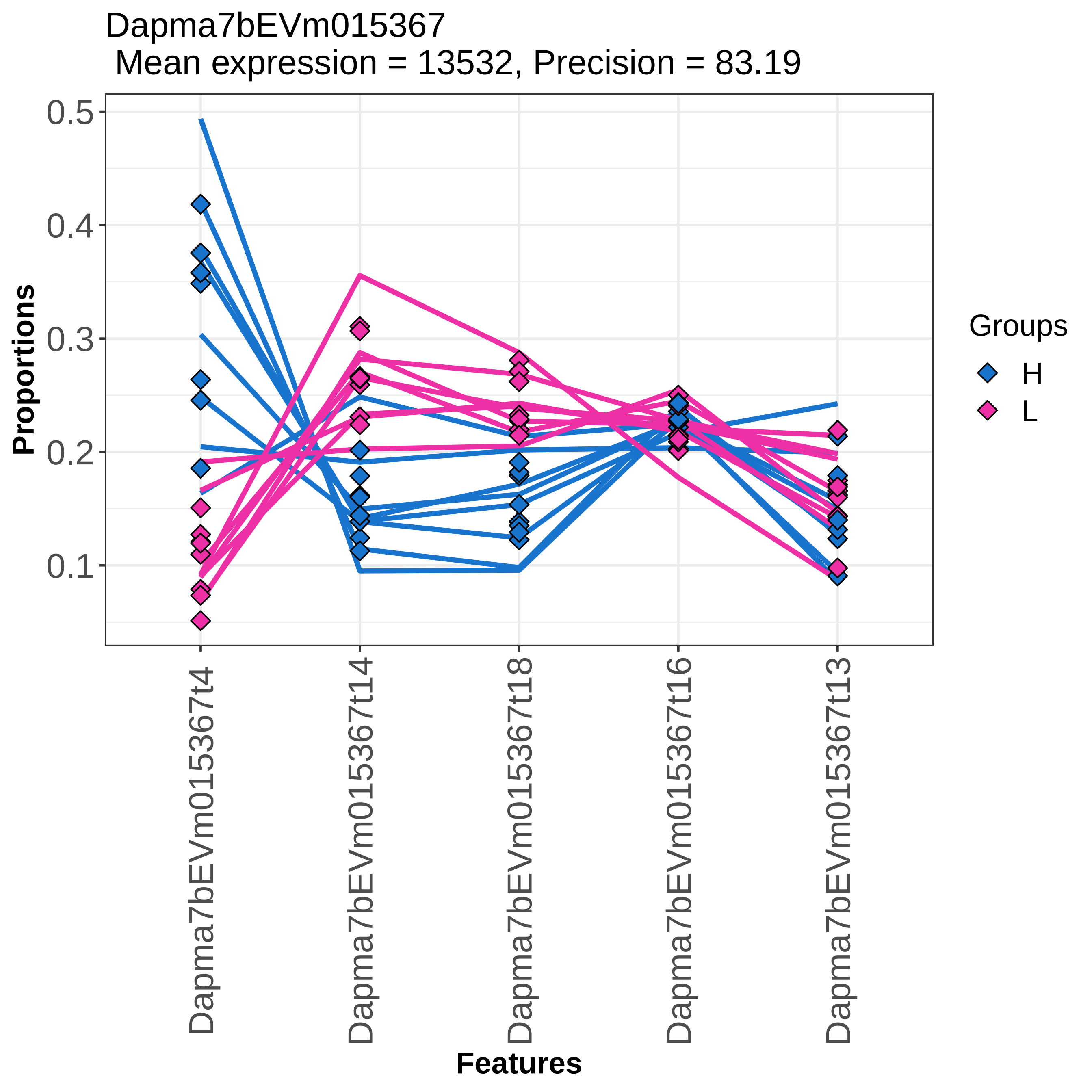

Dapma7bEVm015367
Mean expression = 13532, Precision = 83.19
Proportions
0.5
0.4
0.3
0.2
0.1
Groups
H
L
Dapma7bEVm015367t4
Dapma7bEVm015367t14
Dapma7bEVm015367t18
Dapma7bEVm015367t16
Dapma7bEVm015367t13
Features

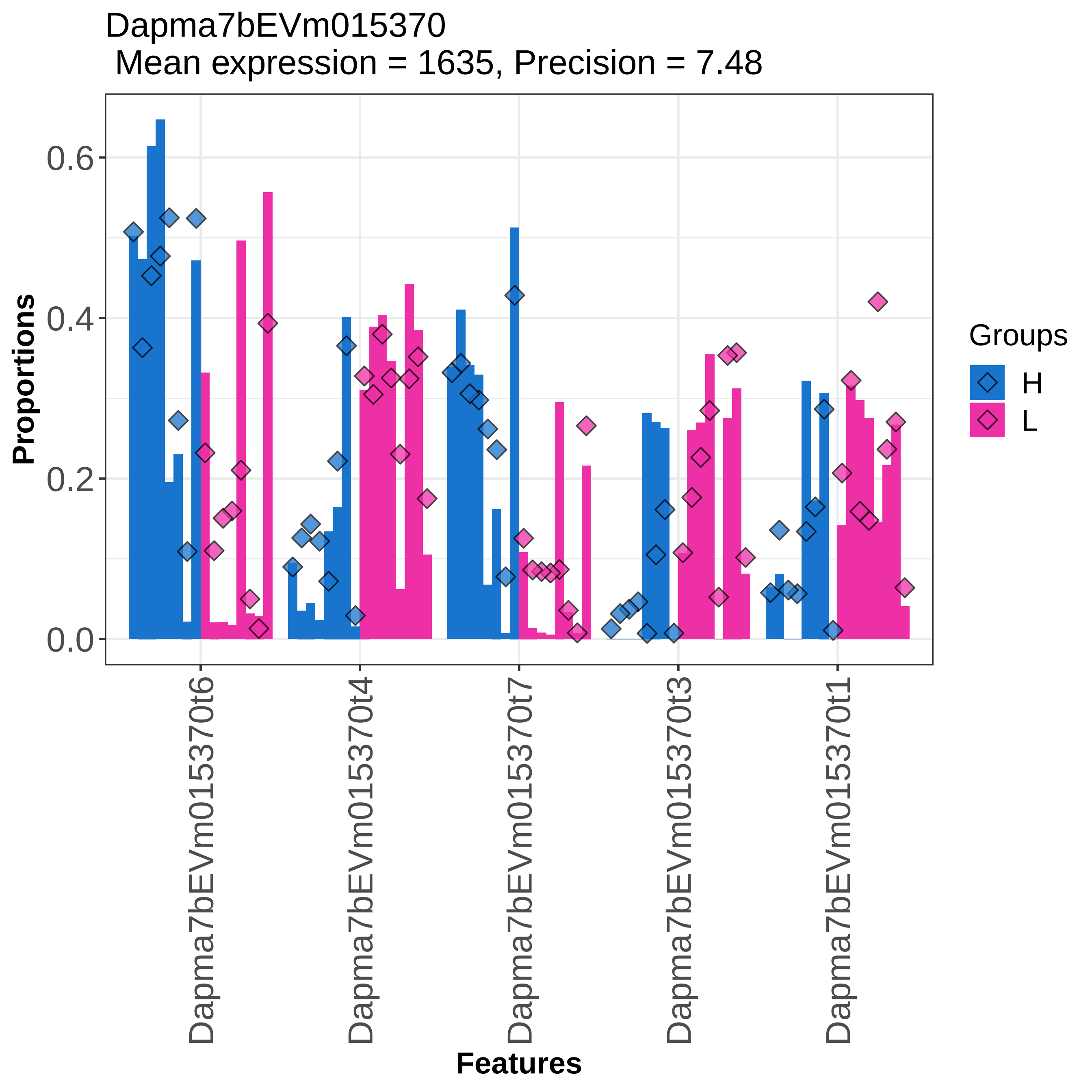

Dapma7bEVm015370
Mean expression = 1635, Precision = 7.48
Proportions
0.0
0.2
0.4
0.6
Features
Dapma7bEVm015370t6
Dapma7bEVm015370t4
Dapma7bEVm015370t7
Dapma7bEVm015370t3
Dapma7bEVm015370t1
Groups
H
L

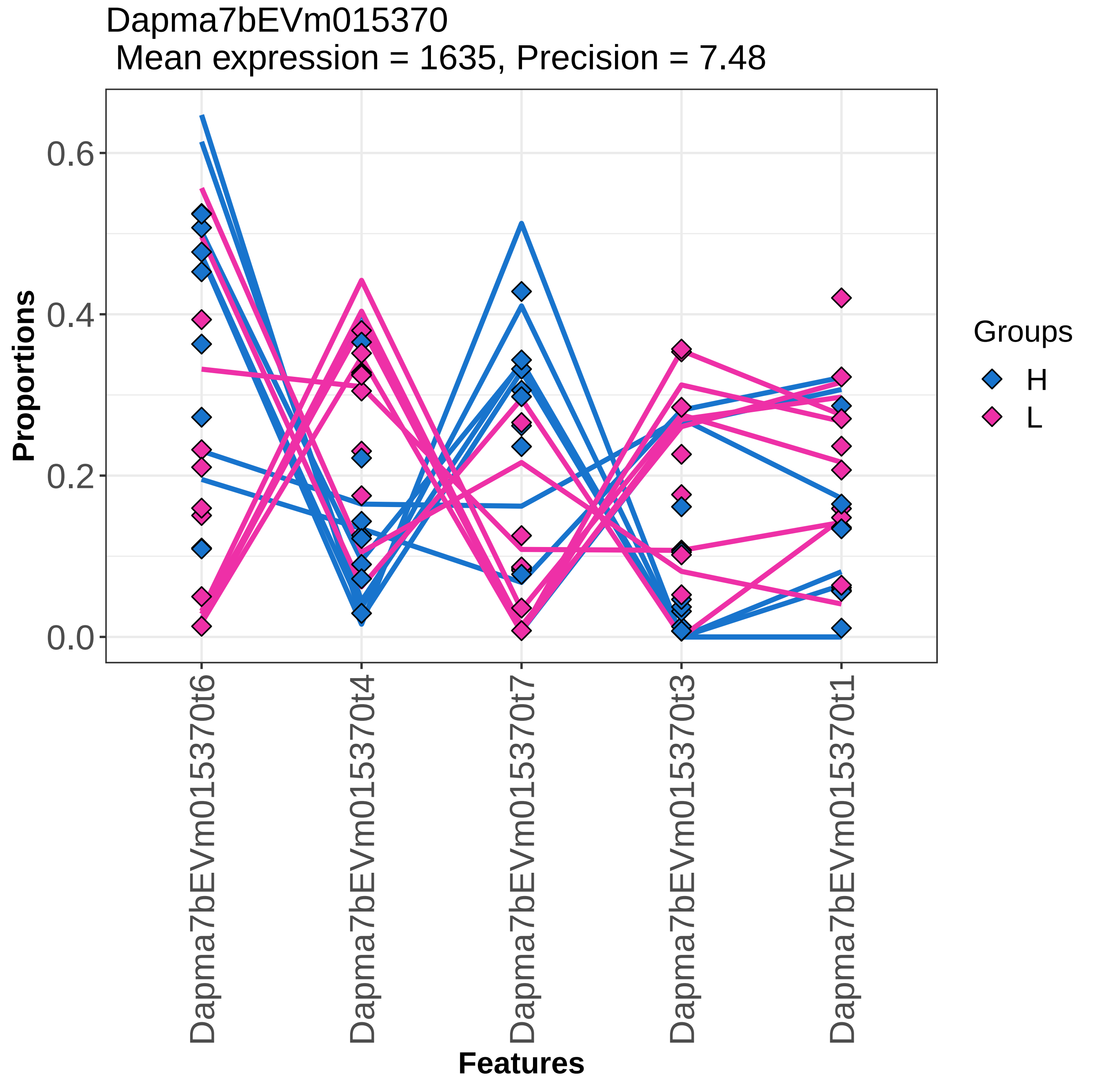

Dapma7bEVm015370
Mean expression = 1635, Precision = 7.48
Proportions
0.0
0.2
0.4
0.6
Features
Dapma7bEVm015370t6
Dapma7bEVm015370t4
Dapma7bEVm015370t7
Dapma7bEVm015370t3
Dapma7bEVm015370t1
Groups
H
L

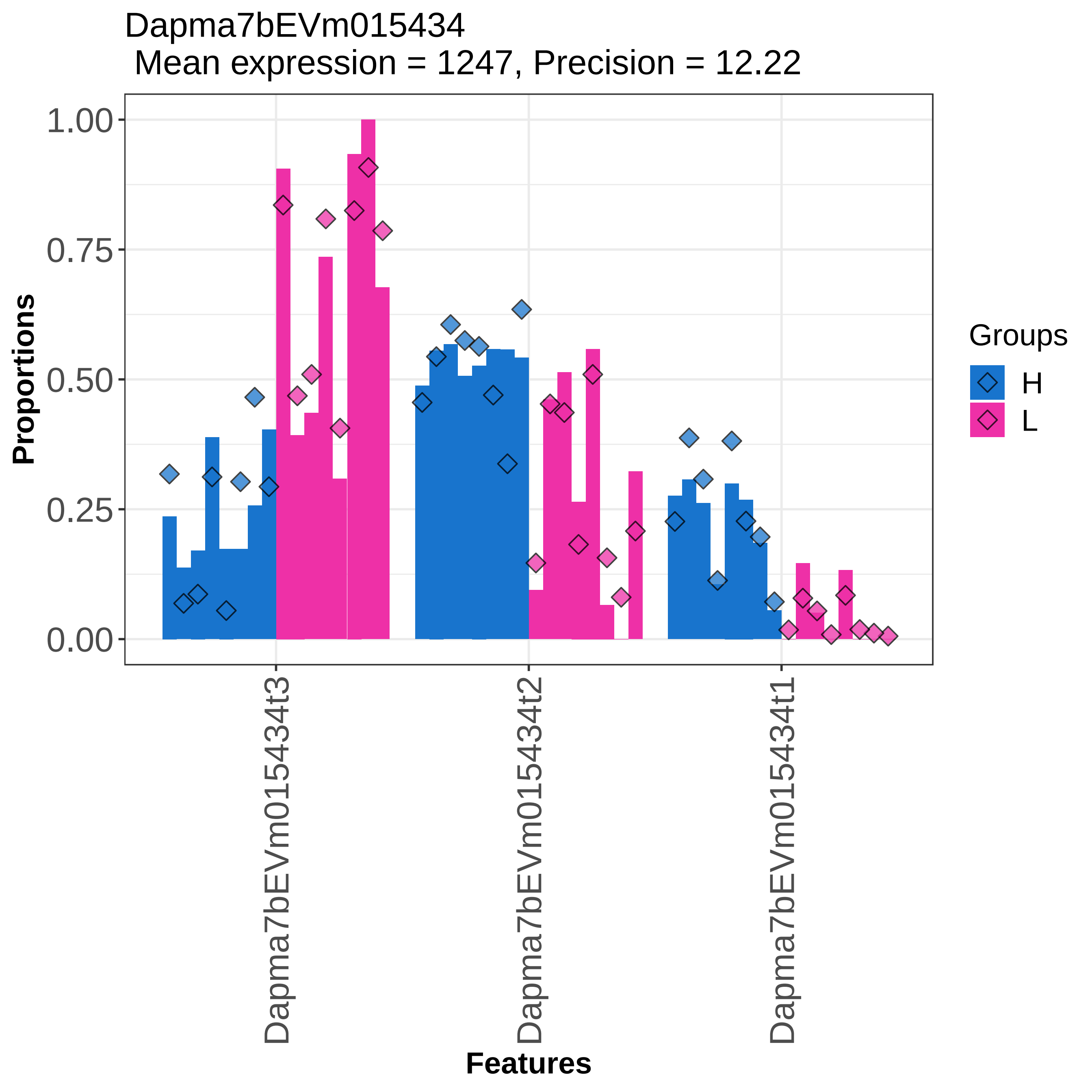

Dapma7bEVm015434
Mean expression = 1247, Precision = 12.22
Proportions
1.00
0.75
0.50
0.25
0.00
Dapma7bEVm015434t3
Dapma7bEVm015434t2
Dapma7bEVm015434t1
Features
Groups
H
L

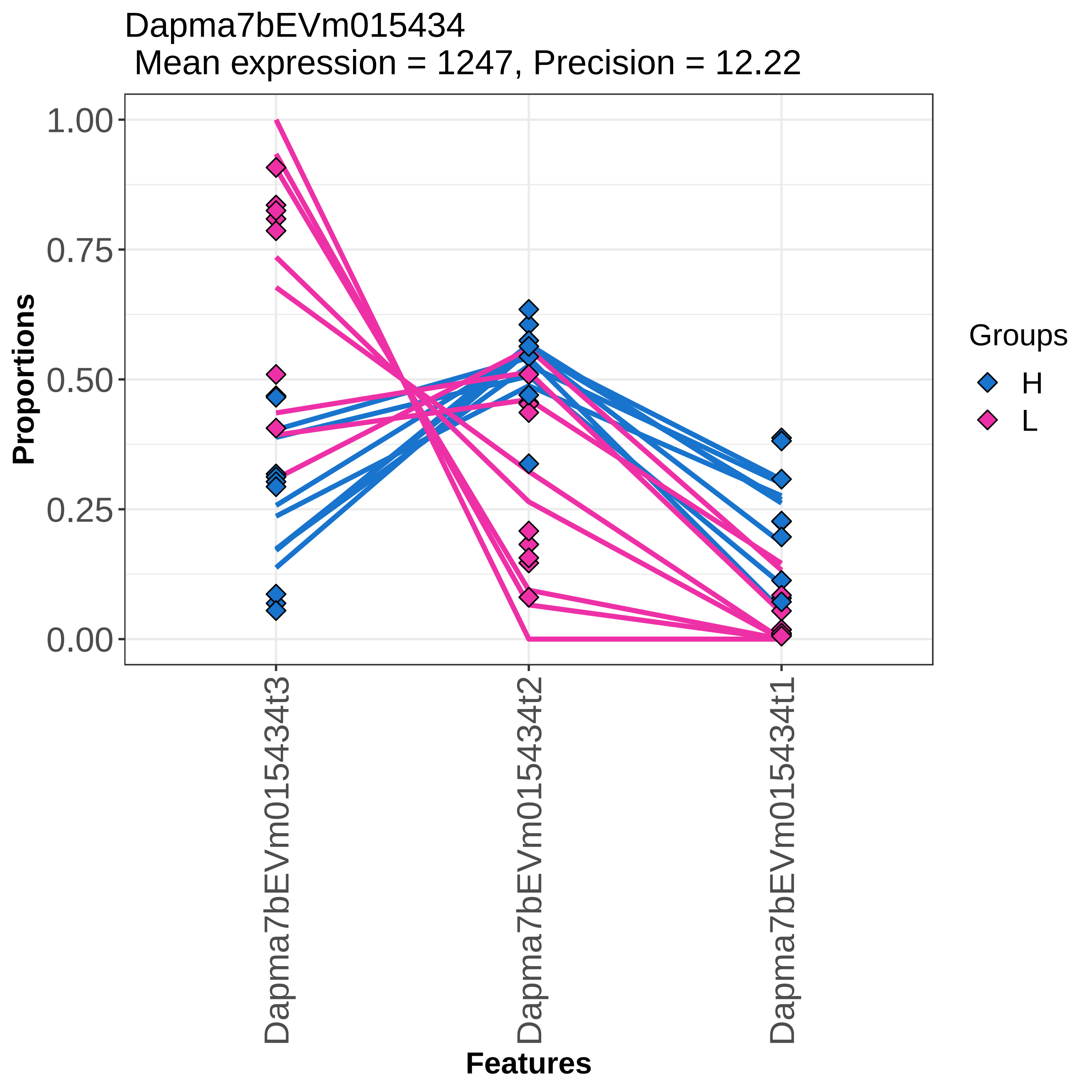

Dapma7bEVm015434
Mean expression = 1247, Precision = 12.22
Proportions
1.00
0.75
0.50
0.25
0.00
Dapma7bEVm015434t3
Dapma7bEVm015434t2
Dapma7bEVm015434t1
Features
Groups
H
L

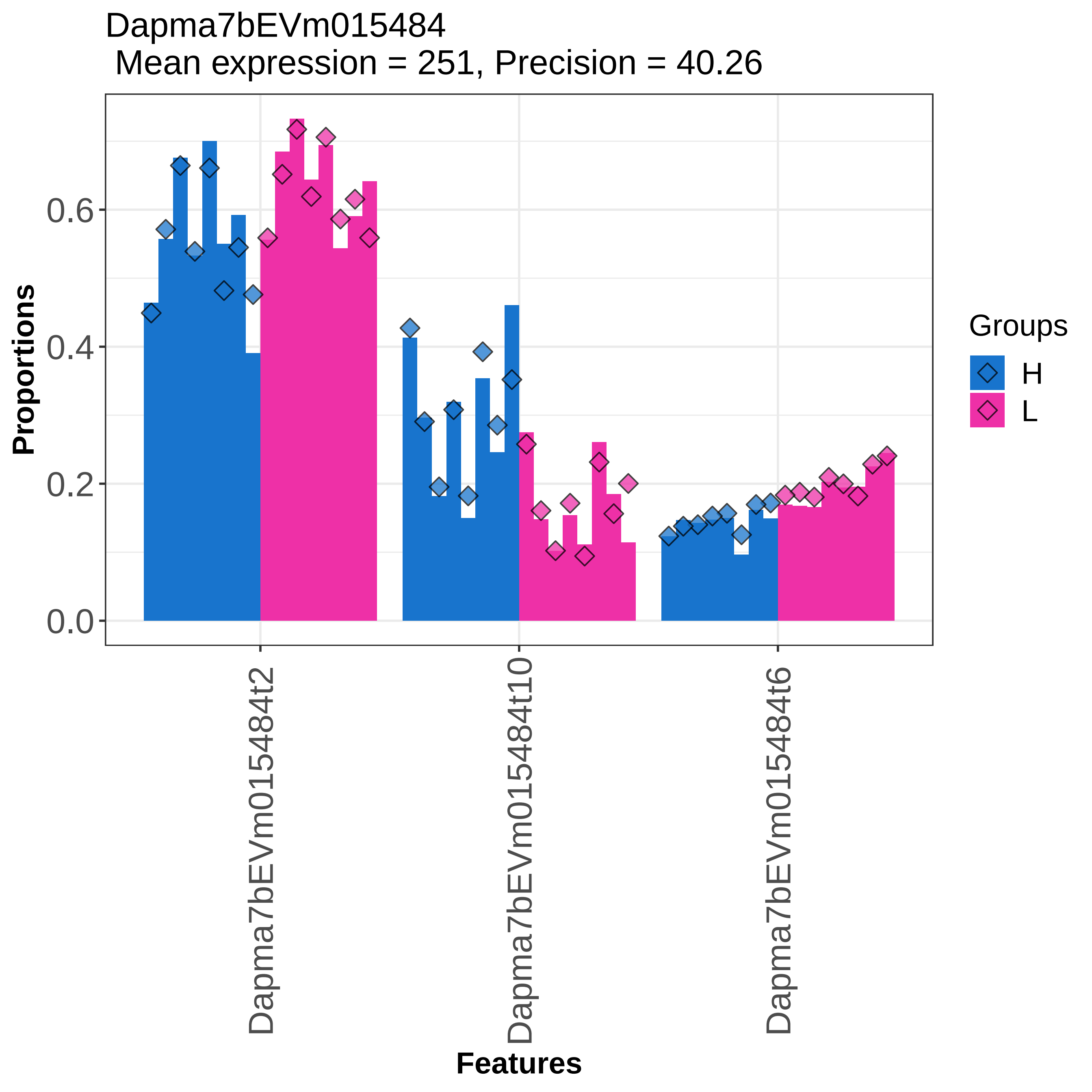
Dapma7bEVm015484
Mean expression = 251, Precision = 40.26
Proportions
0.0
0.2
0.4
0.6
Features
Dapma7bEVm015484t2
Dapma7bEVm015484t10
Dapma7bEVm015484t6
Groups
H
L

Dapma7bEVm015484
 Mean expression = 251, Precision = 40.26

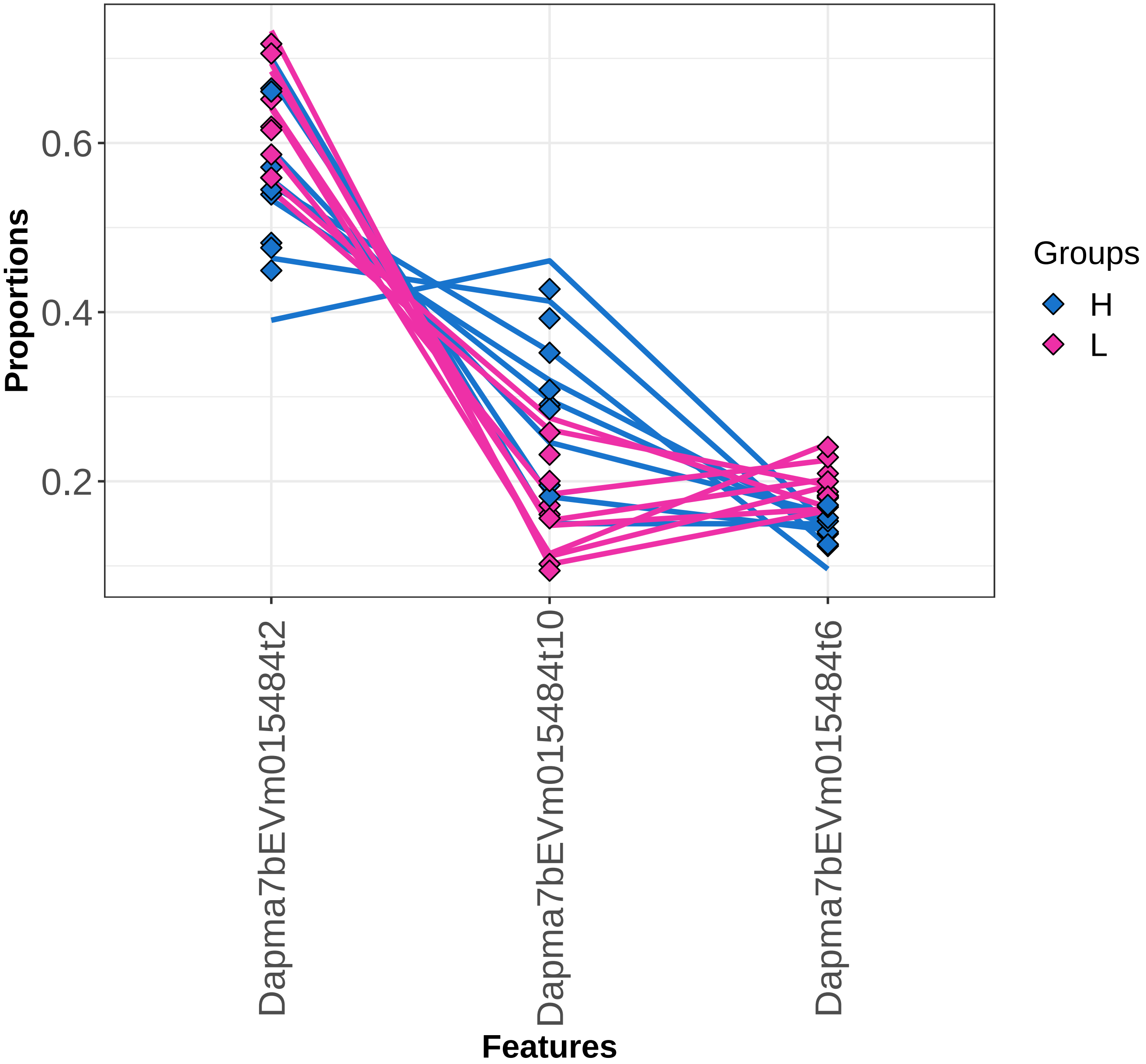

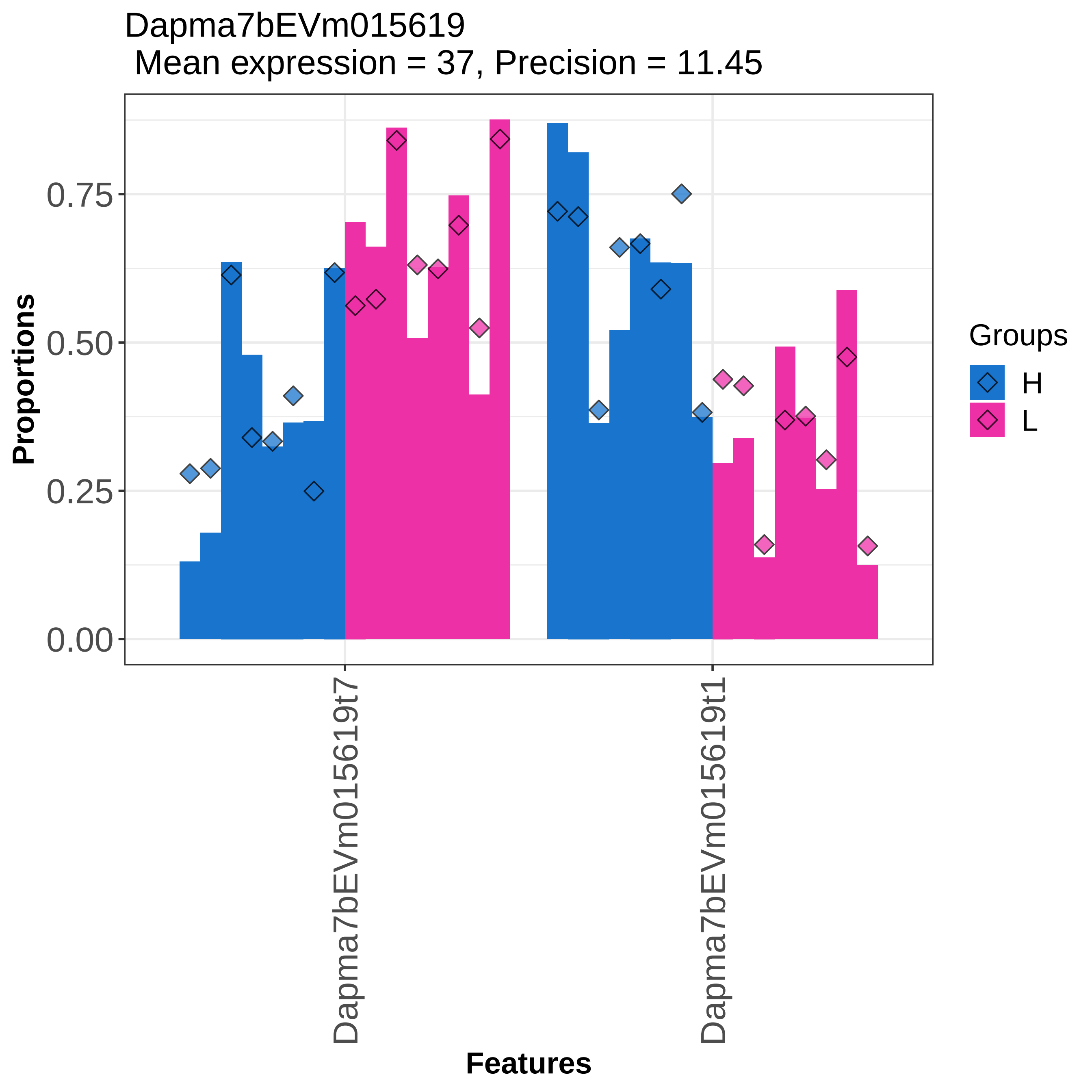

Dapma7bEVm015619
Mean expression = 37, Precision = 11.45
Proportions
0.00
0.25
0.50
0.75
Features
Dapma7bEVm015619t7
Dapma7bEVm015619t1
Groups
H
L

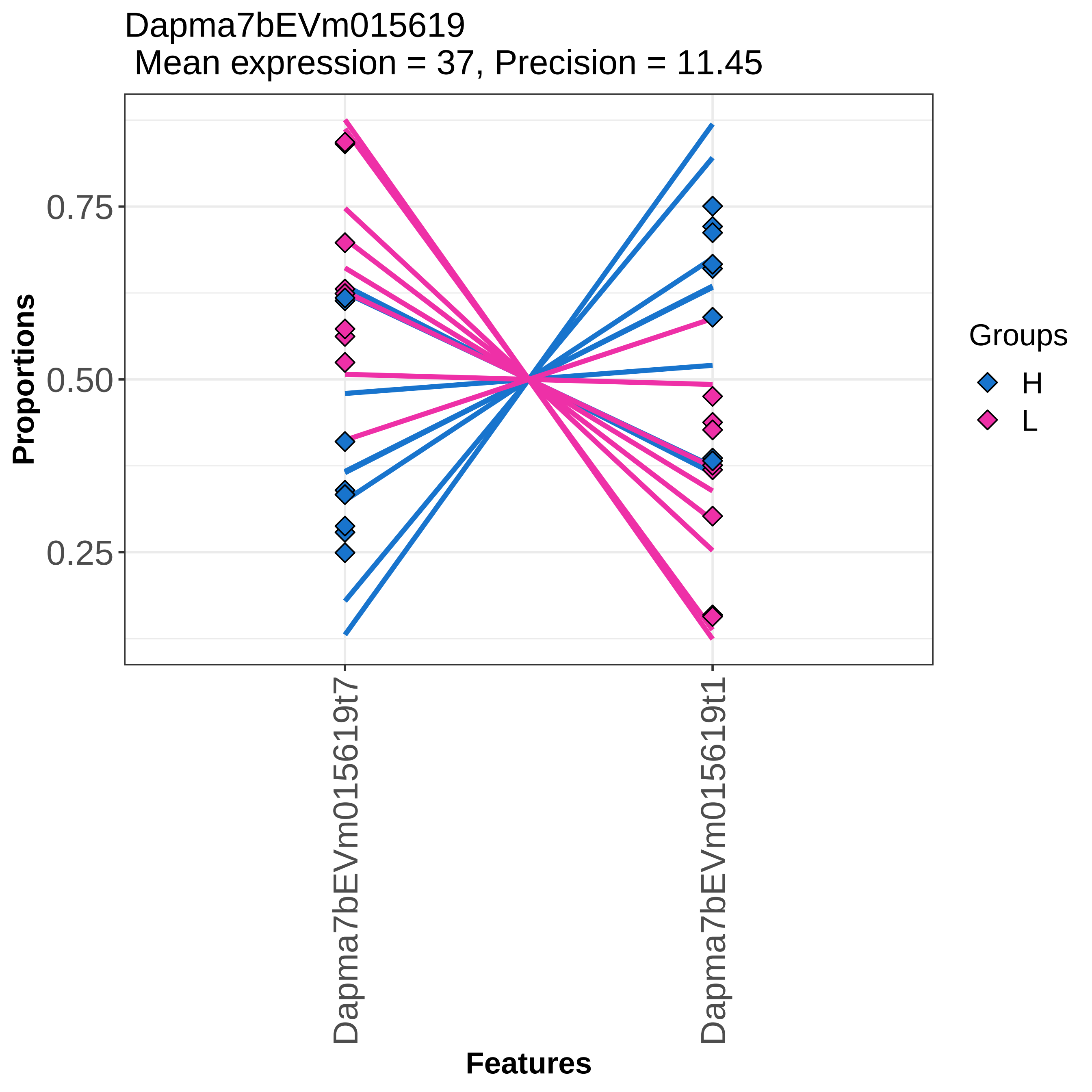
Dapma7bEVm015619
Mean expression = 37, Precision = 11.45
Proportions
0.75
0.50
0.25
Dapma7bEVm015619t7
Dapma7bEVm015619t1
Features
Groups
H
L

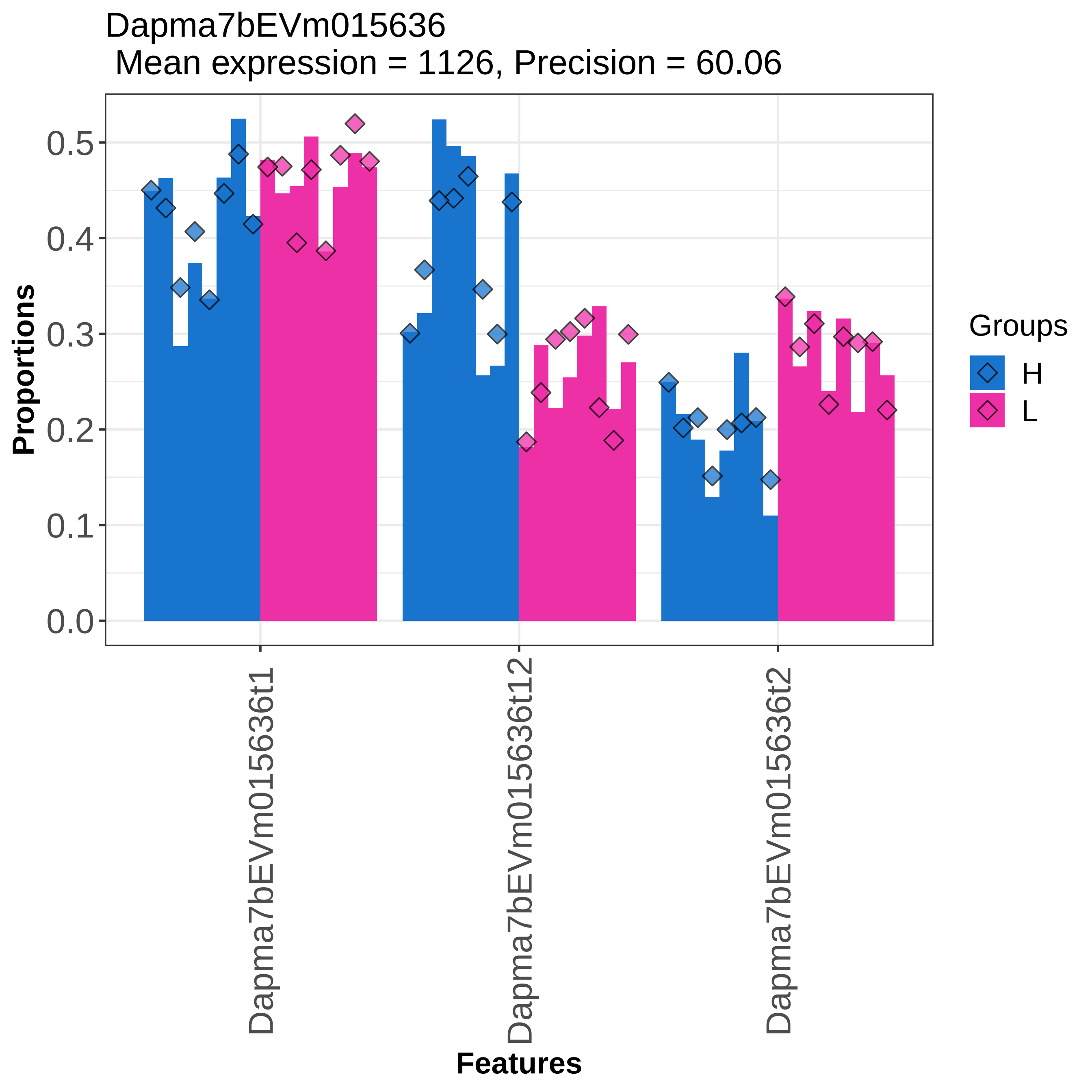

Dapma7bEVm015636
Mean expression = 1126, Precision = 60.06
Proportions
0.0
0.1
0.2
0.3
0.4
0.5
Dapma7bEVm015636t1
Dapma7bEVm015636t12
Dapma7bEVm015636t2
Features
Groups
H
L

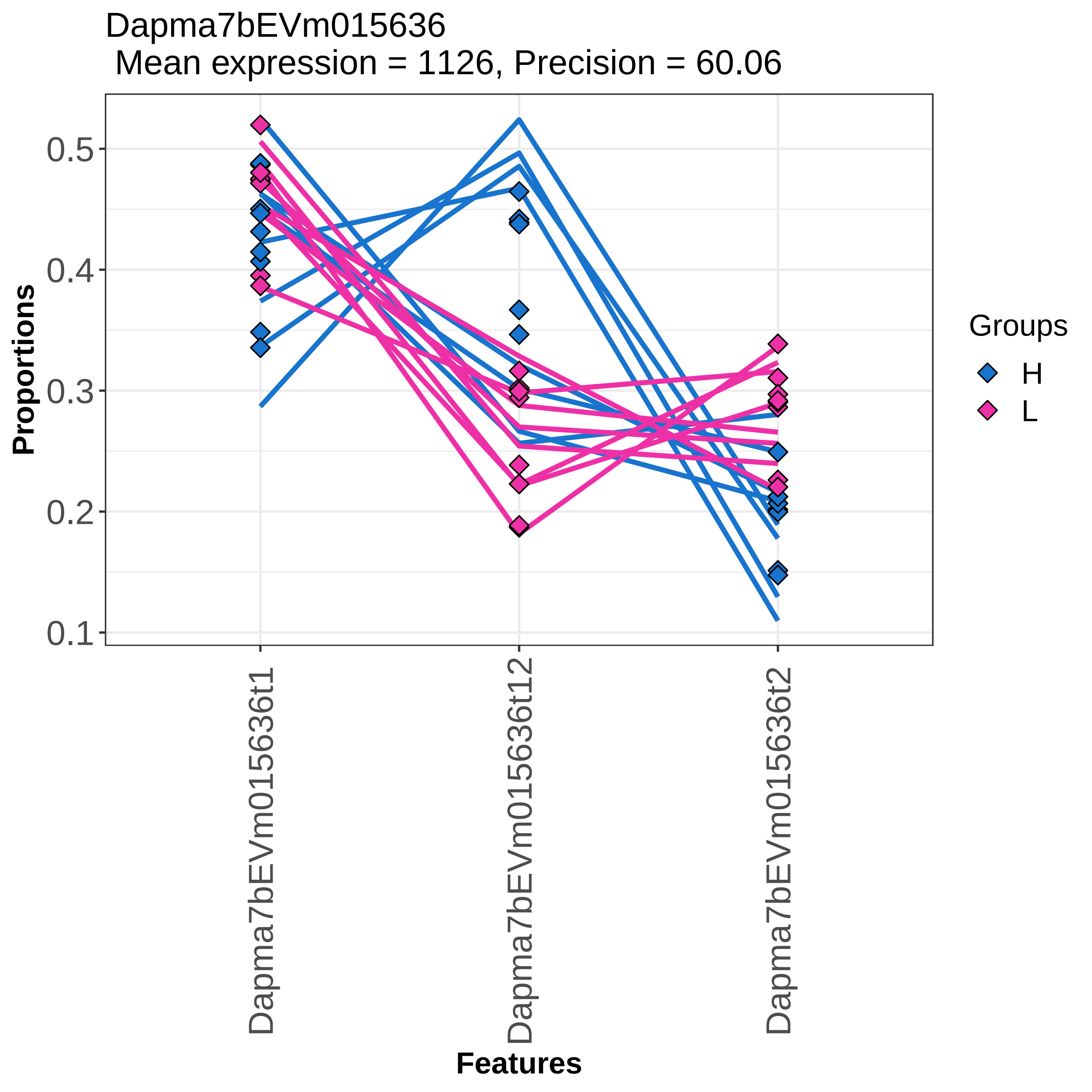

Dapma7bEVm015636
Mean expression = 1126, Precision = 60.06
Proportions
0.5
0.4
0.3
0.2
0.1
Dapma7bEVm015636t1
Dapma7bEVm015636t12
Dapma7bEVm015636t2
Features
Groups
H
L

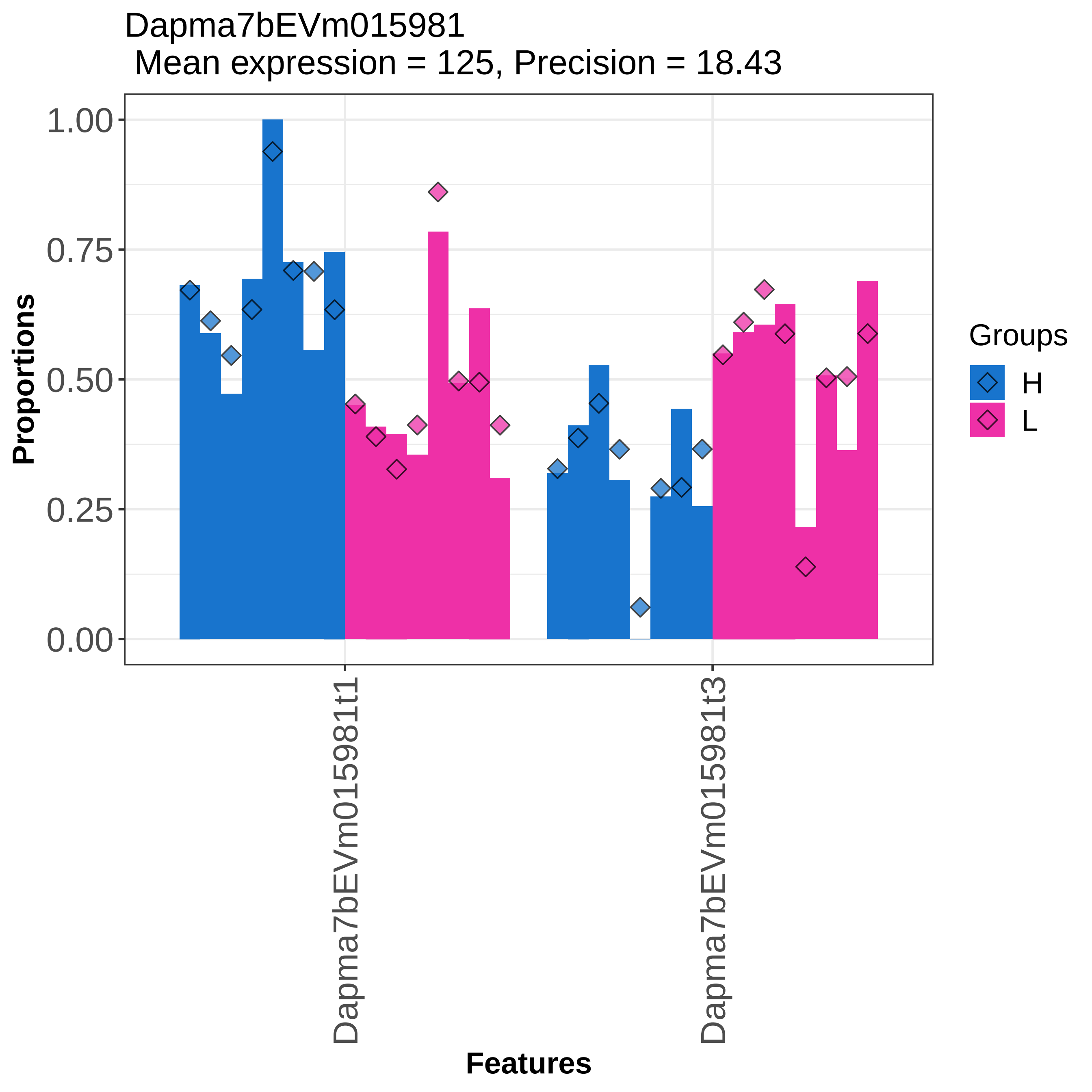

Dapma7bEVm015981
Mean expression = 125, Precision = 18.43
Proportions
1.00
0.75
0.50
0.25
0.00
Dapma7bEVm015981t1
Dapma7bEVm015981t3
Features
Groups
H
L

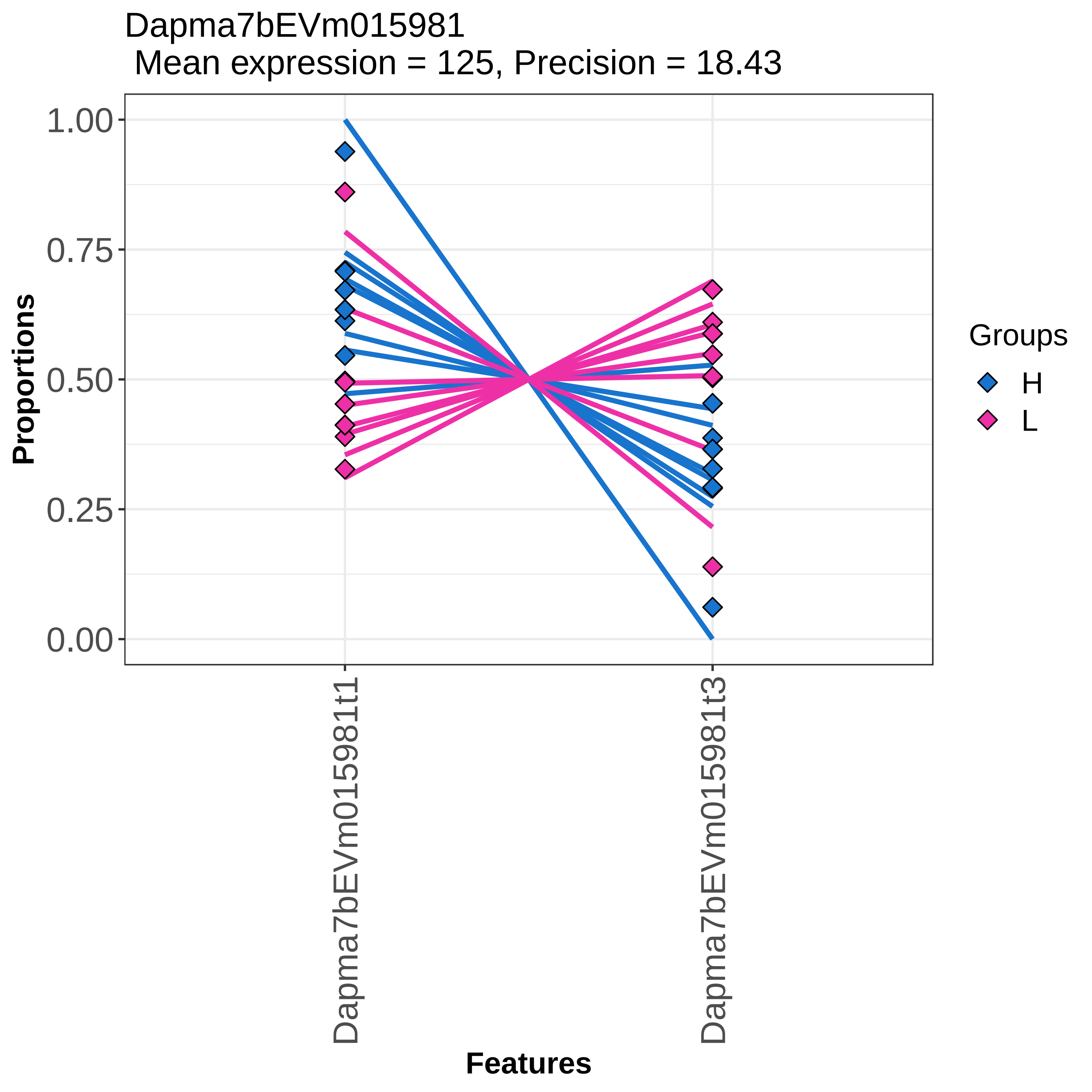
Dapma7bEVm015981
Mean expression = 125, Precision = 18.43
Proportions
1.00
0.75
0.50
0.25
0.00
Dapma7bEVm015981t1
Dapma7bEVm015981t3
Features
Groups
H
L

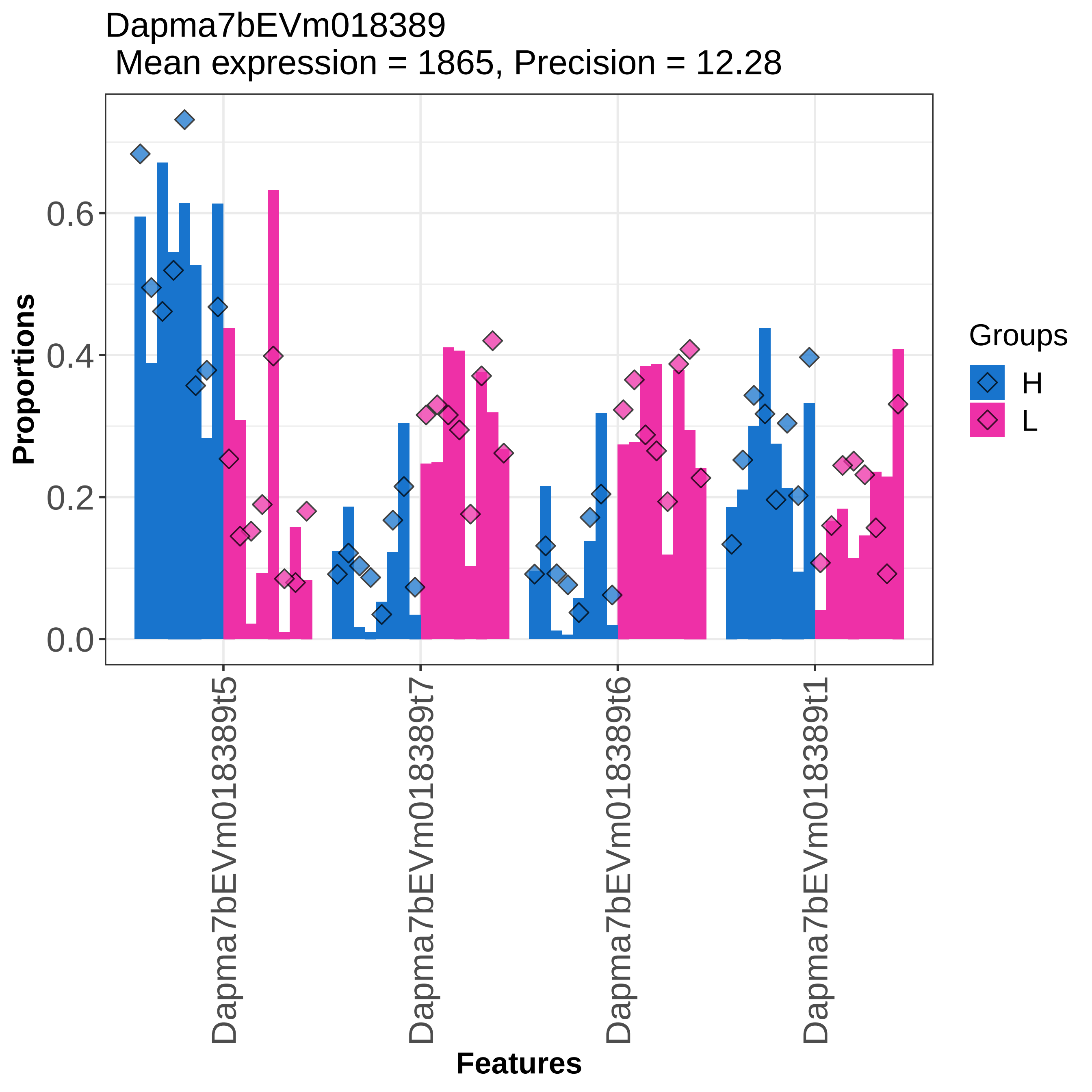

Dapma7bEVm018389
Mean expression = 1865, Precision = 12.28
Proportions
0.0
0.2
0.4
0.6
Dapma7bEVm018389t5
Dapma7bEVm018389t7
Dapma7bEVm018389t6
Dapma7bEVm018389t1
Features
Groups
H
L

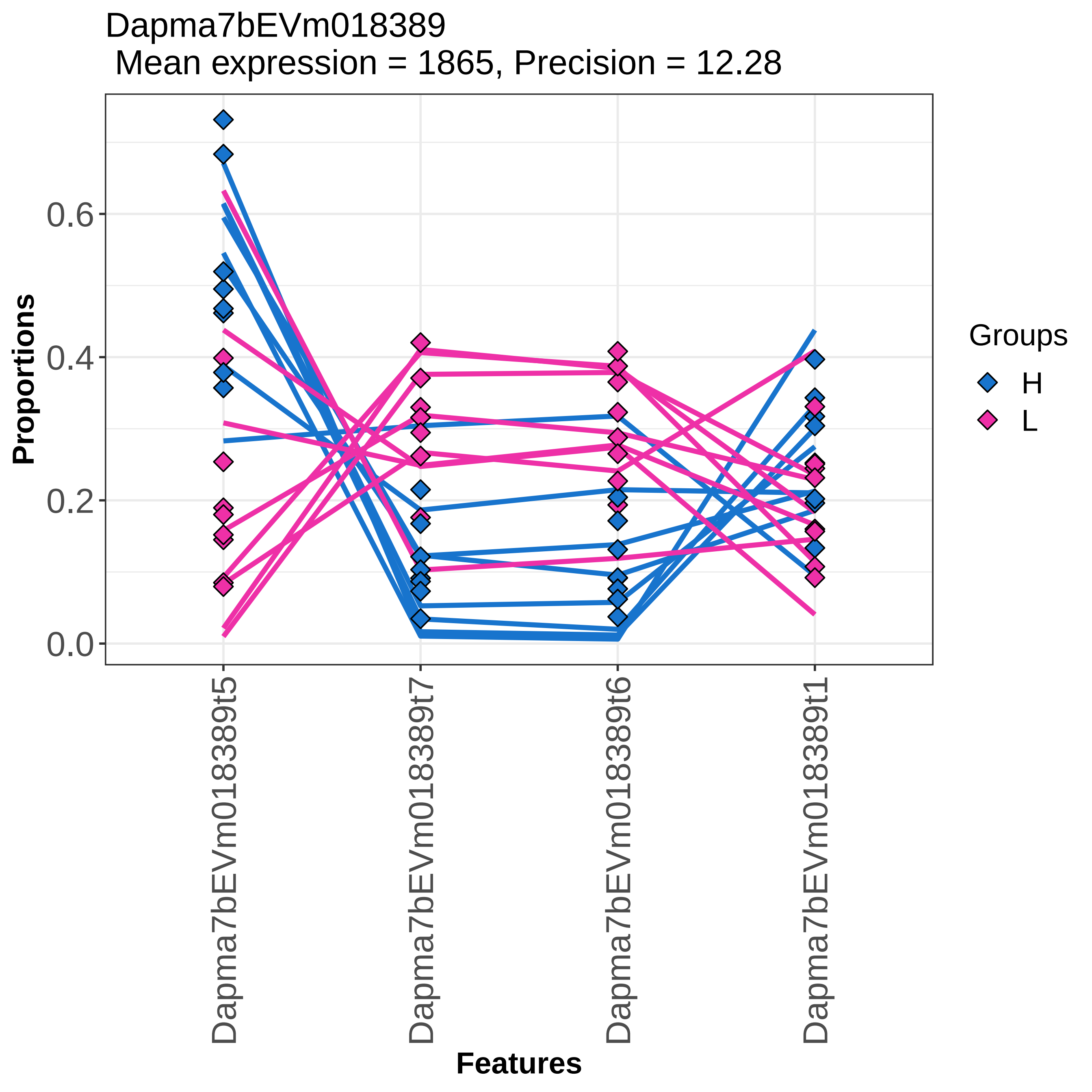

Dapma7bEVm018389
Mean expression = 1865, Precision = 12.28
Proportions
0.0
0.2
0.4
0.6
Dapma7bEVm018389t5
Dapma7bEVm018389t7
Dapma7bEVm018389t6
Dapma7bEVm018389t1
Features
Groups
H
L

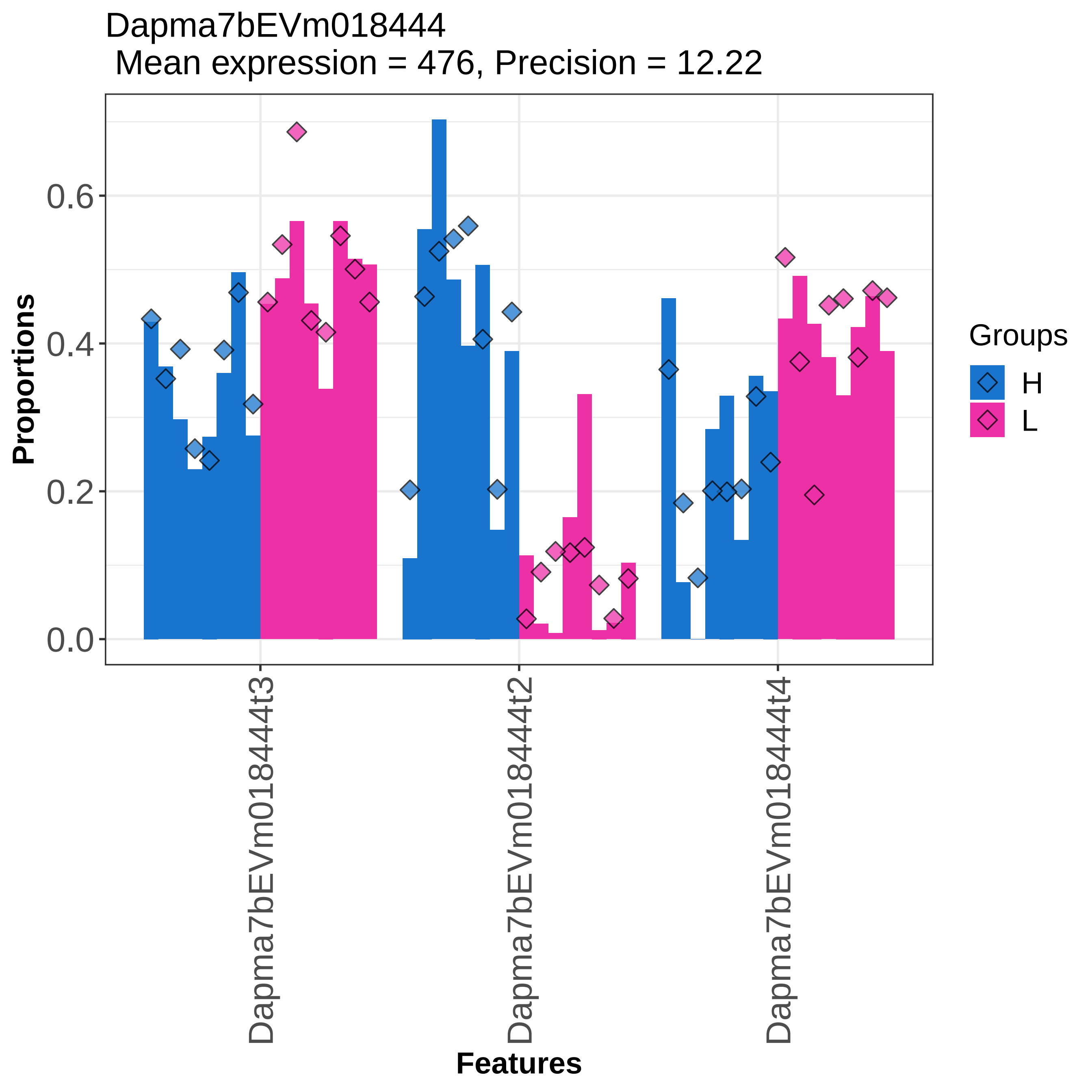

Dapma7bEVm018444
Mean expression = 476, Precision = 12.22
Proportions
0.0
0.2
0.4
0.6
Dapma7bEVm018444t3
Dapma7bEVm018444t2
Dapma7bEVm018444t4
Features
Groups
H
L

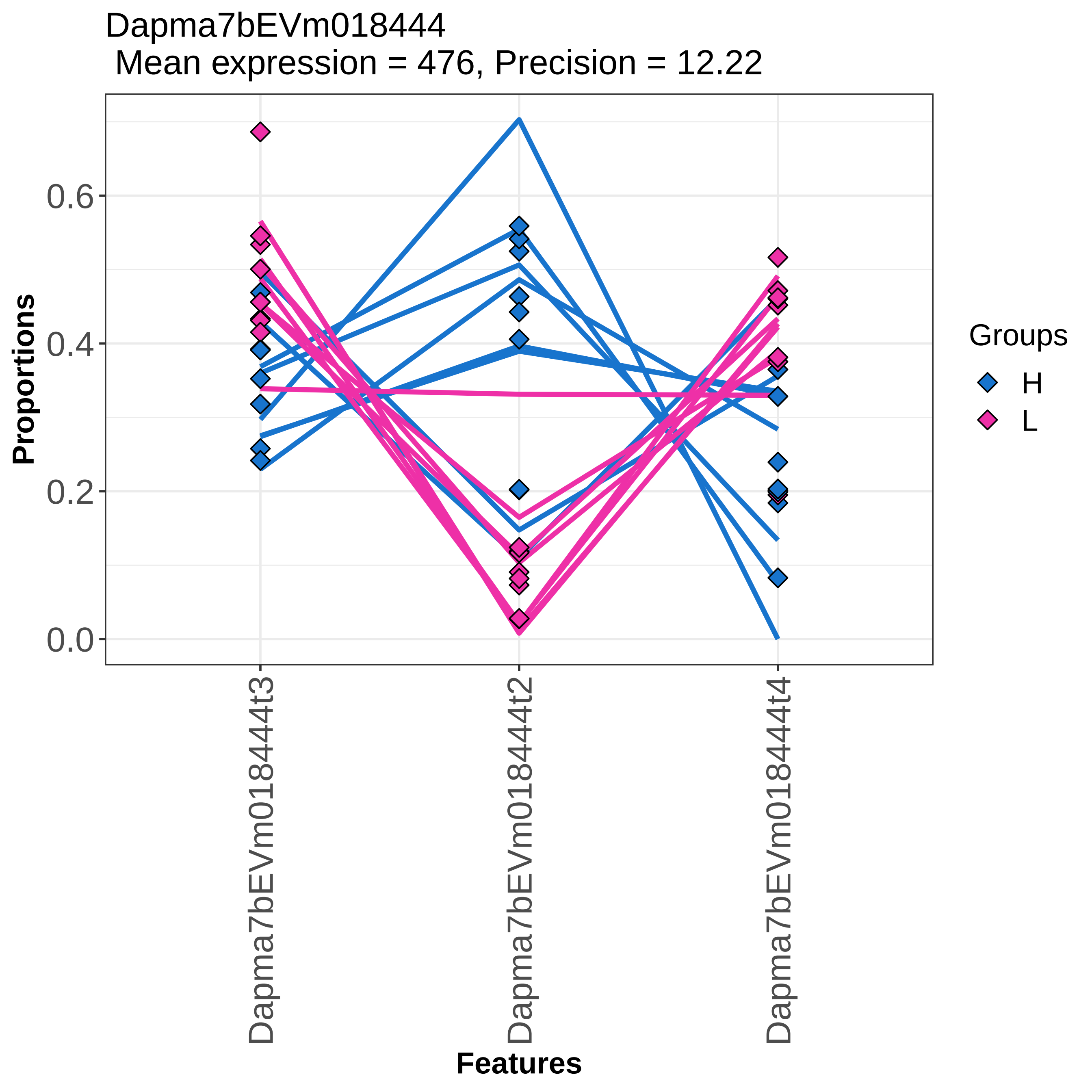

Dapma7bEVm018444
Mean expression = 476, Precision = 12.22
Proportions
0.0
0.2
0.4
0.6
Features
Dapma7bEVm018444t3
Dapma7bEVm018444t2
Dapma7bEVm018444t4
Groups
H
L

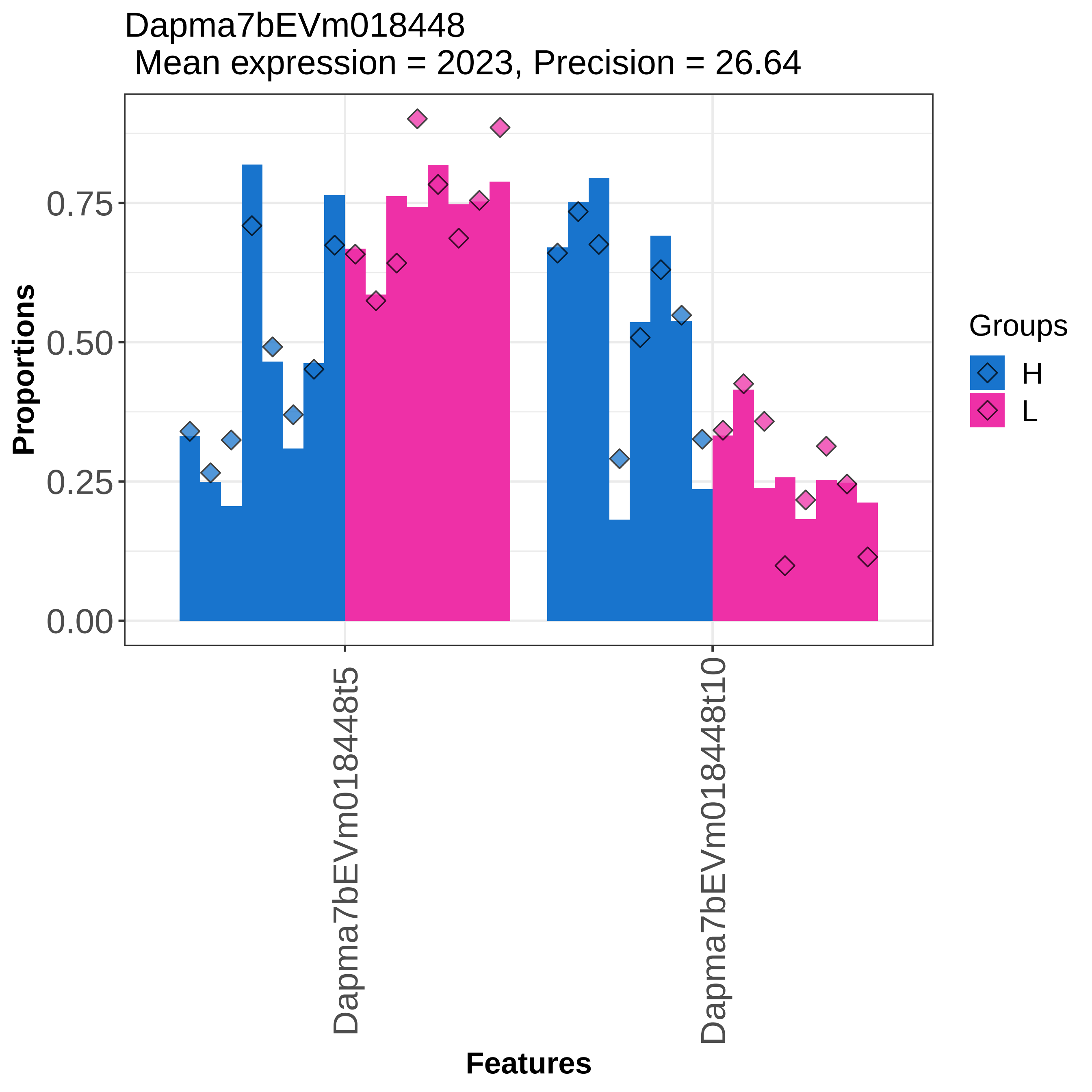

Dapma7bEVm018448
Mean expression = 2023, Precision = 26.64
Proportions
0.75
0.50
0.25
0.00
Dapma7bEVm018448t5
Dapma7bEVm018448t10
Features
Groups
H
L

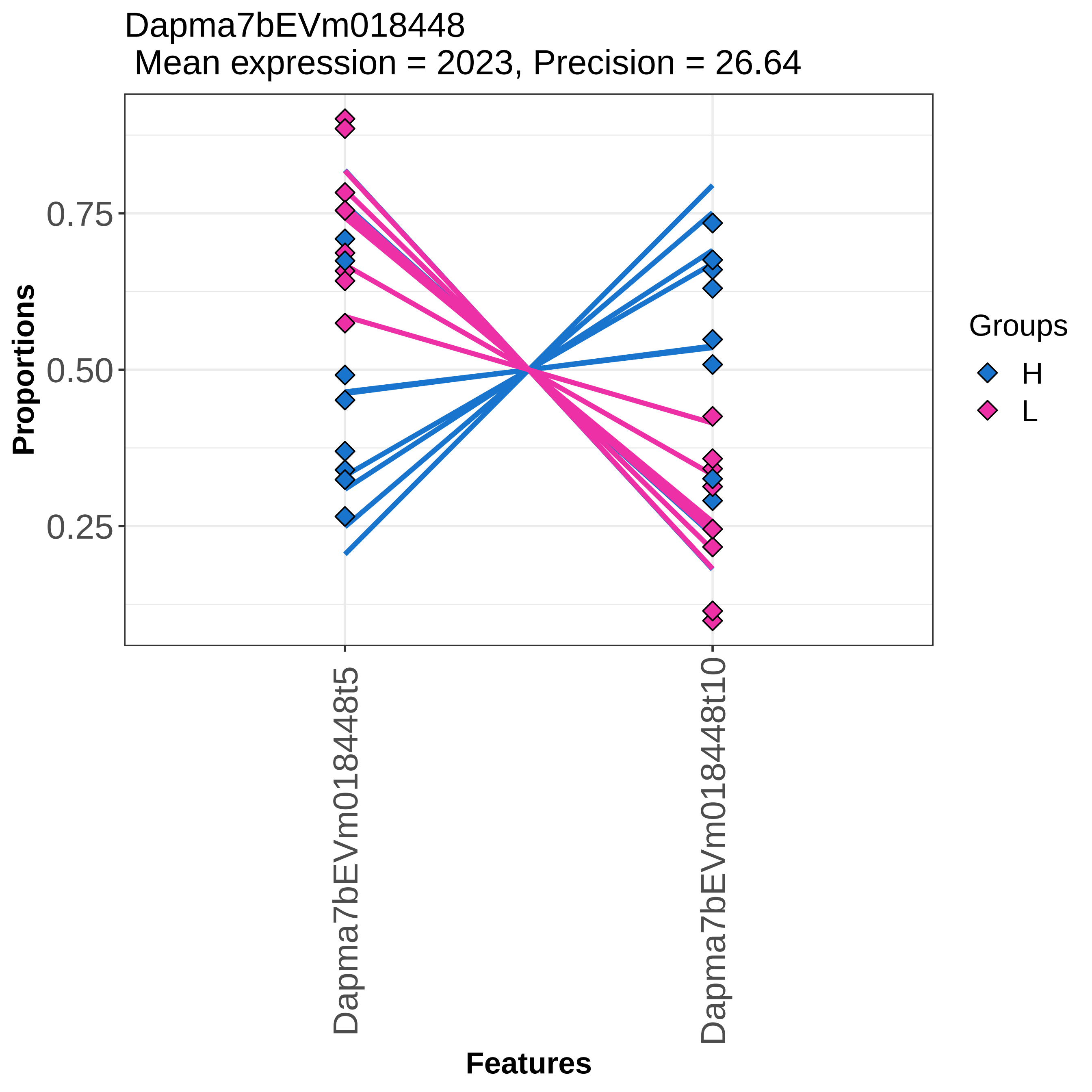
Dapma7bEVm018448
Mean expression = 2023, Precision = 26.64
Proportions
0.75
0.50
0.25
Dapma7bEVm018448t5
Dapma7bEVm018448t10
Features
Groups
H
L

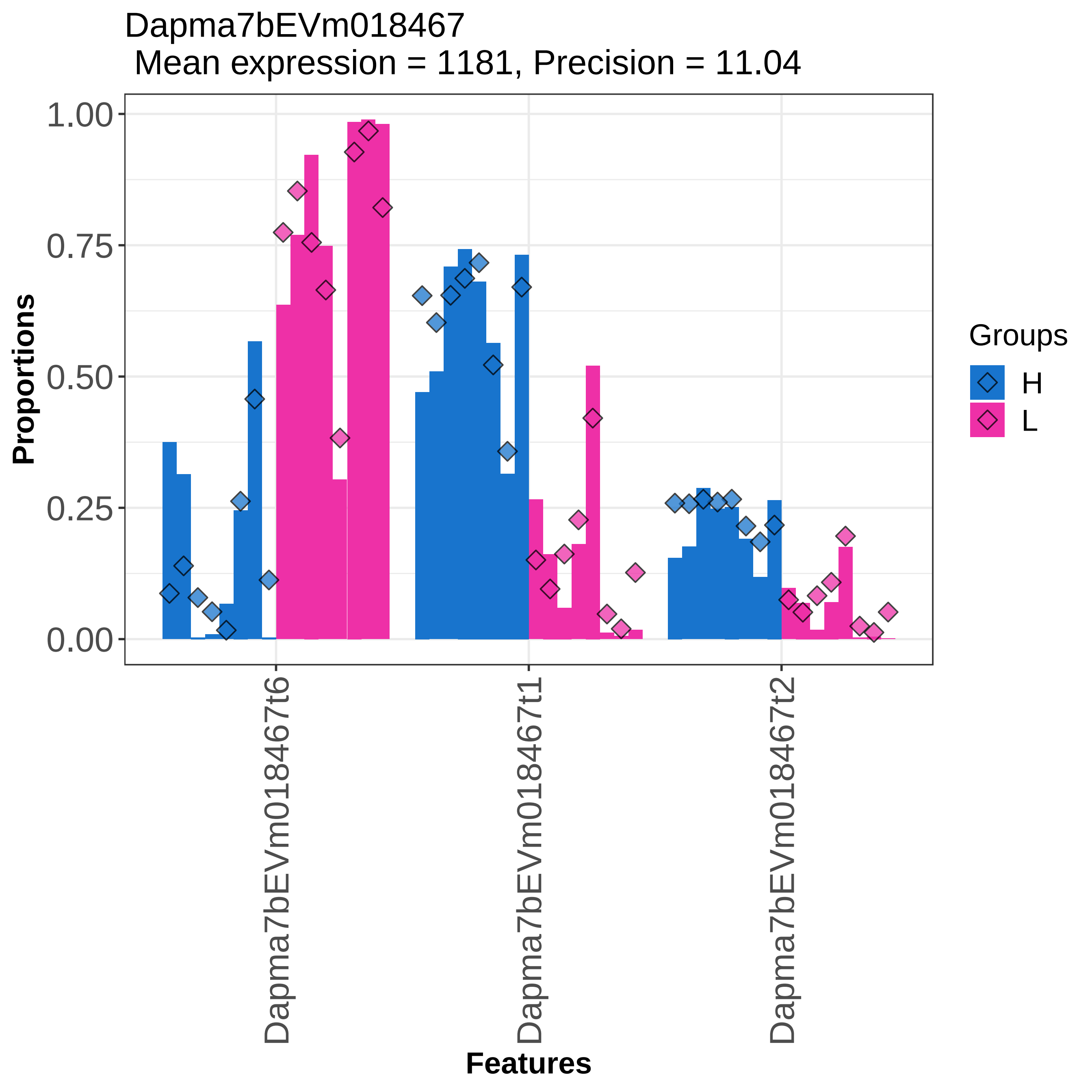
Dapma7bEVm018467
Mean expression = 1181, Precision = 11.04
Proportions
1.00
0.75
0.50
0.25
0.00
Dapma7bEVm018467t6
Dapma7bEVm018467t1
Dapma7bEVm018467t2
Features
Groups
H
L

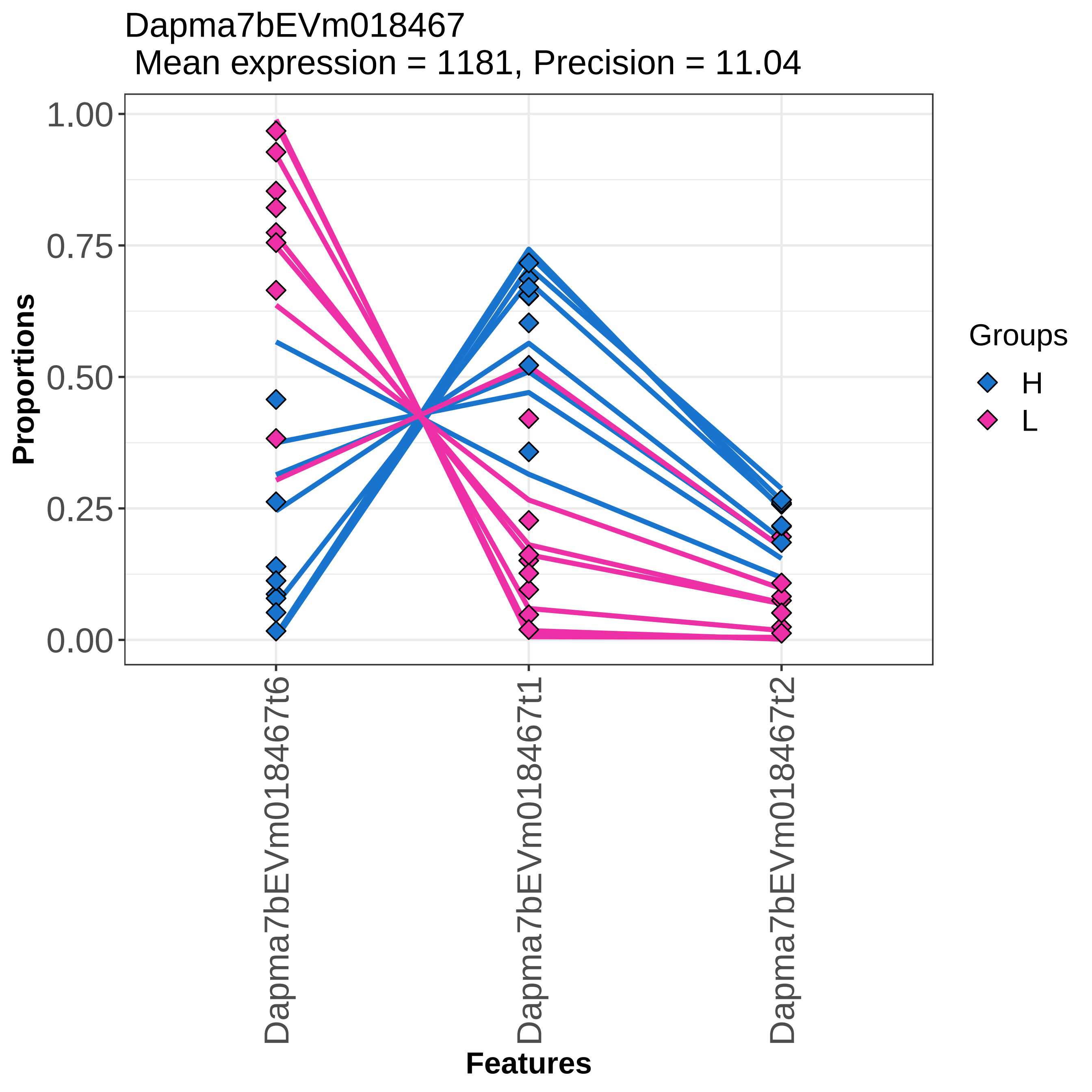

Dapma7bEVm018467
Mean expression = 1181, Precision = 11.04
Proportions
1.00
0.75
0.50
0.25
0.00
Dapma7bEVm018467t6
Dapma7bEVm018467t1
Dapma7bEVm018467t2
Features
Groups
H
L

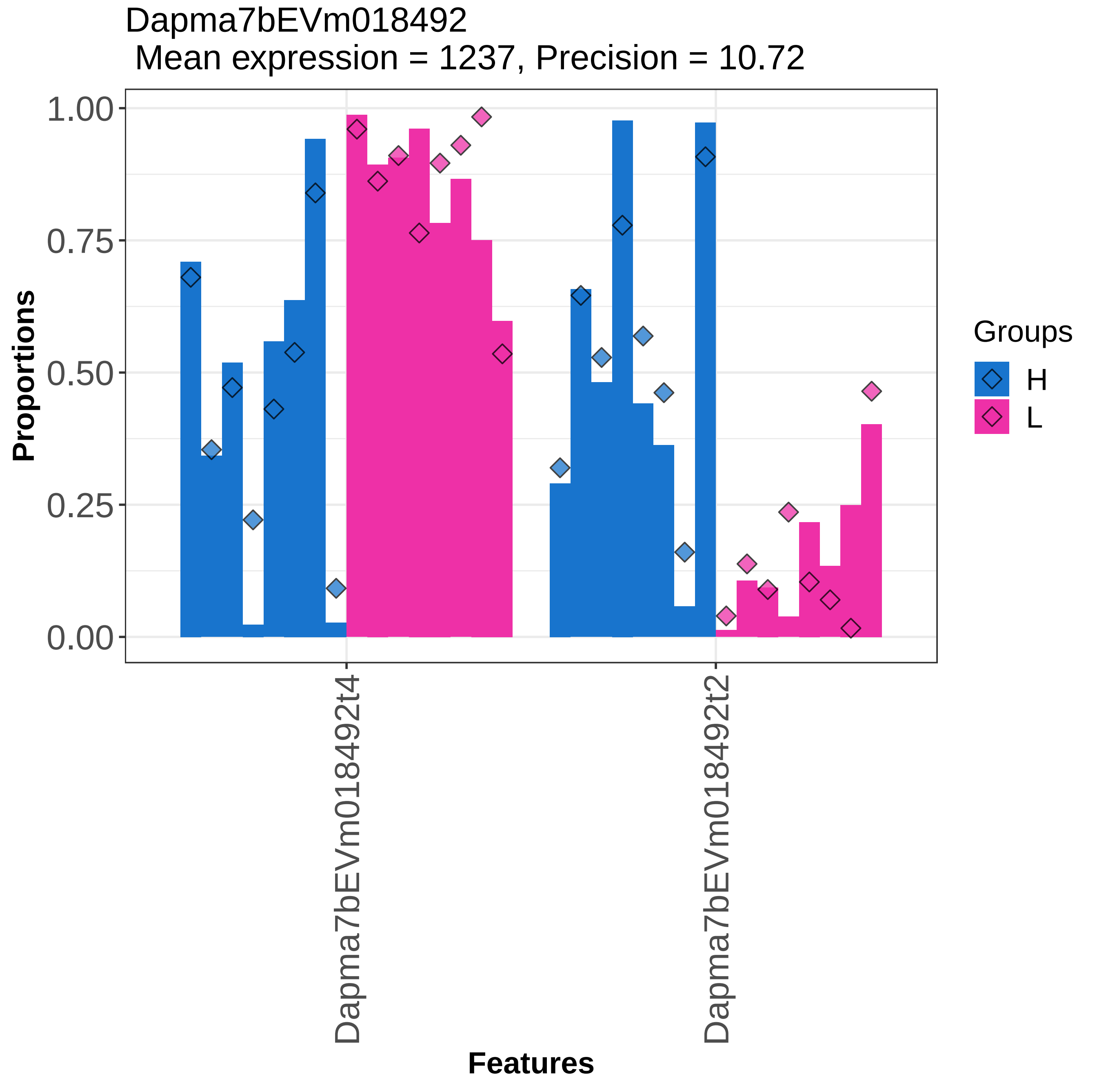
Dapma7bEVm018492
Mean expression = 1237, Precision = 10.72
Proportions
1.00
0.75
0.50
0.25
0.00
Dapma7bEVm018492t4
Dapma7bEVm018492t2
Features
Groups
H
L

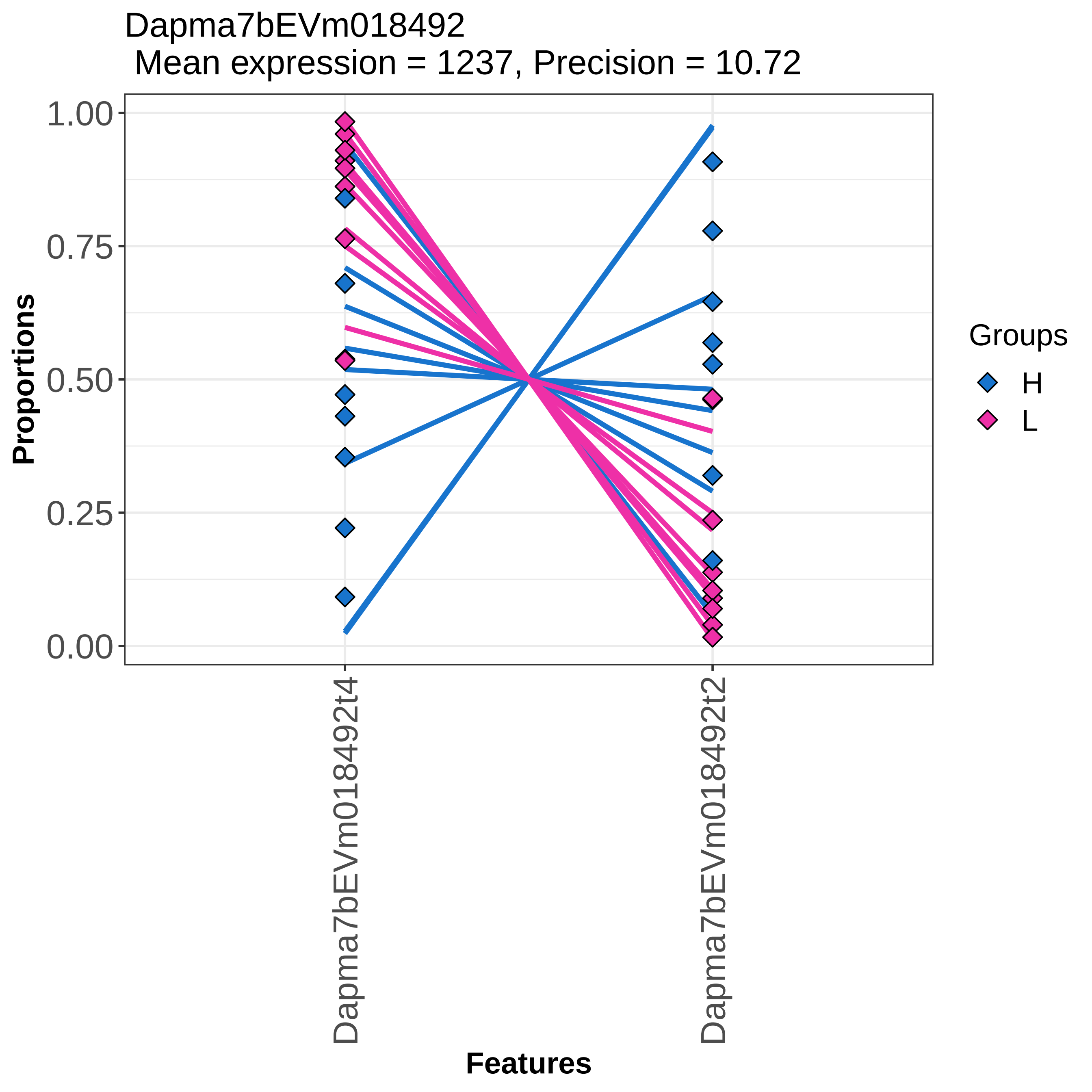
Dapma7bEVm018492
Mean expression = 1237, Precision = 10.72
Proportions
1.00
0.75
0.50
0.25
0.00
Dapma7bEVm018492t4
Dapma7bEVm018492t2
Features
Groups
H
L

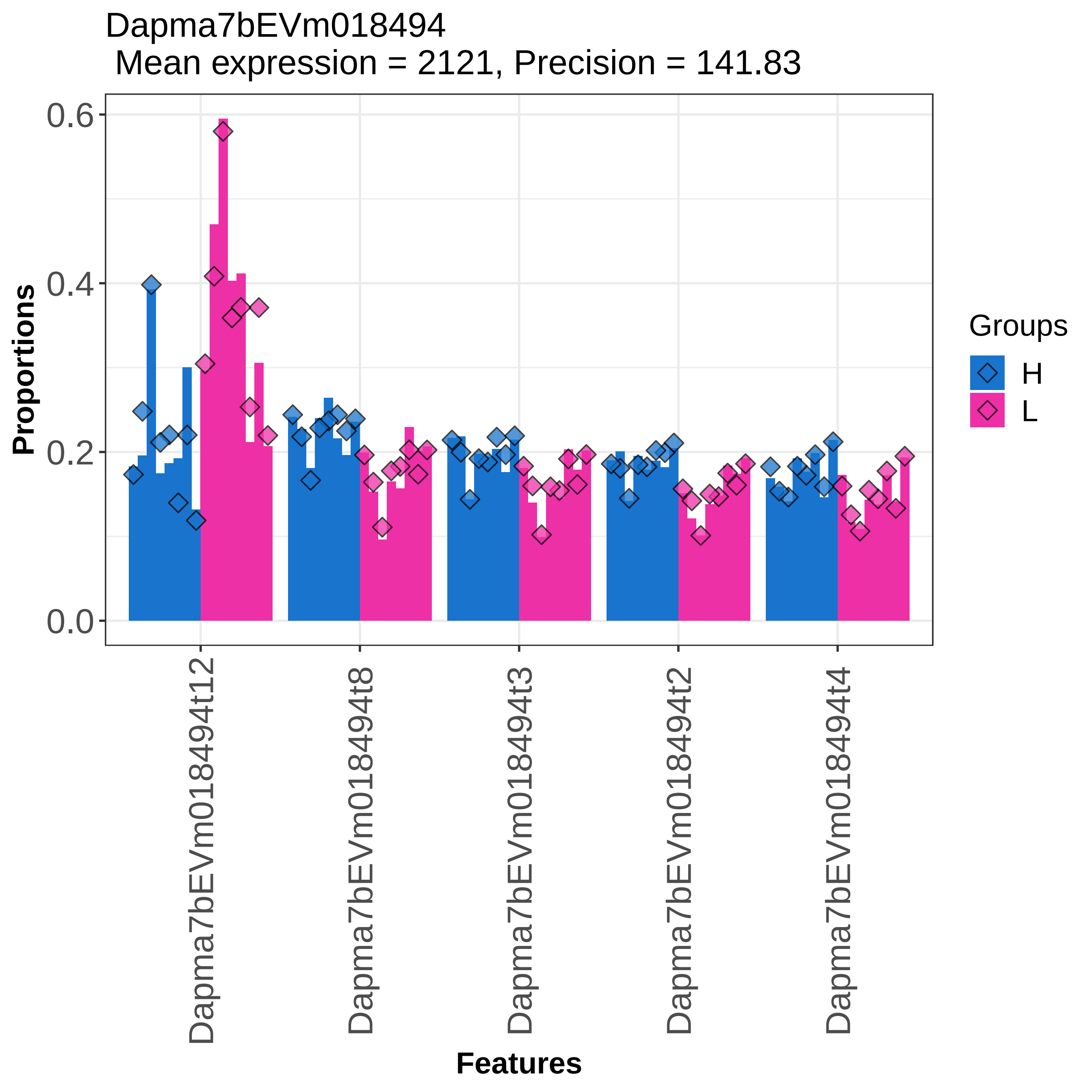
Dapma7bEVm018494
Mean expression = 2121, Precision = 141.83
Proportions
0.0
0.2
0.4
0.6
Groups
H
L
Dapma7bEVm018494t12
Dapma7bEVm018494t8
Dapma7bEVm018494t3
Dapma7bEVm018494t2
Dapma7bEVm018494t4
Features

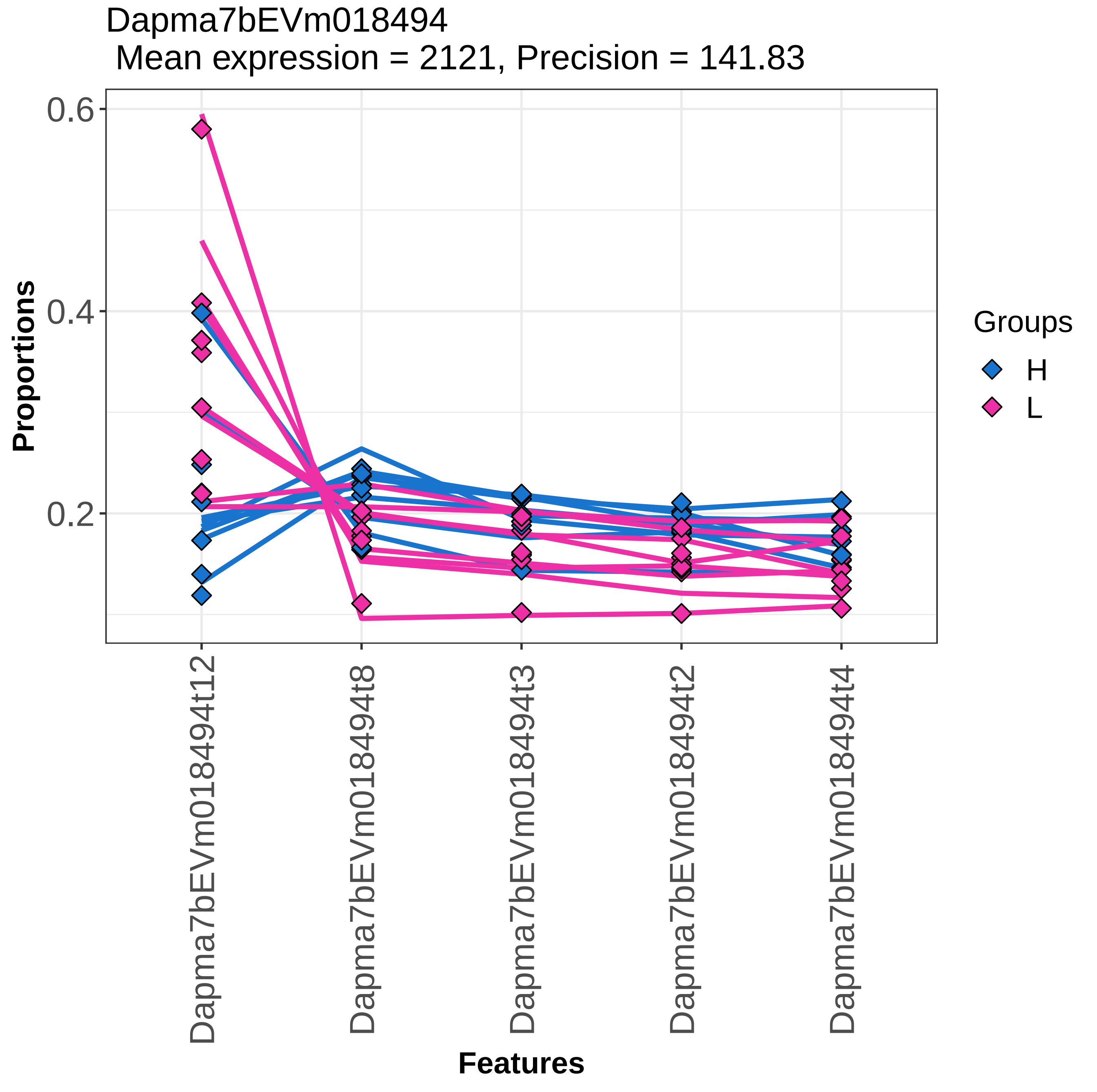

Dapma7bEVm018494
Mean expression = 2121, Precision = 141.83
Proportions
0.6
0.4
0.2
Dapma7bEVm018494t12
Dapma7bEVm018494t8
Dapma7bEVm018494t3
Dapma7bEVm018494t2
Dapma7bEVm018494t4
Features
Groups
H
L

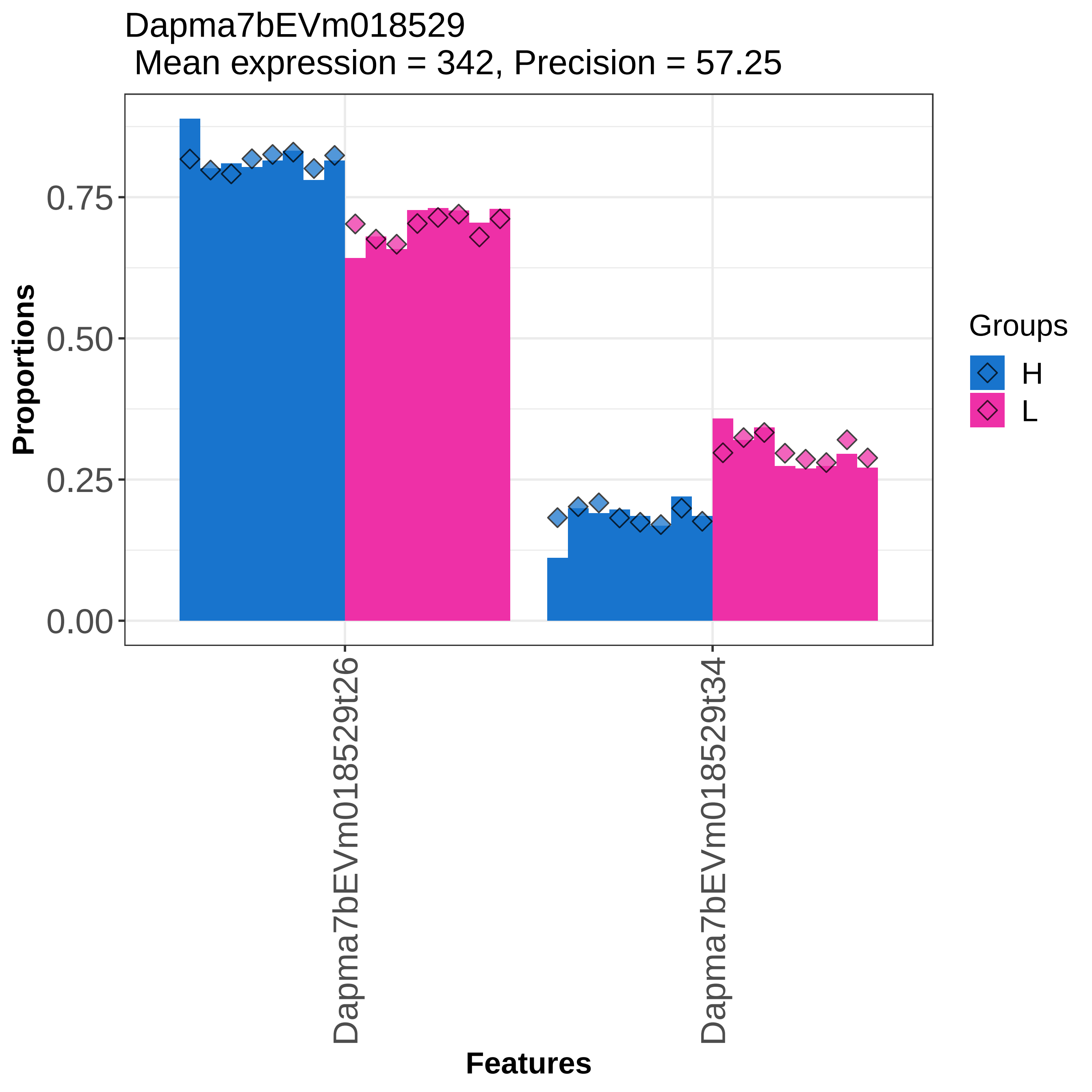

Dapma7bEVm018529
Mean expression = 342, Precision = 57.25
Proportions
0.00
0.25
0.50
0.75
Dapma7bEVm018529t26
Dapma7bEVm018529t34
Features
Groups
H
L

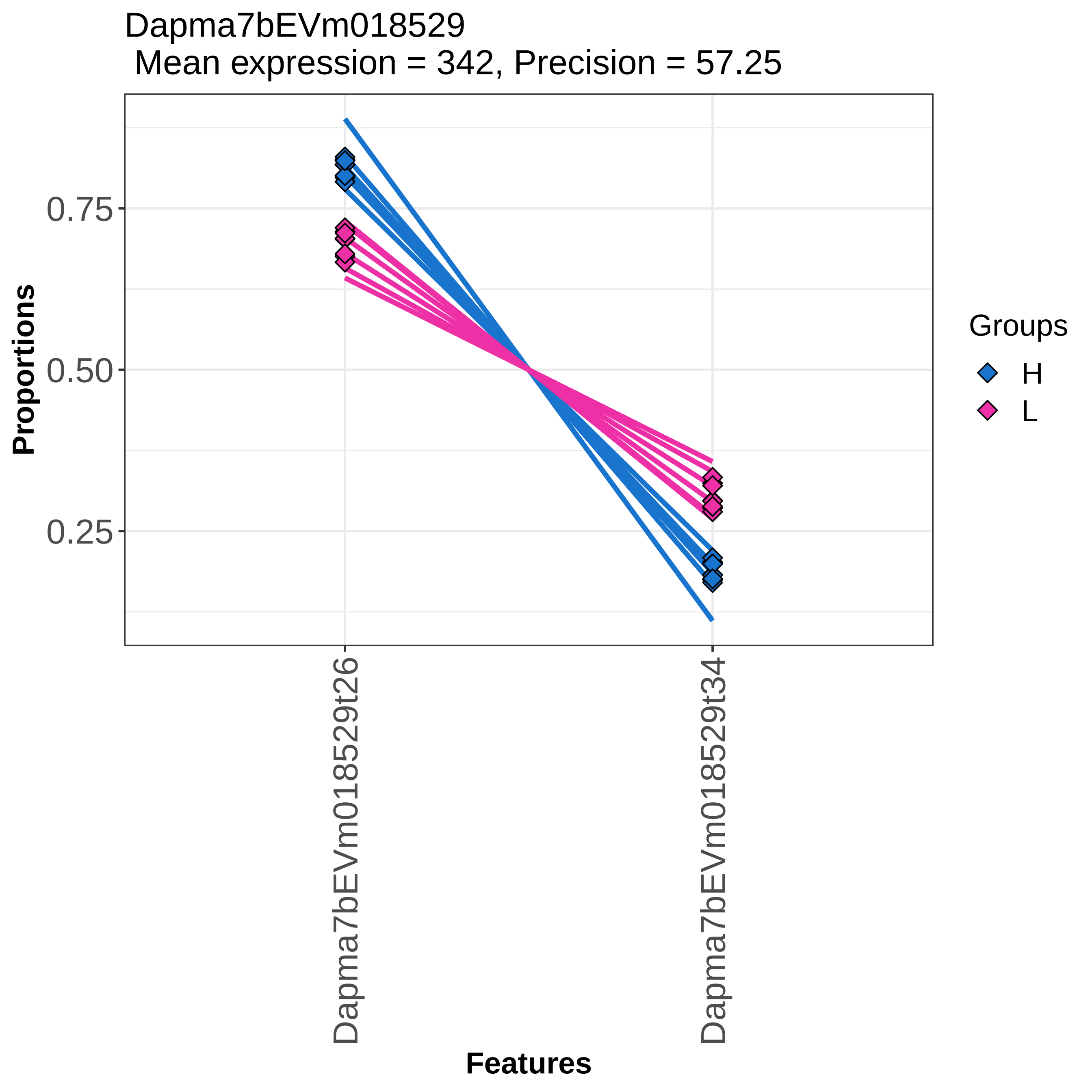

Dapma7bEVm018529
Mean expression = 342, Precision = 57.25
Proportions
0.75
0.50
0.25
Dapma7bEVm018529t26
Dapma7bEVm018529t34
Features
Groups
H
L

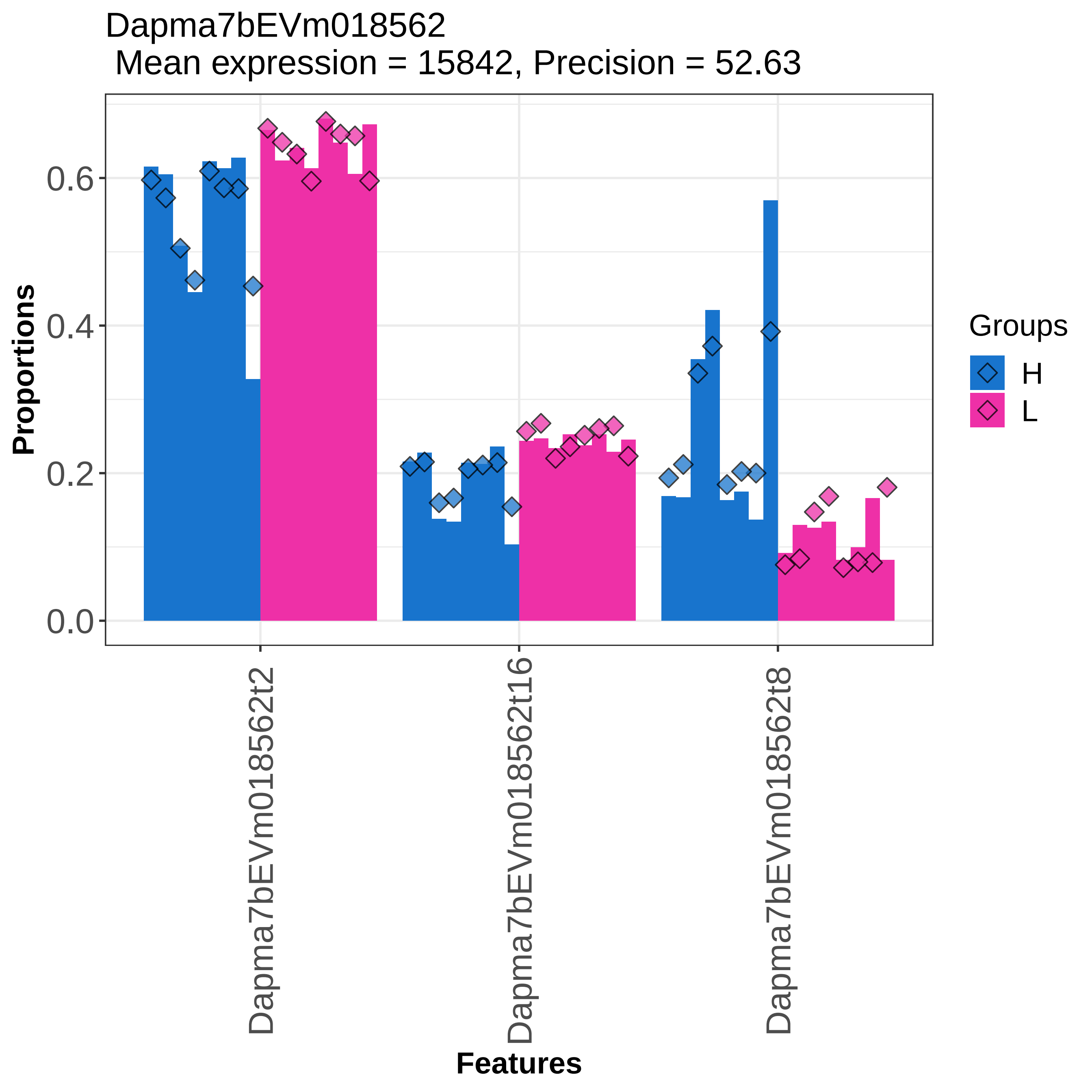
Dapma7bEVm018562
Mean expression = 15842, Precision = 52.63
Proportions
0.0
0.2
0.4
0.6
Features
Dapma7bEVm018562t2
Dapma7bEVm018562t16
Dapma7bEVm018562t8
Groups
H
L

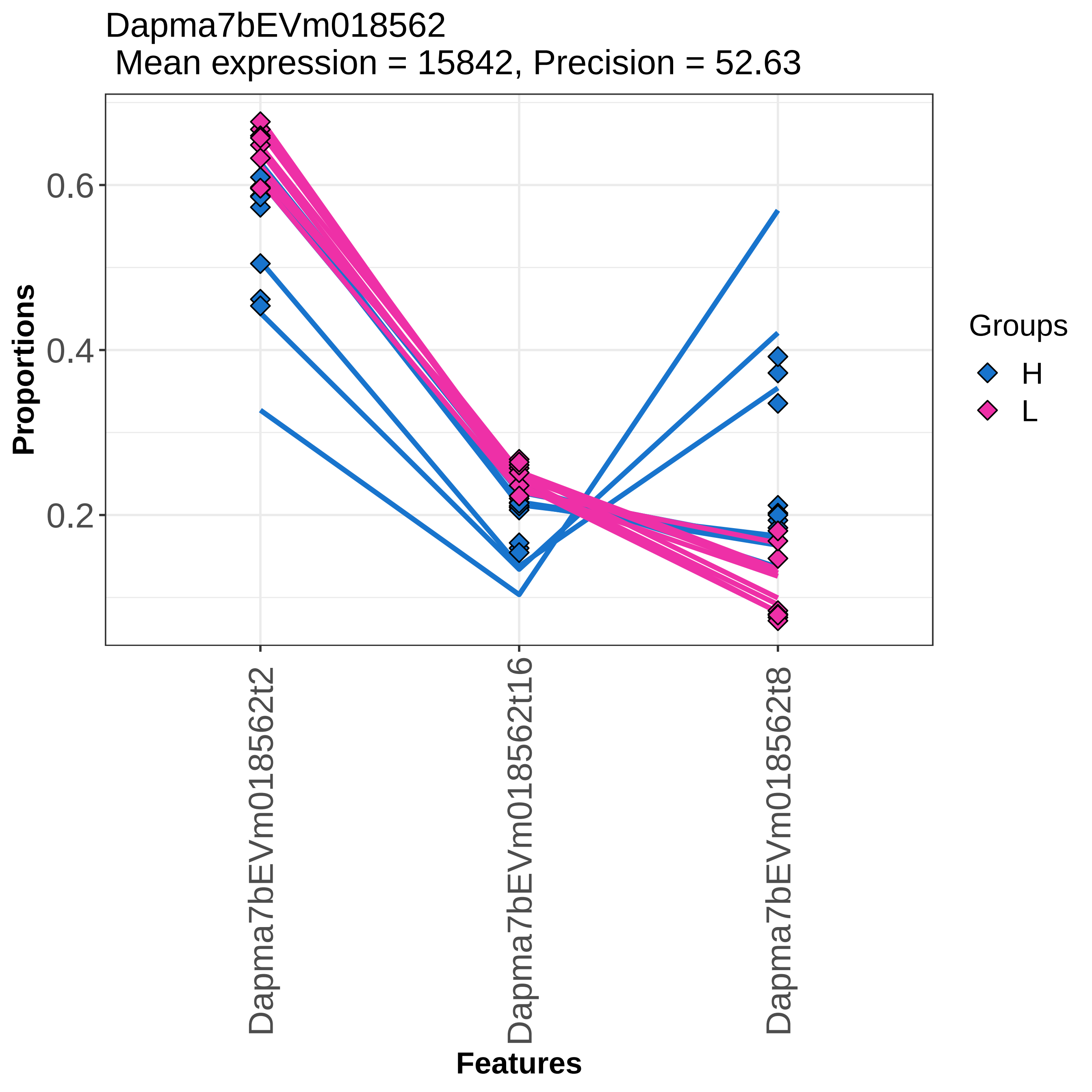

Dapma7bEVm018562
Mean expression = 15842, Precision = 52.63
Proportions
0.6
0.4
0.2
Groups
H
L
Dapma7bEVm018562t2
Dapma7bEVm018562t16
Dapma7bEVm018562t8
Features

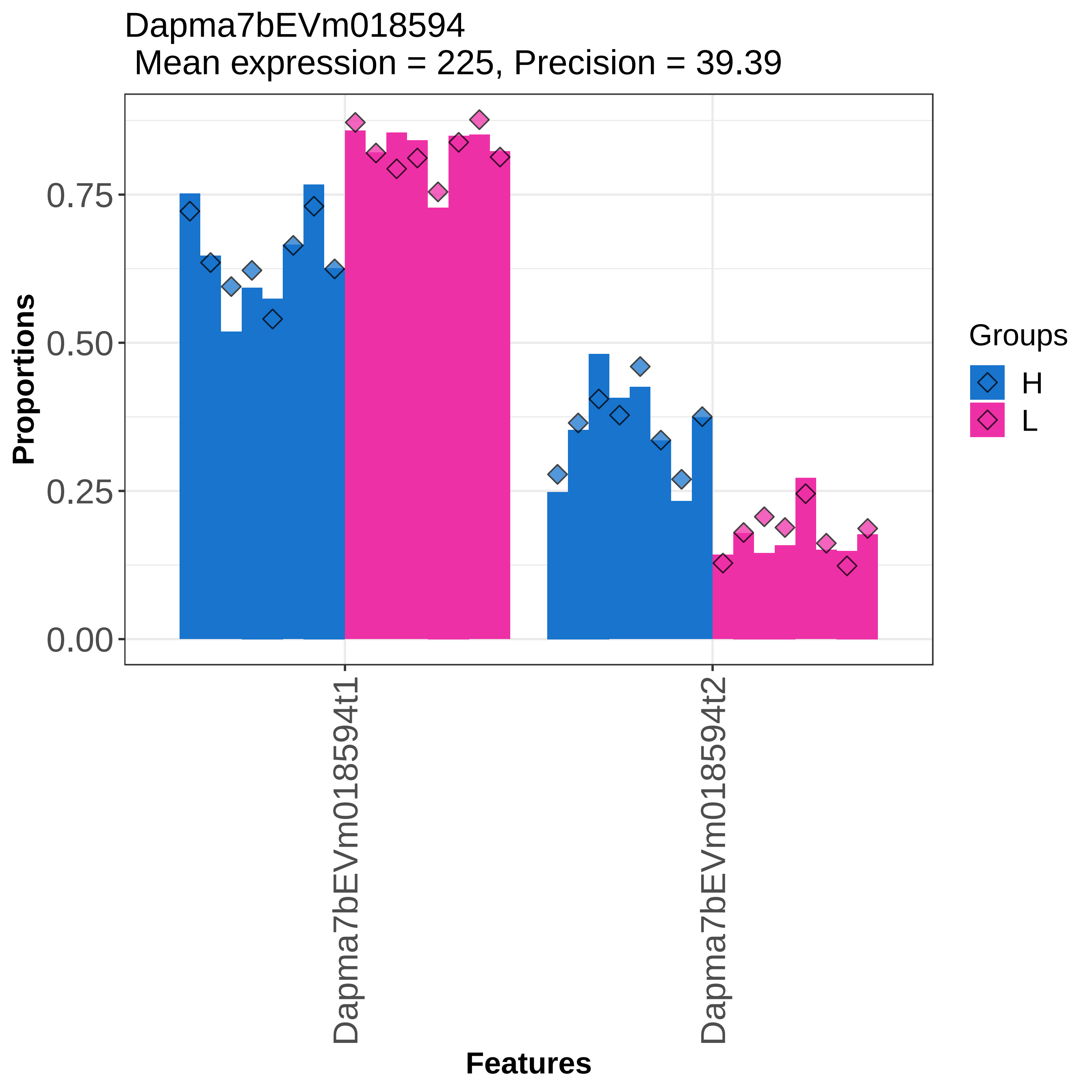
Dapma7bEVm018594
Mean expression = 225, Precision = 39.39
Proportions
0.75
0.50
0.25
0.00
Dapma7bEVm018594t1
Dapma7bEVm018594t2
Features
Groups
H
L

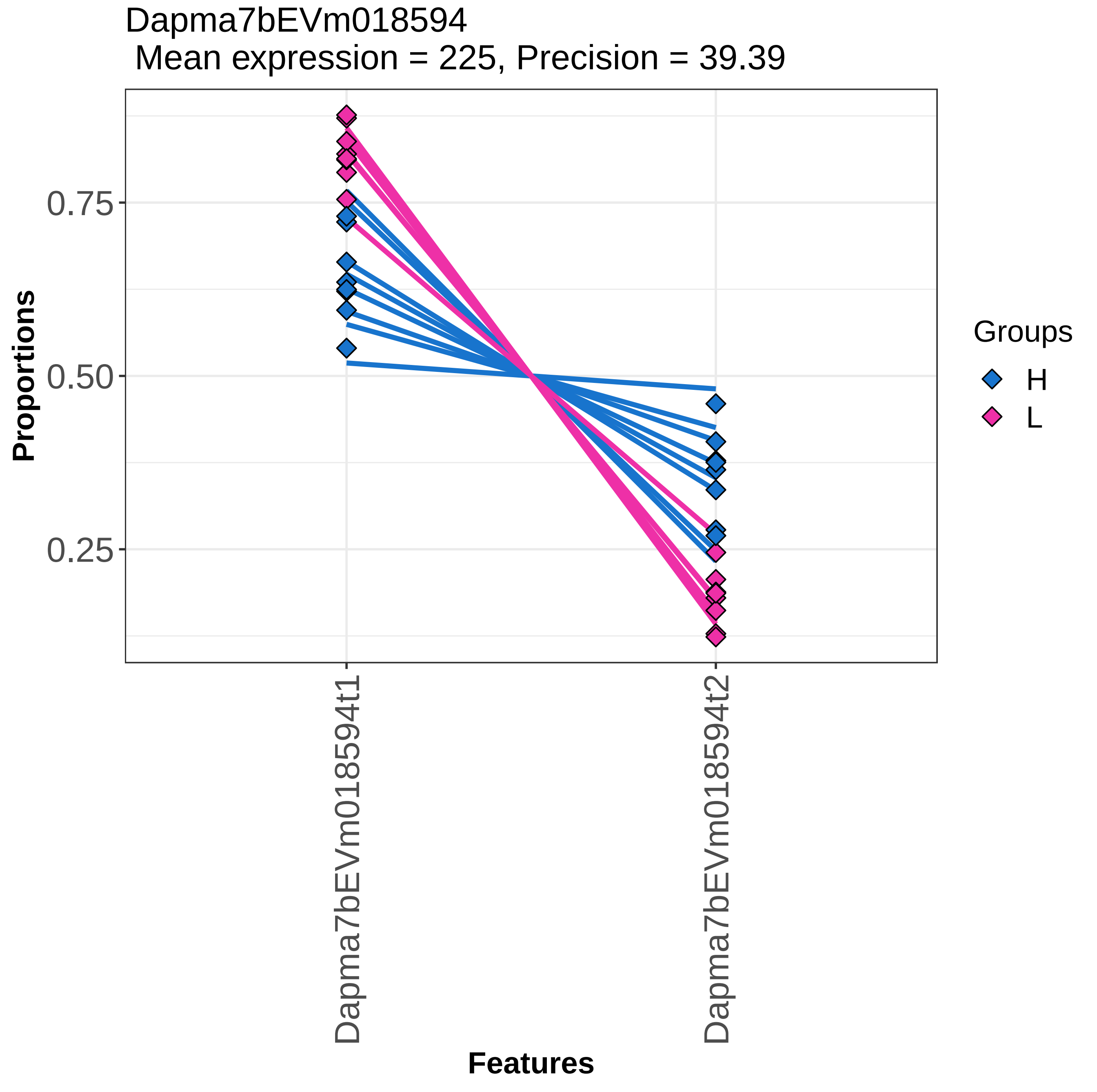

Dapma7bEVm018594
Mean expression = 225, Precision = 39.39
Proportions
0.75
0.50
0.25
Dapma7bEVm018594t1
Dapma7bEVm018594t2
Features
Groups
H
L

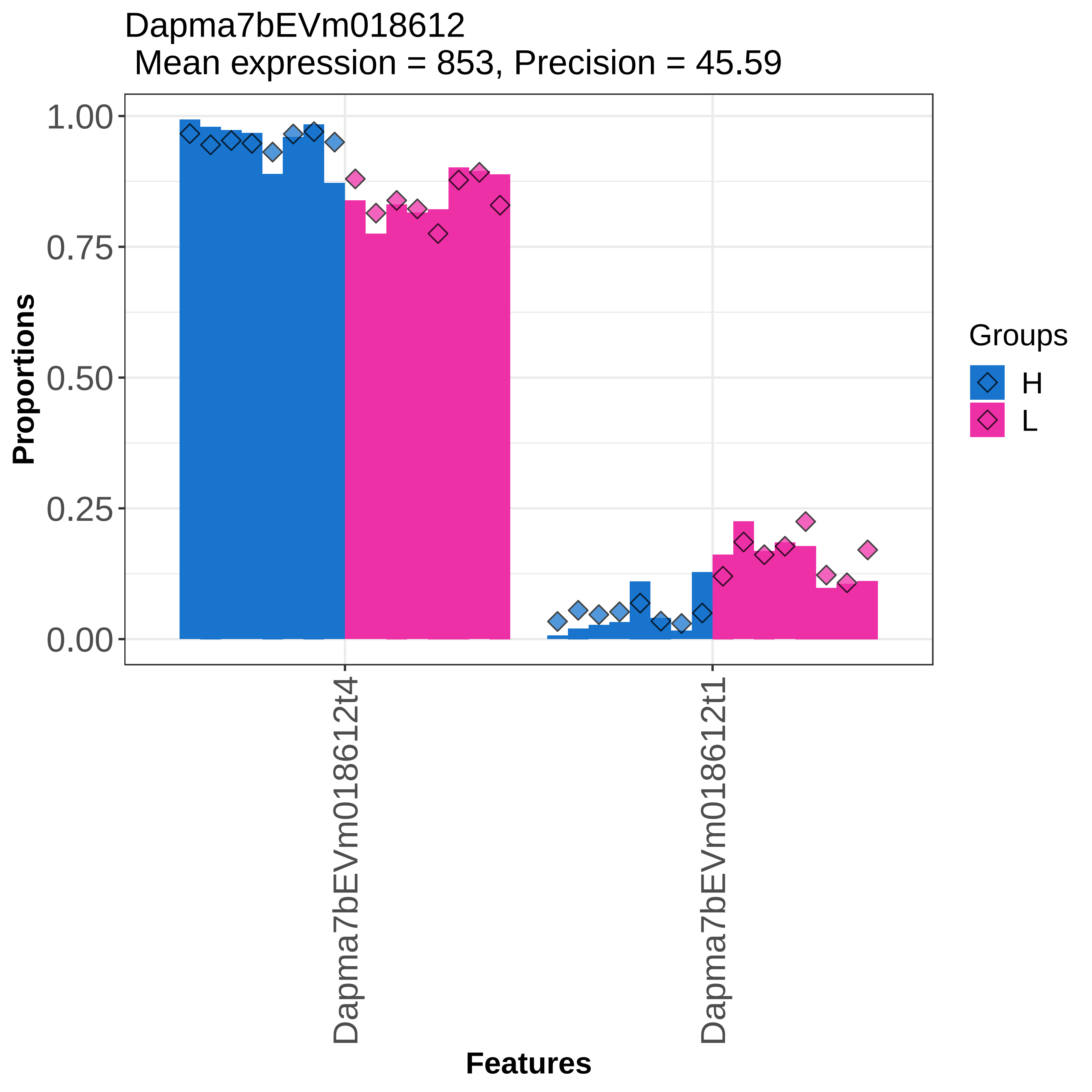

Dapma7bEVm018612
Mean expression = 853, Precision = 45.59
Proportions
1.00
0.75
0.50
0.25
0.00
Dapma7bEVm018612t4
Dapma7bEVm018612t1
Features
Groups
H
L

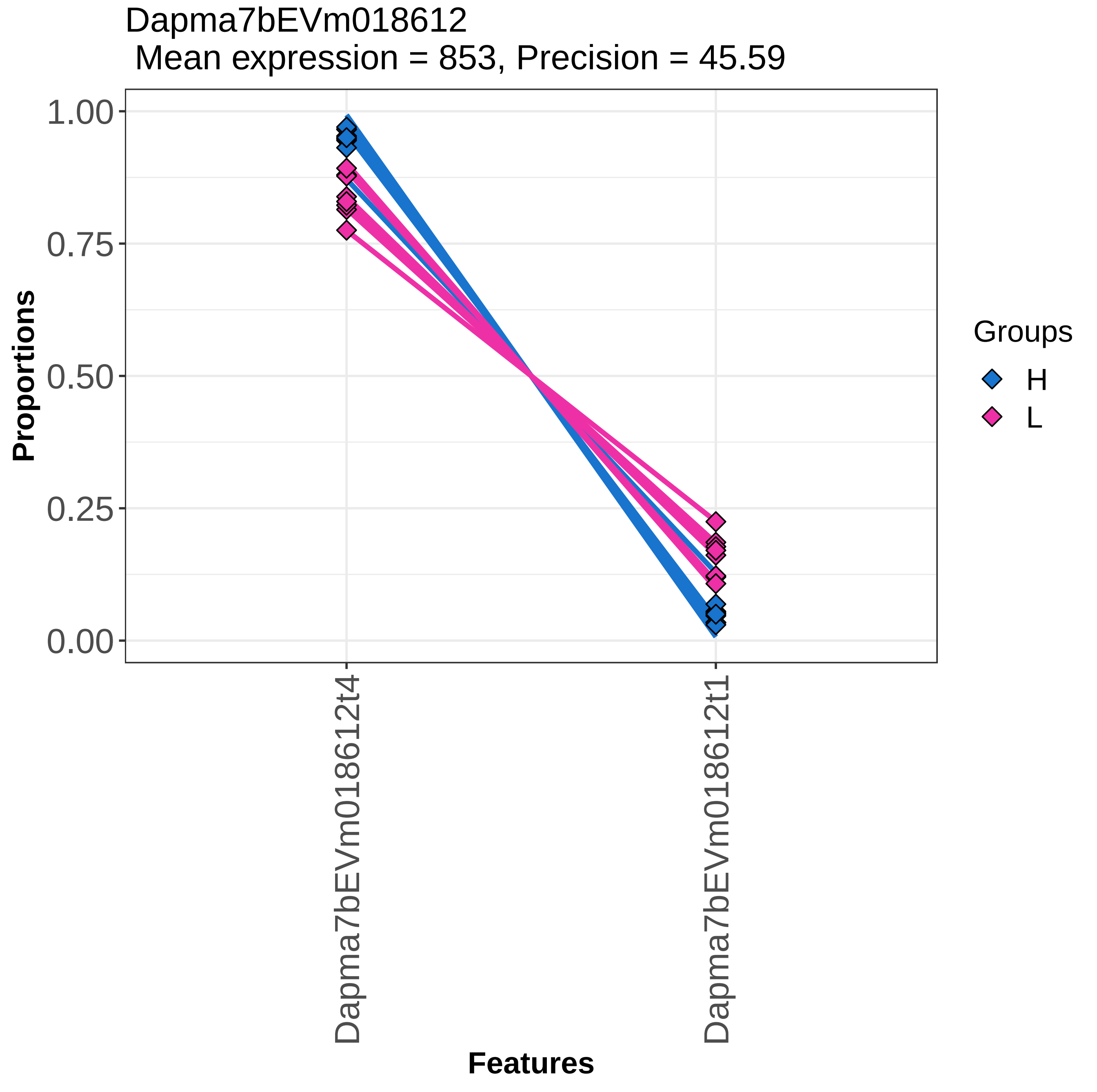

Dapma7bEVm018612
Mean expression = 853, Precision = 45.59
Proportions
1.00
0.75
0.50
0.25
0.00
Dapma7bEVm018612t4
Dapma7bEVm018612t1
Features
Groups
H
L

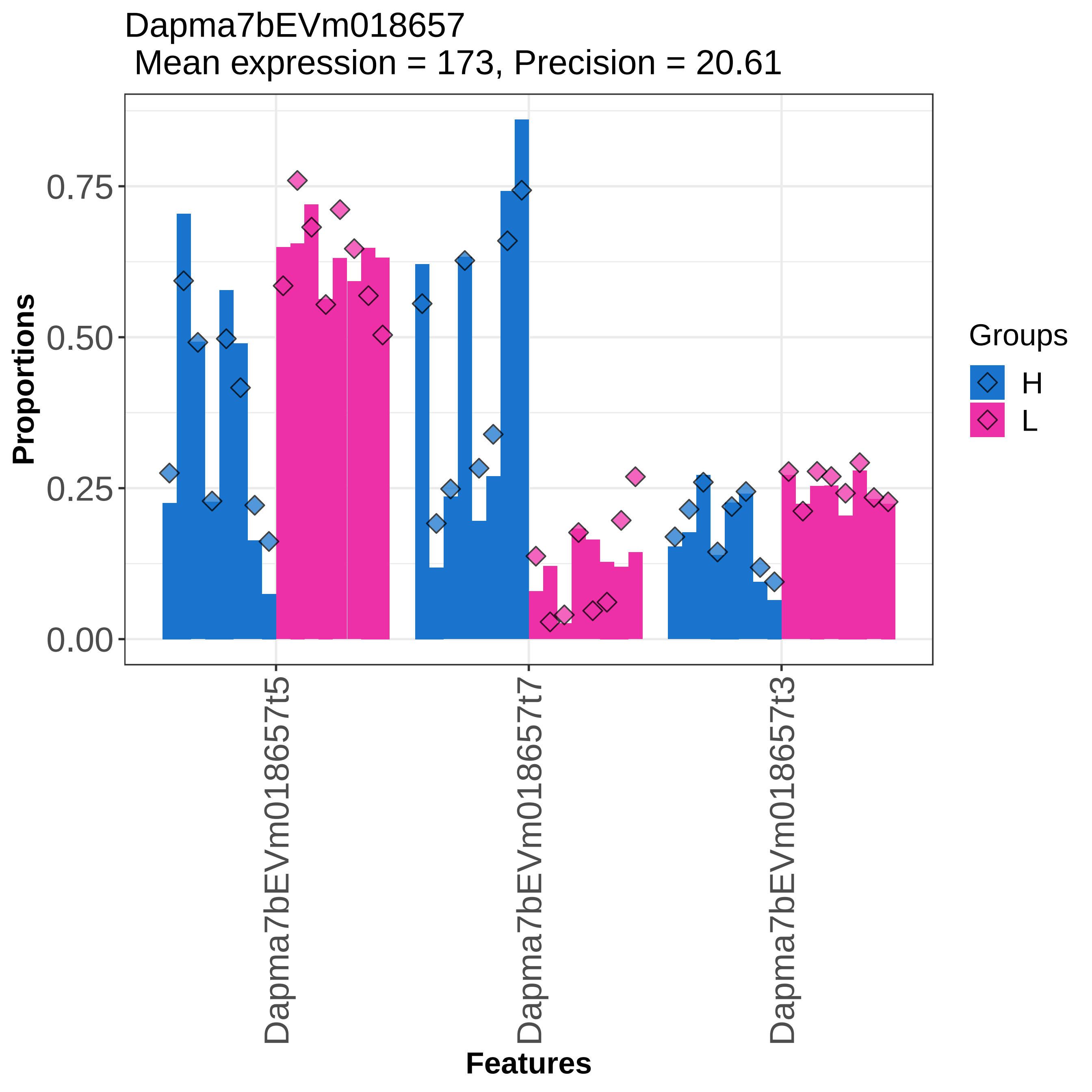

Dapma7bEVm018657
Mean expression = 173, Precision = 20.61
Proportions
0.00
0.25
0.50
0.75
Features
Dapma7bEVm018657t5
Dapma7bEVm018657t7
Dapma7bEVm018657t3
Groups
H
L

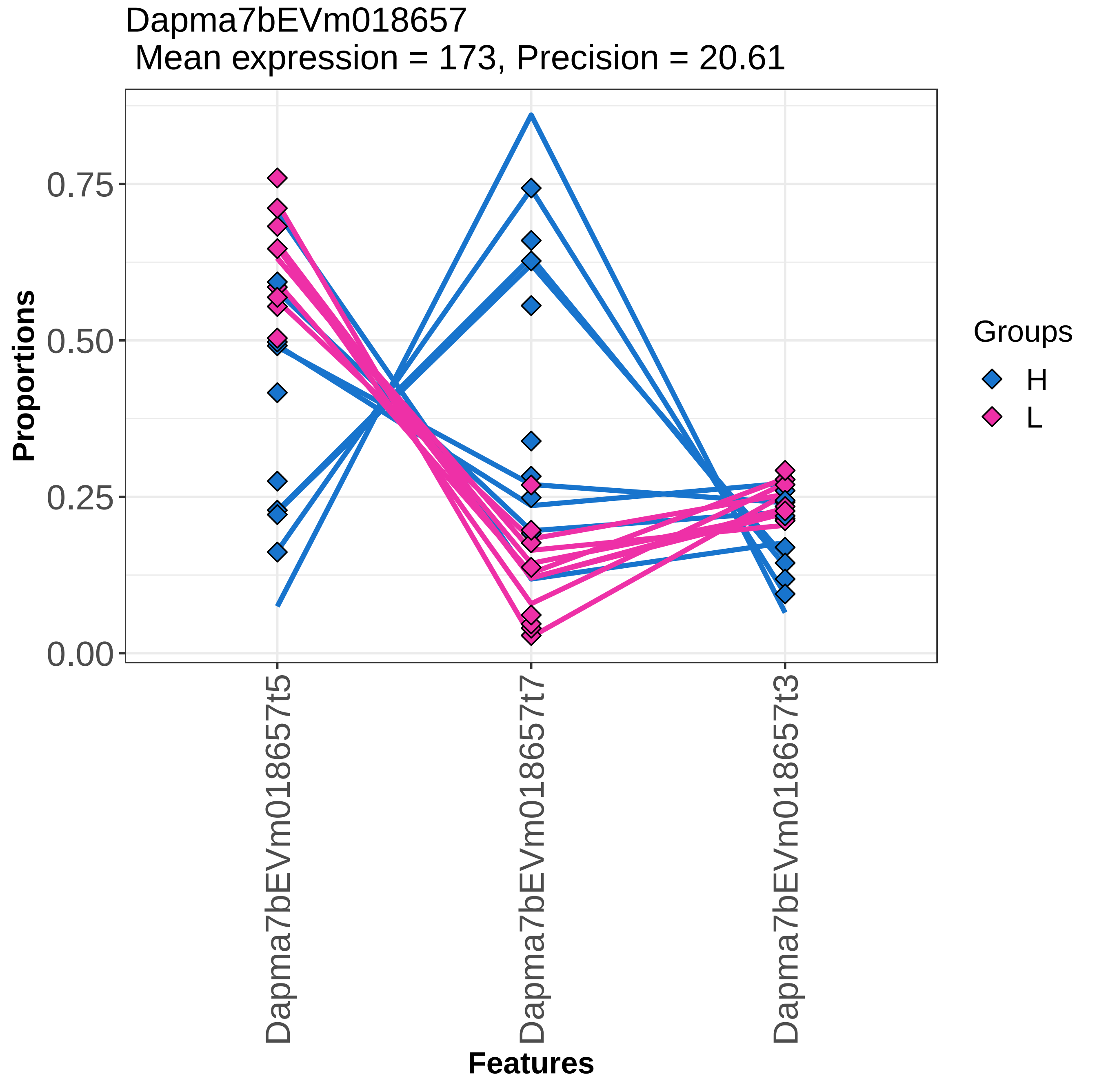
Dapma7bEVm018657
Mean expression = 173, Precision = 20.61
Proportions
0.00
0.25
0.50
0.75
Dapma7bEVm018657t5
Dapma7bEVm018657t7
Dapma7bEVm018657t3
Features
Groups
H
L

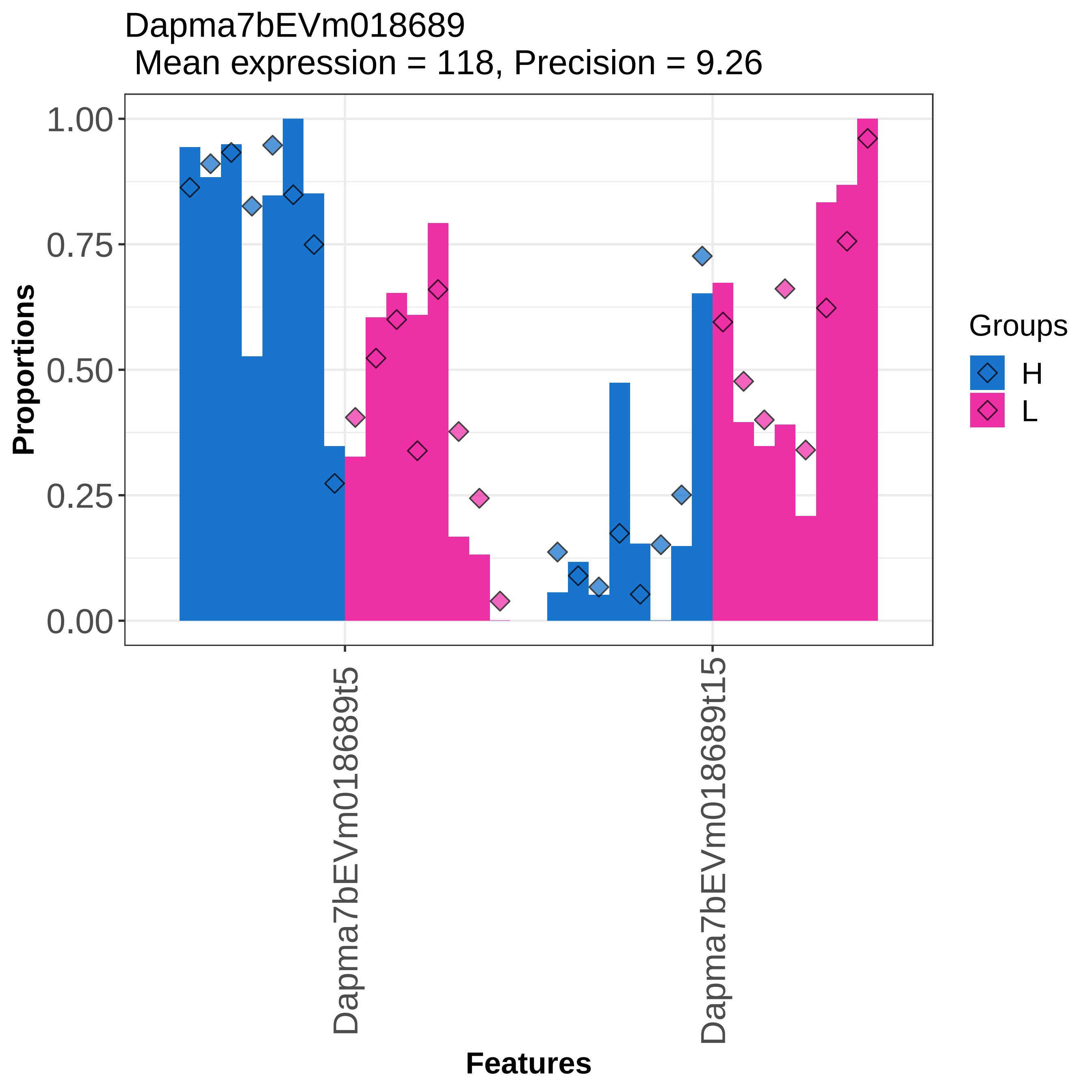
Dapma7bEVm018689
Mean expression = 118, Precision = 9.26
Proportions
1.00
0.75
0.50
0.25
0.00
Dapma7bEVm018689t5
Dapma7bEVm018689t15
Features
Groups
H
L

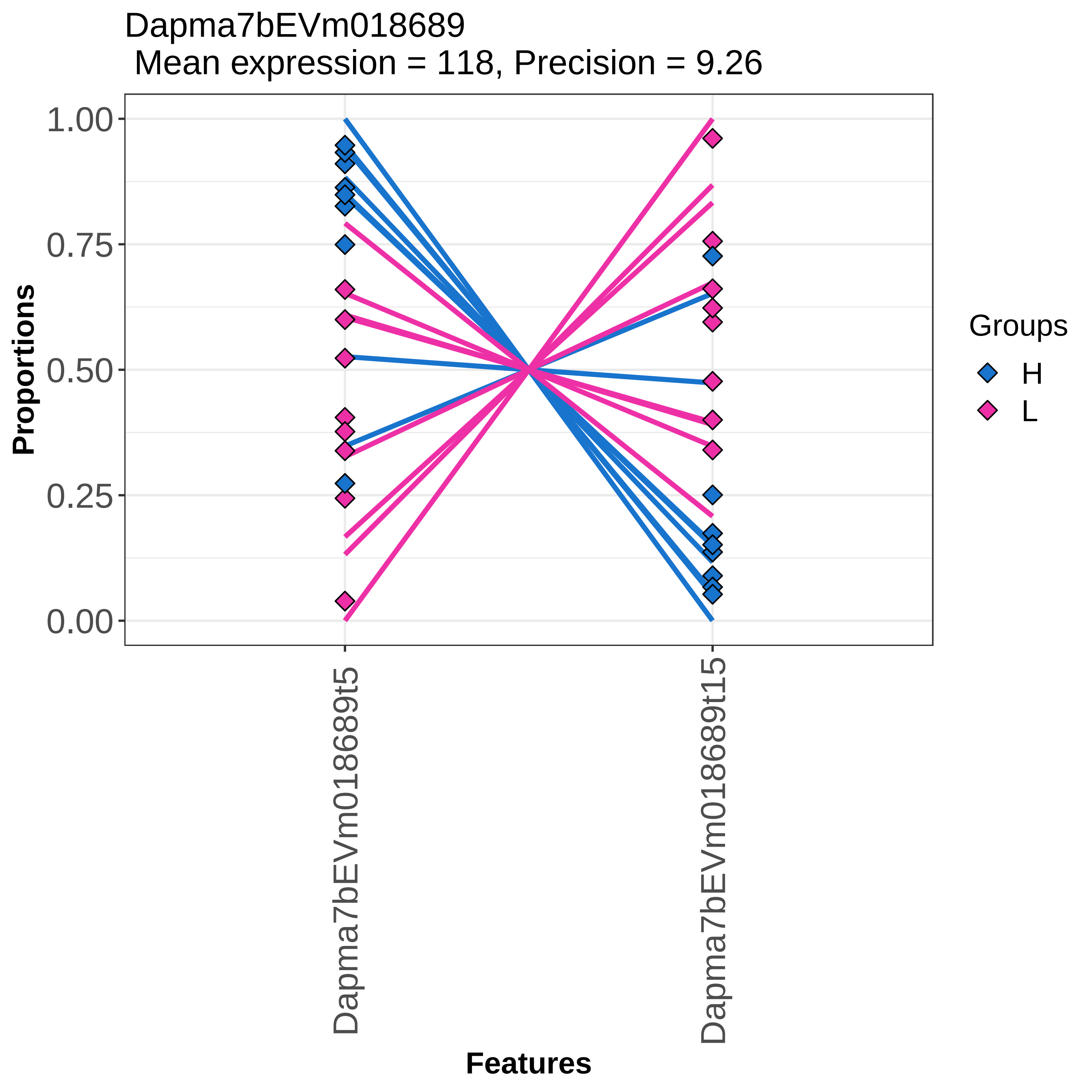
Dapma7bEVm018689
Mean expression = 118, Precision = 9.26
Proportions
1.00
0.75
0.50
0.25
0.00
Dapma7bEVm018689t5
Dapma7bEVm018689t15
Features
Groups
H
L

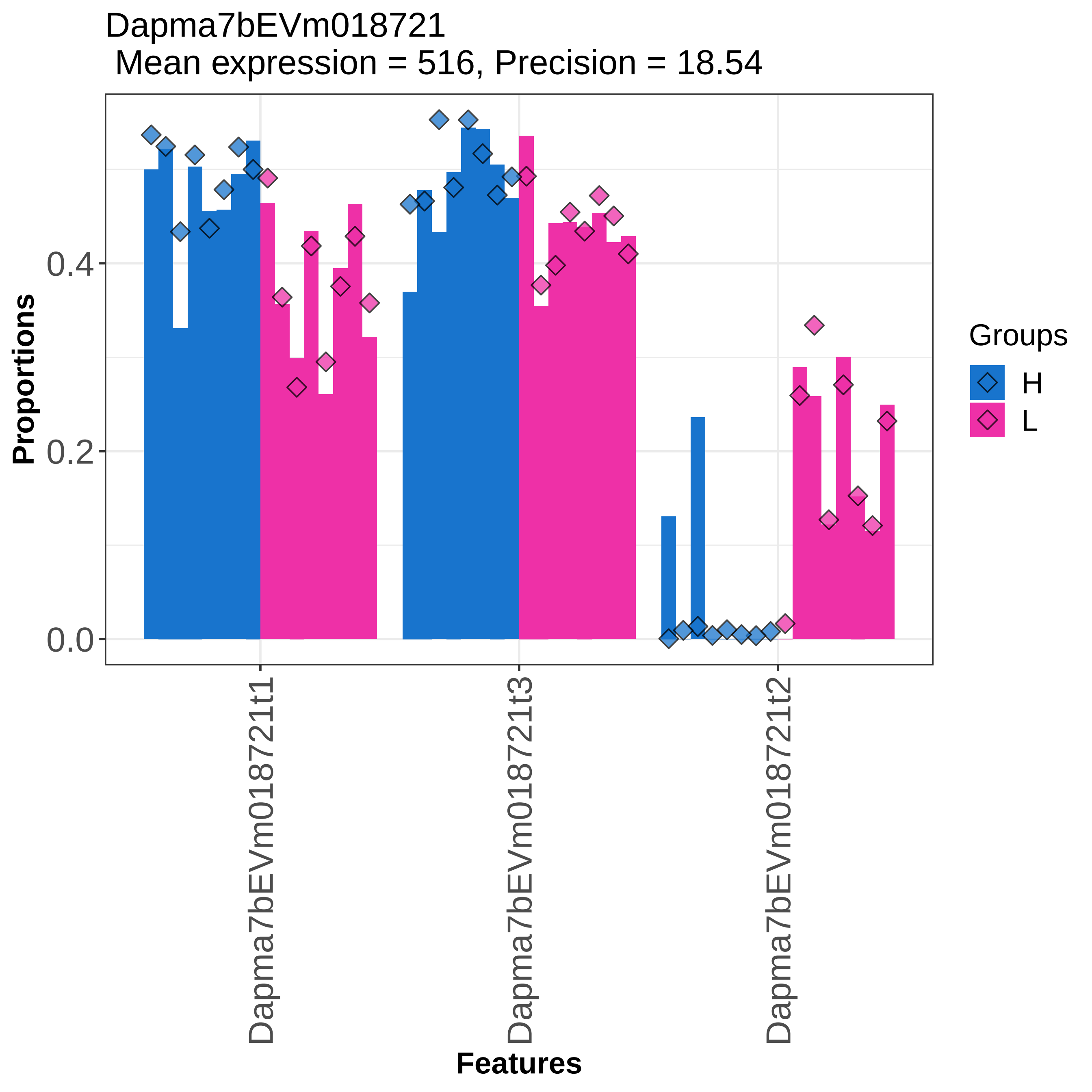
Dapma7bEVm018721
Mean expression = 516, Precision = 18.54
Proportions
0.0
0.2
0.4
Features
Dapma7bEVm018721t1
Dapma7bEVm018721t3
Dapma7bEVm018721t2
Groups
H
L

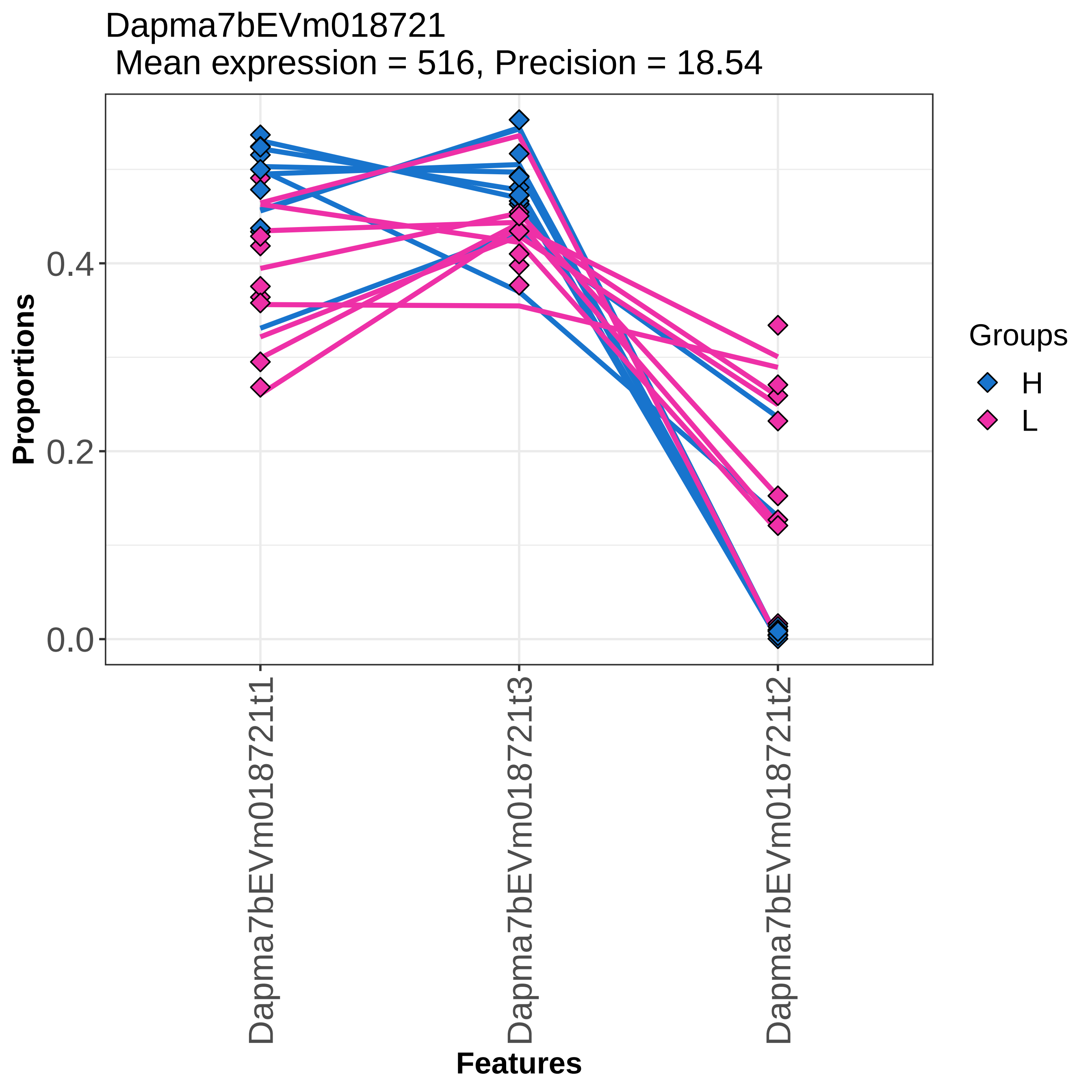

Dapma7bEVm018721
Mean expression = 516, Precision = 18.54
Proportions
0.0
0.2
0.4
Dapma7bEVm018721t1
Dapma7bEVm018721t3
Dapma7bEVm018721t2
Features
Groups
H
L

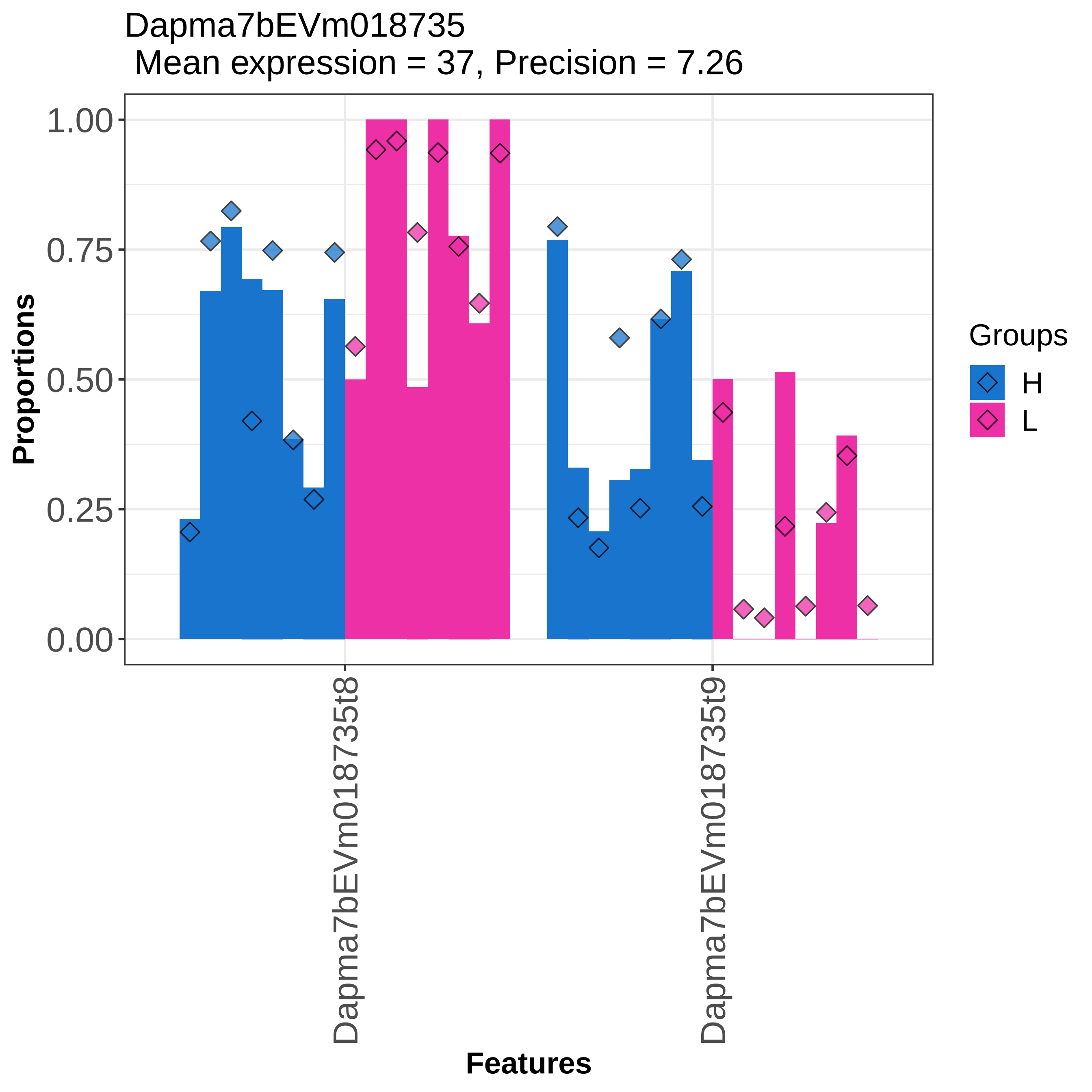

Dapma7bEVm018735
Mean expression = 37, Precision = 7.26
Proportions
1.00
0.75
0.50
0.25
0.00
Dapma7bEVm018735t8
Dapma7bEVm018735t9
Features
Groups
H
L

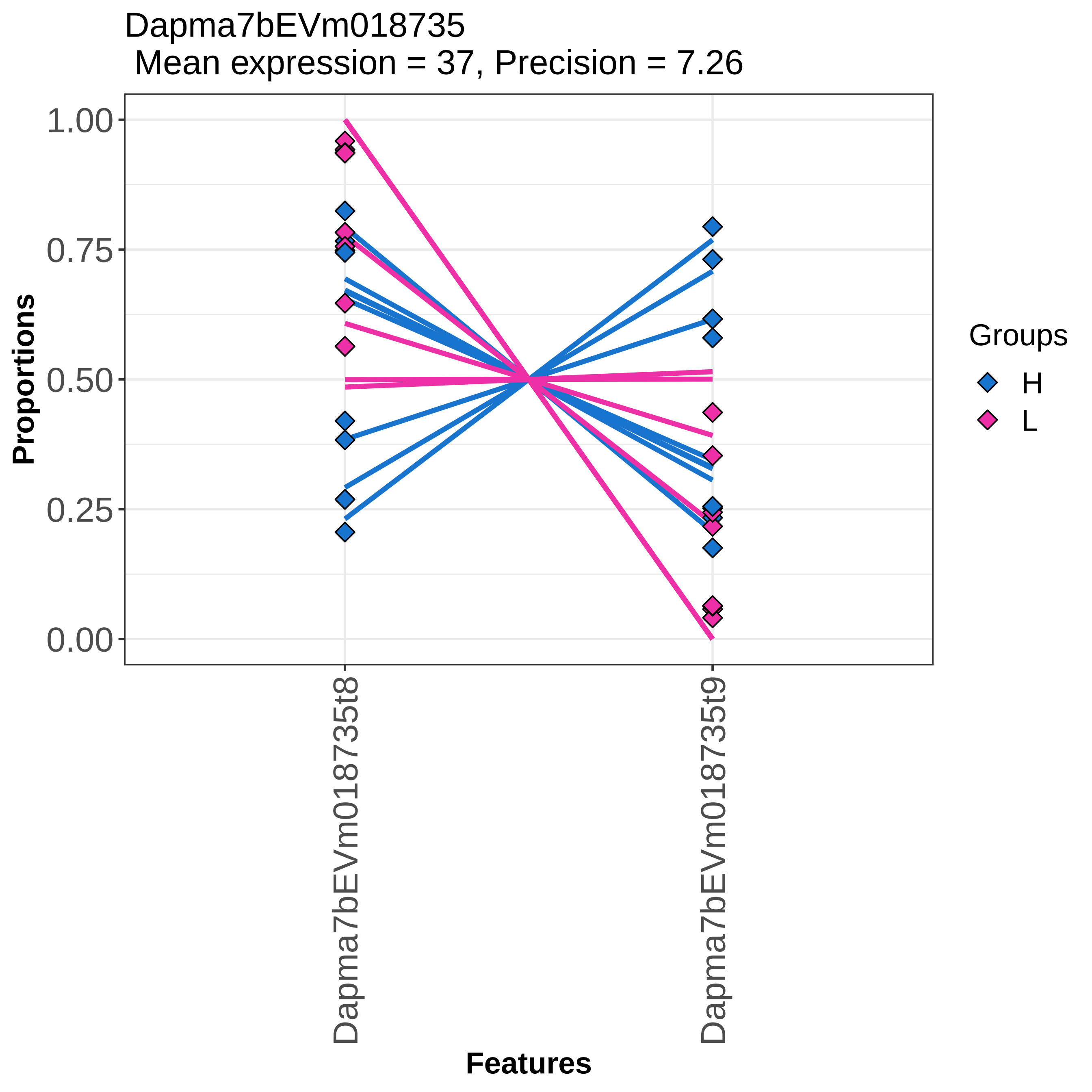

Dapma7bEVm018735
Mean expression = 37, Precision = 7.26
Proportions
1.00
0.75
0.50
0.25
0.00
Dapma7bEVm018735t8
Dapma7bEVm018735t9
Features
Groups
H
L

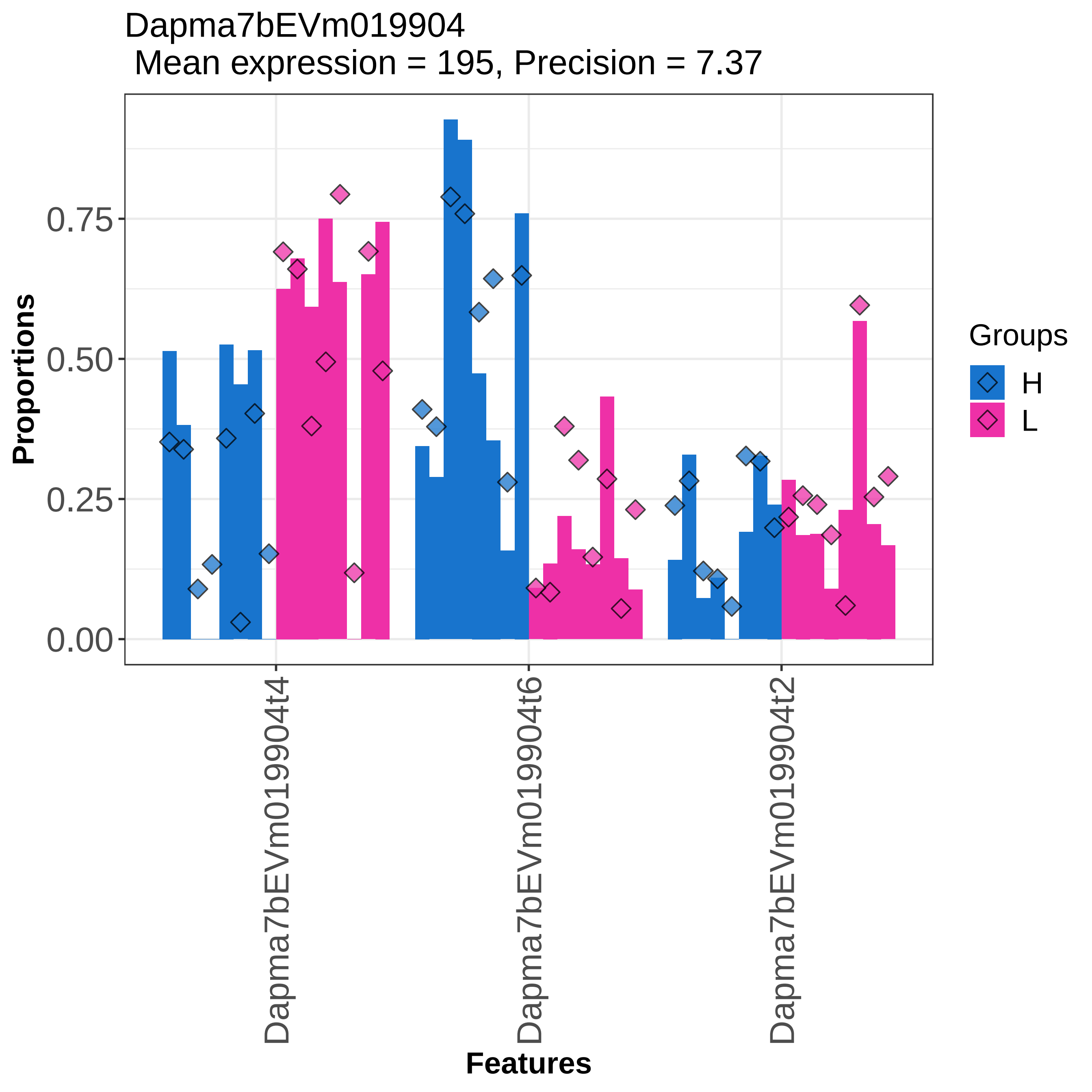

Dapma7bEVm019904
Mean expression = 195, Precision = 7.37
Proportions
0.00
0.25
0.50
0.75
Dapma7bEVm019904t4
Dapma7bEVm019904t6
Dapma7bEVm019904t2
Features
Groups
H
L

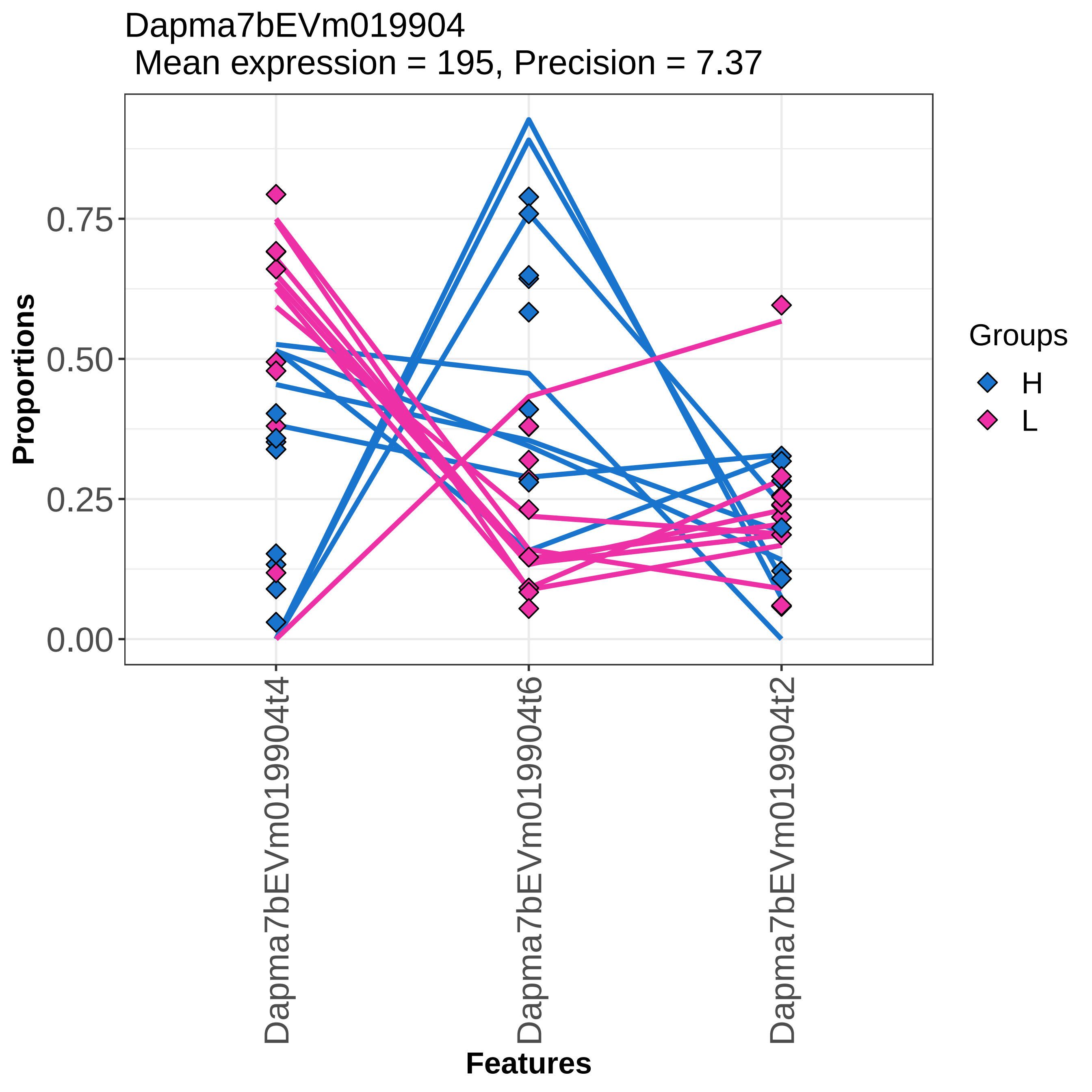
Dapma7bEVm019904
Mean expression = 195, Precision = 7.37
Proportions
0.75
0.50
0.25
0.00
Dapma7bEVm019904t4
Dapma7bEVm019904t6
Dapma7bEVm019904t2
Features
Groups
H
L

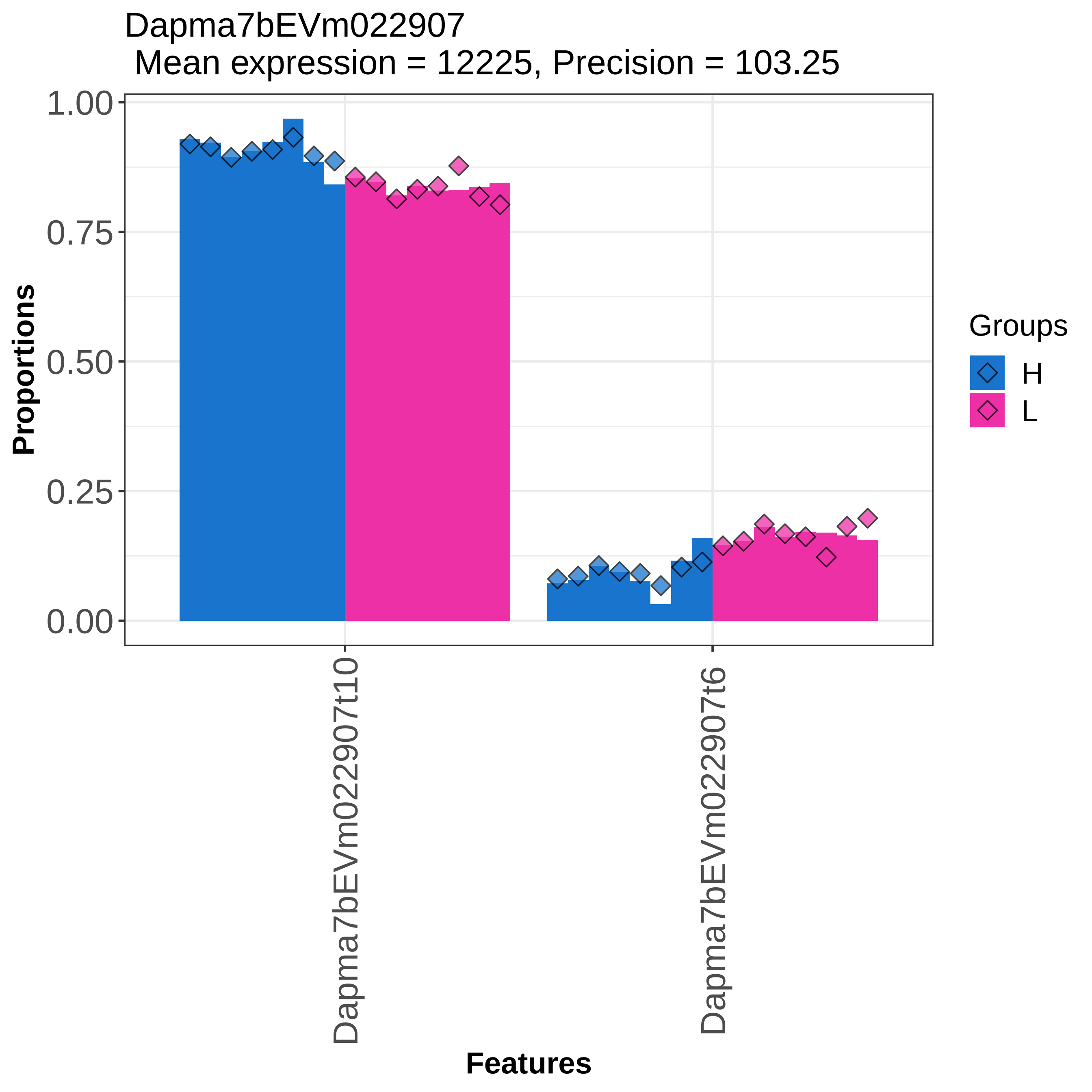

Dapma7bEVm022907
Mean expression = 12225, Precision = 103.25
Proportions
1.00
0.75
0.50
0.25
0.00
Dapma7bEVm022907t10
Dapma7bEVm022907t6
Features
Groups
H
L

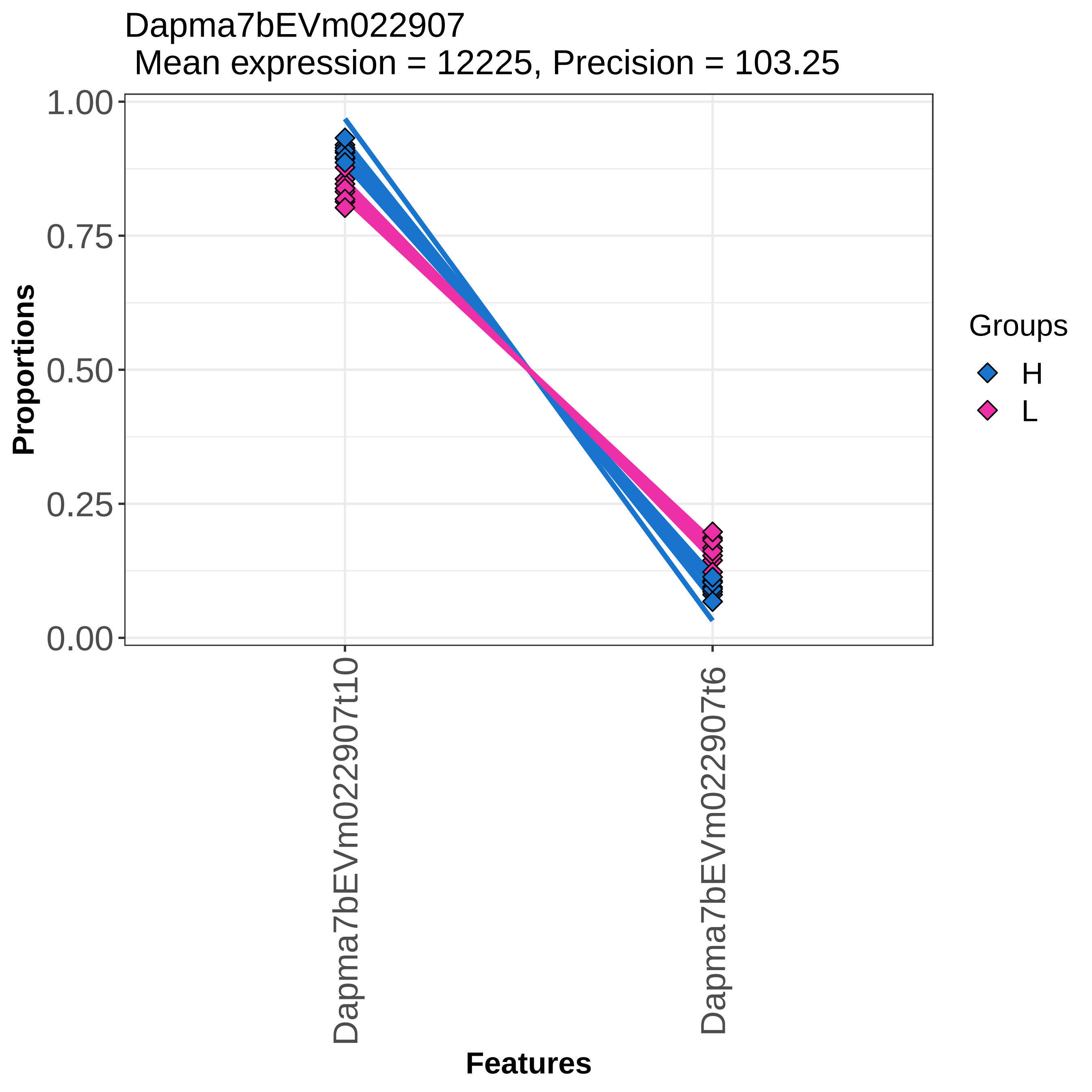
Dapma7bEVm022907
Mean expression = 12225, Precision = 103.25
Proportions
1.00
0.75
0.50
0.25
0.00
Dapma7bEVm022907t10
Dapma7bEVm022907t6
Features
Groups
H
L

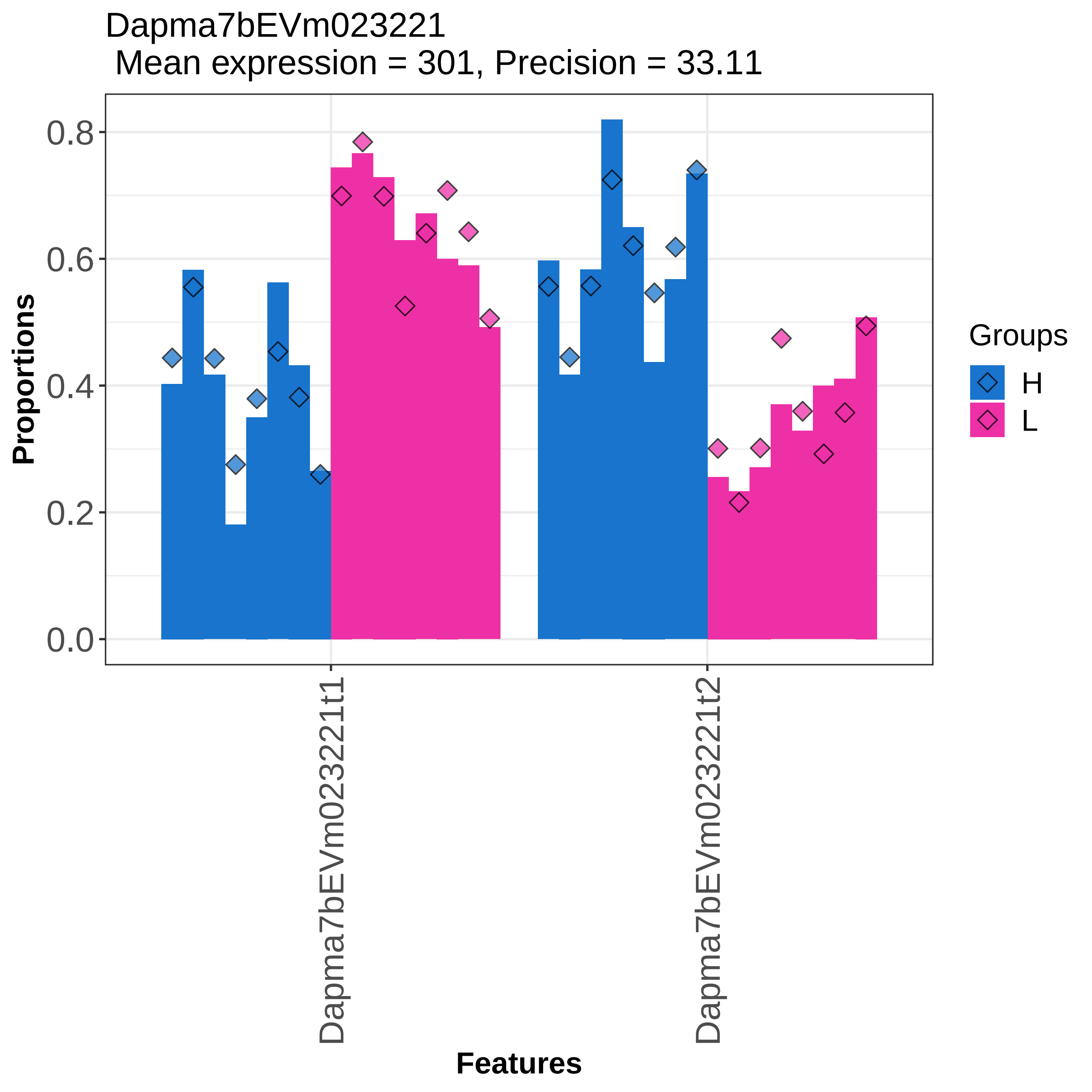

Dapma7bEVm023221
Mean expression = 301, Precision = 33.11
Proportions
0.0
0.2
0.4
0.6
0.8
Dapma7bEVm023221t1
Dapma7bEVm023221t2
Features
Groups
H
L

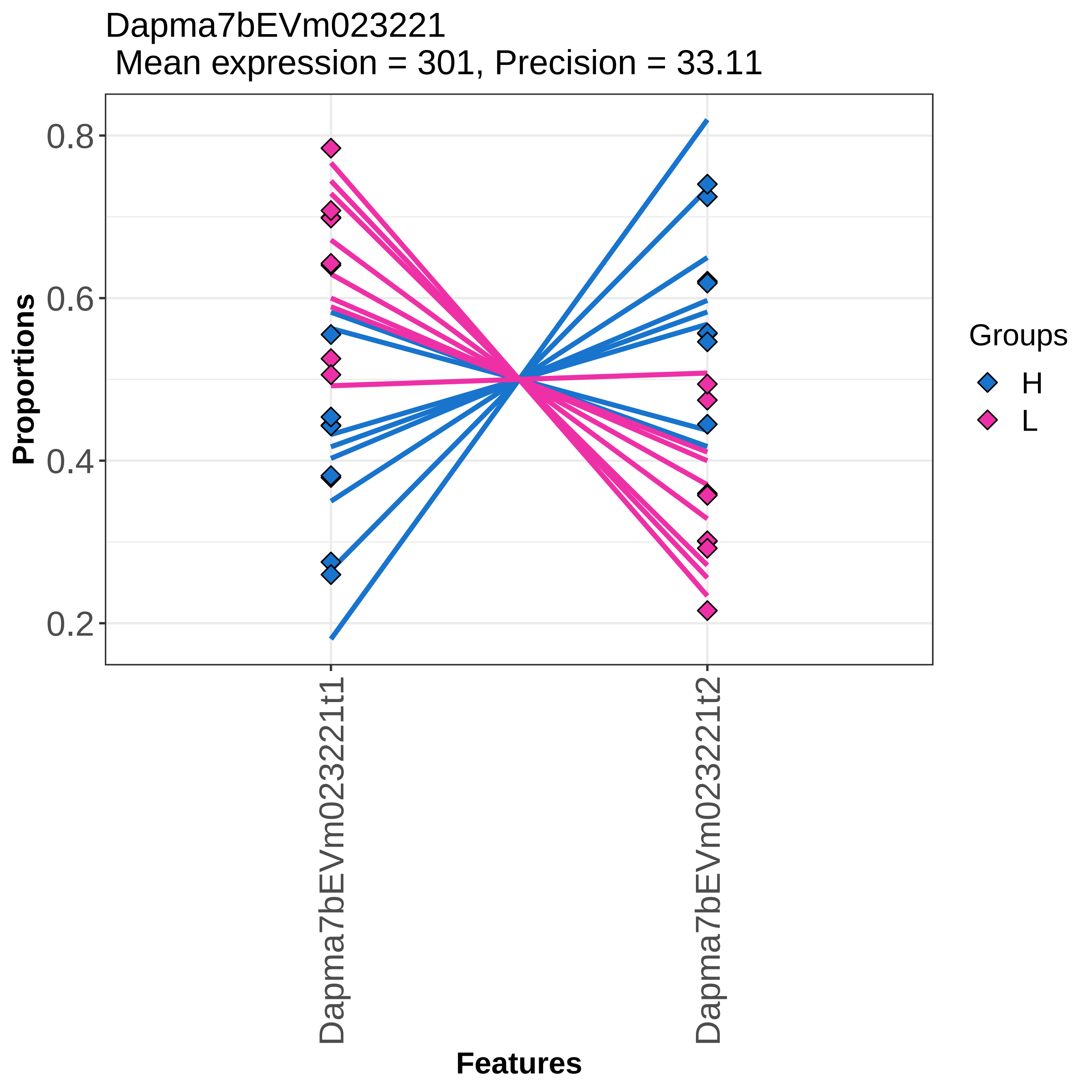

Dapma7bEVm023221
Mean expression = 301, Precision = 33.11
Proportions
0.8
0.6
0.4
0.2
Dapma7bEVm023221t1
Dapma7bEVm023221t2
Features
Groups
H
L

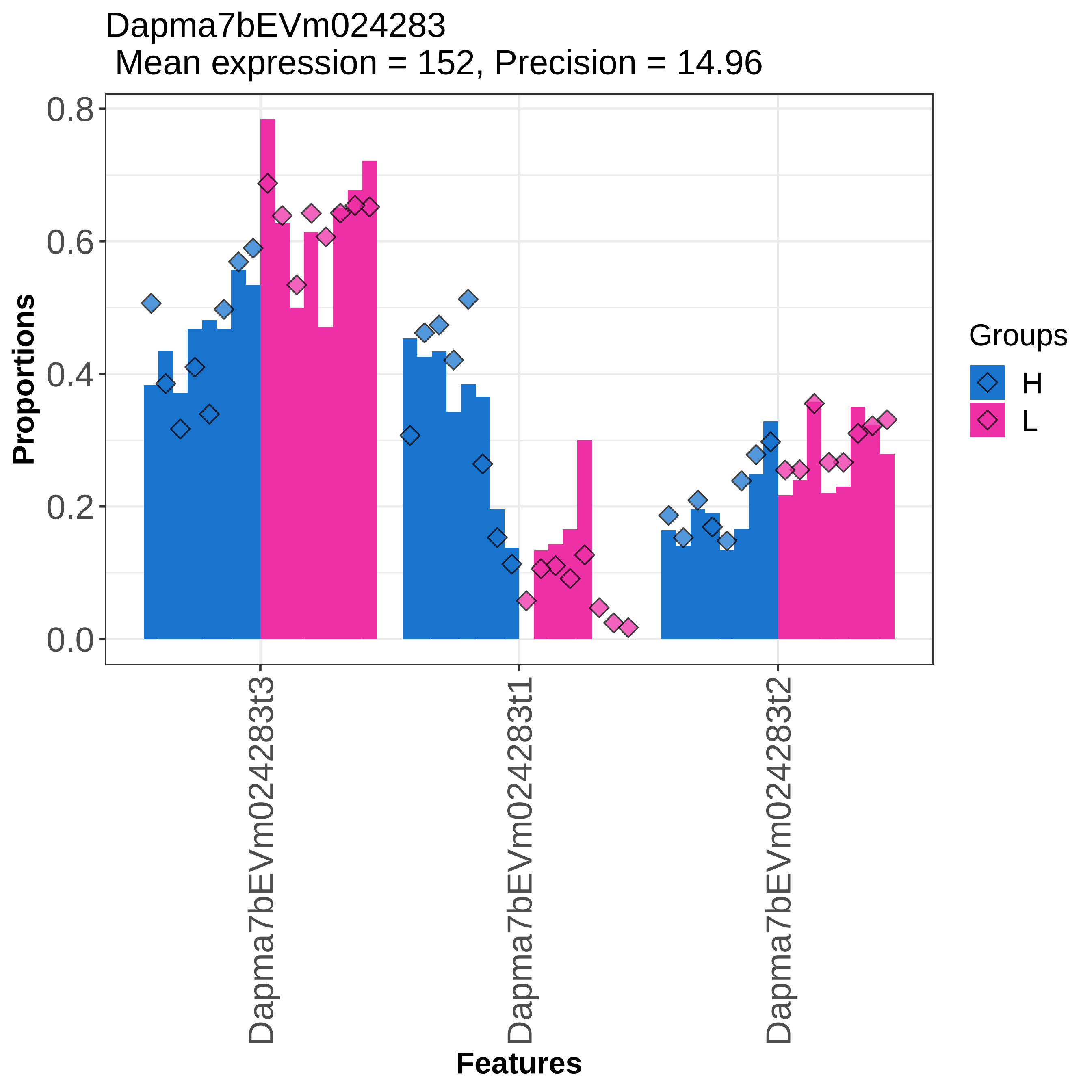

Dapma7bEVm024283
Mean expression = 152, Precision = 14.96
Proportions
0.0
0.2
0.4
0.6
0.8
Dapma7bEVm024283t3
Dapma7bEVm024283t1
Dapma7bEVm024283t2
Features
Groups
H
L

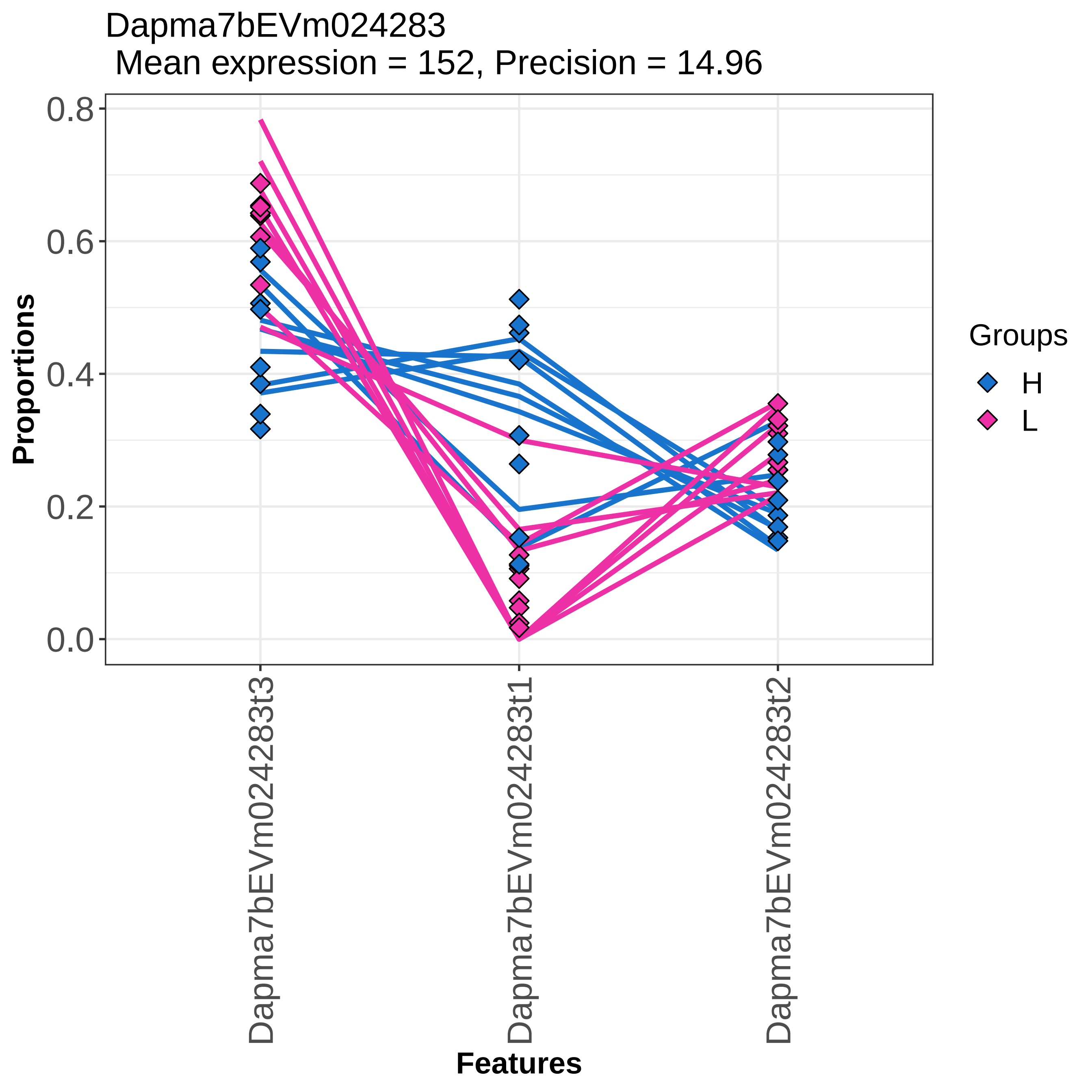

Dapma7bEVm024283
Mean expression = 152, Precision = 14.96
Proportions
0.8
0.6
0.4
0.2
0.0
Dapma7bEVm024283t3
Dapma7bEVm024283t1
Dapma7bEVm024283t2
Features
Groups
H
L

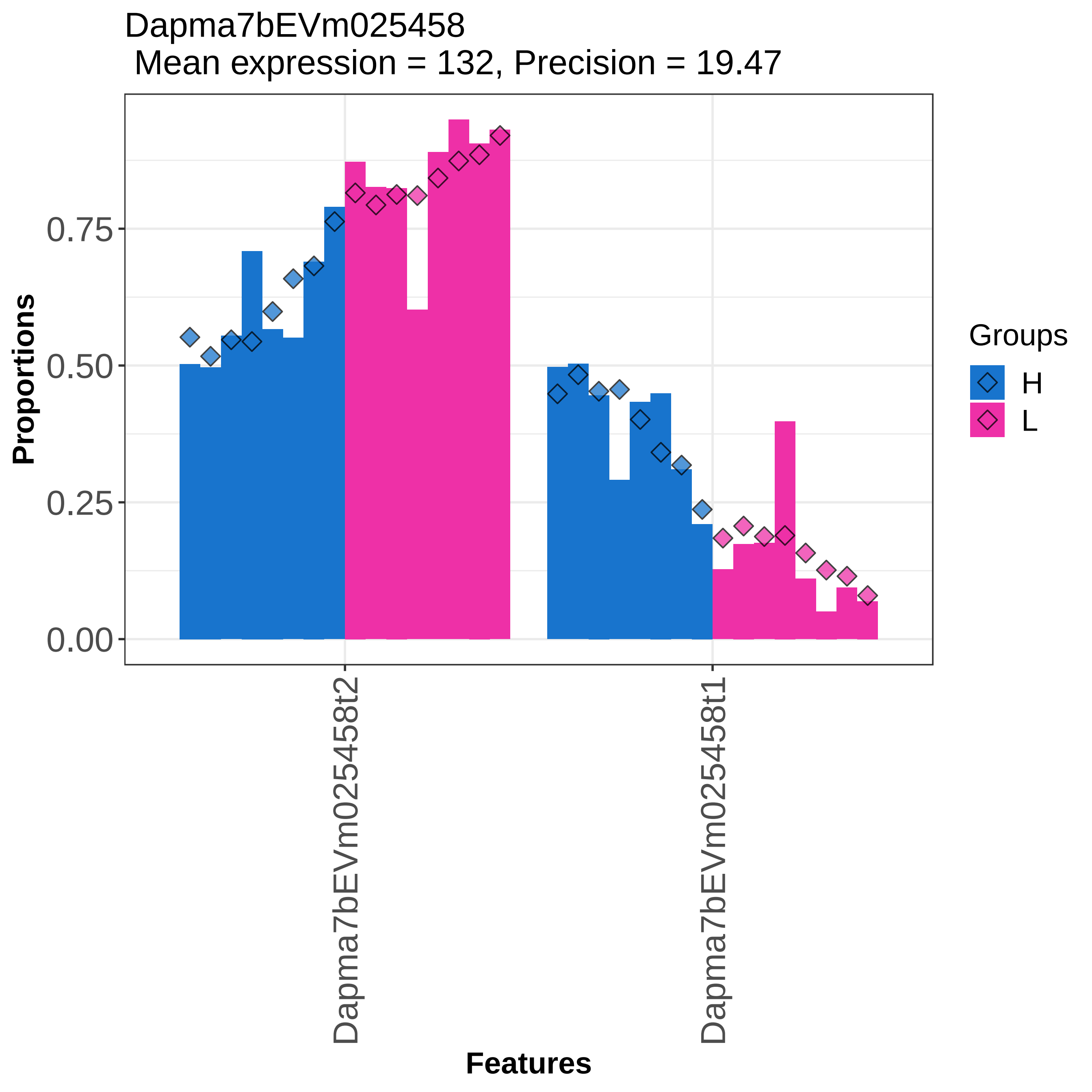

Dapma7bEVm025458
Mean expression = 132, Precision = 19.47
Proportions
0.75
0.50
0.25
0.00
Dapma7bEVm025458t2
Dapma7bEVm025458t1
Features
Groups
H
L

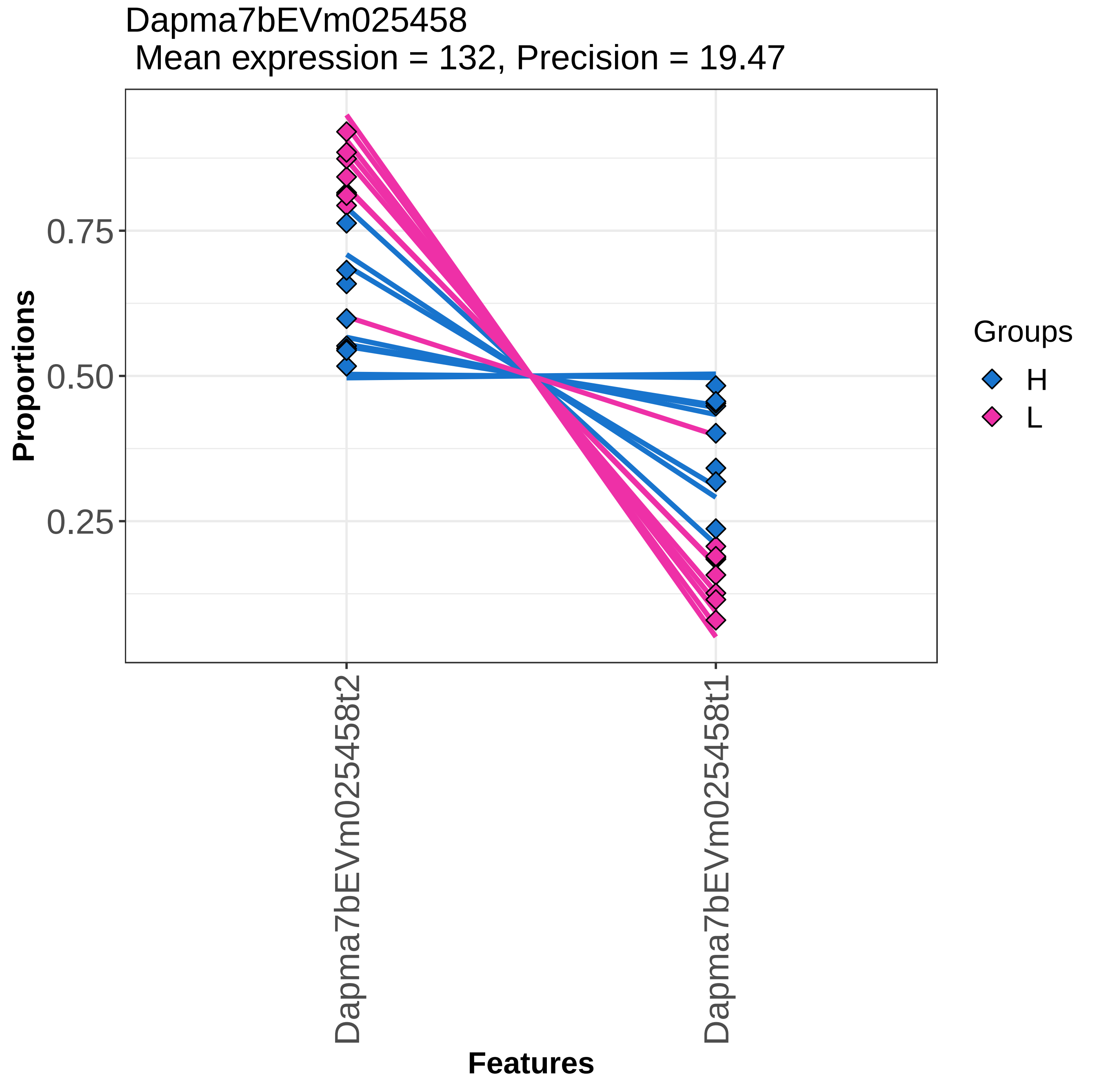

Dapma7bEVm025458
Mean expression = 132, Precision = 19.47
Proportions
0.75
0.50
0.25
Dapma7bEVm025458t2
Dapma7bEVm025458t1
Features
Groups
H
L

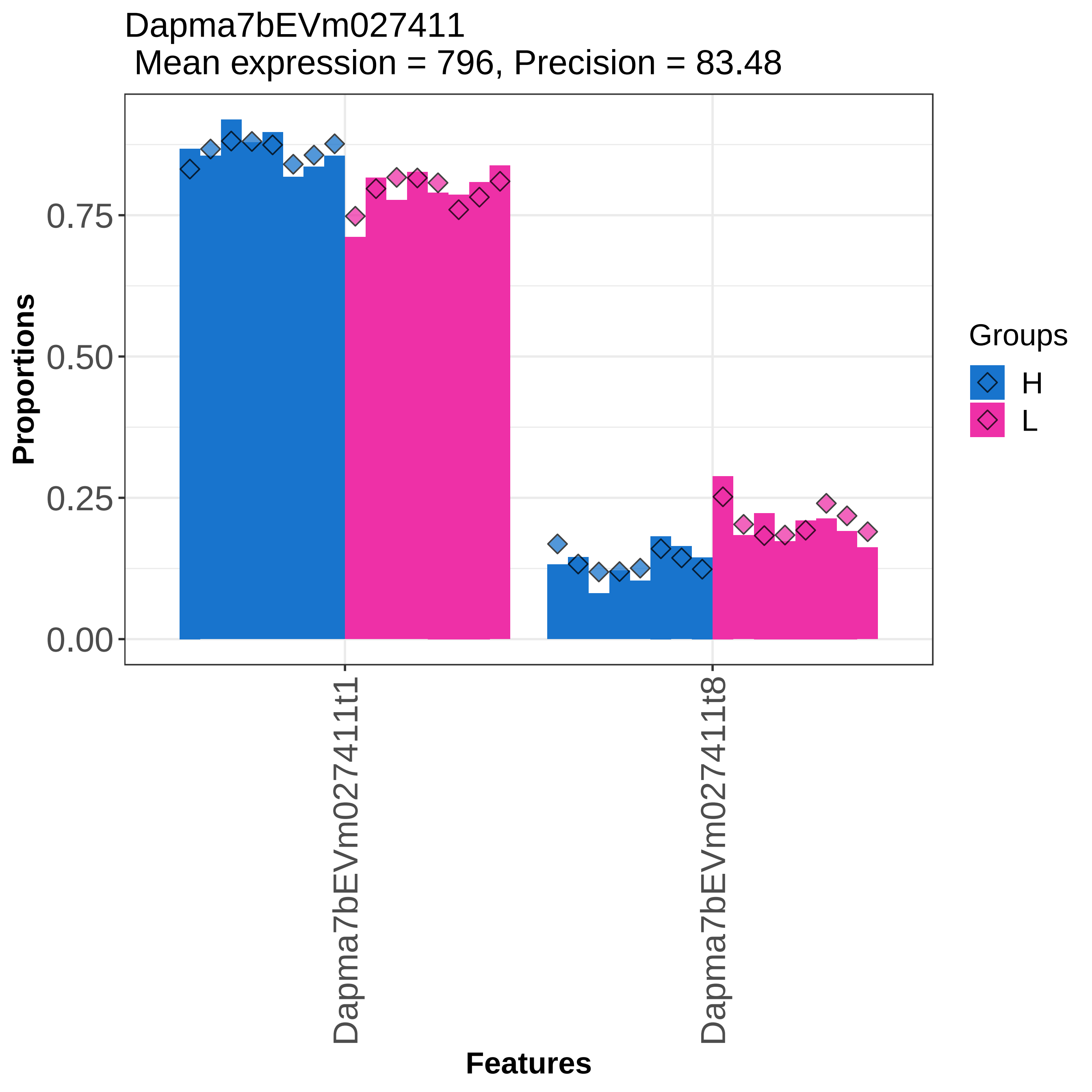

Dapma7bEVm027411
Mean expression = 796, Precision = 83.48
Proportions
0.00
0.25
0.50
0.75
Dapma7bEVm027411t1
Dapma7bEVm027411t8
Features
Groups
H
L

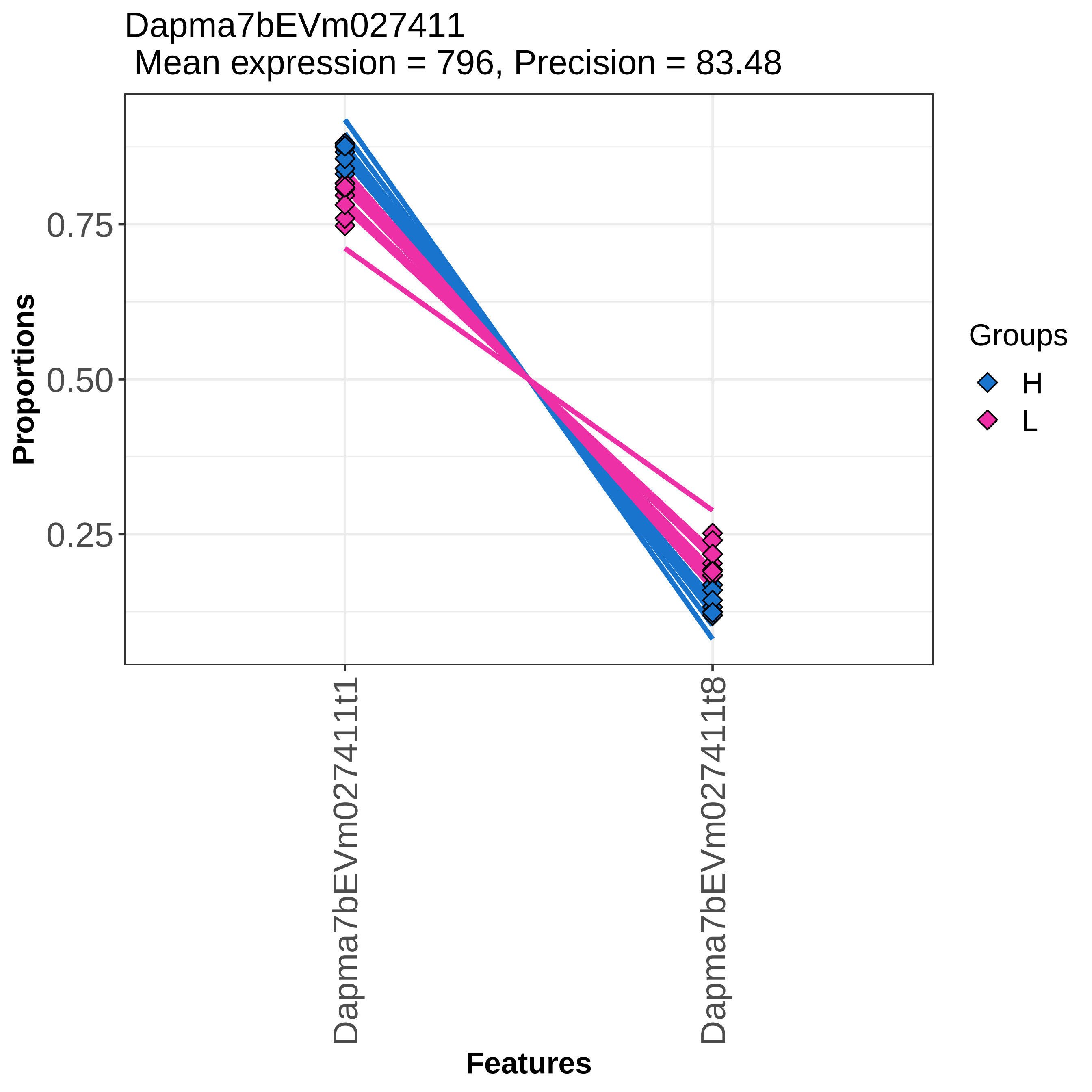

Dapma7bEVm027411
Mean expression = 796, Precision = 83.48
Proportions
0.75
0.50
0.25
Dapma7bEVm027411t1
Dapma7bEVm027411t8
Features
Groups
H
L

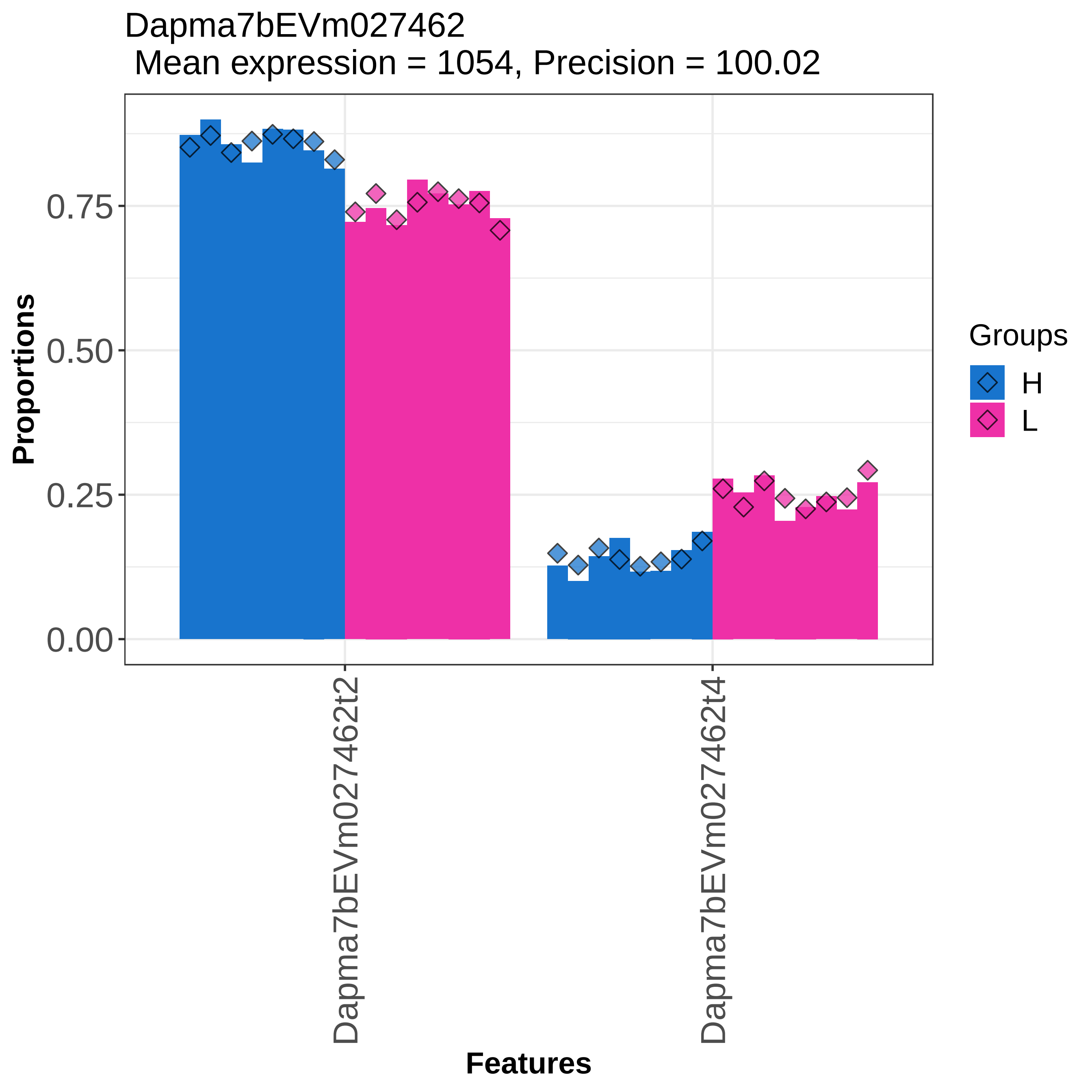

Dapma7bEVm027462
Mean expression = 1054, Precision = 100.02
Proportions
0.75
0.50
0.25
0.00
Dapma7bEVm027462t2
Dapma7bEVm027462t4
Features
Groups
H
L

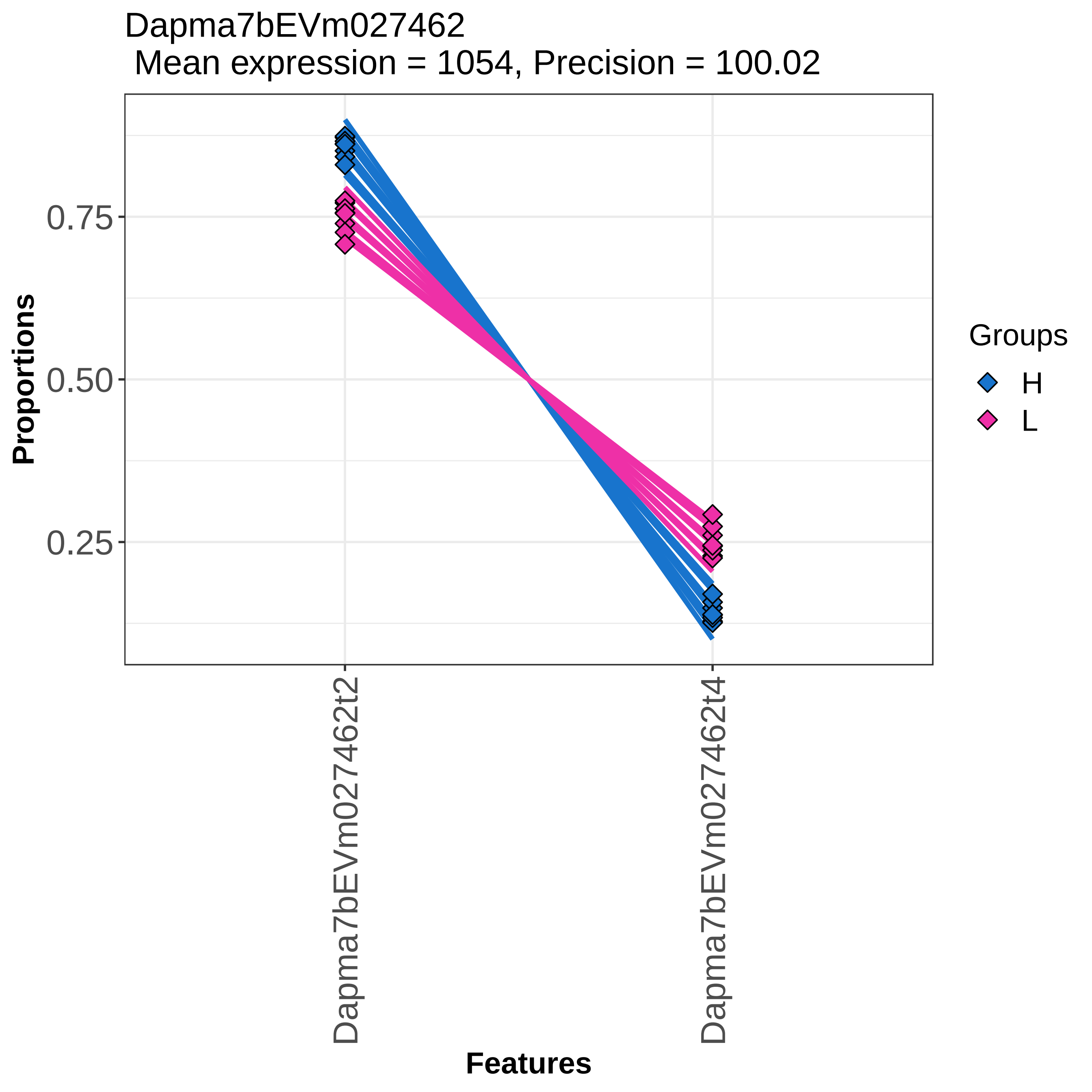
Dapma7bEVm027462
Mean expression = 1054, Precision = 100.02
Proportions
0.75
0.50
0.25
Dapma7bEVm027462t2
Dapma7bEVm027462t4
Features
Groups
H
L

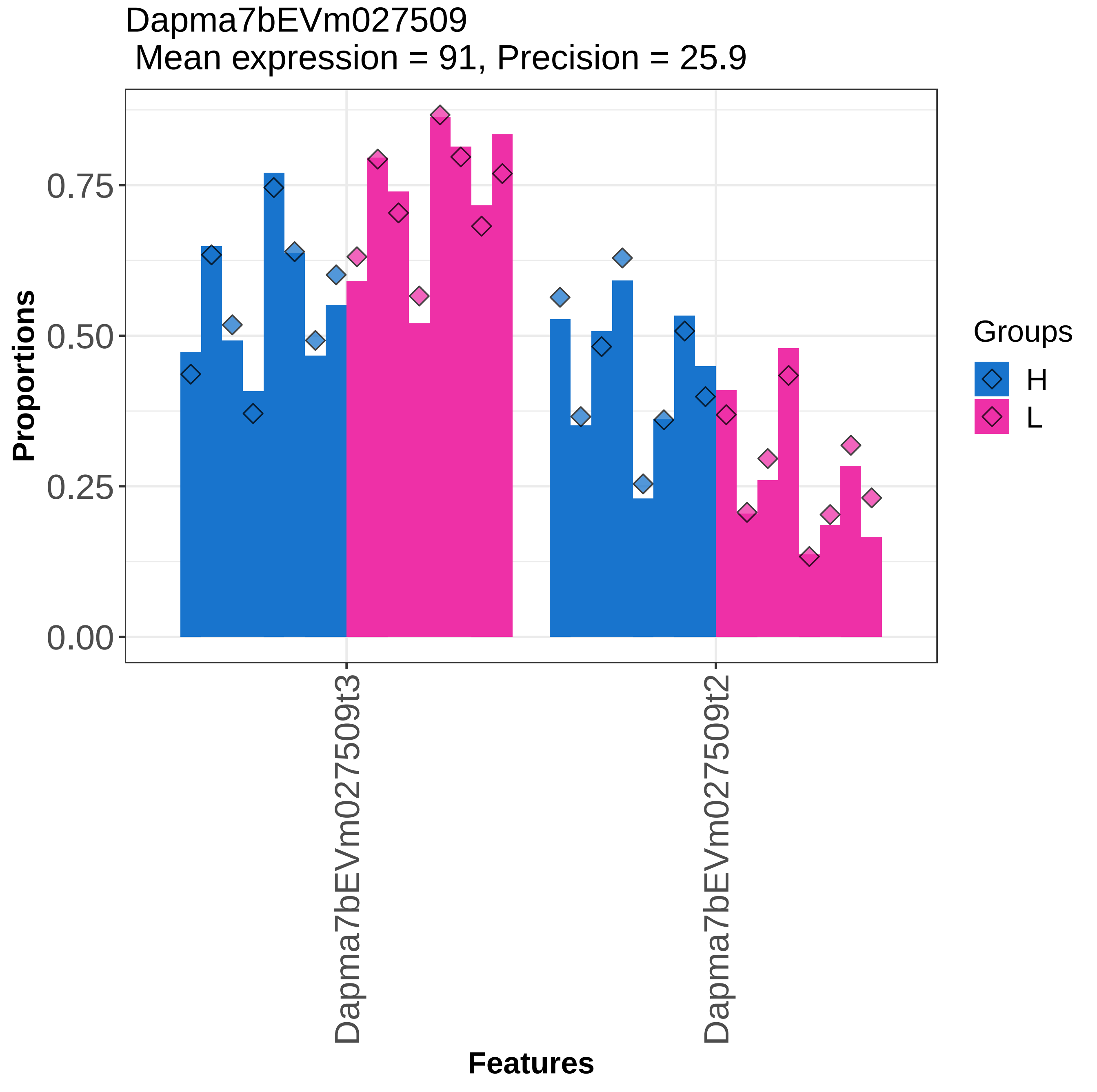

Dapma7bEVm027509
Mean expression = 91, Precision = 25.9
Proportions
0.00
0.25
0.50
0.75
Features
Dapma7bEVm027509t3
Dapma7bEVm027509t2
Groups
H
L

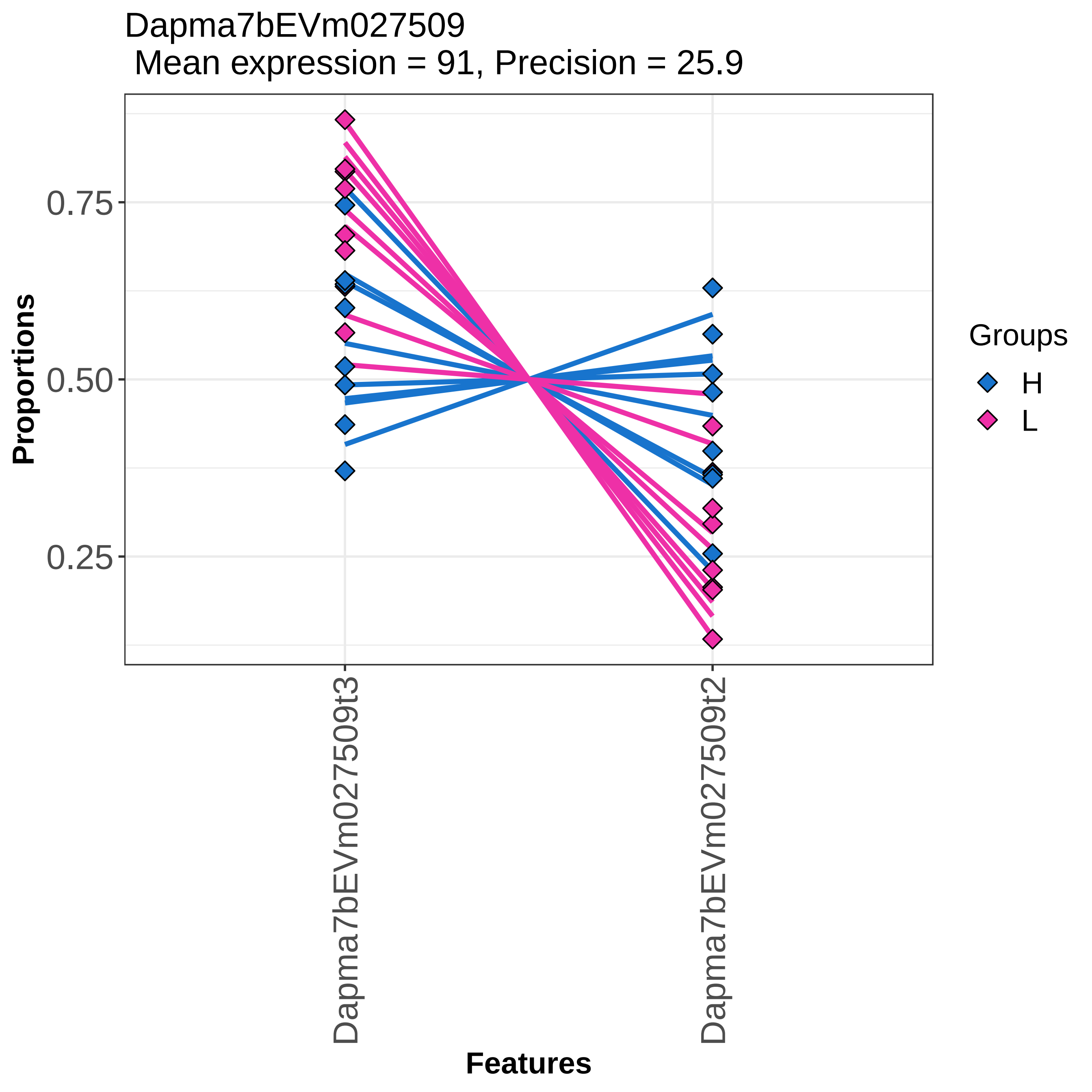

Dapma7bEVm027509
Mean expression = 91, Precision = 25.9
Proportions
0.75
0.50
0.25
Dapma7bEVm027509t3
Dapma7bEVm027509t2
Features
Groups
H
L

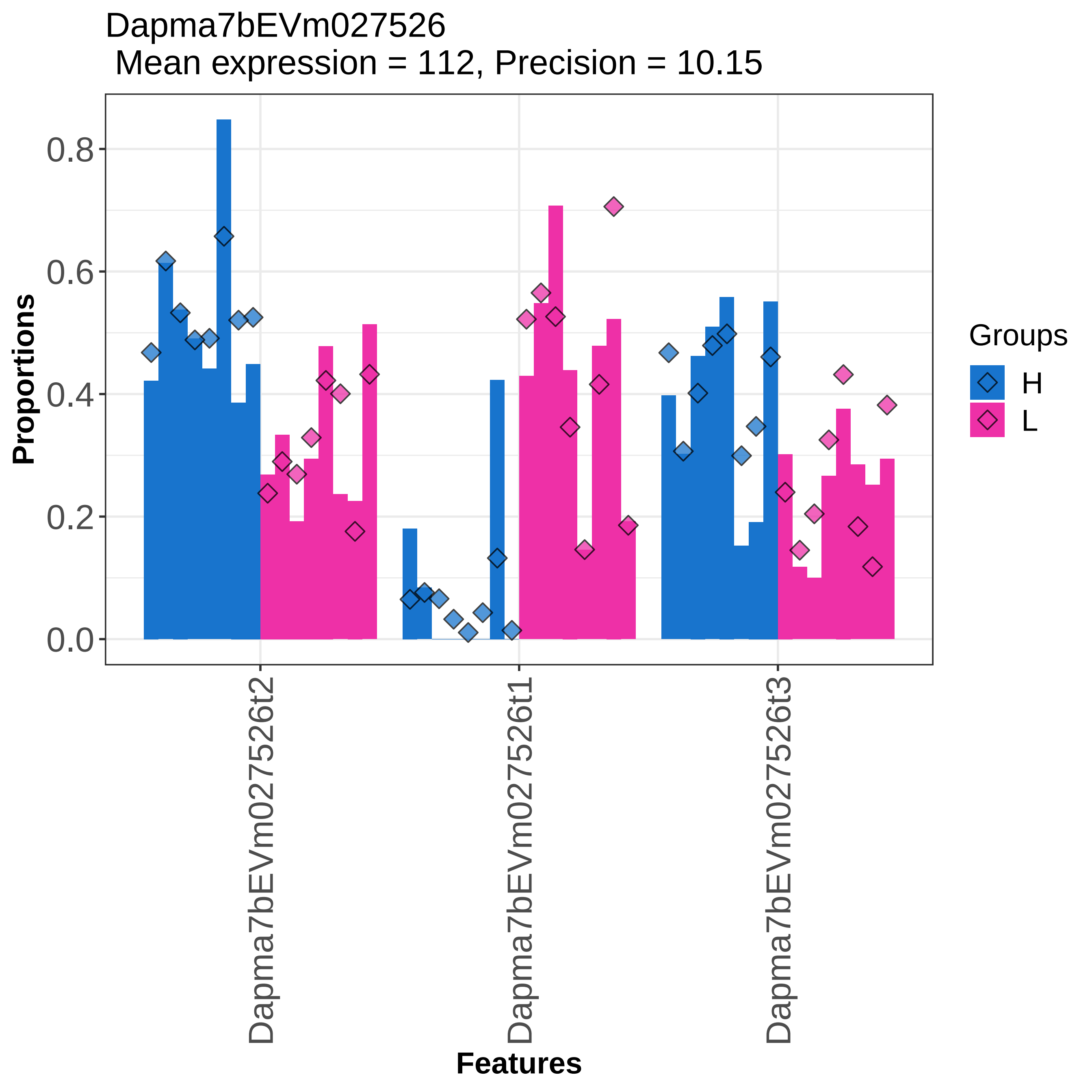

Dapma7bEVm027526
Mean expression = 112, Precision = 10.15
Proportions
0.0
0.2
0.4
0.6
0.8
Dapma7bEVm027526t2
Dapma7bEVm027526t1
Dapma7bEVm027526t3
Features
Groups
H
L

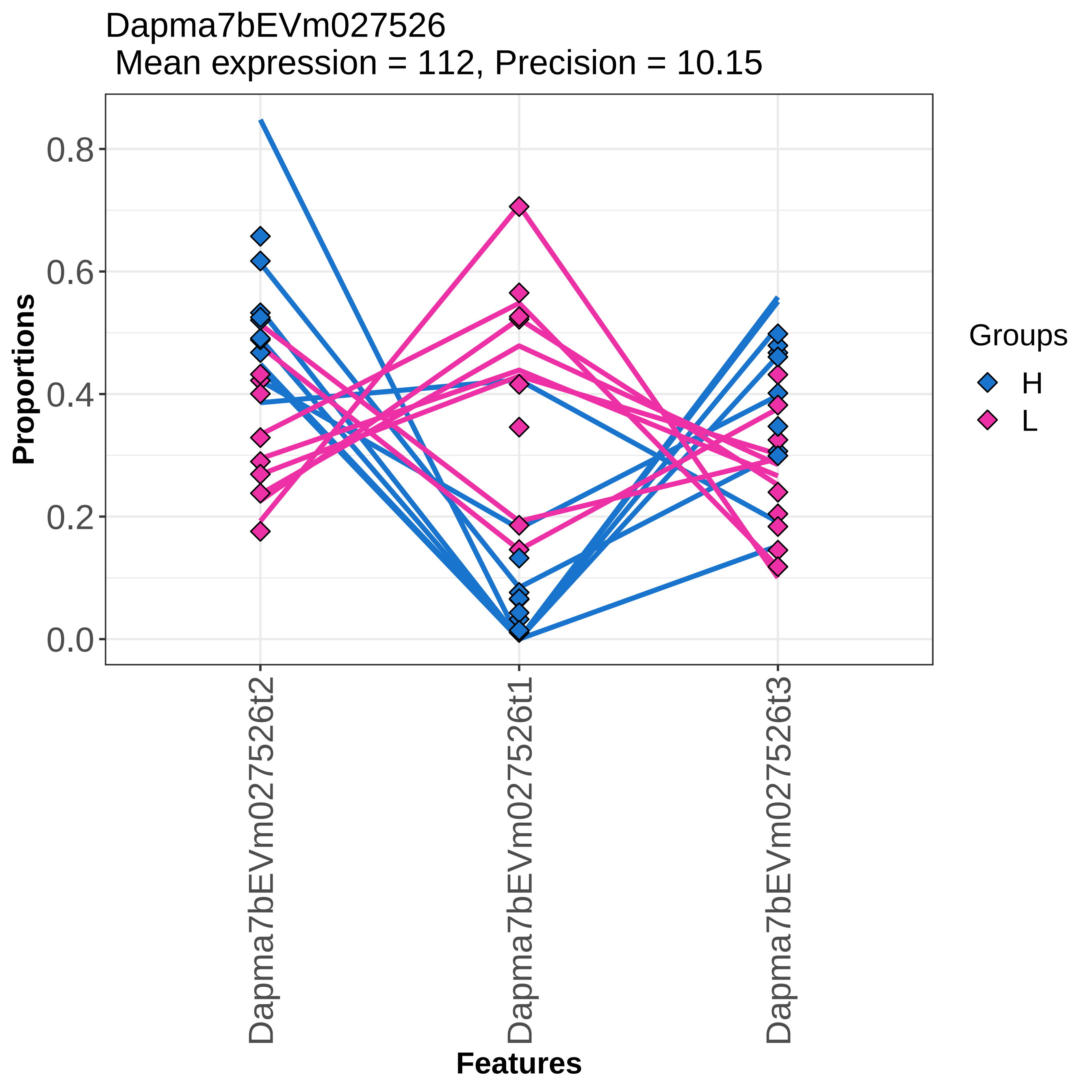
Dapma7bEVm027526
Mean expression = 112, Precision = 10.15
Proportions
0.0
0.2
0.4
0.6
0.8
Dapma7bEVm027526t2
Dapma7bEVm027526t1
Dapma7bEVm027526t3
Features
Groups
H
L

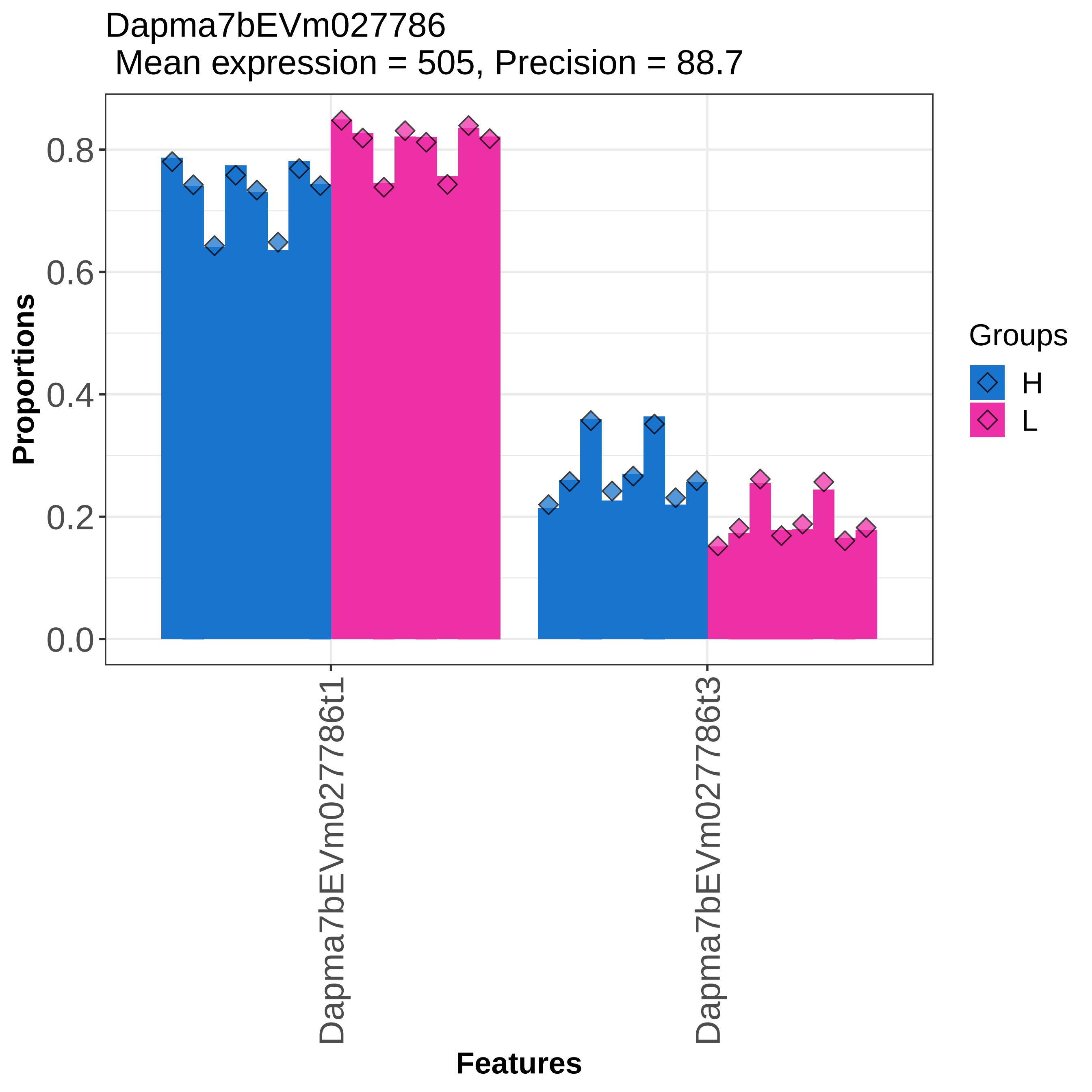

Dapma7bEVm027786
Mean expression = 505, Precision = 88.7
Proportions
0.0
0.2
0.4
0.6
0.8
Dapma7bEVm027786t1
Dapma7bEVm027786t3
Features
Groups
H
L

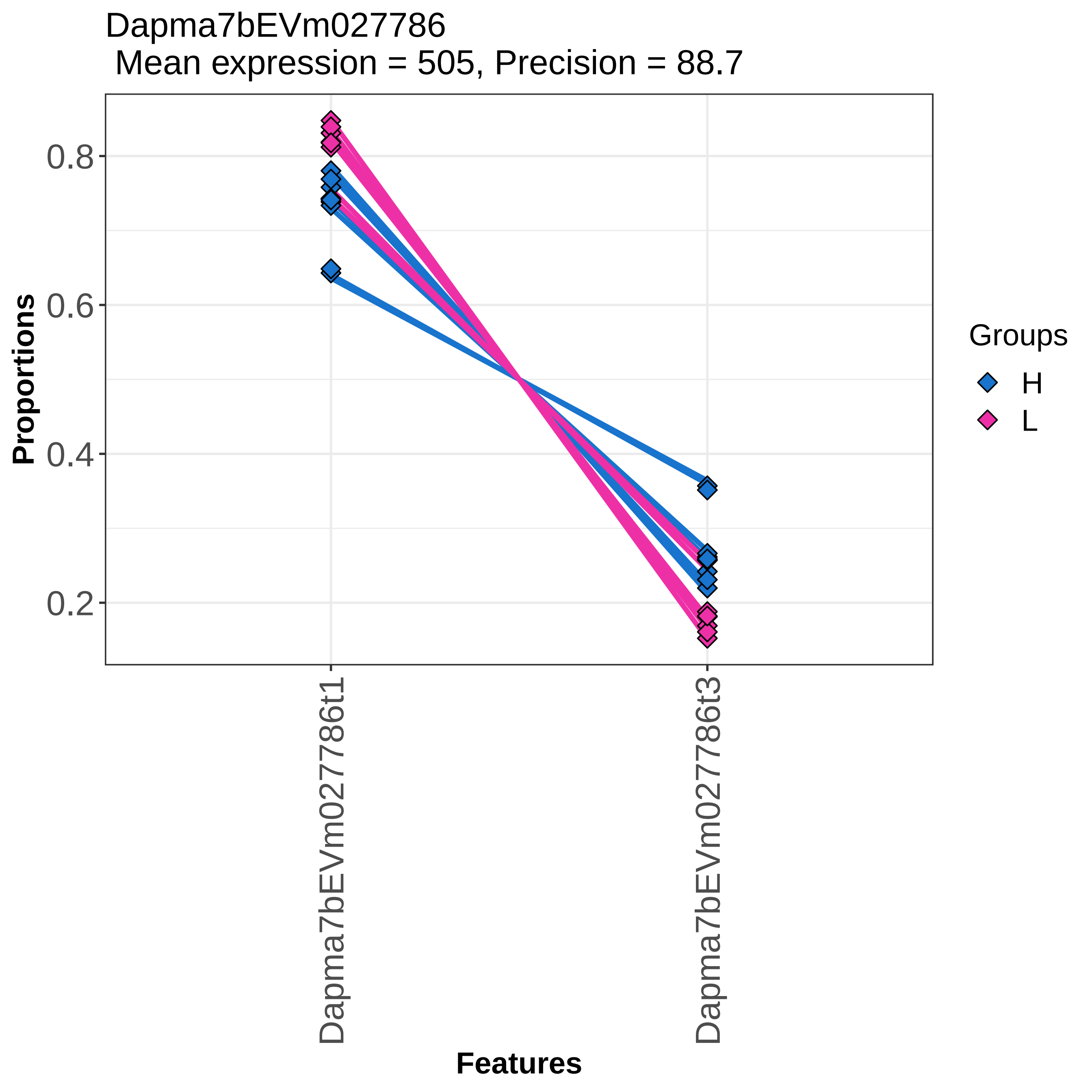

Dapma7bEVm027786
Mean expression = 505, Precision = 88.7
Proportions
0.8
0.6
0.4
0.2
Dapma7bEVm027786t1
Dapma7bEVm027786t3
Features
Groups
H
L

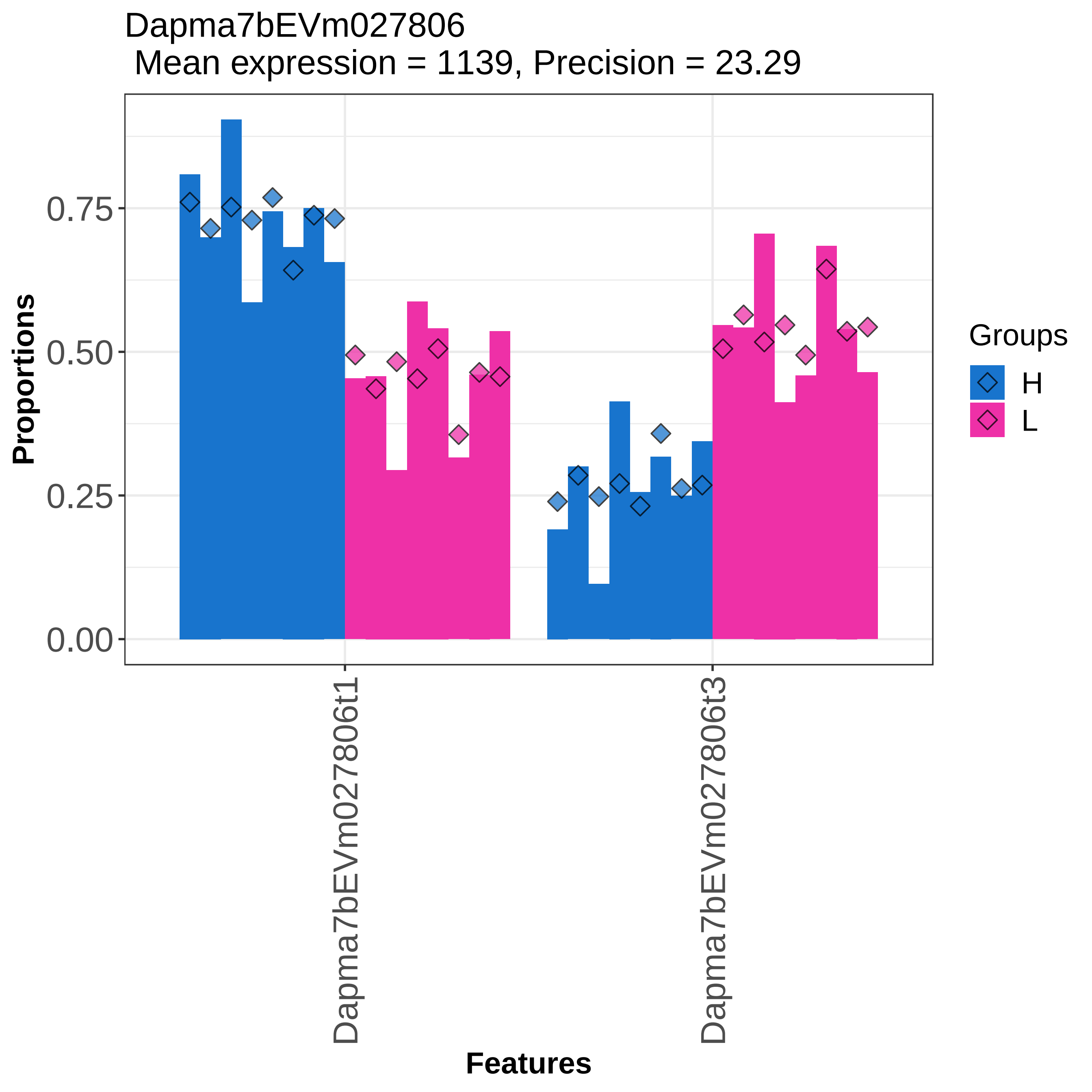
Dapma7bEVm027806
Mean expression = 1139, Precision = 23.29
Proportions
0.00
0.25
0.50
0.75
Dapma7bEVm027806t1
Dapma7bEVm027806t3
Features
Groups
H
L

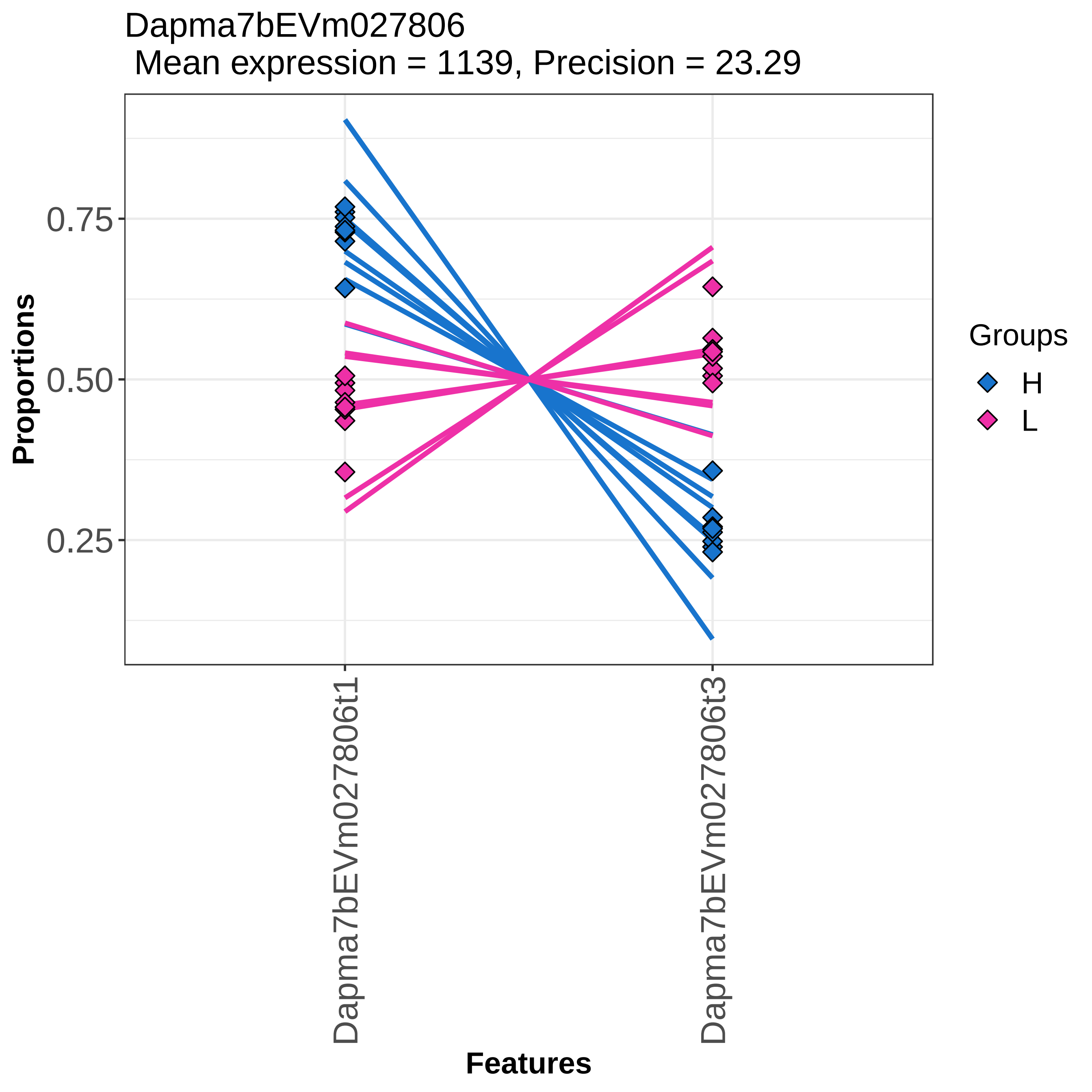

Dapma7bEVm027806
Mean expression = 1139, Precision = 23.29
Proportions
0.75
0.50
0.25
Groups
H
L
Dapma7bEVm027806t1
Dapma7bEVm027806t3
Features

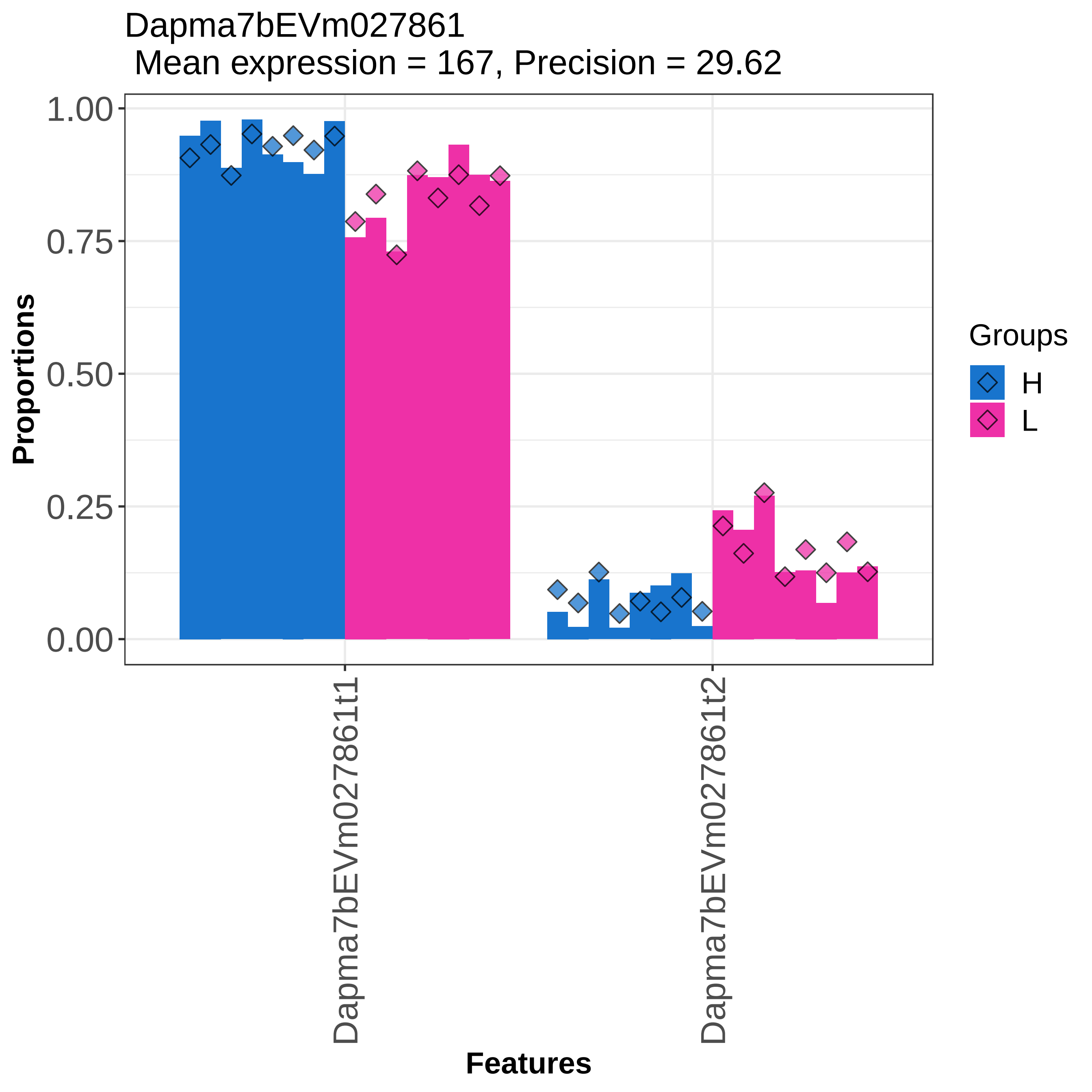
Dapma7bEVm027861
Mean expression = 167, Precision = 29.62
Proportions
1.00
0.75
0.50
0.25
0.00
Dapma7bEVm027861t1
Dapma7bEVm027861t2
Features
Groups
H
L

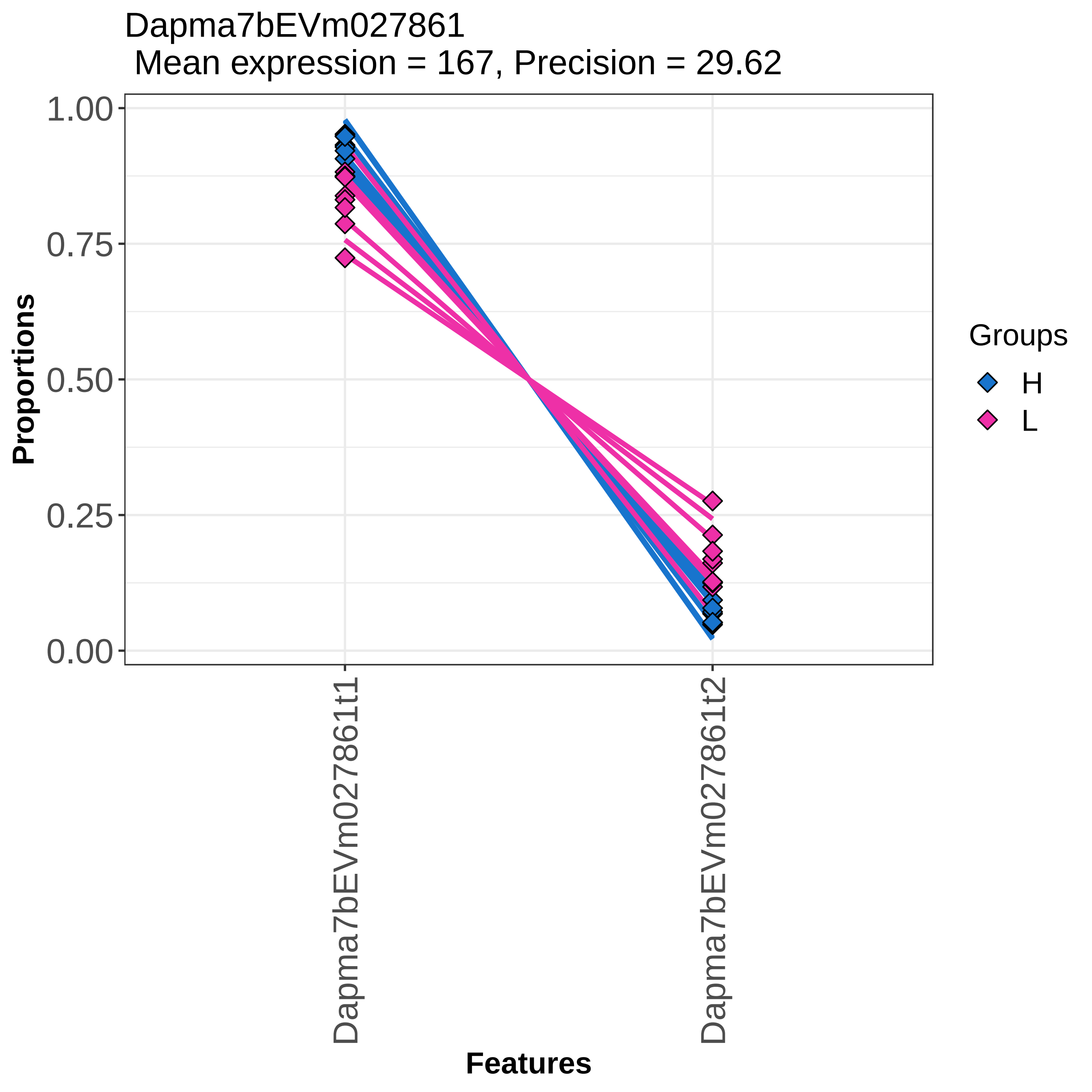

Dapma7bEVm027861
Mean expression = 167, Precision = 29.62
Proportions
1.00
0.75
0.50
0.25
0.00
Dapma7bEVm027861t1
Dapma7bEVm027861t2
Features
Groups
H
L

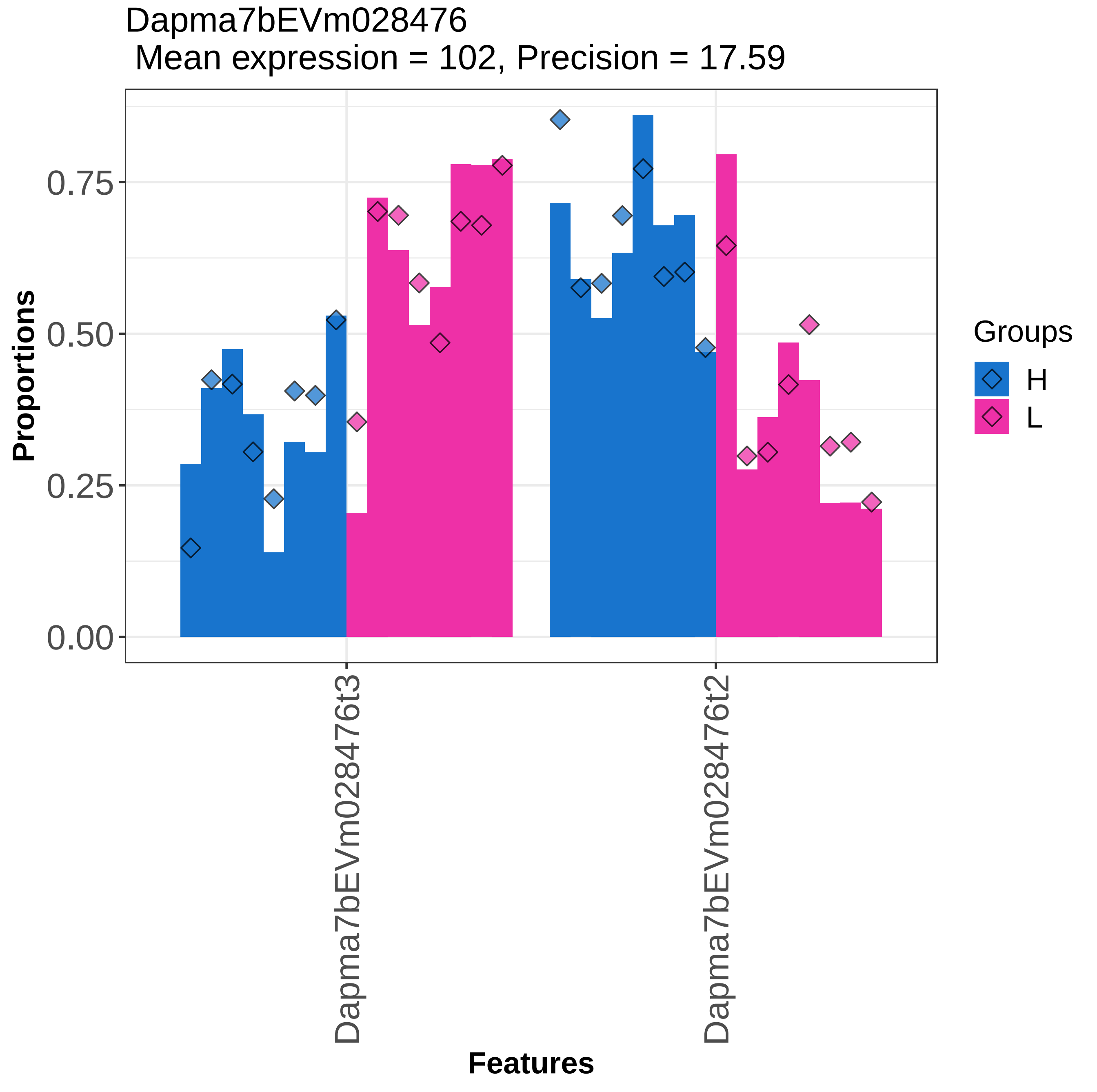

Dapma7bEVm028476
Mean expression = 102, Precision = 17.59
Proportions
0.00
0.25
0.50
0.75
Features
Dapma7bEVm028476t3
Dapma7bEVm028476t2
Groups
H
L

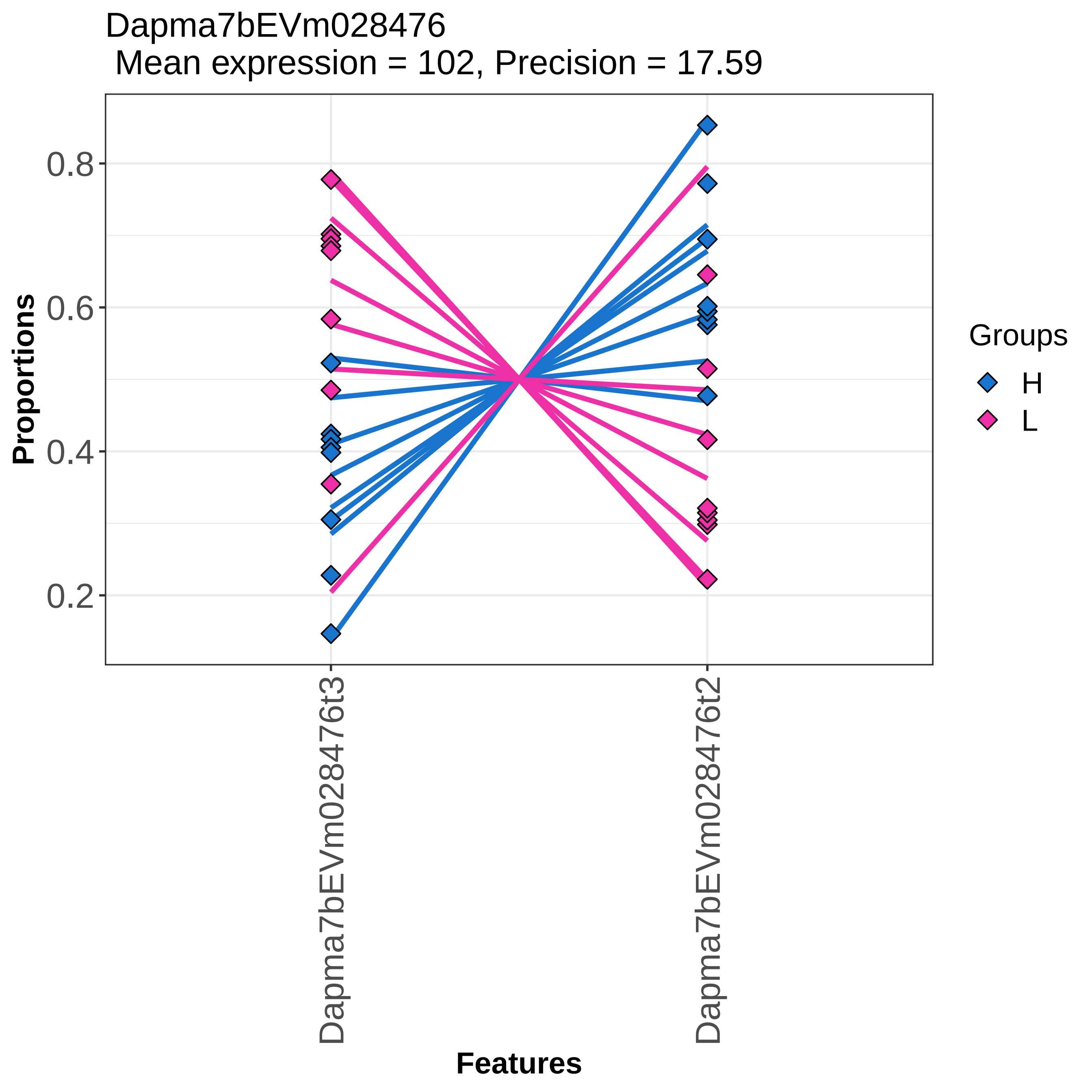

Dapma7bEVm028476
Mean expression = 102, Precision = 17.59
Proportions
0.8
0.6
0.4
0.2
Dapma7bEVm028476t3
Dapma7bEVm028476t2
Features
Groups
H
L

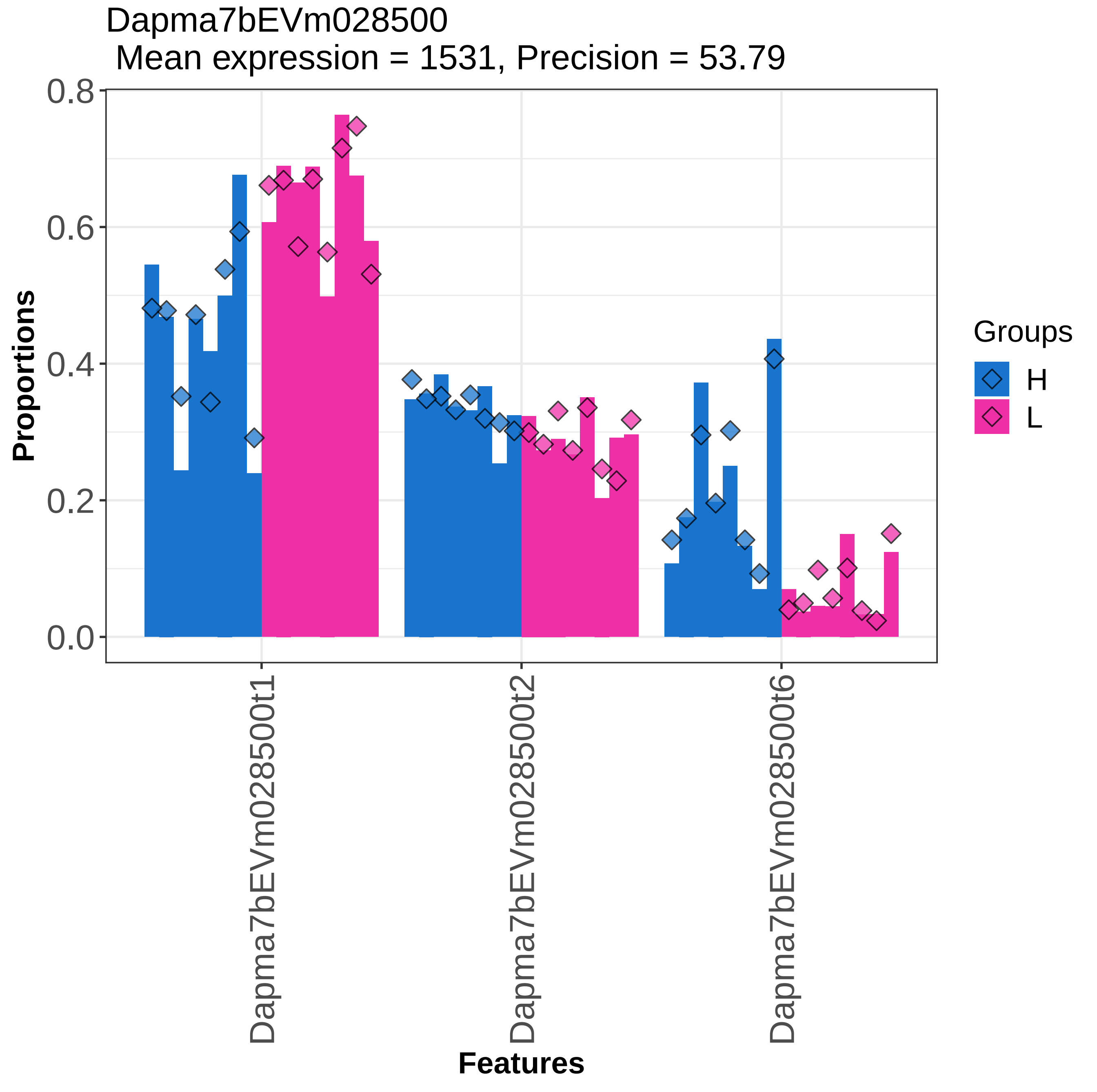
Dapma7bEVm028500
Mean expression = 1531, Precision = 53.79
0.8
0.6
0.4
0.2
0.0
Proportions
Dapma7bEVm028500t1
Dapma7bEVm028500t2
Dapma7bEVm028500t6
Features
Groups
H
L

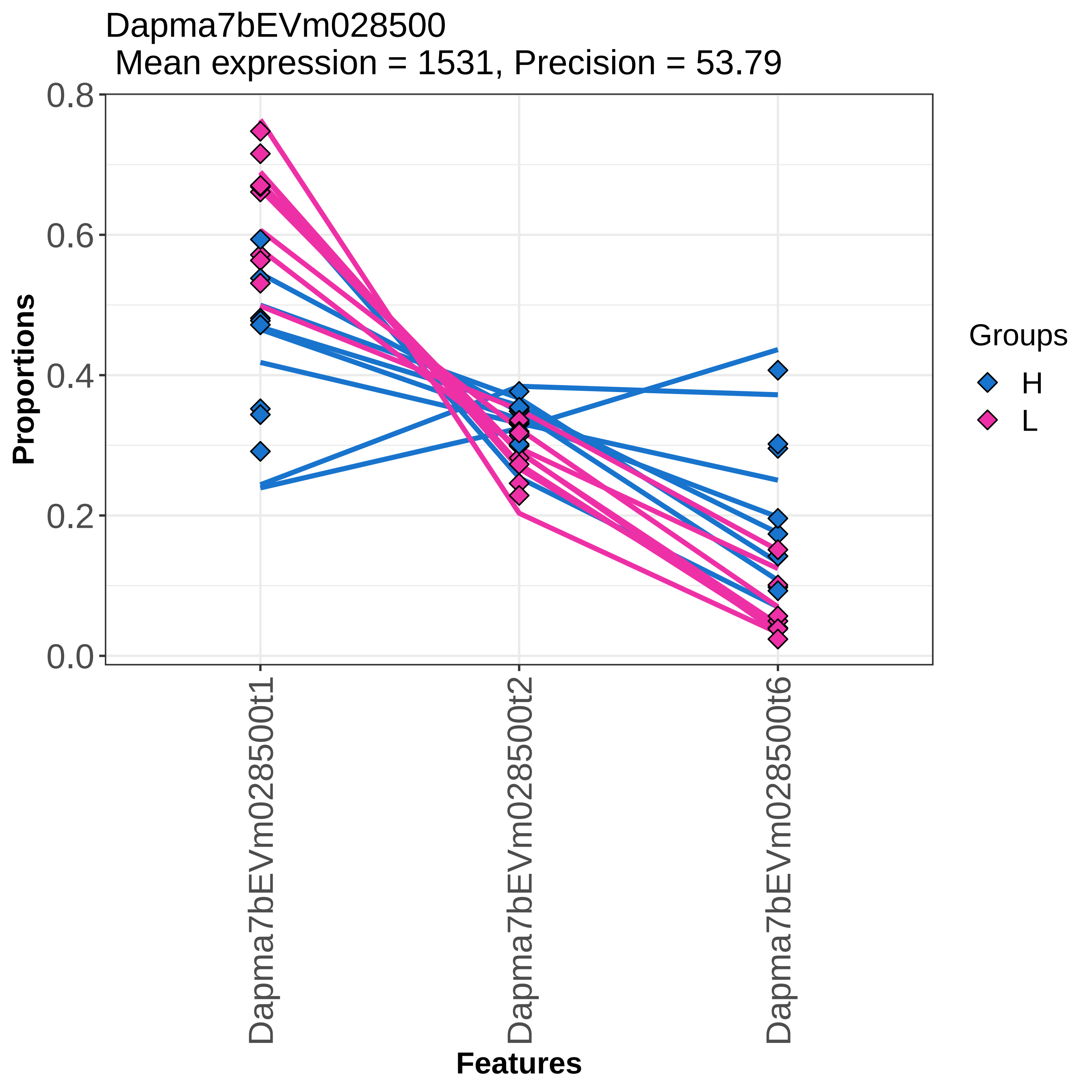
Dapma7bEVm028500
Mean expression = 1531, Precision = 53.79
Proportions
0.0
0.2
0.4
0.6
0.8
Dapma7bEVm028500t1
Dapma7bEVm028500t2
Dapma7bEVm028500t6
Features
Groups
H
L

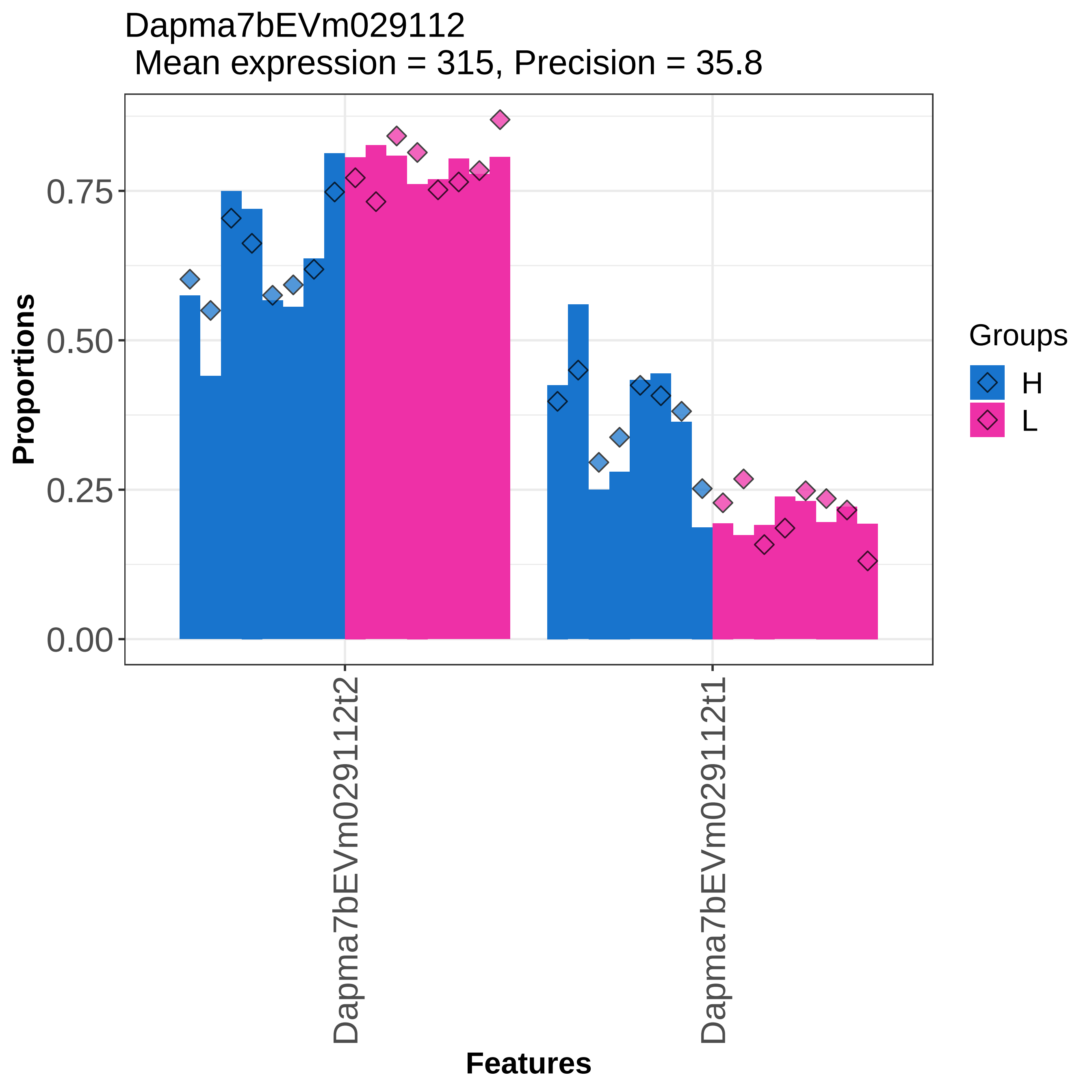
Dapma7bEVm029112
Mean expression = 315, Precision = 35.8
Proportions
0.75
0.50
0.25
0.00
Dapma7bEVm029112t2
Dapma7bEVm029112t1
Features
Groups
H
L

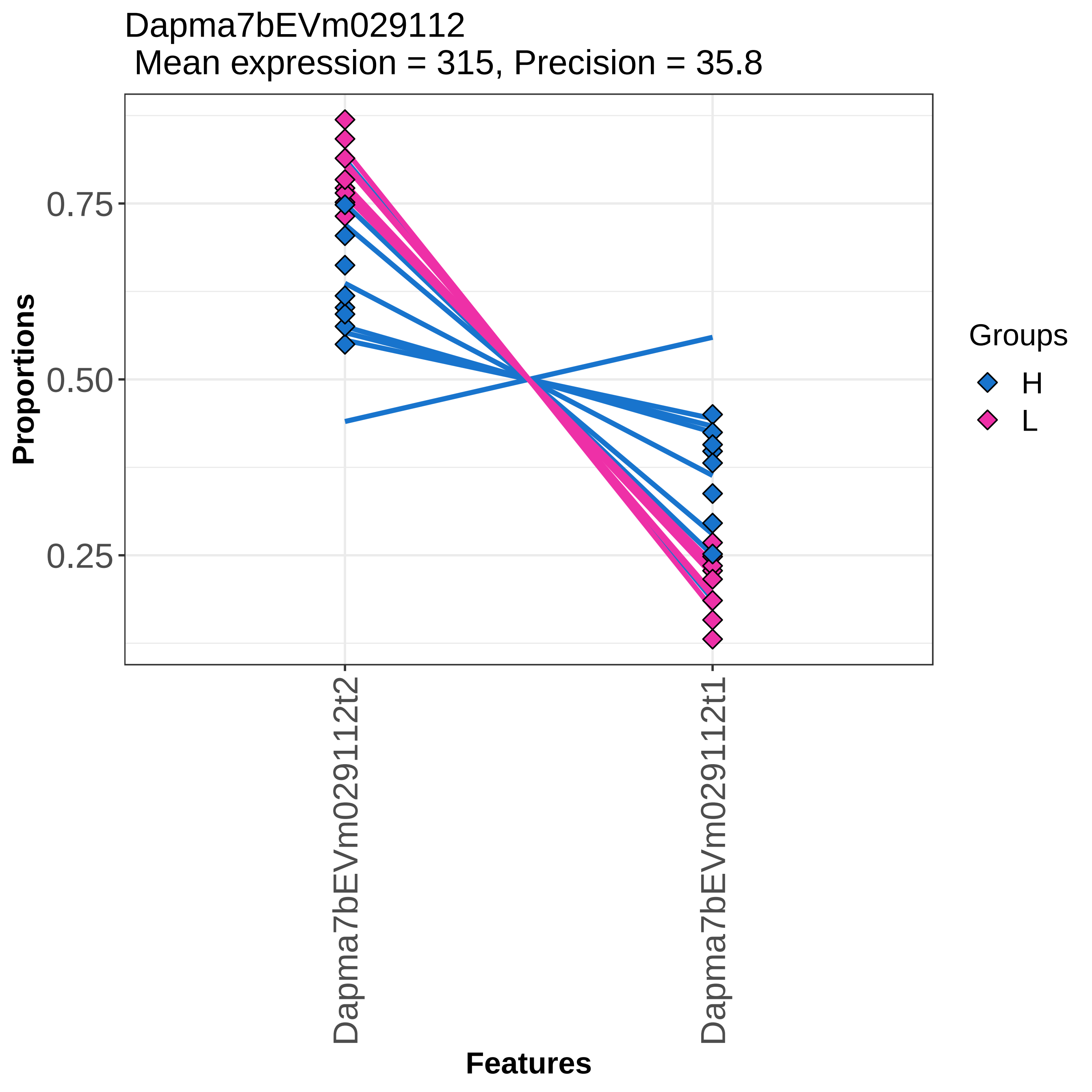

Dapma7bEVm029112
Mean expression = 315, Precision = 35.8
Proportions
0.75
0.50
0.25
Dapma7bEVm029112t2
Dapma7bEVm029112t1
Features
Groups
H
L
