## Supplementary figures and images for "*Daphnia magna* modifies its gene expression extensively in response to caloric restriction revealing a novel effect on haemoglobin isoform preference"

### Supplementary File 5

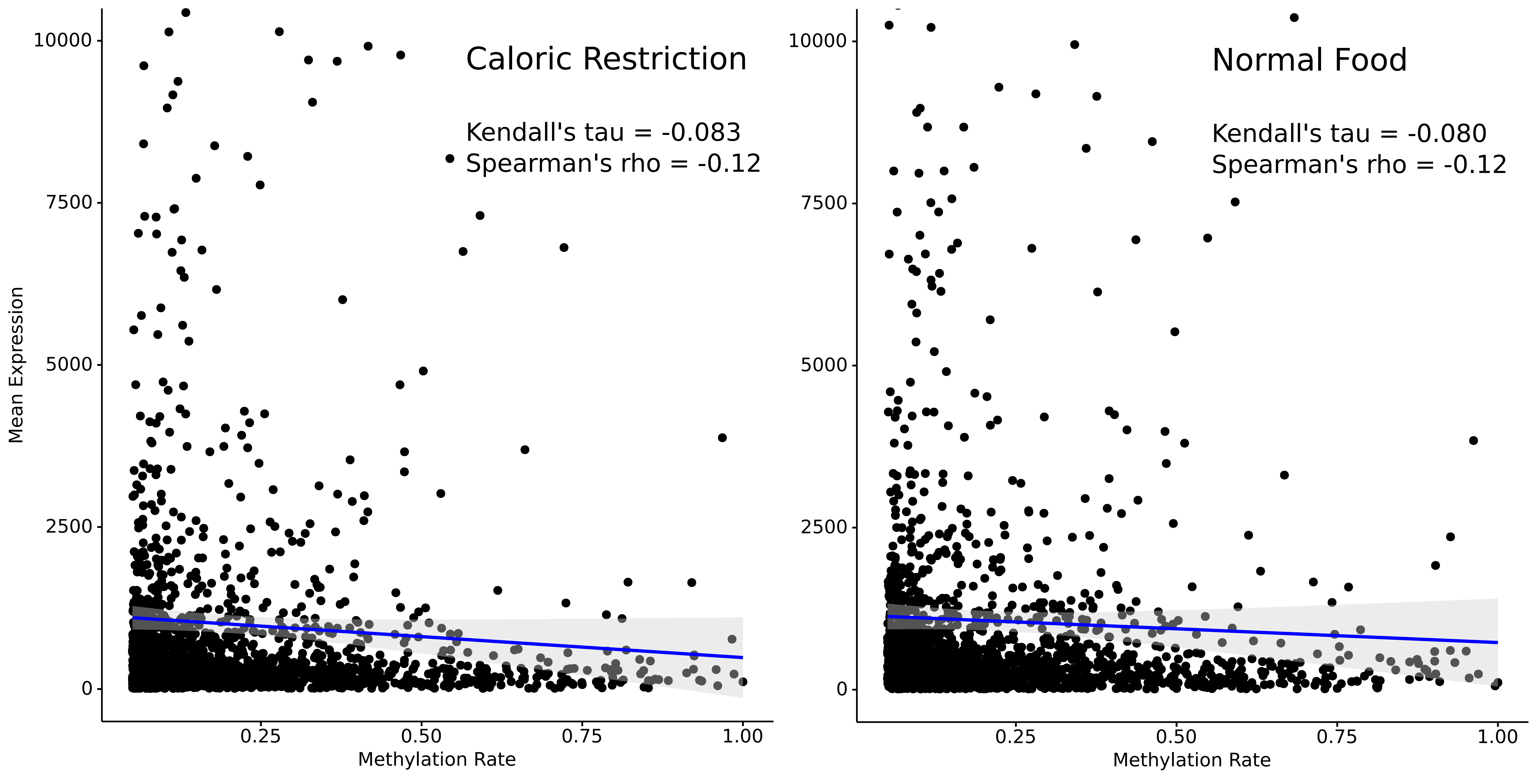
